# Computational Design of Peptides to Block Binding of the SARS-CoV-2 Spike Protein to Human ACE2

**DOI:** 10.1101/2020.03.28.013607

**Authors:** Xiaoqiang Huang, Robin Pearce, Yang Zhang

## Abstract

The outbreak of COVID-19 has now become a global pandemic and it continues to spread rapidly worldwide, severely threatening lives and economic stability. Making the problem worse, there is no specific antiviral drug that can be used to treat COVID-19 to date. SARS-CoV-2 initiates its entry into human cells by binding to angiotensin-converting enzyme 2 (hACE2) via the receptor binding domain (RBD) of its spike protein. Therefore, molecules that can block SARS-CoV-2 from binding to hACE2 may potentially prevent the virus from entering human cells and serve as an effective antiviral drug. Based on this idea, we designed a series of peptides that can strongly bind to SARS-CoV-2 RBD in computational experiments. Specifically, we first constructed a 31-mer peptidic scaffold by linking two fragments grafted from hACE2 (a.a. 22-44 and 351-357) with a linker glycine, and then redesigned the peptide sequence to enhance its binding affinity to SARS-CoV-2 RBD. Compared with several computational studies that failed to identify that SARS-CoV-2 shows higher binding affinity for hACE2 than SARS-CoV, our protein design scoring function, EvoEF2, makes a correct identification, which is consistent with the recently reported experimental data, implying its high accuracy. The top designed peptide binders exhibited much stronger binding potency to hACE2 than the wild-type (−53.35 vs. −46.46 EvoEF2 energy unit for design and wild-type, respectively). The extensive and detailed computational analyses support the high reasonability of the designed binders, which not only recapitulated the critical native binding interactions but also introduced new favorable interactions to enhance binding. Due to the urgent situation created by COVID-19, we share these computational data to the community, which should be helpful to develop potential antiviral peptide drugs to combat this pandemic.

## Introduction

The continuing pandemic of coronavirus disease 2019 (COVID-19) caused by severe acute respiratory syndrome coronavirus 2 (SARS-CoV-2, previously known as 2019-nCoV) has now become an international public health threat, causing inconceivable loss of lives and economic instability^1^. As of March 28, 2020, there have been more than 570000 confirmed cases and over 26000 deaths caused by COVID-19 worldwide^2^. Exacerbating the problem, there is no specific antiviral medication toward COVID-19, though development efforts are underway^3–6^. Although vaccines are thought to be the most powerful weapon to fight against virus invasion, it may take quite a long time to develop and clinically test the safety of a vaccine. Moreover, vaccines are usually limited as preventative measures given to uninfected individuals. Thus, as an emergency measure, it is desirable to develop effective antiviral therapeutics that can take effect rapidly not only to treat COVID-19 and but also to prevent its further transmission.

It has been confirmed that SARS-CoV-2 initiates its entry into host cells by binding to the angiotensin-converting enzyme 2 (ACE2) via the receptor binding domain (RBD) of its spike protein^7, 8^. Therefore, it is possible to develop new therapeutics to block SARS-CoV-2 from binding to ACE2. Although small molecule compounds are commonly preferred as therapeutics, they are not effective at blocking protein-protein interactions (PPIs) where a deep binding pocket may be missing at the interface^9^. On the contrary, peptide binders are more suitable for disrupting PPIs by specifically binding to the interface binding region^10^. More importantly, small peptides have reduced immunogenicity^11^. These positive features make peptides great candidates to serve as therapeutics^12, 13^. Recently, Zhang et al^14^ reported that the natural 23-mer peptide (a.a. 21-43) cut from the human ACE2 (hACE2) α1 helix can strongly bind to SARS-CoV-2 RBD with a disassociation constant (K_d_) of 47 nM, which was comparable to that of the full-length hACE2 binding to SARS-CoV-2 RBD^15^; they also showed that a shorter 12-mer peptide (a.a. 27-38) from the same helix was not able to bind the virus RBD at all. In an earlier report, Han et al^16^ performed a study to identify the critical determinants on hACE2 for SARS-CoV entry, and they found that two natural peptides from hACE2 (a.a. 22-44 and 22-57) exhibited a modest antiviral activity to inhibit the binding of SARS-CoV RBD to hACE2 with IC50 values of about 50 μM and 6 μM, respectively, implying that the peptide composed of residues 22-57 had stronger binding affinity for SARS-CoV RBD. They also generated a hybrid peptide by linking two discontinuous fragments from hACE2 (a.a. 22-44 and 351-357) with a glycine, and this 31-mer exhibited a potent antiviral activity with an IC50 of about 0.1 μM, indicating that this artificial peptide had much stronger binding affinity for SARS-CoV RBD than the peptides composed of residues 22-44 or 22-57. Due to the high similarity of the binding interfaces between SARS-CoV RBD/hACE2 and SARS-CoV-2 RBD/hACE2, we think that this artificial peptide may also bind to SARS-CoV-2 more strongly than the peptide 21-43 tested by Zhang et al^14^, which is similar to the peptide 22-44 from Han et al^16^. Although the natural peptides are promising, it has been argued that the sequence of hACE2 is suboptimal for binding the S protein of SARS-CoV-2^17^. Therefore, further redesign of the natural peptides may significantly enhance its binding affinity to the virus RBD and the improved peptide binders may have the potential to inhibit SARS-CoV-2 from entering human cells and hinder its rapid transmission.

In this work, we computationally designed thousands of peptide binders that exhibited stronger binding affinity for SARS-CoV-2 than the natural peptides through computational examination. Based on the crystal structure of the SARS-CoV-2 RBD/hACE2 complex, we constructed a hybrid peptide by linking two peptidic fragments from hACE2 (a.a. 22-44 and 351-357) with a glycine. Starting from the peptide-protein complex, we used our protein design approaches, EvoEF2^18^ and EvoDesign^19^, to completely redesign the amino acid sequences that match the peptide scaffold while enhancing its binding affinity to SARS-CoV-2. We have selected dozens of designed peptides for wet-lab experimental validation and the experiments are ongoing. Although our experimental results are not available yet, we have performed extensive computational benchmark tests to show the reasonability of the computationally designed peptide binders. Due to the urgent situation caused by COVID-19 and the limited resources in our own laboratory, we share these computational data to the scientific community and hope researchers can work together to test them and to identify potential antiviral peptide therapeutics to combat this pandemic.

## Methods

### Initial peptide scaffold construction

Several experimental SARS-CoV-2 RBD/hACE2 complex structures have been reported^20–22^, and deposited in the Protein Data Bank (PDB)^23^. Specifically, PDB ID 6m17 is a 2.9 Å structure of the SARS-CoV-2 RBD/ACE2-B0AT1 complex determined using the cryogenic electron microscopy (Cryo-EM) technique^22^. Furthermore, PDB ID 6m0j is a 2.45 Å X-ray crystal structure of SRRS-CoV-2 RBD/hACE2^20^, while 6vw1 is a 2.68 Å X-ray structure of SARS-CoV-2 chimeric RBD/hACE2^21^, where the chimeric RBD is comprised of the receptor binding motif (RBM) from SARS-CoV-2 S and the core from SARS-CoV, with the mutation N439R. The three experimental complex structures are quite similar to each other in terms of global folds (Figure 1A). Since 6vw1 does not contain the wild-type SARS-CoV-2 RBD, we did not use it as template. Based on a preliminary examination, we found that structure quality of 6m0j was better than 6m17 (see Results and Discussion), and therefore we only considered 6m0j as the template complex.

**Figure 1.**
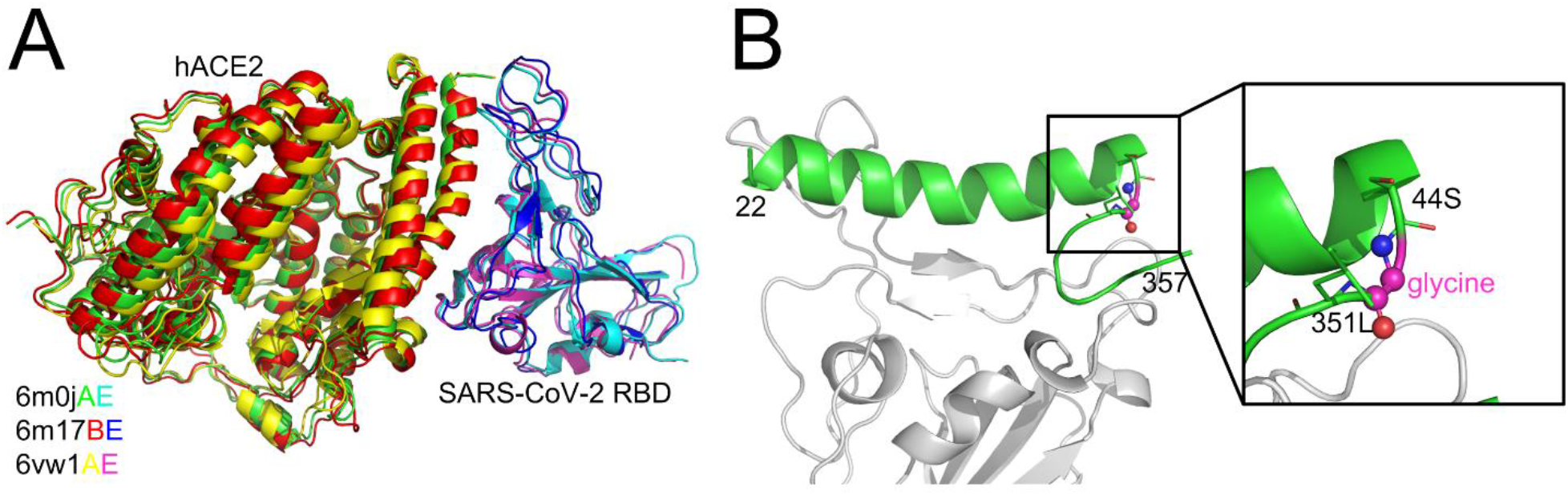
Comparison of the SARS-CoV-2 RBD/hACE2 complex structures (A) and the constructed SARS-CoV-2 RBD/hACE2 peptide complex (B). The superposition of the three complex structures was performed using MM-align^25^; the TM-score^26^ between each complex pair was >0.98.

Two peptide fragments (a.a. 22-44 and 351-357) from hACE2 (6m0j, chain A) were extracted because they were in extensive contact with SARS-CoV-2 RBD (6m0j, chain E). The positions 44 and 351 were chosen because the distance between their Cα atoms was only 5.5 Å, and therefore only one residue was required to link them. To reduce the interference to the surrounding amino acids, the linker residue was initially chosen as glycine. The small loop, 44S-glycine-351L, was then reconstructed using MODELLER^24^, while the other parts of the whole peptide were kept constant; five similar loop conformations were produced and the one with the best DOPE score was selected. For the sake of simplifying the discussion, the initial hybrid peptide constructed in this manner was denoted as the wild-type (note that it was not a truly native peptide), and the complex structure of SARS-CoV-2 RBD/hACE2 hybrid peptide was used as the template for computational peptide design (Figure 1B).

### Peptide design procedure

Based on the constructed protein-peptide complex structure (SARS-CoV-2 RBD/hACE2-22-44G351-357), we performed 1000 independent design trajectories individually, using (1) EvoEF2^18^, a physics- and knowledge-based energy function specifically designed for protein design and (2) a new version of EvoDesign^19^, which combines EvoEF2 and evolutionary profiles for design scoring. A simulated annealing Monte Carlo (SAMC)^27^ protocol was used to search for low total energy sequences as previously described^18^. For each trajectory, only the single lowest energy in that design simulation was selected, and therefore 1000 sequences each were collected from the EvoEF2 and EvoDesign designs. The EvoEF2 and EvoDesign designs were separately analyzed to determine the impact of the physics- and profile-based scores. Since SAMC is a stochastic searching method, some of the 1000 sequences were duplicates and thus excluded from analysis. The backbone conformations of the hACE2 peptide and SARS-CoV-2 RBD were held constant during the protein design simulations, all the residues on the peptide were redesigned, and the side-chains of the interface residues on the virus RBD were repacked without design. The non-redundant designed peptides are listed in Supplementary Tables S1-S5, and the raw data are freely available at https://zhanglab.ccmb.med.umich.edu/EvoEF/COVID-19/.

### Evolutionary profile construction

To construct reliable structural evolutionary profiles, we used the hACE2 protein structure instead of the hybrid peptide to search structural analogs against a non-redundant PDB library. Only structures with a TM-score ≥0.7 to the hACE2 scaffold were collected to build a pairwise multiple sequence alignment (MSA). A total of nine structural analogs were identified. The corresponding alignment for residues 22-44 and 351-357 were directly extracted from the full-length MSA and combined to build an MSA for the hybrid peptide. Since an arbitrary glycine was used to link positions 44 and 351, a gap ‘-’ was inserted in the peptide MSA for the glycine position. The peptide MSA constructed in this manner is described in Supplementary Figure S1. The peptide MSA was used to construct the evolutionary profile position-specific scoring matrix (PSSM) as previously described^28^.

In previous studies, we also proposed incorporating protein-protein interface evolutionary profiles to model PPIs^19, 29, 30^. However, no interface structural analogs were identified from the non-redundant interface library (NIL)^30^, and no interface sequence analogs were found from the STRING^31^ database with a PPI link score ≥0.8. Therefore, the interface evolutionary profile scoring was excluded from design.

## Results and Discussion

### Evaluation of EvoEF2 score on experimental complexes

At the very beginning of the outbreak of SARS-CoV-2, to determine its relative infectivity, many computational studies were performed to compare the binding affinity of SARS-CoV-2 RBD for hACE2 with that of SARS-CoV RBD for hACE2 based on homology modeling structures; all these studies came up with the conclusion that SARS-CoV-2 showed much weaker binding affinity to hACE2 than SARS-CoV and SARS-CoV-2 might not be as infectious as SARS-CoV^32–34^. However, the recent biochemical studies demonstrated that SARS-CoV-2 exhibited much stronger binding affinity to hACE2 than SARS-CoV^3, 15, 21^, implying that the homology models may not have been sufficiently accurate for binding affinity assessment based on atomic-level scoring functions, although the global folds of these models were correct.

Here, we used the EvoEF2 energy function to evaluate the binding affinity of SARS-CoV and/or SARS-CoV-2 (chimeric) RBD for hACE2 based on the experimental structures described above. As shown in Table 1, SARS-CoV-2 RBD showed stronger binding potency (lower EvoEF2 scores indicate stronger binding affinity) to hACE2 than SARS-CoV based on the calculations performed on two X-ray crystal structures (PDB IDs: 2ajf and 6m0j), regardless of whether or not the residues at the protein-protein interfaces were repacked; the computationally estimations were consistent with the experimental results (Table 1). However, the EvoEF2 binding scores calculated using the Cryo-EM structure (i.e. 6m17) were much higher than those obtained from the X-ray structure 6m0j, suggesting that the Cryo-EM structure might not be as high quality as its X-ray counterparts. In fact, we manually inspected 6m0j and 6m17 and found that significantly more steric clashes were present in 6m17 (data not shown). Moreover, Shang et al^21^ demonstrated that the artificial SARS-CoV-2 chimeric RBD showed improved binding affinity to hACE2, compared to the wild-type SARS-CoV-2, and this improvement was also somewhat captured by EvoEF2 (Table 1). Thus, out of the two wild-type SARS-CoV-2 RBD/hACE2 structures (6m0j and 6m17), only 6m0j was used as a template structure for the peptide design study because it was better refined.

**Table 1.** Comparison of binding affinities for different PPIs.

| PPI | Experiment K<sub>d</sub> | EvoEF2 score (EEU) |  |
| --- | --- | --- | --- |
|  |  | Interface not repacked | Interface repacked |
| SARS-CoV RBD/hACE2 | 325.8 nM<sup>15</sup><br>185 nM<sup>21</sup> | -40.73 (2ajfAE) | -51.12 (2ajfAE) |
| SARS-CoV-2 RBD/hACE2 | 14.7 nM<sup>15</sup> | -49.95 (6m0jAE) | -55.67 (6m0jAE) |
|  | 44.2 nM<sup>21</sup> | -19.84 (6m17BE) | -30.50 (6m17BE) |
|  |  | -19.84 (6m17DF) | -30.50 (6m17DF) |
| SARS-CoV-2 chimeric RBD/hACE2 | 23.2 nM<sup>21</sup> | -53.15 (6vw1AE) | -58.81 (6vw1AE) |

EEU stands for EvoEF2 energy unit.

### Peptide design results and computational analyses

Eight out of the 1000 low-energy sequences that were designed using the EvoEF2 energy function were duplicates, resulting in 992 non-redundant designs. The EvoEF2 total energy values of the designed protein complex structures ranged from −829 to −816 EvoEF2 energy unit (EEU), the majority of which varied from −827 to −822 EEU (Figure 2A). The EvoEF2 binding energies of the 992 designed peptides to SARS-CoV-2 RBD ranged from −53 to −40 EEU, centering around −50 to −47 EEU (Figure 2B). The sequence identities between the designed peptides and the wild-type peptide was diversely distributed, varying from 15% to 50% and centering around 37% (Figure 2C), which was much higher than the sequence recapitulation rate obtained for the protein surface residues during the benchmarking of EvoEF2^18^. Although the peptide residues were considered to be highly exposed, the high sequence identity revealed that a large number of critical binding residues should be correctly predicted, indicating that the designed peptides are reasonable.

**Figure 2.**
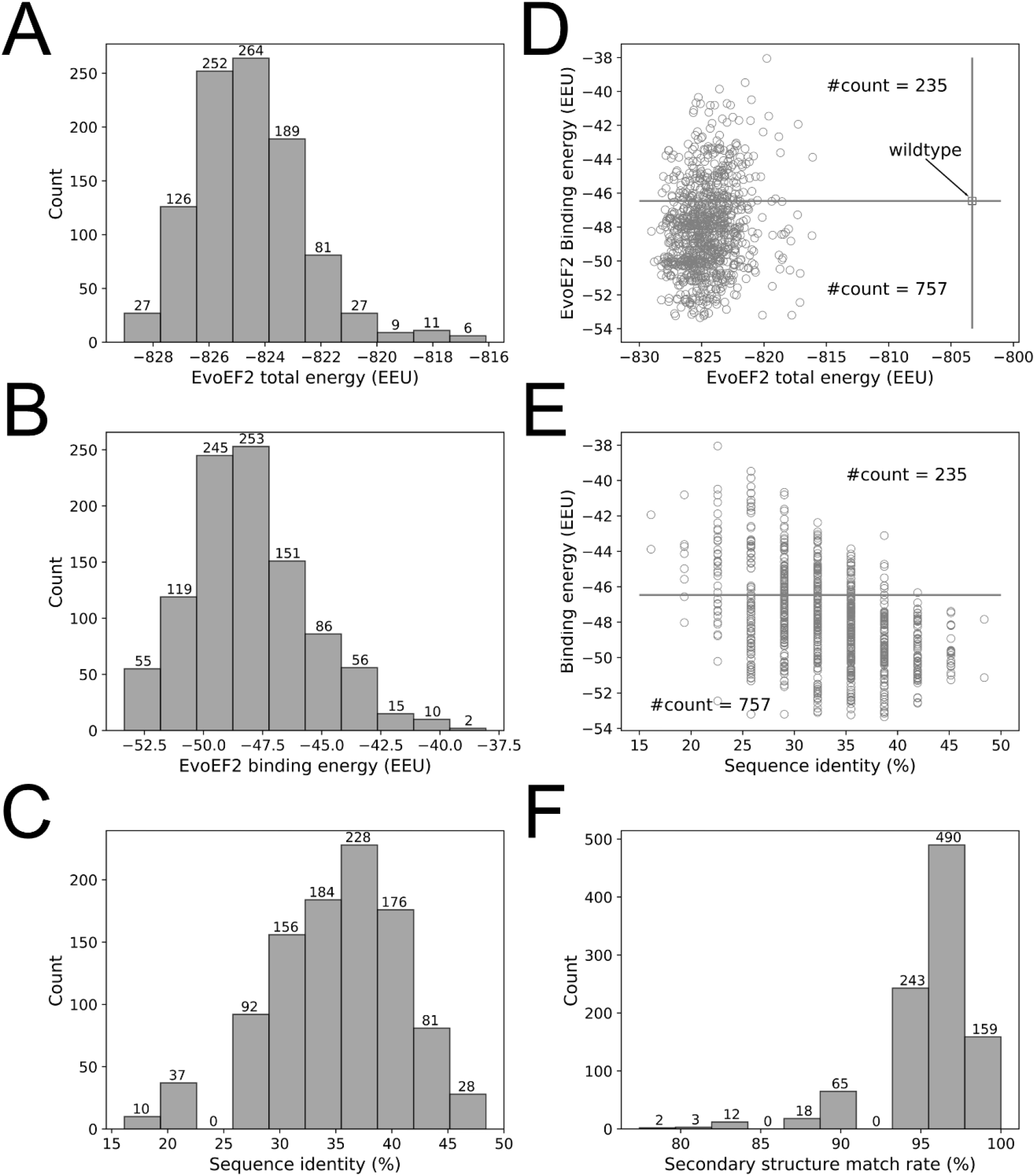
Overview of the characteristics of the EvoEF2 designs. (A) Distribution of total energy, (B) distribution of binding energy, (C) distribution of sequence identity, (D) binding energy as a function of total energy, (E) binding energy as a function of sequence identity, and (F) distribution of secondary structure match rate.

The wild-type peptide showed an EvoEF2 binding energy of −46.3 EEU, whereas the total energy of the wild-type peptide/SARS-CoV-2 RBD complex was −802 EEU (Figure 2D). 757 out of the 992 designs exhibited better binding affinities to SARS-CoV-2 RBD and showed lower total energies than the wild-type, and some designs showed good binding and stability simultaneously (Figure 2D), indicating that the wild-type peptide could be improved through design. Figure 2E illustrates the binding energy as a function of sequence identity for the designed peptides; it illustrates that a majority of the designs showed weaker binding affinity to SARS-CoV-2 than the wild-type peptide when the sequence identity was <25%, whereas most of the designs with sequence identities >35% exhibited stronger binding to SARS-CoV-2. These results suggest that, in general, low sequence identity designs may not be as good as high sequence identity designs. However, we can also see from Figure 2E that it does not necessarily mean that higher sequence identity always ensures better designs, since the two designs with the highest sequence identity (15/31=48.4%) did not always show stronger binding than those around 35%. Thus, the results suggest that good binders showed a high similarity to the wild-type, but the similarity should not be too high in order to leave room for the designs to be improved. This is in line with the common thinking that the critical binding residues (i.e. hot spot residues) should be conserved while some other residues can be mutated to enhance binding. Note that the wild-type peptide was comprised of a helix (a.a. 22-44) and a short loop (a.a. 351-357) with a glycine linker. To ensure good binding to SARS-CoV-2 RBD, the designed peptides should be able to preserve the secondary structure of this motif. To check this point, we used an artificial neural network-based secondary structure predictor^28^ implemented in EvoDesign to predict the secondary structure of the designed peptides; the predictor that we used here was much faster than some other state-of-the-art predictors, e.g. PSIPRED^35^ and PSSpred^36^, but showed similar performance^28^. To quantify the similarity between the secondary structure of a designed peptide and that of the wild-type, we calculated the secondary structure match rate, which was defined as the ratio of the number of residues with correctly assigned secondary structure elements (i.e. helix, strand and coil) to the total number of residues (i.e. 31). As shown in Figure 2F, 892 out of the 992 designed peptides had >90% secondary structure elements predicted to be identical to that of the wild-type peptide, indicating the high accuracy of the designs, although the EvoEF2 scoring function does not include any explicit secondary structure-related energy terms^18^.

We used WebLogo^37^ to perform a sequence logo analysis for the 992 designed sequences to investigate the residue substitutions and the results are shown in Figure 3A. 16 residues from the initial peptide scaffold were at the protein-peptide surface in contact with residues from SARS-CoV-2 RBD; these residues were Q24, T27, F28, D30, K31, H34, E35, E37, D38, F40, Y41, Q42, K353, G354, D355 and R357. Of these residues, Q24, D30, E35, E37, D38, Y41, Q42 and K353 formed hydrogen bonds or ion bridges with the binding partner (i.e. SARS-CoV-2 RBD) and the designed residues at these positions maintained favorable binding interactions. As shown in Figure 3A, the native residue types at these positions were top ranked out of all 20 canonical amino acids, suggesting that these residues may play critical roles in binding. For the nonpolar residues that were originally buried in the hACE2 structure (e.g. A25, L29, F32, L39, L351 and F356), they were likely to be mutated into polar or charged amino acids (Figure 3A), because they were largely exposed to the bulk solvent. The three glycine residues, including the one that was artificially introduced, were conserved probably due to the narrow space at these positions.

**Figure 3.**
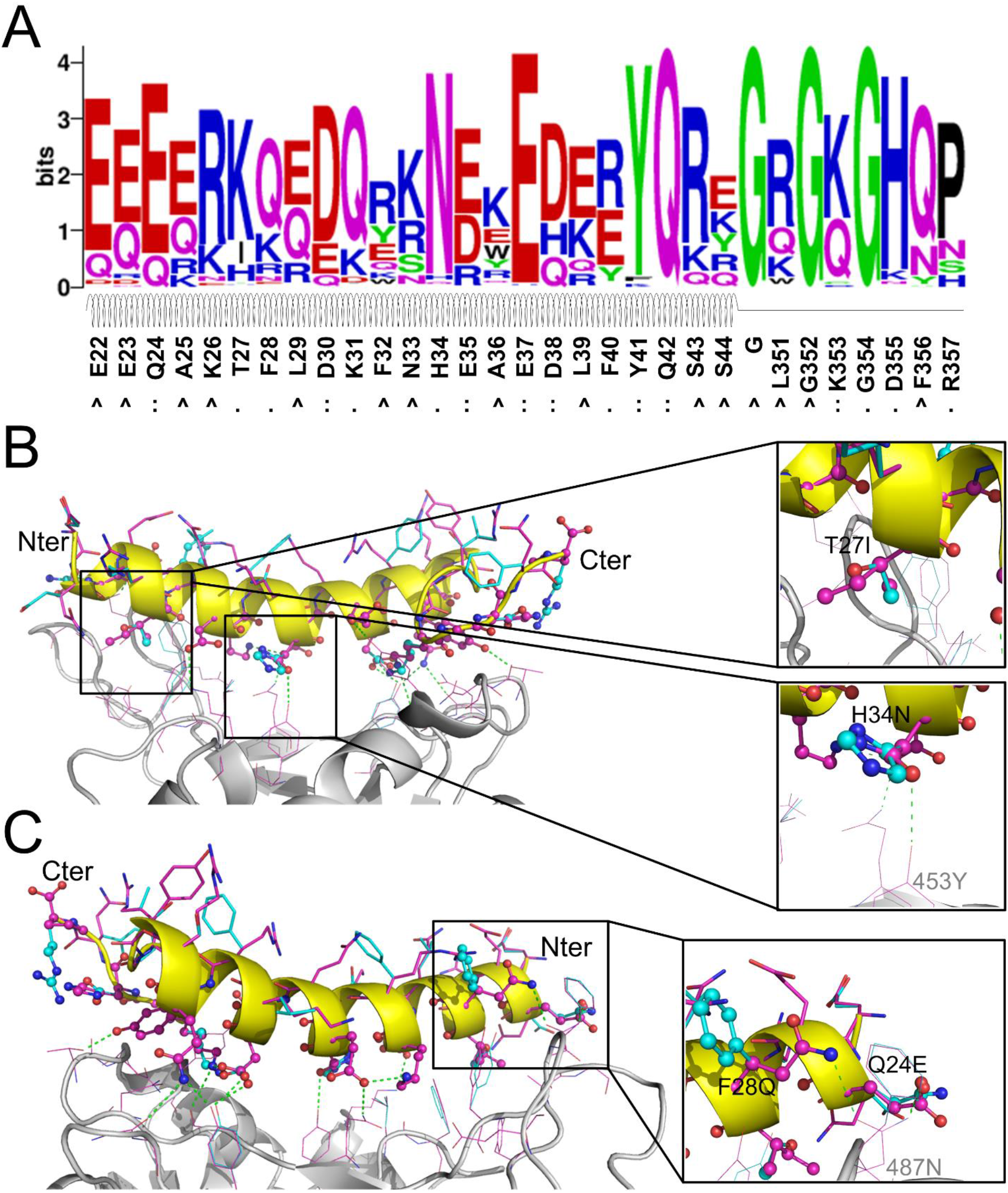
Sequence logo analysis of 992 unique peptide binders designed by EvoEF2 (A) and favorable interactions introduced in the top binder (B and C). In figure (A), the interface residues on the wild-type peptide are marked with ‘:’ if hydrogen bonds or ion bridges exist, or ‘.’ otherwise; non-interface residues are marked with ‘^’. In figures (B) and (C), the residues on the wild-type and designed structures are colored in cyan and magenta, respectively; interface and non-interface residues on the peptide are shown in ball-and-stick and stick models, respectively, while residues on SARS-CoV-2 RBD are shown in lines. Hydrogen bonds and/or ion bridges are shown using green-dashed lines.

To further examine what interactions improved the binding affinity of most designs, we carried out a detailed examination of some designed structures. We found that favorable hydrogen bonds or hydrophobic interactions were introduced in the binder that had the lowest EvoEF2 binding score (Figure 3B-C); the amino acid sequence of this binder was “EQEERIQQDKRKNEQEDKRYQRYGRGKGHQP”. For this design, T27 was mutated to isoleucine (Figure 3B). In the wild-type structure, the threonine was enveloped by four hydrophobic residues on SARS-CoV-2 RBD (i.e. Y489, F456, Y473 and A475), but its hydroxyl group did not form any hydrogen bond with the hydroxyl group of either Y489 or Y473, and the mutation enhanced the favorable burial of nonpolar groups. The interface residue H34 was substituted for asparagine (Figure 3B), introducing a hydrogen bond to Y453 on SARS-CoV-2 RBD. Additionally, two mutations, F28Q and Q24E, simultaneously formed hydrogen bonds with the amide group of N487 from SARS-CoV-2 RBD (Figure 3C). Although the mutation D355H did not form hydrogen bonds with any residues from SARS-CoV-2, it simultaneously formed two hydrogen bonds with the hydroxyl group of Y41 and the main-chain carbonyl group of G45 on the peptide, which may help stabilize the loop region (a.a. 351-357).

In previous studies, we found that evolutionary information can facilitate the design of proteins, improving their ability to fold into desired structures^28, 38^. To examine whether the evolutionary profile is important for peptide design here, we also performed four sets of designs with different weight settings for the evolution energy; for each design set, 1000 independent design simulation trajectories were carried out and the unique sequences out of the 1000 lowest energy designs were analyzed (Table 2). In general, giving a higher weight to the evolutionary energy facilitated the convergence of the design simulations, as indicated by the fewer unique designed sequences. It also helped identify sequences that were closer to the wild-type peptide as demonstrated by the higher sequence identities and the lower average evolutionary energy, which were both much more similar to those of the wild-type than the designs created using the physical score alone. We also found that incorporation of the profile energy moderately increased the ability of the designed sequences to maintain the original secondary structure. However, despite these improvements, giving a higher value to the profile weight clearly hindered the identification of binders that exhibited better binding energy than the wild-type.

**Table 2.** Summary of evolution-based peptide design results.

| Comparison items <sup>a</sup> | Weight of evolutionary profile energy |  |  |  |  |
| --- | --- | --- | --- | --- | --- |
|  | 0.00 | 0.25 | 0.50 | 0.75 | 1.00 |
| Number of unique designs | 992 | 991 | 966 | 877 | 695 |
| Number of better binders <sup>b</sup> | 757 | 636 | 392 | 340 | 226 |
| EvoEF2 binding energy | -48.1 ±2.5 | -47.2 ±2.2 | -46.1 ±1.7 | -45.8 ±1.6 | -45.5 ±1.6 |
| EvoEF2 total energy | -824.6 ±2.0 | -823.4 ±2.2 | -818.4 ±2.3 | -813.3 ±1.9 | -809.7 ±2.3 |
| Profile energy <sup>c</sup> | 6.7 ±2.7 | -0.8 ±3.6 | -13.3 ±3.3 | -21.6 ±2.0 | -25.6 ±1.6 |
| EvoEF2+profile energy | -824.6 ±2.0 | -823.6 ±1.8 | -825.0 ±1.4 | -829.5 ±1.2 | -835.3 ±1.4 |
| Sequence identity (%) | 33.7±5.6 | 39.1±5.5 | 44.2±5.1 | 46.2±5.6 | 48.3±6.0 |
| Sec. Str. match rate (%)<sup>d</sup> | 95.7±3.3 | 96.2±3.0 | 97.5±2.7 | 97.5±2.5 | 97.7±2.4 |

<sup>a</sup> The units for the EvoEF2 and profile energies is EEU. <sup>b</sup> The EvoEF2 binding energy of the wild-type peptide binder was -46.46 EEU; this row shows the number of designed peptide binders with EvoEF2 binding energies lower than -46.46 EEU. <sup>c</sup> The profile energy of wild-type peptide binder was -22.2 EEU.

We performed sequence logo analyses of the four sets of designs obtained from the evolution-based method and the results are illustrated in Figure 4. Overall, the evolutionary profile did not have a dramatic effect on most interface residues (e.g. Q24, K31, H34, E35, E37, D38, Y41, Q42 and K353), because the dominating residue types identified in the EvoEF2-based designs were also top ranked (Figure 3A and Figure 4). However, some interface residues were indeed influenced. For instance, T27 could be substituted for either lysine or isoleucine without evolution (Figure 3A), but it was only mutated to lysine when the evolutionary weight was ≥0.75 (Figure 4C-D). Additionally, without evolutionary profiles, F28 preferred glutamine over all other residues (Figure 3A), but it was conserved as phenylalanine when the evolutionary weight was ≥0.5 (Figure 4B-D). The naturally occurring residues, glutamic acid and arginine never appeared at positions 335 and 337, respectively, without evolutionary profile-guided design (Figure 3A); however, both of them were ranked second when a weight of 1.0 was given to the profiles. The residues that were most affected by evolution were those nonpolar residues that were not at the interface (e.g. A25, L29, F32, A36, L39, L351 and F356); without the evolutionary profile, polar or charged residue types were preferred at these positions (Figure 3A), while nonpolar residues were more frequently chosen for most of them when the weight of the profile energy was high (Figure 4B-D). As discussed above, most of these residues were buried in the original hACE2 structure, but they were solvent exposed in the peptide, and therefore it might not be necessary to maintain the hydrophobic nature at these positions.

**Figure 4.**
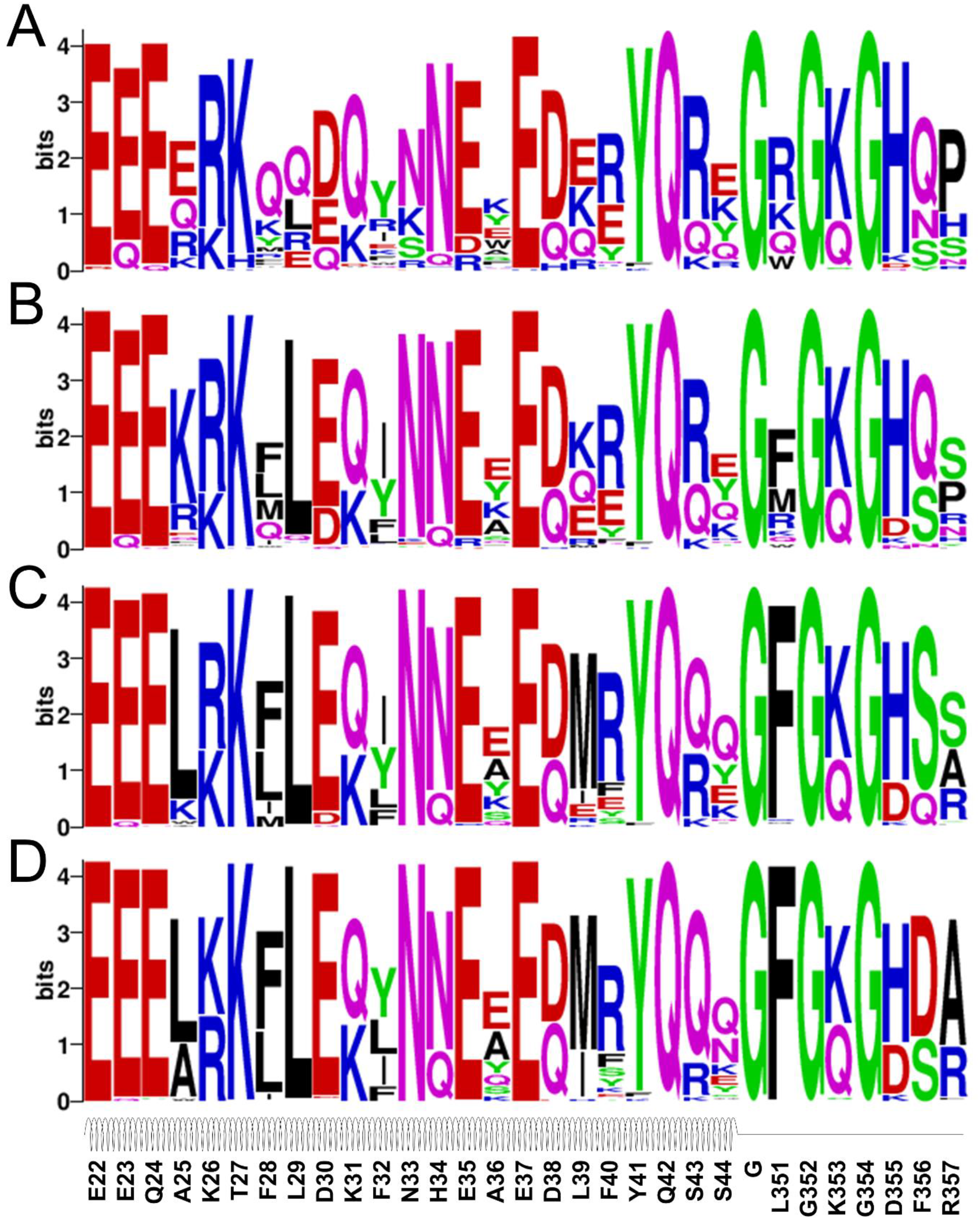
Sequence logo analysis of the evolution-based design results. Four sets of profile energy weight were used: 0.25 (A), 0.50 (B), 0.75 (C) and 1.00 (D).

In summary, detailed computational analyses showed that the designed peptide binders were quite reasonable, as indicated by the recapitulation of critical binding interactions at the protein-peptide interface and the introduction of new favorable binding interactions, as well as the preservation of secondary structures to maintain these interactions. We have selected dozens of designed peptides (with/without evolutionary information) that had good EvoEF2 binding energy to SARS-CoV-2 RBD to perform wet-lab experimental validation on. Specifically, we are carrying out in vitro experiments to determine their binding affinity and inhibitory potency, as well as in vivo experiments to examine their potential to disrupt the binding of SARS-CoV-2 to hACE2. Due to the urgent situation caused by COVID-19 worldwide, we would like to share our computational data to the scientific community so that researchers can work together to test them. Although we do not have experimental data yet, we believe that our computational results are well examined and reasonable, and these data may help favorably combat the COVID-19 pandemic.

## Conclusion

We designed a series of peptide binders that showed enhanced binding affinity to the SARS-CoV-2 RBD in computational experiments. Structural bioinformatics and sequence logo analyses indicated that the good binders to a high extent recapitulated the critical residues that contributed significantly to binding. Detailed inspection confirmed this point and also revealed that some extra favorable interactions were introduced to enhance binding in the top designed binders. Moreover, the designs were predicted to maintain the secondary structure of the motif, which was important for facilitating the protein-peptide binding interaction. Although our experimental results are not available yet, we would like to share these useful data to the community so that people can work together to test them, which should help to develop new antiviral drugs to combat COVID-19.

## Associated Content

### Supporting Information

The supporting Information is available free of charge on the ACS Publications website. Supporting Information Tables S1-S5 and Figure S1 as mentioned in the text.

## Author Information

### Corresponding Author

* Corresponding author: Yang Zhang, tel (734) 647-1549, fax (734) 615-6553

### Notes

The authors declare no competing financial interest.

## Acknowledgement

The work used the XSEDE clusters^39^ which is supported by the National Science Foundation (ACI-1548562).

## Abbreviations

ACE2: angiotensin-converting enzyme 2
COVID-19: coronavirus disease 2019
Cryo-EM: cryogenic electron microscopy
EEU: EvoEF2 energy unit
hACE2: human angiotensin-converting enzyme 2
MSA: multiple sequence alignment
NIL: non-redundant interface library
PDB: protein data bank
PPI: protein-protein interaction
PSSM: position specific scoring matrix
RBD: receptor binding domain
RBM: receptor binding motif
SAMC: simulated annealing Monte Carlo
SARS-CoV: severe acute respiratory syndrome coronavirus
SARS-CoV-2: severe acute respiratory syndrome coronavirus 2

---

For Table of Contents Only

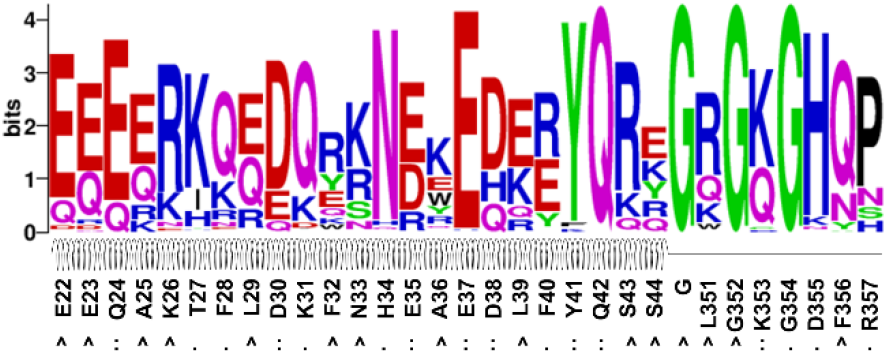

