## Supplementary for "Computational Design of Peptides to Block Binding of the SARS-CoV-2 Spike Protein to Human ACE2"

### Table of Contents

#### Supporting Tables

**Table S1**. Summary of 992 peptide sequences designed using EvoEF2 only.
**Table S2**. Summary of 991 peptide sequences designed using EvoEF2 and the evolutionary profile (weight = 0.25).
**Table S3**. Summary of 966 peptide sequences designed using EvoEF2 and the evolutionary profile (weight = 0.50).
**Table S4**. Summary of 877 peptide sequences designed using EvoEF2 and the evolutionary profile (weight = 0.75).
**Table S5**. Summary of 695 peptide sequences designed using EvoEF2 and the evolutionary profile (weight = 1.00).

#### Supporting Figures

**Figure S1**. Peptide multiple sequence alignment for evolutionary profile construction.

### Supporting Tables

**Table S1**. Summary of 992 peptide sequences designed using EvoEF2 only. The table is sorted according to binding score from the lowest to the highest. '#label' presents the index of the binder among the 1000 low total energy designs.

| #label | #peptide | #binding(EEU) | #SeqID(%) | #Secondary structure |
|---|---|---|---|---|
| WT | EEQAKTFLDKFNHEAEDLFYQSSGLGKGDFR | -46.46 | 100. | HHHHHHHHHHHHHHHHHHHHHHHCCCCCCCC |
| 728 | EQEERIQQDKRKNEQEDKRYQRYGRGKGHQP | -53.35 | 38.7 | CHHHHHHHHHHHHHHHHHHHHHHHCCCCCCC |
| 513 | EQQQRIQEDQYRNDWEDEEYQRKGRGKGHQP | -53.24 | 35.5 | HHHHHHHHHHHHCHHHHHHHHHHHCCCCCCC |
| 213 | EQQQRIQQDQDSNDREDKEYQKRGKGKGHNH | -53.24 | 35.5 | HHHHHHHHHHCCCCHHHHHHHHHHCCCCCCC |
| 830 | QEEQKIQEDQRRNDKEHKRKQRYGRGCGKQN | -53.20 | 25.8 | CHHHHHHHHHHHHHHHHHHHHHHCCCCCCCC |
| 138 | EQEERIQRDKRKNEKEHEEKQRRGRGCGKQN | -53.20 | 29.0 | CHHHHHHHHHHHHHHHHHHHHHHCCCCCCCC |
| 585 | EQEERIQQDKRKNENEDKRYQRYGRGKGHQP | -53.16 | 38.7 | CHHHHHHHHHHHHHHHHHHHHHHHCCCCCCC |
| 818 | EQEERIQQDKRKNEWEDQYYQKYGQGKGHQP | -53.16 | 38.7 | CHHHHHHHHHHHHHHHHHHHHHHHHCCCCCC |
| 111 | EQEERIQRDQYKNDYEDEEYQRKGRGKGHQP | -53.11 | 32.3 | CHHHHHHHHHHHCCCHHHHHHHHHCCCCCCC |
| 280 | EQEQRIQQDKRSNEQEDKRYQREGKGKGHNN | -53.09 | 38.7 | HHHHHHHHHHHHHHHHHHHHHHHCCCCCCCC |
| 406 | EQEERIQQDQRKNDKEDQRYQREGKGKGHNH | -53.04 | 32.3 | CHHHHHHHHHHHCHHHHHHHHHHHCCCCCCC |
| 297 | EQEQRIQEDKRKNEEEDKRYQRYGDGKGHNP | -53.03 | 38.7 | HHHHHHHHHHHHHHHHHHHHHHHCCCCCCCC |
| 909 | QDEKRIQEDKRKQEKEDEEYQRRGRGKGHQP | -52.90 | 35.5 | CHHHHHHHHHHHHHHHHHHHHHHHCCCCCCC |
| 794 | ENEERIQQDQRKNDKEDKRYQREGKGKGHQP | -52.86 | 32.3 | CHHHHHHHHHHHCHHHHHHHHHHCCCCCCCC |
| 384 | EQERRIQEDKERNEKEDEEYQRRGWGKGHYP | -52.80 | 38.7 | HHHHHHHHHHHHHHHHHHHHHHHCCCCCCCC |
| 643 | EEEKKIQEDQRKNDKEDEEYQRRGRGKGHQP | -52.74 | 38.7 | HHHHHHHHHHHHCCHHHHHHHHHHCCCCCCC |
| 369 | EQQQRIQEDQRRNDKEDRQYQQRGKGKGHNH | -52.61 | 35.5 | HHHHHHHHHHHHHHHHHHHHHHHHCCCCCCC |
| 659 | EQEERIQRDKWRHEWEDKYYQKKGQGKGHQP | -52.58 | 41.9 | CHHHHHHHHHHHHHHHHHHHHHHCCCCCCCC |
| 546 | EQEQRIQQDQYNNDWEDEEFQRKGRGKGKNN | -52.56 | 32.3 | HHHHHHHHHHCCCCHHHHHHHHHHCCCCCCC |
| 195 | EQEQRIQEDKRRNEKEDKRYQREGKGKGHNH | -52.54 | 38.7 | HHHHHHHHHHHHHHHHHHHHHHHCCCCCCCC |
| 950 | EQQQRIQQDQRNNEQEDQRYQREGRGKGHQP | -52.53 | 41.9 | HHHHHHHHHHHHHHHHHHHHHHHHCCCCCCC |
| 298 | EQEERIQRDQQKNEREDKRYQRYGRGKGHQP | -52.46 | 35.5 | HHHHHHHHHHHHHHHHHHHHHHHHCCCCCCC |
| 394 | EQEQRIQEDQRKNQKEHDRKQREGKGCGKNN | -52.45 | 22.6 | HHHHHHHHHHHHHHHHHHHHHHCCCCCCCCC |
| 017 | EQQQRIQEDQRKNRKEDERYQREGRGKGHQP | -52.44 | 35.5 | HHHHHHHHHHHHHHHHHHHHHHHHCCCCCCC |
| 770 | EQEQRIQQDQFSNDFEDKRYQQEGRGKGHQP | -52.43 | 35.5 | HHHHHHHHHHHHHHHHHHHHHHHHCCCCCCC |
| 235 | EQEQRIQQDQYNNDYEDEEYQRKGRGKGHQP | -52.43 | 35.5 | HHHHHHHHHHCCCCHHHHHHHHHHCCCCCCC |
| 910 | EQEERIQQDQRKNEKEDEEYQRRGRGKGHQP | -52.40 | 35.5 | CHHHHHHHHHHHHHHHHHHHHHHHCCCCCCC |
| 636 | EEEKRIQEDQRKNDKEDEEYQRYGRGKGHQP | -52.38 | 35.5 | HHHHHHHHHHHHCCHHHHHHHHHHHCCCCCC |
| 377 | QDQERINQDQQKNDKEDKRYQREGRGKGHQP | -52.38 | 32.3 | CHHHHHHHHHHHHHHHHHHHHHHHCCCCCCC |
| 382 | EQEQRIQEDQRKHEKEDQRYQREGRGKGHQP | -52.36 | 38.7 | HHHHHHHHHHHHHHHHHHHHHHHHCCCCCCC |
| 639 | EQEKRIQEDQRKNREEDERYQREGRGKGHQP | -52.31 | 32.3 | HHHHHHHHHHHHHHHHHHHHHHHHCCCCCCC |
| 648 | EQEERIQQDQQKNRKEDEEYQRRGRGKGHQP | -52.30 | 32.3 | CHHHHHHHHHHHHHHHHHHHHHHHCCCCCCC |
| 050 | EQEQRIQEDQRKNDKEDRRYQREGRGKGHQP | -52.29 | 32.3 | HHHHHHHHHHHHCHHHHHHHHHHHCCCCCCC |

| | | | | |
|---|---|---|---|---|
| 361 | EQEERIKQDKQNNEREDKRYQREGKGKGHNN | -52.29 | 41.9 | CHHHHHHHHHHHHHHHHHHHHHHCCCCCCCC |
| 658 | EQEERIQRDQRKNRKEDEEYQKQGRGKGHQP | -52.28 | 32.3 | CHHHHHHHHHHHHHHHHHHHHHHCCCCCCCC |
| 519 | EQEERIQQDQYKNEWEDEEYQRKGRGKGHQP | -52.27 | 35.5 | CHHHHHHHHHHHHHHHHHHHHHHHCCCCCCC |
| 831 | EQEERIQRDQYKNEHEDERYQREGWGKGHYP | -52.27 | 35.5 | CHHHHHHHHHHHHHHHHHHHHHHHCCCCCCC |
| 483 | EQEKRIQEDQRKNEEEDKRYQRYGQGKGHQS | -52.27 | 35.5 | HHHHHHHHHHHHHHHHHHHHHHHHHCCCCCC |
| 542 | QEEEKIQRDQQRHEREDKRYQREGRGKGHQP | -52.26 | 41.9 | CHHHHHHHHHHHHHHHHHHHHHHHCCCCCCC |
| 669 | EQEKRIQEDQRSHEEEDEYYQQYGQGKGHQP | -52.24 | 38.7 | HHHHHHHHHHHHHHHHHHHHHHHHHCCCCCC |
| 903 | EQQQRIQQDQYSNEWEDEEYQRKGRGKGHQP | -52.23 | 38.7 | HHHHHHHHHHCCCHHHHHHHHHHHCCCCCCC |
| 411 | EQEERIQQDQRKNEKEDEEYQRYGRGKGHQP | -52.21 | 35.5 | CHHHHHHHHHHHHHHHHHHHHHHHHCCCCCC |
| 868 | EQEERIQQDQRKNEWEDKYYQKKGQGKGHQP | -52.21 | 35.5 | CHHHHHHHHHHHHHHHHHHHHHHHCCCCCCC |
| 992 | EQEERIQQDQRKNEYEDEEYQRKGRGKGHTP | -52.21 | 35.5 | CHHHHHHHHHHHHHHHHHHHHHHCCCCCCCC |
| 771 | EQEQRIQEDQRKNEKEDKRYQEQGRGKGHQP | -52.21 | 35.5 | HHHHHHHHHHHHHHHHHHHHHHHHCCCCCCC |
| 370 | EQEERIQQDQRKNDNEDRRYQREGDGKGHQP | -52.10 | 32.3 | CHHHHHHHHHHHCHHHHHHHHHHCCCCCCCC |
| 901 | EQEERIKQDKYSNEWEDKKYQREGRGKGHQP | -52.10 | 38.7 | CHHHHHHHHCCCHHHHHHHHHHHHCCCCCCC |
| 108 | EQEKRIQEDQQKNEREDKRYQREGRGKGHQP | -52.09 | 35.5 | HHHHHHHHHHHHHHHHHHHHHHHHCCCCCCC |
| 821 | KREEEIQQDQRRNEQEDQRYQKEGQGKGHQS | -52.02 | 32.3 | CHHHHHHHHHHHHHHHHHHHHHHHCCCCCCC |
| 962 | EQEERIKEDQKRNDEEDKRYQRYGRGKGHQP | -51.98 | 32.3 | HHHHHHHHHHHHHHHHHHHHHHHHCCCCCCC |
| 217 | QREQEIQQDQYNNEWEDERYQRQGRGKGHQP | -51.98 | 35.5 | CHHHHHHHHHCHHHHHHHHHHHHHCCCCCCC |
| 044 | EQQERINQDQQRNEKEDERYQRQGRGKGHQP | -51.98 | 38.7 | HHHHHHHHHHHHHHHHHHHHHHHCCCCCCCC |
| 980 | EEEKKIQEDQRKNDEEDRRYQREGRGKGHQP | -51.97 | 38.7 | HHHHHHHHHHHHCHHHHHHHHHHHCCCCCCC |
| 863 | DQERRIQEDQWRNEYEDEEYQRQGRGKGHQP | -51.90 | 32.3 | HHHHHHHHHHHHHHHHHHHHHHHHCCCCCCC |
| 339 | QDQKRIQEDQRKNREEDEYYQQKGQGKGHQP | -51.88 | 32.3 | CHHHHHHHHHHHHHHHHHHHHHHHCCCCCCC |
| 997 | SDQERIQRDQYKNEYEDEEYQRKGRGKGHQP | -51.84 | 35.5 | CHHHHHHHHHHHHHHHHHHHHHHHCCCCCCC |
| 738 | KREEEIKQDQYNNDYEDKEYQKEGRGKGHQP | -51.80 | 32.3 | CHHHHHHHHCCCCHHCHHHHHHHHCCCCCCC |
| 796 | QDEERIQQDQRKNEYEDEEYQRKGRGKGHQP | -51.78 | 32.3 | CHHHHHHHHHHHHHHHHHHHHHHHCCCCCCC |
| 201 | DEERKIQEDQRRNEKEDEEYQRYGRGKGHQP | -51.67 | 38.7 | HHHHHHHHHHHHHHHHHHHHHHHHHCCCCCC |
| 469 | QDEERIKEDKYRNESEDQRYQRYGRGKGHQP | -51.67 | 35.5 | CHHHHHHHHHCCCHHHHHHHHHHHCCCCCCC |
| 827 | QQEENIQRDQYKNEWEDERYQRQGRGKGHQP | -51.67 | 32.3 | CHHHHHHHHHHHHHHHHHHHHHHHCCCCCCC |
| 447 | QDEKRIQEDQRKNEEEDQQYQKQGRGKGHQP | -51.65 | 32.3 | CHHHHHHHHHHHHHHHHHHHHHHHCCCCCCC |
| 522 | EQQQRIQEDQRRNDKEHKYYQQKGQGQGHQP | -51.65 | 29.0 | HHHHHHHHHHHHHHHHHHHHHHHHCCCCCCC |
| 840 | EQEQRIQQDQKNNEEEDERYQKQGRGKGHQP | -51.60 | 38.7 | HHHHHHHHHHHHHHHHHHHHHHHCCCCCCCC |
| 126 | EQEERIKQDKMKNEKEDEEYQRRGQGKGHQS | -51.58 | 38.7 | CHHHHHHHHHHHHHHHHHHHHHHHHCCCCCC |
| 252 | QDEERIKQDQESNDREDKRYQREGRGKGHQP | -51.55 | 29.0 | CHHHHHHHHHHHHHHHHHHHHHHHCCCCCCC |
| 055 | QDQERIKKDQYDNDWEDERYQQQGRGKGHQP | -51.54 | 32.3 | CHHHHHHHHCCCHHHHHHHHHHHHCCCCCCC |
| 822 | QRQQEIQEDAWRHRYEDEEFQRKGRGKGKQN | -51.50 | 32.3 | HHHHHHHHHHHHHHHHHHHHHHHHCCCCCCC |
| 092 | EEEQRKQEDQRRQDKEDQRYQREGRGKGHQP | -51.44 | 35.5 | HHHHHHHHHHHHHHHHHHHHHHHHCCCCCCC |
| 899 | EEEERKQQDKRKQEKEDEEYQRRGKGKGHNH | -51.43 | 41.9 | HHHHHHHHHHHHHHHHHHHHHHHHCCCCCCC |
| 536 | EQEQRIQQDQKNNEEEDKRYQREGRGKGHQP | -51.42 | 38.7 | HHHHHHHHHHHHHHHHHHHHHHHHCCCCCCC |

| | | | | |
|---|---|---|---|---|
| 250 | KRQQEIQQDQYNNDYEHEEYQRKGRGQGHQP | -51.34 | 29.0 | CHHHHHHHHHCCCHHHHHHHHHHHCCCCCCC |
| 976 | EQEERIQQDQRKNDKEHEEYQRRGWGQGHYP | -51.32 | 25.8 | CHHHHHHHHHHHHHHHHHHHHHHCCCCCCCC |
| 779 | EQEERIQQDQRKNDKEHEYYQQYGQGQGHQP | -51.32 | 25.8 | CHHHHHHHHHHHHHHHHHHHHHHHHCCCCCC |
| 565 | EQEERIQRDQRKNDQEHKRYQREGRGQGHQP | -51.32 | 25.8 | CHHHHHHHHHHHHHHHHHHHHHHHCCCCCCC |
| 854 | EQQQRKQEDQRRNDKEHKRYQREGRGRGHQP | -51.31 | 29.0 | HHHHHHHHHHHHHHHHHHHHHHHHCCCCCCC |
| 227 | EEEERKQQDKRQNEREDKYYQQKGQGKGHQP | -51.31 | 41.9 | HHHHHHHHHHHHHHHHHHHHHHHHCCCCCCC |
| 037 | EEQQRKQEDKRRNEKEDEYYQRKGQGKGHQP | -51.26 | 45.2 | HHHHHHHHHHHHHHHHHHHHHHHHCCCCCCC |
| 591 | EEEQRKQEDKRRNEKEDKYYQKYGQGKGHQP | -51.26 | 41.9 | HHHHHHHHHHHHHHHHHHHHHHHHHCCCCCC |
| 458 | QEEQKKQEDQRRHDWEDSYYQQYGQGKGHQP | -51.24 | 38.7 | HHHHHHHHHHHHHHHHHHHHHHHHHCCCCCC |
| 266 | EEEERKQRDQYKNDWEDEEYQRKGRGKGTNN | -51.21 | 35.5 | HHHHHHHHHHHHHHHHHHHHHHHHCCCCCCC |
| 077 | EEEERKQRDKQKNEKEDKRYQREGRGKGHQP | -51.20 | 41.9 | HHHHHHHHHHHHHHHHHHHHHHHHCCCCCCC |
| 020 | EQEERKQQDKQKNEREDKRYQREGKGKGHNH | -51.19 | 38.7 | HHHHHHHHHHHHHHHHHHHHHHHCCCCCCCC |
| 967 | EQEERIKQDQYNNRYEDEEYQRQGQGKGHQS | -51.18 | 35.5 | CHHHHHHHHHHCCCHHHHHHHHHHCCCCCCC |
| 961 | QREQEIQERQRRERKEDERYQREGQGKGHQS | -51.17 | 25.8 | HHHHHHHHHHHHHHHHHHHHHHHHCCCCCCC |
| 549 | EQEERIKQDKYNNEYEDEEYQRKGRGKGNQN | -51.16 | 41.9 | CHHHHHHHHHCCCCCHHHHHHHHHCCCCCCC |
| 134 | EQEERIKQDQEKNEKEDEEYQRRGRGKGHQP | -51.15 | 35.5 | CHHHHHHHHHHHHHHHHHHHHHHHCCCCCCC |
| 768 | QQEENIKQDQKSNEEEDEYYQRKGQGKGHQP | -51.15 | 32.3 | CHHHHHHHHHHHHHHHHHHHHHHHCCCCCCC |
| 989 | EQEERIQQDQRKNDKEHQRYQKQGRGQGHQP | -51.14 | 25.8 | CHHHHHHHHHHHHHHHHHHHHHHHCCCCCCC |
| 819 | EQEERIKEDQYRNEWEDRRYQREGRGKGHQP | -51.14 | 35.5 | HHHHHHHHHHHHHHHHHHHHHHHHCCCCCCC |
| 764 | EEQQRKQQDKYNNEWEDKKYQREGRGKGHNH | -51.13 | 48.4 | HHHHHHHHHHHHHHHHHHHHHHHHCCCCCCC |
| 883 | EEEERKQRDKRKNEQEDKRYQREGKGKGHNH | -51.13 | 41.9 | HHHHHHHHHHHHHHHHHHHHHHHCCCCCCCC |
| 699 | QEEEKKQQDKRKNENEDKRYQREGRGKGHQP | -51.13 | 41.9 | CHHHHHHHHHHHHHHHHHHHHHHHCCCCCCC |
| 086 | QEEQKKQEDKRKNEKEDKRYQKQGWGKGHYP | -51.13 | 41.9 | HHHHHHHHHHHHHHHHHHHHHHHCCCCCCCC |
| 208 | EEERRKQEDQERNDKEDKRYQREGRGKGHQP | -51.07 | 35.5 | HHHHHHHHHHHHHHHHHHHHHHHHCCCCCCC |
| 448 | QEEQKKQYDQEKNDKEDEEYQRKGRGKGHQP | -51.07 | 35.5 | HHHHHHHHHHHHHHHHHHHHHHHHCCCCCCC |
| 142 | EEERKKQEDQYRNDAEHKRYQREGKGRGHNH | -51.05 | 35.5 | HHHHHHHHHHHHHHHHHHHHHHHCCCCCCCC |
| 047 | EEQQRKQEDQRRNDQEDKRYQRYGRGKGHQP | -51.02 | 38.7 | HHHHHHHHHHHHHHHHHHHHHHHHCCCCCCC |
| 313 | EQQERKQQDQRRNDKEDEEYQRRGQGKGHNP | -51.02 | 35.5 | HHHHHHHHHHHHHHHHHHHHHHHHCCCCCCC |
| 485 | QEQQKKQEDQRKNDKEDKRYQREGRGKGHQP | -51.02 | 38.7 | HHHHHHHHHHHHHHHHHHHHHHHHCCCCCCC |
| 757 | EEEQRKQEDQRKNDREDKYYQREGKGKGHNH | -51.01 | 35.5 | HHHHHHHHHHHHHHHHHHHHHHCCCCCCCCC |
| 043 | EEEEKKQRDKQKNEREDEYYQQYGQGKGHQP | -51.00 | 45.2 | HHHHHHHHHHHHHHHHHHHHHHHHHCCCCCC |
| 388 | EEEQRKQEDKQKNEREDEEYQRRGKGKGHNN | -51.00 | 41.9 | HHHHHHHHHHHHHHHHHHHHHHHCCCCCCCC |
| 189 | QRQQEINEDQRKNEKEDEEYQRRGRGKGHQP | -50.99 | 35.5 | CHHHHCHHHHHHHHHHHHHHHHHHCCCCCCC |
| 200 | QQEEEIKQDQQNNEKEDEEYQRYGQGKGHQS | -50.97 | 35.5 | CHHHHHHHHHHHHHHHHHHHHHHHHCCCCCC |
| 982 | DEERKKQEDKQRNEKEDKRYQREGKGKGHNH | -50.97 | 41.9 | CHHHHHHHHHHHHHHHHHHHHHHCCCCCCCC |
| 988 | DDQRRIQEDQYRNDYEHEEYQRKGRGQGHQP | -50.96 | 25.8 | CHHHHHHHHHHHHHHHHHHHHHHHCCCCCCC |
| 707 | DQERKKQEDKRRNEYEDEEYQRQGRGKGHQP | -50.94 | 38.7 | HHHHHHHHHHHHHHHHHHHHHHHHCCCCCCC |
| 709 | EEEERKQQDKRKNEKEDEEYQRYGRGKGHQP | -50.94 | 41.9 | HHHHHHHHHHHHHHHHHHHHHHHHHCCCCCC |

| | | | | |
|---|---|---|---|---|
| 914 | EEEERKQQDKRKNENEDKRYQREGKGKGHNH | -50.94 | 41.9 | HHHHHHHHHHHHHHHHHHHHHHHCCCCCCCC |
| 759 | EEEQRKQEDKRKNEKEDKRYQRYGQGKGHQS | -50.94 | 41.9 | HHHHHHHHHHHHHHHHHHHHHHHHCCCCCCC |
| 824 | EEEQRKQEDKRKNENEDKSYQRHGQGKGHQS | -50.94 | 41.9 | HHHHHHHHHHHHHHHHHHHHHHHCCCCCCCC |
| 693 | EQQERINQDQQKNDKEHEEYQRRGRGQGHQP | -50.91 | 29.0 | HHHHHHHHHHHHHHHHHHHHHHHHHCCCCCC |
| 629 | QEEEKKNQDQQKNDKEDEEYQRYGQGKGHQS | -50.89 | 35.5 | CHHHHHHHHHHHCHHHHHHHHHHHHCCCCCC |
| 413 | EEEKKKQEDQRKNDEEDEEYQRKGRGKGHQP | -50.89 | 38.7 | HHHHHHHHHHHHHHHHHHHHHHHHCCCCCCC |
| 972 | EEEKRKQEDQRKNDEEDERYQKEGKGKGHNH | -50.89 | 35.5 | HHHHHHHHHHHHHHHHHHHHHHHCCCCCCCC |
| 265 | EEEERKQRDQEKNDKEDEKYQREGRGKGHQP | -50.88 | 35.5 | HHHHHHHHHHHHHHHHHHHHHHHHCCCCCCC |
| 900 | EEEQKKQEDKRRNEKEDEEYQRRGWGKGHYP | -50.87 | 45.2 | HHHHHHHHHHHHHHHHHHHHHHHCCCCCCCC |
| 479 | QEEQKKQQDKRSNEFEDKYYQQKGQGKGHNH | -50.87 | 41.9 | HHHHHHHHHHHHHHHHHHHHHHHHCCCCCCC |
| 871 | QQQQNIQEDQRRNRKEHEYYQKKGQGQGHQP | -50.84 | 25.8 | HHHHHHHHHHHHHHHHHHHHHHHHCCCCCCC |
| 434 | EEEQRKNEDQRKNDKEDEEYQRRGRGKGHQP | -50.83 | 35.5 | HHHHHHHHHHHHCHHHHHHHHHHHCCCCCCC |
| 347 | EQEERIKQDQWKNEWEDEYYQRKGQGKGHQP | -50.82 | 35.5 | CHHHHHHHHHHHHHHHHHHHHHHHCCCCCCC |
| 562 | EEEEKKQQDQRKNDNEDKRYQREGQGKGHQS | -50.82 | 38.7 | CHHHHHHHHHHHHHHHHHHHHHHHCCCCCCC |
| 686 | EEEERKQQDQRKNDKEDEEYQRYGKGKGHNH | -50.82 | 35.5 | HHHHHHHHHHHHHHHHHHHHHHHHHCCCCCC |
| 038 | EEEERKQQDQRKNDKEDQRYQQQGRGKGHQP | -50.82 | 35.5 | HHHHHHHHHHHHHHHHHHHHHHHHCCCCCCC |
| 527 | EEEERKQQDQRKNDNEDERYQQQGRGKGHQP | -50.82 | 35.5 | HHHHHHHHHHHHHCHHHHHHHHHHCCCCCCC |
| 570 | EEEERKQQDQRKNDNEDKRYQREGRGKGHQP | -50.82 | 35.5 | HHHHHHHHHHHHHHHHHHHHHHHHCCCCCCC |
| 104 | EEEERKQQDQRRNDQEDERYQKQGRGKGHQP | -50.82 | 35.5 | HHHHHHHHHHHHHHHHHHHHHHHCCCCCCCC |
| 254 | EEEERKQQDQRRNDQEDKRYQREGRGKGHQP | -50.82 | 35.5 | HHHHHHHHHHHHHHHHHHHHHHHHCCCCCCC |
| 778 | QEEEKKQQDQRKNDWEDKYYQKKGQGKGHQP | -50.82 | 35.5 | CHHHHHHHHHHHHHHHHHHHHHHCCCCCCCC |
| 315 | QEEKKKQEDKRKNEEEDEQYQRYGRGKGHQP | -50.81 | 41.9 | HHHHHHHHHHHHHHHHHHHHHHHHHCCCCCC |
| 416 | NDQRRKQEDKERNEKEDKRYQREGRGKGHQP | -50.80 | 38.7 | CHHHHHHHHHHHHHHHHHHHHHHHCCCCCCC |
| 109 | EEEERKQQDQDKNDREHEEYQRYGRGRGHQP | -50.79 | 29.0 | HHHHHHHHHHHHHHHHHHHHHHHHHCCCCCC |
| 019 | EEEERKQQDQRKNDKEDKRYQREGRGKGHQP | -50.75 | 35.5 | HHHHHHHHHHHHHHHHHHHHHHHHCCCCCCC |
| 893 | EEEEKKQRDQWKNDWEHEYKQKYGQGCGKQN | -50.74 | 29.0 | HHHHHHHHHHHHHHHHHHHHHHHHHCCCCCC |
| 554 | QDEERKQQDKRRNEYEDKQYQKEGRGKGHQP | -50.73 | 35.5 | CHHHHHHHHHHHHHHHHHHHHHHHCCCCCCC |
| 581 | QDQERIQQDQRKNEYEDEEYQRKGRGKGDQR | -50.73 | 41.9 | CHHHHHHHHHHHHHHHHHHHHHHCCCCCCCC |
| 960 | QDEERKQRDKYKNEYEDEEFQRKGRGKGKNN | -50.72 | 32.3 | CHHHHHHHHHHHHHHHHHHHHHHCCCCCCCC |
| 344 | QQQQNINEDQRKNDKEHEEYQRRGWGQGHYP | -50.72 | 25.8 | CHHHHCHHHHHHHHHHHHHHHHHCCCCCCCC |
| 859 | EEEERKQQDKRKNEKEDEEYQRRGRGKGHQP | -50.70 | 41.9 | HHHHHHHHHHHHHHHHHHHHHHHHCCCCCCC |
| 396 | EEQQRKQQDQRSNDKEDREFQRRGQGKGKNN | -50.69 | 35.5 | HHHHHHHHHHHHCHHHHHHHHHHHCCCCCCC |
| 739 | DEERKKQEDQWRNDWEDEYYQRKGQGKGHQP | -50.69 | 35.5 | HHHHHHHHHHHHHHHHHHHHHHHHCCCCCCC |
| 516 | EEEERKQQDQRKNDYEHEEKQRKGRGCGKQN | -50.67 | 25.8 | HHHHHHHHHHHHHHHHHHHHHHHCCCCCCCC |
| 015 | EEEKRKQEDQRKNDKEDEEYQRRGKGKGHNN | -50.65 | 35.5 | HHHHHHHHHHHHHHHHHHHHHHHCCCCCCCC |
| 550 | EQEQRKQEDQRRNEKEDEEFQRRGKGKGKNN | -50.62 | 32.3 | HHHHHHHHHHHHHHHHHHHHHHHHCCCCCCC |
| 736 | NDQERIKQDQKKNEKEDEEYQRRGKGKGHQH | -50.59 | 35.5 | CHHHHHHHHHHHHHHHHHHHHHHHCCCCCCC |
| 751 | QREQEIQEQKRRNEKEDKRYQRYGRGKGHQP | -50.58 | 32.3 | CHHHHHHHHHHHHHHHHHHHHHHHCCCCCCC |

| | | | | |
|---|---|---|---|---|
| 168 | EQQQRIQQDQDSNDREQEEYQRRGRGQGHQP | -50.58 | 29.0 | HHHHHHHHHHHCHHHHHHHHHHHHHCCCCCC |
| 947 | EQEQRIQEDQRKNDKEQKRYQREGRGQGHQP | -50.57 | 25.8 | HHHHHHHHHHHHHHHHHHHHHHHHCCCCCCC |
| 462 | EEEERKQQDQRRHDKEHEEKQRRGRGCGKQN | -50.56 | 29.0 | HHHHHHHHHHHHHHHHHHHHHHHCCCCCCCC |
| 656 | SDEERIKEDQKRNDEEDRRYQRYGRGKGHQP | -50.55 | 29.0 | CHHHHHHHHHHHCHHHHHHHHHHHCCCCCCC |
| 318 | QEEKKKQEDQRKNDEEHQRKQKQGKGCGKNN | -50.54 | 25.8 | HHHHHHHHHHHHHHHHHHHHHHHCCCCCCCC |
| 944 | EQQQRKNNDKYNNEWEDEEYQRKGRGKGHQP | -50.51 | 45.2 | HHHHHHHCHHCCCHHHHHHHHHHHCCCCCCC |
| 844 | SREEQIQREQQSNEREDKRYQREGRGKGHQP | -50.50 | 29.0 | CHHHHHHHHHHHHHHHHHHHHHHHCCCCCCC |
| 497 | EQQERKQQDKRKNEKEDEEYQRRGRGKGHQP | -50.50 | 41.9 | HHHHHHHHHHHHHHHHHHHHHHHHCCCCCCC |
| 898 | EEEQRKQQDQYNNDYEHEEKQRKGRGCGKQN | -50.48 | 29.0 | HHHHHHHHHHHHHHHHHHHHHHHCCCCCCCC |
| 880 | QRQKEIQEQQRSNDEEDEYYQQYGQGKGHQP | -50.46 | 29.0 | HHHHHHHHHHHHCHHHHHHHHHHHHCCCCCC |
| 033 | EEEERKQQDKRRHEYEDQEYQRQGRGKGHQP | -50.45 | 45.2 | HHHHHHHHHHHHHHHHHHHHHHHHCCCCCCC |
| 177 | DNERRIQEEQYRNDWEDERYQRQGKGKGHSN | -50.44 | 25.8 | CHHHHHHHHHHHHHHHHHHHHHCCCCCCCCC |
| 867 | EEEERKQQDKRRNEWEDKYYQKKGQGKGHQP | -50.42 | 41.9 | HHHHHHHHHHHHHHHHHHHHHHHHCCCCCCC |
| 012 | EEEQRKQQDQYNQDWEDERYQRQGWGKGHYP | -50.41 | 38.7 | HHHHHHHHHHHHHHHHHHHHHHHHCCCCCCC |
| 572 | EEEERKQRDKWKNEWEDKYYQKKGQGKGHQP | -50.39 | 41.9 | HHHHHHHHHHHHHHHHHHHHHHHHCCCCCCC |
| 107 | EEEQRKQEDKYRNEYEDKEYQRKGRGKGHQP | -50.39 | 41.9 | HHHHHHHHHHHHHHHHHHHHHHHCCCCCCCC |
| 641 | KRQEEIQRDQRKNDQEQQRYQREGKGQGHNH | -50.39 | 25.8 | CHHHHHHHHHHHHHHHHHHHHHHHCCCCCCC |
| 123 | QEQQKKQEDQRRNDKEDRRYQRYGKGKGHNN | -50.38 | 38.7 | HHHHHHHHHHHHHHHHHHHHHHCCCCCCCCC |
| 968 | EEEQRKQQDKYNNEWEDKKYQREGRGKGHQP | -50.33 | 45.2 | HHHHHHHHHHHHHHHHHHHHHHHHCCCCCCC |
| 223 | EEEQRKQQDKYNNEYEDEEYQRKGRGKGHQP | -50.33 | 45.2 | HHHHHHHHHHHHHHHHHHHHHHHHCCCCCCC |
| 713 | EQEQRIQEEQYRNEYEDEEYQRKGRGKGHQP | -50.32 | 32.3 | HHHHHHHHHHHHHHHHHHHHHHHHCCCCCCC |
| 084 | EEEERKQQDQQKNDREDRRYQRYGRGKGHQP | -50.31 | 35.5 | HHHHHHHHHHHHHHHHHHHHHHHHCCCCCCC |
| 537 | EEQERKQRDQEKNDKEDRRYQREGRGKGHQP | -50.31 | 38.7 | HHHHHHHHHHHHHHHHHHHHHHHHCCCCCCC |
| 955 | EEQQRKQQDQRNNEKEDEEYQRRGRGKGHQP | -50.31 | 45.2 | HHHHHHHHHHHHHHHHHHHHHHHHCCCCCCC |
| 427 | EEQQRKQQDQRSNEQEDKRYQKEGQGKGHQS | -50.31 | 41.9 | HHHHHHHHHHHHHHHHHHHHHHHCCCCCCCC |
| 026 | EQQQRKQEDQRRNEKEDEEYQRKGRGKGHQP | -50.31 | 38.7 | HHHHHHHHHHHHHHHHHHHHHHHHCCCCCCC |
| 920 | EEERRKQEDQERNDKEDRRYQREGKGKGHNN | -50.31 | 35.5 | HHHHHHHHHHHHHHHHHHHHHHHCCCCCCCC |
| 586 | ENEQRIQEDQRKNEKEHKRYQREGRGQGHQP | -50.31 | 29.0 | CHHHHHHHHHHHHHHHHHHHHHHHCCCCCCC |
| 174 | EQERRKQEDQWRNDYEDEEYQRKGRGKGHQP | -50.28 | 32.3 | HHHHHHHHHHHHHHHHHHHHHHHHCCCCCCC |
| 428 | EEQQRKQYDQESNRKEDERYQREGRGKGHQP | -50.28 | 38.7 | HHHHHHHHHHHHHHHHHHHHHHHHCCCCCCC |
| 133 | EQEERIKQDQKNNDEEHERYQQQGRGQGHQP | -50.26 | 29.0 | CHHHHHHHHHHCHHHHHHHHHHHHCCCCCCC |
| 457 | EEQQRKQEDQWRNDYEDREYQRKGRGKGHQP | -50.26 | 38.7 | HHHHHHHHHHHHHHHHHHHHHHHHCCCCCCC |
| 352 | EEQERKQQDQRKNDYEDREYQRKGRGKGHQP | -50.26 | 38.7 | HHHHHHHHHHHHHHHHHHHHHHHHCCCCCCC |
| 373 | EEEQRKQEDQRKNDKEDRRYQREGRGKGHQP | -50.25 | 35.5 | HHHHHHHHHHHHHHHHHHHHHHHHCCCCCCC |
| 990 | QREEQIQREQRSNEKEDQEYQKQGKGKGHNN | -50.25 | 29.0 | CHHHHHHHHHHHHHHHHHHHHHHCCCCCCCC |
| 321 | QREEQIQREQRSNEQEDKRYQKEGRGKGHQP | -50.25 | 29.0 | CHHHHHHHHHHHHHHHHHHHHHHHCCCCCCC |
| 430 | EEEERKQRDQRKQEKEDEEYQRYGRGKGHQP | -50.24 | 38.7 | HHHHHHHHHHHHHHHHHHHHHHHHHCCCCCC |
| 657 | EEEERKQQDQQKNEREDEEYQRRGRGKGHQP | -50.24 | 38.7 | HHHHHHHHHHHHHHHHHHHHHHHHCCCCCCC |

| | | | | |
|---|---|---|---|---|
| 853 | EEEERKQRDQQKNEREDEEYQRRGRGKGHQP | -50.24 | 38.7 | HHHHHHHHHHHHHHHHHHHHHHHHCCCCCCC |
| 552 | EEEQKKQEDQQKNEKEDKRYQREGRGKGHQP | -50.24 | 41.9 | HHHHHHHHHHHHHHHHHHHHHHHHCCCCCCC |
| 152 | QQEKNKQEDQQKNEREDQRYQKEGQGKGHQS | -50.24 | 32.3 | CHHHHHHHHHHHHHHHHHHHHHHCCCCCCCC |
| 594 | EQQQRIQEDQRRNDKEQEEYQRKGRGQGHQP | -50.23 | 29.0 | HHHHHHHHHHHHHHHHHHHHHHHHCCCCCCC |
| 452 | EEEEKKRRDQEKNDKEDEYYQKYGQGKGHNP | -50.22 | 38.7 | CHHHHHHHHHHHHHHHHHHHHHHHHCCCCCC |
| 106 | QREEEIQQQQRKQRNEDERFQREGQGKGKQS | -50.22 | 22.6 | CHHHHHHHHHHHHHHHHHHHHHHHCCCCCCC |
| 207 | EEEQRKQEDQRRNRKEDEEYQRSGRGKGHQP | -50.22 | 38.7 | HHHHHHHHHHHHHHHHHHHHHHHCCCCCCCC |
| 039 | EEEQKKQQDQDNNDREDEEYQRYGRGKGHQP | -50.21 | 41.9 | HHHHHHHHHHCCCCHHHHHHHHHHHCCCCCC |
| 937 | EQEERKNQDQQKNEKEDERYQRQGQGKGHQS | -50.19 | 35.5 | HHHHHHHHHHHHHHHHHHHHHHHHCCCCCCC |
| 309 | EEEEKKQQDKRRNEFEDKYYQQKGQGKGNQN | -50.19 | 45.2 | HHHHHHHHHHHHHHHHHHHHHHHHCCCCCCC |
| 611 | EEQQRKQEDQKRNEEEDERYQRYGRGKGHQP | -50.18 | 41.9 | HHHHHHHHHHHHHHHHHHHHHHHHHCCCCCC |
| 984 | EEQQRKQEDQWRNEWEDEYYQKYGQGKGHQP | -50.18 | 41.9 | HHHHHHHHHHHHHHHHHHHHHHHHHCCCCCC |
| 559 | EEQQRKQEDQWRNEWEDQKYQREGRGKGHQP | -50.18 | 41.9 | HHHHHHHHHHHHHHHHHHHHHHHHCCCCCCC |
| 282 | EEQQRKQQDQYNNEWEDEEYQRKGRGKGHQP | -50.18 | 45.2 | HHHHHHHHHHHHHHHHHHHHHHHHCCCCCCC |
| 590 | EEEQRKQQDQQSNEEEDERYQKQGRGKGHQP | -50.18 | 38.7 | HHHHHHHHHHHHHHHHHHHHHHHCCCCCCCC |
| 877 | EEEERKQQDQRKNEQEDKRYQREGRGKGHQP | -50.18 | 38.7 | HHHHHHHHHHHHHHHHHHHHHHHHCCCCCCC |
| 971 | EEEERKQQDQRKNEYEDEEYQRKGRGKGHQP | -50.18 | 38.7 | HHHHHHHHHHHHHHHHHHHHHHHHCCCCCCC |
| 599 | EEEQRKQEDQRKNEKEDEKYQKEGRGKGHQP | -50.18 | 38.7 | HHHHHHHHHHHHHHHHHHHHHHHHCCCCCCC |
| 381 | EEEQRKQEDQRKNEKEDKRYQREGRGKGHQP | -50.18 | 38.7 | HHHHHHHHHHHHHHHHHHHHHHHHCCCCCCC |
| 122 | EEEQRKQEDQRRNEKEDEEYQRYGRGKGHQP | -50.18 | 38.7 | HHHHHHHHHHHHHHHHHHHHHHHHHCCCCCC |
| 681 | EQEERKQRDQRKNEKEDEEYQRRGRGKGHQP | -50.18 | 35.5 | HHHHHHHHHHHHHHHHHHHHHHHHCCCCCCC |
| 332 | QEEEKKQQDQRKNEKEDEEYQRYGWGKGHYP | -50.18 | 38.7 | CHHHHHHHHHHHHHHHHHHHHHHCCCCCCCC |
| 858 | EQEKRKQEDQRKNEEEDEEFQRRGKGKGKNN | -50.17 | 32.3 | HHHHHHHHHHHHHHHHHHHHHHHCCCCCCCC |
| 953 | EQEKRKQEDQRKNEEEDERFQQQGRGKGKQN | -50.17 | 32.3 | HHHHHHHHHHHHHHHHHHHHHHHHCCCCCCC |
| 468 | QDQERIQQDQWKNEWEHKFYQKEGRGQGHQP | -50.12 | 32.3 | CHHHHHHHHHHHHHHHHHHHHHHHCCCCCCC |
| 647 | EEEERKQQDQKKNDEEDRRYQRYGWGKGHYP | -50.12 | 35.5 | HHHHHHHHHHHHHHHHHHHHHHHCCCCCCCC |
| 132 | EEEERKQQDQYRNDWEDRRYQREGRGKGHQP | -50.12 | 35.5 | HHHHHHHHHHHHHHHHHHHHHHHHCCCCCCC |
| 727 | EEEKRKQEDQRKNDEEDRRYQREGRGKGHQP | -50.12 | 35.5 | HHHHHHHHHHHHHHHHHHHHHHHHCCCCCCC |
| 215 | EEERRKQEDQWRNDWEDRYYQQQGQGKGHQP | -50.12 | 35.5 | HHHHHHHHHHHHHHHHHHHHHHHHCCCCCCC |
| 872 | EEEEKKQRDQDKNRREDERYQREGKGKGHNH | -50.12 | 38.7 | HHHHHHHHHHHHHHHHHHHHHHHCCCCCCCC |
| 506 | EEERRKQEDQYRNRYEDEEYQRKGRGKGHQP | -50.09 | 35.5 | HHHHHHHHHHHHHHHHHHHHHHHHCCCCCCC |
| 621 | EQERRKQEDQKRNREEDERYQRYGRGKGHQP | -50.09 | 32.3 | HHHHHHHHHHHHHHHHHHHHHHHHHCCCCCC |
| 075 | EQERRKQEDQYRNRYEDEEYQRKGRGKGHQP | -50.09 | 32.3 | HHHHHHHHHHHHHHHHHHHHHHHHCCCCCCC |
| 789 | EEQERKRQDQEKNDKEDEEYQRYGQGKGHQS | -50.08 | 38.7 | HHHHHHHHHHHHHCHHHHHHHHHHHCCCCCC |
| 663 | EQEERKQRDQQKNRREDEEYQRRGRGKGHQP | -50.08 | 32.3 | HHHHHHHHHHHHHHHHHHHHHHHHCCCCCCC |
| 410 | EEEERKKQDQYNNDYEDEEFQRKGRGKGKQN | -50.07 | 35.5 | HHHHHHHHHHHCCCHHHHHHHHHHCCCCCCC |
| 085 | DDQRRIQEDQYRNQWEHDRYQREGRGQGHQP | -50.06 | 25.8 | CHHHHHHHHHHHHHHHHHHHHHHCCCCCCCC |
| 939 | EEEERKQQDQRKNDKEDRRYQREGKGKGHNH | -50.06 | 35.5 | HHHHHHHHHHHHHHHHHHHHHHHCCCCCCCC |

| | | | | |
|---|---|---|---|---|
| 587 | EQEERKQQDQRRNDKEDREYQRRGQGKGHNP | -50.06 | 32.3 | HHHHHHHHHHHHHHHHHHHHHHHHCCCCCCC |
| 316 | QEEQKKQEDQRKNDKEDRRYQRYGKGKGHNN | -50.06 | 35.5 | HHHHHHHHHHHHHHHHHHHHHHCCCCCCCCC |
| 510 | QEEEKKQQDQRKNRWEDEYYQKKGQGKGHQP | -50.06 | 35.5 | CHHHHHHHHHHHHHHHHHHHHHHHCCCCCCC |
| 583 | EEEERKQRDQQKNEKEDKRYQREGRGKGHNP | -50.06 | 38.7 | HHHHHHHHHHHHHHHHHHHHHHHCCCCCCCC |
| 626 | DEERKKQEDQYRNEWEDEEYQRKGRGKGHNP | -50.05 | 38.7 | HHHHHHHHHHHHHHHHHHHHHHHCCCCCCCC |
| 633 | EEEKRKQEDQRKNEEEDEEYQRKGRGKGHQP | -50.05 | 38.7 | HHHHHHHHHHHHHHHHHHHHHHHHCCCCCCC |
| 525 | EEEKRKQEDQRKNEEEDEQYQRYGWGKGHYP | -50.05 | 38.7 | HHHHHHHHHHHHHHHHHHHHHHHCCCCCCCC |
| 803 | EEEKRKQEDQRKNEEEDERYQKQGRGKGHQP | -50.05 | 38.7 | HHHHHHHHHHHHHHHHHHHHHHHCCCCCCCC |
| 651 | EEEKRKQEDQRKNEKEDKEYQKQGKGKGHQH | -50.05 | 38.7 | HHHHHHHHHHHHHHHHHHHHHHHHCCCCCCC |
| 916 | EEEKRKQEDQRKNEKEDKEYQKQGWGKGHYP | -50.05 | 38.7 | HHHHHHHHHHHHHHHHHHHHHHHHCCCCCCC |
| 048 | EEERRKQEDQYRNEWEDERYQQQGRGKGHQP | -50.05 | 38.7 | HHHHHHHHHHHHHHHHHHHHHHHHCCCCCCC |
| 523 | EQEKRKQEDQRKNEEEDERYQQQGRGKGHQP | -50.05 | 35.5 | HHHHHHHHHHHHHHHHHHHHHHHHCCCCCCC |
| 838 | QEEENKQRDQYKNEWEDEEYQRKGRGKGHQP | -50.05 | 35.5 | CHHHHHHHHHHHHHHHHHHHHHHHCCCCCCC |
| 933 | QQEKEKQEDQRKNEEEDERYQKQGRGKGHQP | -50.05 | 32.3 | CHHHHHHHHHHHHHHHHHHHHHHCCCCCCCC |
| 744 | EQEQRKNEDQQKNEREDKRYQRYGWGKGHYP | -50.05 | 35.5 | HHHHHHHHHHHHHHHHHHHHHHHCCCCCCCC |
| 804 | EEEERKQRDQWKNDWEDQYYQKKGQGKGHNH | -50.04 | 35.5 | HHHHHHHHHHHHHHHHHHHHHHHHCCCCCCC |
| 262 | EEEERKNQDQYKNRWEDERYQREGRGKGHQP | -50.03 | 35.5 | HHHHHHHHHHHHHHHHHHHHHHHHCCCCCCC |
| 783 | EEEQKKQEDQRKNEYEDEEYQRKGRGKGHNP | -50.03 | 41.9 | HHHHHHHHHHHHHHHHHHHHHHHCCCCCCCC |
| 685 | EEEERKQQDQRKNRKEDEEYQRRGRGKGHQP | -50.02 | 35.5 | HHHHHHHHHHHHHHHHHHHHHHHHCCCCCCC |
| 807 | EEEERKQQDQRKNRWEDEYYQKYGQGKGHQP | -50.02 | 35.5 | HHHHHHHHHHHHHHHHHHHHHHHHHCCCCCC |
| 151 | QEEEKKQQDQWKHEWEDQYYQKYGQGKGHQP | -50.01 | 41.9 | CHHHHHHHHHHHHHHHHHHHHHHHHCCCCCC |
| 153 | EEEERKKEDKYRNEYEDEEYQRKGRGKGTQN | -50.00 | 41.9 | HHHHHHHHHHHHHHHHHHHHHHHHCCCCCCC |
| 205 | EEEQRKNEDQRKNEKEDKEYQKQGQGKGHQS | -50.00 | 38.7 | HHHHHHHHHHHHHHHHHHHHHHHHCCCCCCC |
| 211 | EEQEKKKRDQYKNDWEDERYQKQGRGKGHQP | -50.00 | 41.9 | HHHHHHHHHHHHHHHHHHHHHHHCCCCCCCC |
| 688 | EEEQRKQEDQRRNDKEDRRYQRYGRGKGHQP | -49.99 | 35.5 | HHHHHHHHHHHHHHHHHHHHHHHHCCCCCCC |
| 547 | EEEQRKQQDQRNNRQEDERYQQYGQGKGHQS | -49.99 | 38.7 | HHHHHHHHHHHHHHHHHHHHHHHHHCCCCCC |
| 892 | DEERKKQEDQRRNEQEDKRYQREGKGKGHSH | -49.99 | 38.7 | CHHHHHHHHHHHHHHHHHHHHHHCCCCCCCC |
| 421 | EEEERKQQDQRKNEKEDEEYQRYGQGKGHQS | -49.99 | 38.7 | HHHHHHHHHHHHHHHHHHHHHHHHHCCCCCC |
| 049 | EEEERKQQDQRKNEKEDERYQQQGRGKGHQP | -49.99 | 38.7 | HHHHHHHHHHHHHHHHHHHHHHHHCCCCCCC |
| 726 | EEEERKQQDQRKNEKEDKRYQREGKGKGHNH | -49.99 | 38.7 | HHHHHHHHHHHHHHHHHHHHHHHCCCCCCCC |
| 330 | EEEERKQRDQRKNEEEDKKYQRYGWGKGHYP | -49.99 | 38.7 | HHHHHHHHHHHHHHHHHHHHHHHCCCCCCCC |
| 879 | EQEERKQQDQRKNEHEDQRYQREGRGKGHQP | -49.99 | 35.5 | HHHHHHHHHHHHHHHHHHHHHHHHCCCCCCC |
| 919 | EQEQRKQEDQRKNEKEDERYQKQGWGKGHYP | -49.99 | 35.5 | HHHHHHHHHHHHHHHHHHHHHHHCCCCCCCC |
| 487 | QEEQKKQEDQRKNEYEDEEYQRKGRGKGHQP | -49.99 | 38.7 | HHHHHHHHHHHHHHHHHHHHHHHHCCCCCCC |
| 983 | EEEQRKQQDQQNNEREDERYQRQGRGKGHQP | -49.98 | 41.9 | HHHHHHHHHHHHHHHHHHHHHHHCCCCCCCC |
| 592 | EEEQRKQEDQRKNRKEDERYQREGRGKGHQP | -49.98 | 35.5 | HHHHHHHHHHHHHHHHHHHHHHHHCCCCCCC |
| 847 | EEQEKINRDQKKNDEEHRRYQKEGQGQGHQS | -49.96 | 35.5 | CHHHHHHHHHHCHHHHHHHHHHHHCCCCCCC |
| 120 | EEEQRKQEDQRRNRKEDERYQREGRGKGHQP | -49.95 | 35.5 | HHHHHHHHHHHHHHHHHHHHHHHHCCCCCCC |

| | | | | |
|---|---|---|---|---|
| 190 | EQEQRKQQDQRSNRQEDERYQRYGRGKGHQP | -49.95 | 32.3 | HHHHHHHHHHHHHHHHHHHHHHHHHCCCCCC |
| 340 | EEEERKKQDQEKNDKEDEEYQRKGRGKGHNP | -49.95 | 35.5 | HHHHHHHHHHHHHHHHHHHHHHHCCCCCCCC |
| 635 | EEEERKKQDQEKNDKEDKRYQREGKGKGHNN | -49.95 | 35.5 | HHHHHHHHHHHHHHHHHHHHHHHCCCCCCCC |
| 414 | EEEERKQRDQWKNEWEHEYKQKYGQGCGKQN | -49.93 | 29.0 | HHHHHHHHHHHHHHHHHHHHHHHHHCCCCCC |
| 343 | EEEQKKQEDQRRNEKEDKRYQREGRGKGHQP | -49.92 | 41.9 | HHHHHHHHHHHHHHHHHHHHHHHHCCCCCCC |
| 000 | EEEQKKQRDQRKNEKEDKEYQKQGRGKGHQP | -49.92 | 41.9 | HHHHHHHHHHHHHHHHHHHHHHHHCCCCCCC |
| 024 | EEEQKKQRDQRKNEQEDERYQRQGRGKGHQP | -49.92 | 41.9 | HHHHHHHHHHHHHHHHHHHHHHHHCCCCCCC |
| 365 | EEEQRKQEDQRRNEKEDEYYQKYGQGKGHQP | -49.92 | 38.7 | HHHHHHHHHHHHHHHHHHHHHHHHHCCCCCC |
| 443 | QDQQRKQEDQRRNEKEDEEYQRQGNGKGHQP | -49.92 | 35.5 | CHHHHHHHHHHHHHHHHHHHHHHCCCCCCCC |
| 166 | QDQQRKQEDQRRNEKEDKRYQREGRGKGHQP | -49.92 | 35.5 | CHHHHHHHHHHHHHHHHHHHHHHHCCCCCCC |
| 913 | QRQEEIQRQQRSNDKEDREYQRRGQGKGHQP | -49.92 | 29.0 | CHHHHHHHHHHHCCHHHHHHHHHHCCCCCCC |
| 963 | REEERIQQDQRSQDKQDRRYQRYGRGKGHQP | -49.91 | 29.0 | CHHHHHHHHHHHHHHHHHHHHHHHCCCCCCC |
| 936 | QKQQEIQEQQKRNEEEDKRYQRYGRGKGHQP | -49.90 | 32.3 | CHHHHHHHHHHHHHHHHHHHHHHHCCCCCCC |
| 042 | EEEERKQRDQYKNRWEDERYQREGDGKGHQP | -49.89 | 35.5 | HHHHHHHHHHHHHHHHHHHHHHHCCCCCCCC |
| 644 | EEEEKKKRDKYKNEYEDKEYQRKGRGKGHQP | -49.88 | 45.2 | CHHHHHHHHHHHHHHHHHHHHHHCCCCCCCC |
| 491 | EEEERKKEDKYRNEWEDKRYQREGRGKGHQP | -49.88 | 41.9 | HHHHHHHHHHHHHHHHHHHHHHHHCCCCCCC |
| 717 | EEEERKKQDKYNNEWEDEEYQRKGRGKGHQP | -49.88 | 45.2 | HHHHHHHHHHHHHHHHHHHHHHHHCCCCCCC |
| 627 | EQEERKKQDKQSNEKEDEEYQRRGRGKGHQP | -49.88 | 38.7 | HHHHHHHHHHHHHHHHHHHHHHHHCCCCCCC |
| 811 | DEERKKQEDQWRNEWEDEYYQKYGQGKGHNP | -49.86 | 38.7 | HHHHHHHHHHHHHHHHHHHHHHHHHCCCCCC |
| 385 | EEEKRKQEDQRKNEEEDKRYQRYGRGKGHQP | -49.86 | 38.7 | HHHHHHHHHHHHHHHHHHHHHHHHCCCCCCC |
| 057 | EEEQRKQEDQRKNEEEDKYYQQKGQGKGHQP | -49.86 | 38.7 | HHHHHHHHHHHHHHHHHHHHHHHHCCCCCCC |
| 735 | QEEEKKQRDQYKNEWEDEEYQRKGRGKGHQP | -49.86 | 38.7 | CHHHHHHHHHHHHHHHHHHHHHHHCCCCCCC |
| 476 | QEEKKKQEDQRKNEKEDEEYQRRGKGKGHQH | -49.86 | 38.7 | HHHHHHHHHHHHHHHHHHHHHHHHCCCCCCC |
| 420 | QEEKKKQEDQRKNEKEDEEYQRRGQGKGHQS | -49.86 | 38.7 | HHHHHHHHHHHHHHHHHHHHHHHHHCCCCCC |
| 322 | EEEQRKNEDQKRNREEHERKQKQGKGCGKNN | -49.82 | 25.8 | HHHHHHHHHHHHHHHHHHHHHHHCCCCCCCC |
| 131 | EEEERKKQDQRKNDEEDQRYQKQGKGKGHNN | -49.81 | 35.5 | HHHHHHHHHHHHHHHHHHHHHHHCCCCCCCC |
| 673 | EQEERIQRDQQKNEREQKRYQKEGRGQGHQP | -49.80 | 29.0 | HHHHHHHHHHHHHHHHHHHHHHHHCCCCCCC |
| 080 | QREKEIQEQQRKNDEEDRRYQQQGKGKGHSN | -49.78 | 25.8 | CHHHHHHHHHHHHHHHHHHHHHHCCCCCCCC |
| 016 | EEEEKKKRDQYKNDYEDEEYQRQGRGKGHQP | -49.76 | 38.7 | CHHHHHHHHHHHHHHHHHHHHHHHCCCCCCC |
| 857 | EEEEKKKRDQYNNDAEDKRYQREGRGKGHQP | -49.76 | 45.2 | CHHHHHHHHHHCHHHHHHHHHHHHCCCCCCC |
| 921 | EEEERKKQDQKNNDEEDERYQQQGWGKGHYP | -49.76 | 38.7 | HHHHHHHHHHHHHHHHHHHHHHHCCCCCCCC |
| 946 | EEEERKKQDQQNNDKEDEEYQRRGRGKGHQP | -49.76 | 38.7 | HHHHHHHHHHHCCCHHHHHHHHHHCCCCCCC |
| 124 | EEEERKKQDQYNNDWEDEEYQRKGRGKGHQP | -49.76 | 38.7 | HHHHHHHHHHHCCCHHHHHHHHHHCCCCCCC |
| 593 | EQEERKKEDQYRNDYEDEEYQRKGRGKGHQP | -49.76 | 32.3 | HHHHHHHHHHHHHHHHHHHHHHHHCCCCCCC |
| 773 | EEEERKQQDQRKNEKEDQRYQREGKGKGHNH | -49.75 | 38.7 | HHHHHHHHHHHHHHHHHHHHHHHHCCCCCCC |
| 130 | EEEERKKQDQEKNDKEDERYQKQGKGKGHNH | -49.75 | 35.5 | HHHHHHHHHHHHHHHHHHHHHHHCCCCCCCC |
| 357 | EEEERKKQDQEKNDKEDKRYQKEGKGKGHNH | -49.75 | 35.5 | HHHHHHHHHHHHHHHHHHHHHHHCCCCCCCC |
| 897 | EEEQRKQEDQKRNREEDERYQKQGRGKGHQP | -49.73 | 35.5 | HHHHHHHHHHHHHHHHHHHHHHHCCCCCCCC |

| | | | | |
|---|---|---|---|---|
| 926 | EEQERKNQDQWKHEWEDEYYQKYGQGKGHQP | -49.71 | 45.2 | HHHHHHHHHHHHHHHHHHHHHHHHHCCCCCC |
| 125 | EEQQRKQEDQYRNRYEDEEYQRKGRGKGHQP | -49.70 | 38.7 | HHHHHHHHHHHHHHHHHHHHHHHHCCCCCCC |
| 912 | QDQERKKQDKYKNEWEHEEYQRKGRGRGHNP | -49.69 | 32.3 | CHHHHHHHHHHHHHHHHHHHHHHCCCCCCCC |
| 029 | EQEKRIQEDQRKNDEEQRRYQNEGKGQGHNH | -49.68 | 25.8 | HHHHHHHHHHHHHHHHHHHHHHHCCCCCCCC |
| 059 | EEEEKKKRDKYKNEWEDKRYQREGRGKGHQP | -49.68 | 45.2 | CHHHHHHHHHHHHHHHHHHHHHHHCCCCCCC |
| 179 | QEEQKKQEDQRKNEKEDKRYQRYGRGKGHQP | -49.66 | 38.7 | HHHHHHHHHHHHHHHHHHHHHHHHCCCCCCC |
| 113 | SDQERKQRDQKKNEEEDQRYQRYGWGKGHYP | -49.66 | 35.5 | CHHHHHHHHHHHHHHHHHHHHHHCCCCCCCC |
| 425 | EEEERKKQDKYNNEWEHKRKQREGQGCGKQS | -49.66 | 35.5 | HHHHHHHHHHHHHHHHHHHHHHHCCCCCCCC |
| 729 | EREKRYQEDQRKNDEEDKRYQRYGWGKGHYP | -49.66 | 32.3 | CHHHHHHHHHHHHHHHHHHHHHHCCCCCCCC |
| 878 | EEEEKKKRDKKKNEEEDKRYQKEGRGKGHQP | -49.64 | 45.2 | CHHHHHHHHHHHHHHHHHHHHHHHCCCCCCC |
| 455 | EQERRKQEDQQRNEREDKRYQREGQGKGHQS | -49.63 | 35.5 | HHHHHHHHHHHHHHHHHHHHHHHCCCCCCCC |
| 240 | QQQQNIQEQQRRHEKEDERYQRQGWGKGHYP | -49.63 | 35.5 | HHHHHHHHHHHHHHHHHHHHHHHHCCCCCCC |
| 520 | EEEERKQRDQWKNEWEDQYYQKKGQGKGHNH | -49.62 | 38.7 | HHHHHHHHHHHHHHHHHHHHHHHHCCCCCCC |
| 247 | EEEEKKQRDKYKNEWEDRRYQREGKGKGHNN | -49.61 | 45.2 | CHHHHHHHHHHHHHHHHHHHHHCCCCCCCCC |
| 503 | EEQQKKQRDQKKNEEEDKRYQRYGRGKGHQP | -49.59 | 45.2 | HHHHHHHHHHHHHHHHHHHHHHHHCCCCCCC |
| 139 | EEEQRKQKDQESNEREDEYYQKYGQGKGHQP | -49.56 | 38.7 | HHHHHHHHHHHHHHHHHHHHHHHHHCCCCCC |
| 987 | QREEEIQRDQRKNEKEQEEYQRRGRGQGHQP | -49.56 | 25.8 | CHHHHHHHHHHHHHHHHHHHHHHHHCCCCCC |
| 172 | EQQQRKQEEKRRNEKEDKQYQERGRGKGHQP | -49.55 | 38.7 | HHHHHHHHHHHHHHHHHHHHHHHHCCCCCCC |
| 051 | EQQQRKQEEKRRNEKEDKRYQREGRGKGHQP | -49.55 | 38.7 | HHHHHHHHHHHHHHHHHHHHHHHHCCCCCCC |
| 535 | EEQQRKQQDKRNNENEHKRYQRYGRGQGHQP | -49.54 | 41.9 | HHHHHHHHHHHHHHHHHHHHHHHHCCCCCCC |
| 219 | EEEKRKQEDQRKNEEEDKRYQKEGRGKGHQP | -49.53 | 38.7 | HHHHHHHHHHHHHHHHHHHHHHHHCCCCCCC |
| 069 | EEEEKKQQDQRKNEKEDEEYQRRGQGKGHNP | -49.53 | 41.9 | CHHHHHHHHHHHHHHHHHHHHHHHCCCCCCC |
| 683 | EEERRKQEDQYRNDSEDRRYQRYGRGKGHQP | -49.52 | 35.5 | HHHHHHHHHHHHHHHHHHHHHHHHCCCCCCC |
| 198 | EEEERKQRDQRRHDQEHKRYQREGRGQGHQP | -49.52 | 32.3 | HHHHHHHHHHHHHHHHHHHHHHHHCCCCCCC |
| 001 | QEEQKKQEDQRRHDKEHNRYQREGRGQGHQP | -49.52 | 32.3 | HHHHHHHHHHHHHHHHHHHHHHHCCCCCCCC |
| 400 | EEEKRKQEDQRKNREEDEYYQKYGQGKGHQP | -49.52 | 35.5 | HHHHHHHHHHHHHHHHHHHHHHHHHCCCCCC |
| 745 | EEEERKKQDQQKNEREDKRYQKEGRGKGHQP | -49.49 | 38.7 | HHHHHHHHHHHHHHHHHHHHHHHCCCCCCCC |
| 668 | ENEQRKQQEKRSNEEEDKEYQRKGRGKGTQN | -49.48 | 35.5 | CHHHHHHHHHHHHHHHHHHHHHHHCCCCCCC |
| 825 | EREERYQQDKRKNEWEDKYYQQQGQGKGHQS | -49.47 | 38.7 | CHHHHHHHHHHHHHHHHHHHHHHHCCCCCCC |
| 538 | EEEERKQQDQRKNEKEDKEYQKRGDGKGHQP | -49.47 | 38.7 | HHHHHHHHHHHHHHHHHHHHHHCCCCCCCCC |
| 348 | EEEERKQRDQYKNEYEDEEYQRKGRGKGHQP | -49.46 | 38.7 | HHHHHHHHHHHHHHHHHHHHHHHHCCCCCCC |
| 149 | EEEQRKQQDQESNKREDEEYQRRGRGKGHQP | -49.45 | 35.5 | HHHHHHHHHHHHHHHHHHHHHHHHCCCCCCC |
| 787 | EQQQRKQEDKRKNEEEDKYYQQKGQGKGNQN | -49.43 | 41.9 | HHHHHHHHHHHHHHHHHHHHHHHCCCCCCCC |
| 839 | QQEKNIQEDQRKNEEEQERYQKQGRGQGHQP | -49.43 | 25.8 | CHHHHHHHHHHHHHHHHHHHHHHHCCCCCCC |
| 115 | EEQQKKQRDQRKNDKEHQRYQRYGKGQGHNN | -49.43 | 35.5 | HHHHHHHHHHHHHHHHHHHHHHHCCCCCCCC |
| 624 | DEERKKQEDQRRNEKEDEEYQRKGRGKGHQP | -49.42 | 38.7 | CHHHHHHHHHHHHHHHHHHHHHHHCCCCCCC |
| 614 | EEEERKQQDKRKNEKEHKRYQREGRGQGHQP | -49.41 | 35.5 | HHHHHHHHHHHHHHHHHHHHHHHHCCCCCCC |
| 305 | QEEQKKQEDKRKNEKEHKRYQREGKGQGHQP | -49.41 | 35.5 | HHHHHHHHHHHHHHHHHHHHHHHCCCCCCCC |

| | | | | |
|---|---|---|---|---|
| 232 | EQEQRKQEDQRRNEKEDKRYQREGRGKGHQP | -49.40 | 35.5 | HHHHHHHHHHHHHHHHHHHHHHHHCCCCCCC |
| 031 | EQQQRKNEDQRKNDEEDRRYQNQGRGKGHQP | -49.40 | 35.5 | HHHHHHHHHHHHHHHHHHHHHHCCCCCCCCC |
| 454 | EQEERIQQDDRKNRKEDEEYQRRGRGKGHQP | -49.39 | 32.3 | CHHHHHHHHHHHHHHHHHHHHHHHCCCCCCC |
| 470 | EEEQRKQEDQKRNEEEDERYQRYGWGKGHYP | -49.38 | 38.7 | HHHHHHHHHHHHHHHHHHHHHHHCCCCCCCC |
| 364 | EQEQRKQQDQKNNEEEDERYQQQGRGKGHQP | -49.38 | 38.7 | HHHHHHHHHHHHHHHHHHHHHHHHCCCCCCC |
| 073 | EQEQRKQQDQYNNEWEDEEYQRKGRGKGHNP | -49.38 | 38.7 | HHHHHHHHHHHHHHHHHHHHHHHCCCCCCCC |
| 249 | QQEENIQRDQRKNEKEQEEYQRRGRGQGHQP | -49.37 | 25.8 | CHHHHHHHHHHHHHHHHHHHHHHHHCCCCCC |
| 981 | EEEERKQRDQQKNDREHKRYQREGRGQGHQP | -49.36 | 29.0 | HHHHHHHHHHHHHHHHHHHHHHHHCCCCCCC |
| 743 | EEEKRKQEDQRKNEEEDERYQRQGKGKGTNR | -49.35 | 41.9 | HHHHHHHHHHHHHHHHHHHHHHHCCCCCCCC |
| 935 | EEEEKKQRDQWKNEWEDEYYQRKGQGKGHQP | -49.34 | 41.9 | HHHHHHHHHHHHHHHHHHHHHHHHCCCCCCC |
| 209 | EEEERKQQDKRKNEKEDRRFQRYGRGKGKQN | -49.33 | 38.7 | HHHHHHHHHHHHHHHHHHHHHHHCCCCCCCC |
| 366 | EQQQRKQEDQWRNDYEHEEYQRKGRGQGHQP | -49.30 | 29.0 | HHHHHHHHHHHHHHHHHHHHHHHHCCCCCCC |
| 951 | EEQQRKQEDQRKNDEEHKRYQRYGRGQGHQP | -49.30 | 32.3 | HHHHHHHHHHHHHHHHHHHHHHHHCCCCCCC |
| 437 | EEEERKKQDQYKNEYEDEEYQRKGRGKGHQP | -49.30 | 38.7 | HHHHHHHHHHHHHHHHHHHHHHHHCCCCCCC |
| 985 | EQEERKKRDQWKNEWEDKYYQKQGQGKGHQP | -49.30 | 35.5 | HHHHHHHHHHHHHHHHHHHHHHHHCCCCCCC |
| 790 | EEEEKKQQDQRKNDNEHERYQQQGRGQGHQP | -49.29 | 32.3 | CHHHHHHHHHHHHHHHHHHHHHHHHCCCCCC |
| 096 | EEEERKQQDQRKNDYEHEEYQRQGRGQGHQP | -49.29 | 29.0 | HHHHHHHHHHHHHHHHHHHHHHHHCCCCCCC |
| 870 | EEEERKQRDQRKNDQEHKRYQRYGWGQGHYP | -49.29 | 29.0 | HHHHHHHHHHHHHHHHHHHHHHHCCCCCCCC |
| 224 | EEEQKKQEDQRKNDKEHKRYQREGKGQGHNH | -49.29 | 32.3 | HHHHHHHHHHHHHHHHHHHHHHHHCCCCCCC |
| 389 | EEEQRKQEDQRKNDKEHKRYQREGRGQGHQP | -49.29 | 29.0 | HHHHHHHHHHHHHHHHHHHHHHHHCCCCCCC |
| 814 | EEQQRKQEDQKRNEEEDKRYQREGDGKGNQN | -49.25 | 41.9 | HHHHHHHHHHHHHHHHHHHHHHHCCCCCCCC |
| 776 | EEEEKKKQDQKNNEEEDERFQKQGRGKGKQN | -49.23 | 41.9 | HHHHHHHHHHHHHHHHHHHHHHHCCCCCCCC |
| 074 | EEERRKQEDAYRNRYEDEEYQRKGRGKGTQN | -49.22 | 35.5 | HHHHHHHHHHHHHHHHHHHHHHHHCCCCCCC |
| 993 | EQEKRIQEDQRKNDEEQRRYQRYGRGQGHQP | -49.21 | 25.8 | HHHHHHHHHHHHHHHHHHHHHHHHHCCCCCC |
| 101 | KQQQEKQEDQRRNDKEHEEYQRRGKGQGHNH | -49.20 | 25.8 | CHHHHHHHHHHHHHHHHHHHHHHHCCCCCCC |
| 682 | EEEERKQRQKYSEEWQHKRYQREGRGQGHQP | -49.19 | 29.0 | CHHHHHHHHHHHHHHHHHHHHHHHCCCCCCC |
| 884 | DEERKKQEDQYRNDWEHEEYQRKGRGQGHQP | -49.17 | 29.0 | HHHHHHHHHHHHHHHHHHHHHHHHCCCCCCC |
| 750 | EEEKRKQEDQRKNDEEHKRYQREGWGQGHYP | -49.17 | 29.0 | HHHHHHHHHHHHHHHHHHHHHHHHCCCCCCC |
| 351 | EQEERKQQDQKKNDEEHNRYQRYGQGQGHQS | -49.17 | 25.8 | HHHHHHHHHHHHHHHHHHHHHHHHCCCCCCC |
| 426 | EEEQRKQQDKRNNEKEHKEYQKQGRGQGHQP | -49.15 | 38.7 | HHHHHHHHHHHHHHHHHHHHHHHCCCCCCCC |
| 374 | EEEERKKQDQEKNRKEDERYQREGRGKGHQP | -49.15 | 35.5 | HHHHHHHHHHHHHHHHHHHHHHHHCCCCCCC |
| 192 | EEEERKQRDQQKNDREHERYQQQGRGQGHQP | -49.13 | 29.0 | HHHHHHHHHHHHHHHHHHHHHHHHCCCCCCC |
| 558 | EQEERKQQDQRKNDKEDRRYQREGKGKGNNN | -49.13 | 32.3 | HHHHHHHHHHHHHHHHHHHHHHHCCCCCCCC |
| 832 | EQEQRKQEEQRSNDEEDEYYQKYGQGKGHQP | -49.12 | 29.0 | HHHHHHHHHHHHHHHHHHHHHHHHHCCCCCC |
| 541 | DEERKKNEDKYRNEYEHEEYQRKGRGQGHQP | -49.11 | 35.5 | HHHHHHHHHHHHHHHHHHHHHHHHCCCCCCC |
| 998 | EEEEKKKQDQEKNEKEDKRYQREGRGKGHQP | -49.11 | 41.9 | CHHHHHHHHHHHHHHHHHHHHHHHCCCCCCC |
| 304 | EEEERKKKDQEKNEREDEEYQRRGRGKGHQP | -49.11 | 38.7 | CHHHHHHHHHHHHHHHHHHHHHHHCCCCCCC |
| 924 | EEEQKKQEDQRKNDKEHEEYQRYGRGQGHQP | -49.10 | 32.3 | HHHHHHHHHHHHHHHHHHHHHHHHHCCCCCC |

| | | | | |
|---|---|---|---|---|
| 253 | EQEERKQQDQRKNEWEDEEYQRKGRGKGNQN | -49.05 | 35.5 | HHHHHHHHHHHHHHHHHHHHHHHHCCCCCCC |
| 786 | EEEERKKRDQWKNRWEDEYYQKEGQGKGHQP | -49.05 | 35.5 | HHHHHHHHHHHHHHHHHHHHHHHHCCCCCCC |
| 267 | EQEQRKQEDQRRNDKEHERYQQQGWGQGHYP | -49.03 | 25.8 | HHHHHHHHHHHHHHHHHHHHHHHCCCCCCCC |
| 283 | EQEERIQRDQWKNEWEQQYYQKKGQGQGHQS | -49.00 | 29.0 | HHHHHHHHHHHHHHHHHHHHHHHHCCCCCCC |
| 948 | EEEERKWRDQQSNDKEHEEYQRRGRGRGHQP | -49.00 | 29.0 | CHHHHHHHHHHHCHHHHHHHHHHHCCCCCCC |
| 474 | EEEEKKKRDQYKNDWEDRRYQREGKGKGHNN | -49.00 | 38.7 | CHHHHHHHHHHHHHHHHHHHHHCCCCCCCCC |
| 368 | EQEERKKQDQQNNDKEDREYQKRGWGKGHYP | -49.00 | 35.5 | HHHHHHHHHHHHCHHHHHHHHHHCCCCCCC |
| 431 | QEEENKKEDQRKNDKEDREYQKQGRGKGHQP | -49.00 | 32.3 | CHHHHHHHHHHHHHHHHHHHHHHHCCCCCCC |
| 379 | EQEERKNQDKKKNEEEHKRYQKEGQGQGHQS | -49.00 | 32.3 | HHHHHHHHHHHHHHHHHHHHHHHHCCCCCCC |
| 164 | EQEERKKEDQWRNRWEDEYYQKYGQGKGHQP | -49.00 | 32.3 | HHHHHHHHHHHHHHHHHHHHHHHHHCCCCCC |
| 338 | QQEENKKQDQYNNRYEDEEYQRKGRGKGHQP | -49.00 | 32.3 | CHHHHHHHHHHHHHHHHHHHHHHHCCCCCCC |
| 004 | QEQENKKQDQYNNDWEDRRYQREGRGKGHQP | -48.99 | 38.7 | CHHHHHHHHHHCCHHHHHHHHHHCCCCCCCC |
| 689 | EEEERKQQDKRKNENEHKQYQQRGQGQGHQS | -48.99 | 35.5 | HHHHHHHHHHHHHHHHHHHHHHHHHCCCCCC |
| 860 | EREERYQQDQRKNRKEDERYQREGRGKGHQP | -48.99 | 32.3 | CHHHHHHHHHHHHHHHHHHHHHHHCCCCCCC |
| 582 | EEEERKKQDQKKNEEEDERYQQQGRGKGHQP | -48.98 | 38.7 | HHHHHHHHHHHHHHHHHHHHHHHHCCCCCCC |
| 119 | EEEERKKQDQYKNEAEDERYQQQGQGKGHQS | -48.98 | 41.9 | HHHHHHHHHHHHHHHHHHHHHHHHCCCCCCC |
| 652 | EEEERKKQDQYRNEWEDEEYQRKGRGKGHQP | -48.98 | 38.7 | HHHHHHHHHHHHHHHHHHHHHHHHCCCCCCC |
| 788 | EEEKRKQEDQRKNDKEHKEYQKQGKGQGHNN | -48.97 | 29.0 | HHHHHHHHHHHHHHHHHHHHHHHCCCCCCCC |
| 734 | EQEERKQRDQYKNDWEHEEYQRKGRGQGHQP | -48.97 | 25.8 | HHHHHHHHHHHHHHHHHHHHHHHHCCCCCCC |
| 440 | EQEERIQQDQRKNDKEQRRYQREGKGQGHNH | -48.97 | 25.8 | CHHHHHHHHHHHHHHHHHHHHHHHCCCCCCC |
| 505 | ERQQRYQQDQYNNEWEDKRYQREGRGKGHQP | -48.96 | 41.9 | CHHHHHHHHHHCCHHHHHHHHHHHCCCCCCC |
| 864 | ERQQRYQQDQYNNEYEDEEYQRKGRGKGHNP | -48.96 | 41.9 | CHHHHHHHHHHCCCHHHHHHHHHCCCCCCCC |
| 970 | EEQERKKEDQYRNRFEDEYYQQKGQGKGHQP | -48.95 | 38.7 | HHHHHHHHHHHHHHHHHHHHHHHHCCCCCCC |
| 571 | EEEERKKEDKKRNEEEDKRYQRYGRGKGNQN | -48.94 | 41.9 | HHHHHHHHHHHHHHHHHHHHHHHCCCCCCCC |
| 359 | EEEERKQRDQYKNEWEDEEYQRKGRGKGHQP | -48.93 | 38.7 | HHHHHHHHHHHHHHHHHHHHHHHHCCCCCCC |
| 645 | EEEEKKKRDQYNNEWEDEYYQRKGQGKGHNH | -48.93 | 45.2 | CHHHHHHHHHHHHHHHHHHHHHHHCCCCCCC |
| 465 | EEEERKKEDQWKNEWEDSYYQQYGQGKGHQP | -48.93 | 38.7 | HHHHHHHHHHHHHHHHHHHHHHHHHCCCCCC |
| 975 | EEEERKKEDQYRNEYEDKSYQRYGRGKGHQP | -48.93 | 38.7 | HHHHHHHHHHHHHHHHHHHHHHHCCCCCCCC |
| 680 | EEEERKKQDQYNNEWEDEEYQRKGRGKGHQP | -48.93 | 41.9 | HHHHHHHHHHHHHHHHHHHHHHHHCCCCCCC |
| 324 | EEEERKKQDQYNNEYEDEEYQRKGRGKGHQP | -48.93 | 41.9 | HHHHHHHHHHHHHHHHHHHHHHHHCCCCCCC |
| 071 | EQEERKKQDQYNNEYEDEEYQRKGRGKGHQP | -48.93 | 38.7 | HHHHHHHHHHHHHHHHHHHHHHHHCCCCCCC |
| 451 | QEEEKKKEDQYRNEWEDERYQQEGRGKGHQP | -48.93 | 38.7 | CHHHHHHHHHHHHHHHHHHHHHHHCCCCCCC |
| 169 | EEEEKKRQDMKNNREEDERYQRYGWGKGHYP | -48.91 | 41.9 | CHHHHHHHHHHHHHHHHHHHHHHCCCCCCCC |
| 515 | EEEERKQRQQRKQDQEDRRYQREGKGKGHNH | -48.91 | 32.3 | HHHHHHHHHHHHHHHHHHHHHHHCCCCCCCC |
| 724 | QDQERKQQDQRKNDYEHEEYQRKGRGQGHQP | -48.90 | 25.8 | CHHHHHHHHHHHHHHHHHHHHHHHCCCCCCC |
| 911 | EEERRKQEDAYRNRYEDEEYQRKGRGKGHQP | -48.90 | 35.5 | HHHHHHHHHHHHHHHHHHHHHHHHCCCCCCC |
| 845 | QEEKKKQEDARKNRQEDQRYQREGRGKGHQP | -48.90 | 35.5 | HHHHHHHHHHHHHHHHHHHHHHHHCCCCCCC |
| 170 | EEEQRKNEQARQERREDEEYQRKGRGKGHQP | -48.87 | 32.3 | HHHHHHHHHHHHHHHHHHHHHHHHCCCCCCC |

| | | | | |
|---|---|---|---|---|
| 696 | EEQEKKNQDQQKNDKEHKRYQRYGRGQGHQP | -48.87 | 35.5 | HHHHHHHHHHHHHHHHHHHHHHHHCCCCCCC |
| 661 | EEEERKQQDARKNRKEDEEYQRRGRGKGHQP | -48.83 | 35.5 | HHHHHHHHHHHHHHHHHHHHHHHHCCCCCCC |
| 902 | EEEERKKQDQWKNEWEHKYKQKKGQGCGKQN | -48.83 | 29.0 | HHHHHHHHHHHHHHHHHHHHHHCCCCCCCCC |
| 337 | EREERYQRDQYKNEYEDEEYQRKGRGKGHQP | -48.82 | 35.5 | CHHHHHHHHHHHHHHHHHHHHHHHCCCCCCC |
| 705 | EQQERKNREKYKNESEDKRYQKEGKGKGHNN | -48.80 | 38.7 | HHHHHHHHHHHHHHHHHHHHHHCCCCCCCCC |
| 424 | QDEERKQRDQQKNDKEHKRYQREGQGQGHQS | -48.77 | 22.6 | CHHHHHHHHHHHHHHHHHHHHHHHCCCCCCC |
| 945 | DEERKRQEDQRRNDKEHEEYQRYGQGQGHQS | -48.77 | 29.0 | HHHHHHHHHHHHHHHHHHHHHHHHHCCCCCC |
| 766 | EEEERKQQDQRRHRNEHERYQREGRGQGHQP | -48.75 | 32.3 | HHHHHHHHHHHHHHHHHHHHHHHHCCCCCCC |
| 354 | EEQQRKQEQQRRNDKEDEEYQRYGWGKGHYP | -48.74 | 35.5 | HHHHHHHHHHHHHHHHHHHHHHHCCCCCCCC |
| 829 | EEEEKKKRDQESNEKEDEEYQRRGQGKGHQP | -48.72 | 41.9 | CHHHHHHHHHHHHHHHHHHHHHHHHCCCCCC |
| 826 | EQEERIKQDQQKNEKEQEEYQRRGRGQGHQP | -48.72 | 29.0 | CHHHHHHHHHHHHHHHHHHHHHHHHCCCCCC |
| 595 | EEEQRKKEDQRKNDEEDEEYQRKGRGKGHNP | -48.71 | 35.5 | HHHHHHHHHHHHHHHHHHHHHHHHCCCCCCC |
| 753 | EEEQRKKEDQRKNDEEDERYQQEGRGKGHQP | -48.71 | 35.5 | HHHHHHHHHHHHHHHHHHHHHHHHCCCCCCC |
| 110 | QDEQRKQEDQRKNDKEHEEYQRRGRGQGHQP | -48.71 | 22.6 | CHHHHHHHHHHHHHHHHHHHHHHHHCCCCCC |
| 954 | EEEERKSRDQYKNDYEDREYQRKGRGKGHQP | -48.70 | 35.5 | CHHHHHHHHHHHHHHHHHHHHHHHCCCCCCC |
| 191 | EEERRKQEDKYRNEWEHEEYQRKGRGQGHQP | -48.67 | 35.5 | HHHHHHHHHHHHHHHHHHHHHHHHCCCCCCC |
| 046 | QEQQNKQEDQRRNDKEHRRYQREGKGQGHNH | -48.66 | 29.0 | HHHHHHHHHHHHHHHHHHHHHHHHCCCCCCC |
| 866 | DNERRIQEEKYRNEYEQEEYQRKGRGQGHQP | -48.66 | 25.8 | CHHHHHHHHHHHHHHHHHHHHHHHCCCCCCC |
| 642 | EEEERKKQDKWKNEWEDEYYQRKGQGKGHNP | -48.65 | 41.9 | HHHHHHHHHHHHHHHHHHHHHHHHCCCCCCC |
| 556 | EEEKKKQEDQRKNEEEHKYFQKEGQGQGKQN | -48.65 | 32.3 | HHHHHHHHHHHHHHHHHHHHHHHHCCCCCCC |
| 706 | QREKQIQEEKRSNEEEQKRYQRYGWGQGHYP | -48.65 | 25.8 | HHHHHHHHHHHHHHHHHHHHHHHCCCCCCCC |
| 999 | EEERRKQEDKERNEKEHKRYQREGRGRGHQP | -48.65 | 35.5 | HHHHHHHHHHHHHHHHHHHHHHHHCCCCCCC |
| 598 | DNQRRKNHEQESNDKEDKRYQREGRGKGHQP | -48.63 | 29.0 | CHHHHHHHHHHHCHHHHHHHHHHHCCCCCCC |
| 679 | EEEERKRQDKEKNEKEDKRYQREGRGKGHQP | -48.63 | 41.9 | HHHHHHHHHHHHHHHHHHHHHHHHCCCCCCC |
| 243 | EEEERKKEDQRKNDEEDRRYQRYGRGKGHQP | -48.61 | 35.5 | HHHHHHHHHHHHHHHHHHHHHHHHCCCCCCC |
| 763 | EEEERKQRDKEKNEKEHKRYQREGRGRGHQP | -48.57 | 35.5 | HHHHHHHHHHHHHHHHHHHHHHHHCCCCCCC |
| 808 | QDQERKKQDQYNNRWEDERYQREGRGKGHQP | -48.56 | 35.5 | CHHHHHHHHHHHHHHHHHHHHHHHCCCCCCC |
| 501 | EEEEKKQRDQYKNDSEHQRYQRYGRGQGHQP | -48.56 | 32.3 | HHHHHHHHHHHCHHHHHHHHHHHHCCCCCCC |
| 187 | EEEERKQRDQYKNDWEHEEYQRKGRGQGHQP | -48.56 | 29.0 | HHHHHHHHHHHHHHHHHHHHHHHHCCCCCCC |
| 233 | EEEERKQRDQYKNDWEHEEYQRQGRGQGHQP | -48.56 | 29.0 | HHHHHHHHHHHHHHHHHHHHHHHHCCCCCCC |
| 986 | EEERRKQEDQYRNDYEHEEYQRKGRGQGHQP | -48.56 | 29.0 | HHHHHHHHHHHHHHHHHHHHHHHHCCCCCCC |
| 795 | EQEQRINEDFRKNREEQERYQREGQGQGHQS | -48.55 | 25.8 | HHHHHHHHHHHHHHHHHHHHHHHHHCCCCCC |
| 930 | EEERKKQEDQRSNDEEDERYQRQGRGKGHQP | -48.54 | 38.7 | HHHHHHHHHHHHHHHHHHHHHHHHCCCCCCC |
| 290 | EEEERKKEDQRKNEEEDEYYQKYGQGKGHQP | -48.53 | 38.7 | HHHHHHHHHHHHHHHHHHHHHHHHHCCCCCC |
| 435 | EEEERKQRDKQKNEREQEEYQRYGRGQGHQP | -48.53 | 35.5 | HHHHHHHHHHHHHHHHHHHHHHHHHCCCCCC |
| 261 | QEEEKKQRDKQKNEREQKRYQRYGRGQGHQP | -48.53 | 35.5 | CHHHHHHHHHHHHHHHHHHHHHHHHCCCCCC |
| 875 | EQEERKKEDQYRNESEHQRKQNQGKGCGKNH | -48.51 | 25.8 | HHHHHHHHHHHHHHHHHHHHHHCCCCCCCCC |
| 154 | EEEQKKQQDQYSNDYEHEEYQRKGRGQGHQP | -48.49 | 32.3 | HHHHHHHHHHHHHHHHHHHHHHHHCCCCCCC |

| | | | | |
|---|---|---|---|---|
| 248 | EQEERIKKDQEKNEKEQEEYQRRGRGQGHQP | -48.49 | 29.0 | CHHHHHHHHHHHHHHHHHHHHHHHHCCCCCC |
| 070 | EQEERIKQDQKSNEEEQQRYQRYGQGQGHQS | -48.49 | 29.0 | CHHHHHHHHHCHHHHHHHHHHHHHHCCCCCC |
| 613 | EQEERIKQDQQNNEAEQKRYQREGKGQGHNN | -48.49 | 35.5 | CHHHHHHHHHHHHHHHHHHHHHHCCCCCCCC |
| 310 | EEQQRKQEDQRRNDKEQEEYQRYGKGQGHNN | -48.49 | 32.3 | HHHHHHHHHHHHHHHHHHHHHHHHHCCCCCC |
| 701 | EEEERKRQDKEKNEKEHKRKQRYGKGCGKSN | -48.48 | 32.3 | HHHHHHHHHHHHHHHHHHHHHHCCCCCCCCC |
| 387 | EEQQRKQEDKWRNEWEQKYYQKKGQGQGHQP | -48.47 | 38.7 | HHHHHHHHHHHHHHHHHHHHHHHHCCCCCCC |
| 161 | EEEQRKQQDKYNNEWEQKRYQREGRGQGHQP | -48.47 | 38.7 | HHHHHHHHHHHHHHHHHHHHHHHHCCCCCCC |
| 575 | EEEERKQRDQEKNDKEDERYQREGKGKGHSH | -48.47 | 35.5 | HHHHHHHHHHHHHHHHHHHHHHHCCCCCCCC |
| 062 | EEEEKKQQDQRKNEKEHEEYQRRGKGQGHNH | -48.46 | 35.5 | CHHHHHHHHHHHHHHHHHHHHHHHCCCCCCC |
| 022 | EQEERKQQDQRRHENEHKSYQRYGRGQGHQP | -48.45 | 32.3 | HHHHHHHHHHHHHHHHHHHHHHHHCCCCCCC |
| 032 | EQEQRKQEEQRENEREDKEYQREGRGKGHQP | -48.45 | 32.3 | HHHHHHHHHHHHHHHHHHHHHHHHCCCCCCC |
| 230 | EEQEKKNQDQQKNDKEQEEYQRRGWGQGHYP | -48.44 | 35.5 | HHHHHHHHHHHHHHHHHHHHHHHCCCCCCCC |
| 299 | EEEERKNQDQYKNEWEDEEYQRKGRGKGHQP | -48.44 | 38.7 | HHHHHHHHHHHHHHHHHHHHHHHHCCCCCCC |
| 978 | EEEERKNQDQYKNEYEDEEYQRKGRGKGHQP | -48.44 | 38.7 | HHHHHHHHHHHHHHHHHHHHHHHHCCCCCCC |
| 888 | EQEQRKKEDKRKNEEEHKQKQKQGQGCGKQS | -48.44 | 29.0 | HHHHHHHHHHHHHHHHHHHHHHHCCCCCCCC |
| 143 | EEQRRKQEDQYRNDWEHQRKQREGRGCGKQN | -48.44 | 29.0 | HHHHHHHHHHHHHHHHHHHHHHCCCCCCCCC |
| 011 | EEEQRKQEQQRKNDKEDEEYQRRGRGKGHQP | -48.42 | 32.3 | HHHHHHHHHHHHHHHHHHHHHHHHCCCCCCC |
| 691 | EEEERKQRDAWKNRWEDQYYQKQGQGKGHQP | -48.38 | 35.5 | HHHHHHHHHHHHHHHHHHHHHHHHCCCCCCC |
| 848 | EEEQRKQEDQRKNDKEQKRYQREGKGQGHQP | -48.35 | 29.0 | HHHHHHHHHHHHHHHHHHHHHHHHCCCCCCC |
| 371 | QEEQNKQEDQRKNDKEQKRYQREGRGQGHQP | -48.35 | 25.8 | HHHHHHHHHHHHHHHHHHHHHHHHCCCCCCC |
| 843 | EQEERKKQDKEKNEKEHEEYQRRGRGQGHQP | -48.34 | 32.3 | HHHHHHHHHHHHHHHHHHHHHHHHCCCCCCC |
| 419 | EEEERKQQDQRKNDKEHRRYQREGRGQGHQP | -48.34 | 29.0 | HHHHHHHHHHHHHHHHHHHHHHHHCCCCCCC |
| 767 | EEEERKKEDKRKNEKEHKRYQRYGRGQGHQP | -48.33 | 35.5 | HHHHHHHHHHHHHHHHHHHHHHHHCCCCCCC |
| 477 | EEEQRKHEDQNRNRKEDERYQREGKGKGHNN | -48.33 | 35.5 | HHHHHHHHHHHHHHHHHHHHHHHCCCCCCCC |
| 553 | QEQQKKQEDFRRNKKEDEEYQRRGRGKGNQN | -48.32 | 38.7 | HHHHHHHHHHHHHHHHHHHHHHHHCCCCCCC |
| 820 | EEEQRKQEDQERNDKEHQRKQRYGKGCGKNN | -48.32 | 25.8 | HHHHHHHHHHHHHHHHHHHHHHCCCCCCCCC |
| 904 | EEEQRKQHDQQKNEKEHEEYQRRGKGQGHNH | -48.32 | 32.3 | HHHHHHHHHHHHHHHHHHHHHHHHCCCCCCC |
| 399 | EQEERIKEDQYRNEYEQKSYQRYGRGQGHQP | -48.31 | 29.0 | HHHHHHHHHHHHHHHHHHHHHHHHCCCCCCC |
| 392 | EQEERKQQEQRKNDREDRYYQREGKGKGHNH | -48.29 | 29.0 | HHHHHHHHHHHHHHHHHHHHHHCCCCCCCCC |
| 493 | EEQQRKQEQQRRNDKEDRRFQRYGRGKGKQN | -48.29 | 32.3 | HHHHHHHHHHHHHHHHHHHHHHHHCCCCCCC |
| 817 | ENEERKKQEQFNQDFEDEYYQKYGQGKGHQP | -48.29 | 35.5 | CHHHHHHHHHHHHHHHHHHHHHHHHCCCCCC |
| 835 | QEERKKQEDQRSNEKEHKEYQKQGKGRGTSN | -48.29 | 32.3 | HHHHHHHHHHHHHHHHHHHHHHHCCCCCCCC |
| 402 | EEEKRKQEEQRSNEEEDKYYQKKGQGKGHQP | -48.29 | 35.5 | HHHHHHHHHHHHHHHHHHHHHHHHCCCCCCC |
| 662 | EEEERKWQDQKRNREEDERFQKQGNGKGKQH | -48.28 | 32.3 | CHHHHHHHHHHHHHHHHHHHHHHCCCCCCCC |
| 850 | EEEQRKQQDQRNNKKEHEEYQRRGWGQGHYP | -48.27 | 32.3 | HHHHHHHHHHHHHHHHHHHHHHHCCCCCCCC |
| 894 | EEEERKRQDQESNDKEDEEYQRRGRGKGHQP | -48.26 | 35.5 | HHHHHHHHHHHHCCHHHHHHHHHHCCCCCCC |
| 460 | EEEERKRQDQESNDREDERYQREGKGKGHNH | -48.26 | 35.5 | HHHHHHHHHHHHCHHHHHHHHHHCCCCCCCC |
| 030 | EEQQRKQQDHYNQRYEDEEYQRKGRGKGHQP | -48.25 | 41.9 | HHHHHHHHHHHHHHHHHHHHHHHHCCCCCCC |

| | | | | |
|---|---|---|---|---|
| 188 | NDQRRIQEDQERNRKEDEEYQRRGQGKGHQS | -48.23 | 32.3 | CHHHHHHHHHHHHHHHHHHHHHHHHCCCCCC |
| 637 | DQERKKQEDQYRNDYEQQEYQRKGRGQGHQP | -48.23 | 25.8 | HHHHHHHHHHHHHHHHHHHHHHHHCCCCCCC |
| 391 | EQEERKRQDQEKNEKEDEEFQRRGRGKGKQN | -48.22 | 32.3 | HHHHHHHHHHHHHHHHHHHHHHHHCCCCCCC |
| 896 | EEEERKREDKRKNEAEDKRYQREGRGKGHQP | -48.20 | 45.2 | HHHHHHHHHHHHHHHHHHHHHHHHCCCCCCC |
| 303 | EEEERKRQDKENNEREDKRYQREGKGKGHNH | -48.20 | 45.2 | HHHHHHHHHHHHHHHHHHHHHHHCCCCCCCC |
| 206 | EQEQRKQEEQRRNRKEDERYQREGRGKGHQP | -48.19 | 29.0 | HHHHHHHHHHHHHHHHHHHHHHHHCCCCCCC |
| 165 | EEEQRKQQDKYNNEWEDKRYQREGRGKGHQP | -48.18 | 45.2 | HHHHHHHHHHHHHHHHHHHHHHHHCCCCCCC |
| 928 | EQEERKRQDQEKNDKEDRRFQRYGQGKGKQS | -48.16 | 29.0 | HHHHHHHHHHHHHHHHHHHHHHHHHCCCCCC |
| 615 | QEQEKKKQDQQKNDREHEEYQRRGRGQGHQP | -48.16 | 32.3 | CHHHHHHHHHHHHHHHHHHHHHHHHCCCCCC |
| 577 | QEEQKKEEDQRKNDKEDEEYQRRGQGKGHQS | -48.16 | 35.5 | CHHHHHHHHHHHHHHHHHHHHHHHHCCCCCC |
| 194 | EEEERKKQDKYNNEWEHKRYQREGRGQGHQP | -48.16 | 38.7 | HHHHHHHHHHHHHHHHHHHHHHHHCCCCCCC |
| 271 | EQEQRKQEEQRRNEKEDEEYQRRGRGKGHQP | -48.15 | 32.3 | HHHHHHHHHHHHHHHHHHHHHHHHCCCCCCC |
| 929 | EEEERKKQDQEKNEREDEEYQRRGQGKGHQS | -48.15 | 38.7 | HHHHHHHHHHHHHHHHHHHHHHHHHCCCCCC |
| 439 | EEEERKKQDQEKNEREDEEYQRRGRGKGHQP | -48.15 | 38.7 | HHHHHHHHHHHHHHHHHHHHHHHHCCCCCCC |
| 398 | EEEKKKEEDQQRHEREDEEYQRRGQGKGHQS | -48.14 | 45.2 | HHHHHHHHHHHHHHHHHHHHHHHHHCCCCCC |
| 010 | EEEQKKEEDQKKNEQEDKRYQREGRGKGHQP | -48.13 | 41.9 | CHHHHHHHHHHHHHHHHHHHHHHHCCCCCCC |
| 286 | EEEQRKEEDQKKNEKEDEEYQRRGWGKGHYP | -48.13 | 38.7 | HHHHHHHHHHHHHHHHHHHHHHHCCCCCCCC |
| 257 | EEEQRKEEDQKKNEQEDQRYQREGQGKGHQS | -48.13 | 38.7 | HHHHHHHHHHHHHHHHHHHHHHHHCCCCCCC |
| 193 | EEEEKKKRDQESNDKEHRRYQREGRGRGHQP | -48.12 | 32.3 | CHHHHHHHHHHHHHHHHHHHHHHHCCCCCCC |
| 481 | QDQERKRQDQEKNDKEHKRKQRYGRGCGKQN | -48.12 | 22.6 | CHHHHHHHHHHHHHHHHHHHHHHCCCCCCCC |
| 147 | EEEERKKQDQYRNEWEHEEYQRKGRGRGHQP | -48.11 | 32.3 | HHHHHHHHHHHHHHHHHHHHHHHHCCCCCCC |
| 957 | EEQRRKQEDQERNDKEDKRYQREGKGKGHNH | -48.10 | 38.7 | HHHHHHHHHHHHHHHHHHHHHHHCCCCCCCC |
| 289 | EQQERKNQEQWKNEWEDQYFQQKGQGKGKQN | -48.10 | 32.3 | HHHHHHHHHHHHHHHHHHHHHHHHCCCCCCC |
| 977 | EEEQRKQEQQYRNEFEDQMYQRYGKGKGTQN | -48.10 | 35.5 | HHHHHHHHHHHHHHHHHHHHHHHHHCCCCCC |
| 319 | EEEQRKQQDQRSNDKEQQEYQKQGRGQGHQP | -48.09 | 29.0 | HHHHHHHHHHHHHHHHHHHHHHHHCCCCCCC |
| 212 | EEEERKKQDQWKNDWEHKYYQKKGQGQGHNH | -48.09 | 29.0 | HHHHHHHHHHHHHHHHHHHHHHHHCCCCCCC |
| 569 | EEQERKKQDQKKNDEEHERYQQQGKGQGHNN | -48.08 | 32.3 | HHHHHHHHHHHHHHHHHHHHHHHCCCCCCCC |
| 088 | EEEERKRQDQEKNEKEHSEYQQRGRGRGHQP | -48.08 | 32.3 | HHHHHHHHHHHHHHHHHHHHHHHHCCCCCCC |
| 529 | EEERKKQEDQYRNEWEDEEFQRKGRGKGKQN | -48.08 | 38.7 | HHHHHHHHHHHHHHHHHHHHHHHHCCCCCCC |
| 159 | DDERRKQHDQEKNDKEDEQYQRRGRGKGHQP | -48.07 | 29.0 | CHHHHHHHHHHHCHHHHHHHHHHHCCCCCCC |
| 444 | EEERRKQEDQERQEKEDKRYQREGDGKGHQP | -48.07 | 38.7 | HHHHHHHHHHHHHHHHHHHHHHHCCCCCCCC |
| 630 | QEERKKQEDQERQEKEDQYYQKEGQGKGHQP | -48.07 | 38.7 | HHHHHHHHHHHHHHHHHHHHHHHHCCCCCCC |
| 625 | EEEERKKEDQKRNDKEHEEYQRRGRGQGHQP | -48.04 | 29.0 | HHHHHHHHHHHHHHHHHHHHHHHHHCCCCCC |
| 341 | EEEERKKQDQYNNDWEHERYQREGWGQGHYP | -48.04 | 32.3 | HHHHHHHHHHHCHHHHHHHHHHHHCCCCCCC |
| 995 | QEEEKKKEDQKRNDEEHKRYQREGRGQGHQP | -48.04 | 29.0 | CHHHHHHHHHHHHHHHHHHHHHHHCCCCCCC |
| 237 | EEEERKRQDQDSNDREHEEKQRRGQGCGKQS | -48.04 | 25.8 | HHHHHHHHHHCHCHHHHHHHHHHCCCCCCCC |
| 353 | QREKEIQEQQRSNREEHERYQKQGRGQGHQP | -48.03 | 19.4 | CHHHHHHHHHHHHHHHHHHHHHHCCCCCCCC |
| 931 | NDEQRKKEDQRKNDEEHKRYQREGRGQGHQP | -48.02 | 22.6 | CHHHHHHHHHHCHHHHHHHHHHHHCCCCCCC |

| | | | | |
|---|---|---|---|---|
| 148 | QEEEKKQQDKRKNEQEHRRYQREGRGQGHQP | -48.02 | 35.5 | CHHHHHHHHHHHHHHHHHHHHHHHHCCCCCCC |
| 317 | EEQQRKQQDHRSNRQEDQRYQKEGWGKGHYP | -48.00 | 38.7 | HHHHHHHHHHHHHHHHHHHHHHHHHCCCCCCC |
| 404 | EEEKRKQEDFQKNRKEHERYQKEGQGQGHQS | -47.99 | 29.0 | HHHHHHHHHHHHHHHHHHHHHHHHHCCCCCCC |
| 275 | EEQQRKQEQQRRNDKEDRRYQREGRGKGHQP | -47.98 | 35.5 | HHHHHHHHHHHHHHHHHHHHHHHHHCCCCCCC |
| 810 | QKEQEIQEQKRRNEKEQKRYQREGKGQGHNH | -47.97 | 25.8 | CHHHHHHHHHHHHHHHHHHHHHHHHCCCCCCC |
| 609 | EEERRKQEDKERNEKEDKRYQREGKGKGHNH | -47.97 | 41.9 | HHHHHHHHHHHHHHHHHHHHHHHHCCCCCCCC |
| 530 | EEEERKKQDQQKNDKEDREYQRRGQGKGHQP | -47.95 | 35.5 | HHHHHHHHHHHHHHHHHHHHHHHHHCCCCCCC |
| 887 | EQEERKKQDQEKNDKEDRRYQREGRGKGHNH | -47.95 | 32.3 | HHHHHHHHHHHHHHHHHHHHHHHHHCCCCCCC |
| 617 | EEERKKQEDQERNDKEHRRYQREGRGRGHQP | -47.95 | 32.3 | HHHHHHHHHHHHHHHHHHHHHHHHHCCCCCCC |
| 409 | EEEEKKKQDQQKNEKEDEEYQRRGQGKGHQS | -47.94 | 41.9 | CHHHHHHHHHHHHHHHHHHHHHHHHHCCCCCC |
| 349 | EEEERKKQDQIKNEKEDEEYQRYGRGKGHQP | -47.94 | 38.7 | HHHHHHHHHHHHHHHHHHHHHHHHHHCCCCCC |
| 045 | EEEERKKQDQYKNEYEDQEYQRKGRGKGHQP | -47.94 | 38.7 | HHHHHHHHHHHHHHHHHHHHHHHHHCCCCCCC |
| 710 | EEEQRKKEDQRKNEEEDKRYQREGRGKGHQP | -47.94 | 38.7 | HHHHHHHHHHHHHHHHHHHHHHHHHCCCCCCC |
| 375 | EEEQRKKEDQRKNEEEDQRYQKEGRGKGHQP | -47.94 | 38.7 | HHHHHHHHHHHHHHHHHHHHHHHHHCCCCCCC |
| 548 | EEEQRKKEDQRKNEEEDQRYQREGRGKGHQP | -47.94 | 38.7 | HHHHHHHHHHHHHHHHHHHHHHHHHCCCCCCC |
| 616 | EQEERKKQDQEKNEREDERYQRQGRGKGHQP | -47.94 | 35.5 | HHHHHHHHHHHHHHHHHHHHHHHHCCCCCCCC |
| 259 | EQEERKKQDQQKNEKEDEEYQQKGRGKGHQP | -47.94 | 35.5 | HHHHHHHHHHHHHHHHHHHHHHHHHCCCCCCC |
| 725 | QEEEKKKQDQEKNEKEDKRYQREGRGKGHQP | -47.94 | 38.7 | CHHHHHHHHHHHHHHHHHHHHHHHHCCCCCCC |
| 792 | QEEEKKKQDQQKNEKEDEEYQRRGKGKGHNN | -47.94 | 38.7 | CHHHHHHHHHHHHHHHHHHHHHHHCCCCCCCC |
| 362 | QEEEKKKQDQYRNEWEDEEYQRQGRGKGHQP | -47.94 | 38.7 | CHHHHHHHHHHHHHHHHHHHHHHHHCCCCCCC |
| 823 | QQEEEKKQDQYKNEWEDEEYQRKGRGKGHQP | -47.94 | 32.3 | CHHHHHHHHHHHHHHHHHHHHHHHHCCCCCCC |
| 566 | EEERRKQEDQERNDKEQKRYQREGKGQGHNH | -47.94 | 29.0 | HHHHHHHHHHHHHHHHHHHHHHHHHCCCCCCC |
| 461 | EEEERKRQDQEKNDKEDRRYQRYGQGKGHQS | -47.93 | 35.5 | HHHHHHHHHHHHCHHHHHHHHHHHHCCCCCCC |
| 612 | EEQERKHQDQNKNRKEDEEYQKQGRGKGHQP | -47.92 | 38.7 | HHHHHHHHHHHHHHHHHHHHHHHHCCCCCCCC |
| 560 | EQEQRKQEDKRRHEKEQKRYQREGRGQGHQP | -47.92 | 35.5 | HHHHHHHHHHHHHHHHHHHHHHHHHCCCCCCC |
| 714 | EEQQKKNEEQQSNDREDRRYQREGRGKGHQP | -47.92 | 38.7 | HHHHHHHHHHHHHHHHHHHHHHHHHCCCCCCC |
| 417 | EEERRKQEDQERHEKEDKRYQRYGRGKGHQP | -47.92 | 41.9 | HHHHHHHHHHHHHHHHHHHHHHHHHCCCCCCC |
| 539 | EEERRKQEDQYRHEWEDEYYQRKGQGKGHQP | -47.92 | 41.9 | HHHHHHHHHHHHHHHHHHHHHHHHHCCCCCCC |
| 749 | EEEEKKRQDQEKNEKEDEEYQRRGKGKGHNH | -47.91 | 41.9 | CHHHHHHHHHHHHHHHHHHHHHHHHCCCCCCC |
| 144 | EEEEKKRQDQEKNEKEDKRYQREGRGKGHQP | -47.91 | 41.9 | CHHHHHHHHHHHHHHHHHHHHHHHHCCCCCCC |
| 356 | EEEERKRQDQEKNEKEDERYQREGWGKGHYP | -47.91 | 38.7 | HHHHHHHHHHHHHHHHHHHHHHHHHCCCCCCC |
| 793 | EEEERKRQDQEKNEKEDKRYQRYGRGKGHQP | -47.91 | 38.7 | HHHHHHHHHHHHHHHHHHHHHHHHHCCCCCCC |
| 403 | EEEERKRQDQEKNEKEDKRYQRYGWGKGHYP | -47.91 | 38.7 | HHHHHHHHHHHHHHHHHHHHHHHHCCCCCCCC |
| 296 | EEEERKRQDQEKNEKEDQRYQREGRGKGHQP | -47.91 | 38.7 | HHHHHHHHHHHHHHHHHHHHHHHHHCCCCCCC |
| 802 | EQEERKRQDQEKNEKEDEEYQRKGRGKGHQP | -47.91 | 35.5 | HHHHHHHHHHHHHHHHHHHHHHHHHCCCCCCC |
| 874 | QEEEKKRQDQEKNEKEDEEYQRRGRGKGHQP | -47.91 | 38.7 | CHHHHHHHHHHHHHHHHHHHHHHHHCCCCCCC |
| 666 | QEEEKKRQDQEKNEKEDKRYQREGKGKGHNP | -47.91 | 38.7 | CHHHHHHHHHHHHHHHHHHHHHHHCCCCCCCC |
| 622 | EEQQRKQQQQRSNEKEDEEYQRRGRGKGHQP | -47.91 | 38.7 | HHHHHHHHHHHHHHHHHHHHHHHHHCCCCCCC |

| | | | | |
|---|---|---|---|---|
| 756 | EEEQRKKQDQINNEKEDEEYQRRGRGKGHQP | -47.87 | 41.9 | HHHHHHHHHHHHHHHHHHHHHHHHCCCCCCC |
| 518 | EREQRYQEDQRKNDQEHKRYQRYGRGQGHQP | -47.87 | 25.8 | HHHHHHHHHHHHHHHHHHHHHHHHCCCCCCC |
| 450 | EEEKRKQEQQRENDREHREYQREGKGRGHNP | -47.86 | 25.8 | HHHHHHHHHHHHHHHHHHHHHHHCCCCCCCC |
| 494 | EEEERKQQDQRKHDYEQKEYQKEGRGQGHQP | -47.85 | 32.3 | HHHHHHHHHHHHHHHHHHHHHHHHCCCCCCC |
| 882 | EEQQRKQQQKYSNEYEDKEYQRKGRGKGNQN | -47.85 | 41.9 | HHHHHHHHHHHHHHHHHHHHHHHHCCCCCCC |
| 378 | EEEQRKQYDQKKNEEEDKRYQREGRGKGHQP | -47.84 | 38.7 | HHHHHHHHHHHHHHHHHHHHHHHHCCCCCCC |
| 966 | EEEKRKQLDQENHEKEDEEYQRRGRGKGHQP | -47.84 | 48.4 | HHHHHHHHHHHHHHHHHHHHHHHHCCCCCCC |
| 733 | EEQQRKQEDKRKNEKEQKRYQREGRGQGHQP | -47.84 | 38.7 | HHHHHHHHHHHHHHHHHHHHHHHHCCCCCCC |
| 453 | EEEERKSRDQQKNEKEDQEYQREGSGKGHQR | -47.83 | 41.9 | CHHHHHHHHHHHHHHHHHHHHHHCCCCCCCC |
| 528 | EEEERKKQDKKKNEEEHKRYQRYGQGQGHQS | -47.82 | 35.5 | HHHHHHHHHHHHHHHHHHHHHHHHCCCCCCC |
| 229 | EEQERKRQDQESNDREDQYYQKKGQGKGHQP | -47.82 | 38.7 | HHHHHHHHHHHHHHHHHHHHHHHHCCCCCCC |
| 605 | EEEQRKQEQQRKNRKEDERYQREGQGKGHQS | -47.81 | 32.3 | HHHHHHHHHHHHHHHHHHHHHHHHCCCCCCC |
| 140 | EEEKKKQEDQRKNDEEHRRYQRYGRGQGHQP | -47.80 | 32.3 | HHHHHHHHHHHHHHHHHHHHHHHHHCCCCCC |
| 480 | EEERKKQQDQYSNEYEDEEYQRKGRGKGHQP | -47.77 | 41.9 | HHHHHHHHHHHHHHHHHHHHHHHHCCCCCCC |
| 273 | EEERRKQEDQERNEKEDERYQQQGRGKGHQP | -47.77 | 38.7 | HHHHHHHHHHHHHHHHHHHHHHHHCCCCCCC |
| 256 | EEERRKQEDQERNEKEDKYYQKKGQGKGHQP | -47.77 | 38.7 | HHHHHHHHHHHHHHHHHHHHHHHHCCCCCCC |
| 653 | EEERRKQEDQERNEKEDQRYQNQGQGKGHQS | -47.77 | 38.7 | HHHHHHHHHHHHHHHHHHHHHHHCCCCCCCC |
| 781 | EEERRKQEDQRKNEEEDQRYQRYGRGKGHQP | -47.77 | 38.7 | HHHHHHHHHHHHHHHHHHHHHHHHCCCCCCC |
| 741 | EEERRKQLDQESNEREDEEYQRRGKGKGHNH | -47.77 | 41.9 | HHHHHHHHHHHHHHHHHHHHHHHHCCCCCCC |
| 634 | EEEERKQRDQYKNDYEQQEYQKQGQGQGHQS | -47.75 | 29.0 | HHHHHHHHHHHHHHHHHHHHHHHHCCCCCCC |
| 231 | QEEENKQRDQWKNDWEQQYYQKQGQGQGHQP | -47.75 | 25.8 | CHHHHHHHHHHHHHHHHHHHHHHHCCCCCCC |
| 395 | EQEERKQREQKSNREEDERYQRYGQGKGHQS | -47.75 | 29.0 | HHHHHHHHHHHHHHHHHHHHHHHHHCCCCCC |
| 812 | EEEERKQQQRENEREDEEYQRKGQGKGHFS | -47.75 | 38.7 | HHHHHHHHHHHHHHHHHHHHHHHHCCCCCCC |
| 068 | QEERKKQEDQERNEKEHEEYQRRGRGRGHQP | -47.75 | 32.3 | HHHHHHHHHHHHHHHHHHHHHHHHCCCCCCC |
| 137 | EEEERKKQDQEKNRKEDERYQREGKGKGHNH | -47.73 | 35.5 | HHHHHHHHHHHHHHHHHHHHHHHCCCCCCCC |
| 782 | EEEQKIQEQQRRNQKEHERYQRQGRGQGHQP | -47.71 | 29.0 | HHHHHHHHHHHHHHHHHHHHHHHCCCCCCCC |
| 567 | EEEQRKKEDQRKNEKEDQEYQKQGQGKGHQS | -47.71 | 38.7 | HHHHHHHHHHHHHHHHHHHHHHHHCCCCCCC |
| 099 | DEERKKQHDQESNDKEDRRYQREGRGKGHQP | -47.70 | 35.5 | HHHHHHHHHHHHCHHHHHHHHHHHCCCCCCC |
| 573 | DEERKKQEDQERNEKEDERYQREGQGKGHNS | -47.69 | 38.7 | CHHHHHHHHHHHHHHHHHHHHHHCCCCCCCC |
| 155 | DEERKKQHDQESNEKEDKRYQRYGKGKGHNH | -47.69 | 38.7 | HHHHHHHCHHHHHHHHHHHHHHHHCCCCCCC |
| 041 | EEEERKQRDQESNEKEDERYQQQGRGKGHQP | -47.69 | 38.7 | HHHHHHHHHHHHHHHHHHHHHHHHCCCCCCC |
| 895 | EQEQRKQEEQYRNDSEDRRYQQQGRGKGHQP | -47.68 | 29.0 | HHHHHHHHHHHHHHHHHHHHHHHHCCCCCCC |
| 502 | EQEKRKQEDKRKNEEEQKRYQRQGRGQGHQP | -47.67 | 32.3 | HHHHHHHHHHHHHHHHHHHHHHHHCCCCCCC |
| 852 | EEEERKQRDQEKNRKEDEYYQKYGQGKGHQP | -47.67 | 35.5 | HHHHHHHHHHHHHHHHHHHHHHHHHCCCCCC |
| 035 | QEERKKQHDQESNRKEDERYQREGRGKGHQP | -47.67 | 35.5 | HHHHHHHHHHHHHHHHHHHHHHHHCCCCCCC |
| 376 | EEEQKKQQDKYSNEYEQEEYQRQGRGQGHQP | -47.67 | 38.7 | HHHHHHHHHHHHHHHHHHHHHHHHCCCCCCC |
| 021 | EEEEKKKRDQQKNDREQEEYQRRGRGQGHQP | -47.66 | 32.3 | CHHHHHHHHHHHHHHHHHHHHHHHHCCCCCC |
| 463 | EEQQKKQEDQRRNEKEQQRYQREGRGQGHQP | -47.65 | 38.7 | HHHHHHHHHHHHHHHHHHHHHHHHCCCCCCC |

| | | | | |
|---|---|---|---|---|
| 655 | EEEKKKQEQQRSNEKEDEEYQRRGQGKGHQS | -47.65 | 38.7 | HHHHHHHHHHHHHHHHHHHHHHHHHCCCCCC |
| 162 | QREEQIQREQYKNEYEQEEYQRKGRGQGHQP | -47.65 | 22.6 | CHHHHHHHHHHHHHHHHHHHHHHHCCCCCCC |
| 889 | QREKQIQEEQRSNEEEQQRYQRQGKGQGHNN | -47.65 | 22.6 | HHHHHHHHHHHHHHHHHHHHHHHCCCCCCCC |
| 263 | EQEQRKQEDQRRNEKEQKRYQREGKGQGHNN | -47.65 | 29.0 | HHHHHHHHHHHHHHHHHHHHHHHCCCCCCCC |
| 089 | EEEERKREDQWRHEYEDEEYQRKGRGKGHQP | -47.63 | 41.9 | CHHHHHHHHHHHHHHHHHHHHHHHCCCCCCC |
| 979 | EEEKRKEEDQRKNDEEDRRYQRYGRGKGHQP | -47.63 | 35.5 | HHHHHHHHHHHHHHHHHHHHHHHHCCCCCCC |
| 777 | EEEERKSRDQQKNEREDEEYQRRGRGKGHQP | -47.59 | 38.7 | CHHHHHHHHHHHHHHHHHHHHHHHCCCCCCC |
| 007 | EQEERLNRQQESNDKEDQRYQREGRGKGHQP | -47.59 | 29.0 | HHHHHHHHHHHHCHHHHHHHHHHHCCCCCCC |
| 620 | EEEERKQRDQQKNEKEQKRYQREGRGQGHQP | -47.59 | 32.3 | HHHHHHHHHHHHHHHHHHHHHHHHCCCCCCC |
| 160 | EEQERKQQDQQSNEREQERYQQQGRGQGHQP | -47.58 | 35.5 | HHHHHHHHHHHHHHHHHHHHHHHHCCCCCCC |
| 604 | QDEQRKNEDQQKNDKEDRRYQREGRGKGHQP | -47.56 | 29.0 | CHHHHHHHHHHHHHHHHHHHHHHHCCCCCCC |
| 716 | EEQQRKQEDQYRNRYEQEEYQRKGRGQGHQP | -47.56 | 32.3 | HHHHHHHHHHHHHHHHHHHHHHHHCCCCCCC |
| 755 | EEERRKQEDQWRNDWEDRYYQKEGQGKGHQP | -47.54 | 35.5 | HHHHHHHHHHHHHHHHHHHHHHHCCCCCCCC |
| 721 | EQQQRKQEEQYRNDWEHQRYQREGRGQGHQP | -47.53 | 25.8 | HHHHHHHHHHHHHHHHHHHHHHHHCCCCCCC |
| 674 | EQEERKQREQRNNDYEHEEYQRKGRGQGHQP | -47.53 | 25.8 | HHHHHHHHHHHHHHHHHHHHHHHHCCCCCCC |
| 504 | EEEERKKQQKYSNEYEDEEYQRKGRGKGHQP | -47.53 | 38.7 | HHHHHHHHHHHHHHHHHHHHHHHHCCCCCCC |
| 202 | EEQQRKQEDQYRNEWEQKRYQREGKGQGHNN | -47.52 | 35.5 | HHHHHHHHHHHHHHHHHHHHHHHCCCCCCCC |
| 327 | EEEQKKQRDQYKNEWEQEEYQRQGRGQGHQP | -47.52 | 35.5 | HHHHHHHHHHHHHHHHHHHHHHHHCCCCCCC |
| 432 | EEEERKQQDQRKNEKEQEEYQRRGRGQGHQP | -47.52 | 32.3 | HHHHHHHHHHHHHHHHHHHHHHHHHCCCCCC |
| 514 | QEEEKKQQDQRRNEKEQEEYQRRGRGQGHQP | -47.52 | 32.3 | CHHHHHHHHHHHHHHHHHHHHHHHHCCCCCC |
| 429 | EEQQRKKEDQRKNEEEDERYQQQGKGKGHSS | -47.50 | 41.9 | HHHHHHHHHHHHHHHHHHHHHHHCCCCCCCC |
| 772 | EEEERKREDQWRNDWEDRYYQKKGQGKGHQP | -47.49 | 35.5 | HHHHHHHHHHHHHHHHHHHHHHHCCCCCCCC |
| 855 | EEEQKKQEDDRRNRKEDQRYQREGRGKGHQP | -47.49 | 38.7 | HHHHHHHHHHHHHHHHHHHHHHHHCCCCCCC |
| 555 | EEEQRKEEDQKRNEREDEYYQQQGRGKGHQP | -47.49 | 38.7 | HHHHHHHHHHHHHHHHHHHHHHHHCCCCCCC |
| 128 | EEEEKKRKDQYDNEWEDEEYQRKGRGKGHQP | -47.48 | 41.9 | CHHHHHHHHHCCCHHHHHHHHHHHCCCCCCC |
| 906 | EEEEKKRRDQESNEEEDKRYQRYGRGKGHQP | -47.48 | 41.9 | CHHHHHHHHHHHHHHHHHHHHHHHCCCCCCC |
| 197 | EEEERKRQDQENNEREDKRYQREGKGKGHNH | -47.48 | 41.9 | HHHHHHHHHHHHHHHHHHHHHHHCCCCCCCC |
| 342 | EEEERKRQDQESNEREDQRYQREGRGKGHQP | -47.48 | 38.7 | HHHHHHHHHHHHHHHHHHHHHHHHCCCCCCC |
| 942 | EEEERKRQDQESNEYEDEEYQRKGRGKGHQN | -47.48 | 38.7 | HHHHHHHHHHHHHHHHHHHHHHHHCCCCCCC |
| 141 | EEEEKKKQDQQKNDKEDEEYQRYGRGKGDQR | -47.47 | 45.2 | CHHHHHHHHHHHHHHHHHHHHHHCCCCCCCC |
| 511 | EEEKRKQEDQRKNDEEQRRYQRYGRGQGHQP | -47.46 | 29.0 | HHHHHHHHHHHHHHHHHHHHHHHHHCCCCCC |
| 484 | EEEERKREDQQRNREEDERYQKQGRGKGHQP | -47.46 | 35.5 | HHHHHHHHHHHHHHHHHHHHHHHCCCCCCCC |
| 638 | EEEKRKQEQQRSNEEEDKRYQREGRGKGHQP | -47.46 | 35.5 | HHHHHHHHHHHHHHHHHHHHHHHHCCCCCCC |
| 949 | EEEKRKQEQQRSNEEEDKRYQRYGRGKGHQP | -47.46 | 35.5 | HHHHHHHHHHHHHHHHHHHHHHHHCCCCCCC |
| 066 | EQEQRKQQDQQNNRREDERYQKQGRGKGHQP | -47.45 | 35.5 | HHHHHHHHHHHHHHHHHHHHHHHCCCCCCCC |
| 780 | EEEQRKKNDQYNNDWEDRYYQQYGQGKGHQP | -47.44 | 38.7 | HHHHHHHHHHCCHHHHHHHHHHHHHCCCCCC |
| 216 | EEEKRKQEDQRKNREEQEYYQKYGQGQGHQS | -47.43 | 29.0 | HHHHHHHHHHHHHHHHHHHHHHHHHCCCCCC |
| 167 | EEEQRKQEDDQKNRKEDQRYQREGRGKGHQP | -47.43 | 35.5 | HHHHHHHHHHHHHHHHHHHHHHHHCCCCCCC |

| | | | | |
|---|---|---|---|---|
| 355 | EEQRRKQEDQQKNRREDEYYQKYGQGKGHQP | -47.42 | 38.7 | HHHHHHHHHHHHHHHHHHHHHHHHHCCCCCC |
| 475 | EEEERKKQQQYKNDWEDEEYQRKGRGKGHQP | -47.41 | 32.3 | HHHHHHHHHHHHHHHHHHHHHHHHCCCCCCC |
| 079 | EEEEKKKRDKQKNEREQEEYQRRGRGQGHQP | -47.41 | 38.7 | CHHHHHHHHHHHHHHHHHHHHHHHHCCCCCC |
| 264 | QQQQNHQEEKRRNEKEDKRYQREGRGKGHQP | -47.40 | 35.5 | CHHHHHHHHHHHHHHHHHHHHHHHCCCCCCC |
| 464 | EEEERKKQDKEKNEKEQKRYQREGRGQGHQP | -47.40 | 35.5 | HHHHHHHHHHHHHHHHHHHHHHHHCCCCCCC |
| 876 | EEEKRKQEEQRKNDEEHKRYQKEGQGQGHQS | -47.40 | 25.8 | HHHHHHHHHHHHHHHHHHHHHHHHCCCCCCC |
| 175 | EQEKRKQEEQRSNDEEHKRYQREGRGQGHQP | -47.40 | 22.6 | HHHHHHHHHHHHHHHHHHHHHHHHCCCCCCC |
| 405 | EEEEKRRRDKEKNEREDKRYQREGRGKGHQP | -47.40 | 45.2 | HHHHHHHHHHHHHHHHHHHHHHHHCCCCCCC |
| 908 | EEQKRKQEDQRKNEEEQERYQKQGRGQGHQP | -47.39 | 35.5 | HHHHHHHHHHHHHHHHHHHHHHHHCCCCCCC |
| 006 | EEQKRKQEDQRKNEKEQEEYQRYGKGQGHNN | -47.39 | 35.5 | HHHHHHHHHHHHHHHHHHHHHHHHHCCCCCC |
| 563 | EEEKKKEEDKRKNEQEDKRYQREGDGKGNSN | -47.39 | 45.2 | HHHHHHHHHHHHHHHHHHHHHHHCCCCCCCC |
| 588 | EEEKRKQEDQRKNEEEQEYYQRKGQGQGHQP | -47.39 | 32.3 | HHHHHHHHHHHHHHHHHHHHHHHHCCCCCCC |
| 606 | EEEKRKQEDQRSNEKEQEEYQRRGQGQGHQS | -47.39 | 32.3 | HHHHHHHHHHHHHHHHHHHHHHHHHCCCCCC |
| 526 | EEEQRKQEDQRKNEEEQNYYQKEGQGQGHNH | -47.39 | 32.3 | HHHHHHHHHHHHHHHHHHHHHHHHCCCCCCC |
| 545 | EEEERKQQDDRKNRKEDEEYQRYGKGKGHNN | -47.36 | 35.5 | HHHHHHHHHHHHHHHHHHHHHHHHCCCCCCC |
| 959 | EQEQRKNEDQRNNEEEQERYQRYGKGQGHNH | -47.34 | 32.3 | HHHHHHHHHHHCHHHHHHHHHHHHHCCCCCC |
| 380 | EQEERKQQEQRKNDKEHEEYQRRGRGQGHQP | -47.34 | 22.6 | HHHHHHHHHHHHHHHHHHHHHHHHHCCCCCC |
| 423 | EQQERKKQDQQKNDREQEEYQRRGRGQGHQP | -47.33 | 29.0 | HHHHHHHHHHHHHHHHHHHHHHHHHCCCCCC |
| 112 | EEEERKQQDQRKNEKEQKEYQRSGQGQGHQP | -47.33 | 35.5 | HHHHHHHHHHHHHHHHHHHHHHHCCCCCCCC |
| 856 | EEEQKKQEDQRKNEKEQQRYQKEGRGQGHQP | -47.33 | 35.5 | HHHHHHHHHHHHHHHHHHHHHHHHCCCCCCC |
| 076 | EEEQRKQEDQRKNEKEQQRYQREGRGQGHQP | -47.33 | 32.3 | HHHHHHHHHHHHHHHHHHHHHHHHCCCCCCC |
| 118 | EQEERKQQDQRKNENEQKRYQREGKGQGHNN | -47.33 | 29.0 | HHHHHHHHHHHHHHHHHHHHHHHCCCCCCCC |
| 712 | QEEEKKQQDQRKNEKEQEEYQRRGRGQGHQP | -47.33 | 32.3 | CHHHHHHHHHHHHHHHHHHHHHHHHCCCCCC |
| 551 | EEEQKKQRQQRSHEKEDEEYQRRGWGKGHYP | -47.28 | 41.9 | HHHHHHHHHHHHHHHHHHHHHHHCCCCCCCC |
| 925 | EEEQRKQEDDQKNRREHEEKQRRGQGCGKQN | -47.28 | 25.8 | HHHHHHHHHHHHHHHHHHHHHHHCCCCCCCC |
| 719 | EQEERKKEDQRRNDKEQKYYQQQGQGQGHNP | -47.27 | 25.8 | HHHHHHHHHHHHHHHHHHHHHHHHCCCCCCC |
| 102 | EQEQRHQEEKRKNEKEDEEYQRRGKGKGHNH | -47.27 | 35.5 | HHHHHHHHHHHHHHHHHHHHHHHHCCCCCCC |
| 492 | EQEERKQRDQWKNDWEQRYYQKKGQGQGHNH | -47.27 | 25.8 | HHHHHHHHHHHHHHHHHHHHHHHHCCCCCCC |
| 579 | EQEERKKQDQQKNEEEHKRYQRYGRGQGHQP | -47.26 | 29.0 | HHHHHHHHHHHHHHHHHHHHHHHHCCCCCCC |
| 486 | NDQQRKQEDQRRNEKEQEEYQRRGRGQGHQP | -47.26 | 29.0 | CHHHHHHHHHHHHHHHHHHHHHHHHCCCCCC |
| 973 | EEERKKQEDQERNEKEDQRYQRYGRGKGHQP | -47.25 | 41.9 | HHHHHHHHHHHHHHHHHHHHHHHHCCCCCCC |
| 210 | EQEERHQQEKRENEREDKEYQRYGRGKGHQP | -47.24 | 35.5 | CHHHHHHHHHHHHHHHHHHHHHHHCCCCCCC |
| 568 | EEEQRKNEDQRKNDEEHERYRQQGRGQGHQP | -47.22 | 25.8 | HHHHHHHHHHHCHHHHHHHHHHHCCCCCCCC |
| 938 | EQQQRIQEQQRRHEKEQEEYQRRGRGQGHQP | -47.22 | 32.3 | HHHHHHHHHHHHHHHHHHHHHHHHHCCCCCC |
| 907 | EEEERKKEDKRRNEEEQKRYQRYGRGQGHQP | -47.22 | 35.5 | HHHHHHHHHHHHHHHHHHHHHHHHHCCCCCC |
| 097 | EEEERKRRDQERHDREHEYYQQQGRGQGHQP | -47.20 | 32.3 | HHHHHHHHHHHHHHHHHHHHHHHHCCCCCCC |
| 760 | EEEKKKQEDQRKNEEEQEEYQRKGRGQGHQP | -47.20 | 35.5 | HHHHHHHHHHHHHHHHHHHHHHHHCCCCCCC |
| 094 | EQEKRKQEDQRKNEEEQKSYQKEGRGQGHQP | -47.20 | 29.0 | HHHHHHHHHHHHHHHHHHHHHHHHCCCCCCC |

| | | | | |
|---|---|---|---|---|
| 557 | QEEKNKQEDQRKNEEEQERYQREGWGQGHYP | -47.20 | 29.0 | CHHHHHHHHHHHHHHHHHHHHHHHCCCCCCC |
| 281 | EEEQRKEEDQKKNDKEHEEYQRYGRGQGHQP | -47.19 | 29.0 | HHHHHHHHHHHHHHHHHHHHHHHHHCCCCCC |
| 524 | EEEKRKQEDARKNREEHERYQQQGRGQGHQP | -47.18 | 29.0 | HHHHHHHHHHHHHHHHHHHHHHHHCCCCCCC |
| 228 | EEEERKQQDDRKNRWEDEYYQRKGQGKGHNH | -47.17 | 35.5 | HHHHHHHHHHHHHHHHHHHHHHHHCCCCCCC |
| 805 | EQEERKQRDDRKNRQEDQRYQREGRGKGHQP | -47.17 | 32.3 | HHHHHHHHHHHHHHHHHHHHHHHHCCCCCCC |
| 564 | EEERRKQEDQERNDKEDEEYQRRGRGKGNQN | -47.16 | 35.5 | HHHHHHHHHHHHHHHHHHHHHHHHCCCCCCC |
| 672 | EQEKRHQEEKRSNEEEDKRYQREGQGKGHQS | -47.14 | 35.5 | HHHHHHHHHHHHHHHHHHHHHHHHCCCCCCC |
| 072 | EQERRHQEEKKRNEEEDKRYQKQGRGKGHQP | -47.14 | 35.5 | HHHHHHHHHHHHHHHHHHHHHHHCCCCCCCC |
| 183 | EQEKRIQEDHRKNREEQQRYQRYGRGQGHQP | -47.12 | 25.8 | HHHHHHHHHHHHHHHHHHHHHHHHHCCCCCC |
| 245 | EEEERKQQDARKNRKEHEEYQRYGRGQGHQP | -47.11 | 29.0 | HHHHHHHHHHHHHHHHHHHHHHHHHCCCCCC |
| 813 | EEEQRKNEDDRKNREEDQRYQREGDGKGHQP | -47.11 | 35.5 | HHHHHHHHHHHHHHHHHHHHHHHCCCCCCCC |
| 186 | EQQQRHQQEQYSNDYEDEEYQRKGRGKGHQP | -47.11 | 32.3 | HHHHHHHHHHHHCCHHHHHHHHHHCCCCCCC |
| 507 | EEEERKKQDQKNNDEEQKRYQQYGRGQGHQP | -47.10 | 32.3 | HHHHHHHHHHHCHHHHHHHHHHHHHCCCCCC |
| 013 | EEQERKKQDQYNNDWEQKRYQREGRGQGHQP | -47.09 | 35.5 | HHHHHHHHHHHHHHHHHHHHHHHHCCCCCCC |
| 740 | EEQQRKQEQKKRNEEEHEEYQRKGRGQGHQP | -47.06 | 35.5 | HHHHHHHHHHHHHHHHHHHHHHHHCCCCCCC |
| 956 | EQEQRIQEDDRRNRKEQERYQQQGKGQGHNN | -47.05 | 25.8 | HHHHHHHHHHHHHHHHHHHHHHHCCCCCCCC |
| 415 | EEQQRKQEDQYRNEWEQERYQQQGKGQGHNN | -47.05 | 35.5 | HHHHHHHHHHHHHHHHHHHHHHHCCCCCCCC |
| 135 | EEEERKQRDDYKNRYEDEEYQRKGRGKGHQP | -47.04 | 35.5 | HHHHHHHHHHHHHHHHHHHHHHHHCCCCCCC |
| 531 | EQEERKKQEQYSNDSEDRRYQRYGRGKGHQP | -47.04 | 29.0 | HHHHHHHHHHHCCHHHHHHHHHHHCCCCCCC |
| 060 | EQEKRKQEDQRKNREEQERYQREGKGQGHNH | -47.01 | 25.8 | HHHHHHHHHHHHHHHHHHHHHHHHCCCCCCC |
| 865 | EEEKRKQEDDRKNRAEDQRYQKEGRGKGHQP | -47.00 | 38.7 | HHHHHHHHHHHHHHHHHHHHHHHHCCCCCCC |
| 009 | EEEQRKQEQQRRNEKEDQSYQRHGRGKGHQP | -47.00 | 35.5 | HHHHHHHHHHHHHHHHHHHHHHHCCCCCCCC |
| 276 | SNEERLQRDQESNEKEDEEYQRRGQGKGNQN | -47.00 | 32.3 | CHHHHHHHHHHHHHHHHHHHHHHHCCCCCCC |
| 083 | QEEEKKKQDQWRNDYEQEEYQRKGRGQGHQP | -47.00 | 29.0 | CHHHHHHHHHHHHHHHHHHHHHHHCCCCCCC |
| 692 | QREEEIQQQQRKNEKEQKRYQREGKGQGHNH | -46.99 | 22.6 | CHHHHHHHHHHHHHHHHHHHHHHHCCCCCCC |
| 670 | EQEERHQREQYSNDWEDEEYQRKGRGKGHQP | -46.98 | 29.0 | CHHHHHHHHHHCCCHHHHHHHHHHCCCCCCC |
| 578 | QEEQNKQQQQYSNEYEDEEYQRKGRGKGHQP | -46.97 | 32.3 | HHHHHHHHHHHHHHHHHHHHHHHHCCCCCCC |
| 334 | EEEERKQRQKYSNEFEHKYYQQKGQGQGHQP | -46.97 | 32.3 | CHHHHHHHHHHHHHHHHHHHHHHHCCCCCCC |
| 105 | EEEERKRQDQEKNDREHQYYQREGQGQGHQS | -46.97 | 29.0 | HHHHHHHHHHHHHHHHHHHHHHHHCCCCCCC |
| 482 | QEEEKKRQDQEKNDKEHKRYQKEGKGQGHNN | -46.97 | 29.0 | CHHHHHHHHHHHHHHHHHHHHHHCCCCCCCC |
| 163 | EQEKRHQEEQRSNDKEHEEYQRRGRGRGHQP | -46.95 | 22.6 | HHHHHHHHHHHHHHHHHHHHHHHHCCCCCCC |
| 065 | EEEKRKQEQQRSNEKEDEEYQRRGQGKGHQS | -46.94 | 35.5 | HHHHHHHHHHHHHHHHHHHHHHHHHCCCCCC |
| 801 | EEEERKKQDKQKNEKEHEEYQRYGRGQGHQP | -46.93 | 35.5 | HHHHHHHHHHHHHHHHHHHHHHHHHCCCCCC |
| 798 | EEEERKKQDKWKNEWEHKYYQKKGQGQGHQP | -46.93 | 35.5 | HHHHHHHHHHHHHHHHHHHHHHHHCCCCCCC |
| 325 | EREERYQQDQRKNDKEQEEYQRRGRGQGHQP | -46.93 | 25.8 | CHHHHHHHHHHHHHHHHHHHHHHHHCCCCCC |
| 422 | EEEERKQRQKKKNEEEHKRYQRYGRGQGHQP | -46.93 | 32.3 | HHHHHHHHHHHHHHHHHHHHHHHHCCCCCCC |
| 287 | EREERYRQDKENNEREDKRYQREGRGKGHNP | -46.90 | 41.9 | CHHHHHHHHHHHHHHHHHHHHHHCCCCCCCC |
| 996 | EEEQRKEEDMKKNRQEQQRYQREGRGQGHQP | -46.90 | 29.0 | HHHHHHHHHHHHHHHHHHHHHHHHCCCCCCC |

| | | | | |
|---|---|---|---|---|
| 927 | EEEQRKQKQQYDNDWEHEEYQRKGRGQGHNP | -46.89 | 25.8 | HHHHHHHHHHHHHHHHHHHHHHHCCCCCCCC |
| 258 | EEEQKKYEDQRKNEEEHEEFQRKGRGQGKQN | -46.89 | 32.3 | CHHHHHHHHHHCHHHHHHHHHHHHHCCCCCC |
| 061 | QEEKKKQEDQRSNDKEHEEYQRYGRGQGHQP | -46.82 | 29.0 | HHHHHHHHHHHHHHHHHHHHHHHHHCCCCCC |
| 623 | EQERRKQEEKRSNEEEDKRYQKQGRGKGHQP | -46.77 | 35.5 | HHHHHHHHHHHHHHHHHHHHHHHCCCCCCCC |
| 708 | EEERKKQEDKRKNEKEHEEYQRYGRGQGHQP | -46.76 | 38.7 | HHHHHHHHHHHHHHHHHHHHHHHHHCCCCCC |
| 489 | EEERRKQEDKYRNEWEHKKYQREGWGQGHYP | -46.76 | 35.5 | HHHHHHHHHHHHHHHHHHHHHHHHCCCCCCC |
| 269 | EEEEKRKRDQYKNESEHKRYQRYGKGQGHNH | -46.76 | 35.5 | HHHHHHHHHHHCHHHHHHHHHHHHCCCCCCC |
| 114 | EEEEKKQQDQRKNDNEQRYYQEYGQGQGNQN | -46.75 | 32.3 | CHHHHHHHHHHHHHHHHHHHHHHHHCCCCCC |
| 809 | EEEERKQRDQEKNDKEHEEYQRRGKGQGHNN | -46.75 | 29.0 | HHHHHHHHHHHHHHHHHHHHHHHCCCCCCCC |
| 078 | EQQRRKQEDKERNEKEHKRYQREGKGQGHNH | -46.72 | 35.5 | HHHHHHHHHHHHHHHHHHHHHHHHCCCCCCC |
| 917 | EEEQKKQRDQKKNEEEQERYQRYGTGQGHQP | -46.72 | 35.5 | HHHHHHHHHHHHHHHHHHHHHHHCCCCCCCC |
| 543 | EEEQRKQEDQWRNEWEQEYYQRKGQGQGHQP | -46.72 | 32.3 | HHHHHHHHHHHHHHHHHHHHHHHHCCCCCCC |
| 236 | EQEQRKQKDQEKNEREQKRYQREGRGQGHQP | -46.72 | 29.0 | HHHHHHHHHHHHHHHHHHHHHHHHCCCCCCC |
| 350 | QDEERKKQDQDSNDREQKRYQREGRGQGHQP | -46.70 | 22.6 | CHHHHHHHHHHHHHHHHHHHHHHHCCCCCCC |
| 116 | DNERRKQHEKESNEKEDKRYQRQGRGKGHQP | -46.69 | 32.3 | CHHHHHHHHHHHHHHHHHHHHHHCCCCCCCC |
| 498 | EQEQRKEEEQKKNEKEDEEYQRRGKGKGTNN | -46.69 | 32.3 | HHHHHHHHHHHHHHHHHHHHHHHHCCCCCCC |
| 628 | EEEQRKQHQQQSNDKEHEEYQRRGRGQGHQP | -46.69 | 25.8 | HHHHHHHHHHHHHHHHHHHHHHHHHCCCCCC |
| 675 | EEEERKRRDAEKNRREDQRYQREGKGKGHNH | -46.69 | 35.5 | HHHHHHHHHHHHHHHHHHHHHHHCCCCCCCC |
| 676 | EDEERRNQQQWKNDYEDEEYQRKGRGKGHNP | -46.68 | 29.0 | CHHHHHHHHHHHHHHHHHHHHHHCCCCCCCC |
| 631 | EQEERKNQEQWKNEWEDEYYQQYGQGKGHQP | -46.67 | 32.3 | HHHHHHHHHHHHHHHHHHHHHHHHHCCCCCC |
| 301 | EEQERKKQDQQSNDKEQKEYQKQGRGQGHQP | -46.66 | 32.3 | HHHHHHHHHHHHHHHHHHHHHHHHCCCCCCC |
| 145 | EREQRYKEDQRKNDKEDREYQKQGRGKGHQP | -46.66 | 32.3 | CHHHHHHHHHHHCHHHHHHHHHHHCCCCCCC |
| 737 | EEEERKRQDQEKNEKEHKRYQREGRGQGHQP | -46.64 | 32.3 | HHHHHHHHHHHHHHHHHHHHHHHHCCCCCCC |
| 664 | EEQERKNREQYSNDWEDKRYQREGRGKGHQP | -46.64 | 35.5 | HHHHHHHHHHHHHHHHHHHHHHHCCCCCCCC |
| 698 | EQEQRKQEDQRKNEKEQKRYQREGKGQGHNN | -46.60 | 29.0 | HHHHHHHHHHHHHHHHHHHHHHHCCCCCCCC |
| 495 | EEEKRKEEDQKENEREHEEYQRKGRGQGHQP | -46.57 | 32.3 | HHHHHHHHHHHHHHHHHHHHHHHHCCCCCCC |
| 881 | DRQRQYHEDQNRNEKEDEEYQRRGQGKGHQP | -46.57 | 35.5 | CHHHHHHHHHHCCHHHHHHHHHHHHCCCCCC |
| 091 | EQEKRHQEEQRKNDKEHEEKQRRGQGCGKQS | -46.57 | 19.4 | HHHHHHHHHHHHHHHHHHHHHHHCCCCCCCC |
| 386 | QEERKKQEDQERNQKEHDRYQREGRGQGHQP | -46.56 | 29.0 | HHHHHHHHHHHHHHHHHHHHHHHCCCCCCCC |
| 393 | EEEEKKRQDQDNNDREHEEYQRRGRGQGHQP | -46.54 | 35.5 | CHHHHHHHHHCCCHHHHHHHHHHHHCCCCCC |
| 117 | EEEEKKRRDQESNDKEHQRYQREGRGQGHQP | -46.54 | 32.3 | CHHHHHHHHHHHHHHHHHHHHHHHCCCCCCC |
| 302 | EQEERKREDQKRNEEEQQRYQRYGRGQGHQP | -46.54 | 29.0 | HHHHHHHHHHHHHHHHHHHHHHHHHCCCCCC |
| 288 | EEEERKKKQQYDNEYEDEEYQRKGRGKGHQP | -46.52 | 35.5 | CHHHHHHHHHHHHHHHHHHHHHHHCCCCCCC |
| 058 | EEEERKKRDQYKNDSEHERYRQQGQGQGHQS | -46.52 | 25.8 | HHHHHHHHHHCCCHHHHHHHHHHHCCCCCCC |
| 799 | EEEERKQQDQRKNENEQKRYQREGKGQGHNH | -46.50 | 32.3 | HHHHHHHHHHHHHHHHHHHHHHHHCCCCCCC |
| 584 | QEQQKKQEDARRNRFEQQYYQQKGQGQGHQP | -46.50 | 32.3 | HHHHHHHHHHHHHHHHHHHHHHHHCCCCCCC |
| 601 | QEEEKKKQDQEKNRKEQEEYQRRGRGQGHQP | -46.49 | 29.0 | CHHHHHHHHHHHHHHHHHHHHHHHHCCCCCC |
| 098 | QEEQKKKEDQRKNDAEHRRFQREGQGQGKQS | -46.48 | 29.0 | HHHHHHHHHHHHHHHHHHHHHHHHCCCCCCC |

| | | | | |
|---|---|---|---|---|
| 326 | EEEEKKKQDDWRNRWEDQYYQKKGQGKGHQP | -46.48 | 38.7 | CHHHHHHHHHHHHHHHHHHHHHHHCCCCCCC |
| 687 | EQQQRHQEEKRRNEKEDKYYQKEGQGKGNQN | -46.47 | 38.7 | HHHHHHHHHHHHHHHHHHHHHHHCCCCCCCC |
| 052 | EQQQRHQQEQQSNEREDEEYQRRGKGKGHNN | -46.47 | 35.5 | HHHHHHHHHHHHHHHHHHHHHHHCCCCCCCC |
| 974 | EEEERKLQDQQKNDKEQRRFQRYGKGQGKNN | -46.42 | 25.8 | CHHHHHHHHHHHHHHHHHHHHHHHHCCCCCC |
| 797 | EEEEKKQQQQREDEYRDEEYQRKGRGKGHNP | -46.40 | 35.5 | HHHHHHHHHHHHHHHHHHHHHHHCCCCCCCC |
| 574 | EQQQRHQEEQRQNEKEDKRYQKEGRGKGHQP | -46.40 | 35.5 | HHHHHHHHHHHHHHHHHHHHHHHCCCCCCCC |
| 040 | QQEQNHQEEQRRNEKEDEEYQRRGRGKGHQP | -46.40 | 29.0 | CHHHHHHHHHHHHHHHHHHHHHHHCCCCCCC |
| 005 | ENEERKNQEKQKHEKEQKRYQREGRGQGHQP | -46.39 | 32.3 | CHHHHHHHHHHHHHHHHHHHHHHHCCCCCCC |
| 255 | EEEQRIQEQQRENEREQEEYQQKGWGQGHYP | -46.39 | 29.0 | HHHHHHHHHHHHHHHHHHHHHHHCCCCCCCC |
| 279 | EQEERKWRDQKKNEEEHERYQQQGRGQGHQP | -46.34 | 29.0 | CHHHHHHHHHHHHHHHHHHHHHHHCCCCCCC |
| 646 | QEEEKKRRDQESNEREHKYFQKKGQGQGKNN | -46.33 | 29.0 | CHHHHHHHHHHHHHHHHHHHHHHHCCCCCCC |
| 358 | EEERRKQEDKKRNEEEDKRYQKQGRGKGHQP | -46.33 | 41.9 | HHHHHHHHHHHHHHHHHHHHHHHCCCCCCCC |
| 722 | EEQERKKQDQYNNDWEQRRYQREGDGQGHQP | -46.33 | 35.5 | HHHHHHHHHHHHHHHHHHHHHHHCCCCCCCC |
| 014 | EQEERKKQEQWKNDWEHEYYQKYGQGQGHQP | -46.33 | 22.6 | HHHHHHHHHHHHHHHHHHHHHHHHHCCCCCC |
| 018 | EQERRKQEQKYKQEWEDEEYQRKGRGKGHQP | -46.28 | 35.5 | HHHHHHHHHHHHHHHHHHHHHHHHCCCCCCC |
| 833 | EEQERKKQDQWKNDYEQRSYQREGRGQGHQP | -46.27 | 32.3 | HHHHHHHHHHHHHHHHHHHHHHHCCCCCCCC |
| 784 | EQEQRHQEEQRKNEKEDKRYQREGRGKGHQP | -46.27 | 32.3 | HHHHHHHHHHHHHHHHHHHHHHHHCCCCCCC |
| 702 | EQEQRHQEEQRKNEKEDKRYQRYGRGKGHQP | -46.27 | 32.3 | HHHHHHHHHHHHHHHHHHHHHHHHCCCCCCC |
| 436 | DNQRRRNEEQWRNDWEDQYYQKKGQGKGHNH | -46.27 | 29.0 | CHHHHHHHHHHHHHHHHHHHHHHHCCCCCCC |
| 158 | EEEEKKKRDQYKNESEQERYQRYGRGQGHQP | -46.27 | 35.5 | CHHHHHHHHHHCHHHHHHHHHHHHHCCCCCC |
| 649 | EEEERKKEDQRKNEEEQQRYQRYGKGQGHNN | -46.27 | 32.3 | HHHHHHHHHHHHHHHHHHHHHHHHCCCCCCC |
| 746 | EEEERKKEDQYRNEWEQEEYQRKGRGQGHQP | -46.27 | 32.3 | HHHHHHHHHHHHHHHHHHHHHHHHCCCCCCC |
| 412 | EEEERKKQDQYNNEWEQKRYQREGQGQGHQS | -46.27 | 35.5 | HHHHHHHHHHHHHHHHHHHHHHHHCCCCCCC |
| 222 | EQEERHQREQKSNEEEDKRFQREGDGKGKQN | -46.26 | 29.0 | CHHHHHHHHHHHHHHHHHHHHHHCCCCCCCC |
| 372 | EQEKRHQEEQRSQEQEDEEYQRRGQGKGHQS | -46.24 | 32.3 | HHHHHHHHHHHHHHHHHHHHHHHHHCCCCCC |
| 496 | EQEKRHQEEQRSNDEEDRRYQRYGQGKGHYS | -46.22 | 29.0 | HHHHHHHHHHHHCHHHHHHHHHHHCCCCCCC |
| 711 | ENEERKQREQESQRKEDERYQREGRGKGHQP | -46.20 | 29.0 | CHHHHHHHHHHHHHHHHHHHHHHHCCCCCCC |
| 196 | EEEERKKQDQYNNEWEQEEFQRKGRGQGKQN | -46.20 | 32.3 | HHHHHHHHHHHHHHHHHHHHHHHHHCCCCCC |
| 508 | EQEERKRQDQEKNEKEHKRYQREGRGQGHQP | -46.19 | 29.0 | HHHHHHHHHHHHHHHHHHHHHHHHCCCCCCC |
| 036 | EEEQRKEEDKKKNEREQKEYQQRGRGQGHQP | -46.19 | 35.5 | CHHHHHHHHHHHHHHHHHHHHHHHCCCCCCC |
| 311 | EEEERKKQEQWKNEWEDQYYQKQGQGKGHQP | -46.18 | 35.5 | HHHHHHHHHHHHHHHHHHHHHHHHCCCCCCC |
| 932 | EQEERKKQEQWKNEWEDQFYQKQGRGKGHQP | -46.18 | 35.5 | HHHHHHHHHHHHHHHHHHHHHHHHCCCCCCC |
| 390 | EQEERHKEEQYRNDYEDEEYQRKGRGKGTQN | -46.17 | 29.0 | HHHHHHHHHHHHHHHHHHHHHHHHCCCCCCC |
| 121 | EEEERKQQDARKNRYEQEEYQRKGRGQGHQP | -46.17 | 29.0 | HHHHHHHHHHHHHHHHHHHHHHHHCCCCCCC |
| 576 | EEEQRKRYDQEKNRKEHENYQREGRGQGHQP | -46.17 | 29.0 | HHHHHHHHHHHHHHHHHHHHHHHCCCCCCCC |
| 081 | EEERRKQEDQQKNEREHEEYQRKGRGQGHNP | -46.16 | 32.3 | HHHHHHHHHHHHHHHHHHHHHHHCCCCCCCC |
| 087 | EEEERKQRQQYSNDWEHERYQREGRGQGHQP | -46.16 | 25.8 | HHHHHHHHHHHHHHHHHHHHHHHHCCCCCCC |
| 890 | EQQQRHQEEQRRNDKEDERYQQQGWGKGHYP | -46.15 | 32.3 | HHHHHHHHHHHHHHHHHHHHHHHCCCCCCCC |

| | | | | |
|---|---|---|---|---|
| 360 | ENEERKRQEQEKNEKEDEEYQRRGWGKGHYP | -46.15 | 32.3 | CHHHHHHHHHHHHHHHHHHHHHHCCCCCCCC |
| 284 | EQEKRHQEEQRSNEEEDKRYQKEGRGKGHQP | -46.14 | 32.3 | HHHHHHHHHHHHHHHHHHHHHHHHCCCCCCC |
| 136 | QQEKNHQEEQRSNEAEDKQYQQRGRGKGHQP | -46.14 | 32.3 | CHHHHHHHHHHHHHHHHHHHHHHHCCCCCCC |
| 401 | EQEERHQQEQRKNRWEDEYYQQKGQGKGHQP | -46.11 | 29.0 | CHHHHHHHHHHHHHHHHHHHHHHHCCCCCCC |
| 181 | DEERKKQEEKERNEKEDKRYQQQGWGKGHYP | -46.10 | 38.7 | HHHHHHHHHHHHHHHHHHHHHHHHCCCCCCC |
| 834 | EEEERKKQDQEKNDKEQRRYQREGRGQGHQP | -46.05 | 29.0 | HHHHHHHHHHHHHHHHHHHHHHHHCCCCCCC |
| 103 | QEERKKQEDQWRNEWEHEYYQRKGQGQGHQP | -46.05 | 32.3 | HHHHHHHHHHHHHHHHHHHHHHHHCCCCCCC |
| 294 | EQEERKKQEQFSNDFEHKYYQQKGQGQGHQP | -46.05 | 25.8 | HHHHHHHHHHHHHHHHHHHHHHHHCCCCCCC |
| 500 | EHEERRQQEQRKNEYEDEEYQRKGRGKGHQP | -46.04 | 32.3 | CHHHHHHHHHHHHHHHHHHHHHHHCCCCCCC |
| 274 | EQEERKRQDQERNDKEQEEYQRKGRGQGHQP | -46.03 | 25.8 | HHHHHHHHHHHHHHHHHHHHHHHHCCCCCCC |
| 774 | EQEQRHNEEQRSNREEDERYQREGKGKGHNN | -46.01 | 29.0 | HHHHHHHHHHHHHHHHHHHHHHHCCCCCCCC |
| 418 | EEERRKQEEQERNEKEDKRYQRYGRGKGHQP | -46.00 | 35.5 | HHHHHHHHHHHHHHHHHHHHHHHHCCCCCCC |
| 815 | EEEQRKKEDKRKNEEEQKKYQEQGRGQGHQP | -45.99 | 35.5 | HHHHHHHHHHHHHHHHHHHHHHHHCCCCCCC |
| 748 | EQERRKQEEQRSNRKEDEEYQRRGRGKGHQP | -45.98 | 29.0 | HHHHHHHHHHHHHHHHHHHHHHHHCCCCCCC |
| 220 | EEEERKQRDQEKNEKEHERYQKQGRGQGHQP | -45.97 | 32.3 | HHHHHHHHHHHHHHHHHHHHHHHCCCCCCCC |
| 891 | QEEEKKRQDKEKNEKEQKRYQRYGRGQGHQP | -45.97 | 35.5 | CHHHHHHHHHHHHHHHHHHHHHHHHCCCCCC |
| 723 | ERQERYQRDQESNEKEDKRYQRYGRGKGHQP | -45.96 | 38.7 | CHHHHHHHHHHHHHHHHHHHHHHHCCCCCCC |
| 769 | EQEQRKQEEQRRNDKEQRRYQREGKGQGHNN | -45.96 | 22.6 | HHHHHHHHHHHHHHHHHHHHHHHCCCCCCCC |
| 521 | EQEERIQRDDYKHRWWHEEYQRKGRGQGHQP | -45.95 | 25.8 | CHHHHHHHHHHHHHHHHHHHHHHCCCCCCCC |
| 023 | EQEKRHQEEQRKNEEEDERYQRQGQGKGHQS | -45.95 | 32.3 | HHHHHHHHHHHHHHHHHHHHHHHHCCCCCCC |
| 407 | EEEQKKRQDQENNEKEHEEYQRRGRGQGHQP | -45.94 | 38.7 | HHHHHHHHHHHHHHHHHHHHHHHHCCCCCCC |
| 607 | EEEQRKHEQQNRNEKEDKRYQRYGQGKGHQS | -45.91 | 35.5 | HHHHHHHHHHHHHHHHHHHHHHHHCCCCCCC |
| 408 | EEEQKKEEDQKSNEQEHERYQREGQGQGHQP | -45.89 | 35.5 | CHHHHHHHHHHHHHHHHHHHHHHHCCCCCCC |
| 226 | EEERKKQRDQEKNDKEQEEYQRRGQGQGHNP | -45.88 | 32.3 | HHHHHHHHHHHHHHHHHHHHHHHHHCCCCCC |
| 184 | EQEERHKKEQEKNDKEDEEYQRYGRGKGHQP | -45.85 | 29.0 | CHHHHHHHHHHHCHHHHHHHHHHHHCCCCCC |
| 905 | EEERRKQEDKERNEKEQKQYQQRGKGQGHNN | -45.82 | 35.5 | HHHHHHHHHHHHHHHHHHHHHHHCCCCCCCC |
| 180 | ERQQRYQEDQYRNEYEQEEYQRKGRGQGHQP | -45.82 | 32.3 | HHHHHHHHHHHHHHHHHHHHHHHHCCCCCCC |
| 791 | EEEERKRQDQEKNDREQKRYQREGRGQGHQP | -45.76 | 29.0 | HHHHHHHHHHHHHHHHHHHHHHHHCCCCCCC |
| 886 | EEEERKRQDQESNEREHKRYQREGRGQGHQP | -45.76 | 32.3 | HHHHHHHHHHHHHHHHHHHHHHHHCCCCCCC |
| 964 | EEERRKQEDKERNEKEQKRYQREGRGQGHQP | -45.74 | 35.5 | HHHHHHHHHHHHHHHHHHHHHHHHCCCCCCC |
| 034 | EQEKRHQEEQRSNEKEDEEYQRRGKGKGHNH | -45.72 | 32.3 | HHHHHHHHHHHHHHHHHHHHHHHHCCCCCCC |
| 095 | QQQQNHQEEKRRNEKEHKRYQRYGKGQGHNN | -45.68 | 29.0 | CHHHHHHHHHHHHHHHHHHHHHHCCCCCCCC |
| 345 | EQERRLQEDQYRNEWEQEEYQRKGRGQGHQP | -45.66 | 29.0 | HHHHHHHHHHHHHHHHHHHHHHHHCCCCCCC |
| 800 | EEEQRKQEDDRKNRKEHERYQREGRGQGHQP | -45.64 | 29.0 | HHHHHHHHHHHHHHHHHHHHHHHHCCCCCCC |
| 632 | EQEERHQQEQRKNDKEDREYQKQGRGKGHQP | -45.63 | 29.0 | CHHHHHHHHHHHCHHHHHHHHHHHCCCCCCC |
| 241 | EQEERHKEEQYRNDWEDERYQREGQGKGHQS | -45.61 | 29.0 | HHHHHHHHHHHHHHHHHHHHHHHHCCCCCCC |
| 752 | EQEKRHQEEQRKNDEEHEEFQRKGRGQGKQN | -45.58 | 19.4 | HHHHHHHHHHHCHHHHHHHHHHHHHCCCCCC |
| 025 | EQEERHQREQWKNRWEDEFYQKQGRGKGHQP | -45.57 | 32.3 | CHHHHHHHHHHHHHHHHHHHHHHHCCCCCCC |

| | | | | |
|---|---|---|---|---|
| 941 | EQEQRHQQEQDSNRREDEEYQRRGQGKGHQP | -45.54 | 29.0 | HHHHHHHHHHHHHHHHHHHHHHHHCCCCCCC |
| 246 | EEEERKKQQQYKNEWEDKRYQREGDGKGHNP | -45.54 | 35.5 | HHHHHHHHHHHHHHHHHHHHHHCCCCCCCCC |
| 678 | EEEQRKQEDQQKNEREDEEYQRRGQGKGHQS | -45.50 | 38.7 | HHHHHHHHHHHHHHHHHHHHHHHHHCCCCCC |
| 602 | EQEERKQQQKRKNEKEQKRYQREGRGQGHQP | -45.49 | 29.0 | HHHHHHHHHHHHHHHHHHHHHHHHCCCCCCC |
| 533 | QEERKKQHDQEKNQKEQDRYQREGRGQGHQP | -45.41 | 29.0 | HHHHHHHHHHHHHHHHHHHHHHHHCCCCCCC |
| 544 | QEQEKKKEDQNRNRKEQERYQREGRGQGHQP | -45.39 | 32.3 | CHHHHHHHHHHHHHHHHHHHHHHHCCCCCCC |
| 671 | EQEQRHQEEQRKNDEEHRQYQERGRGQGHQP | -45.38 | 22.6 | HHHHHHHHHHHHHHHHHHHHHHHHCCCCCCC |
| 336 | EQEERHQQEQRSNEKEDEEYQRRGQGKGNQN | -45.33 | 32.3 | CHHHHHHHHHHHHHHHHHHHHHHHCCCCCCC |
| 715 | DEERKKQEDQERNDKEQKRYQREGQGQGHQP | -45.33 | 29.0 | CHHHHHHHHHHHHHHHHHHHHHHHCCCCCCC |
| 816 | EEEQRKQHDQEKNDKEQEEYQKRGRGQGHQP | -45.33 | 29.0 | HHHHHHHHHHHHHHHHHHHHHHHHCCCCCCC |
| 182 | EEERRRQEEQYRNEWEDERYQRQGKGKGHNN | -45.32 | 35.5 | HHHHHHHHHHHHHHHHHHHHHHHCCCCCCCC |
| 090 | EQQERKRQDQESNRREHERYQKQGRGQGHQP | -45.30 | 29.0 | HHHHHHHHHHHHHHHHHHHHHHHCCCCCCCC |
| 459 | EQEQRHQQEKRSNEKEHEEYQRRGRGQGHQP | -45.29 | 29.0 | HHHHHHHHHHHHHHHHHHHHHHHHHCCCCCC |
| 314 | EEEERKRQDQEKNEKEQKRYQREGRGQGHQP | -45.25 | 32.3 | HHHHHHHHHHHHHHHHHHHHHHHHCCCCCCC |
| 214 | EEEQRKQEQQRRNEKEQERYQRQGWGQGHYP | -45.25 | 29.0 | HHHHHHHHHHHHHHHHHHHHHHHCCCCCCCC |
| 129 | QREKEKEEQQRSNEKEDEEYQRRGRGKGHQP | -45.24 | 29.0 | CHHHHHHHHHHHHHHHHHHHHHHHCCCCCCC |
| 063 | EEEQRKQEQHRKNRKEDEEYQRRGRGKGHQP | -45.23 | 32.3 | HHHHHHHHHHHHHHHHHHHHHHHHCCCCCCC |
| 225 | EEEKRKEEQQRKNEEEDQRYQRYGRGKGHQP | -45.21 | 35.5 | HHHHHHHHHHHHHHHHHHHHHHHHCCCCCCC |
| 234 | EQEQRKQEEQKRNEEEQKRFQREGKGQGKNH | -45.20 | 22.6 | HHHHHHHHHHHHHHHHHHHHHHHHCCCCCCC |
| 064 | EEEERKRQQQESNEREDRRYQREGRGKGHQP | -45.19 | 35.5 | HHHHHHHHHHHHHHHHHHHHHHHHCCCCCCC |
| 650 | EQEERHKQEQYKNDYEDREYQRKGRGKGHQP | -45.14 | 29.0 | HHHHHHHHHHHHHHHHHHHHHHHHCCCCCCC |
| 178 | EEEERKRQDDEKQRKEDEEYQRRGRGKGHQP | -45.13 | 35.5 | HHHHHHHHHHHHHHHHHHHHHHHHCCCCCCC |
| 660 | EQEQRHQEEQRRNDKEHQRYQREGRGQGHQP | -45.12 | 22.6 | HHHHHHHHHHHHHHHHHHHHHHHHCCCCCCC |
| 600 | EEEERKRQDQENNDREQERYQRQGRGQGHQP | -45.12 | 32.3 | HHHHHHHHHHHHHHHHHHHHHHHHCCCCCCC |
| 597 | EREQRYEEDQKKNEQEHSRYQQQGKGQGHNH | -45.12 | 29.0 | HHHHHHHHHHHHHHHHHHHHHHHCCCCCCCC |
| 270 | EEERKKQEDQERNEKEQQRYQREGKGQGHNH | -45.11 | 35.5 | HHHHHHHHHHHHHHHHHHHHHHHHCCCCCCC |
| 054 | EEERKKQEDQYRNEYEQEEYQRKGRGQGHQP | -45.11 | 35.5 | HHHHHHHHHHHHHHHHHHHHHHHHCCCCCCC |
| 171 | EEEKRKQEDHRKNREEQEKYQREGKGQGHNH | -45.08 | 29.0 | HHHHHHHHHHHHHHHHHHHHHHHHCCCCCCC |
| 695 | EEEERKRRDQEKNEKEQEEYQRRGRGQGHQP | -45.07 | 32.3 | HHHHHHHHHHHHHHHHHHHHHHHHHCCCCCC |
| 561 | EEEERKRRDQESNEKEQKEYQKQGRGQGHQP | -45.07 | 32.3 | HHHHHHHHHHHHHHHHHHHHHHHHCCCCCCC |
| 610 | EQEERHKQEQWKNEWEDKYYQKKGQGKGHQS | -45.07 | 32.3 | HHHHHHHHHHHHHHHHHHHHHHHHCCCCCCC |
| 785 | QQEQNKEEDDKRHRYEDEEYQRKGRGKGHQP | -45.03 | 32.3 | CHHHHHHHHHHHHHHHHHHHHHHHCCCCCCC |
| 150 | DEERKKQHDQESNEKEQKRYQRYGQGQGHQS | -45.03 | 32.3 | HHHHHHHCHHHHHHHHHHHHHHHHHCCCCCC |
| 278 | EEEERKKQDDRKNREEHQRYQREGWGRGHYP | -45.02 | 29.0 | HHHHHHHHHHHHHHHHHHHHHHHHCCCCCCC |
| 608 | EQEERHKEEQRSNEKEDEEYQRRGRGKGHQP | -45.02 | 32.3 | HHHHHHHHHHHHHHHHHHHHHHHHCCCCCCC |
| 204 | EQEERHKQEQYNNEYEDEEYQRKGRGKGHQP | -45.02 | 35.5 | HHHHHHHHHHHHHHHHHHHHHHHHCCCCCCC |
| 700 | EEEERKQQDHRKNRYEQQRYQREGKGQGHSN | -45.01 | 29.0 | HHHHHHHHHHHHHHHHHHHHHHHCCCCCCCC |
| 293 | EEEQRKEEDQKKNEKEQEEYQRRGQGQGHQP | -45.00 | 32.3 | HHHHHHHHHHHHHHHHHHHHHHHHHCCCCCC |

| | | | | |
|---|---|---|---|---|
| 346 | DNERRKQHEQESNDKEHKRYQREGKGQGHNH | -44.98 | 19.4 | CHHHHHHHHHHHHHHHHHHHHHHCCCCCCCC |
| 331 | EEQRRKQEDQYRNDWEQHRYQNEGRGQGHQP | -44.97 | 32.3 | HHHHHHHHHHHHHHHHHHHHHCCCCCCCCCC |
| 694 | EQEKRHQEEQREDEKRDEEYQRRGQGKGHQS | -44.96 | 29.0 | HHHHHHHHHHHHHHHHHHHHHHHHHCCCCCC |
| 441 | EQQERHNQEQQKNDKEHQRYQREGKGQGHNH | -44.96 | 25.8 | HHHHHHHHHHHHHHHHHHHHHHHHCCCCCCC |
| 473 | QEEKKKEEDQRKNEEEQKRYQREGRGQGHQP | -44.95 | 32.3 | HHHHHHHHHHHHHHHHHHHHHHHHCCCCCCC |
| 146 | EQEQRKQEEQWRNEWEQQYYQKQGQGQGHQP | -44.95 | 25.8 | HHHHHHHHHHHHHHHHHHHHHHHHCCCCCCC |
| 176 | EEERRKQEDQERQEKEQQEYQQRGQGQGHQS | -44.94 | 32.3 | HHHHHHHHHHHHHHHHHHHHHHHHHCCCCCC |
| 433 | EEQQRKQKDHQDNRREQEEYQRRGKGQGHNN | -44.93 | 32.3 | HHHHHHHHHHHHHHHHHHHHHHHCCCCCCCC |
| 596 | QQEQNHEEEKKSNEKEDEEYQRRGKGKGHNH | -44.92 | 32.3 | CHHHHHHHHHHHHHHHHHHHHHHHCCCCCCC |
| 312 | EEEERRKEEQRRNDEEDRRYQRYGQGKGHQS | -44.88 | 32.3 | HHHHHHHHHHHHHHHHHHHHHHHHHCCCCCC |
| 837 | EEEERKQQDDRENRREQQEYQREGKGQGHNH | -44.87 | 29.0 | HHHHHHHHHHHHHHHHHHHHHHHHCCCCCCC |
| 603 | EEEERKKEQKYRNEWEQKRYQREGRGQGHQP | -44.87 | 32.3 | HHHHHHHHHHHHHHHHHHHHHHHHCCCCCCC |
| 203 | EEEEKKKQDDEKNRKEDEQYQRYGRGKGHQP | -44.86 | 38.7 | CHHHHHHHHHHHHHHHHHHHHHHHHCCCCCC |
| 100 | EEEQKKKEDDRKNREEDQRYQREGQGKGHQS | -44.86 | 38.7 | HHHHHHHHHHHHHHHHHHHHHHHHCCCCCCC |
| 918 | EEEERKRQDQESNDKEQREYQKQGQGQGHQS | -44.83 | 29.0 | HHHHHHHHHHHHHHHHHHHHHHHHCCCCCCC |
| 093 | EEEERKKQQQYSNDWEQEEFQRKGRGQGKQN | -44.82 | 22.6 | HHHHHHHHHHHHHHHHHHHHHHHHHCCCCCC |
| 008 | EQEERHKQEQKSNEEEDKRYQRYGRGKGHQP | -44.82 | 32.3 | HHHHHHHHHHHHHHHHHHHHHHHHCCCCCCC |
| 684 | EEEEKKKQDQQKNEKEQEEYQRRGRGQGHQP | -44.81 | 35.5 | CHHHHHHHHHHHHHHHHHHHHHHHHCCCCCC |
| 003 | EEEERKQRQQESNEREQERYQKQGRGQGHQP | -44.80 | 29.0 | HHHHHHHHHHHHHHHHHHHHHHHCCCCCCCC |
| 329 | EQQERHKEEQWRNEYEDEEYQRKGRGKGHQP | -44.77 | 35.5 | CHHHHHHHHHHHHHHHHHHHHHHHCCCCCCC |
| 239 | QEEEKKQRDDEKHRKEDQRYQREGRGKGHQP | -44.73 | 38.7 | CHHHHHHHHHHHHHHHHHHHHHHHCCCCCCC |
| 509 | EQQERKREDQWRNEYEHEEYQRKGRGQGHQP | -44.73 | 32.3 | HHHHHHHHHHHHHHHHHHHHHHHHCCCCCCC |
| 238 | QQQQNHQEEQRRNEKEHQRYQREGRGQGHQP | -44.68 | 25.8 | CHHHHHHHHHHHHHHHHHHHHHHHCCCCCCC |
| 532 | EQEERHRQEKEKNEKEDKRYQREGKGKGHNH | -44.68 | 35.5 | HHHHHHHHHHHHHHHHHHHHHHHCCCCCCCC |
| 934 | QDEERKQRDQEKNEKEQEEYQRRGKGQGHSS | -44.64 | 25.8 | CHHHHHHHHHHHHHHHHHHHHHHHCCCCCCC |
| 994 | EEERRKQEDQERNEKEQEEYQRYGWGQGHFP | -44.63 | 35.5 | HHHHHHHHHHHHHHHHHHHHHHHCCCCCCCC |
| 471 | EQERRHQHEQESNDKEHEEYQRRGWGRGHYP | -44.63 | 22.6 | HHHHHHHHHHHHHHHHHHHHHHHCCCCCCCC |
| 923 | ENEERRKREQYSNDWEDEEYQRKGRGKGHQP | -44.59 | 29.0 | CHHHHHHHHHHCCHHHHHHHHHHHCCCCCCC |
| 703 | EQEERHRQEQEKNDKEHERYQRQGWGQGHYP | -44.59 | 22.6 | HHHHHHHHHHHHHHHHHHHHHHHHCCCCCCC |
| 456 | EQEERKKQEQYSNRYEQEEYQRKGRGQGHQP | -44.54 | 22.6 | HHHHHHHHHHHHHHHHHHHHHHHHCCCCCCC |
| 367 | EEEERKKRDQWKNEWEQKYYQQKGQGQGHQP | -44.44 | 32.3 | HHHHHHHHHHHHHHHHHHHHHHHHCCCCCCC |
| 438 | EEEEKKKQEKEKNEKEHKRYQREGRGQGHQP | -44.43 | 35.5 | HHHHHHHHHHHHHHHHHHHHHHHHCCCCCCC |
| 251 | EEEERKKEDDRKNREEHERYQQQGKGQGHNN | -44.39 | 29.0 | CHHHHHHHHHHHHHHHHHHHHHHCCCCCCCC |
| 244 | EQEERKKEDDWKNRWEHEYYQRKGQGQGHQP | -44.39 | 25.8 | HHHHHHHHHHHHHHHHHHHHHHHHCCCCCCC |
| 333 | QQEENHRQEQEKNRKEDERYQREGKGKGHNN | -44.37 | 25.8 | CHHHHHHHHHHHHHHHHHHHHHHCCCCCCCC |
| 540 | EEERRRQQEQQSNDKEDEEYQRYGRGKGHQP | -44.37 | 32.3 | HHHHHHHHHHHHCCHHHHHHHHHHHCCCCCC |
| 067 | EEEQKKQEQQKRNEEEQKRYQQYGKGQGHNN | -44.31 | 32.3 | HHHHHHHHHHHHHHHHHHHHHHHHCCCCCCC |
| 873 | EQEKRHNQEKYSNEWEQEEYQRQGRGQGHQP | -44.31 | 29.0 | HHHHHHHHHHHHHHHHHHHHHHHHCCCCCCC |

| | | | | |
|---|---|---|---|---|
| 446 | EEEERKQQDDRSNRKEQQEYQRSGKGQGHNN | -44.28 | 32.3 | HHHHHHHHHHHHHHHHHHHHHHHCCCCCCCC |
| 720 | EEEQRKEEQQKKNEQEDQRYQRYGRGKGNQN | -44.27 | 35.5 | HHHHHHHHHHHHHHHHHHHHHHHCCCCCCCC |
| 242 | EEEERKQQQQREDDWRQEYYQKYGQGQGHQP | -44.17 | 22.6 | HHHHHHHHHHHHHHHHHHHHHHHHHCCCCCC |
| 306 | EEERKKQRDQESNEKEQEKYQQEGKGQGHNH | -44.16 | 35.5 | HHHHHHHHHHHHHHHHHHHHHHHHCCCCCCC |
| 499 | EEEERKREQQERNDKEHEEYQRYGRGQGHQP | -44.13 | 25.8 | HHHHHHHHHHHHHHHHHHHHHHHHHCCCCCC |
| 958 | EQEERHKEEQWRNDYEHEEYQRKGRGQGHQP | -44.13 | 22.6 | HHHHHHHHHHHHHHHHHHHHHHHHCCCCCCC |
| 885 | QQEENHKEEQRSNDKEHEEYQRRGRGQGHQP | -44.13 | 19.4 | CHHHHHHHHHHHHHHHHHHHHHHHHCCCCCC |
| 667 | EEEERKKKQQEKDEKRDEEYQRRGRGKGHQP | -44.12 | 32.3 | CHHHHHHHHHHHHHHHHHHHHHHHCCCCCCC |
| 775 | EEEERKKQQVQKNRKEHEEYQRRGRGQGHQP | -44.12 | 25.8 | HHHHHHHHHHHHHHHHHHHHHHHHCCCCCCC |
| 449 | EQEERHRQEQEKNEKEHKRYQREGKGRGHNN | -44.08 | 25.8 | HHHHHHHHHHHHHHHHHHHHHHHCCCCCCCC |
| 965 | EQEERHKQEQIKNEKEDEEYQRRGRGKGHQP | -43.94 | 32.3 | HHHHHHHHHHHHHHHHHHHHHHHHCCCCCCC |
| 383 | EQEERHKQEQYKNEWEDKRYQREGRGKGHQP | -43.94 | 32.3 | HHHHHHHHHHHHHHHHHHHHHHHHCCCCCCC |
| 862 | QQQENHWREQESNDREDRRYQREGQGKGHQS | -43.94 | 29.0 | CHHHHHHHHHHHCCHHHHHHHHHHCCCCCCC |
| 861 | ENEERRKEEQKRNDKEHEEYQRRGRGQGHQP | -43.91 | 22.6 | CHHHHHHHHHHHHHHHHHHHHHHHHCCCCCC |
| 185 | EQEERHRREQESNRKEDERYQREGRGKGHQP | -43.89 | 29.0 | HHHHHHHHHHHHHHHHHHHHHHHHCCCCCCC |
| 943 | QQEQNKEEEQKEEEKRHEEKQRRGKGCGKNN | -43.88 | 16.1 | CHHHHHHHHHHHHHHHHHHHHHHCCCCCCCC |
| 589 | EEEERKKQEDQSNRREDEEYQRRGRGKGHQP | -43.87 | 32.3 | HHHHHHHHHHHHHHHHHHHHHHHHCCCCCCC |
| 260 | EEQEKKKRDQYKEEWRHEEYQRKGRGQGHQP | -43.86 | 35.5 | HHHHHHHHHHHHHHHHHHHHHHHCCCCCCCC |
| 697 | EEEERKQQQKRDQENRQKRYQKEGWGQGHYP | -43.82 | 29.0 | HHHHHHHHHHHHHHHHHHHHHHHCCCCCCCC |
| 619 | EQERRHQEEQRSNDEEDRRYQREGRGKGHQP | -43.78 | 29.0 | HHHHHHHHHHHHCHHHHHHHHHHHCCCCCCC |
| 952 | QQERNHQEEQRSNDEEDRRYQREGRGKGHQP | -43.78 | 25.8 | CHHHHHHHHHHHCHHHHHHHHHHHCCCCCCC |
| 156 | EQERRHQEEQERNRKEDERYQREGQGKGHQS | -43.74 | 29.0 | HHHHHHHHHHHHHHHHHHHHHHHHCCCCCCC |
| 754 | EQEERHKQEQYKFDWWHEEYQRKGRGQGHQP | -43.73 | 19.4 | CHHHHHHHHHHHHHHHHHHHHHHCCCCCCCC |
| 842 | EQEKRHQEEQRNNQEEQDRYQREGKGQGHSH | -43.73 | 25.8 | HHHHHHHHHHHHHHHHHHHHHHHCCCCCCCC |
| 490 | EQEQRKEEEQKKNEYEQEEYQRKGRGQGHQP | -43.71 | 25.8 | HHHHHHHHHHHHHHHHHHHHHHHHCCCCCCC |
| 027 | EQEQRHQEEKRRNEKEQEEYQRKGRGQGHQP | -43.69 | 29.0 | HHHHHHHHHHHHHHHHHHHHHHHHCCCCCCC |
| 654 | QQQENHQQEQQKNEKEQEEYQRRGKGQGHQH | -43.68 | 25.8 | CHHHHHHHHHHHHHHHHHHHHHHHHCCCCCC |
| 488 | QWERWLQELQERNDKEQQRYQKQGRGQGHQP | -43.62 | 19.4 | CHHHHHHHHHHHHHHHHHHHHHHHCCCCCCC |
| 618 | EQEQRHQEEQRKNEREQEEYQQRGRGQGHQP | -43.61 | 25.8 | HHHHHHHHHHHHHHHHHHHHHHHHHCCCCCC |
| 478 | EQERRHQEEQRRNEQEQKRYQREGRGQGHNP | -43.61 | 25.8 | HHHHHHHHHHHHHHHHHHHHHHHHCCCCCCC |
| 841 | NDEERKRQDDEKNRKEDERYQRQGRGKGNQN | -43.50 | 29.0 | CHHHHHHHHHHHHHHHHHHHHHHCCCCCCCC |
| 335 | EQQERHQREQWKNEWEQKYYQKEGQGQGHQS | -43.49 | 29.0 | CHHHHHHHHHHHHHHHHHHHHHHHCCCCCCC |
| 445 | EQEERHREEQYRNEWEDEEYQRKGRGKGHSN | -43.48 | 32.3 | HHHHHHHHHHHHHHHHHHHHHHHCCCCCCCC |
| 732 | EQEERHRQEQESNEREDKRYQREGRGKGHQP | -43.48 | 32.3 | HHHHHHHHHHHHHHHHHHHHHHHHCCCCCCC |
| 272 | EQEKREQEKQRSEEEEQEEYQRKGRGQGHNP | -43.46 | 25.8 | HHHHHHHHHHHHHHHHHHHHHHHHCCCCCCC |
| 320 | EQQQRHQQEDRSNRQEDQRYQRYGRGKGHQP | -43.44 | 32.3 | HHHHHHHHHHHHHHHHHHHHHHHHCCCCCCC |
| 758 | EEQERKKEDDWRNRWEQEYYQRKGQGQGHQP | -43.44 | 32.3 | HHHHHHHHHHHHHHHHHHHHHHHHCCCCCCC |
| 277 | EQEQRHQEEQRRNDKEQRRYQRYGRGQGHQP | -43.42 | 22.6 | HHHHHHHHHHHHHHHHHHHHHHHHHCCCCCC |

| | | | | |
|---|---|---|---|---|
| 761 | EQEERHQQEQRKNEKEQKRYQREGRGQGHQP | -43.42 | 25.8 | CHHHHHHHHHHHHHHHHHHHHHHHHCCCCCCC |
| 082 | EEEERKQRQQESNDKEQKRYQREGWGQGHYP | -43.40 | 25.8 | HHHHHHHHHHHHHHHHHHHHHHHHHCCCCCCC |
| 969 | DNERRKHEEDNKNRKEDEEYQRYGQGKGHQS | -43.39 | 25.8 | CHHHHHHHHHHHHHHHHHHHHHHHHHCCCCCC |
| 053 | EQQERHKEEKQRNEKEQKRYQRYGRGQGHQP | -43.38 | 32.3 | CHHHHHHHHHHHHHHHHHHHHHHHHHCCCCCC |
| 300 | EEQERKKQDDYKNRWEQEYYQQYGQGQGHQP | -43.38 | 32.3 | HHHHHHHHHHHHHHHHHHHHHHHHHHCCCCCC |
| 308 | EQEERHKEEQRSNDEEHRYYQKYGRGQGHQP | -43.37 | 22.6 | HHHHHHHHHHHHHHHHHHHHHHHHHCCCCCCC |
| 665 | EQEERHKEEQWRNRWEHEYYQKKGQGQGHNH | -43.37 | 22.6 | HHHHHHHHHHHHHHHHHHHHHHHHHCCCCCCC |
| 328 | EQEERKKQEKYKNEYEQEEYQRKGRGQGNQN | -43.35 | 29.0 | HHHHHHHHHHHHHHHHHHHHHHHHHCCCCCCC |
| 806 | EQERRKQEEQRSNEEEQQRYQREGWGQGHYP | -43.34 | 25.8 | HHHHHHHHHHHHHHHHHHHHHHHHHCCCCCCC |
| 704 | EQQQRHQEEQRRNEKEQEEYQRRGWGQGHYP | -43.27 | 29.0 | HHHHHHHHHHHHHHHHHHHHHHHHCCCCCCCC |
| 765 | QQQENHQREQESNEKEDKRYQKEGRGKGHQP | -43.25 | 32.3 | CHHHHHHHHHHHHHHHHHHHHHHHCCCCCCCC |
| 363 | EQEQRHEEEQKSNDWEHKYYQQKGQGQGHQP | -43.21 | 22.6 | HHHHHHHHHHHHHHHHHHHHHHHHHCCCCCCC |
| 285 | EEQERKQRDDQKNRREQERYQRQGRGQGHQP | -43.13 | 32.3 | HHHHHHHHHHHHHHHHHHHHHHHHHCCCCCCC |
| 291 | EQEERHQQEDRSNRQEDEQYQRYGKGKGHNN | -43.12 | 29.0 | CHHHHHHHHHHHHHHHHHHHHHHHHCCCCCCC |
| 292 | EEEERRRQEKEKNEKEDRRYQRYGRGKGHQP | -43.11 | 38.7 | HHHHHHHHHHHHHHHHHHHHHHHHHCCCCCCC |
| 849 | QQQENHRQEQQSNEWEDKYYQKKGQGKGHQP | -43.04 | 32.3 | CHHHHHHHHHHHHHHHHHHHHHHHHCCCCCCC |
| 869 | EEEERRQQEDRKNRHEDERYQRQGRGKGHQP | -42.90 | 32.3 | HHHHHHHHHHHHHHHHHHHHHHHHHCCCCCCC |
| 157 | EQERRHQEEQERNDKEHEEYQRRGKGQGHNN | -42.82 | 22.6 | HHHHHHHHHHHHHHHHHHHHHHHHCCCCCCCC |
| 828 | EQEERHQREQESNDKEHKRYQRYGRGQGHQP | -42.75 | 22.6 | CHHHHHHHHHHHHHHHHHHHHHHHHCCCCCCC |
| 730 | EEEERKRQDDESNRKEHEEYQRRGQGQGHQS | -42.68 | 29.0 | CHHHHHHHHHHHHHHHHHHHHHHHHHCCCCCC |
| 846 | QQEENHHQEQNKNDQEDRRYQRYGQGKGNQN | -42.67 | 25.8 | CHHHHHHHHHHHCHHHHHHHHHHHCCCCCCCC |
| 580 | EQQQRHNEEQRSNDKEQREYQRRGRGQGHQP | -42.59 | 25.8 | HHHHHHHHHHHHCHHHHHHHHHHHHCCCCCCC |
| 268 | EQEERKRQEQESNEKEQKEYQRSGKGQGHNN | -42.58 | 29.0 | HHHHHHHHHHHHHHHHHHHHHHHHCCCCCCCC |
| 762 | NQERQKQEEKREDEERQKSYQREGKGQGHNN | -42.39 | 22.6 | CHHHHHHHHHHHHHHHHHHHHHHHCCCCCCCC |
| 915 | EQQRRHQEEKYRNEYEHEEYQRKGRGQGHQP | -42.38 | 32.3 | HHHHHHHHHHHHHHHHHHHHHHHHHCCCCCCC |
| 002 | EQQERKRQQQEKNEKEQEEYQRRGKGQGHNH | -42.18 | 29.0 | HHHHHHHHHHHHHHHHHHHHHHHHHCCCCCCC |
| 512 | EQEERHREEQREEDERHRRKQKEGRGCGKQN | -41.94 | 16.1 | HHHHHHHHHHHHHHHHHHHHHHHHCCCCCCCC |
| 534 | EQERRHQEEQERNDKEQQKYQKEGRGQGHQP | -41.88 | 22.6 | HHHHHHHHHHHHHHHHHHHHHHHHHCCCCCCC |
| 295 | EQEERHNQEQQKNEREQKRYQREGKGQGHNN | -41.83 | 25.8 | HHHHHHHHHHHHHHHHHHHHHHHHCCCCCCCC |
| 397 | EEQERKKQEDQSNRREQERYQQQGRGQGHQP | -41.72 | 29.0 | HHHHHHHHHHHHHHHHHHHHHHHHHCCCCCCC |
| 677 | EEEERKKQQQKEDEERHKRYQRYGRGQGNQN | -41.71 | 25.8 | HHHHHHHHHHHHHHHHHHHHHHHHHCCCCCCC |
| 442 | QQQENHKEEDYRNRWEDQRYQREGRGKGHQP | -41.62 | 29.0 | CHHHHHHHHHHHHHHHHHHHHHHHHCCCCCCC |
| 218 | QQEENHQQEQERNRKEDERYQREGWGKGHYP | -41.52 | 25.8 | CHHHHHHHHHHHHHHHHHHHHHHHHCCCCCCC |
| 836 | EEQRRKQEQQYRDDFRQRMYQQYGKGQGHQP | -41.46 | 25.8 | HHHHHHHHHHHHHHHHHHHHHHHHHHCCCCCC |
| 173 | EQEERHRQEQEKNEKEQEEYQRQGRGQGHQP | -41.25 | 25.8 | HHHHHHHHHHHHHHHHHHHHHHHHHCCCCCCC |
| 690 | EQERRHQEEQQKNDREQRRYQREGRGQGHQP | -41.23 | 22.6 | HHHHHHHHHHHHHHHHHHHHHHHHHCCCCCCC |
| 466 | EQERRHQEEQERNEKEQEEYQRRGRGQGHQP | -41.11 | 25.8 | HHHHHHHHHHHHHHHHHHHHHHHHHHCCCCCC |
| 467 | QQEENHREEQERNEKEQEEYQRRGRGQGHQP | -40.82 | 22.6 | CHHHHHHHHHHHHHHHHHHHHHHHHHCCCCCC |

| | | | | |
|---|---|---|---|---|
| 127 | EQQERHRQEQEKNEKEQKRYQREGQGQGHQS | -40.81 | 29.0 | HHHHHHHHHHHHHHHHHHHHHHHHCCCCCCC |
| 199 | EQEERHQREQEEDDKRHRRYQREGKGQGHNN | -40.81 | 19.4 | CHHHHHHHHHHHHHHHHHHHHHHCCCCCCCC |
| 922 | EQEQRHEEEQRKNEKEQEEYQRYGRGQGHQP | -40.72 | 25.8 | HHHHHHHHHHHHHHHHHHHHHHHHHCCCCCC |
| 747 | QQEENHKQEDYKNRYEDEEYQRKGRGKGHQP | -40.71 | 25.8 | CHHHHHHHHHHHHHHHHHHHHHHHCCCCCCC |
| 640 | EQQRRHQEEQYRNEYEQEEYQRKGRGQGHQP | -40.67 | 29.0 | HHHHHHHHHHHHHHHHHHHHHHHHCCCCCCC |
| 851 | EQERRHQEEVWRNRYEQEEYQRKGRGQGHQP | -40.49 | 22.6 | HHHHHHHHHHHHHHHHHHHHHHHHCCCCCCC |
| 517 | EQEERHRQEQESNEREQQRYQREGRGQGHQP | -40.35 | 25.8 | HHHHHHHHHHHHHHHHHHHHHHHHCCCCCCC |
| 028 | EQQQRHNEEHRKNREEQEEYQRKGRGQGHNP | -39.86 | 25.8 | HHHHHHHHHHHHHHHHHHHHHHHHCCCCCCC |
| 221 | EEERRKQEQDESNRKEQERYQKQGRGQGHQP | -39.48 | 25.8 | HHHHHHHHHHHHHHHHHHHHHHHCCCCCCCC |
| 307 | EQEERHKQEDQKNRREHHEYQNRGRGQGNQN | -38.05 | 22.6 | HHHHHHHHHHHHHHHHHHHHHHCCCCCCCCC |

**Table S2**. Summary of 991 peptide sequences designed using EvoEF2 and the evolutionary profile (weight = 0.25).

| #label | #peptide | #binding(EEU) | #SeqID(%) | #Secondary structure |
|---|---|---|---|---|
| WT | EEQAKTFLDKFNHEAEDLFYQSSGLGKGDFR | -46.46 | 100. | HHHHHHHHHHHHHHHHHHHHHHHCCCCCCCC |
| 185 | EQEQRIQQDQFSNEFEDKYYQQKGQGKGHSH | -52.40 | 38.7 | HHHHHHHHHHHHHHHHHHHHHHHHCCCCCCC |
| 416 | EEERRKQLDQKKQDEEDSRFQQQGRGKGKQN | -51.50 | 35.5 | HHHHHHHHHHHHHHHHHHHHHHHHCCCCCCC |
| 731 | EEEQRKQQDKYNQESEDKRYQRYGWGKGHNP | -51.43 | 45.2 | HHHHHHHHHHHHHHHHHHHHHHHCCCCCCCC |
| 208 | EEQQRKQQDKYNNESEDKRFQRYGKGKGKSR | -51.41 | 48.4 | HHHHHHHHHHHCHHHHHHHHHHHCCCCCCCC |
| 148 | EEEQKKQQDKDNNEREDKRYQREGQGKGHQS | -51.13 | 48.4 | HHHHHHHHHHHHHHHHHHHHHHHCCCCCCCC |
| 765 | EEEQKKQQDKYNNEWEDKKYQREGRGKGHQP | -51.13 | 48.4 | HHHHHHHHHHHHHHHHHHHHHHHHCCCCCCC |
| 859 | EEEQKKQQDKYNNEYEDKQYQKEGWGKGHYP | -51.13 | 48.4 | HHHHHHHHHHHHHHHHHHHHHHHHCCCCCCC |
| 840 | EEEQKKQRDKYNNEHEDKRYQQQGKGKGHNH | -51.13 | 48.4 | HHHHHHHHHHHHHHHHHHHHHHHCCCCCCCC |
| 350 | EEEQKKQRDKYNNESEDKRYQRYGWGKGHNP | -51.13 | 48.4 | HHHHHHHHHHHCHHHHHHHHHHCCCCCCCCC |
| 666 | EEEQRKQQDKKNNEEEDKRYQRYGKGKGHSN | -51.13 | 45.2 | HHHHHHHHHHHHHHHHHHHHHHHCCCCCCCC |
| 725 | EEEQRKQQDKYNNEEEDKRYQREGWGKGHNP | -51.13 | 45.2 | HHHHHHHHHHHCHHHHHHHHHHHHCCCCCCC |
| 830 | EEEQRKQQDKYNNEYEDEEYQRKGRGKGHQP | -51.13 | 45.2 | HHHHHHHHHHHHHHHHHHHHHHHHCCCCCCC |
| 119 | EEEQRKQQDKYNNEYEDKEYQRKGRGKGHQP | -51.13 | 45.2 | HHHHHHHHHHHHHHHHHHHHHHHHCCCCCCC |
| 835 | EQEQRKQQDKYNNEYEDKEYQRKGRGKGHQP | -51.13 | 41.9 | HHHHHHHHHHHHHHHHHHHHHHHHCCCCCCC |
| 609 | EEEQRKQEDKRNNEQEDKRYQRYGRGKGHQP | -51.13 | 45.2 | HHHHHHHHHHHHHHHHHHHHHHHHCCCCCCC |
| 697 | EEEEKKQQDKFKNEFEDKYYQQKGQGKGHNH | -51.00 | 48.4 | CHHHHHHHHHHHHHHHHHHHHHHHCCCCCCC |
| 791 | EEEERKQRDKFNNEYEDKYYQQKGQGKGHQS | -51.00 | 48.4 | HHHHHHHHHHHHHHHHHHHHHHHHCCCCCCC |
| 342 | EEEERKQRDKYNNEAEDKRYQREGRGKGHQP | -51.00 | 48.4 | HHHHHHHHHHHHHHHHHHHHHHHHCCCCCCC |
| 359 | EEEERKQRDKYNNEYEDKQYQKEGRGKGHQP | -51.00 | 45.2 | HHHHHHHHHHHHHHHHHHHHHHHHCCCCCCC |
| 499 | EEERKKQLDKWKNEFEDKYYQQQGQGKGHQS | -51.00 | 48.4 | HHHHHHHHHHHHHHHHHHHHHHHHCCCCCCC |
| 659 | EEERRKQEDKYRNEWEDEEYQRKGRGKGHQP | -51.00 | 41.9 | HHHHHHHHHHHHHHHHHHHHHHHHCCCCCCC |
| 393 | EEERRKQLDKFKNEFEDKYYQQQGQGKGHSS | -51.00 | 48.4 | HHHHHHHHHHHHHHHHHHHHHHHHCCCCCCC |
| 453 | EEERRKQLDKQSNEEEDKRYQRYGQGKGHQS | -51.00 | 45.2 | HHHHHHHHHHHHHHHHHHHHHHHHHCCCCCC |
| 368 | EEERRKQLDKWKNEFEDKRYQQEGRGKGHQP | -51.00 | 45.2 | HHHHHHHHHHHHHHHHHHHHHHHHCCCCCCC |
| 085 | EEERRKQLDKYKNEAEDKRYQRYGKGKGHSN | -51.00 | 48.4 | HHHHHHHHHHHHHHHHHHHHHHHCCCCCCCC |
| 384 | EEERRKQLDKYNNEAEDKRYQREGQGKGHQS | -51.00 | 51.6 | HHHHHHHHHHHHHHHHHHHHHHHHCCCCCCC |
| 683 | EEEKRKQEDKRRHEKEDEEYQRRGWGKGHNP | -50.98 | 45.2 | HHHHHHHHHHHHHHHHHHHHHHHCCCCCCCC |
| 805 | EEEERKQQDKRNNEKEDKRYQRYGWGKGHYP | -50.94 | 45.2 | HHHHHHHHHHHHHHHHHHHHHHHCCCCCCCC |
| 959 | DEERKKQEDQYRNDWEDEEYQRKGRGKGHQP | -50.89 | 35.5 | HHHHHHHHHHHHHHHHHHHHHHHHCCCCCCC |
| 186 | EEEERKQRDQMKNDKEDKEYQKQGRGKGHQP | -50.89 | 35.5 | HHHHHHHHHHHHHHHHHHHHHHHHCCCCCCC |
| 471 | EQEERKNQDKYNNEYEDEEYQRKGRGKGHNP | -50.83 | 41.9 | HHHHHHHHHHHHHHHHHHHHHHHCCCCCCCC |
| 726 | EEEKRKQEDKRNNEEEDKRYQREGWGKGHNP | -50.81 | 45.2 | HHHHHHHHHHHHHHHHHHHHHHHHCCCCCCC |
| 321 | EEEKRKQLDKYKNEAEDKRYQQQGWGKGHNP | -50.81 | 48.4 | HHHHHHHHHHHHHHHHHHHHHHHCCCCCCCC |
| 279 | EEEQRKQQDKYNNEYEDKMYQREGKGKGHSH | -50.74 | 45.2 | HHHHHHHHHHHHHHHHHHHHHHHCCCCCCCC |

| | | | | |
|---|---|---|---|---|
| 407 | EEERRKQLDQESQEKEDEQYQRYGMGKGHQP | -50.54 | 41.9 | HHHHHHHHHHHHHHHHHHHHHHHHHCCCCCC |
| 669 | EEEQRKQQDQYNNDWEDQRYQREGRGKGHQP | -50.50 | 38.7 | HHHHHHHHHHHHHHHHHHHHHHHHCCCCCCC |
| 793 | EEEQKKQRDQYNNDSEDRRFQREGKGKGKNH | -50.49 | 38.7 | HHHHHHHHHHCCCCHHHHHHHHHCCCCCCCC |
| 087 | EEEQRKQQDQYNQEWEDEYYQRKGQGKGHQP | -50.48 | 41.9 | HHHHHHHHHHHHHHHHHHHHHHHHCCCCCCC |
| 909 | EEEQRKQQDQYNQEWEDEKYQREGKGKGHSS | -50.48 | 41.9 | HHHHHHHHHHHHHHHHHHHHHHHCCCCCCCC |
| 467 | EEEQRKQQDQYNQEYEDEEYQRKGRGKGHQP | -50.48 | 41.9 | HHHHHHHHHHHHHHHHHHHHHHHHCCCCCCC |
| 943 | EEEERKQQDQRNQEKEDERYQQQGRGKGHQP | -50.48 | 41.9 | HHHHHHHHHHHHHHHHHHHHHHHHCCCCCCC |
| 993 | EEEQRKQQDQQNNEREDEYYQQQGRGKGHQP | -50.37 | 41.9 | HHHHHHHHHHHHHHHHHHHHHHHHCCCCCCC |
| 658 | EEEQKKQRDQFNNEFEHEYYQQKGQGRGHNH | -50.35 | 41.9 | HHHHHHHHHHHHHHHHHHHHHHHHCCCCCCC |
| 962 | EEEQKKQQDKYNNESEDKRYQRYGRGKGHQP | -50.33 | 48.4 | HHHHHHHHHHHCHHHHHHHHHHHHCCCCCCC |
| 296 | EEEQKKQRDKYNNEYEDEQYQRKGKGKGHNN | -50.33 | 48.4 | HHHHHHHHHHHHHHHHHHHHHHHCCCCCCCC |
| 102 | EEEQRKQQDKYNNEWEDEEYQRKGRGKGHSN | -50.33 | 45.2 | HHHHHHHHHHHHHHHHHHHHHHHCCCCCCCC |
| 438 | EEEERKKQDKQNNEKEDKRFQREGQGKGKQS | -50.31 | 41.9 | HHHHHHHHHHHHHHHHHHHHHHHHCCCCCCC |
| 526 | EEEQKKQRDQRNNEQEDKRYQRYGRGKGHQP | -50.31 | 45.2 | HHHHHHHHHHHHHHHHHHHHHHHHCCCCCCC |
| 677 | EEEQRKQEDQRKNEKEDKRYQREGKGKGHSH | -50.31 | 38.7 | HHHHHHHHHHHHHHHHHHHHHHHCCCCCCCC |
| 893 | EEERRKQLDQKNNDEEHRRYQRYGWGRGHNP | -50.29 | 35.5 | HHHHHHHHHHHCHHHHHHHHHHHCCCCCCCC |
| 776 | EEEQRKQLDKKNNEEEDKRYQREGKGKGHSH | -50.29 | 48.4 | HHHHHHHHHHHHHHHHHHHHHHHCCCCCCCC |
| 966 | EEEEKKQQDQRKQEEEDERYQQQGRGKGHQP | -50.29 | 41.9 | CHHHHHHHHHHHHHHHHHHHHHHHCCCCCCC |
| 429 | EEQQRKQEDQYRNDHEDRKYQQEGRGKGHQP | -50.26 | 38.7 | HHHHHHHHHHHHHHHHHHHHHHHHCCCCCCC |
| 343 | EEEQRKQQDQRNQRQEDERYQREGKGKGHNH | -50.26 | 38.7 | HHHHHHHHHHHHHHHHHHHHHHHCCCCCCCC |
| 808 | EEEQKKQRDQYNNDWEDRRYQREGKGKGHSS | -50.25 | 41.9 | HHHHHHHHHHHHCHHHHHHHHHCCCCCCCCC |
| 377 | EEEERKKQDKYKNESEDKRYQQQGKGKGHNH | -50.25 | 41.9 | HHHHHHHHHHHHHHHHHHHHHHHHCCCCCCC |
| 380 | EEERKKQLDQQKNEREDSRYQQQGKGKGHSN | -50.24 | 45.2 | HHHHHHHHHHHHHHHHHHHHHHHCCCCCCCC |
| 614 | EEERRKQLDQENNEKEDQRYQREGKGKGHNH | -50.24 | 45.2 | HHHHHHHHHHHHHHHHHHHHHHHCCCCCCCC |
| 202 | EEEQKKQRDQRNQEKEDEEYQRYGWGKGHYP | -50.22 | 45.2 | HHHHHHHHHHHHHHHHHHHHHHHCCCCCCCC |
| 442 | EEEQKKQRDQYNNRYEDEEYQRKGRGKGHQP | -50.22 | 41.9 | HHHHHHHHHHHHHHHHHHHHHHHHCCCCCCC |
| 918 | EQQQRKQQDQYNNESEDQRYQRYGWGKGHNP | -50.18 | 41.9 | HHHHHHHHHHHCHCHHHHHHHHCCCCCCCCC |
| 328 | EEEERKNRDQYNQEWEDEEYQRKGRGKGHQP | -50.18 | 41.9 | HHHHHHHHHHHHHHHHHHHHHHHHCCCCCCC |
| 092 | EEEQKKQRDQFNNEEEDKRYQREGRGKGHQP | -50.18 | 48.4 | HHHHHHHHHHHCHHHHHHHHHHHHCCCCCCC |
| 530 | EEEQKKQRDQYNNEAEDERYQRQGNGKGHSH | -50.18 | 48.4 | HHHHHHHHHHHHHHHHHHHHHHHCCCCCCCC |
| 262 | EEEQKKQRDQYNNEAEDKRYQRYGWGKGHNP | -50.18 | 48.4 | HHHHHHHHHHHHHHHHHHHHHHCCCCCCCCC |
| 820 | EEEQKKQRDQYNNEWEDEEYQRKGRGKGHQP | -50.18 | 45.2 | HHHHHHHHHHHHHHHHHHHHHHHHCCCCCCC |
| 678 | EEEQKKQRDQYNNEWEDERYQRQGWGKGHYP | -50.18 | 45.2 | HHHHHHHHHHHHHHHHHHHHHHHHCCCCCCC |
| 418 | EEEQKKQRDQYNNEYEDEEYQRKGRGKGHNP | -50.18 | 45.2 | HHHHHHHHHHHHHHHHHHHHHHHCCCCCCCC |
| 130 | EEEQKKQRDQYNNEYEDQEYQRTGRGKGHQP | -50.18 | 45.2 | HHHHHHHHHHHHHHHHHHHHHHCCCCCCCCC |
| 914 | EEEQRKQEDQYRNEYEDQEYQRKGRGKGHQP | -50.18 | 38.7 | HHHHHHHHHHHHHHHHHHHHHHHHCCCCCCC |
| 014 | EEEQRKQQDQKNNEEEDERYQQQGWGKGHYP | -50.18 | 41.9 | HHHHHHHHHHHHHHHHHHHHHHHCCCCCCCC |
| 802 | EEEQRKQQDQKNNEEEDKRYQRYGRGKGHQP | -50.18 | 41.9 | HHHHHHHHHHHHHHHHHHHHHHHHCCCCCCC |

| | | | | |
|---|---|---|---|---|
| 487 | EEEQRKQQDQYNNEAEDKRYQRYGRGKGHQP | -50.18 | 45.2 | HHHHHHHHHHHHHHHHHHHHHHHHCCCCCCC |
| 144 | EEEQRKQQDQYNNEAEDQRYQQQGRGKGHQP | -50.18 | 45.2 | HHHHHHHHHHHHHHHHHHHHHHHHCCCCCCC |
| 055 | EEEQRKQQDQYNNEAEDQRYQREGKGKGHNH | -50.18 | 45.2 | HHHHHHHHHHHHHHHHHHHHHHHHCCCCCCC |
| 861 | EEEQRKQQDQYNNEFEDQMYQRYGKGKGHQP | -50.18 | 41.9 | HHHHHHHHHHHHHHHHHHHHHHHHHCCCCCC |
| 941 | EEEQRKQQDQYNNESEDQRYQQYGWGKGHYP | -50.18 | 41.9 | HHHHHHHHHHHCHCHHHHHHHHHCCCCCCCC |
| 196 | EEEQRKQQDQYNNEWEDKRYQREGRGKGHQP | -50.18 | 41.9 | HHHHHHHHHHHHHHHHHHHHHHHHCCCCCCC |
| 767 | EEEQRKQQDQYNNEYEDEEYQRKGRGKGHQP | -50.18 | 41.9 | HHHHHHHHHHHHHHHHHHHHHHHHCCCCCCC |
| 833 | EEEQRKQQDQYNNEYEDKYYQKQGQGKGHQS | -50.18 | 41.9 | HHHHHHHHHHHHHHHHHHHHHHHHCCCCCCC |
| 829 | EEEQRKQQDQYNNEYEDQQYQKEGQGKGHQS | -50.18 | 41.9 | HHHHHHHHHHHHHHHHHHHHHHHHCCCCCCC |
| 040 | EEEQRKQQDQYSNEYEDEEYQRKGRGKGHQP | -50.18 | 38.7 | HHHHHHHHHHHHHHHHHHHHHHHHCCCCCCC |
| 382 | EEEQRKQQDQYSNEYEDKEYQKEGKGKGHNH | -50.18 | 38.7 | HHHHHHHHHHHHHHHHHHHHHHHCCCCCCCC |
| 379 | EEEERKQQDQRSNEKEDEEYQRRGKGKGHNH | -50.18 | 38.7 | HHHHHHHHHHHHHHHHHHHHHHHHCCCCCCC |
| 929 | EEEERKQQDQRSNEQEDQRYQREGKGKGHSS | -50.18 | 38.7 | HHHHHHHHHHHHHHHHHHHHHHHCCCCCCCC |
| 167 | EQEQRKQEDQRKNEKEDKRYQREGKGKGHSH | -50.18 | 35.5 | HHHHHHHHHHHHHHHHHHHHHHHCCCCCCCC |
| 063 | EEEARKQQDQYNQEWEDQRYQREGKGKGHNH | -50.16 | 45.2 | HHHHHHHHHHHHHHHHHHHHHHHHCCCCCCC |
| 094 | DEERKKQEDQYRNDYEDREYQRKGRGKGHQP | -50.12 | 35.5 | HHHHHHHHHHHHHHHHHHHHHHHHCCCCCCC |
| 509 | EEERRKQLDQMKNDKEDREYQRSGQGKGHQS | -50.12 | 41.9 | HHHHHHHHHHHHHHHHHHHHHHHCCCCCCCC |
| 173 | EEERRKQLDQWKNDYEDRYYQQQGQGKGHQP | -50.12 | 38.7 | HHHHHHHHHHHHHHHHHHHHHHHHCCCCCCC |
| 632 | EEERRKQQDQYSNDYEDREYQRKGRGKGHQP | -50.12 | 35.5 | HHHHHHHHHHHCCHHHHHHHHHHCCCCCCCC |
| 657 | EQEERKQRDQYNNEAEDQRFQREGKGKGKNH | -50.10 | 38.7 | HHHHHHHHHHHHHHHHHHHHHHHHCCCCCCC |
| 934 | EEEKKKQEDQRNNREEDERYQREGKGKGHSS | -50.09 | 41.9 | HHHHHHHHHHHHHHHHHHHHHHHCCCCCCCC |
| 860 | EEEERKQQDQRKNRKEDEEYQKQGRGKGHQP | -50.06 | 35.5 | HHHHHHHHHHHHHHHHHHHHHHHCCCCCCCC |
| 690 | EQEERKQQDQRKNEQEDEEYQRKGRGKGHQP | -50.05 | 35.5 | HHHHHHHHHHHHHHHHHHHHHHHHCCCCCCC |
| 336 | DEERKKQEDQYRNEWEDEEYQRKGRGKGHNP | -50.05 | 38.7 | HHHHHHHHHHHHHHHHHHHHHHHCCCCCCCC |
| 243 | EEEERKQQDQYNNESEDKRYQRYGWGKGHNP | -50.05 | 41.9 | HHHHHHHHHHHCHHHHHHHHHHCCCCCCCCC |
| 968 | EEEERKQRDQFNNEFEDQYYQRYGQGKGHNP | -50.05 | 45.2 | HHHHHHHHHHHHHHHHHHHHHHHHHCCCCCC |
| 383 | EEEERKQRDQQSNEEEDQRYQRYGWGKGHNP | -50.05 | 38.7 | HHHHHHHHHHHHHHHHHHHHHHHCCCCCCCC |
| 661 | EEEERKQRDQYNNEAEDKRYQREGKGKGHSS | -50.05 | 45.2 | HHHHHHHHHHHHHHHHHHHHHHHCCCCCCCC |
| 149 | EEEERKQRDQYNNEYEDEEYQRKGRGKGHQP | -50.05 | 41.9 | HHHHHHHHHHHHHHHHHHHHHHHHCCCCCCC |
| 203 | EEEKKKQEDQRNNEEEDKRYQREGRGKGHQP | -50.05 | 45.2 | HHHHHHHHHHHHHHHHHHHHHHHHCCCCCCC |
| 994 | EEEKRKQEDQRNNEEEDERYQRYGWGKGHYP | -50.05 | 41.9 | HHHHHHHHHHHHHHHHHHHHHHHCCCCCCCC |
| 054 | EEEKRKQEDQRNNEEEDKRYQREGKGKGHNN | -50.05 | 41.9 | HHHHHHHHHHHHHHHHHHHHHHHCCCCCCCC |
| 200 | EEERKKQLDQQNNEEEDKRYQRYGKGKGHSS | -50.05 | 48.4 | HHHHHHHHHHHHHHHHHHHHHHHCCCCCCCC |
| 713 | EEERKKQLDQYNNEWEDEEYQRKGRGKGHQP | -50.05 | 48.4 | HHHHHHHHHHHHHHHHHHHHHHHHCCCCCCC |
| 583 | EEERKKQLDQYNNEYEDEYYQQKGQGKGHSH | -50.05 | 48.4 | HHHHHHHHHHHHHHHHHHHHHHHHCCCCCCC |
| 058 | EEERRKQLDQKNNEEEDQRYQKQGWGKGHNP | -50.05 | 45.2 | HHHHHHHHHHHHHHHHHHHHHHHCCCCCCCC |
| 596 | EEERRKQLDQQSNEEEDSRYQQQGKGKGHSN | -50.05 | 41.9 | HHHHHHHHHHHHHHHHHHHHHHHCCCCCCCC |
| 366 | EEERRKQLDQYNNEEEDKRYQREGKGKGHNH | -50.05 | 45.2 | HHHHHHHHHHHCCHHHHHHHHHHHCCCCCCC |

| | | | | |
|---|---|---|---|---|
| 510 | EEERRKQLDQYNNEYEDEEYQRKGRGKGHQP | -50.05 | 45.2 | HHHHHHHHHHHHHHHHHHHHHHHHCCCCCCC |
| 740 | EEEERKNQDQQKNEEEDQRYQREGRGKGHQP | -50.00 | 38.7 | HHHHHHHHHHHHHHHHHHHHHHHHCCCCCCC |
| 867 | DEERKKQEDQRRNEKEDKRYQREGKGKGHNH | -49.99 | 38.7 | CHHHHHHHHHHHHHHHHHHHHHHCCCCCCCC |
| 386 | EEEEKKQRDQRSNEQEDQRYQRYGRGKGHQP | -49.99 | 41.9 | CHHHHHHHHHHHHHHHHHHHHHHHCCCCCCC |
| 266 | EEEERKQQDQRKNEKEDEEYQRRGQGKGHQS | -49.99 | 38.7 | HHHHHHHHHHHHHHHHHHHHHHHHHCCCCCC |
| 255 | EEEERKQQDQRKNEKEDEEYQRRGRGKGHQP | -49.99 | 38.7 | HHHHHHHHHHHHHHHHHHHHHHHHCCCCCCC |
| 400 | EEEERKQQDQRNNEAEDKRYQKEGWGKGHNP | -49.99 | 45.2 | HHHHHHHHHHHHHHHHHHHHHHHHCCCCCCC |
| 038 | EEEQKKQEDQRKNEKEDKRYQRYGWGKGHNP | -49.99 | 41.9 | HHHHHHHHHHHHHHHHHHHHHHCCCCCCCCC |
| 110 | EEEQRKQMDQMKNEKEDEEYQRRGRGKGHQP | -49.94 | 38.7 | HHHHHHHHHHHHHHHHHHHHHHHHCCCCCCC |
| 494 | EEEQRKQQDQFNHEFEDQYYQQKGQGKGHSH | -49.94 | 48.4 | HHHHHHHHHHHHHHHHHHHHHHHHCCCCCCC |
| 898 | EQEERKQRDQYNNDWEDRRYQREGKGKGHNH | -49.93 | 35.5 | HHHHHHHHHHHCCHHHHHHHHHCCCCCCCCC |
| 406 | EEEQRKQQDQRNNEEEDKKYQRYGKGKGHSS | -49.92 | 41.9 | HHHHHHHHHHHHHHHHHHHHHHHHCCCCCCC |
| 206 | EEEQRKQQDQRNNEQEDQRYQRYGRGKGHQP | -49.92 | 41.9 | HHHHHHHHHHHHHHHHHHHHHHHHCCCCCCC |
| 884 | EEEEKKNRDQYNNEYEDEEYQRKGRGKGHNP | -49.88 | 45.2 | HHHHHHHHHHHHHHHHHHHHHHHCCCCCCCC |
| 370 | EEEQRKNEDQFRNEFEDEFYQQQGKGKGHNH | -49.88 | 45.2 | HHHHHHHHHHHHHHHHHHHHHHHHCCCCCCC |
| 458 | EEEEKKKRDKYNNEWEDEEYQRKGRGKGHQP | -49.88 | 48.4 | CHHHHHHHHHHHHHHHHHHHHHHHCCCCCCC |
| 839 | EEEERKKEDKKRNEEEDKRYQRYGRGKGHQP | -49.88 | 41.9 | HHHHHHHHHHHHHHHHHHHHHHHHCCCCCCC |
| 762 | EEEERKKQDKFNNEFEDKFYQQQGKGKGHSH | -49.88 | 51.6 | HHHHHHHHHHHHHHHHHHHHHHHHCCCCCCC |
| 333 | EEEERKKQDKYNNEYEDEEYQRKGRGKGHQP | -49.88 | 45.2 | HHHHHHHHHHHHHHHHHHHHHHHHCCCCCCC |
| 771 | DEERKKQEDQYRNESEDQRYQRYGRGKGHQP | -49.86 | 38.7 | HHHHHHHHHHHHHHHHHHHHHHHHCCCCCCC |
| 907 | EEEAKKQRDQYNNEYEDEEYQRQGRGKGHQS | -49.86 | 48.4 | HHHHHHHHHHHHHHHHHHHHHHHHCCCCCCC |
| 750 | EEEARKQLDQWKNEFEDKYYQKEGQGKGHQP | -49.86 | 45.2 | HHHHHHHHHHHHHHHHHHHHHHHHCCCCCCC |
| 979 | EEEEKKQRDQKNNEEEDERYQKQGRGKGHQP | -49.86 | 45.2 | HHHHHHHHHHHHHHHHHHHHHHHCCCCCCCC |
| 062 | EEEERKQQDQQKNEEEDSRYQQEGKGKGHSH | -49.86 | 38.7 | HHHHHHHHHHHHHHHHHHHHHHHCCCCCCCC |
| 032 | EEEERKQRDQFNNEKEDQRYQQQGQGKGHQS | -49.86 | 45.2 | HHHHHHHHHHHHHHHHHHHHHHHHCCCCCCC |
| 591 | EEEERKQRDQYNNEAEDQRYQREGRGKGHQP | -49.86 | 45.2 | HHHHHHHHHHHHHHHHHHHHHHHHCCCCCCC |
| 976 | EEEERKQRDQYNNESEDKRYQRYGWGKGHNP | -49.86 | 41.9 | HHHHHHHHHHHCCHHHHHHHHHCCCCCCCCC |
| 913 | EEEKRKQLDQWKNEYEDKSYQREGKGKGHNH | -49.86 | 41.9 | HHHHHHHHHHHHHHHHHHHHHHCCCCCCCCC |
| 760 | EDEQRKQQDQYNNRAEDERYQKEGQGKGHQS | -49.86 | 38.7 | CHHHHHHHHHHHHHHHHHHHHHHHCCCCCCC |
| 170 | EEEERKQQDQRNNEKEHKRKQRYGRGCGKQN | -49.84 | 32.3 | HHHHHHHHHHHHHHHHHHHHHHHCCCCCCCC |
| 423 | EEEERKQRDQFNNEFEDQYYQQKGQGKGHSH | -49.82 | 45.2 | HHHHHHHHHHHHHHHHHHHHHHHHCCCCCCC |
| 460 | EEEQRKQEDQYRNEHEDERYQQQGRGKGHQP | -49.79 | 38.7 | HHHHHHHHHHHHHHHHHHHHHHHHCCCCCCC |
| 610 | EEEQRKQQDQRNNEKEDQEYQRSGKGKGHSS | -49.79 | 45.2 | HHHHHHHHHHHHHHHHHHHHHHCCCCCCCCC |
| 781 | EQEERKQRDKYNQEYEDEEYQRKGRGKGNQN | -49.76 | 41.9 | HHHHHHHHHHHHHHHHHHHHHHHHCCCCCCC |
| 078 | EEEQRKNEDQRNNEKEDQEYQKQGKGKGHSS | -49.76 | 41.9 | HHHHHHHHHHHHHHHHHHHHHHHCCCCCCCC |
| 222 | EQEQRKQQEKYSQEYEDKEYQRQGRGKGHQP | -49.71 | 35.5 | HHHHHHHHHHHHHHHHHHHHHHHHCCCCCCC |
| 900 | EEEERKIQDQQKNEKEDEEYQRRGRGKGHQP | -49.71 | 38.7 | CHHHHHHHHHHHHHHHHHHHHHHHCCCCCCC |
| 463 | EEEERKSRDKFNQEFEDKYYQKEGQGKGHSH | -49.71 | 48.4 | CHHHHHHHHHHHHHHHHHHHHHHCCCCCCCC |

| | | | | |
|---|---|---|---|---|
| 937 | EEEQKKQQDQKNNREEDERYQKEGKGKGHNH | -49.70 | 41.9 | HHHHHHHHHHHHHHHHHHHHHHHCCCCCCCC |
| 702 | EEEQRKQQDQYNQEWEDKRYQREGRGKGHQP | -49.68 | 41.9 | HHHHHHHHHHHHHHHHHHHHHHHHCCCCCCC |
| 153 | EEEQRKQQDQYSNEFEDQYYQQQGQGKGHQS | -49.66 | 38.7 | HHHHHHHHHHHHHHHHHHHHHHHHCCCCCCC |
| 890 | EEEEKKKRDKYNNEWEDEYYQRKGQGKGHQH | -49.64 | 48.4 | CHHHHHHHHHHHHHHHHHHHHHHHCCCCCCC |
| 313 | EEEEKKQQDQRNNDKEHQYKQKYGQGSGKQN | -49.62 | 32.3 | CHHHHHHHHHHHHHHHHHHHHHHCCCCCCCC |
| 489 | EEEAKKFRDMFNNRFEDQYYQQQGQGKGHQP | -49.61 | 51.6 | HHHHHHHHHHHHHHHHHHHHHHHHCCCCCCC |
| 143 | EEERRKQLDQWKNEWEDRYYQQKGQGKGHNH | -49.61 | 41.9 | HHHHHHHHHHHHHHHHHHHHHHHHCCCCCCC |
| 164 | EEEQRKQQDQRNNDKEDRRYQREGKGKGHNH | -49.58 | 38.7 | HHHHHHHHHHHHCHHHHHHHHHHCCCCCCCC |
| 667 | EEEERKQRDQFNNRFEDEFYQQKGRGKGHQP | -49.52 | 45.2 | HHHHHHHHHHHHHHHHHHHHHHHHCCCCCCC |
| 125 | EEEQRKQEDKRRNEKEDRRYQREGRGKGHQP | -49.47 | 41.9 | HHHHHHHHHHHHHHHHHHHHHHHHCCCCCCC |
| 777 | EEEQRKQQDQYNNDWEDRKYQREGRGKGHQP | -49.45 | 38.7 | HHHHHHHHHHHCCHHHHHHHHHHHCCCCCCC |
| 000 | EEERRKQLDQFKNEYEDQSYQREGRGKGHQP | -49.44 | 45.2 | HHHHHHHHHHHHHHHHHHHHHHHCCCCCCCC |
| 613 | EEERRKQLDQWKNEFEDKYYQQKGQGKGHQP | -49.44 | 41.9 | HHHHHHHHHHHHHHHHHHHHHHHHCCCCCCC |
| 733 | EQERRKQLDQMKNEKEDEEYQRRGRGKGHQP | -49.44 | 38.7 | HHHHHHHHHHHHHHHHHHHHHHHHCCCCCCC |
| 674 | EEEQKKQQEKYNNESEDKRYQRYGWGKGHNP | -49.42 | 45.2 | HHHHHHHHHHHHHHHHHHHHHHCCCCCCCCC |
| 939 | EEEQKKQQEKYSNEYEDKEYQRNGRGKGHQY | -49.42 | 41.9 | HHHHHHHHHHHHHHHHHHHHHHCCCCCCCCC |
| 216 | EQEQRKQEEKRNNEKEDKRYQREGRGKGHQP | -49.42 | 38.7 | HHHHHHHHHHHHHHHHHHHHHHHHCCCCCCC |
| 331 | EEEERKQRDKYNNESEDRRYQRYGRGKGHQP | -49.41 | 45.2 | HHHHHHHHHHHCCHHHHHHHHHHHCCCCCCC |
| 050 | EEEQKKQQDQYNNEEEDKKYQREGHGKGHQR | -49.38 | 48.4 | HHHHHHHHHHHCHHHHHHHHHHHHCCCCCCC |
| 699 | EEEQKKQRDQYNNESEDQRYQRYGRGKGHQP | -49.38 | 45.2 | HHHHHHHHHHHCCCHHHHHHHHHHCCCCCCC |
| 735 | EEEQKKQRDQYNNEYEDQEYQRQGRGKGHQP | -49.38 | 45.2 | HHHHHHHHHHHHHHHHHHHHHHHHCCCCCCC |
| 045 | EEEQRKQQDQKNNEEEDKRYQQEGKGKGHNH | -49.38 | 41.9 | HHHHHHHHHHHHHHHHHHHHHHHHCCCCCCC |
| 177 | EEEQRKQQDQYNNESEDERYQQQGKGKGHNH | -49.38 | 41.9 | HHHHHHHHHHHCCCHHHHHHHHHHCCCCCCC |
| 705 | EEEQRKQQDQYNNESEDERYQQQGWGKGHYP | -49.38 | 41.9 | HHHHHHHHHHHCCCHHHHHHHHHHCCCCCCC |
| 572 | EEEQRKQQDQYNNESEDQRYQRYGRGKGHQP | -49.38 | 41.9 | HHHHHHHHHHHCCCHHHHHHHHHHCCCCCCC |
| 549 | EEEQRKQQDQYNNEWEDKRYQREGKGKGHSS | -49.38 | 41.9 | HHHHHHHHHHHHHHHHHHHHHHCCCCCCCCC |
| 511 | EEEERKKQDQWKNDWEDRYYQQKGQGKGHNH | -49.37 | 35.5 | HHHHHHHHHHHHHHHHHHHHHHHHCCCCCCC |
| 800 | DEERKKQEDQQRNDKEHKRYQREGKGQGHNH | -49.36 | 29.0 | CHHHHHHHHHHHHHHHHHHHHHHCCCCCCCC |
| 935 | EEEEKKKRDQWKNRWEDEYYQQKGQGKGHNH | -49.33 | 38.7 | HHHHHHHHHHHHHHHHHHHHHHHHCCCCCCC |
| 687 | EEEERKYQDKYNQESEDKRYQRYGWGKGHNP | -49.30 | 45.2 | CHHHHHHHHHHHHHHHHHHHHHCCCCCCCCC |
| 529 | EEEERKKQDQYKNEYEDEEYQRKGRGKGHQP | -49.30 | 38.7 | HHHHHHHHHHHHHHHHHHHHHHHHCCCCCCC |
| 844 | EEEERKKRDQYNNEAEDQRYQREGWGKGHYP | -49.30 | 45.2 | HHHHHHHHHHHHHHHHHHHHHHHHCCCCCCC |
| 107 | EQERRKQLEKWKNEWEDKYYQKEGQGKGHQP | -49.29 | 38.7 | HHHHHHHHHHHHHHHHHHHHHHHCCCCCCCC |
| 540 | EEEQKKQQDQYNNEYEDEEYQRKGRGKGNQN | -49.24 | 45.2 | HHHHHHHHHHHHHHHHHHHHHHHHCCCCCCC |
| 001 | EEEERKKQDQYNNESEDQRFQKEGKGKGKSN | -49.23 | 38.7 | HHHHHHHHHHHCCCHHHHHHHHHCCCCCCCC |
| 084 | EEEEKKKRDQYNQESEDERYQQQGWGKGHNP | -49.23 | 45.2 | CHHHHHHHHHHHHHHHHHHHHHHCCCCCCCC |
| 573 | EEEERKKLDQWKQEYEDEEYQRNGSGKGHQY | -49.23 | 41.9 | CHHHHHHHHHHHHHHHHHHHHHCCCCCCCCC |
| 067 | EEEERKKQDQYNQEYEDEEYQRKGRGKGHQP | -49.23 | 41.9 | HHHHHHHHHHHHHHHHHHHHHHHHCCCCCCC |

| | | | | |
|---|---|---|---|---|
| 891 | EEEEKKQREKRSNEKEDEEYQRRGRGKGHQP | -49.22 | 41.9 | CHHHHHHHHHHHHHHHHHHHHHHHCCCCCCC |
| 214 | EEEKKKMLDMKNNREEDERYQQQGWGKGHNP | -49.22 | 45.2 | HHHHHHHHHHHHHHHHHHHHHHHCCCCCCCC |
| 498 | EEEKRKMLDMYKNRYEDEEYQRKGRGKGHQP | -49.22 | 38.7 | HHHHHHHHHHHHHHHHHHHHHHHHCCCCCCC |
| 473 | EEERRKQLDMINNRQEDQRYQREGKGKGHSS | -49.20 | 41.9 | HHHHHHHHHHHHHHHHHHHHHHHCCCCCCCC |
| 623 | EQEQRKQQEKYSNEYEDEEYQRKGRGKGHSN | -49.18 | 35.5 | HHHHHHHHHHHHHHHHHHHHHHHCCCCCCCC |
| 451 | EEEMRKLQDQINNEEEDERYQQQGWGKGHYP | -49.17 | 41.9 | HHHHHHHHHHHHHHHHHHHHHHHCCCCCCCC |
| 592 | EDEQRKQQDQRSNRKEDEEYQKRGWGKGHYP | -49.15 | 32.3 | CHHHHHHHHHHHHHHHHHHHHHHCCCCCCCC |
| 325 | EEEKRKQLEKYNNESEDKRYQQYGQGKGHQS | -49.10 | 45.2 | HHHHHHHHHHHHHHHHHHHHHHHHCCCCCCC |
| 480 | EEEQRKHEDKNRHEQEDKRYQREGKGKGHSH | -49.08 | 45.2 | HHHHHHHHHHHHHHHHHHHHHHHCCCCCCCC |
| 204 | EQQQRKQQEKYSNEYEQKEYQKEGRGRGHQP | -49.06 | 32.3 | HHHHHHHHHHHHHHHHHHHHHHHHCCCCCCC |
| 034 | EEEEKKYQDQYNNDYEDEEYQRKGRGKGHQP | -49.06 | 41.9 | CHHHHHHHHHHHCCCHHHHHHHHHCCCCCCC |
| 231 | EEEERKNRDKYNNEAEDKRYQREGKGKGHSS | -49.05 | 48.4 | HHHHHHHHHHHCHHHHHHHHHHHCCCCCCCC |
| 502 | EEEERKKQDQYNNEYEDEEFQRKGRGKGKQN | -49.04 | 38.7 | HHHHHHHHHHHHHHHHHHHHHHHHCCCCCCC |
| 068 | EEEQRKQQDAYNNRAEDERYQRYGRGKGHQP | -49.03 | 41.9 | HHHHHHHHHHHHHHHHHHHHHHHHCCCCCCC |
| 320 | EQERRKQQEKYSNESEDKRYQRYGKGKGHNS | -49.02 | 35.5 | HHHHHHHHHHHHHHHHHHHHHHHHCCCCCCC |
| 069 | EEEKRKWLDKLNNEKEDEKFQREGQGKGKQS | -49.01 | 45.2 | HHHHHHHHHHHHHHHHHHHHHHHHCCCCCCC |
| 599 | EEEERKYQDKKNNEEEDKRYQRYGKGKGHSS | -49.00 | 45.2 | CHHHHHHHHHHHHHHHHHHHHHHCCCCCCCC |
| 175 | EEEERKYQDKYKNEWEDKKYQEEGKGKGHNH | -49.00 | 41.9 | CHHHHHHHHHHHHHHHHHHHHHHHCCCCCCC |
| 426 | EEEERKQQDQRNNDSEHRRKQKEGWGSGKYP | -48.99 | 29.0 | HHHHHHHHHHHHHHHHHHHHHHHHCCCCCCC |
| 281 | EEEQRKQQDQYNNEWEHEEKQRKGRGSGKQN | -48.98 | 32.3 | HHHHHHHHHHHHHHHHHHHHHHHCCCCCCCC |
| 799 | EEEERKKQDQYNNEAEDQRYQREGKGKGHNH | -48.98 | 45.2 | HHHHHHHHHHHHHHHHHHHHHHHHCCCCCCC |
| 395 | EEEERKKQDQYNNRYEDEEYQRKGRGKGHQP | -48.96 | 38.7 | HHHHHHHHHHHHHHHHHHHHHHHHCCCCCCC |
| 420 | EEEARKNQDQYNQESEDERYQQQGKGKGDSR | -48.94 | 51.6 | HHHHHHHHHHHHHHHHHHHHHHHCCCCCCCC |
| 369 | EEEQRKQQDQYNNEAEDQRYQREGKGKGDSR | -48.94 | 51.6 | HHHHHHHHHHHHHHHHHHHHHHHCCCCCCCC |
| 345 | EEEERKQQQKKNNEEEDKRFQRYGKGKGKNR | -48.93 | 41.9 | HHHHHHHHHHHHHHHHHHHHHHHCCCCCCCC |
| 653 | EEEEKKKRDQYNNEWEDQKYQREGWGKGHNP | -48.93 | 45.2 | CHHHHHHHHHHHHHHHHHHHHHHHCCCCCCC |
| 753 | EEEEKKKRDQYNNEYEDEEYQRKGRGKGHQP | -48.93 | 45.2 | CHHHHHHHHHHHHHHHHHHHHHHHCCCCCCC |
| 348 | EEEEKKKRDQYNNEYEDQEYQRKGRGKGHQP | -48.93 | 45.2 | CHHHHHHHHHHHHHHHHHHHHHHHCCCCCCC |
| 636 | EEEERKKEDQWRNEEEDKKYQREGRGKGHQP | -48.93 | 38.7 | HHHHHHHHHHHCHHHHHHHHHHHHCCCCCCC |
| 845 | EEEERKKQDQYNNESEDQRYQQQGWGKGHNP | -48.93 | 41.9 | HHHHHHHHHHHCCCHHHHHHHHHHCCCCCCC |
| 588 | EEEERKKQDQYNNESEDQRYQRYGWGKGHYP | -48.93 | 41.9 | HHHHHHHHHHHCCCHHHHHHHHHCCCCCCCC |
| 565 | EEEERKKQDQYNNESEDSRYQQQGKGKGHNH | -48.93 | 41.9 | HHHHHHHHHHHCCCHHHHHHHHHCCCCCCCC |
| 166 | EEEERKKQDQYNNEWEDQRYQREGWGKGHYP | -48.93 | 41.9 | HHHHHHHHHHHHHHHHHHHHHHHHCCCCCCC |
| 305 | EEEERKKQDQYSNEYEDEEYQRKGRGKGHQP | -48.93 | 38.7 | HHHHHHHHHHHHHHHHHHHHHHHHCCCCCCC |
| 156 | QEEEKKKQDQYNNEHEDERYQRQGWGKGHYP | -48.93 | 41.9 | CHHHHHHHHHHHHHHHHHHHHHHHCCCCCCC |
| 822 | EEEKRKQEDQRNNEEEDQYYQQKGQGKGNQN | -48.92 | 41.9 | HHHHHHHHHHHHHHHHHHHHHHHHCCCCCCC |
| 543 | EEEKRKLLDQIENEREDQYYQQKGQGKGHQP | -48.91 | 41.9 | HHHHHHHHHHHHHHHHHHHHHHHHCCCCCCC |
| 049 | EEEQRKQEQKRRNEKEDKQYQRSGRGKGHQP | -48.91 | 41.9 | HHHHHHHHHHHHHHHHHHHHHHCCCCCCCCC |

| | | | | |
|---|---|---|---|---|
| 104 | EEEQRKFEDKRNNEEEDKRYQQQGRGKGHQP | -48.90 | 48.4 | CHHHHHHHHHHHHHHHHHHHHHHHCCCCCCC |
| 459 | EEEEKKQRDQYNNEYEHQEKQRKGRGSGKSN | -48.86 | 35.5 | HHHHHHHHHHHHHHHHHHHHHHCCCCCCCCC |
| 165 | EEEERKQRDQYNNEAEDQRYQREGKGKGHNH | -48.85 | 45.2 | HHHHHHHHHHHHHHHHHHHHHHHHCCCCCCC |
| 536 | EEEERKKQDQYNNRYEHEEKQKEGRGCGKQN | -48.85 | 29.0 | HHHHHHHHHHHHHHHHHHHHHHHCCCCCCCC |
| 385 | EQEERKQQEQRSQEKEDKEFQKQGRGKGKQN | -48.82 | 29.0 | HHHHHHHHHHHHHHHHHHHHHHHHCCCCCCC |
| 183 | EEEARKSQDKQNNEREDKRFQREGKGKGKNH | -48.78 | 45.2 | HHHHHHHHHHHHHHHHHHHHHHHCCCCCCCC |
| 264 | EEEQRKQQQKYNNEYEDEEYQRKGRGKGHQP | -48.78 | 41.9 | HHHHHHHHHHHHHHHHHHHHHHHHCCCCCCC |
| 712 | EEEQRKQQQKYSNEWEDEEYQRKGRGKGHQP | -48.78 | 38.7 | HHHHHHHHHHHHHHHHHHHHHHHHCCCCCCC |
| 933 | EEEERKQQQKRSNEQEDKRYQREGRGKGHQP | -48.78 | 38.7 | HHHHHHHHHHHHHHHHHHHHHHHHCCCCCCC |
| 041 | DEERKKWEDKKRNEEEDKRYQRYGKGKGHSH | -48.78 | 41.9 | CHHHHHHHHHHHHHHHHHHHHHHHCCCCCCC |
| 047 | EEEQKKYEDKRNNEEEDKSYQRHGKGKGHSH | -48.77 | 48.4 | CHHHHHHHHHHHHHHHHHHHHHHCCCCCCCC |
| 436 | EEEQRKLLDQINNDAEDRRYQREGRGKGHQP | -48.76 | 45.2 | HHHHHHHHHHHHHHHHHHHHHHHHCCCCCCC |
| 028 | EEQRRKQLDMINNRAEDQRYQRYGRGKGHQP | -48.76 | 48.4 | HHHHHHHHHHHHHHHHHHHHHHHHCCCCCCC |
| 598 | EEEERKLQDQKKNEAEDQRYQREGRGKGHQP | -48.75 | 41.9 | CHHHHHHHHHHHHHHHHHHHHHHHCCCCCCC |
| 519 | EEEKRKLLDQINNEQEDQRYQREGKGKGHNH | -48.75 | 45.2 | HHHHHHHHHHHHHHHHHHHHHHHHCCCCCCC |
| 557 | EEEQRKLEDQKKNEQEDQRYQRYGRGKGHQP | -48.75 | 38.7 | HHHHHHHHHHHHHHHHHHHHHHHHCCCCCCC |
| 225 | EEEQRKLEDQKSNEKEDEEYQRRGKGKGHNH | -48.75 | 38.7 | HHHHHHHHHHHHHHHHHHHHHHHHCCCCCCC |
| 757 | EEEQRKLYDQINNEKEDEEYQRYGWGKGHNP | -48.75 | 41.9 | HHHHHHHHHHHHHHHHHHHHHHHCCCCCCCC |
| 244 | EEEQRKQEEQRSQRKEDERYQREGQGKGHQS | -48.75 | 32.3 | HHHHHHHHHHHHHHHHHHHHHHHHCCCCCCC |
| 738 | EEEERKKQDQQNNEREDEEYQRKGRGKGHNP | -48.74 | 41.9 | HHHHHHHHHHHHHHHHHHHHHHHCCCCCCCC |
| 455 | EEEERKKQDQYNNEAEDQRYQREGKGKGHSH | -48.73 | 45.2 | HHHHHHHHHHHHHHHHHHHHHHHCCCCCCCC |
| 322 | EEEEKKKRDQQNNEREDEEYQRRGRGKGHQP | -48.72 | 45.2 | CHHHHHHHHHHHHHHHHHHHHHHHCCCCCCC |
| 746 | EEEERKKQEKQSNEREDEEYQRKGRGKGHQP | -48.72 | 38.7 | HHHHHHHHHHHHHHHHHHHHHHHHCCCCCCC |
| 848 | EEEKRKQEEKRNNEEEDKRYQREGKGKGHNN | -48.69 | 41.9 | HHHHHHHHHHHHHHHHHHHHHHHCCCCCCCC |
| 938 | DEERKKSEDQYNNEAEDERYQQQGKGKGHNH | -48.69 | 45.2 | CHHHHHHHHHHHHHHHHHHHHHHHCCCCCCC |
| 274 | EEERKKQLEKYNNESEDKRYQRYGRGKGHQP | -48.68 | 48.4 | HHHHHHHHHHHHHHHHHHHHHHHHCCCCCCC |
| 742 | EEEERKKQDKYKNEYEDKEYQRKGRGKGHQP | -48.65 | 41.9 | HHHHHHHHHHHHHHHHHHHHHHHCCCCCCCC |
| 555 | EEEERKQRQKYNNEWEDKRYQREGRGKGHQP | -48.65 | 41.9 | CHHHHHHHHHHHHHHHHHHHHHHHCCCCCCC |
| 696 | EEERRKQLQKYKNEYEDEEYQRKGRGKGHQP | -48.65 | 41.9 | HHHHHHHHHHHHHHHHHHHHHHHHCCCCCCC |
| 970 | EEERRKQLQKYNNESEDKRYQRYGRGKGHQP | -48.65 | 45.2 | HHHHHHHHHHHHHHHHHHHHHHHHCCCCCCC |
| 745 | EEERRKQQQKYSNEYEDEEYQRKGRGKGHQP | -48.65 | 38.7 | HHHHHHHHHHHHHHHHHHHHHHHHCCCCCCC |
| 975 | EQEERKKQDKIRHEEEDKRYQQQGKGKGHSH | -48.63 | 41.9 | HHHHHHHHHHHHHHHHHHHHHHHCCCCCCCC |
| 478 | EEEQRKQQQQYSNDAEDKQYQQRGRGKGHQP | -48.61 | 35.5 | HHHHHHHHHHHHHHHHHHHHHHHHCCCCCCC |
| 945 | EQEQRKQQEKYNNEYEDKQYQKEGKGKGHSS | -48.61 | 38.7 | HHHHHHHHHHHHHHHHHHHHHHCCCCCCCCC |
| 184 | EEEEKKKRDQFNNDFEDRYYQQQGQGKGHQS | -48.61 | 45.2 | CHHHHHHHHHHHHHHHHHHHHHHHCCCCCCC |
| 988 | EEEERKKQDQQNNDEEDRRYQRYGWGKGHYP | -48.61 | 38.7 | HHHHHHHHHHHHCHHHHHHHHHHCCCCCCCC |
| 011 | EEQQKKQQDQRSNEKEHEEYQRRGQGQGHQS | -48.59 | 38.7 | HHHHHHHHHHHHHHHHHHHHHHHHHCCCCCC |
| 656 | EEEERKYQDQYKQEWEDKKYQREGKGKGHNH | -48.59 | 38.7 | CHHHHHHHHHHHHHHHHHHHHHHCCCCCCCC |

| | | | | |
|---|---|---|---|---|
| 969 | EEEQRKYEDQRNQEKEDERYQQQGWGKGHNP | -48.59 | 41.9 | CHHHHHHHHHHHHHHHHHHHHHHCCCCCCCC |
| 303 | EEERKKQLEQYKQEWEDQRYQREGQGKGHQS | -48.58 | 41.9 | HHHHHHHHHHHHHHHHHHHHHHHHCCCCCCC |
| 607 | EEEERKMRDQYNNDWEDQRYQREGKGKGHNH | -48.57 | 38.7 | HHHHHHHHHHHHCHHHHHHHHHHCCCCCCCC |
| 353 | EEEERKKQDQYNNEYEDEEYQRKGRGKGHQP | -48.53 | 41.9 | HHHHHHHHHHHHHHHHHHHHHHHHCCCCCCC |
| 734 | EEEERKKQDQYNNEYEDQEYQRNGRGKGHQY | -48.53 | 41.9 | HHHHHHHHHHHHHHHHHHHHHHHCCCCCCCC |
| 951 | EEEEKKKREKFSNEFEDKYYQQKGQGKGHQH | -48.53 | 45.2 | CHHHHHHHHHHHHHHHHHHHHHHHCCCCCCC |
| 245 | EEQRRKQLDKINNEAEDKRYQREGKGKGHSS | -48.52 | 54.8 | HHHHHHHHHHHHHHHHHHHHHHHCCCCCCCC |
| 020 | EEEERKYQDQYNNEYEDQYFQKKGQGKGKNH | -48.52 | 38.7 | CHHHHHHHHHHHHHHHHHHHHHHHCCCCCCC |
| 339 | EEEQRKQEQKRRNEKEDKRYQRYGRGKGHQP | -48.52 | 38.7 | HHHHHHHHHHHHHHHHHHHHHHHHCCCCCCC |
| 603 | EEEKKKLLDQINNEAEDKRYQRYGQGKGHQS | -48.52 | 51.6 | HHHHHHHHHHHHHHHHHHHHHHHHCCCCCCC |
| 770 | EQEERKKQDQYNQEYEDEEYQRKGRGKGHQP | -48.51 | 38.7 | HHHHHHHHHHHHHHHHHHHHHHHHCCCCCCC |
| 991 | DEERKKMLDKKNNEEEDERYQQQGRGKGHQP | -48.51 | 48.4 | HHHHHHHHHHHHHHHHHHHHHHHHCCCCCCC |
| 652 | EEERRKQLDQWKNEYEDQSYQREGRGKGNQN | -48.51 | 41.9 | HHHHHHHHHHHHHHHHHHHHHHHCCCCCCCC |
| 318 | EEEQKKQEEQYRNDYEDREYQRKGRGKGHNP | -48.49 | 35.5 | HHHHHHHHHHHHHHHHHHHHHHCCCCCCCCC |
| 139 | EEEERKQQEQRSNDQEDRRYQKEGRGKGHQP | -48.49 | 32.3 | HHHHHHHHHHHHHHHHHHHHHHHHCCCCCCC |
| 434 | EEEQRKQEEQRRNDQEDRRYQQYGRGKGHQP | -48.49 | 32.3 | HHHHHHHHHHHHHHHHHHHHHHHHCCCCCCC |
| 522 | EQEERKQQEQRSNRQEDERYQREGRGKGHQP | -48.49 | 29.0 | HHHHHHHHHHHHHHHHHHHHHHHHCCCCCCC |
| 584 | EEERRKQLDKIKNEQEDKSYQRYGQGKGHQS | -48.48 | 45.2 | HHHHHHHHHHHHHHHHHHHHHHHHCCCCCCC |
| 691 | EEERRKQLDKKNNEEEDKRYQQQGRGKGHQP | -48.48 | 48.4 | HHHHHHHHHHHHHHHHHHHHHHHHCCCCCCC |
| 301 | EEEERKKQDKYNNESEDRRYQRYGRGKGHQP | -48.48 | 45.2 | HHHHHHHHHHHCCHHHHHHHHHHHCCCCCCC |
| 561 | EEEQKKQQDKYSNEYEQKQYQKEGQGQGHSS | -48.47 | 38.7 | HHHHHHHHHHHHHHHHHHHHHHHCCCCCCCC |
| 774 | EEEQKKQRDKFNNEFEQKYYQQKGQGQGHNH | -48.47 | 45.2 | HHHHHHHHHHHHHHHHHHHHHHHHCCCCCCC |
| 521 | EEEQKKQRDKYNNEWEQKRYQREGKGQGHNH | -48.47 | 41.9 | HHHHHHHHHHHHHHHHHHHHHHHHCCCCCCC |
| 689 | EEEQRKQQDKFNNEKEQQRYQQQGRGQGHQP | -48.47 | 41.9 | HHHHHHHHHHHHHHHHHHHHHHHHCCCCCCC |
| 875 | EEEQRKQQDKYSNEYEQKEYQKEGRGQGHQP | -48.47 | 35.5 | HHHHHHHHHHHHHHHHHHHHHHHHCCCCCCC |
| 737 | EEEERKYQDQYNNEYEHEEYQRKGRGRGHQP | -48.45 | 35.5 | CHHHHHHHHHHHHHHHHHHHHHHHCCCCCCC |
| 874 | EQEQRKQQDHYNNRWEDEEYQRKGRGKGHQP | -48.45 | 35.5 | HHHHHHHHHHHHCHHHHHHHHHHHCCCCCCC |
| 747 | EEEQRKRQDQYNNDWEDEEYQRKGRGKGHNP | -48.44 | 38.7 | HHHHHHHHHHHCCCHHHHHHHHHCCCCCCCC |
| 782 | EEEERKFQDQWKNEFEDEYFQRKGQGKGKSH | -48.42 | 38.7 | CHHHHHHHHHHHHHHHHHHHHHHHCCCCCCC |
| 075 | EQQQRKQQEQYSNEYEDEEYQRKGRGKGHNP | -48.42 | 35.5 | HHHHHHHHHHHHHHHHHHHHHHHCCCCCCCC |
| 405 | EEEQRKQQEQFSNEYEDQYYQQQGQGKGHNH | -48.42 | 38.7 | HHHHHHHHHHHHHHHHHHHHHHHHCCCCCCC |
| 432 | EEEEKKQQEQRSNEKEDEEYQRRGQGKGHQS | -48.41 | 38.7 | HHHHHHHHHHHHHHHHHHHHHHHHHCCCCCC |
| 567 | EEEERKQQEQRSNEQEDKRYQREGKGKGHNH | -48.41 | 35.5 | HHHHHHHHHHHHHHHHHHHHHHHCCCCCCCC |
| 792 | EEEERKKRDQYNNDYEHEEYQRKGRGQGHQP | -48.41 | 32.3 | HHHHHHHHHHHCHHHHHHHHHHHHCCCCCCC |
| 189 | EEEEKKKRDQYNNEFEDEYYQQKGQGKGHQH | -48.41 | 45.2 | CHHHHHHHHHHHHHHHHHHHHHHHCCCCCCC |
| 488 | EEEERKQRDKESNEKEDKRYQREGWGKGHYP | -48.41 | 41.9 | HHHHHHHHHHHHHHHHHHHHHHHHCCCCCCC |
| 818 | EEEERKYQDQYKNEWEDRRYQREGNGKGHSH | -48.39 | 38.7 | CHHHHHHHHHHHHHHHHHHHHHCCCCCCCCC |
| 928 | EEEQKKQEQKYRNEWEDKKYQREGRGKGHQP | -48.39 | 41.9 | HHHHHHHHHHHHHHHHHHHHHHHHCCCCCCC |

| | | | | |
|---|---|---|---|---|
| 626 | EEEQRKQQDQYNNDWEQQRYQREGRGQGHQP | -48.35 | 32.3 | HHHHHHHHHHHHHHHHHHHHHHHHCCCCCCC |
| 355 | EEEQRKQQDQYNNDYEQEEYQQQGRGQGHQP | -48.35 | 32.3 | HHHHHHHHHHHHHHHHHHHHHHHHCCCCCCC |
| 670 | EEEERKKQDQFNNEFENKYYQQKGQGKGHQH | -48.34 | 41.9 | HHHHHHHHHHHHHHHHHHHHHHHHCCCCCCC |
| 879 | EEEERKQRDKYNNESEQKRYQRYGRGQGHQP | -48.34 | 38.7 | HHHHHHHHHHHCHHHHHHHHHHHHHCCCCCC |
| 862 | EEERRKQLDKWKNEWEQKYYQQKGQGQGHSH | -48.34 | 38.7 | HHHHHHHHHHHHHHHHHHHHHHHHCCCCCCC |
| 329 | EEERRKQLDKYKNEYEQKEYQRKGRGQGHQP | -48.34 | 38.7 | HHHHHHHHHHHHHHHHHHHHHHHHCCCCCCC |
| 520 | EEEERKMRDQYNNDYEDEEYQRKGRGKGHSN | -48.33 | 38.7 | HHHHHHHHHHHCCCHHHHHHHHHCCCCCCCC |
| 002 | EEEERKYQDQRNNDKEDREYQRRGRGKGHQP | -48.30 | 38.7 | CHHHHHHHHHHHCCHHHHHHHHHHCCCCCCC |
| 129 | EEEERKYQDQYNNDSEDRRYQRYGRGKGHQP | -48.30 | 38.7 | CHHHHHHHHHHCCCHHHHHHHHHHCCCCCCC |
| 464 | EEEERKQQEQRNNRKEDERYQRYGRGKGHQP | -48.29 | 35.5 | HHHHHHHHHHHHHHHHHHHHHHHHCCCCCCC |
| 801 | EEEEKKQREQYNNESEDERYQKQGRGKGHQP | -48.29 | 41.9 | CHHHHHHHHHHHCCHHHHHHHHHCCCCCCCC |
| 954 | EEEERKQQEQYSNEWEDERYQQQGWGKGHYP | -48.29 | 35.5 | HHHHHHHHHHHHHHHHHHHHHHHHCCCCCCC |
| 570 | EEEERKQREQFNNEFEDKYYQQKGQGKGHQP | -48.29 | 41.9 | HHHHHHHHHHHHHHHHHHHHHHHHCCCCCCC |
| 044 | EEERKKQLEQWKNEWEDQYYQKEGQGKGHQH | -48.29 | 41.9 | HHHHHHHHHHHHHHHHHHHHHHHHCCCCCCC |
| 230 | EEERKKQLEQYKNEAEDKRYQQYGWGKGHNP | -48.29 | 45.2 | HHHHHHHHHHHHHHHHHHHHHHHCCCCCCCC |
| 146 | EEERKKQLEQYNNESEDQRYQRYGRGKGHQP | -48.29 | 45.2 | HHHHHHHHHHHHHHHHHHHHHHHHCCCCCCC |
| 985 | EEERRKQEEQQRNEEEDQRYQRYGRGKGHQP | -48.29 | 35.5 | HHHHHHHHHHHHHHHHHHHHHHHHCCCCCCC |
| 832 | EEERRKQLEQYNNEYEDEEYQRKGRGKGHQP | -48.29 | 41.9 | HHHHHHHHHHHHHHHHHHHHHHHHCCCCCCC |
| 732 | EEERRKQLEQYSNEYEDQEYQRVGRGKGHQP | -48.29 | 38.7 | HHHHHHHHHHHHHHHHHHHHHHHHCCCCCCC |
| 780 | EQEERKQREQYNNEAEDQRYQREGKGKGHSH | -48.29 | 38.7 | HHHHHHHHHHHHHHHHHHHHHHHCCCCCCCC |
| 417 | EEEEKKYQDQYNNESEDQRYQRYGRGKGHQP | -48.28 | 45.2 | CHHHHHHHHHHCCCHHHHHHHHHHCCCCCCC |
| 157 | EEEERKYQDQKNNEEEDQRYQRYGRGKGHQP | -48.28 | 41.9 | CHHHHHHHHHHHHHHHHHHHHHHHCCCCCCC |
| 415 | EEEERKYQDQYKNEWEDKRYQREGRGKGHQP | -48.28 | 38.7 | CHHHHHHHHHHHHHHHHHHHHHHHCCCCCCC |
| 600 | EEEERKYQDQYKNEYEDQYYQKKGQGKGHSH | -48.28 | 38.7 | CHHHHHHHHHHHHHHHHHHHHHHHCCCCCCC |
| 421 | EEEERKYQDQYNNESEDKRYQKEGWGKGHYP | -48.28 | 41.9 | CHHHHHHHHHHCCHHHHHHHHHHHCCCCCCC |
| 698 | EEEERKYQDQYNNEYEDQEYQRNGRGKGHQY | -48.28 | 41.9 | CHHHHHHHHHHHHHHHHHHHHHCCCCCCCCC |
| 984 | EEEQKKYEDQRNNEEEDEEYQRKGRGKGHQP | -48.28 | 45.2 | CHHHHHHHHHHCHHHHHHHHHHHHCCCCCCC |
| 341 | EEEQRKYEDQRNNEAEDQRYQRQGWGKGHNP | -48.28 | 45.2 | CHHHHHHHHHHHHHHHHHHHHHHHCCCCCCC |
| 587 | EEEQRKYEDQRNNEEEDEEYQRKGRGKGHQP | -48.28 | 41.9 | CHHHHHHHHHHCHHHHHHHHHHHHCCCCCCC |
| 286 | EEEQRKYEDQRNNEEEDERYQQQGKGKGHSS | -48.28 | 41.9 | CHHHHHHHHHHHHHHHHHHHHHHCCCCCCCC |
| 290 | EEEQRKYEDQRNNEEEDKRYQREGKGKGHSS | -48.28 | 41.9 | CHHHHHHHHHHHHHHHHHHHHHHCCCCCCCC |
| 790 | EEEQRKYEDQRNNEEEDQQYQKQGRGKGHQP | -48.28 | 41.9 | CHHHHHHHHHHHHHHHHHHHHHHHCCCCCCC |
| 027 | EEEQRKYEDQRNNEEEDQRYQREGWGKGHNP | -48.28 | 41.9 | CHHHHHHHHHHHHHHHHHHHHHHHCCCCCCC |
| 198 | EEEQRKYEDQRNNEEEDQRYQRYGRGKGHQP | -48.28 | 41.9 | CHHHHHHHHHHHHHHHHHHHHHHHCCCCCCC |
| 784 | KQEEEKYQDQYNNEWEDEEYQRKGRGKGHQP | -48.28 | 35.5 | CHHHHHHHHHHHHHHHHHHHHHHHCCCCCCC |
| 841 | EEEERKRQDQENNDKEDEEYQRRGWGKGHNP | -48.26 | 38.7 | HHHHHHHHHHHHCHHHHHHHHHHCCCCCCCC |
| 273 | EEQARKSQDQYNNEHEDERYQQQGRGKGHQP | -48.25 | 48.4 | HHHHHHHHHHHHHHHHHHHHHHHHCCCCCCC |
| 036 | EEERKKQLDKESNEKEDKRYQREGQGKGHQS | -48.25 | 48.4 | HHHHHHHHHHHHHHHHHHHHHHHHCCCCCCC |

| | | | | |
|---|---|---|---|---|
| 496 | EEEERKFQQQYSEEYEDEEYQRKGRGKGHQP | -48.25 | 38.7 | CHHHHHHHHHHHHHHHHHHHHHHHCCCCCCC |
| 887 | EEERRKQLDKKNNEEEDKRYQQQGKGKGHNH | -48.25 | 48.4 | HHHHHHHHHHHHHHHHHHHHHHHCCCCCCCC |
| 121 | EEEQRKLYDQINNRKEDEEYQRAGRGKGHQP | -48.24 | 38.7 | HHHHHHHHHHHHHHHHHHHHHHHHHCCCCCC |
| 115 | EEEQRKQQEQRSNDAEDRQYQQRGRGKGHQP | -48.23 | 35.5 | HHHHHHHHHHHHCHHHHHHHHHHHCCCCCCC |
| 606 | EQEERKQQEQRSNEEEDKRYQREGWGKGHYP | -48.22 | 32.3 | HHHHHHHHHHHHHHHHHHHHHHHHCCCCCCC |
| 012 | EQEERKKQEKYNQEAEDKRYQRYGRGKGHQP | -48.22 | 41.9 | HHHHHHHHHHHHHHHHHHHHHHHHCCCCCCC |
| 477 | EEEERKREDKYRNEYEDEYYQQQGQGKGHQP | -48.20 | 41.9 | HHHHHHHHHHHHHHHHHHHHHHHHCCCCCCC |
| 213 | EEEERKRQDKENNEREDKRYQREGQGKGHQS | -48.20 | 45.2 | HHHHHHHHHHHHHHHHHHHHHHHCCCCCCCC |
| 924 | EEEERKRQDKQNNESEDKSYQREGRGKGHQP | -48.20 | 45.2 | HHHHHHHHHHHHHHHHHHHHHHHCCCCCCCC |
| 905 | EEEERKKQDQYNNEYEDEEYQRKGRGKGDQR | -48.19 | 48.4 | HHHHHHHHHHHHHHHHHHHHHHHCCCCCCCC |
| 081 | EEEEKKFRDQYNNEYEDQEYQRKGRGKGHQP | -48.19 | 48.4 | CHHHHHHHHHHHHHHHHHHHHHHHCCCCCCC |
| 195 | EEEERKFRDQYNNEYEDQQYQKEGRGKGHQP | -48.19 | 45.2 | CHHHHHHHHHHHHHHHHHHHHHHHCCCCCCC |
| 548 | EEEKKKFLDQYNNEFEDKMYQRYGKGKGHQP | -48.19 | 51.6 | HHHHHHHHHHHHHHHHHHHHHHHHHCCCCCC |
| 338 | EEEKRKFLDQYKNEYEDEEYQRKGRGKGHQP | -48.19 | 45.2 | HHHHHHHHHHHHHHHHHHHHHHHHCCCCCCC |
| 638 | EEEQRKFEDQRNNEEEDKRYQREGRGKGHQP | -48.19 | 45.2 | CHHHHHHHHHHHHHHHHHHHHHHHCCCCCCC |
| 917 | EEEQKKQRDQFNNEWEHKRKQREGRGSGKQN | -48.18 | 38.7 | HHHHHHHHHHHHHHHHHHHHHHHCCCCCCCC |
| 718 | DEERKKQEEQYRNDYEDREYQRKGRGKGHQP | -48.17 | 32.3 | HHHHHHHHHHHHHHHHHHHHHHHHCCCCCCC |
| 294 | EEEQKKQRQQYNNDAEDRRFQRYGRGKGKQN | -48.16 | 38.7 | HHHHHHHHHHHHHHHHHHHHHHHCCCCCCCC |
| 685 | EEEQRKQQDQYNNEWEDKRYQREGKGKGDNR | -48.13 | 48.4 | HHHHHHHHHHHHHHHHHHHHHHCCCCCCCCC |
| 501 | EEERRKQLQVESERKEDQRYQRYGWGKGHYP | -48.13 | 35.5 | HHHHHHHHHHHHHHHHHHHHHHHCCCCCCCC |
| 931 | EEEQKKFRDQFNNEFEDEFYQQQGRGKGHQP | -48.12 | 54.8 | HHHHHHHHHHHHHHHHHHHHHHHHCCCCCCC |
| 986 | EEEQKKFRDQYNNEWEDEEYQRKGRGKGHQP | -48.12 | 48.4 | HHHHHHHHHHHHCHHHHHHHHHHHCCCCCCC |
| 263 | EEEQRKFQDQYNNEEEDKRYQREGKGKGHSS | -48.12 | 45.2 | HHHHHHHHHHHHHHHHHHHHHHHCCCCCCCC |
| 425 | EEEERKKQDQIENEREDSRYQQQGWGKGHYP | -48.10 | 38.7 | HHHHHHHHHHHHHHHHHHHHHHHCCCCCCCC |
| 664 | DEERKKMLDQKNNEEEDQRFQREGKGKGKNN | -48.10 | 41.9 | HHHHHHHHHHCCHHHHHHHHHHHCCCCCCCC |
| 617 | EEEKKKMLDQFNQEFEDQYYQQKGQGKGHNH | -48.09 | 51.6 | HHHHHHHHHHHHHHHHHHHHHHHHCCCCCCC |
| 404 | QEEEKKMRDQYNQEWEDKRYQREGKGKGHNH | -48.09 | 41.9 | CHHHHHHHHHHHHHHHHHHHHHCCCCCCCCC |
| 247 | EEEARKQQEQYSNEWEDKRYQREGRGKGHQP | -48.09 | 38.7 | HHHHHHHHHHHHHHHHHHHHHHHHCCCCCCC |
| 534 | EEEWKKQMEQKNNEEEDKRYQREGRGKGHQP | -48.09 | 41.9 | HHHHHHHHHHHHHHHHHHHHHHHHCCCCCCC |
| 122 | EEEQRKQQDQYNNEYEHQEYQKEGRGQGHQP | -48.07 | 35.5 | HHHHHHHHHHHHHHHHHHHHHHHHCCCCCCC |
| 217 | EEEERKQQQKRSNEKEDQRYQQQGWGKGHNP | -48.07 | 38.7 | HHHHHHHHHHHHHHHHHHHHHHHCCCCCCCC |
| 051 | EEEERKWRDQKNNEEEDERYQRYGWGKGHNP | -48.06 | 41.9 | CHHHHHHHHHHCHHHHHHHHHHHCCCCCCCC |
| 372 | EEEYRKWQDQKNNEEEDERYQQQGRGKGHQP | -48.06 | 41.9 | HHHHHHHHHHHHHHHHHHHHHHHHCCCCCCC |
| 812 | EEEWRKWIDQLKNEKEDKRYQREGRGKGHQP | -48.05 | 38.7 | HHHHHHHHHHHHHHHHHHHHHHHHCCCCCCC |
| 337 | EEERKKQLDQWKNRWEDEYFQQQGRGKGKQN | -48.05 | 38.7 | HHHHHHHHHHHHHHHHHHHHHHHHCCCCCCC |
| 224 | EEEKKKMLDQINNEKEDEYFQQYGQGKGKSH | -48.03 | 45.2 | HHHHHHHHHHHHHHHHHHHHHHHHHCCCCCC |
| 741 | EEEQRKYEDQRNNREEDERYQRYGRGKGHQP | -48.00 | 38.7 | CHHHHHHHHHHHHHHHHHHHHHHHHCCCCCC |
| 813 | EEEQKKYEDQRNQEEEDKRYQREGQGKGHQS | -47.99 | 45.2 | CHHHHHHHHHHHHHHHHHHHHHHHCCCCCCC |

| | | | | |
|---|---|---|---|---|
| 424 | EEEERKKMQKMEQEREDKEYQQKGRGKGHQP | -47.98 | 38.7 | CHHHHHHHHHHHHHHHHHHHHHHHCCCCCCC |
| 031 | EEERRKQEEQYRQEAEDQRYQREGWGKGHYP | -47.97 | 38.7 | HHHHHHHHHHHHHHHHHHHHHHHHCCCCCCC |
| 287 | EEEQRKQQDKYNQESEQKRYQRYGQGQGHQS | -47.97 | 38.7 | HHHHHHHHHHHHHHHHHHHHHHHHHCCCCCC |
| 229 | EEEERKKQDKYNNEFEHKYYQQKGQGQGHSH | -47.96 | 38.7 | HHHHHHHHHHHHHHHHHHHHHHHHCCCCCCC |
| 665 | EEEERKQRQQYNNEFEDQYFQQKGQGKGKQN | -47.96 | 35.5 | HHHHHHHHHHHHHHHHHHHHHHHHCCCCCCC |
| 486 | EEERRKQLEQQSNDREDRYYQREGKGKGHSH | -47.95 | 35.5 | HHHHHHHHHHHHHHHHHHHHHHCCCCCCCCC |
| 066 | EEEEKKKQDQINNDKEDREYQRYGKGKGHSS | -47.95 | 41.9 | CHHHHHHHHHHHHHHHHHHHHHHCCCCCCCC |
| 413 | EEEERKKQDQWKNDWEDRFYQKEGKGKGHNH | -47.95 | 38.7 | HHHHHHHHHHHHHHHHHHHHHHCCCCCCCCC |
| 877 | EEQQRKQKQQFDNEKEHQRYQREGQGRGHQS | -47.95 | 35.5 | HHHHHHHHHHHCHHHHHHHHHHHHCCCCCCC |
| 387 | EEEKRKFLDQYNNESEDERYQQQGRGKGHQP | -47.95 | 48.4 | HHHHHHHHHHHCCCHHHHHHHHHHCCCCCCC |
| 127 | EEEERKQQQQYKQEWEDEEYQRKGRGKGHQP | -47.94 | 35.5 | HHHHHHHHHHHHHHHHHHHHHHHHCCCCCCC |
| 352 | EEEEKKKQDQEKNEREDKRYQRYGRGKGHQP | -47.94 | 41.9 | CHHHHHHHHHHHHHHHHHHHHHHHCCCCCCC |
| 717 | EEEEKKKQDQIKNEKEDEEYQRRGRGKGHQP | -47.94 | 41.9 | CHHHHHHHHHHHHHHHHHHHHHHHCCCCCCC |
| 112 | EEEERKKQDQIKNEEEDKKYQREGRGKGHQP | -47.94 | 38.7 | HHHHHHHHHHHHHHHHHHHHHHHHCCCCCCC |
| 363 | EEEERKKQDQIKNEEEDQRYQQYGRGKGHQP | -47.94 | 38.7 | HHHHHHHHHHHHHHHHHHHHHHHHHCCCCCC |
| 727 | EEEERKKQDQIKNEKEDEEYQRYGKGKGHSS | -47.94 | 38.7 | HHHHHHHHHHHHHHHHHHHHHHHHCCCCCCC |
| 644 | EEEERKKQDQIKNEKEDQEYQKQGKGKGHSH | -47.94 | 38.7 | HHHHHHHHHHHHHHHHHHHHHHHCCCCCCCC |
| 147 | EEEERKKQDQINNEAEDKRYQKEGKGKGHNH | -47.94 | 45.2 | HHHHHHHHHHHHHHHHHHHHHHHCCCCCCCC |
| 285 | EEEWKKKMDQFNNEKEDKRYQQQGWGKGHNP | -47.94 | 48.4 | HHHHHHHHHHHHHHHHHHHHHHHCCCCCCCC |
| 989 | EEEERKKQDQLNNRKEDERYQREGRGKGHQP | -47.92 | 38.7 | HHHHHHHHHHHHHHHHHHHHHHHHCCCCCCC |
| 783 | EEEERKRQDQEKNEKEDKRYQREGKGKGHNH | -47.91 | 38.7 | HHHHHHHHHHHHHHHHHHHHHHHCCCCCCCC |
| 516 | EEEERKRQDQEKNEKEDKRYQRYGRGKGHQP | -47.91 | 38.7 | HHHHHHHHHHHHHHHHHHHHHHHHCCCCCCC |
| 091 | EEEERKRQDQEKNEKEDQRYQREGKGKGHSH | -47.91 | 38.7 | HHHHHHHHHHHHHHHHHHHHHHHCCCCCCCC |
| 550 | EEEEKKQQEQRNNEKEQEEYQRRGRGRGHQP | -47.86 | 35.5 | HHHHHHHHHHHHHHHHHHHHHHHHCCCCCCC |
| 601 | EEEERKQQQQRNNDREDREYQQRGRGKGHQP | -47.85 | 35.5 | HHHHHHHHHHHHHHHHHHHHHHHHCCCCCCC |
| 089 | EEEQRKQEQQRNNDKEDRRYQREGKGKGHNH | -47.85 | 35.5 | HHHHHHHHHHHHHHHHHHHHHHHCCCCCCCC |
| 660 | EEEQRKQEQQRNNDKEDRRYQRYGRGKGHQP | -47.85 | 35.5 | HHHHHHHHHHHHHHHHHHHHHHHHCCCCCCC |
| 220 | EEEYRKWLDQKKNEEEDQRYQKQGRGKGHQP | -47.83 | 41.9 | HHHHHHHHHHHHHHHHHHHHHHHCCCCCCCC |
| 469 | EEEQRKQQDQYSQEYEQQEYQKEGRGQGHQP | -47.82 | 32.3 | HHHHHHHHHHHHHHHHHHHHHHHHCCCCCCC |
| 289 | DEERKKMLDQIKNEQEDQRYQREGRGKGHQP | -47.79 | 41.9 | HHHHHHHHHHHHHHHHHHHHHHHHCCCCCCC |
| 484 | DEERKKMLDQYKNEYEDEYYQQNGSGKGHQY | -47.79 | 41.9 | HHHHHHHHHHHHHHHHHHHHHHHCCCCCCCC |
| 199 | EEEERKMRDQYNNEAEDERYQRKGMGKGHQP | -47.79 | 45.2 | HHHHHHHHHHHHHHHHHHHHHHHHHCCCCCC |
| 923 | EEEERKMRDQYNNEFEDQYYQKEGQGKGHSH | -47.79 | 41.9 | HHHHHHHHHHHHHHHHHHHHHHHCCCCCCCC |
| 766 | EEEKKKMLDQMKNEKEDEEYQRYGRGKGHQP | -47.79 | 45.2 | HHHHHHHHHHHHHHHHHHHHHHHHHCCCCCC |
| 978 | EEEKKKMLDQWKNEWEDQFYQKQGRGKGHQP | -47.79 | 48.4 | HHHHHHHHHHHHHHHHHHHHHHHHCCCCCCC |
| 608 | EEEKKKMLDQYKNEWEDEEYQRKGRGKGHQP | -47.79 | 45.2 | HHHHHHHHHHHHHHHHHHHHHHHHCCCCCCC |
| 722 | EEEKKKMLDQYKNEYEDEQYQQQGQGKGHQS | -47.79 | 45.2 | HHHHHHHHHHHHHHHHHHHHHHHHCCCCCCC |
| 997 | EEEKRKMLDQIKNEKEDQEYQKQGRGKGHQP | -47.79 | 41.9 | HHHHHHHHHHHHHHHHHHHHHHHHCCCCCCC |

| | | | | |
|---|---|---|---|---|
| 960 | EEEKRKMLDQMKNEKEDEEYQRRGKGKGHSH | -47.79 | 41.9 | HHHHHHHHHHHHHHHHHHHHHHHHCCCCCCC |
| 639 | EEEKRKMLDQMKNEKEDKEYQKQGRGKGHQP | -47.79 | 41.9 | HHHHHHHHHHHHHHHHHHHHHHHHCCCCCCC |
| 647 | EEEKRKMLDQMKNEKEDQEYQKQGRGKGHQP | -47.79 | 41.9 | HHHHHHHHHHHHHHHHHHHHHHHHCCCCCCC |
| 008 | EEEKRKMLDQYNNESEDKRYQRYGRGKGHQP | -47.79 | 45.2 | HHHHHHHHHHHHHHHHHHHHHHHHCCCCCCC |
| 553 | EEERRKQLDQIKNDKEDRDYQREGKGKGHNH | -47.78 | 38.7 | HHHHHHHHHHHHHHHHHHHHHHHCCCCCCCC |
| 064 | EEERRKQLDQIKNDQEDRRYQREGRGKGHQP | -47.78 | 38.7 | HHHHHHHHHHHHHHHHHHHHHHHHCCCCCCC |
| 514 | EEERRKQLDQKNNDEEDRRYQRYGQGKGHQS | -47.78 | 41.9 | HHHHHHHHHHHCCHHHHHHHHHHHHCCCCCC |
| 246 | EEERKKQLDQLKNRQEDERYQREGKGKGHNH | -47.78 | 41.9 | HHHHHHHHHHHHHHHHHHHHHHHCCCCCCCC |
| 545 | EEEQRKQEQQYRNEWEDERYQRQGRGKGHQP | -47.78 | 35.5 | HHHHHHHHHHHHHHHHHHHHHHHHCCCCCCC |
| 908 | EEEQRKQQQQYSNEWEDEEYQRKGRGKGHNP | -47.78 | 35.5 | HHHHHHHHHHHHHHHHHHHHHHHHCCCCCCC |
| 113 | EEEQRKQEQQRNNEKEDKRYQREGKGKGHSS | -47.78 | 38.7 | HHHHHHHHHHHHHHHHHHHHHHHCCCCCCCC |
| 192 | EEEEKKKRDAKNNREEDERYQQQGRGKGHQP | -47.77 | 41.9 | CHHHHHHHHHHHHHHHHHHHHHHHCCCCCCC |
| 005 | EEEERKKQDAYNNRAEDQRYQREGWGKGHYP | -47.77 | 41.9 | HHHHHHHHHHHHHHHHHHHHHHHHCCCCCCC |
| 886 | EEEKRKQEQQRNEEAEDQRYQREGKGKGHNH | -47.77 | 41.9 | HHHHHHHHHHHHHHHHHHHHHHHHCCCCCCC |
| 752 | EEERKKQLDQINNEKEDEEYQRYGRGKGHQP | -47.77 | 48.4 | HHHHHHHHHHHHHHHHHHHHHHHHHCCCCCC |
| 293 | EEERKKQLDQYKNEWEDEEYQRKGRGKGHQP | -47.77 | 45.2 | HHHHHHHHHHHHHHHHHHHHHHHHCCCCCCC |
| 215 | EEERRKQLDQEKNEKEDERYQQQGKGKGHSS | -47.77 | 41.9 | HHHHHHHHHHHHHHHHHHHHHHHCCCCCCCC |
| 360 | EEERRKQLDQEKNEKEDQRYQREGRGKGHQP | -47.77 | 41.9 | HHHHHHHHHHHHHHHHHHHHHHHHCCCCCCC |
| 070 | EEERRKQLDQIKNEKEDEEYQRRGQGKGHQS | -47.77 | 41.9 | HHHHHHHHHHHHHHHHHHHHHHHHHCCCCCC |
| 826 | EEERRKQLDQWKNEFEDKYYQEYGQGKGHQP | -47.77 | 41.9 | HHHHHHHHHHHHHHHHHHHHHHHHHCCCCCC |
| 507 | EEERRKQLDQWKNEYEDEEYQRKGRGKGHQP | -47.77 | 41.9 | HHHHHHHHHHHHHHHHHHHHHHHHCCCCCCC |
| 077 | EEERRKQLDQYKNEYEDQEYQRQGRGKGHQP | -47.77 | 41.9 | HHHHHHHHHHHHHHHHHHHHHHHHCCCCCCC |
| 163 | EEERRKQLDQYNNEYEDEEYQRKGRGKGHNP | -47.77 | 45.2 | HHHHHHHHHHHHHHHHHHHHHHHCCCCCCCC |
| 729 | EEEERKYQDQRNNEEEDQRYQREGRGKGHQP | -47.77 | 41.9 | CHHHHHHHHHHHHHHHHHHHHHHHCCCCCCC |
| 340 | EQEERKYQDQYNNEAEDKRYQREGKGKGHSH | -47.77 | 41.9 | CHHHHHHHHHHHHHHHHHHHHHHCCCCCCCC |
| 672 | EREQRYQQDQRSNEKEDEEYQRRGQGKGNSN | -47.75 | 35.5 | HHHHHHHHHHHHHHHHHHHHHHHHCCCCCCC |
| 505 | EEEQRKLEEKKSNEQEDKRYQRYGRGKGHQP | -47.75 | 38.7 | HHHHHHHHHHHHHHHHHHHHHHHHCCCCCCC |
| 827 | EQEERKRQEQYNNDWEDKRYQREGDGKGHQP | -47.75 | 32.3 | HHHHHHHHHHHHHHHHHHHHHHCCCCCCCCC |
| 347 | EEEERKQRQQRENEREDQEYQRYGRGKGHQP | -47.75 | 35.5 | HHHHHHHHHHHHHHHHHHHHHHHHHCCCCCC |
| 571 | EEEERKQRDKFNNEKEQQRYQQQGKGQGHSS | -47.73 | 41.9 | HHHHHHHHHHHHHHHHHHHHHHHCCCCCCCC |
| 261 | EEEERKKQDQYNNEWEDEEYQRKGRGKGDQR | -47.73 | 48.4 | HHHHHHHHHHHHHHHHHHHHHHHCCCCCCCC |
| 100 | EEEQKKQQDQKNNDEEHRRYQQYGRGQGHQP | -47.73 | 35.5 | HHHHHHHHHHHCHHHHHHHHHHHHHCCCCCC |
| 535 | EEQEKKKREKYNNEFEDKYYQQKGQGKGHQP | -47.72 | 48.4 | HHHHHHHHHHHHHHHHHHHHHHHHCCCCCCC |
| 648 | EEERRKQLQQYKNRYEDEEYQRQGRGKGHQP | -47.72 | 35.5 | HHHHHHHHHHHHHHHHHHHHHHHHCCCCCCC |
| 227 | EEERRKQQEQYSNRYEDEEYQREGRGKGHSR | -47.72 | 35.5 | HHHHHHHHHHHHHHHHHHHHHHHCCCCCCCC |
| 651 | EEEERKKQDQIKNEKEDQEYQRSGKGKGHNH | -47.71 | 41.9 | HHHHHHHHHHHHHHHHHHHHHHCCCCCCCCC |
| 868 | EEEERKQRDQEKNDKEDRRYQREGKGKGHSH | -47.70 | 35.5 | HHHHHHHHHHHHHHHHHHHHHHHCCCCCCCC |
| 715 | DEERKKQEDQERNEKEDEEYQRYGWGKGHNP | -47.69 | 38.7 | CHHHHHHHHHHHHHHHHHHHHHHCCCCCCCC |

| | | | | |
|---|---|---|---|---|
| 779 | DEERKKQHDQESNEKEDKRYQREGKGKGHNH | -47.69 | 38.7 | HHHHHHHCHHHHHHHHHHHHHHHCCCCCCCC |
| 394 | EEEQRKQYDQEKNEKEDERYQQQGWGKGHNP | -47.69 | 38.7 | HHHHHHHHHHHHHHHHHHHHHHHCCCCCCCC |
| 398 | EEEERKQRQQYNNRAEDERYQREGRGKGHQP | -47.69 | 38.7 | HHHHHHHHHHHHHHHHHHHHHHHHCCCCCCC |
| 541 | EEEEKKKQDQYNNEFEDEYYQQYGQGKGDSR | -47.68 | 51.6 | CHHHHHHHHHHHHHHHHHHHHHHHHCCCCCC |
| 086 | EEERKKQLEQYKNEWEDEEYQRKGRGKGHQP | -47.68 | 41.9 | HHHHHHHHHHHHHHHHHHHHHHHHCCCCCCC |
| 866 | EEERKKQLEQYKNEWEDKRYQREGRGKGHQP | -47.68 | 41.9 | HHHHHHHHHHHHHHHHHHHHHHHHCCCCCCC |
| 093 | EEEERKQQDQRNHEKEQERYQQQGRGQGHQP | -47.67 | 38.7 | HHHHHHHHHHHHHHHHHHHHHHHHCCCCCCC |
| 846 | EEEQRKFYDQYNNEYEDQEYQRKGRGKGHQP | -47.67 | 45.2 | HHHHHHHHHHHHHHHHHHHHHHHHCCCCCCC |
| 278 | EEEQKKQRDKFNNEFEQKFYQKEGRGQGHQP | -47.67 | 48.4 | HHHHHHHHHHHHHHHHHHHHHHHHCCCCCCC |
| 461 | EEEERKNQEKYSNEWEDKRYQREGRGKGHQP | -47.66 | 38.7 | HHHHHHHHHHHHHHHHHHHHHHHHCCCCCCC |
| 744 | EEERRKQLDQWKNEWEDQYYQQQGQGKGHSS | -47.66 | 41.9 | HHHHHHHHHHHHHHHHHHHHHHHHCCCCCCC |
| 716 | EEEQRKQQDQYNNEWEHEYYQRKGQGQGHQH | -47.66 | 35.5 | HHHHHHHHHHHHHHHHHHHHHHHHCCCCCCC |
| 357 | DEQRKKMLDQYKNEYEDEEYQRQGRGKGHQP | -47.66 | 45.2 | HHHHHHHHHHHHHHHHHHHHHHHHCCCCCCC |
| 479 | EEEERKQRQQYNNEAEDQRYQREGRGKGHQP | -47.65 | 41.9 | HHHHHHHHHHHHHHHHHHHHHHHHCCCCCCC |
| 977 | EEEERKQRQQYNNESEDERYQQQGKGKGHNH | -47.65 | 38.7 | HHHHHHHHHHHCCCHHHHHHHHHHCCCCCCC |
| 579 | EEEERKQRQQYNNEYEDEEYQRKGRGKGHQP | -47.65 | 38.7 | HHHHHHHHHHHHHHHHHHHHHHHHCCCCCCC |
| 720 | EEEKRKQEQQRNNEEEDERYQRQGRGKGHQP | -47.65 | 38.7 | HHHHHHHHHHHHHHHHHHHHHHHHCCCCCCC |
| 180 | EEERRKQEQQYRNEFEDQYYQQKGQGKGHSH | -47.65 | 35.5 | HHHHHHHHHHHHHHHHHHHHHHHHCCCCCCC |
| 123 | EEERRKQLQQKNNEEEDQRYQRYGWGKGHNP | -47.65 | 41.9 | HHHHHHHHHHHHHHHHHHHHHHHCCCCCCCC |
| 257 | EEEQRKQEDKRRNEKEQKRYQREGKGQGHSH | -47.65 | 35.5 | HHHHHHHHHHHHHHHHHHHHHHHHCCCCCCC |
| 374 | EEEQRKLQDAINNRAEDQRYQREGRGKGHQP | -47.64 | 41.9 | HHHHHHHHHHHHHHHHHHHHHHHHCCCCCCC |
| 462 | EEEQRKHHEKESNEREDEEYQRKGRGKGHQP | -47.62 | 38.7 | HHHHHHHHHHHHHHHHHHHHHHHHCCCCCCC |
| 037 | EEEQRKQQQQYNHDWEDRRYQREGWGKGHNP | -47.61 | 38.7 | HHHHHHHHHHHHCHHHHHHHHHHHCCCCCCC |
| 141 | EEEQKKQQEQYSNESEDQRYQRYGQGKGHQS | -47.61 | 38.7 | HHHHHHHHHHHHHCHHHHHHHHHHHCCCCCC |
| 927 | EQEQRKQQEQYNNESEDERYQQQGRGKGHQP | -47.61 | 35.5 | HHHHHHHHHHHHHCHHHHHHHHHHCCCCCCC |
| 967 | EEEERKQQQQRNNEKEDQRYQREGKGKGHNN | -47.58 | 38.7 | HHHHHHHHHHHHHHHHHHHHHHHCCCCCCCC |
| 154 | EEEERKQQQQRNNEKEDQRYQREGKGKGHSN | -47.58 | 38.7 | HHHHHHHHHHHHHHHHHHHHHHHCCCCCCCC |
| 795 | EEEERKKQQKKNNEEEDKRYQREGWGKGHNP | -47.58 | 41.9 | HHHHHHHHHHHHHHHHHHHHHHHHCCCCCCC |
| 039 | EEEQRKYEDQRNNEEEDQRYQRYGKGKGDSR | -47.55 | 48.4 | CHHHHHHHHHHHHHHHHHHHHHHCCCCCCCC |
| 876 | EQEERKKQEQYKNEWEDKRYQREGKGKGHNH | -47.53 | 32.3 | HHHHHHHHHHHHHHHHHHHHHHCCCCCCCCC |
| 430 | EEERRKQLDQYNNESEDQRYQRYGRGKGHQP | -47.53 | 45.2 | HHHHHHHHHHHHHCHHHHHHHHHHCCCCCCC |
| 620 | EEEERKKQQKFSNEAEDEEYQRRGKGKGHSS | -47.53 | 45.2 | HHHHHHHHHHHHHHHHHHHHHHHHCCCCCCC |
| 381 | EEEQKKQRDQYNNEAEQKRYQREGKGQGHNN | -47.52 | 41.9 | HHHHHHHHHHHCHHHHHHHHHHHCCCCCCCC |
| 622 | EEEQRKQQDQYNNEAEQQRYQREGRGQGHQP | -47.52 | 38.7 | HHHHHHHHHHHHHHHHHHHHHHHHCCCCCCC |
| 940 | EEEQRKQQDQYNNEHEQERYQKQGWGQGHNP | -47.52 | 35.5 | HHHHHHHHHHHHHHHHHHHHHHHCCCCCCCC |
| 949 | EEEQRKQQDQYNNESEQKRYQKEGKGQGHSS | -47.52 | 35.5 | HHHHHHHHHHHCHHHHHHHHHHHCCCCCCCC |
| 021 | EEEQRKQQDQYNNEWEQKRYQREGKGQGHNH | -47.52 | 35.5 | HHHHHHHHHHHHHHHHHHHHHHHHCCCCCCC |
| 300 | EEEQRKQQDQYNNEYEQEEYQRQGRGQGHQP | -47.52 | 35.5 | HHHHHHHHHHHHHHHHHHHHHHHHCCCCCCC |

| | | | | |
|---|---|---|---|---|
| 631 | EEEQRKQEQKRRNEKEDRRYQKEGRGKGHQP | -47.51 | 38.7 | HHHHHHHHHHHHHHHHHHHHHHHCCCCCCCC |
| 242 | EEEKRKFLDKMKNEAEDRRYQRYGRGKGHQP | -47.51 | 51.6 | HHHHHHHHHHHHHHHHHHHHHHHHCCCCCCC |
| 150 | EEEERKRQDQYNNDSEDRRYQRYGRGKGHQP | -47.49 | 38.7 | HHHHHHHHHHCCCCHHHHHHHHHHCCCCCCC |
| 754 | EEEEKKRQDQYNNEWEDEEYQRKGRGKGHQP | -47.48 | 45.2 | CHHHHHHHHHHHHHHHHHHHHHHHCCCCCCC |
| 982 | EEEERKRQDQYNNEWEDKRYQEEGRGKGHQP | -47.48 | 41.9 | HHHHHHHHHHHHHHHHHHHHHHHHCCCCCCC |
| 391 | EEEERKRQDQYNNEWEDKRYQREGRGKGHQP | -47.48 | 41.9 | HHHHHHHHHHHHHHHHHHHHHHHHCCCCCCC |
| 619 | EQEERKRQDQYNNEWEDEEYQRKGRGKGHQP | -47.48 | 38.7 | HHHHHHHHHHHHHHHHHHHHHHHHCCCCCCC |
| 528 | EEEERKKQEQYSNDYEDEEYQRKGRGKGHQP | -47.48 | 32.3 | HHHHHHHHHHHHHHHHHHHHHHHHCCCCCCC |
| 472 | QDQQRKQQDHYNNRWEDEEYQRKGRGKGHQP | -47.47 | 35.5 | CHHHHHHHHHHHCHHHHHHHHHHHCCCCCCC |
| 513 | EEERRKQLDQQNNDEEQRRYQQYGWGQGHYP | -47.46 | 35.5 | HHHHHHHHHHHCCHHHHHHHHHHCCCCCCCC |
| 992 | EEEERKQRQQFNNEFEDQYYQQKGQGKGHSH | -47.46 | 41.9 | HHHHHHHHHHHHHHHHHHHHHHHHCCCCCCC |
| 259 | EEERRKQLDQYNNRSEQERYQKQGKGQGHSN | -47.43 | 35.5 | HHHHHHHHHHHHHHHHHHHHHHHCCCCCCCC |
| 447 | EEEERKQKQKWKNEWEDKFYQKEGKGKGDSR | -47.41 | 48.4 | CHHHHHHHHHHHHHHHHHHHHHHCCCCCCCC |
| 390 | EEEERKRQDQEKNRKEDERYQREGRGKGHQP | -47.41 | 35.5 | HHHHHHHHHHHHHHHHHHHHHHHHCCCCCCC |
| 103 | EEEKRKQEDQRNNEKEQEEYQRRGKGQGHSS | -47.40 | 35.5 | HHHHHHHHHHHHHHHHHHHHHHHHCCCCCCC |
| 304 | EEEERKQQDQYKNEWEQEYYQRKGQGQGHSH | -47.39 | 32.3 | HHHHHHHHHHHHHHHHHHHHHHHHCCCCCCC |
| 856 | EEEERKQQDQYNNEWEQEEYQRKGRGQGHQP | -47.39 | 35.5 | HHHHHHHHHHHHHHHHHHHHHHHHCCCCCCC |
| 577 | EEEERKQRDQYNNESEQKRYQRYGQGQGHQS | -47.39 | 35.5 | HHHHHHHHHHHCHHHHHHHHHHHHHCCCCCC |
| 332 | EEEERKQRDQYNNEWEQEEYQRKGRGQGHQP | -47.39 | 35.5 | HHHHHHHHHHHHHHHHHHHHHHHHCCCCCCC |
| 646 | EEERRKQLDQYNNEHEQQRYQREGKGQGHNH | -47.39 | 38.7 | HHHHHHHHHHHHHHHHHHHHHHHHCCCCCCC |
| 542 | EEEKKKQEDQRNNEEEQRRYQRYGRGQGHQP | -47.37 | 38.7 | HHHHHHHHHHHHHHHHHHHHHHHHHCCCCCC |
| 676 | EEEQKKQREQREDEKRDQEYQRSGRGKGHQP | -47.36 | 38.7 | HHHHHHHHHHHHHHHHHHHHHHHCCCCCCCC |
| 302 | EEEERKQQDQRKNRKEQEEYQRRGRGQGHQP | -47.36 | 29.0 | HHHHHHHHHHHHHHHHHHHHHHHHHCCCCCC |
| 749 | EEERRKQLQQYKQESEDQRYQRYGRGKGHQP | -47.34 | 38.7 | HHHHHHHHHHHHHHHHHHHHHHHHCCCCCCC |
| 371 | EEQRKKQLDQKNNEEEDKRYQKEGWGKGHYP | -47.33 | 51.6 | HHHHHHHHHHHHHHHHHHHHHHHHCCCCCCC |
| 878 | EEEKRKQEQQQNNEKEDSRYQQQGWGKGHNP | -47.32 | 38.7 | HHHHHHHHHHHHHHHHHHHHHHHCCCCCCCC |
| 897 | EEEEKKYQEQYNNDYEDEEYQRKGRGKGHQP | -47.29 | 38.7 | CHHHHHHHHHHHCCCHHHHHHHHHCCCCCCC |
| 972 | EQEQRKYEEQRNNDEEDQRYQREGHGKGHQR | -47.29 | 35.5 | HHHHHHHHHHHHHHHHHHHHHHHHCCCCCCC |
| 990 | EEEEKKYQEKYKNEFEDKYYQQKGQGKGHQP | -47.29 | 41.9 | CHHHHHHHHHHHHHHHHHHHHHHHCCCCCCC |
| 401 | EEEERKYQEKYNNESEDKRYQQQGRGKGHQP | -47.29 | 41.9 | CHHHHHHHHHHHHHHHHHHHHHHHCCCCCCC |
| 073 | EQEERKYQEKRNNEEEDKRYQRYGWGKGHYP | -47.29 | 38.7 | CHHHHHHHHHHHHHHHHHHHHHHCCCCCCCC |
| 450 | EQEQRHQQEKFSNEFEDQYYQKYGQGKGHQP | -47.27 | 38.7 | HHHHHHHHHHHHHHHHHHHHHHHHHCCCCCC |
| 311 | EQEQRHQQEKYSNEYEDEQYQRKGKGKGHSH | -47.27 | 35.5 | HHHHHHHHHHHHHHHHHHHHHHCCCCCCCCC |
| 527 | EEEEKKRQDQENNEREDKRYQREGKGKGHSH | -47.25 | 45.2 | CHHHHHHHHHHHHHHHHHHHHHHCCCCCCCC |
| 327 | EQEERKFRDQYNNDYEHEEYQRKGRGQGHQP | -47.24 | 32.3 | CHHHHHHHHHHHCHHHHHHHHHHHCCCCCCC |
| 399 | EEEERKQQDDYNNRWEDQRYQREGKGKGHSH | -47.24 | 38.7 | HHHHHHHHHHHHHHHHHHHHHHCCCCCCCCC |
| 145 | EEERRKQLDDYKNRSEDQRYQRYGRGKGHQP | -47.24 | 38.7 | HHHHHHHHHHHHCCCHHHHHHHHHCCCCCCC |
| 930 | EQEERKKEEQFRNDFEDRRYQQEGRGKGHQP | -47.23 | 32.3 | HHHHHHHHHHHHHHHHHHHHHHHHCCCCCCC |

| | | | | |
|---|---|---|---|---|
| 517 | EEEKRKQLDQYKNRYEQEEYQRKGRGQGHQP | -47.23 | 32.3 | HHHHHHHHHHHHHHHHHHHHHHHHCCCCCCC |
| 211 | EEEMRKLQDKINNEKEQEEYQRRGRGQGHQP | -47.23 | 38.7 | HHHHHHHHHHHHHHHHHHHHHHHHHCCCCCC |
| 232 | EEEEKKKRDKYNNEWEQKRYQREGRGQGHQP | -47.22 | 41.9 | CHHHHHHHHHHHHHHHHHHHHHHHCCCCCCC |
| 440 | EEEERKFQEKWKNEWEDKYYQQKGQGKGHSH | -47.19 | 41.9 | CHHHHHHHHHHHHHHHHHHHHHHHCCCCCCC |
| 474 | EEEERKQQDDRNNRKEDERYQQQGWGKGHNP | -47.17 | 38.7 | HHHHHHHHHHHHHHHHHHHHHHHCCCCCCCC |
| 952 | EEEERKQQDDRNNRKEDQRYQREGKGKGHNH | -47.17 | 38.7 | HHHHHHHHHHHHHHHHHHHHHHHCCCCCCCC |
| 221 | EEERRKQLEQYKNEWEDEEYQRKGRGKGHQP | -47.16 | 38.7 | HHHHHHHHHHHHHHHHHHHHHHHHCCCCCCC |
| 556 | EEEEKKKREQYNNEHEDQRYQQQGRGKGHQP | -47.16 | 41.9 | CHHHHHHHHHHHHHHHHHHHHHHHCCCCCCC |
| 375 | EEEEKKKREQYNNEYEDEEYQRKGRGKGHQP | -47.16 | 41.9 | CHHHHHHHHHHHHHHHHHHHHHHHCCCCCCC |
| 334 | EEEERKKEEQKRNEEEDQRYQRYGWGKGHNP | -47.16 | 35.5 | HHHHHHHHHHHHHHHHHHHHHHHCCCCCCCC |
| 981 | EEQERKNQQQYNNEYEDEEYQRKGRGKGHNP | -47.16 | 41.9 | HHHHHHHHHHHHHHHHHHHHHHHCCCCCCCC |
| 736 | EEEQKKQREQFNNEYEHQYKQQQGQGSGKQS | -47.15 | 35.5 | HHHHHHHHHHHHHHHHHHHHHHHCCCCCCCC |
| 668 | EQERRHQLEKYKNEWEDEYYQQYGQGKGHQP | -47.14 | 38.7 | HHHHHHHHHHHHHHHHHHHHHHHHHCCCCCC |
| 209 | EEERRKQLDKYKNEYEHEEKQRKGRGSGKQN | -47.14 | 35.5 | HHHHHHHHHHHHHHHHHHHHHHHCCCCCCCC |
| 444 | EEEEKKKQEKFSNEFEDRRYQQEGQGKGHQS | -47.14 | 45.2 | HHHHHHHHHHHHHHHHHHHHHHHHCCCCCCC |
| 010 | EEQRRKQLQQWKNEYEDQSYQREGRGKGHQP | -47.14 | 41.9 | HHHHHHHHHHHHHHHHHHHHHHHCCCCCCCC |
| 483 | EEEQRKQQDQYNNEWEQEEYQRKGRGQGHQP | -47.13 | 35.5 | HHHHHHHHHHHHHHHHHHHHHHHHCCCCCCC |
| 903 | EEEQRKQQDQYSNDFEQRYYQQQGQGQGHQS | -47.12 | 29.0 | HHHHHHHHHHHHHHHHHHHHHHHHCCCCCCC |
| 268 | EQEERKQQEQYEDEYRDEQYQQQGQGKGHQS | -47.11 | 29.0 | HHHHHHHHHHHHHHHHHHHHHHHHCCCCCCC |
| 581 | EQEQRHQQEQYNNDYEDEEYQRKGRGKGHQP | -47.11 | 32.3 | HHHHHHHHHHHHCCCHHHHHHHHHCCCCCCC |
| 282 | EQEERHNQEKFNNEYEDKSYQREGRGKGHQP | -47.09 | 41.9 | CHHHHHHHHHHHHHHHCHHHHHCCCCCCCCC |
| 389 | EQEERHQQEKRSNEKEDKEYQKQGRGKGHQP | -47.08 | 35.5 | CHHHHHHHHHHHHHHHHHHHHHHHCCCCCCC |
| 794 | EEEERKWREQKNNDEEDKRYQRYGRGKGHQP | -47.07 | 35.5 | CHHHHHHHHHHCHHHHHHHHHHHHCCCCCCC |
| 612 | EEEERKQQQQRNNEEEDQRYQREGKGKGHSS | -47.07 | 38.7 | HHHHHHHHHHHHHHHHHHHHHHHCCCCCCCC |
| 019 | EEEKKKWLEKLNNEKEDKRYQREGRGKGHQP | -47.06 | 48.4 | HHHHHHHHHHHHHHHHHHHHHHHHCCCCCCC |
| 641 | EEEARKQQEQYSNEHEDQRYQRQGKGKGDSR | -47.04 | 45.2 | HHHHHHHHHHHHHHHHHHHHHHHCCCCCCCC |
| 863 | EEERRKQQQQYSNEYEDQYYQQKGQGKGHQP | -47.04 | 35.5 | HHHHHHHHHHHHHHHHHHHHHHHHCCCCCCC |
| 065 | EEEERKYQDQKNNEEEDQRYQRYGKGKGDSR | -47.04 | 48.4 | CHHHHHHHHHHHHHHHHHHHHHHCCCCCCCC |
| 637 | EEEERKQRDQYNQEAEQERYQQQGKGQGHNH | -47.03 | 38.7 | HHHHHHHHHHHHHHHHHHHHHHHHCCCCCCC |
| 798 | EEEQKKQRDQFNQEKEQERYQQQGRGQGHQP | -47.02 | 41.9 | HHHHHHHHHHHHHHHHHHHHHHHHCCCCCCC |
| 602 | EEEKRKFLQKMEQEREDKEYQQKGRGKGHQP | -47.01 | 45.2 | HHHHHHHHHHHHHHHHHHHHHHHHCCCCCCC |
| 769 | EEEEKKKRDQQKNEEEHERYQQQGKGQGHNH | -47.01 | 35.5 | CHHHHHHHHHHHHHHHHHHHHHHHCCCCCCC |
| 168 | EEEERKKQDQINNEAEDKRYQREGRGKGNQN | -47.00 | 45.2 | HHHHHHHHHHHHHHHHHHHHHHHHCCCCCCC |
| 365 | QDEERKQQDQRNNDNEQRQYQRSGRGQGHQP | -47.00 | 29.0 | CHHHHHHHHHHHHHHHHHHHHHHCCCCCCCC |
| 412 | EEEKKKLLEQISNDKEDREYQRRGRGKGHQP | -47.00 | 38.7 | HHHHHHHHHHHHHHHHHHHHHHHHCCCCCCC |
| 205 | EEEQRKLYEQISNEKEDQEYQKQGRGKGHQP | -46.99 | 35.5 | HHHHHHHHHHHHHHHHHHHHHHHHCCCCCCC |
| 506 | EEEEKKQRDQQNNEKEQKRYQREGRGQGHQP | -46.98 | 38.7 | HHHHHHHHHHHHHHHHHHHHHHHHCCCCCCC |
| 284 | EQEERHQQEQYNNDWEDQRYQREGWGKGHNP | -46.98 | 32.3 | CHHHHHHHHHHHCHHHHHHHHHHHCCCCCCC |

| | | | | |
|---|---|---|---|---|
| 319 | EEEQRKQEQQWQNEWEDQYYQQKGQGKGHNH | -46.97 | 35.5 | HHHHHHHHHHHHHHHHHHHHHHHHHCCCCCCC |
| 169 | EEEERKKRQQYSNDFEDRYYQQKGQGKGHNH | -46.97 | 32.3 | CHHHHHHHHHHHHHHHHHHHHHHHHCCCCCCC |
| 850 | EEEEKKKREQYNNEAEDQRYQREGRGKGHQP | -46.96 | 45.2 | CHHHHHHHHHHHHHHHHHHHHHHHHCCCCCCC |
| 838 | EEEERKREDQFRQEFEDEYYQQKGQGKGHNH | -46.95 | 41.9 | HHHHHHHHHHHHHHHHHHHHHHHHHCCCCCCC |
| 456 | EEEERKKQEKWKNEFEDKYYQQKGQGKGHSH | -46.94 | 38.7 | HHHHHHHHHHHHHHHHHHHHHHHHHCCCCCCC |
| 778 | EEERRKQLQKMKNEKEHEEYQRRGRGQGHQP | -46.93 | 35.5 | HHHHHHHHHHHHHHHHHHHHHHHHHHCCCCCC |
| 174 | EEERKKQLDQYNNESEQQRYQKQGKGQGHSS | -46.92 | 41.9 | HHHHHHHHHHHHCHHHHHHHHHHHCCCCCCCC |
| 837 | EEERRKQLDQMKNEKEQEEYQRYGRGQGHQP | -46.92 | 35.5 | HHHHHHHHHHHHHHHHHHHHHHHHHHCCCCCC |
| 090 | EEEERKRQEKENNEKEDKRYQREGRGKGHQP | -46.91 | 41.9 | HHHHHHHHHHHHHHHHHHHHHHHHHCCCCCCC |
| 309 | EEEERKRQEKENNEKEDKRYQRYGKGKGHNS | -46.91 | 41.9 | HHHHHHHHHHHHHHHHHHHHHHHHCCCCCCCC |
| 650 | EEEEKKNREQYNNESEDQRYQRYGKGKGHSS | -46.90 | 41.9 | HHHHHHHHHHHCCCHHHHHHHHHHCCCCCCCC |
| 234 | EEEERKNQEQFSNEFEDQYYQQKGQGKGHQP | -46.90 | 38.7 | HHHHHHHHHHHHHHHHHHHHHHHHHCCCCCCC |
| 963 | EEEERKKQDKFNNEFEQKFYQQKGRGQGHQP | -46.82 | 45.2 | HHHHHHHHHHHHHHHHHHHHHHHHHCCCCCCC |
| 362 | EEEERKQRQQYNNEWEDKRYQREGKGKGHSH | -46.81 | 38.7 | HHHHHHHHHHHHHHHHHHHHHHHCCCCCCCCC |
| 276 | EQEKRKMLEKISNEKEDEEYQRRGQGKGHQS | -46.79 | 38.7 | HHHHHHHHHHHHHHHHHHHHHHHHHHCCCCCC |
| 597 | EEEQRKQQDQYNNDYEQREYQRKGRGQGHQP | -46.79 | 32.3 | HHHHHHHHHHHHHHHHHHHHHHHHHCCCCCCC |
| 815 | EEEEKKQRDQFNNEYEQQSYQREGRGQGHQP | -46.78 | 41.9 | HHHHHHHHHHHHHHHHHHHHHHHHCCCCCCCC |
| 191 | EEEKRKQEDQRNNEKEQEEYQRYGRGQGHQP | -46.78 | 35.5 | HHHHHHHHHHHHHHHHHHHHHHHHHHCCCCCC |
| 015 | EEERRKQLEKISNEEEDKEYQRQGRGKGHQP | -46.77 | 41.9 | HHHHHHHHHHHHHHHHHHHHHHHHHCCCCCCC |
| 942 | EEERRKQLEKYNNESEDKRYQRYGRGKGHQP | -46.77 | 45.2 | HHHHHHHHHHHHHHHHHHHHHHHHHCCCCCCC |
| 237 | EQERRKQLEKESNEKEDKSYQRHGWGKGHNP | -46.77 | 38.7 | HHHHHHHHHHHHHHHHHHHHHHHCCCCCCCCC |
| 825 | EEERRKQLDKESNEKEHKRYQREGRGQGHQP | -46.76 | 38.7 | HHHHHHHHHHHHHHHHHHHHHHHHHCCCCCCC |
| 580 | EEERRKQLDKINNEQEHKRYQREGWGQGHYP | -46.76 | 41.9 | HHHHHHHHHHHHHHHHHHHHHHHHHCCCCCCC |
| 902 | EQQQRKQQEKYNNEWEQKRYQREGRGQGHQP | -46.76 | 35.5 | HHHHHHHHHHHHHHHHHHHHHHHHHCCCCCCC |
| 431 | EEEQRKQEEKRNNEKEQKRYQKEGWGQGHNP | -46.76 | 35.5 | HHHHHHHHHHHHHHHHHHHHHHHHHCCCCCCC |
| 161 | EEEQRRQEDQRNNEEEQKRYQREGRGQGHQP | -46.75 | 35.5 | HHHHHHHHHHHHHHHHHHHHHHHHHCCCCCCC |
| 361 | EEEQKKQRDQYNNEWEQERYQQQGRGQGHQP | -46.72 | 38.7 | HHHHHHHHHHHHHHHHHHHHHHHHHCCCCCCC |
| 114 | EQEERKKQEQYNNDWEDRRYQREGRGKGHQP | -46.70 | 32.3 | HHHHHHHHHHHHHHHHHHHHHHHHHCCCCCCC |
| 016 | EEEQRKQQDDYNHRWEDQRYQREGRGKGHQP | -46.68 | 41.9 | HHHHHHHHHHHHHHHHHHHHHHHHHCCCCCCC |
| 604 | QQEEEKRQRQYNEEYEHEEYQRKGRGRGHQP | -46.66 | 25.8 | CHHHHHHHHHHHHHHHHHHHHHHHHCCCCCCC |
| 936 | EEERRKQLDQYKNEFEQQYYQQKGQGQGHQP | -46.65 | 35.5 | HHHHHHHHHHHHHHHHHHHHHHHHHCCCCCCC |
| 249 | EQEQRHQQEKYNNEWEDKRYQREGQGKGHQS | -46.65 | 38.7 | HHHHHHHHHHHHHHHHHHHHHHHHHCCCCCCC |
| 563 | EEEERKYQQKYNNEEEDKKYQKEGRGKGHQP | -46.65 | 41.9 | CHHHHHHHHHHHHHHHHHHHHHHHHCCCCCCC |
| 283 | EEEERKYQQKYNNESEDKRYQQQGRGKGHQP | -46.65 | 41.9 | CHHHHHHHHHHHHHHHHHHHHHHHHCCCCCCC |
| 854 | EEEERKQQDAEKNRKEDQRYQREGRGKGHQP | -46.65 | 35.5 | HHHHHHHHHHHHHHHHHHHHHHHHHCCCCCCC |
| 809 | EEEERKRQDQYNNEYEDEEYQRKGRGKGDSR | -46.64 | 48.4 | HHHHHHHHHHHHHHHHHHHHHHHHCCCCCCCC |
| 468 | EEEQKKYEDKRNQEEEQKRYQQQGWGQGHNP | -46.64 | 41.9 | CHHHHHHHHHHHHHHHHHHHHHHHCCCCCCCC |
| 126 | EEEERKQRDDYNNRWEDERYQRQGRGKGHQP | -46.63 | 38.7 | HHHHHHHHHHHHHHHHHHHHHHHHHCCCCCCC |

| | | | | |
|---|---|---|---|---|
| 346 | EEEEKKQREKFSNEFEQKYYQQKGQGQGHQP | -46.63 | 38.7 | CHHHHHHHHHHHHHHHHHHHHHHHCCCCCCC |
| 662 | EEEERKKQDKYNNEAEQKRYQREGKGQGHSH | -46.62 | 41.9 | HHHHHHHHHHHHHHHHHHHHHHHHCCCCCCC |
| 983 | EQEQRKQQDQYNNEYEQEQYQRQGRGQGNQN | -46.58 | 32.3 | HHHHHHHHHHHHHHHHHHHHHHHCCCCCCCC |
| 880 | EEEERKKQQQWKNEWEDKYYQQQGQGKGHSH | -46.58 | 35.5 | HHHHHHHHHHHHHHHHHHHHHHHHCCCCCCC |
| 492 | EEERKKQLDQIKNEKEHEEKQRRGKGSGKSN | -46.57 | 35.5 | HHHHHHHHHHHHHHHHHHHHHHHCCCCCCCC |
| 758 | EQEQRKQEEQRNNEEEHKRYQQQGWGQGHNP | -46.57 | 29.0 | HHHHHHHHHHHHHHHHHHHHHHHCCCCCCCC |
| 201 | EQEQRHQQEQYNQEAEDQRYQREGKGKGHSH | -46.57 | 38.7 | HHHHHHHHHHHHHHHHHHHHHHHCCCCCCCC |
| 367 | EEEERKYRDQKNNEEEHERYQQQGRGQGHQP | -46.56 | 35.5 | CHHHHHHHHHHCHHHHHHHHHHHHCCCCCCC |
| 240 | EEEQRKYEDQRNNEEEHEEYQRKGRGQGHQP | -46.56 | 35.5 | CHHHHHHHHHHCHHHHHHHHHHHHCCCCCCC |
| 142 | EEEERKKQEQQSNEKEDEEYQRRGWGKGHYP | -46.56 | 35.5 | HHHHHHHHHHHHHHHHHHHHHHHCCCCCCCC |
| 435 | EEEERKKQEQYNNEWEDEEYQRKGRGKGHQP | -46.56 | 38.7 | HHHHHHHHHHHHHHHHHHHHHHHHCCCCCCC |
| 414 | EEEERKSRDQQKNEKEHEEKQRRGQGSGKQS | -46.56 | 29.0 | CHHHHHHHHHHHHHHHHHHHHHHCCCCCCCC |
| 503 | EEEEKKRQDQYNNEAEDQRYQREGKGKGNSN | -46.55 | 48.4 | CHHHHHHHHHHHHHHHHHHHHHHCCCCCCCC |
| 764 | EEERRKQLDQKNNDEEDRRYQQYGKGKGDSR | -46.54 | 48.4 | HHHHHHHHHHHCCHHHHHHHHHHCCCCCCCC |
| 649 | EEERRKQLDMIKNRKEQEEYQRYGRGQGHQP | -46.54 | 32.3 | HHHHHHHHHHHHHHHHHHHHHHHHHCCCCCC |
| 701 | EEEERKKEQQYRNEYEDEEYQRKGRGKGHQP | -46.52 | 35.5 | HHHHHHHHHHHHHHHHHHHHHHHHCCCCCCC |
| 810 | EDERRKQLDQKNNEEEQQRYQRYGRGQGHQP | -46.52 | 35.5 | HHHHHHHHHHHCHHHHHHHHHHHHHCCCCCC |
| 096 | EEEEKKYQEQFNNEFEDQYYQQQGRGKGHQP | -46.52 | 45.2 | CHHHHHHHHHHHHHHHHHHHHHHHCCCCCCC |
| 611 | EEEEKKYQEQYNNEWEDKRYQREGRGKGHQP | -46.52 | 41.9 | CHHHHHHHHHHHHHHHHHHHHHHHCCCCCCC |
| 693 | EEEERKYQEQYKNEYEDEEYQRKGRGKGHQP | -46.52 | 35.5 | CHHHHHHHHHHHHHHHHHHHHHHHCCCCCCC |
| 009 | EEEERKYQEQYNNEYEDEEYQRKGRGKGHQP | -46.52 | 38.7 | CHHHHHHHHHHHHHHHHHHHHHHHCCCCCCC |
| 356 | EEEQKKYEEQRNNEAEDKRYQRYGRGKGHQP | -46.52 | 45.2 | HHHHHHHHHHHHHHHHHHHHHHHHCCCCCCC |
| 152 | EEEERKLRQQIENEREDKYYQQKGQGKGHSH | -46.51 | 35.5 | HHHHHHHHHHHHHHHHHHHHHHHHCCCCCCC |
| 376 | EEERRKQLDQFNNEFEHKYKQQKGQGSGKSH | -46.50 | 38.7 | HHHHHHHHHHHHHHHHHHHHHHHCCCCCCCC |
| 128 | EEEQKKQQDQYNNEAEDQRYQRYGRGKGHQP | -46.50 | 48.4 | HHHHHHHHHHHHHHHHHHHHHHHHCCCCCCC |
| 824 | EEEERKQRDQYNNEFEQQYYQQKGQGQGHNH | -46.49 | 35.5 | HHHHHHHHHHHHHHHHHHHHHHHHCCCCCCC |
| 373 | EEEEKKRQEKDSNEREDKRYQREGRGKGHQP | -46.48 | 41.9 | HHHHHHHHHHHHHHHHHHHHHHHHCCCCCCC |
| 828 | EEEERKREEKYRNEHEDKRYQRQGWGKGHFP | -46.48 | 41.9 | HHHHHHHHHHHHHHHHHHHHHHHHCCCCCCC |
| 258 | EEQEKKRQEKEKNEKEDKRYQREGRGKGHQP | -46.48 | 45.2 | HHHHHHHHHHHHHHHHHHHHHHHHCCCCCCC |
| 188 | EEEERKKQEQIKQEAEDQRYQREGKGKGHNH | -46.47 | 38.7 | HHHHHHHHHHHHHHHHHHHHHHHHCCCCCCC |
| 476 | EEEERKQQQQKEDEERDQRYQRYGRGKGHQP | -46.47 | 32.3 | HHHHHHHHHHHHHHHHHHHHHHHHCCCCCCC |
| 007 | EEEERKQRQQYEDEWRDERYQREGQGKGHSS | -46.47 | 32.3 | HHHHHHHHHHHHHHHHHHHHHHHCCCCCCCC |
| 560 | EEEAKKFREQYNNEWEDEEYQRKGRGKGHQP | -46.42 | 48.4 | HHHHHHHHHHHHHHHHHHHHHHHHCCCCCCC |
| 239 | EEEERKFQEQYRNEWEDKRYQREGKGKGHNH | -46.42 | 38.7 | CHHHHHHHHHHHHHHHHHHHHHCCCCCCCCC |
| 006 | EEEERKFQEQYRNEYEDEEYQRKGRGKGHQP | -46.42 | 38.7 | CHHHHHHHHHHHHHHHHHHHHHHHCCCCCCC |
| 408 | EEEERKFREQKNNEEEDERYQQQGKGKGHNH | -46.42 | 41.9 | CHHHHHHHHHHHHHHHHHHHHHHHCCCCCCC |
| 786 | EEEQRKFEEQRNNEEEDEEYQRKGRGKGHQP | -46.42 | 41.9 | HHHHHHHHHHHHHHHHHHHHHHHHCCCCCCC |
| 605 | EEEQRKFEEQYRNEYEDQEYQKEGQGKGHQS | -46.42 | 38.7 | HHHHHHHHHHHHHHHHHHHHHHHHCCCCCCC |

| | | | | |
|---|---|---|---|---|
| 682 | EEEQRKFYEQFNNEFEDQYYQQKGQGKGHNH | -46.42 | 45.2 | HHHHHHHHHHHHHHHHHHHHHHHHCCCCCCC |
| 176 | EEEKRKLLEQIKNEKEDEEYQRRGQGKGHSS | -46.39 | 38.7 | HHHHHHHHHHHHHHHHHHHHHHHHHCCCCCC |
| 559 | EEQQRKQQDAFSNRFEQEYYQRKGQGQGHNH | -46.37 | 35.5 | HHHHHHHHHHHHHHHHHHHHHHHHCCCCCCC |
| 101 | EEEERKKQEQYNNEAEDERYQRQGRGKGHQP | -46.37 | 41.9 | HHHHHHHHHHHHHHHHHHHHHHHHCCCCCCC |
| 118 | EEEQKKLEDKKKNEQEHRRYQREGKGQGHNH | -46.35 | 38.7 | HHHHHHHHHHHHHHHHHHHHHHHHCCCCCCC |
| 763 | EEEKRKLLQQISNEKEDEEYQRRGWGKGHNP | -46.35 | 38.7 | HHHHHHHHHHHHHHHHHHHHHHHCCCCCCCC |
| 901 | EEERRKQLQQMKDEKRDEEYQRYGQGKGHQS | -46.35 | 35.5 | HHHHHHHHHHHHHHHHHHHHHHHHHCCCCCC |
| 881 | EQEQRHQKEQEKNRREDEEYQRKGRGKGHQP | -46.34 | 29.0 | HHHHHHHHHHHHHHHHHHHHHHHHCCCCCCC |
| 095 | EEEKRKLLDKIKNEKEQKEYQKQGRGQGHQP | -46.33 | 38.7 | HHHHHHHHHHHHHHHHHHHHHHHHCCCCCCC |
| 193 | KQEQEKQYEQEKQEKEDRRYQREGRGKGHQP | -46.33 | 29.0 | HHHHHHHHHHHHHHHHHHHHHHHHCCCCCCC |
| 140 | EEEQRKREQKLRQEKEDKRYQREGRGKGHQP | -46.32 | 38.7 | HHHHHHHHHHHHHHHHHHHHHHHHCCCCCCC |
| 396 | EQEERKWQEQRSNDEEDRRYQRYGRGKGHQP | -46.31 | 29.0 | CHHHHHHHHHHHCHHHHHHHHHHHCCCCCCC |
| 807 | EEEERKQRDQYNNESEQQRYQRYGRGQGHQP | -46.31 | 35.5 | HHHHHHHHHHHCHHHHHHHHHHHHHCCCCCC |
| 445 | EEEERKKQQKINNEAEDKRYQRYGQGKGHQS | -46.30 | 45.2 | HHHHHHHHHHHHHHHHHHHHHHHHCCCCCCC |
| 814 | EEEERKKQQKINNEAEDKRYQRYGRGKGHQP | -46.30 | 45.2 | HHHHHHHHHHHHHHHHHHHHHHHHCCCCCCC |
| 419 | EEEERKKQDQFNNRYEQESYQRYGKGQGHSS | -46.30 | 35.5 | HHHHHHHHHHHHHHHHHHHHHHHHCCCCCCC |
| 159 | EQEYRKWQEQYNNEFEDQYYQQKGQGKGHQP | -46.30 | 35.5 | HHHHHHHHHHHHHHHHHHHHHHHHCCCCCCC |
| 710 | EQEYRKWQEQYSNESEDQRYQRYGQGKGHQS | -46.30 | 32.3 | HHHHHHHHHHHHCCHHHHHHHHHHHCCCCCC |
| 235 | EEEKKKWLEQLSNEKEDKRYQREGKGKGHQP | -46.29 | 41.9 | HHHHHHHHHHHHHHHHHHHHHHHCCCCCCCC |
| 190 | EQQQRHQQEQYSNESEDQRYQKQGRGKGHQP | -46.28 | 35.5 | HHHHHHHHHHHCCHHHHHHHHHHHCCCCCCC |
| 855 | EQEQRHQQEQYSNEYEDEEYQRKGRGKGHQP | -46.27 | 32.3 | HHHHHHHHHHHHHHHHHHHHHHHHCCCCCCC |
| 388 | EEEEKKKRDQYNNEWEQEEYQRKGRGQGHQP | -46.27 | 38.7 | CHHHHHHHHHHHHHHHHHHHHHHHCCCCCCC |
| 743 | EEEERKKQDQYNNESEQQRYQRYGQGQGHQS | -46.27 | 35.5 | HHHHHHHHHHHCHHHHHHHHHHHHHCCCCCC |
| 292 | KQEEEKKQDQYNNEYEQEEYQRKGRGQGHQP | -46.27 | 29.0 | CHHHHHHHHHHHHHHHHHHHHHHHCCCCCCC |
| 512 | EEEEKKNQDQWKNEFEQKYFQQKGQGQGKSH | -46.25 | 32.3 | HHHHHHHHHHHHHHHHHHHHHHHHCCCCCCC |
| 574 | EEERKKQLDAYNNRYEQQYYQKKGQGQGHQP | -46.24 | 38.7 | HHHHHHHHHHHHHHHHHHHHHHHHCCCCCCC |
| 308 | EEEQRRQQEQRNNEKEDEEYQRRGMGKGHQP | -46.19 | 38.7 | HHHHHHHHHHHHHHHHHHHHHHHHHCCCCCC |
| 437 | EEEERKKQQQYNNEAEDQRYQRYGRGKGHQP | -46.18 | 41.9 | HHHHHHHHHHHHHHHHHHHHHHHHCCCCCCC |
| 655 | EEEERKYQQQQKQEEEDKRYQRYGRGKGHQP | -46.18 | 35.5 | HHHHHHHHHHHHHHHHHHHHHHHHCCCCCCC |
| 882 | EEEEKKKQEQESNEKEDKRYQKQGQGKGHQS | -46.18 | 38.7 | HHHHHHHHHHHHHHHHHHHHHHHCCCCCCCC |
| 590 | EEEEKKKREQINNEAEDQRYQREGRGKGHQP | -46.18 | 45.2 | HHHHHHHHHHHHHHHHHHHHHHHHCCCCCCC |
| 932 | EQEERKKLEQISNEKEDQEYQKQGWGKGHNP | -46.18 | 35.5 | CHHHHHHHHHHHHHHHHHHHHHHCCCCCCCC |
| 083 | EEEERKKQDDFKNRKEDERYQQQGRGKGHQP | -46.16 | 38.7 | HHHHHHHHHHHHHHHHHHHHHHHHCCCCCCC |
| 595 | EEEERKMRQKYNNESEDKRYQRYGKGKGHSS | -46.16 | 41.9 | HHHHHHHHHHHHHHHHHHHHHHHCCCCCCCC |
| 920 | EEEKRKMLQKKKNEEEDKRYQQQGWGKGHNP | -46.16 | 41.9 | HHHHHHHHHHHHHHHHHHHHHHHCCCCCCCC |
| 525 | EEEKRKMLQKMKNEKEDEEYQRRGKGKGHSH | -46.16 | 41.9 | HHHHHHHHHHHHHHHHHHHHHHHHCCCCCCC |
| 719 | EEQRRKQLDQYKNESEQQRYQRYGKGQGDSR | -46.16 | 45.2 | HHHHHHHHHHHHHHHHHHHHHHHCCCCCCCC |
| 728 | DEERKKMLDKKNQEEEQKRYQQYGRGQGHQP | -46.15 | 41.9 | HHHHHHHHHHHHHHHHHHHHHHHHHCCCCCC |

| | | | | |
|---|---|---|---|---|
| 998 | EEEERKRQEQEKNEKEDQRYQQQGWGKGHNP | -46.15 | 35.5 | HHHHHHHHHHHHHHHHHHHHHHHCCCCCCCC |
| 761 | EQEERHQQEQYSNEYEDQEYQKQGRGKGHQP | -46.14 | 32.3 | CHHHHHHHHHHHHHHHHHHHHHHHCCCCCCC |
| 847 | EQERRHQLEQMKNEKEDQEYQRAGRGKGHQP | -46.14 | 35.5 | HHHHHHHHHHHHHHHHHHHHHHHHHCCCCCC |
| 618 | EQERRHQLEQQKNEEEDKRYQRYGRGKGHQP | -46.14 | 35.5 | HHHHHHHHHHHHHHHHHHHHHHHHCCCCCCC |
| 457 | EQERRHQLEQWKNEWEDKYYQKEGQGKGHQS | -46.14 | 35.5 | HHHHHHHHHHHHHHHHHHHHHHHCCCCCCCC |
| 218 | EEERRKQLQKIKNEKEDEEYQRRGQGKGHQS | -46.13 | 41.9 | HHHHHHHHHHHHHHHHHHHHHHHHHCCCCCC |
| 061 | EEERRKQLQKINNEAEDKRYQREGRGKGHQP | -46.13 | 48.4 | HHHHHHHHHHHHHHHHHHHHHHHHCCCCCCC |
| 297 | EEERRKQLQKYKNEFEDKYYQQKGQGKGHSH | -46.13 | 41.9 | HHHHHHHHHHHHHHHHHHHHHHHHCCCCCCC |
| 944 | EEERRKQLEQQNNEKEDQRYQREGRGKGHQP | -46.12 | 41.9 | HHHHHHHHHHHHHHHHHHHHHHHHCCCCCCC |
| 915 | EEEQRKQQDQYNNEWEQEEYQRKGRGQGNQN | -46.11 | 35.5 | HHHHHHHHHHHHHHHHHHHHHHHHCCCCCCC |
| 739 | EQQQRKYEEQRNNREEDERYQRYGQGKGHQS | -46.09 | 35.5 | HHHHHHHHHHHHHHHHHHHHHHHHHCCCCCC |
| 135 | EEEERKLRDQINNEKEQEEYQRRGKGQGHNH | -46.09 | 35.5 | HHHHHHHHHHHHHHHHHHHHHHHHCCCCCCC |
| 378 | EEEERKLRDQINNEKEQEEYQRYGRGQGHQP | -46.09 | 35.5 | HHHHHHHHHHHHHHHHHHHHHHHHHCCCCCC |
| 817 | EQEERHQQEQRNNEKEDEEYQRRGWGKGHNP | -46.08 | 35.5 | CHHHHHHHHHHHHHHHHHHHHHHCCCCCCCC |
| 315 | EEEKRKMLDQINNEKEHEEYQRRGQGQGHQS | -46.07 | 38.7 | HHHHHHHHHHHHHHHHHHHHHHHHHCCCCCC |
| 443 | EEEQRKQYQKESNEKEDEEYQRKGRGKGHQP | -46.06 | 38.7 | HHHHHHHHHHHHHHHHHHHHHHHHCCCCCCC |
| 853 | EEEQRKQYQKESNEKEDKRYQREGKGKGHQP | -46.06 | 38.7 | HHHHHHHHHHHHHHHHHHHHHHHCCCCCCCC |
| 796 | EEEERKYQQQKENEREDQEYQRKGRGKGHQP | -46.05 | 35.5 | CHHHHHHHHHHHHHHHHHHHHHHHCCCCCCC |
| 270 | EEEERKKQDQFSNEFEQQYYQQKGQGQGHSH | -46.04 | 35.5 | HHHHHHHHHHHHHHHHHHHHHHHHCCCCCCC |
| 136 | DEERKKMLEQIKNEAEDQRYQRYGWGKGHNP | -46.03 | 41.9 | HHHHHHHHHHHHHHHHHHHHHHHCCCCCCCC |
| 098 | EEEKRKMLEQKNNEEEDQRYQRYGQGKGHQS | -46.03 | 41.9 | HHHHHHHHHHHCHHHHHHHHHHHHHCCCCCC |
| 022 | EQEKRKMLEQWKNEWEDKYYQQQGQGKGHQS | -46.03 | 35.5 | HHHHHHHHHHHHHHHHHHHHHHHHCCCCCCC |
| 323 | EQEKRHQLEQMKNDKEDREYQRSGKGKGHSN | -46.02 | 35.5 | HHHHHHHHHHHHHHHHHHHHHHCCCCCCCCC |
| 576 | EQERRKQLEQINNDAEDRRYQQYGKGKGHSS | -46.02 | 38.7 | HHHHHHHHHHHHHHHHHHHHHHHCCCCCCCC |
| 775 | EEEERKKEQQKRNEEEDQRYQQYGRGKGHQP | -46.01 | 35.5 | HHHHHHHHHHHHHHHHHHHHHHHHHCCCCCC |
| 823 | EEERKKQLEQESNEKEDQKYQRQGQGKGHQS | -46.00 | 41.9 | HHHHHHHHHHHHHHHHHHHHHHHHCCCCCCC |
| 490 | EEERKKQLEQINNEAEDQRYQREGWGKGHYP | -46.00 | 48.4 | HHHHHHHHHHHHHHHHHHHHHHHHCCCCCCC |
| 946 | EEERKKQLEQWKNEWEDKYYQQKGQGKGHSH | -46.00 | 41.9 | HHHHHHHHHHHHHHHHHHHHHHHHCCCCCCC |
| 392 | EEERRKQLEQINNEAEDKRYQREGKGKGHSS | -46.00 | 45.2 | HHHHHHHHHHHHHHHHHHHHHHHCCCCCCCC |
| 621 | EEERRKQLEQYKNEHEDRRYQEQGKGKGHSS | -46.00 | 38.7 | HHHHHHHHHHHHHHHHHHHHHHHCCCCCCCC |
| 071 | EQEERKYQEQKNNEEEDERYQRYGQGKGHQS | -46.00 | 35.5 | CHHHHHHHHHHHHHHHHHHHHHHHHCCCCCC |
| 288 | EEEERKKQDKIKNEKEQEEYQRRGQGQGHQS | -45.99 | 35.5 | HHHHHHHHHHHHHHHHHHHHHHHHHCCCCCC |
| 029 | EEEEKKRQEQENNDKEDRRYQREGQGKGHSS | -45.99 | 38.7 | HHHHHHHHHHHHCHHHHHHHHHHCCCCCCCC |
| 804 | EEERRKQLQKWKNEYEQKSYQREGKGQGHNH | -45.99 | 35.5 | HHHHHHHHHHHHHHHHHHHHHHCCCCCCCCC |
| 465 | EQEERKKQEQINNEAEDKRYQRYGQGKGHQS | -45.98 | 38.7 | HHHHHHHHHHHHHHHHHHHHHHHHCCCCCCC |
| 558 | EEEEKKRQDKEKNEKEQKRYQREGRGQGHQP | -45.97 | 38.7 | CHHHHHHHHHHHHHHHHHHHHHHHCCCCCCC |
| 508 | EEEERKRQEQFSNEFEDQYYQQKGQGKGHSH | -45.97 | 38.7 | HHHHHHHHHHHHHHHHHHHHHHHHCCCCCCC |
| 470 | EQEQRKRQEQYNNEYEDEEYQRKGRGKGHQP | -45.97 | 35.5 | HHHHHHHHHHHHHHHHHHHHHHHHCCCCCCC |

| | | | | |
|---|---|---|---|---|
| 964 | EQEERHQQEQYNNEWEDKRYQREGRGKGHQP | -45.95 | 35.5 | CHHHHHHHHHHHHHHHHHHHHHHHCCCCCCC |
| 233 | EEEERKRQDQEKNEKEHQRYQREGRGQGHQP | -45.93 | 32.3 | HHHHHHHHHHHHHHHHHHHHHHHHCCCCCCC |
| 497 | EEEKRKQEQQRNNEEEHQRYQREGKGQGHSH | -45.93 | 32.3 | HHHHHHHHHHHHHHHHHHHHHHHHCCCCCCC |
| 133 | EEEQKKQREQYNNRYEHEYYQEKGQGQGHQP | -45.93 | 32.3 | HHHHHHHHHHHHHHHHHHHHHHHHCCCCCCC |
| 072 | EEEEKKQREQESNEKEDERYQREGQGKGHQS | -45.93 | 38.7 | HHHHHHHHHHHHHHHHHHHHHHHHCCCCCCC |
| 564 | EQEERKQREQENNEKEDRRYQRYGWGKGHNP | -45.93 | 35.5 | HHHHHHHHHHHHHHHHHHHHHHCCCCCCCCC |
| 482 | EQEERKQREQWKQDWEQQYYQQQGQGQGHQS | -45.93 | 22.6 | HHHHHHHHHHHHHHHHHHHHHHHHCCCCCCC |
| 252 | EEEQRKYEDQRNQEEEQQRYQREGKGQGHQS | -45.93 | 35.5 | CHHHHHHHHHHHHHHHHHHHHHHHCCCCCCC |
| 634 | EEEQRKYEDQRSQEEEQKRYQREGRGQGHQP | -45.93 | 32.3 | CHHHHHHHHHHHHHHHHHHHHHHHCCCCCCC |
| 899 | EEEEKKYQEQYSNEWEDEEYQRKGRGKGHQP | -45.92 | 38.7 | CHHHHHHHHHHHHHHHHHHHHHHHCCCCCCC |
| 250 | EEEEKKKREQYNNEAEDKRYQREGKGKGDSR | -45.92 | 51.6 | CHHHHHHHHHHHHHHHHHHHHHHCCCCCCCC |
| 108 | EQEERKQREQESNRKEDERYQREGKGKGHNH | -45.90 | 29.0 | HHHHHHHHHHHHHHHHHHHHHHHCCCCCCCC |
| 026 | EQEQRKQEEQRSQEKEQEEYQRYGRGQGHQP | -45.90 | 25.8 | HHHHHHHHHHHHHHHHHHHHHHHHHCCCCCC |
| 500 | EEEERKIQQQLNNEKEDEEYQRKGRGKGDSR | -45.89 | 45.2 | HHHHHHHHHHHHHHHHHHHHHHHCCCCCCCC |
| 219 | EEEERKYQQQKNNEEEDEEYQRKGRGKGHQP | -45.88 | 38.7 | CHHHHHHHHHHHHHHHHHHHHHHHCCCCCCC |
| 896 | EEEERKYQQQKNNEEEDERYQQQGRGKGHQP | -45.88 | 38.7 | CHHHHHHHHHHHHHHHHHHHHHHHCCCCCCC |
| 082 | EEEERKYQQQKNNEEEDQRYQKQGWGKGHNP | -45.88 | 38.7 | CHHHHHHHHHHHHHHHHHHHHHHCCCCCCCC |
| 117 | EEEKRKYLQQYNNESEDERYQQQGKGKGHNH | -45.88 | 41.9 | HHHHHHHHHHHHCCHHHHHHHHHHCCCCCCC |
| 759 | EEEQRKYEQQRNNEEEDQRYQKQGRGKGHQP | -45.88 | 38.7 | HHHHHHHHHHHHHHHHHHHHHHHCCCCCCCC |
| 568 | EQEQRHQQEQYSNEWEDEEYQRKGRGKGHQP | -45.88 | 32.3 | HHHHHHHHHHHHHHHHHHHHHHHHCCCCCCC |
| 441 | EEEERKKQDQYNNESEQQRYQRYGRGQGHQP | -45.87 | 35.5 | HHHHHHHHHHHCHHHHHHHHHHHHHCCCCCC |
| 013 | EEEERKQQQKKKQEEEQKRYQREGRGQGHQP | -45.86 | 32.3 | HHHHHHHHHHHHHHHHHHHHHHHHCCCCCCC |
| 692 | EEEERKRQQQFSNDYEDKFYQQQGRGKGHQP | -45.85 | 38.7 | HHHHHHHHHHHHHHHHHHHHHHHHCCCCCCC |
| 182 | EEEERKKEDQYRNEWEQKRYQREGRGQGHQP | -45.79 | 32.3 | HHHHHHHHHHHHHHHHHHHHHHHHCCCCCCC |
| 306 | EEEERKFQQQYKNEHEDQRYQQQGWGKGHNP | -45.78 | 38.7 | CHHHHHHHHHHHHHHHHHHHHHHCCCCCCCC |
| 594 | EEERKKQLEQESNEKEDERYQQQGQGKGHQS | -45.77 | 41.9 | HHHHHHHHHHHHHHHHHHHHHHHHCCCCCCC |
| 074 | EEEKRKFLDKYKNEYEQEEYQRKGRGQGHQP | -45.77 | 41.9 | HHHHHHHHHHHHHHHHHHHHHHHHCCCCCCC |
| 842 | EQEERHQQEQRSNEQEDKRYQRYGRGKGHQP | -45.75 | 32.3 | CHHHHHHHHHHHHHHHHHHHHHHHCCCCCCC |
| 883 | EEEQRKEEQQKNNDSEDRRYQREGQGKGHQS | -45.74 | 35.5 | HHHHHHHHHHHHCHHHHHHHHHHHCCCCCCC |
| 312 | EEEQRKQEEQRRNDKEQKRYQREGRGQGHQP | -45.73 | 25.8 | HHHHHHHHHHHHHHHHHHHHHHHHCCCCCCC |
| 566 | EEERRKQLEQESNDKEDRYFQQYGQGKGKQN | -45.73 | 32.3 | HHHHHHHHHHHHHHHHHHHHHHHHHCCCCCC |
| 097 | EEEERKRQEQYSNEYEDEEYQRKGRGKGHQP | -45.72 | 35.5 | HHHHHHHHHHHHHHHHHHHHHHHHCCCCCCC |
| 706 | EEERRKQLQQISQEKEDEEYQRYGKGKGHNH | -45.66 | 38.7 | HHHHHHHHHHHHHHHHHHHHHHHHCCCCCCC |
| 181 | EEEERKKQEQINNEEEDKRYQREGRGKGHQP | -45.66 | 38.7 | HHHHHHHHHHHCHHHHHHHHHHHHCCCCCCC |
| 060 | EEEKRKWLQQLSNEKEDEQYQQRGQGKGHQS | -45.65 | 38.7 | HHHHHHHHHHHHHHHHHHHHHHHHHCCCCCC |
| 454 | EEEERKYQQQYNNEWEDKRYQREGRGKGHQP | -45.65 | 38.7 | CHHHHHHHHHHHHHHHHHHHHHHHCCCCCCC |
| 132 | EEEKRKYLDQYNNDYEQREYQRKGRGQGHQP | -45.64 | 35.5 | HHHHHHHHHHHHHHHHHHHHHHHHCCCCCCC |
| 686 | EEEKRKWLDKLSNEKEQKRYQRQGRGQGHQP | -45.63 | 38.7 | HHHHHHHHHHHHHHHHHHHHHHHHCCCCCCC |

| | | | | |
|---|---|---|---|---|
| 043 | EQEKRHQEEQRNNEEEDQRYQREGQGKGHQS | -45.62 | 35.5 | HHHHHHHHHHHHHHHHHHHHHHHHCCCCCCC |
| 785 | EEEERKYQDQKNNEEEQQRYQKQGRGQGHQP | -45.62 | 35.5 | CHHHHHHHHHHCHHHHHHHHHHHHCCCCCCC |
| 106 | EEEERKYQDQYNNEYEQEEYQRKGRGQGHQP | -45.62 | 35.5 | CHHHHHHHHHHHHHHHHHHHHHHHCCCCCCC |
| 197 | EEEQKKYEDQRNNEEEQQYYQKYGQGQGHNH | -45.62 | 38.7 | CHHHHHHHHHHCHHHHHHHHHHHHHCCCCCC |
| 971 | EEEERRKKEKESNEKEDKRYQREGKGKGHSS | -45.60 | 38.7 | HHHHHHHHHHHHHHHHHHHHHHHCCCCCCCC |
| 448 | EEEERKQQEQRSNEKEQEEYQRRGRGQGHQP | -45.56 | 29.0 | HHHHHHHHHHHHHHHHHHHHHHHHHCCCCCC |
| 885 | EEEEKKKQEKYKNEYEQEEYQRKGRGQGHQP | -45.55 | 35.5 | HHHHHHHHHHHHHHHHHHHHHHHHCCCCCCC |
| 851 | EEEERKKQQQINNDKEDREYQRRGRGKGHQP | -45.55 | 35.5 | HHHHHHHHHHHHHHHHHHHHHHHHCCCCCCC |
| 475 | EEEQKKLQDQINNEQEQERYQRQGWGQGHNP | -45.55 | 38.7 | HHHHHHHHHHHHHHHHHHHHHHHCCCCCCCC |
| 295 | EEERRKQLEKINNEEEHKRKQQQGKGSGKSR | -45.54 | 38.7 | HHHHHHHHHHHHHHHHHHHHHHHCCCCCCCC |
| 892 | EEEERKRQDKYNNEYEHEEYQRKGRGQGNSN | -45.54 | 38.7 | HHHHHHHHHHHHHHHHHHHHHHHCCCCCCCC |
| 466 | EEEERKKQQQENNEKEDQRYQREGKGKGHSS | -45.54 | 38.7 | HHHHHHHHHHHHHHHHHHHHHHHCCCCCCCC |
| 870 | EEEERKKQQQINNEAEDKRYQREGKGKGHQH | -45.54 | 41.9 | HHHHHHHHHHHHHHHHHHHHHHHHCCCCCCC |
| 755 | EEEERKKQQQINNEEEDKRYQREGRGKGHQP | -45.54 | 38.7 | HHHHHHHHHHHCHHHHHHHHHHHHCCCCCCC |
| 524 | EEEERKKQQQINNEEEDQKYQREGQGKGHQS | -45.54 | 38.7 | HHHHHHHHHHHHHHHHHHHHHHHHCCCCCCC |
| 025 | EEEERKKQQQISNEKEDEEYQRRGKGKGHSS | -45.54 | 35.5 | HHHHHHHHHHHHHHHHHHHHHHHHCCCCCCC |
| 625 | EEEERKRQDKENNEREQKRYQREGQGQGHQS | -45.54 | 38.7 | HHHHHHHHHHHHHHHHHHHHHHHHCCCCCCC |
| 210 | EQERRKQLEQINNDAEDKRYQKEGKGKGDSR | -45.53 | 45.2 | HHHHHHHHHHHHHHHHHHHHHHHCCCCCCCC |
| 950 | EEEEKKFRDQYNNEYEQEEYQRKGRGQGHQP | -45.52 | 41.9 | CHHHHHHHHHHHHHHHHHHHHHHHCCCCCCC |
| 629 | EEEKRKFLDQYKNEWEQKRYQREGRGQGHQP | -45.52 | 38.7 | HHHHHHHHHHHHHHHHHHHHHHHHCCCCCCC |
| 871 | EEEQRKFEDQYRNEWEQKRYQREGWGQGHYP | -45.52 | 35.5 | CHHHHHHHHHHHHHHHHHHHHHHHCCCCCCC |
| 552 | EEEKRKMLDKWKNEWEQKYYQQKGQGQGHQP | -45.52 | 38.7 | HHHHHHHHHHHHHHHHHHHHHHHHCCCCCCC |
| 953 | EEERRKQLEQINNDAEDRRYQRYGKGKGHNH | -45.50 | 41.9 | HHHHHHHHHHHHHHHHHHHHHHHCCCCCCCC |
| 645 | EEERRKQLDAINNRKEDEEYQRRGKGKGDSR | -45.48 | 48.4 | HHHHHHHHHHHHHHHHHHHHHHHCCCCCCCC |
| 269 | EEEKRKQEEQRNNEEEQQRYQRYGRGQGHQP | -45.43 | 32.3 | HHHHHHHHHHHHHHHHHHHHHHHHHCCCCCC |
| 056 | EEEQRKQEQQYRQEWEQEEYQRKGRGQGHQP | -45.41 | 29.0 | HHHHHHHHHHHHHHHHHHHHHHHHCCCCCCC |
| 615 | EEEERKMRDQYNQEYEQQQYQKEGKGQGHSH | -45.41 | 35.5 | HHHHHHHHHHHHHHHHHHHHHHHHCCCCCCC |
| 048 | EEERRKQLEQYKNEWEDEEYQRKGRGKGHNP | -45.41 | 38.7 | HHHHHHHHHHHHHHHHHHHHHHHHCCCCCCC |
| 633 | EEEERKQQQQRSNEEEHKQYQQRGWGQGHYP | -45.41 | 29.0 | HHHHHHHHHHHHHHHHHHHHHHHCCCCCCCC |
| 681 | EEEYRKWLDQKKNEEEQERYQQQGRGQGHQP | -45.40 | 35.5 | HHHHHHHHHHHHHHHHHHHHHHHHCCCCCCC |
| 921 | EQEERKWRDQKNNEEEQQRYQRYGKGQGHSS | -45.40 | 32.3 | CHHHHHHHHHHCHHHHHHHHHHHHCCCCCCC |
| 207 | EEEERKMRQQYNNEYEDQEYQRNGRGKGHQY | -45.39 | 38.7 | HHHHHHHHHHHHHHHHHHHHHHHCCCCCCCC |
| 904 | EEEKRKMLQQYKNEYEDEEYQRKGRGKGHQP | -45.39 | 38.7 | HHHHHHHHHHHHHHHHHHHHHHHHCCCCCCC |
| 427 | EEERRKQLDQEKQEKEQEEYQRRGKGQGHSS | -45.39 | 35.5 | HHHHHHHHHHHHHHHHHHHHHHHCCCCCCCC |
| 124 | EEERKKQLQQKNNEEEDKRYQREGKGKGHNH | -45.37 | 45.2 | HHHHHHHHHHHHHHHHHHHHHHHHCCCCCCC |
| 099 | EEERRKQEQQLRNEKEDSRYQQQGWGKGHNP | -45.37 | 35.5 | HHHHHHHHHHHHHHHHHHHHHHHCCCCCCCC |
| 111 | EEERRKQLQQIKNEKEDEEYQRRGQGKGHQS | -45.37 | 38.7 | HHHHHHHHHHHHHHHHHHHHHHHHHCCCCCC |
| 551 | EEERRKQLQQIKNEKEDQEYQKQGRGKGHQP | -45.37 | 38.7 | HHHHHHHHHHHHHHHHHHHHHHHHCCCCCCC |

| | | | | |
|---|---|---|---|---|
| 643 | EEERRKQLQQINNEEEDKRYQREGMGKGHQP | -45.37 | 41.9 | HHHHHHHHHHHHHHHHHHHHHHHHCCCCCCC |
| 042 | EEERRKQLQQWKNEWEDEYYQQYGQGKGHSH | -45.37 | 38.7 | HHHHHHHHHHHHHHHHHHHHHHHHHCCCCCC |
| 995 | EEERRKQLQQYKNEWEDKRYQREGQGKGHQS | -45.37 | 38.7 | HHHHHHHHHHHHHHHHHHHHHHHHCCCCCCC |
| 708 | EEEERKQRQQESNEKEDEEYQRKGRGKGHQP | -45.29 | 35.5 | HHHHHHHHHHHHHHHHHHHHHHHHCCCCCCC |
| 965 | EEEQRKQHQQESNEKEDKRYQREGRGKGHQP | -45.29 | 35.5 | HHHHHHHHHHHHHHHHHHHHHHHHCCCCCCC |
| 703 | EEEKRKMLDQYKHEWEQEEYQRKGRGQGHQP | -45.28 | 38.7 | HHHHHHHHHHHHHHHHHHHHHHHHCCCCCCC |
| 194 | EEEERKKQDQIKNEEEQKRYQRYGKGQGHSS | -45.28 | 32.3 | HHHHHHHHHHHHHHHHHHHHHHHHCCCCCCC |
| 291 | EEEERKKQDQIKNEKEQEEYQRYGWGQGHYP | -45.28 | 32.3 | HHHHHHHHHHHHHHHHHHHHHHHCCCCCCCC |
| 254 | EEEERKKQDQINNEKEQEEYQRRGWGQGHNP | -45.28 | 35.5 | HHHHHHHHHHHHHHHHHHHHHHHCCCCCCCC |
| 138 | EEEERKKQDQINNEQEQERYQQQGRGQGHQP | -45.28 | 35.5 | HHHHHHHHHHHHHHHHHHHHHHHHCCCCCCC |
| 238 | EEERRKQLDQINHREEQERYQREGRGQGHQP | -45.27 | 38.7 | HHHHHHHHHHHHHHHHHHHHHHHHHCCCCCC |
| 212 | EEEERKKQQKYKNEWEQKRYQREGRGQGHQP | -45.24 | 32.3 | HHHHHHHHHHHHHHHHHHHHHHHHCCCCCCC |
| 919 | EEEQRKYEDDRNNREEDQRYQRYGRGKGHQP | -45.20 | 38.7 | HHHHHHHHHHHHHHHHHHHHHHHHCCCCCCC |
| 616 | EEEKRKQEEQRNNEEEQQRYQRQGWGQGHNP | -45.15 | 32.3 | HHHHHHHHHHHHHHHHHHHHHHHCCCCCCCC |
| 586 | EEEERKYQDQKNNEEEQKRYQRYGWGQGHNP | -45.15 | 35.5 | CHHHHHHHHHHCHHHHHHHHHHHCCCCCCCC |
| 351 | DEERKKMEDQFRNEYEQQSYQREGKGQGHSS | -45.13 | 35.5 | HHHHHHHHHHHHHHHHHHHHHHCCCCCCCCC |
| 671 | EEEERKMRDQYNNEYEQEEYQRKGRGQGHQP | -45.13 | 35.5 | HHHHHHHHHHHHHHHHHHHHHHHHCCCCCCC |
| 926 | EEEEKKKREKYNNEAEQKRYQRYGQGQGHQS | -45.11 | 41.9 | CHHHHHHHHHHCHHHHHHHHHHHHHCCCCCC |
| 872 | EEEEKKIQEQISNEKEHEEYQRRGKGQGNSN | -45.11 | 32.3 | CHHHHHHHHHHHHHHHHHHHHHHCCCCCCCC |
| 956 | EEERRKQLDQEKNEREQQYYQREGKGQGHNH | -45.11 | 35.5 | HHHHHHHHHHHHHHHHHHHHHHHHCCCCCCC |
| 948 | EEERRKQLDQIKNEQEQQRYQREGKGQGHSS | -45.11 | 35.5 | HHHHHHHHHHHHHHHHHHHHHHHCCCCCCCC |
| 428 | EEERRKQLDQINNEAEQKRYQRYGKGQGHNH | -45.11 | 41.9 | HHHHHHHHHHHHHHHHHHHHHHHHCCCCCCC |
| 873 | EEERRKQLDQYNNEWEQEEYQRKGRGQGHQP | -45.11 | 38.7 | HHHHHHHHHHHHHHHHHHHHHHHHCCCCCCC |
| 888 | EEEKRKFLDDYNNRSEDQRYQRYGQGKGHQS | -45.10 | 45.2 | HHHHHHHHHHHHCCCHHHHHHHHHHCCCCCC |
| 709 | EEEQRKYEDKRNNEEEQKRYQRYGKGQGDSR | -45.10 | 45.2 | CHHHHHHHHHHCHHHHHHHHHHHCCCCCCCC |
| 260 | EQEKRKLLEKISNEKEQKEYQKQGWGQGHNP | -45.09 | 32.3 | HHHHHHHHHHHHHHHHHHHHHHHCCCCCCCC |
| 134 | EEEEKKKRDQENNEKEQQRYQREGQGQGHQS | -45.09 | 38.7 | CHHHHHHHHHHHHHHHHHHHHHHHCCCCCCC |
| 539 | EEEERKRQQQESNEREDKRYQREGQGKGHSS | -45.08 | 35.5 | HHHHHHHHHHHHHHHHHHHHHHHCCCCCCCC |
| 849 | EQEERKWREQKNNEEEDKRYQREGKGKGDSR | -45.05 | 41.9 | CHHHHHHHHHHHHHHHHHHHHHHCCCCCCCC |
| 018 | EQEERHYQEKKNNEEEDKRYQRYGQGKGHQS | -45.05 | 38.7 | CHHHHHHHHHHHHHHHHHHHHHHHHCCCCCC |
| 869 | EEEEKKQQEQRSNDQEQRRYQREGRGQGHQP | -45.05 | 29.0 | HHHHHHHHHHHHHHHHHHHHHHHHCCCCCCC |
| 409 | EQEQRHQQEQYNNEYEDQEYQRKGRGKGDSR | -45.03 | 41.9 | HHHHHHHHHHHHHHHHHHHHHHHCCCCCCCC |
| 889 | EEEEKKQREQYNNEYEQEEYQRKGRGQGHQP | -45.02 | 35.5 | CHHHHHHHHHHHHHHHHHHHHHHHCCCCCCC |
| 120 | EEERKKQLEQYKNENEQKQYQRSGRGQGHQP | -45.02 | 38.7 | HHHHHHHHHHHHHHHHHHHHHHHCCCCCCCC |
| 017 | EQEERHKQEQKNNEEEDERYQQQGKGKGHSS | -45.02 | 35.5 | HHHHHHHHHHHHHHHHHHHHHHHCCCCCCCC |
| 635 | EQEERHKQEQYNNEYEDEYYQQKGQGKGHSH | -45.02 | 35.5 | HHHHHHHHHHHHHHHHHHHHHHHHCCCCCCC |
| 314 | EEEERKQQDHRKNRKEQEEYQRRGRGQGHQP | -45.01 | 29.0 | HHHHHHHHHHHHHHHHHHHHHHHHHCCCCCC |
| 695 | QDEQRKYEDQRNNEEEQQRYQQYGKGQGHNN | -45.00 | 29.0 | CHHHHHHHHHHCHHHHHHHHHHHHHCCCCCC |

| | | | | |
|---|---|---|---|---|
| 723 | EEQERKFQDQYNNEFEQKYYQQQGQGQGHSH | -45.00 | 41.9 | CHHHHHHHHHHHHHHHHHHHHHHHCCCCCCC |
| 532 | EEEERKQRDHESNRKEDQRYQRQGKGKGHSH | -44.99 | 35.5 | HHHHHHHHHHHHHHHHHHHHHHHCCCCCCCC |
| 955 | EEEKKKMLQQIEDDKRDEEYQRRGWGKGHNP | -44.99 | 35.5 | HHHHHHHHHHHHHHHHHHHHHHHCCCCCCCC |
| 253 | EEEERKRQEQENNEKEDQRYQREGKGKGDSR | -44.90 | 45.2 | HHHHHHHHHHHHHHHHHHHHHHHCCCCCCCC |
| 307 | EEQQRRLYEKISNEKEDEEYQRRGRGKGHQP | -44.90 | 41.9 | HHHHHHHHHHHHHHHHHHHHHHHHCCCCCCC |
| 433 | EEERRKQLDQIKNEKEQEEYQRYGKGQGHNH | -44.88 | 35.5 | HHHHHHHHHHHHHHHHHHHHHHHHHCCCCCC |
| 179 | EEERRKQLDQKNNEEEQERYQQQGRGQGHQP | -44.88 | 38.7 | HHHHHHHHHHHCHHHHHHHHHHHHCCCCCCC |
| 187 | EEEERRYQEKYNNEWEDKRYQREGQGKGHQS | -44.88 | 41.9 | CHHHHHHHHHHHHHHHHHHHHHHHCCCCCCC |
| 711 | EEEERKKQDDYRNRWEDEEYQRKGRGKGHQP | -44.86 | 35.5 | HHHHHHHHHHHHHHHHHHHHHHHHCCCCCCC |
| 811 | EEQKRKQLQQESNEKEDKRYQRYGRGKGHQP | -44.85 | 41.9 | HHHHHHHHHHHHHHHHHHHHHHHHCCCCCCC |
| 894 | EEEQKKYEEQRSNEEEHKRYQRYGWGQGHNP | -44.80 | 32.3 | HHHHHHHHHHHHHHHHHHHHHHHCCCCCCCC |
| 241 | EEEERKFRQAFNNRFEDQYFQQQGQGKGKQS | -44.75 | 38.7 | CHHHHHHHHHHHHHHHHHHHHHHHCCCCCCC |
| 748 | EQEERHKQEKYNNEAEHKRKQREGKGSGKNH | -44.75 | 32.3 | HHHHHHHHHHHHHHHHHHHHHHHCCCCCCCC |
| 947 | EEEKRKLLQKIKQEKEQKEYQRNGSGQGHQY | -44.75 | 35.5 | HHHHHHHHHHHHHHHHHHHHHHHCCCCCCCC |
| 105 | EQEERHLQEQQSNEKEDEKYQQEGMGKGHQP | -44.75 | 32.3 | HHHHHHHHHHHHHHHHHHHHHHHHCCCCCCC |
| 797 | EQERRKQLEAKNNREEDERYQQQGWGKGHNP | -44.73 | 35.5 | HHHHHHHHHHHHHHHHHHHHHHHCCCCCCCC |
| 912 | EEEEKKYQDQYNNEYEQEEYQRKGRGQGHQP | -44.73 | 38.7 | CHHHHHHHHHHHHHHHHHHHHHHHCCCCCCC |
| 226 | EEERKKQLDQFNQEFEQQYYQQKGQGQGHNH | -44.71 | 45.2 | HHHHHHHHHHHHHHHHHHHHHHHHCCCCCCC |
| 999 | EEEEKKQQEKRNNEKEQRRYQRYGRGQGHQP | -44.71 | 38.7 | HHHHHHHHHHHHHHHHHHHHHHHHHCCCCCC |
| 515 | EQEQRHQEEQERNDKEDKRFQREGKGKGKNH | -44.70 | 25.8 | HHHHHHHHHHHHHHHHHHHHHHHHCCCCCCC |
| 684 | EEEQRKYEEQREEEERDERYQQQGRGKGHQP | -44.69 | 32.3 | HHHHHHHHHHHHHHHHHHHHHHHCCCCCCCC |
| 046 | EQEERKKQEQESNEKEQKRYQREGKGQGHSH | -44.69 | 25.8 | HHHHHHHHHHHHHHHHHHHHHHHCCCCCCCC |
| 857 | EEERKKQLDDYNNRWEDQRYQREGRGKGHQP | -44.69 | 45.2 | HHHHHHHHHHHHHHHHHHHHHHHHCCCCCCC |
| 751 | EEEERKMRDQYNNEHEQSRYQQQGRGQGHQP | -44.66 | 35.5 | HHHHHHHHHHHHHHHHHHHHHHHHCCCCCCC |
| 922 | EEEKKKMLDQINNEAEQKRYQREGKGQGHSH | -44.66 | 45.2 | HHHHHHHHHHHHHHHHHHHHHHHHCCCCCCC |
| 162 | EEEQRKEEDQRRNEKEQKRYQREGKGQGHNH | -44.66 | 32.3 | HHHHHHHHHHHHHHHHHHHHHHHHCCCCCCC |
| 663 | EEEKRKYEQQRNNEAEDKRYQREGKGKGDSR | -44.64 | 48.4 | HHHHHHHHHHHHHHHHHHHHHHHCCCCCCCC |
| 223 | EEERRKQLDQIKNEEEQQRYQRYGMGQGHQP | -44.63 | 35.5 | HHHHHHHHHHHHHHHHHHHHHHHCCCCCCCC |
| 030 | EEERRKQLDQIKNEKEQKEYQRSGRGQGHQP | -44.63 | 38.7 | HHHHHHHHHHHHHHHHHHHHHHHCCCCCCCC |
| 980 | EEEERKYQEKYNNEAEQKRYQRYGKGQGHNH | -44.63 | 38.7 | CHHHHHHHHHHHHHHHHHHHHHHHCCCCCCC |
| 724 | EEEERKKQDQESNRKEQERYQRQGRGQGHQP | -44.63 | 29.0 | HHHHHHHHHHHHHHHHHHHHHHHHCCCCCCC |
| 155 | EEEERKKQEKINNEAEHKRYQREGKGQGHSS | -44.63 | 38.7 | HHHHHHHHHHHHHHHHHHHHHHHCCCCCCCC |
| 052 | EQERRHQLEQYKNEYEDEEYQRKGRGKGNQN | -44.60 | 35.5 | HHHHHHHHHHHHHHHHHHHHHHHHCCCCCCC |
| 957 | EEERRKQLQQWKNDWEQRYYQQKGQGQGHSH | -44.59 | 29.0 | HHHHHHHHHHHHHHHHHHHHHHHHCCCCCCC |
| 582 | EQEKRHMLEKYKNEYEDEEYQRKGRGKGHQP | -44.56 | 38.7 | HHHHHHHHHHHHHHHHHHHHHHHHCCCCCCC |
| 585 | EQEERKKQEQQNNEREQEEYQRRGRGQGHQP | -44.55 | 29.0 | HHHHHHHHHHHHHHHHHHHHHHHHHCCCCCC |
| 789 | EEERKKQLDDYKNRWEDEEYQRKGRGKGHQP | -44.54 | 41.9 | HHHHHHHHHHHHHHHHHHHHHHHHCCCCCCC |
| 316 | EQERRHQLEKYKNEYEDEEYQRKGRGKGHQP | -44.53 | 38.7 | HHHHHHHHHHHHHHHHHHHHHHHHCCCCCCC |

| | | | | |
|---|---|---|---|---|
| 554 | EEEQRRQEEKRRNEKEQKYYQQKGQGQGHNH | -44.53 | 32.3 | HHHHHHHHHHHHHHHHHHHHHHHHCCCCCCC |
| 787 | EEEERKKEEQWRNEWEQQMYQRYGKGQGHQP | -44.50 | 29.0 | HHHHHHHHHHHHHHHHHHHHHHHHHCCCCCC |
| 996 | EEEKRKLLQQIEERKRDEEYQRRGRGKGHQP | -44.49 | 32.3 | HHHHHHHHHHHHHHHHHHHHHHHHCCCCCCC |
| 158 | EEQERRKQEQYNNRWEDERYQREGWGKGHNP | -44.45 | 38.7 | HHHHHHHHHHHHHHHHHHHHHHHHCCCCCCC |
| 640 | EQERRHLLEQISNEKEDEEYQRRGQGKGHQS | -44.43 | 35.5 | HHHHHHHHHHHHHHHHHHHHHHHHHCCCCCC |
| 531 | EEEERKWREKKNNEEEQKRYQRYGRGQGHQP | -44.40 | 35.5 | CHHHHHHHHHHCHHHHHHHHHHHHHCCCCCC |
| 533 | EEEARKSEQKYRNEAEQKRYQREGKGQGHNH | -44.39 | 38.7 | HHHHHHHHHHHHHHHHHHHHHHHHCCCCCCC |
| 806 | EEEQRKQEQQYRNDSEQRRYQRYGQGQGHQS | -44.39 | 25.8 | HHHHHHHHHHHHHHHHHHHHHHHHHCCCCCC |
| 925 | EEEKRKQEDDRNNREEQQRYQRYGRGQGHQP | -44.38 | 32.3 | HHHHHHHHHHHHHHHHHHHHHHHHHCCCCCC |
| 172 | EEEKRKQEQQRNNEKEQEEYQRRGQGQGHQS | -44.38 | 32.3 | HHHHHHHHHHHHHHHHHHHHHHHHHCCCCCC |
| 364 | EEEERKKEQQREEERRDEEYQRRGWGKGHNP | -44.37 | 32.3 | HHHHHHHHHHHHHHHHHHHHHHHCCCCCCCC |
| 397 | EEEQRKQQQQYSNRFEQEYYQQKGQGQGHSH | -44.35 | 25.8 | HHHHHHHHHHHHHHHHHHHHHHHHCCCCCCC |
| 911 | EEEQRKQQQQYSNESEQERYQQQGWGQGHNP | -44.31 | 29.0 | HHHHHHHHHHHCHHHHHHHHHHHCCCCCCCC |
| 326 | EQEQRHYEEQRNNEEEDQRYQRYGKGKGHSS | -44.28 | 35.5 | HHHHHHHHHHHHHHHHHHHHHHHHCCCCCCC |
| 265 | EEEEKKRQEKESNEKEQEEYQRKGRGQGHQP | -44.25 | 35.5 | HHHHHHHHHHHHHHHHHHHHHHHHCCCCCCC |
| 493 | EEQRRKQLDDIKNRKEDEEYQRRGWGKGHNP | -44.25 | 41.9 | HHHHHHHHHHHHHHHHHHHHHHHCCCCCCCC |
| 452 | EQEERHQQEQYNNESEHERYQQQGQGQGHSS | -44.23 | 29.0 | CHHHHHHHHHHCHHHHHHHHHHHCCCCCCCC |
| 057 | EEERRRQLEKIKNEKEDEEYQRYGKGKGHSN | -44.21 | 41.9 | HHHHHHHHHHHHHHHHHHHHHHHCCCCCCCC |
| 578 | EEERRKQLDQESNEKEQKRYQREGQGQGHSS | -44.21 | 35.5 | HHHHHHHHHHHHHHHHHHHHHHHCCCCCCCC |
| 504 | EEEQKKMYEKINNEKEQEEYQRRGRGQGHQP | -44.20 | 38.7 | HHHHHHHHHHHHHHHHHHHHHHHHHCCCCCC |
| 654 | EEERRKQLQKISNEEEHKRYQREGKGQGHSH | -44.18 | 35.5 | HHHHHHHHHHHCHHHHHHHHHHHHCCCCCCC |
| 547 | EEERRKQLEQISNEEEHKRKQREGKGSGKSR | -44.18 | 32.3 | HHHHHHHHHHHHHHHHHHHHHHHCCCCCCCC |
| 624 | EEEKKKMLEKINNEAEQKRYQREGRGQGHQP | -44.13 | 45.2 | HHHHHHHHHHHHHHHHHHHHHHHHCCCCCCC |
| 310 | EEEERRYQEQYNNDYEDREYQRKGRGKGHQP | -44.12 | 35.5 | CHHHHHHHHHHHCHHHHHHHHHHHCCCCCCC |
| 852 | EEERRKQLQQESNEKEHEQKQQRGQGSGKQS | -44.10 | 29.0 | HHHHHHHHHHHHHHHHHHHHHHHCCCCCCCC |
| 171 | EEEEKKKREHYNNRWEDERYQQQGQGKGHQS | -44.07 | 38.7 | HHHHHHHHHHHHHHHHHHHHHHHHCCCCCCC |
| 895 | EEEERKFQQQFNNEFEHEYYQQKGQGQGHNH | -44.06 | 38.7 | CHHHHHHHHHHHHHHHHHHHHHHHCCCCCCC |
| 730 | EQERRHQLEQIKQEKEDEEYQRRGRGKGHQP | -44.06 | 35.5 | HHHHHHHHHHHHHHHHHHHHHHHHCCCCCCC |
| 906 | EQERRHQLEQWKQEWEDEYYQRKGQGKGHNH | -44.06 | 35.5 | HHHHHHHHHHHHHHHHHHHHHHHHCCCCCCC |
| 858 | EQEERHWQEQMKNEKEDEEYQRRGWGKGHNP | -44.06 | 32.3 | CHHHHHHHHHHHHHHHHHHHHHHCCCCCCCC |
| 151 | EEEQRKYEEKRNNEEEQKRYQREGWGQGHNP | -44.03 | 35.5 | CHHHHHHHHHHHHHHHHHHHHHHHCCCCCCC |
| 987 | EQEKRKLLEQIENEREQQEYQREGRGQGHQP | -44.01 | 29.0 | HHHHHHHHHHHHHHHHHHHHHHHHCCCCCCC |
| 544 | EEEKRKLLQQIKNEEEQKRFQRYGRGQGKQN | -44.01 | 29.0 | HHHHHHHHHHHHHHHHHHHHHHHHHCCCCCC |
| 721 | EEEERKKEQKYRNEAEQKRYQREGKGQGHSS | -43.97 | 35.5 | HHHHHHHHHHHHHHHHHHHHHHHCCCCCCCC |
| 109 | EEERRKQLQKESQEKEQKRYQREGRGQGHQP | -43.95 | 35.5 | HHHHHHHHHHHHHHHHHHHHHHHHCCCCCCC |
| 537 | EQEQRKEEEKKRNEKEQKRYQREGRGQGHQP | -43.93 | 29.0 | HHHHHHHHHHHHHHHHHHHHHHHHCCCCCCC |
| 236 | EEEERKKQEQYNNEYEQEEYQRKGRGQGHQP | -43.90 | 32.3 | HHHHHHHHHHHHHHHHHHHHHHHHCCCCCCC |
| 481 | EEEQRKQQQQQSNEEEQQRYQRYGKGQGDSR | -43.88 | 35.5 | HHHHHHHHHHHHHHHHHHHHHHHCCCCCCCC |

| | | | | |
|---|---|---|---|---|
| 714 | EEEERKKQQQYNNEYEQEEYQRQGRGQGHQP | -43.86 | 32.3 | HHHHHHHHHHHHHHHHHHHHHHHHCCCCCCC |
| 958 | EEQERKKQQQINNEKEDEEYQRRGKGKGDSR | -43.86 | 48.4 | HHHHHHHHHHHHHHHHHHHHHHHCCCCCCCC |
| 836 | EQEERKYQEQFNNRFEQEYYQQKGQGQGHQP | -43.84 | 29.0 | CHHHHHHHHHHHHHHHHHHHHHHHCCCCCCC |
| 485 | EEERRKQLQAEKNRKEDQRYQREGKGKGHSH | -43.83 | 35.5 | HHHHHHHHHHHHHHHHHHHHHHHCCCCCCCC |
| 756 | EEEERKKQQQINNEKEHQEYQKQGQGQGHQS | -43.82 | 32.3 | HHHHHHHHHHHHHHHHHHHHHHHCCCCCCCC |
| 275 | EQEKRHMLEQKNNEEEDKRYQREGRGKGHQP | -43.79 | 38.7 | HHHHHHHHHHHHHHHHHHHHHHHHCCCCCCC |
| 569 | EQERRHQLEQESNDKEDRRYQREGRGKGHQP | -43.78 | 32.3 | HHHHHHHHHHHHCCHHHHHHHHHHCCCCCCC |
| 916 | EEEQRKFEEQYRNDAEQRRYQREGRGQGHQP | -43.77 | 32.3 | HHHHHHHHHHHHHHHHHHHHHHHHCCCCCCC |
| 324 | EQEKRHMLEQINNRAEDERYQREGKGKGHNH | -43.77 | 38.7 | HHHHHHHHHHHHHHHHHHHHHHHCCCCCCCC |
| 280 | EQERRHQLEQQSNEEEDERYQQQGWGKGHNP | -43.77 | 35.5 | HHHHHHHHHHHHHHHHHHHHHHHCCCCCCCC |
| 694 | EQERRHQLEQYKNEWEDEEYQRKGRGKGHQP | -43.77 | 35.5 | HHHHHHHHHHHHHHHHHHHHHHHHCCCCCCC |
| 248 | EQERRHQLEQYSNEYEDKQYQKEGKGKGHNH | -43.77 | 35.5 | HHHHHHHHHHHHHHHHHHHHHHHCCCCCCCC |
| 546 | EQEERHKQEQFSNRFEDEYYQKYGQGKGHSH | -43.74 | 32.3 | HHHHHHHHHHHHHHHHHHHHHHHHHCCCCCC |
| 344 | EEEQKKQEEQYRNEYEQEEYQRKGRGQGDSR | -43.71 | 38.7 | HHHHHHHHHHHHHHHHHHHHHHHCCCCCCCC |
| 834 | EEEQRKLYQQLNNEKEQKRYQREGRGQGHQP | -43.69 | 32.3 | HHHHHHHHHHHHHHHHHHHHHHHHCCCCCCC |
| 088 | EEERRKLLQQISNEKEQEEYQRRGWGQGHYP | -43.69 | 32.3 | HHHHHHHHHHHHHHHHHHHHHHHCCCCCCCC |
| 354 | EQEERHQREQQSNEREQKRYQREGRGQGHQP | -43.67 | 25.8 | CHHHHHHHHHHHHHHHHHHHHHHHCCCCCCC |
| 410 | EQERRHQLEQISNEEEDRRYQREGKGKGHSH | -43.64 | 35.5 | HHHHHHHHHHHHHHHHHHHHHHHCCCCCCCC |
| 523 | EEERKKQLEQIKQEAEQQRYQREGKGQGHSS | -43.64 | 38.7 | HHHHHHHHHHHHHHHHHHHHHHHCCCCCCCC |
| 700 | EEERKKQLEKIKNEKEQKEYQRSGRGQGHQP | -43.64 | 41.9 | HHHHHHHHHHHHHHHHHHHHHHHCCCCCCCC |
| 330 | EQERRHQLEQISNEKEDERYQQQGRGKGHQP | -43.62 | 35.5 | HHHHHHHHHHHHHHHHHHHHHHHHCCCCCCC |
| 023 | EQQQRHQEEQYRNEEEQQRYQRQGRGQGHQP | -43.61 | 29.0 | HHHHHHHHHHHHHHHHHHHHHHHHHCCCCCC |
| 961 | EQEQRHQEEQWRNEYEQEEYQRKGRGQGHQP | -43.61 | 25.8 | HHHHHHHHHHHHHHHHHHHHHHHHCCCCCCC |
| 298 | EEEERKRQDQENNEREQKRYQREGKGQGDSR | -43.58 | 41.9 | HHHHHHHHHHHHHHHHHHHHHHHCCCCCCCC |
| 768 | EEEKRRQLEQESNDKEDRRYQREGWGKGHNP | -43.53 | 35.5 | HHHHHHHHHHHHHHHHHHHHHHHHCCCCCCC |
| 704 | EQQERHKQEQINNEKEDEEYQRRGRGKGHQP | -43.50 | 38.7 | CHHHHHHHHHHHHHHHHHHHHHHHCCCCCCC |
| 642 | EEEERKMRQKFNNEYEQKSYQREGRGQGHQP | -43.50 | 38.7 | HHHHHHHHHHHHHHHHHHHHHHHCCCCCCCC |
| 024 | EEEKRKMLQKKNNEEEQKRYQRYGRGQGHQP | -43.50 | 38.7 | HHHHHHHHHHHCCHHHHHHHHHHHHCCCCCC |
| 628 | EQEERHRLEQMKNEKEDEEYQRRGRGKGHQP | -43.48 | 35.5 | HHHHHHHHHHHHHHHHHHHHHHHHCCCCCCC |
| 449 | EEEQRKQQEQYSEEWRQKRYQREGRGQGHQP | -43.40 | 25.8 | HHHHHHHHHHHHHHHHHHHHHHHHCCCCCCC |
| 974 | EEEERKMREQYNNDYEQRQYQQQGRGQGHQP | -43.38 | 29.0 | HHHHHHHHHHHHHHHHHHHHHHHHCCCCCCC |
| 059 | EQEARHQQEQYSNDAEQRRYQREGRGQGHQP | -43.36 | 29.0 | HHHHHHHHHHHCHHHHHHHHHHHHCCCCCCC |
| 317 | EEERKKQLEQYKNEWEQEEYQRKGRGQGHQP | -43.34 | 35.5 | HHHHHHHHHHHHHHHHHHHHHHHHCCCCCCC |
| 819 | EEEERRRQEQYNNEYEDEEYQRKGRGKGHQP | -43.31 | 38.7 | HHHHHHHHHHHHHHHHHHHHHHHHCCCCCCC |
| 053 | EEEERKYQEDYNNREEDQKYQREGRGKGHQP | -43.29 | 35.5 | HHHHHHHHHHHHHHHHHHHHHHHHCCCCCCC |
| 403 | EQEERHQREQYNNEYEQQQYQKEGRGQGHQP | -43.29 | 29.0 | CHHHHHHHHHHHHHHHHHHHHHHHCCCCCCC |
| 491 | EQEKRHMLEQISNEKEDEEYQRYGRGKGHQP | -43.27 | 35.5 | HHHHHHHHHHHHHHHHHHHHHHHHHCCCCCC |
| 803 | EEEQRKYEQQRNNDEEQRRYQRYGWGQGHNP | -43.24 | 29.0 | HHHHHHHHHHHHHHHHHHHHHHHCCCCCCCC |

| | | | | |
|---|---|---|---|---|
| 773 | EQEARHNQEDYNQRSEDERYQQQGRGKGHQP | -43.24 | 35.5 | HHHHHHHHHHHHHHHHHHHHHHHHCCCCCCC |
| 411 | EEEERKYQQQFKNEFEQKYYQQKGQGQGHNH | -43.22 | 32.3 | CHHHHHHHHHHHHHHHHHHHHHHHCCCCCCC |
| 772 | EEEEKKFQQKYNHEYEQEEYQRKGRGQGHQP | -43.18 | 45.2 | CHHHHHHHHHHHHHHHHHHHHHHHCCCCCCC |
| 033 | EQEQRHYEEQRSNDEEHRRKQREGRGSGKQN | -43.10 | 19.4 | HHHHHHHHHHHHHHHHHHHHHHHCCCCCCCC |
| 358 | EQEERHKQEQYSNEYEDEEYQRKGRGKGHSN | -43.09 | 32.3 | HHHHHHHHHHHHHHHHHHHHHHHCCCCCCCC |
| 518 | EEEEKKREEQYRNEWEQKRYQREGKGQGHNH | -43.06 | 32.3 | HHHHHHHHHHHHHHHHHHHHHHHHCCCCCCC |
| 973 | EEEEKKRQEQQSNEEEQQRYQRYGWGQGHYP | -43.06 | 32.3 | HHHHHHHHHHHHHHHHHHHHHHHCCCCCCCC |
| 831 | EEERRKQLQQESQEKEQQRYQREGKGQGHNH | -43.00 | 32.3 | HHHHHHHHHHHHHHHHHHHHHHHHCCCCCCC |
| 816 | EEEERKKEQQYRNEYEQKQYQQQGQGQGHQS | -43.00 | 29.0 | HHHHHHHHHHHHHHHHHHHHHHHHCCCCCCC |
| 131 | EEEEKKYQQQFNHEYEQQSYQREGRGQGHQP | -42.98 | 41.9 | CHHHHHHHHHHHHHHHHHHHHHHCCCCCCCC |
| 675 | EEEKKKMLEQYKNEWEQEEYQRKGRGQGHQP | -42.92 | 35.5 | HHHHHHHHHHHHHHHHHHHHHHHCCCCCCCC |
| 495 | EEEERKKMEQMKNEAEQQSYQQYGRGQGHQP | -42.92 | 32.3 | CHHHHHHHHHHHHHHHHHHHHHHHHCCCCCC |
| 267 | EEEERKMREQFNNEYEQQRYQQEGQGQGHQS | -42.89 | 35.5 | HHHHHHHHHHHHHHHHHHHHHHHHCCCCCCC |
| 335 | EQERRKQLEQYKNEWEQEYYQRKGQGQGHNH | -42.87 | 29.0 | HHHHHHHHHHHHHHHHHHHHHHHHCCCCCCC |
| 679 | EEEERKKRQAYNNRSEQQRYQRYGRGQGHQP | -42.86 | 29.0 | HHHHHHHHHHHHHHHHHHHHHHHHHCCCCCC |
| 422 | EEEEKKQQEDRSNRKEQEEYQRRGRGQGHQP | -42.79 | 29.0 | HHHHHHHHHHHHHHHHHHHHHHHHHCCCCCC |
| 137 | EEEEKKKREQINNEEEQQRYQRYGRGQGHQP | -42.73 | 35.5 | HHHHHHHHHHHCHHHHHHHHHHHHHCCCCCC |
| 277 | EEERRKQEQQRNNEEEQQRYQRYGQGQGHQS | -42.71 | 32.3 | HHHHHHHHHHHHHHHHHHHHHHHHHCCCCCC |
| 003 | EEERRKQLQQESNEKEQKRYQREGKGQGHQP | -42.71 | 32.3 | HHHHHHHHHHHHHHHHHHHHHHHHCCCCCCC |
| 080 | EEEKKRQEEQRSNDEEQRRYQREGQGQGHQS | -42.67 | 29.0 | HHHHHHHHHHHCHHHHHHHHHHHHCCCCCCC |
| 251 | EQERRHQLEQYKNESEDERYQQQGKGKGDSR | -42.52 | 41.9 | HHHHHHHHHHHHHHHHHHHHHHHCCCCCCCC |
| 788 | EEERRKQLQHYKNRWEQEEYQRKGRGQGHQP | -42.52 | 29.0 | HHHHHHHHHHHHHHHHHHHHHHHHCCCCCCC |
| 035 | EQERRHQLEQIKNDKEHREKQRNGSGSGKQY | -42.51 | 22.6 | HHHHHHHHHHHHHHHHHHHHHHCCCCCCCCC |
| 627 | EQEQRKQQEAYSERYRQEEYQRKGRGQGHQP | -42.43 | 19.4 | HHHHHHHHHHHHHHHHHHHHHHHHCCCCCCC |
| 272 | EEEERKRQEQESNRKEQERYQREGRGQGHQP | -42.39 | 25.8 | HHHHHHHHHHHHHHHHHHHHHHHHCCCCCCC |
| 160 | EQEERHNQEQYNNESEQQRYQRYGRGQGNQN | -42.38 | 29.0 | CHHHHHHHHHHCCHHHHHHHHHHHCCCCCCC |
| 538 | EQEERHKKEQYDNEWEQEKYQKEGRGQGHQP | -42.36 | 25.8 | CHHHHHHHHHHHHHHHHHHHHHHHCCCCCCC |
| 402 | EEEQRKYEQQRNNEEEQEEYQRKGRGQGNQN | -42.29 | 32.3 | HHHHHHHHHHHHHHHHHHHHHHHHCCCCCCC |
| 079 | EQEERHKQEQIKNEKEDEEYQRRGKGKGDSR | -42.18 | 38.7 | HHHHHHHHHHHHHHHHHHHHHHHCCCCCCCC |
| 228 | EEEERKMRDDYNNRYEQEEYQRKGRGQGHQP | -42.05 | 32.3 | HHHHHHHHHHHHHHHHHHHHHHHHCCCCCCC |
| 864 | EQSERHKQEKWKNEWEQKYYQQQGRGQGHQP | -41.89 | 29.0 | CHHHHHHHHHHHHHHHHHHHHHHHCCCCCCC |
| 116 | EEQERKARQQFNNEFEQQYYQKYGQGQGHNH | -41.85 | 38.7 | HHHHHHHHHHHHHHHHHHHHHHHHHCCCCCC |
| 299 | EEEKRKYLQDYNNRWEDQRYQREGRGKGHQP | -41.81 | 38.7 | HHHHHHHHHHHHHHHHHHHHHHHHCCCCCCC |
| 446 | EQEQRHYEEQRNNEEEQQRYQREGWGQGHNP | -41.62 | 29.0 | HHHHHHHHHHHHHHHHHHHHHHHHCCCCCCC |
| 593 | EEERKKQLEDYNNRSEQERYQQQGQGQGHQS | -41.58 | 35.5 | HHHHHHHHHHHHHHHHHHHHHHHHHCCCCCC |
| 271 | EEEERKRQQQESNEKEQKRYQREGRGQGNQN | -41.48 | 29.0 | HHHHHHHHHHHHHHHHHHHHHHHHCCCCCCC |
| 910 | EQERRHQLEKINNEAEQKRYQRYGRGQGHQP | -41.24 | 38.7 | HHHHHHHHHHHHHHHHHHHHHHHHHCCCCCC |
| 178 | EQQQRHYEEQRSNDEEQRYYQQQGQGQGHNH | -41.20 | 25.8 | HHHHHHHHHHHHHHHHHHHHHHHHCCCCCCC |

| | | | | |
|---|---|---|---|---|
| 673 | EQEERHMREQYNNEYEQEEYQQQGRGQGHQP | -41.13 | 29.0 | HHHHHHHHHHHHHHHHHHHHHHHHCCCCCCC |
| 349 | EQERRHQLEQINNDQEQREYQRRGRGQGHQP | -41.12 | 29.0 | HHHHHHHHHHHHHHHHHHHHHHHHHCCCCCC |
| 707 | EQEERKYQEDRNNREEQQRYQREGRGQGHQP | -40.63 | 25.8 | CHHHHHHHHHHHHHHHHHHHHHHHCCCCCCC |
| 821 | EEEERRRQEQYNNRWEQERYQREGKGQGHSH | -40.62 | 29.0 | HHHHHHHHHHHHHHHHHHHHHHHCCCCCCCC |
| 439 | EEEERKRQQDFSNRYEQQYYQKQGQGQGHSH | -39.19 | 29.0 | HHHHHHHHHHHHHHHHHHHHHHHHCCCCCCC |
| 843 | EEEERKYQQHFEERFRQQYYQQKGQGQGHNH | -38.10 | 25.8 | HHHHHHHHHHHHHHHHHHHHHHHHCCCCCCC |
| 680 | EQEERHYQEDYNNRYEQEEYQRKGRGQGHSN | -37.92 | 25.8 | CHHHHHHHHHHHHHHHHHHHHHHCCCCCCCC |

**Table S3**. Summary of 966 peptide sequences designed using EvoEF2 and the evolutionary profile (weight = 0.50).

| #label | #peptide | #binding(EEU) | #SeqID(%) | #Secondary structure |
|---|---|---|---|---|
| WT | EEQAKTFLDKFNHEAEDLFYQSSGLGKGDFR | -46.46 | 100. | HHHHHHHHHHHHHHHHHHHHHHHCCCCCCCC |
| 866 | EEERRKQLDKYNNEHEDKRYQREGMGKGHQP | -51.00 | 48.4 | HHHHHHHHHHHHHHHHHHHHHHHHCCCCCCC |
| 986 | EEEARKQLDKYKNEEEDKRYQREGQGKGHQS | -50.81 | 48.4 | HHHHHHHHHHHHHHHHHHHHHHHHCCCCCCC |
| 473 | EEEKKKILDKINQEAEDKRYQREGWGKGHNP | -50.78 | 54.8 | HHHHHHHHHHHHHHHHHHHHHHHHCCCCCCC |
| 266 | EEEKRKILDKINNEAEDKRYQREGMGKGHQP | -50.48 | 51.6 | HHHHHHHHHHHHHHHHHHHHHHHHHCCCCCC |
| 685 | EEEQRKQQDKYNNEYEDKYYQQKGFGKGHQS | -50.33 | 45.2 | HHHHHHHHHHHHHHHHHHHHHHHCCCCCCCC |
| 477 | EEERKKQLDQYNNEYEDREYQRNGRGKGHQY | -50.22 | 48.4 | HHHHHHHHHHHHHHHHHHHHHHCCCCCCCCC |
| 130 | EEERRKQLDQFNNRKEDERYQQQGMGKGHQP | -50.09 | 45.2 | HHHHHHHHHHHHHHHHHHHHHHHHCCCCCCC |
| 514 | EEERKKQLDQYNNEHEDQRYQREGQGKGHSS | -50.05 | 48.4 | HHHHHHHHHHHHHHHHHHHHHHHCCCCCCCC |
| 440 | EEERRKQLDQFNNEYEDSFYQQKGFGKGHQS | -50.05 | 51.6 | HHHHHHHHHHHHHHHHHHHHHHHCCCCCCCC |
| 824 | EEERRKQLDQYNNEEEDKRYQREGMGKGHQP | -50.05 | 45.2 | HHHHHHHHHHHCHHHHHHHHHHHHCCCCCCC |
| 565 | EEELKKILDQINQEEEDRRYQRYGFGKGHQS | -50.00 | 48.4 | HHHHHHHHHHHHHHHHHHHHHHHCCCCCCCC |
| 448 | EEEEKKKRDKYNNEYEDEEYQRKGRGKGHQP | -49.88 | 48.4 | CHHHHHHHHHHHHHHHHHHHHHHHCCCCCCC |
| 591 | EEEKRKILDQLNQEKEDQRYQREGMGKGHQP | -49.83 | 45.2 | HHHHHHHHHHHHHHHHHHHHHHHHHCCCCCC |
| 776 | EEELRKILDQINQEEEDQRYQQYGFGKGHQN | -49.83 | 45.2 | HHHHHHHHHHHHHHHHHHHHHHHCCCCCCCC |
| 884 | EEEKRKILDQINNEQEDQSFQRHGFGKGKSR | -49.81 | 45.2 | HHHHHHHHHHHHHHHHHHHHHHHCCCCCCCC |
| 426 | EEEKRKLLDKINNEEEDKRFQREGFGKGKQN | -49.77 | 45.2 | HHHHHHHHHHHHHHHHHHHHHHHHCCCCCCC |
| 190 | EEERKKQLEKYNQEHEDKRYQREGMGKGHQP | -49.58 | 48.4 | HHHHHHHHHHHHHHHHHHHHHHHHCCCCCCC |
| 031 | EEEKRKILDQINNEEEDKRYQRYGFGKGHQS | -49.53 | 45.2 | HHHHHHHHHHHHHHHHHHHHHHHCCCCCCCC |
| 080 | EEEKRKILDQYNNEYEDQEYQRNGRGKGHSY | -49.53 | 45.2 | HHHHHHHHHHHHHHHHHHHHHHCCCCCCCCC |
| 566 | EEERRKQLDQFNNRFEDEYYQQYGFGKGHQS | -49.48 | 45.2 | HHHHHHHHHHHHHHHHHHHHHHHCCCCCCCC |
| 205 | EEEKKKLLDKINNEEEDKRYQQYGFGKGHQS | -49.46 | 51.6 | HHHHHHHHHHHHHHHHHHHHHHHCCCCCCCC |
| 283 | EEEKRKLLDKINNEAEDKRYQKEGMGKGHQP | -49.46 | 51.6 | HHHHHHHHHHHHHHHHHHHHHHHHCCCCCCC |
| 207 | EEEKRKLLDKINNEEEDKRYQREGFGKGHQS | -49.46 | 48.4 | HHHHHHHHHHHHHHHHHHHHHHHHCCCCCCC |
| 969 | EEEKRKLLDKLNNEKEDKRYQREGQGKGHQS | -49.46 | 48.4 | HHHHHHHHHHHHHHHHHHHHHHHHCCCCCCC |
| 179 | EEERRKQLDQYNNEYEDEYYQQQGQGKGHSY | -49.44 | 45.2 | HHHHHHHHHHHHHHHHHHHHHHHHCCCCCCC |
| 724 | EEEQRKQQDQYNNEYEDQKYQEQGFGKGHQS | -49.38 | 41.9 | HHHHHHHHHHHHHHHHHHHHHHHCCCCCCCC |
| 297 | EEERRKQLEKFNNEFEDKYYQQQGFGKGHSS | -49.29 | 48.4 | HHHHHHHHHHHHHHHHHHHHHHHCCCCCCCC |
| 066 | EEERRKQLEKFNNEYEDKYYQQQGFGKGHQS | -49.29 | 48.4 | HHHHHHHHHHHHHHHHHHHHHHHCCCCCCCC |
| 933 | EEERKKQLDQYNNEEEDMRYQREGMGKGHQP | -49.21 | 48.4 | HHHHHHHHHHHCCHHHHHHHHHHHHCCCCCC |
| 732 | EQEKRKLLDQINQEKEDREYQRYGFGKGHQS | -49.16 | 41.9 | HHHHHHHHHHHHHHHHHHHHHHHCCCCCCCC |
| 956 | EEERKKQLEKMKNEKEDKEYQRNGSGKGHSY | -49.10 | 45.2 | HHHHHHHHHHHHHHHHHHHHHHCCCCCCCCC |
| 745 | EEEAKKQQEKYSNEYEDKEYQRNGRGKGHSY | -49.10 | 45.2 | HHHHHHHHHHHHHHHHHHHHHHCCCCCCCCC |
| 148 | EEEKRKQLEKYNNEEEDKRYQREGMGKGHQP | -49.10 | 45.2 | HHHHHHHHHHHCHHHHHHHHHHHHCCCCCCC |
| 303 | EEEKRKLLDQINQEAEDERYQQQGMGKGHQP | -49.05 | 48.4 | HHHHHHHHHHHHHHHHHHHHHHHHHCCCCCC |

| | | | | |
|---|---|---|---|---|
| 689 | EEEKKKLLDQIENEREDQEYQRYGFGKGHQS | -48.91 | 45.2 | HHHHHHHHHHHHHHHHHHHHHHHHCCCCCCCC |
| 238 | EEEKKKFLDKYNNEEEDMRYQREGMGKGHQP | -48.90 | 54.8 | HHHHHHHHHHHCCHHHHHHHHHHHHHCCCCCC |
| 688 | EEEKRKFLDKYNNEYEDKYYQQKGFGKGHSS | -48.90 | 51.6 | HHHHHHHHHHHHHHHHHHHHHHHHCCCCCCCC |
| 836 | EEEKRKLLDQINNEEEDRRYQREGFGKGHQS | -48.86 | 45.2 | HHHHHHHHHHHHHHHHHHHHHHHHHCCCCCCC |
| 653 | EEEEKKIQEKRNNEEEDKQYQRSGMGKGHQP | -48.77 | 48.4 | CHHHHHHHHHHHHHHHHHHHHHHHHCCCCCCC |
| 293 | EEEKKKILEKINNEAEDKRYQRQGFGKGHSS | -48.77 | 51.6 | HHHHHHHHHHHHHHHHHHHHHHHHCCCCCCCC |
| 001 | EEEKRKILEKINNEEEDKRYQREGMGKGHQP | -48.77 | 45.2 | HHHHHHHHHHHHHHHHHHHHHHHHHCCCCCCC |
| 861 | EEEKRKILEKINNEKEDKEYQRAGRGKGHQP | -48.77 | 45.2 | HHHHHHHHHHHHHHHHHHHHHHHHHCCCCCCC |
| 379 | EEEKRKILEKINNEKEDKEYQRSGMGKGHQP | -48.77 | 48.4 | HHHHHHHHHHHHHHHHHHHHHHHHHCCCCCCC |
| 664 | EEEKRKILEKLNNEKEDKRYQRYGFGKGHQS | -48.77 | 45.2 | HHHHHHHHHHHHHHHHHHHHHHHHCCCCCCCC |
| 580 | EEEKKKLLDQINNEAEDKRYQRYGQGKGHQS | -48.75 | 51.6 | HHHHHHHHHHHHHHHHHHHHHHHHHCCCCCCC |
| 326 | EEEKKKLLDQINNEKEDEEYQRYGFGKGHSS | -48.75 | 48.4 | HHHHHHHHHHHHHHHHHHHHHHHHCCCCCCCC |
| 469 | EEEKKKLLDQINNEKEDQEYQKQGMGKGHQP | -48.75 | 48.4 | HHHHHHHHHHHHHHHHHHHHHHHHHCCCCCCC |
| 413 | EEEKKKLLDQLNNEKEDKRYQREGRGKGHQP | -48.75 | 48.4 | HHHHHHHHHHHHHHHHHHHHHHHHHCCCCCCC |
| 530 | EEEKRKLLDQINNEAEDKRYQREGKGKGHSA | -48.75 | 48.4 | HHHHHHHHHHHHHHHHHHHHHHHHCCCCCCCC |
| 160 | EEEKRKLLDQINNEAEDQRYQREGMGKGHQP | -48.75 | 48.4 | HHHHHHHHHHHHHHHHHHHHHHHHHHCCCCCC |
| 803 | EEEKRKLLDQINNEEEDERYQQQGMGKGHQP | -48.75 | 45.2 | HHHHHHHHHHHHHHHHHHHHHHHHHCCCCCCC |
| 142 | EEEKRKLLDQINNEEEDKRYQQQGFGKGHQS | -48.75 | 45.2 | HHHHHHHHHHHHHHHHHHHHHHHHHCCCCCCC |
| 046 | EEEKRKLLDQINNEEEDQRYQQQGFGKGHQS | -48.75 | 45.2 | HHHHHHHHHHHHHHHHHHHHHHHHCCCCCCCC |
| 394 | EEEKRKLLDQINNEEEDQRYQQYGFGKGHQS | -48.75 | 45.2 | HHHHHHHHHHHHHHHHHHHHHHHHCCCCCCCC |
| 733 | EEEKRKLLDQINNEEEDQRYQRYGMGKGHQP | -48.75 | 45.2 | HHHHHHHHHHHHHHHHHHHHHHHHHCCCCCCC |
| 858 | EEEKRKLLDQLNNEKEDQRYQRYGFGKGHSS | -48.75 | 45.2 | HHHHHHHHHHHHHHHHHHHHHHHHCCCCCCCC |
| 282 | EEERRKLLDQINNEEEDSRYQQQGMGKGHQP | -48.75 | 45.2 | HHHHHHHHHHHHHHHHHHHHHHHHHCCCCCCC |
| 792 | EEERRKQLEKFNNEFEDKYYQQKGFGKGHQS | -48.69 | 48.4 | HHHHHHHHHHHHHHHHHHHHHHHHCCCCCCCC |
| 746 | EEERKKQLEQFNQREEDERYQREGFGKGHSS | -48.65 | 45.2 | HHHHHHHHHHHHHHHHHHHHHHHHHCCCCCCC |
| 094 | EQEKRKILEQYNNDYEDEEYQRKGRGKGHQP | -48.60 | 35.5 | HHHHHHHHHHHHHHHHHHHHHHHHHCCCCCCC |
| 362 | EQEQRKQQEQFSNEYEDRFYQQKGFGKGHSS | -48.58 | 38.7 | HHHHHHHHHHHHHHHHHHHHHHHHCCCCCCCC |
| 601 | EEEKRKFLDKKNNEEEQKRYQREGKGRGHSH | -48.54 | 45.2 | HHHHHHHHHHCCCHHHHHHHHHHHCCCCCCCC |
| 049 | EEEKRKLLDQINNEAEDKRYQREGKGKGHSH | -48.52 | 48.4 | HHHHHHHHHHHHHHHHHHHHHHHHCCCCCCCC |
| 352 | DEERKKMLDKINNEAEDKRYQREGKGKGHSH | -48.51 | 51.6 | HHHHHHHHHHHHHHHHHHHHHHHHCCCCCCCC |
| 414 | EEEKKKMLDKKNNEEEDKRYQREGMGKGHQP | -48.51 | 51.6 | HHHHHHHHHHCCHHHHHHHHHHHHHHCCCCCC |
| 044 | EEEKRKMLDKINNEEEDKRYQREGMGKGHQP | -48.51 | 48.4 | HHHHHHHHHHHHHHHHHHHHHHHHHCCCCCCC |
| 344 | EEEKRKMLDKINNEQEDKRYQQQGFGKGHQS | -48.51 | 48.4 | HHHHHHHHHHHHHHHHHHHHHHHHHCCCCCCC |
| 248 | EEEKRKFLDQFNQEYEDEEYQRKGRGKGHQS | -48.49 | 51.6 | HHHHHHHHHHHHHHHHHHHHHHHHHCCCCCCC |
| 177 | EEEKRKFLDQYNQESEDERYQQQGFGKGHQS | -48.49 | 48.4 | HHHHHHHHHHHHHHHHHHHHHHHHCCCCCCCC |
| 461 | EEERKKQLDKENNEKEDKRYQRYGFGKGHQS | -48.48 | 51.6 | HHHHHHHHHHHHHHHHHHHHHHHHCCCCCCCC |
| 102 | EEERRKQLDKINNEQEDEEYQRKGRGKGHQS | -48.48 | 48.4 | HHHHHHHHHHHHHHHHHHHHHHHHHCCCCCCC |
| 705 | EEERRKQLDKYNNEYEDKYYQQKGFGKGHQS | -48.48 | 48.4 | HHHHHHHHHHHHHHHHHHHHHHHHCCCCCCCC |

| | | | | |
|---|---|---|---|---|
| 254 | EQERRKQLDKFNNEYEDKFYQQKGFGKGHQS | -48.48 | 51.6 | HHHHHHHHHHHHHHHHHHHHHHHCCCCCCCC |
| 292 | EEEWRKILEKLNNEKEDKRYQREGMGKGHQP | -48.44 | 45.2 | HHHHHHHHHHHHHHHHHHHHHHHHCCCCCCC |
| 670 | EEEQKKQREQFNNEYEDQRYQQEGFGKGHSS | -48.42 | 45.2 | HHHHHHHHHHHHHHHHHHHHHHHHCCCCCCC |
| 908 | EEEERKFRDKYNNEYEDKEYQRTGRGKGHQF | -48.38 | 48.4 | CHHHHHHHHHHHHHHCCHHHHHCCCCCCCCC |
| 810 | EEEKKKWLDQLNQEKEDQRYQREGMGKGHQP | -48.36 | 48.4 | HHHHHHHHHHHHHHHHHHHHHHHHHCCCCCC |
| 481 | EQERRKQLEKYNNEYEDKEYQRTGRGKGNSN | -48.35 | 41.9 | HHHHHHHHHHHHHHHHHHHHHHCCCCCCCCC |
| 005 | EEERRKQLDKYNNEAEQKRYQREGMGQGHQP | -48.34 | 45.2 | HHHHHHHHHHHHHHHHHHHHHHHHHCCCCCC |
| 527 | EEEKRKFLDQFNNEYEDRYYQQKGFGKGHQS | -48.29 | 51.6 | HHHHHHHHHHHHHHHHHHHHHHHCCCCCCCC |
| 161 | EEEKRKFLDQYNNEYEDRYYQQKGQGKGHSH | -48.29 | 48.4 | HHHHHHHHHHHHHHHHHHHHHHHHCCCCCCC |
| 304 | EEEKRKQLEQFNNEYEDQFYQQKGFGKGHSS | -48.29 | 48.4 | HHHHHHHHHHHHHHHHHHHHHHHCCCCCCCC |
| 189 | EEERKKQLEQKNNEEEDQRYQREGMGKGHQP | -48.29 | 45.2 | HHHHHHHHHHHHHHHHHHHHHHHHCCCCCCC |
| 091 | EEERKKQLEQYNNEAEDERYQQQGFGKGHQS | -48.29 | 48.4 | HHHHHHHHHHHHHHHHHHHHHHHCCCCCCCC |
| 458 | EEERKKQLEQYNNEAEDQRYQRQGFGKGHQN | -48.29 | 48.4 | HHHHHHHHHHHHHHHHHHHHHHHHCCCCCCC |
| 062 | EEERKKQLEQYNNEAEDQRYQRYGFGKGHSS | -48.29 | 48.4 | HHHHHHHHHHHHHHHHHHHHHHHCCCCCCCC |
| 937 | EEERKKQLEQYNNEEEDMRYQREGMGKGHQP | -48.29 | 45.2 | HHHHHHHHHHHCHHHHHHHHHHHHHCCCCCC |
| 048 | EEERKKQLEQYNNEYEDEEYQRTGRGKGHQF | -48.29 | 45.2 | HHHHHHHHHHHHHHHHHHHHHHHCCCCCCCC |
| 312 | EEERKKQLEQYNNEYEDQFYQQKGFGKGHQS | -48.29 | 48.4 | HHHHHHHHHHHHHHHHHHHHHHHCCCCCCCC |
| 020 | EEERKKQLEQYSNEEEDMRYQREGKGKGHSS | -48.29 | 41.9 | HHHHHHHHHHHHHHHHHHHHHHHCCCCCCCC |
| 552 | EEERRKQLEQFNNEFEDQYYQQKGQGKGHSH | -48.29 | 45.2 | HHHHHHHHHHHHHHHHHHHHHHHHCCCCCCC |
| 520 | EEERRKQLEQINNEEEDQRYQRYGFGKGHQS | -48.29 | 41.9 | HHHHHHHHHHHHHHHHHHHHHHHCCCCCCCC |
| 634 | EEERRKQLEQYNNESEDQRYQQYGFGKGHSS | -48.29 | 41.9 | HHHHHHHHHHHHHHHHHHHHHHHCCCCCCCC |
| 790 | EEERRKQLEQYNNEYEDEEYQRKGRGKGHQP | -48.29 | 41.9 | HHHHHHHHHHHHHHHHHHHHHHHHCCCCCCC |
| 561 | EQERRKQLEQYNNEAEDQRYQRYGFGKGHQS | -48.29 | 41.9 | HHHHHHHHHHHHHHHHHHHHHHHCCCCCCCC |
| 510 | EEERRKQLEQYNNQSEDERYQRYGFGKGHQA | -48.27 | 38.7 | HHHHHHHHHHHHHCCHHHHHHHHCCCCCCCC |
| 298 | EEEQKKQREKYNNEAEQKRYQREGKGRGHSH | -48.26 | 41.9 | HHHHHHHHHHHHHHHHHHHHHHHCCCCCCCC |
| 502 | EEEKRKMLDQINQEEEDRRYQQQGMGKGHQP | -48.20 | 45.2 | HHHHHHHHHHHHHHHHHHHHHHHHCCCCCCC |
| 735 | EEEERKFRDQFNNEAEDEEYQRRGMGKGHQP | -48.19 | 51.6 | CHHHHHHHHHHHHHHHHHHHHHHHHCCCCCC |
| 300 | EEEKKKFLDQFNNEYEDQFYQQKGFGKGHQS | -48.19 | 58.1 | HHHHHHHHHHHHHHHHHHHHHHHCCCCCCCC |
| 863 | EEEKKKFLDQFNNEYEDQYYQQKGQGKGHSH | -48.19 | 54.8 | HHHHHHHHHHHHHHHHHHHHHHHHCCCCCCC |
| 614 | EEEKKKFLDQYNNEEEDQRYQREGMGKGHQP | -48.19 | 51.6 | HHHHHHHHHHHCCHHHHHHHHHHHCCCCCCC |
| 076 | EEEKKKFLDQYNNESEDERYQQQGWGKGHNP | -48.19 | 51.6 | HHHHHHHHHHHCCCHHHHHHHHHCCCCCCCC |
| 704 | EEEKRKFLDQFNNEEEDKRYQREGFGKGHQS | -48.19 | 51.6 | HHHHHHHHHHHHHHHHHHHHHHHHCCCCCCC |
| 759 | EEEKRKFLDQFNNEFEDQYYQQKGQGKGHSH | -48.19 | 51.6 | HHHHHHHHHHHHHHHHHHHHHHHHCCCCCCC |
| 267 | EEEKRKFLDQFNNEYEDKYYQQQGFGKGHQS | -48.19 | 51.6 | HHHHHHHHHHHHHHHHHHHHHHHCCCCCCCC |
| 574 | EEEKRKFLDQFNNEYEDQYYQQKGFGKGHQS | -48.19 | 51.6 | HHHHHHHHHHHHHHHHHHHHHHHCCCCCCCC |
| 771 | EEEKRKFLDQFNNEYEDQYYQQKGQGKGHSH | -48.19 | 51.6 | HHHHHHHHHHHHHHHHHHHHHHHHCCCCCCC |
| 642 | EEEKRKFLDQYNNEAEDQRYQREGMGKGHQP | -48.19 | 51.6 | HHHHHHHHHHHHHHHHHHHHHHHHHCCCCCC |
| 146 | EEEKRKFLDQYNNESEDQRYQQYGFGKGHQS | -48.19 | 48.4 | HHHHHHHHHHHCCCHHHHHHHHHCCCCCCCC |

| | | | | |
|---|---|---|---|---|
| 657 | EEEKRKFLDQYNNESEDQRYQRYGFGKGHSS | -48.19 | 48.4 | HHHHHHHHHHHCCCCHHHHHHHHCCCCCCCC |
| 368 | EEEKRKFLDQYNNEYEDEEYQQTGRGKGHSS | -48.19 | 48.4 | HHHHHHHHHHHHHHHHHHHHHHCCCCCCCCC |
| 825 | EEEKRKFLDQYNNEYEDEEYQRNGRGKGHSY | -48.19 | 48.4 | HHHHHHHHHHHHHHHHHHHHHHCCCCCCCCC |
| 874 | EEEKRKFLDQYNNEYEDKEYQKEGRGKGHQP | -48.19 | 48.4 | HHHHHHHHHHHHHHHHHHHHHHHHCCCCCCC |
| 498 | EEEQRKFEDQRNNEEEDKRYQQYGFGKGHQS | -48.19 | 45.2 | CHHHHHHHHHHHHHHHHHHHHHHCCCCCCCC |
| 783 | EEEKKKFLDQMKNRKEDEEYQKQGWGKGHNP | -48.16 | 45.2 | HHHHHHHHHHHHHHHHHHHHHHHCCCCCCCC |
| 123 | EEEKRKMLDQINNEKEDEEYQRYGFGKGTQN | -48.12 | 45.2 | HHHHHHHHHHHHHHHHHHHHHHHCCCCCCCC |
| 321 | EEEERKQREQYNNEAEDQRYQREGKGKGHSS | -48.09 | 41.9 | HHHHHHHHHHHHHHHHHHHHHHHCCCCCCCC |
| 622 | EEEKRKLLDKINNEAEDRRYQKEGMGKGHQP | -48.07 | 51.6 | HHHHHHHHHHHHHHHHHHHHHHHHCCCCCCC |
| 995 | EEEKRKLLDKINNEEEDRRYQREGVGKGHNP | -48.07 | 48.4 | HHHHHHHHHHHHHHHHHHHHHHHHCCCCCCC |
| 750 | EEEYRKWLDQYKNEAEDQRYQREGQGKGHQS | -48.06 | 45.2 | HHHHHHHHHHHHHHHHHHHHHHHHCCCCCCC |
| 488 | EQEKRKILEQLNQEKEDKRYQREGKGKGHSS | -48.06 | 38.7 | HHHHHHHHHHHHHHHHHHHHHHHCCCCCCCC |
| 671 | EEEKKKLLEKINNEEEDKRFQQQGFGKGKQN | -48.06 | 45.2 | HHHHHHHHHHHHHHHHHHHHHHHCCCCCCCC |
| 350 | EEERRKQLEKYNNEEEDKRYQREGFGKGDSR | -48.04 | 51.6 | HHHHHHHHHHHCHHHHHHHHHHHHCCCCCCC |
| 957 | EEEKRKLLDQINNEEEDQRYQQQGFGKGDSR | -48.01 | 51.6 | HHHHHHHHHHHHHHHHHHHHHHHCCCCCCCC |
| 096 | EEEKRKLLDQINNEEEDQRYQQYGFGKGDSR | -48.01 | 51.6 | HHHHHHHHHHHHHHHHHHHHHHHCCCCCCCC |
| 357 | EEEKKKFLDQYNNEWEDMRYQREGKGKGHSH | -47.95 | 51.6 | HHHHHHHHHHHHHHHHHHHHHHHCCCCCCCC |
| 285 | EEEKRKLLEKLNNEKEDKRYQQQGRGKGHQP | -47.94 | 45.2 | HHHHHHHHHHHHHHHHHHHHHHHHCCCCCCC |
| 719 | EEEARKNQEKYNNEYEDEEYQRKGRGKGDSR | -47.87 | 51.6 | HHHHHHHHHHHHHHHHHHHHHHHCCCCCCCC |
| 173 | EEELKKILEQISQEEEDKRYQREGFGKGHSN | -47.83 | 41.9 | HHHHHHHHHHHHHHHHHHHHHHHHCCCCCCC |
| 960 | EEEKRKLLDQINNEQEDEEYQRKGRGKGNQN | -47.81 | 45.2 | HHHHHHHHHHHHHHHHHHHHHHHHCCCCCCC |
| 665 | EEEKRKLLDQINQEEEDQRYQQQGFGKGDSR | -47.81 | 51.6 | HHHHHHHHHHHHHHHHHHHHHHHCCCCCCCC |
| 968 | EEEKKKMLDQFNNEYEDQFYQQKGFGKGHQN | -47.79 | 54.8 | HHHHHHHHHHHHHHHHHHHHHHHCCCCCCCC |
| 243 | EEEKKKMLDQINNEAEDKRYQKEGFGKGHSS | -47.79 | 51.6 | HHHHHHHHHHHHHHHHHHHHHHHHCCCCCCC |
| 870 | EEEKKKMLDQINNEEEDERYQQQGFGKGHQS | -47.79 | 48.4 | HHHHHHHHHHHHHHHHHHHHHHHCCCCCCCC |
| 087 | EEEKKKMLDQINNEEEDKRYQQYGMGKGHQP | -47.79 | 48.4 | HHHHHHHHHHHHHHHHHHHHHHHHCCCCCCC |
| 325 | EEEKRKMLDQINNEAEDKRYQREGKGKGHSH | -47.79 | 48.4 | HHHHHHHHHHHHHHHHHHHHHHHCCCCCCCC |
| 311 | EEEKRKMLDQINNEAEDKRYQREGMGKGHQP | -47.79 | 48.4 | HHHHHHHHHHHHHHHHHHHHHHHHHCCCCCC |
| 416 | EEEKRKMLDQINNEAEDKRYQRYGFGKGHQS | -47.79 | 48.4 | HHHHHHHHHHHHHHHHHHHHHHHCCCCCCCC |
| 571 | EEEKRKMLDQINNEKEDEEYQRRGKGKGHSS | -47.79 | 45.2 | HHHHHHHHHHHHHHHHHHHHHHHCCCCCCCC |
| 615 | EEEKRKMLDQINNEKEDKEYQRSGMGKGHQP | -47.79 | 48.4 | HHHHHHHHHHHHHHHHHHHHHHHHCCCCCCC |
| 931 | EEEKRKMLDQINNREEDERYQQQGFGKGHQS | -47.77 | 41.9 | HHHHHHHHHHHHHHHHHHHHHHHCCCCCCCC |
| 902 | EEERKKQLDQINNEKEDEEYQRYGFGKGHQN | -47.77 | 48.4 | HHHHHHHHHHHHHHHHHHHHHHHCCCCCCCC |
| 221 | EEERRKQLDQINNEEEDERYQQQGMGKGHQP | -47.77 | 45.2 | HHHHHHHHHHHHHHHHHHHHHHHHCCCCCCC |
| 006 | EEERRKQLDQINNEEEDKRYQQYGFGKGHSS | -47.77 | 45.2 | HHHHHHHHHHHHHHHHHHHHHHHCCCCCCCC |
| 455 | EEERRKQLDQINNEEEDKRYQREGFGKGHQS | -47.77 | 45.2 | HHHHHHHHHHHHHHHHHHHHHHHHCCCCCCC |
| 098 | EEERRKQLDQINNEEEDKRYQREGMGKGHQP | -47.77 | 45.2 | HHHHHHHHHHHHHHHHHHHHHHHHCCCCCCC |
| 181 | EEERRKQLDQINNEKEDEEYQRRGWGKGHNP | -47.77 | 45.2 | HHHHHHHHHHHHHHHHHHHHHHHCCCCCCCC |

| | | | | |
|---|---|---|---|---|
| 909 | EEERRKQLDQINNEKEDEEYQRRGWGKGHTP | -47.77 | 45.2 | HHHHHHHHHHHHHHHHHHHHHHHCCCCCCCC |
| 725 | EEERRKQLDQYNNEYEDEEYQRTGRGKGHQF | -47.77 | 45.2 | HHHHHHHHHHHHHHHHHHHHHHHCCCCCCCC |
| 758 | EEEKKKILEQINNEEEDQRYQQYGWGKGHNP | -47.77 | 45.2 | HHHHHHHHHHHHHHHHHHHHHHHCCCCCCCC |
| 537 | EEEKKKILEQINNEKEDEEYQRYGQGKGHQS | -47.77 | 45.2 | HHHHHHHHHHHHHHHHHHHHHHHHHCCCCCC |
| 953 | EEEKRKILEQINNEAEDQRYQREGMGKGHQP | -47.77 | 45.2 | HHHHHHHHHHHHHHHHHHHHHHHHHCCCCCC |
| 794 | EEEKRKILEQINNEEEDERYQRYGFGKGHQS | -47.77 | 41.9 | HHHHHHHHHHHHHHHHHHHHHHHCCCCCCCC |
| 236 | EEEKRKILEQINNEEEDQRYQQYGFGKGHSS | -47.77 | 41.9 | HHHHHHHHHHHHHHHHHHHHHHHCCCCCCCC |
| 443 | EEEKRKILEQINNEQEDQSYQRYGFGKGHQS | -47.77 | 41.9 | HHHHHHHHHHHHHHHHHHHHHHHCCCCCCCC |
| 749 | EEEKRKILEQLNNEKEDKRYQREGMGKGHQP | -47.77 | 41.9 | HHHHHHHHHHHHHHHHHHHHHHHHCCCCCCC |
| 489 | EEELKKILEQINNEEEDERYQQQGWGKGHTP | -47.77 | 45.2 | HHHHHHHHHHHHHHHHHHHHHHHCCCCCCCC |
| 738 | EEELKKILEQLNNEQEDQKYQRYGFGKGHQS | -47.77 | 45.2 | HHHHHHHHHHHHHHHHHHHHHHHCCCCCCCC |
| 122 | EEELRKILEQINNEEEDKRYQREGMGKGHQP | -47.77 | 41.9 | HHHHHHHHHHHHHHHHHHHHHHHHCCCCCCC |
| 731 | EEEWRKILEQLNNEKEDQRYQREGMGKGHQP | -47.77 | 41.9 | HHHHHHHHHHHHHHHHHHHHHHHHHCCCCCC |
| 891 | EQEKRKILEQINNEEEDQRYQRYGMGKGHQP | -47.77 | 38.7 | HHHHHHHHHHHHHHHHHHHHHHHHCCCCCCC |
| 403 | EEEERKLQEKINNEAEDKRYQREGWGKGHNP | -47.75 | 45.2 | HHHHHHHHHHHHHHHHHHHHHHHHCCCCCCC |
| 407 | EEEKKKLLEKINNEAEDKRYQREGMGKGHQP | -47.75 | 51.6 | HHHHHHHHHHHHHHHHHHHHHHHHHCCCCCC |
| 721 | EEEKKKLLEKINNEEEDKRYQQQGFGKGHQS | -47.75 | 48.4 | HHHHHHHHHHHHHHHHHHHHHHHHCCCCCCC |
| 700 | EEEKKKLLEKINNEEEDKRYQQQGFGKGHSS | -47.75 | 48.4 | HHHHHHHHHHHHHHHHHHHHHHHCCCCCCCC |
| 774 | EEEKKKLLEKLNNEKEDEEYQRRGMGKGHQP | -47.75 | 48.4 | HHHHHHHHHHHHHHHHHHHHHHHHHCCCCCC |
| 010 | EEEKKKLLEKLNNEKEDKRYQRQGFGKGHQS | -47.75 | 48.4 | HHHHHHHHHHHHHHHHHHHHHHHHCCCCCCC |
| 912 | EEEKRKLLEKINNEAEDKRYQKEGFGKGHQS | -47.75 | 48.4 | HHHHHHHHHHHHHHHHHHHHHHHHCCCCCCC |
| 934 | EEEKRKLLEKINNEAEDKRYQQQGMGKGHQP | -47.75 | 48.4 | HHHHHHHHHHHHHHHHHHHHHHHHCCCCCCC |
| 391 | EEEKRKLLEKINNEAEDKRYQREGMGKGHQP | -47.75 | 48.4 | HHHHHHHHHHHHHHHHHHHHHHHHHCCCCCC |
| 569 | EEEKRKLLEKINNEAEDKRYQRYGFGKGHQS | -47.75 | 48.4 | HHHHHHHHHHHHHHHHHHHHHHHCCCCCCCC |
| 660 | EEEKRKLLEKINNEAEDKRYQRYGFGKGHSS | -47.75 | 48.4 | HHHHHHHHHHHHHHHHHHHHHHHCCCCCCCC |
| 880 | EEEKRKLLEKINNEEEDKRYQQQGMGKGHQP | -47.75 | 45.2 | HHHHHHHHHHHHHHHHHHHHHHHHCCCCCCC |
| 674 | EEEKRKLLEKINNEEEDKRYQREGFGKGHQS | -47.75 | 45.2 | HHHHHHHHHHHHHHHHHHHHHHHHCCCCCCC |
| 028 | EEEKRKLLEKINNEEEDKRYQREGWGKGHNP | -47.75 | 45.2 | HHHHHHHHHHHHHHHHHHHHHHHHCCCCCCC |
| 944 | EEEKRKLLEKINNEEEDKRYQRYGFGKGHSS | -47.75 | 45.2 | HHHHHHHHHHHHHHHHHHHHHHHCCCCCCCC |
| 804 | EEEKRKLLEKINNEKEDKEYQKQGWGKGHNP | -47.75 | 45.2 | HHHHHHHHHHHHHHHHHHHHHHHCCCCCCCC |
| 214 | EEEKRKLLEKINNEQEDEEYQRKGRGKGHQP | -47.75 | 45.2 | HHHHHHHHHHHHHHHHHHHHHHHHCCCCCCC |
| 492 | EEEKRKLLEKINNEQEDKSYQRYGFGKGHQS | -47.75 | 45.2 | HHHHHHHHHHHHHHHHCHHHHHHCCCCCCCC |
| 315 | EEEKRKLLEKLNNEKEDKKYQQEGHGKGHSR | -47.75 | 48.4 | HHHHHHHHHHHHHHHHHHHHHHHCCCCCCCC |
| 786 | EEEKRKLLEKLNNEKEDKRYQQQGFGKGHSS | -47.75 | 45.2 | HHHHHHHHHHHHHHHHHHHHHHHCCCCCCCC |
| 279 | EEEKRKLLEKLNNEKEDKRYQRYGMGKGHQP | -47.75 | 45.2 | HHHHHHHHHHHHHHHHHHHHHHHHCCCCCCC |
| 462 | EEEKRKLLEKLNNEQEDKRYQRQGFGKGHQS | -47.75 | 45.2 | HHHHHHHHHHHHHHHHHHHHHHHHCCCCCCC |
| 380 | EEERRKLLEKINNEEEDKRYQQQGFGKGHQS | -47.75 | 45.2 | HHHHHHHHHHHHHHHHHHHHHHHHCCCCCCC |
| 948 | EQEKRKLLEKINNEQEDKSYQRYGFGKGHQS | -47.75 | 41.9 | HHHHHHHHHHHHHHHHCHHHHHHCCCCCCCC |

| | | | | |
|---|---|---|---|---|
| 513 | EEERRKQLDQYNNRYEDEYYQQKGQGKGHSH | -47.74 | 41.9 | HHHHHHHHHHHHHHHHHHHHHHHHCCCCCCC |
| 482 | EEEQRKILEKINNEEEDRRKQRYGFGSGKQN | -47.73 | 38.7 | HHHHHHHHHHHHHHHHHHHHHHCCCCCCCCC |
| 791 | EEEKRKLLDAINNREEDERYQQQGMGKGHQP | -47.71 | 41.9 | HHHHHHHHHHHHHHHHHHHHHHHHCCCCCCC |
| 782 | EEERKKQLEQYNNEYEDEEYQRKGRGKGHQP | -47.68 | 45.2 | HHHHHHHHHHHHHHHHHHHHHHHHCCCCCCC |
| 013 | EEERRKQLEQFNNEFEDKYYQQKGFGKGHQS | -47.68 | 45.2 | HHHHHHHHHHHHHHHHHHHHHHHCCCCCCCC |
| 164 | EEERRKQLEQYNNEYEDEEYQRKGRGKGHQS | -47.68 | 41.9 | HHHHHHHHHHHHHHHHHHHHHHHHCCCCCCC |
| 363 | EEERRKQLEQYNNEYEDQRYQQEGMGKGHQP | -47.68 | 41.9 | HHHHHHHHHHHHHHHHHHHHHHHHCCCCCCC |
| 474 | EEERRKQLEKYNNEAEDRRYQREGMGKGHQP | -47.68 | 48.4 | HHHHHHHHHHHHHHHHHHHHHHHHHCCCCCC |
| 951 | EEEERKFQEKYNQEWEDMRYQREGFGKGHQS | -47.67 | 45.2 | CHHHHHHHHHHHHHHHHHHHHHHHHCCCCCC |
| 845 | EEEKRKFLDQQKNEEEDQRYQQYGFGKGHQS | -47.67 | 45.2 | HHHHHHHHHHHHHHHHHHHHHHHCCCCCCCC |
| 780 | EEEQKKLYEKLNNEKEDKRYQREGKGKGHSH | -47.64 | 45.2 | HHHHHHHHHHHHHHHHHHHHHHHCCCCCCCC |
| 699 | EEERKKQLDQLNNEKEDQRYQRYGFGKGHSP | -47.62 | 48.4 | HHHHHHHHHHHHHHHHHHHHHHHCCCCCCCC |
| 751 | EEEQRKQQEQYNNEYEDQFYQQQGFGKGHQS | -47.61 | 41.9 | HHHHHHHHHHHHHHHHHHHHHHHCCCCCCCC |
| 014 | EEEKRKFLDQKNNEEEDQRYQQYGFGKGHQP | -47.59 | 48.4 | HHHHHHHHHHHHHHHHHHHHHHHCCCCCCCC |
| 865 | EEEKRKMLDKFNNEYEDKYYQQQGFGKGNQN | -47.57 | 51.6 | HHHHHHHHHHHHHHHHHHHHHHHCCCCCCCC |
| 406 | EEERRKQLDQINQEAEDKRYQRYGMGKGHQP | -47.55 | 48.4 | HHHHHHHHHHHHHHHHHHHHHHHHCCCCCCC |
| 584 | EEERRKQLDDYNQRWEDMRYQREGMGKGHQP | -47.54 | 41.9 | HHHHHHHHHHHHHHHHHHHHHHHHHCCCCCC |
| 656 | EQELRKILEQYNNEEEDMRYQREGFGKGHQN | -47.53 | 38.7 | HHHHHHHHHHHCHHHHHHHHHHHHCCCCCCC |
| 683 | EEEKRKLLEKINNEEEDERYQQQGKGKGHSH | -47.52 | 45.2 | HHHHHHHHHHHHHHHHHHHHHHHCCCCCCCC |
| 427 | EEEEKKFQEKYNNESEDKRFQRYGFGKGKQN | -47.50 | 45.2 | CHHHHHHHHHHHHHHHHHHHHHHCCCCCCCC |
| 649 | EEEKRKFLEKFNNEQEDKSFQRHGFGKGKQN | -47.50 | 48.4 | HHHHHHHHHHHHHHHHHHHHHHHCCCCCCCC |
| 914 | EEEKRKFLEKFNQEEEDKEYQRVGRGKGHQF | -47.48 | 51.6 | HHHHHHHHHHHHHHHHHHHHHHHHCCCCCCC |
| 616 | EEEKKKLLEKINNEKEDKEFQKQGQGKGKQS | -47.47 | 45.2 | HHHHHHHHHHHHHHHHHHHHHHHHCCCCCCC |
| 138 | EEEQKKQREQYNQEYEDEEYQRKGRGKGDSR | -47.47 | 48.4 | HHHHHHHHHHHHHHHHHHHHHHHCCCCCCCC |
| 581 | EEEKRKFLEKYNNESEDQRFQRYGFGKGKSR | -47.46 | 48.4 | HHHHHHHHHHHHHCHHHHHHHHHCCCCCCCC |
| 686 | EEEKRKFLDQYNNEYEDEEYQRKGRGKGHQS | -47.46 | 48.4 | HHHHHHHHHHHHHHHHHHHHHHHHCCCCCCC |
| 099 | EEEERKQRQQMNNEKEDEEYQRRGMGKGHQP | -47.46 | 38.7 | HHHHHHHHHHHHHHHHHHHHHHHHHCCCCCC |
| 939 | EEEQKKFEDARNQREEDQRYQQQGFGKGHQS | -47.44 | 45.2 | CHHHHHHHHHHHHHHHHHHHHHHCCCCCCCC |
| 932 | EEEKRKLLQKINQEKEDKEYQKQGFGKGHQS | -47.41 | 45.2 | HHHHHHHHHHHHHHHHHHHHHHHHCCCCCCC |
| 433 | EEERRKQLDQFNNEKEQQRYQQQGFGQGHSS | -47.39 | 41.9 | HHHHHHHHHHHHHHHHHHHHHHHCCCCCCCC |
| 212 | EEEKRKILEKINNEEEDRRYQQYGMGKGHQP | -47.37 | 45.2 | HHHHHHHHHHHHHHHHHHHHHHHHCCCCCCC |
| 135 | EEEKRKILEKINNEKEDREYQKQGMGKGHQP | -47.37 | 45.2 | HHHHHHHHHHHHHHHHHHHHHHHHCCCCCCC |
| 768 | EEEKKKLLEQINQEEEDQRYQQQGFGKGHQS | -47.28 | 45.2 | HHHHHHHHHHHHHHHHHHHHHHHCCCCCCCC |
| 729 | EEEKKKLLEQLNQEKEDQSYQKEGFGKGHQS | -47.28 | 45.2 | HHHHHHHHHHHHHHHHHHHHHHHHCCCCCCC |
| 648 | EEEKRKLLEQINQEQEDEEYQRKGRGKGHQS | -47.28 | 41.9 | HHHHHHHHHHHHHHHHHHHHHHHHCCCCCCC |
| 964 | EEERRKLLEQINQEEEDKRYQREGMGKGHQP | -47.28 | 41.9 | HHHHHHHHHHHHHHHHHHHHHHHHCCCCCCC |
| 259 | EEEKRKMLDQINQEAEDQRYQRQGFGKGHQS | -47.25 | 48.4 | HHHHHHHHHHHHHHHHHHHHHHHHCCCCCCC |
| 621 | EEEKRKFLDQYNNEEEDMRYQREGFGKGNQN | -47.25 | 48.4 | HHHHHHHHHHHCHHHHHHHHHHHHCCCCCCC |

| | | | | |
|---|---|---|---|---|
| 139 | EEEKRKLLEKINNEKEDEEYQRYGFGKGHQN | -47.20 | 45.2 | HHHHHHHHHHHHHHHHHHHHHHHCCCCCCCC |
| 253 | EEEKRKFLEQLNNDEEDERYQQQGFGKGHSS | -47.20 | 41.9 | HHHHHHHHHHHHHHHHHHHHHHHCCCCCCCC |
| 400 | EEEQRKFEEQRNNDEEDQRYQQQGMGKGHQP | -47.20 | 38.7 | HHHHHHHHHHHHHHHHHHHHHHHHCCCCCCC |
| 501 | EEEKKKFLEKQNNEEEDKRYQQQGMGKGHQP | -47.19 | 51.6 | HHHHHHHHHHHHHHHHHHHHHHHHCCCCCCC |
| 152 | EEEKKKFLEKYNNESEDQRYQRYGFGKGHQN | -47.19 | 51.6 | HHHHHHHHHHHHHCHHHHHHHHHCCCCCCCC |
| 272 | EEEKKKFLEKYNNEYEDEEYQRKGRGKGHQS | -47.19 | 51.6 | HHHHHHHHHHHHHHHHHHHHHHHHCCCCCCC |
| 000 | EEEKKKFLEKYSNESEDKRYQQQGFGKGHQS | -47.19 | 48.4 | HHHHHHHHHHHHHHHHHHHHHHHHCCCCCCC |
| 017 | EEEKRKFEEKRNNEEEDKRYQQYGMGKGHQP | -47.19 | 45.2 | HHHHHHHHHHHHHHHHHHHHHHHHCCCCCCC |
| 976 | EEEKRKFLEKFNNEFEDKYYQQKGFGKGHQP | -47.19 | 51.6 | HHHHHHHHHHHHHHHHHHHHHHHCCCCCCCC |
| 526 | EEEKRKFLEKFNNEYEDKYYQQKGFGKGHQS | -47.19 | 51.6 | HHHHHHHHHHHHHHHHHHHHHHHCCCCCCCC |
| 392 | EEEKRKFLEKFNNEYEDKYYQQKGQGKGHSH | -47.19 | 51.6 | HHHHHHHHHHHHHHHHHHHHHHHHCCCCCCC |
| 079 | EEEKRKFLEKKNNEEEDKRYQQYGMGKGHQP | -47.19 | 48.4 | HHHHHHHHHHHHHHHHHHHHHHHHCCCCCCC |
| 464 | EEEKRKFLEKMKNEKEDEEYQRYGFGKGHSS | -47.19 | 45.2 | HHHHHHHHHHHHHHHHHHHHHHHCCCCCCCC |
| 360 | EEEKRKFLEKYNNEAEDERYQRYGFGKGHQS | -47.19 | 51.6 | HHHHHHHHHHHHHHHHHHHHHHHCCCCCCCC |
| 945 | EEEKRKFLEKYNNEAEDKRYQREGKGKGHSH | -47.19 | 51.6 | HHHHHHHHHHHHHHHHHHHHHHHCCCCCCCC |
| 280 | EEEKRKFLEKYNNEAEDKRYQREGMGKGHQP | -47.19 | 51.6 | HHHHHHHHHHHHHHHHHHHHHHHHHCCCCCC |
| 387 | EEEKRKFLEKYNNEAEDKRYQRQGFGKGHQS | -47.19 | 51.6 | HHHHHHHHHHHHHHHHHHHHHHHHCCCCCCC |
| 651 | EEEKRKFLEKYNNEAEDKRYQRYGFGKGHQS | -47.19 | 51.6 | HHHHHHHHHHHHHHHHHHHHHHHCCCCCCCC |
| 274 | EEEKRKFLEKYNNESEDKRYQQQGMGKGHQP | -47.19 | 48.4 | HHHHHHHHHHHHHHHHHHHHHHHHCCCCCCC |
| 991 | EQEKRKFLEKYNNEYEDKYYQQNGQGKGHSY | -47.19 | 45.2 | HHHHHHHHHHHHHHHHHHHHHHHCCCCCCCC |
| 895 | EQEWRKFIEKMKNEKEDKEYQKQGFGKGHSS | -47.19 | 38.7 | CHHHHHHHHHHHHHHHHHHHHHHCCCCCCCC |
| 922 | EEEKRKFLDKYNNESEHQRYQQQGWGQGHNP | -47.18 | 45.2 | HHHHHHHHHHHHHHHHHHHHHHHHCCCCCCC |
| 920 | EEELKKFLEKMNNEQEDKRYQKEGFGKGHSS | -47.17 | 51.6 | HHHHHHHHHHHHHHHHHHHHHHHHCCCCCCC |
| 353 | EEERRKQLEQFNNEYEDEEYQRKGRGKGHSS | -47.16 | 45.2 | HHHHHHHHHHHHHHHHHHHHHHHCCCCCCCC |
| 107 | EEERRKQLEQYNNEYEDQEYQRNGRGKGHSY | -47.16 | 41.9 | HHHHHHHHHHHHHHHHHHHHHHCCCCCCCCC |
| 955 | EQEERKKLEQYKNEYEDKQYQKEGFGKGHQS | -47.16 | 35.5 | CHHHHHHHHHHHHHHHHHHHHHHHCCCCCCC |
| 093 | EEEKKKLLEQIENEREDQEYQRKGMGKGHQP | -47.15 | 41.9 | HHHHHHHHHHHHHHHHHHHHHHHHCCCCCCC |
| 002 | EEEKRKMLEQINQDEEDKRYQREGFGKGHQS | -47.10 | 38.7 | HHHHHHHHHHHHHHHHHHHHHHHHCCCCCCC |
| 620 | EEEEKKLQEQINNEEEDRRYQREGQGKGHQS | -47.09 | 41.9 | CHHHHHHHHHHHHHHHHHHHHHHHCCCCCCC |
| 340 | EEEERKLQEQINNEEEDRRYQQYGKGKGHSH | -47.09 | 38.7 | HHHHHHHHHHHHHHHHHHHHHHHHCCCCCCC |
| 562 | EEEKKKLLEQINNEAEDRRYQRYGFGKGHQS | -47.09 | 48.4 | HHHHHHHHHHHHHHHHHHHHHHHCCCCCCCC |
| 593 | EEEKKKLLEQLNNEKEDRRYQKEGQGKGHQS | -47.09 | 45.2 | HHHHHHHHHHHHHHHHHHHHHHHCCCCCCCC |
| 195 | EEEKRKLLEQINNEEEDRNYQREGKGKGHSH | -47.09 | 41.9 | HHHHHHHHHHHHHHHHHHHHHHHCCCCCCCC |
| 839 | EEEKKKMLEKINQEAEDKRYQRYGFGKGHQN | -47.09 | 51.6 | HHHHHHHHHHHHHHHHHHHHHHHCCCCCCCC |
| 542 | EEEKKKMLEKINQEKEDEEYQRRGMGKGHQP | -47.09 | 48.4 | HHHHHHHHHHHHHHHHHHHHHHHHHCCCCCC |
| 814 | EQERRKQLEKFNNEYEDKFFQKQGFGKGKQN | -47.08 | 45.2 | HHHHHHHHHHHHHHHHHHHHHHHCCCCCCCC |
| 952 | EEERRKQLEQINQDAEDQRYQKEGFGKGHQS | -47.07 | 41.9 | HHHHHHHHHHHHHHHHHHHHHHHHCCCCCCC |
| 546 | EEEYKKWLEKFKNEYEDKYYQQKGFGKGHQS | -47.06 | 48.4 | HHHHHHHHHHHHHHHHHHHHHHHCCCCCCCC |

| | | | | |
|---|---|---|---|---|
| 972 | EEEKRKMLDQYNNEWEDEEYQRKGRGKGDSR | -47.06 | 51.6 | HHHHHHHHHHHHHHHHHHHHHHHCCCCCCCC |
| 289 | EEEKKKWLEKLNNEKEDKRYQRQGFGKGHQS | -47.06 | 48.4 | HHHHHHHHHHHHHHHHHHHHHHHHCCCCCCC |
| 628 | EEEKRKWLEKLNNEKEDKEYQKQGFGKGHQS | -47.06 | 45.2 | HHHHHHHHHHHHHHHHHHHHHHHHCCCCCCC |
| 258 | EEERKKQLEQYNNEYEDEEYQRKGRGKGDSR | -47.04 | 51.6 | HHHHHHHHHHHHHHHHHHHHHHHCCCCCCCC |
| 159 | EEERRKQLQQYNNESEDQRYQQQGMGKGHQP | -47.04 | 41.9 | HHHHHHHHHHHHHCHHHHHHHHHHCCCCCCC |
| 196 | EEELRKILDQINNEKEQREYQRYGFGQGHQS | -47.04 | 38.7 | HHHHHHHHHHHHHHHHHHHHHHHCCCCCCCC |
| 549 | EQEKRKILEQLNNEKEDQRYQRYGFGKGDSR | -47.03 | 45.2 | HHHHHHHHHHHHHHHHHHHHHHHCCCCCCCC |
| 954 | EEEKRKFLEQFNQEYEDQFFQQKGFGKGKQN | -47.02 | 48.4 | HHHHHHHHHHHHHHHHHHHHHHHCCCCCCCC |
| 815 | EEEKRKLLEKINNEEEDKRYQQYGFGKGDSR | -47.02 | 51.6 | HHHHHHHHHHHHHHHHHHHHHHHCCCCCCCC |
| 041 | EEEKRKLLEQINNDEEDRRYQRYGFGKGHQN | -47.00 | 38.7 | HHHHHHHHHHHHHHHHHHHHHHHCCCCCCCC |
| 193 | EEEKRKLLEQINNRKEDEEYQRNGSGKGHSY | -47.00 | 38.7 | HHHHHHHHHHHHHHHHHHHHHHCCCCCCCCC |
| 355 | EEEKRKFLDQFNNEYEHQFKQQKGFGSGKQN | -46.99 | 45.2 | HHHHHHHHHHHHHHHHHHHHHHHCCCCCCCC |
| 008 | EEEKKKLLEQINNEAEDKRYQREGKGKGHSH | -46.99 | 48.4 | HHHHHHHHHHHHHHHHHHHHHHHCCCCCCCC |
| 890 | EEEKKKLLEQINNEAEDKRYQRYGFGKGHSN | -46.99 | 48.4 | HHHHHHHHHHHHHHHHHHHHHHHCCCCCCCC |
| 831 | EEEKKKLLEQINNEEEDERYQQQGFGKGHQS | -46.99 | 45.2 | HHHHHHHHHHHHHHHHHHHHHHHCCCCCCCC |
| 166 | EEEKKKLLEQINNEEEDQRYQQYGFGKGHQS | -46.99 | 45.2 | HHHHHHHHHHHHHHHHHHHHHHHCCCCCCCC |
| 706 | EEEKKKLLEQINNEEEDQRYQQYGWGKGHTP | -46.99 | 45.2 | HHHHHHHHHHHHHHHHHHHHHHHCCCCCCCC |
| 369 | EEEKKKLLEQINNEEEDQRYQREGFGKGHQS | -46.99 | 45.2 | HHHHHHHHHHHHHHHHHHHHHHHHCCCCCCC |
| 260 | EEEKKKLLEQINNEKEDEEYQRRGKGKGHSH | -46.99 | 45.2 | HHHHHHHHHHHHHHHHHHHHHHHHCCCCCCC |
| 170 | EEEKKKLLEQINNEYEDEFYQQKGFGKGHQS | -46.99 | 48.4 | HHHHHHHHHHHHHHHHHHHHHHHCCCCCCCC |
| 219 | EEEKKKLLEQLNNEKEDKRYQRQGFGKGHSS | -46.99 | 45.2 | HHHHHHHHHHHHHHHHHHHHHHHCCCCCCCC |
| 180 | EEEKRKLLEQINNEAEDERYQQQGFGKGHQS | -46.99 | 45.2 | HHHHHHHHHHHHHHHHHHHHHHHCCCCCCCC |
| 905 | EEEKRKLLEQINNEAEDKRYQRYGFGKGHQS | -46.99 | 45.2 | HHHHHHHHHHHHHHHHHHHHHHHCCCCCCCC |
| 288 | EEEKRKLLEQINNEAEDQRYQRQGFGKGHSN | -46.99 | 45.2 | HHHHHHHHHHHHHHHHHHHHHHHCCCCCCCC |
| 892 | EEEKRKLLEQINNEEEDERYQQQGKGKGHSS | -46.99 | 41.9 | HHHHHHHHHHHHHHHHHHHHHHHCCCCCCCC |
| 519 | EEEKRKLLEQINNEEEDKRYQQYGFGKGHQS | -46.99 | 41.9 | HHHHHHHHHHHHHHHHHHHHHHHCCCCCCCC |
| 390 | EEEKRKLLEQINNEEEDKRYQREGKGKGHSH | -46.99 | 41.9 | HHHHHHHHHHHHHHHHHHHHHHHCCCCCCCC |
| 878 | EEEKRKLLEQINNEEEDKRYQREGMGKGHQP | -46.99 | 41.9 | HHHHHHHHHHHHHHHHHHHHHHHHCCCCCCC |
| 679 | EEEKRKLLEQINNEEEDKRYQREGQGKGHQS | -46.99 | 41.9 | HHHHHHHHHHHHHHHHHHHHHHHHCCCCCCC |
| 869 | EEEKRKLLEQINNEEEDKRYQRYGMGKGHQP | -46.99 | 41.9 | HHHHHHHHHHHHHHHHHHHHHHHHCCCCCCC |
| 370 | EEEKRKLLEQINNEEEDQRYQQQGWGKGHNP | -46.99 | 41.9 | HHHHHHHHHHHHHHHHHHHHHHHCCCCCCCC |
| 769 | EEEKRKLLEQINNEEEDQRYQQYGWGKGHNP | -46.99 | 41.9 | HHHHHHHHHHHHHHHHHHHHHHHCCCCCCCC |
| 154 | EEEKRKLLEQINNEEEDQRYQREGFGKGHQS | -46.99 | 41.9 | HHHHHHHHHHHHHHHHHHHHHHHCCCCCCCC |
| 547 | EEEKRKLLEQINNEEEDQRYQRYGFGKGHSN | -46.99 | 41.9 | HHHHHHHHHHHHHHHHHHHHHHHCCCCCCCC |
| 109 | EEEKRKLLEQINNEKEDEEYQRRGMGKGHQP | -46.99 | 41.9 | HHHHHHHHHHHHHHHHHHHHHHHHHCCCCCC |
| 023 | EEEKRKLLEQINNEKEDEEYQRYGMGKGHQP | -46.99 | 41.9 | HHHHHHHHHHHHHHHHHHHHHHHHHCCCCCC |
| 284 | EEEKRKLLEQINNEKEDKEYQKQGFGKGHQS | -46.99 | 41.9 | HHHHHHHHHHHHHHHHHHHHHHHHCCCCCCC |
| 476 | EEEKRKLLEQINNEKEDQEYQKQGMGKGHQP | -46.99 | 41.9 | HHHHHHHHHHHHHHHHHHHHHHHHCCCCCCC |

| | | | | |
|---|---|---|---|---|
| 762 | EEEKRKLLEQINNEKEDQEYQRAGFGKGHSN | -46.99 | 41.9 | HHHHHHHHHHHHHHHHHHHHHHHHCCCCCCC |
| 147 | EEEKRKLLEQINNEQEDERYQRQGFGKGHQS | -46.99 | 41.9 | HHHHHHHHHHHHHHHHHHHHHHHHCCCCCCC |
| 744 | EEEKRKLLEQINNEQEDQRYQREGMGKGHQP | -46.99 | 41.9 | HHHHHHHHHHHHHHHHHHHHHHHHCCCCCCC |
| 038 | EEEKRKLLEQLNNEKEDKRYQREGMGKGHQP | -46.99 | 41.9 | HHHHHHHHHHHHHHHHHHHHHHHHCCCCCCC |
| 770 | EEEKRKLLEQLNNEKEDMRYQQQGFGKGHQS | -46.99 | 41.9 | HHHHHHHHHHHHHHHHHHHHHHHHCCCCCCC |
| 073 | EEEKRKLLEQLNNEKEDQRYQKEGFGKGHQS | -46.99 | 41.9 | HHHHHHHHHHHHHHHHHHHHHHHHCCCCCCC |
| 150 | EEEKRKLLEQLNNEKEDQRYQRYGFGKGHQS | -46.99 | 41.9 | HHHHHHHHHHHHHHHHHHHHHHHCCCCCCCC |
| 595 | EEEKRKLLEQLNNEKEDQRYQRYGFGKGHSS | -46.99 | 41.9 | HHHHHHHHHHHHHHHHHHHHHHHCCCCCCCC |
| 365 | EEEWRKLMEQYNNEAEDQRYQREGMGKGHQP | -46.99 | 41.9 | HHHHHHHHHHHHHHHHHHHHHHHHHCCCCCC |
| 343 | EEEWRKLMEQYNNEYEDQEYQRTGRGKGHSF | -46.99 | 38.7 | HHHHHHHHHHHHHHHHHHHHHHCCCCCCCCC |
| 225 | EQEARKLQEQINNEEEDKRYQREGVGKGHNP | -46.99 | 38.7 | HHHHHHHHHHHHHHHHHHHHHHHHCCCCCCC |
| 779 | EQEKRKLLEQINNEAEDKRYQKEGWGKGHNP | -46.99 | 41.9 | HHHHHHHHHHHHHHHHHHHHHHHHCCCCCCC |
| 108 | EQEKRKLLEQINNEEEDQRYQRYGFGKGHQS | -46.99 | 38.7 | HHHHHHHHHHHHHHHHHHHHHHHCCCCCCCC |
| 974 | EEEKRKMLEKINNEKEHEEYQRYGFGRGHQS | -46.96 | 38.7 | HHHHHHHHHHHHHHHHHHHHHHHCCCCCCCC |
| 691 | EEEKRKFLEKYNNEYEDEEYQRNGRGKGHSY | -46.95 | 48.4 | HHHHHHHHHHHHHHHHHHHHHHCCCCCCCCC |
| 672 | EEEQRKYEQKRNQEEEDKYYQQYGQGKGHSH | -46.94 | 41.9 | CHHHHHHHHHHHHHHHHHHHHHHHHCCCCCC |
| 301 | EEEERKFQDQYNNEYEDEEYQRKGRGKGDSR | -46.94 | 51.6 | CHHHHHHHHHHHHHHHHHHHHHHCCCCCCCC |
| 439 | EEEKKKLLEQIEQEREDSEYQQKGFGKGHQS | -46.93 | 41.9 | HHHHHHHHHHHHHHHHHHHHHHHCCCCCCCC |
| 373 | EEERRKQLEKFNQEYEDKFYQKEGFGKGHQN | -46.92 | 51.6 | HHHHHHHHHHHHHHHHHHHHHHHHCCCCCCC |
| 536 | EEEKRKFLQQYNQDYEDQEYQRNGRGKGHQY | -46.85 | 41.9 | HHHHHHHHHHHHHHHHHHHHHHHCCCCCCCC |
| 393 | EEEKKKLLEKINNEAEDKRYQRQGFGKGNSN | -46.82 | 51.6 | HHHHHHHHHHHHHHHHHHHHHHHCCCCCCCC |
| 582 | EEEKRKLLDKINNEAEQKRYQREGWGQGHNP | -46.80 | 45.2 | HHHHHHHHHHHHHHHHHHHHHHHHCCCCCCC |
| 587 | EEEKRKLLDKINNEKEQEEYQRRGMGQGHQP | -46.80 | 41.9 | HHHHHHHHHHHHHHHHHHHHHHHHHCCCCCC |
| 451 | DEERKKMLEKINNEEEDKRYQQQGMGKGHQP | -46.79 | 45.2 | HHHHHHHHHHHHHHHHHHHHHHHHCCCCCCC |
| 167 | DEERKKMLEKINNEKEDKEYQKRGWGKGHNP | -46.79 | 45.2 | HHHHHHHHHHHHHHHHHHHHHHHCCCCCCCC |
| 785 | EEEKKKMLEKFNNEYEDKYYQQKGQGKGHSH | -46.79 | 51.6 | HHHHHHHHHHHHHHHHHHHHHHHHCCCCCCC |
| 460 | EEEKKKMLEKINNEAEDKRYQRYGFGKGHSS | -46.79 | 51.6 | HHHHHHHHHHHHHHHHHHHHHHHCCCCCCCC |
| 318 | EEEKKKMLEKINNEAEDKRYQRYGMGKGHQP | -46.79 | 51.6 | HHHHHHHHHHHHHHHHHHHHHHHHCCCCCCC |
| 398 | EEEKKKMLEKINNEKEDKEYQRGGFGKGHQP | -46.79 | 48.4 | HHHHHHHHHHHHHHHHHHHHHHCCCCCCCCC |
| 252 | EEEKKKMLEKINNEKEDKEYQRSGMGKGHQP | -46.79 | 51.6 | HHHHHHHHHHHHHHHHHHHHHHHHCCCCCCC |
| 617 | EEEKKKMLEKYNNEAEDIYYQKQGFGKGHSS | -46.79 | 51.6 | HHHHHHHHHHHHHHHHHHHHHHHHCCCCCCC |
| 711 | EEEKKKMLEKYNNEYEDEEYQRKGRGKGHQP | -46.79 | 48.4 | HHHHHHHHHHHHHHHHHHHHHHHHCCCCCCC |
| 638 | EEEKRKMLEKFNNEFEDKYYQQYGFGKGHSS | -46.79 | 48.4 | HHHHHHHHHHHHHHHHHHHHHHHCCCCCCCC |
| 958 | EEEKRKMLEKFNNEYEDKFYQQQGFGKGHQS | -46.79 | 51.6 | HHHHHHHHHHHHHHHHHHHHHHHCCCCCCCC |
| 378 | EEEKRKMLEKFNNEYEDKYYQQKGQGKGHSH | -46.79 | 48.4 | HHHHHHHHHHHHHHHHHHHHHHHHCCCCCCC |
| 993 | EEEKRKMLEKINNEAEDKRYQRYGFGKGHQS | -46.79 | 48.4 | HHHHHHHHHHHHHHHHHHHHHHHCCCCCCCC |
| 901 | EEEKRKMLEKINNEEEDKRYQQYGMGKGHQP | -46.79 | 45.2 | HHHHHHHHHHHHHHHHHHHHHHHHCCCCCCC |
| 468 | EEEKRKMLEKINNEKEDKEYQKQGFGKGHQS | -46.79 | 45.2 | HHHHHHHHHHHHHHHHHHHHHHHHCCCCCCC |

| | | | | |
|---|---|---|---|---|
| 539 | EEEQRKMLEKINNEEEDKRYQRQGFGKGHQN | -46.79 | 45.2 | HHHHHHHHHHHHHHHHHHHHHHHHHCCCCCCC |
| 643 | EQEKRKMLEKINNEKEDKEYQKQGMGKGHQP | -46.79 | 41.9 | HHHHHHHHHHHHHHHHHHHHHHHHHCCCCCCC |
| 377 | EEERRKQLDQMKNEKEQEEYQRYGFGQGHSN | -46.78 | 35.5 | HHHHHHHHHHHHHHHHHHHHHHHHCCCCCCCC |
| 702 | EEERKKQLEKINNEAEDKRYQREGMGKGHQP | -46.77 | 51.6 | HHHHHHHHHHHHHHHHHHHHHHHHHHCCCCCC |
| 058 | EEERKKQLEKYNNEEEDMRYQREGFGKGHQS | -46.77 | 48.4 | HHHHHHHHHHHCCHHHHHHHHHHHHCCCCCCC |
| 913 | EEERRKQLEKINNEAEDKRYQQYGFGKGHQS | -46.77 | 48.4 | HHHHHHHHHHHHHHHHHHHHHHHHCCCCCCCC |
| 234 | EEERRKQLEKINNEAEDKRYQQYGFGKGHSS | -46.77 | 48.4 | HHHHHHHHHHHHHHHHHHHHHHHHCCCCCCCC |
| 882 | EEERRKQLEKINNEEEDKRYQREGQGKGHQS | -46.77 | 45.2 | HHHHHHHHHHHHHHHHHHHHHHHHHCCCCCCC |
| 975 | EEERRKQLEKYNNESEDKRYQRYGFGKGHSS | -46.77 | 45.2 | HHHHHHHHHHHHHHHHHHHHHHHCCCCCCCCC |
| 856 | EEERRKQLEKYNNEYEDEEYQRKGRGKGHQS | -46.77 | 45.2 | HHHHHHHHHHHHHHHHHHHHHHHHHCCCCCCC |
| 813 | EQERRKQLEKINNEAEDKRYQREGMGKGHQP | -46.77 | 45.2 | HHHHHHHHHHHHHHHHHHHHHHHHHHCCCCCC |
| 739 | EEQKRKFLEKKNNEEEDKRYQQQGWGKGHTP | -46.75 | 51.6 | HHHHHHHHHHHHHHHHHHHHHHHHCCCCCCCC |
| 555 | EEEKRKLLEQINNEAEDKRYQREGMGKGHQP | -46.75 | 45.2 | HHHHHHHHHHHHHHHHHHHHHHHHHHCCCCCC |
| 320 | EEEKKKWLEKLNNEKEDMRYQREGMGKGHQP | -46.73 | 48.4 | HHHHHHHHHHHHHHHHHHHHHHHHHHCCCCCC |
| 296 | EEEEKKFQEQYNNEYEDEEFQRKGRGKGKSN | -46.73 | 41.9 | CHHHHHHHHHHHHHHHHHHHHHHHCCCCCCCC |
| 231 | EEEKKKFLEQYNNEYEDQQFQKEGFGKGKSN | -46.73 | 45.2 | HHHHHHHHHHHHHHHHHHHHHHHHHCCCCCCC |
| 802 | EEEKRKFLEQMKNEKEDQEFQRAGFGKGKQN | -46.73 | 38.7 | HHHHHHHHHHHHHHHHHHHHHHHHHCCCCCCC |
| 973 | EEEERKFREQYNQEYEDEEYQRKGRGKGHQS | -46.72 | 41.9 | CHHHHHHHHHHHHHHHHHHHHHHHHCCCCCCC |
| 484 | EEEKKKFLEQYNQEYEDEEYQRKGRGKGHQS | -46.72 | 48.4 | HHHHHHHHHHHHHHHHHHHHHHHHHCCCCCCC |
| 894 | EEEKRKFLEQFNQEFEDQFYQQKGFGKGHQS | -46.72 | 51.6 | HHHHHHHHHHHHHHHHHHHHHHHHCCCCCCCC |
| 512 | EEEKRKFLEQYNQEAEDERYQQQGMGKGHQP | -46.72 | 48.4 | HHHHHHHHHHHHHHHHHHHHHHHHHCCCCCCC |
| 401 | EEEKRKFLEQYNQEEEDMRYQREGFGKGHSS | -46.72 | 45.2 | HHHHHHHHHHHHHHHHHHHHHHHHHHCCCCCC |
| 129 | EEEKRKFLEQYNQESEDQRYQQQGMGKGHQP | -46.72 | 45.2 | HHHHHHHHHHHHHHHHHHHHHHHHHCCCCCCC |
| 893 | EEEKRKFLEQYNQESEDQRYQRYGFGKGHQS | -46.72 | 45.2 | HHHHHHHHHHHHHHHHHHHHHHHHCCCCCCCC |
| 755 | EEEKRKFLEQYNQEYEDEEYQRVGRGKGHSY | -46.72 | 45.2 | HHHHHHHHHHHHHHHHHHHHHHHHHCCCCCCC |
| 250 | EEELKKLLEQIEQEREDQEYQQKGFGKGDSR | -46.71 | 48.4 | HHHHHHHHHHHHHHHHHHHHHHHHCCCCCCCC |
| 329 | EEEKRKLLEQINNEKEDEEYQRRGWGKGHTP | -46.66 | 41.9 | HHHHHHHHHHHHHHHHHHHHHHHHCCCCCCCC |
| 990 | EEEKRKLLEQLNNEKEDQRYQREGMGKGHQP | -46.66 | 41.9 | HHHHHHHHHHHHHHHHHHHHHHHHHHCCCCCC |
| 144 | EQEKRKFLEQMKNEKEDEEFQRRGMGKGKQP | -46.66 | 35.5 | HHHHHHHHHHHHHHHHHHHHHHHHHCCCCCCC |
| 603 | EEEKKKMLDQINNEKEDRRYQQQGFGKGDSR | -46.66 | 54.8 | HHHHHHHHHHHHHHHHHHHHHHHHCCCCCCCC |
| 198 | EEEKKKLLEKINQEAEDRRYQREGMGKGHQP | -46.65 | 51.6 | HHHHHHHHHHHHHHHHHHHHHHHHHHCCCCCC |
| 644 | EEEKRKMLEQINQEEEDQRYQRYGFGKGTQN | -46.65 | 41.9 | HHHHHHHHHHHHHHHHHHHHHHHHCCCCCCCC |
| 800 | EEERKKQLEKINNEAEDKRYQRYGFGKGHQS | -46.62 | 51.6 | HHHHHHHHHHHHHHHHHHHHHHHHCCCCCCCC |
| 075 | EEEKRKWLEQLNNEKEDKRKQREGRGSGKQR | -46.61 | 38.7 | HHHHHHHHHHHHHHHHHHHHHHHHCCCCCCCC |
| 467 | EEEKRKFLEKYNNEYEDKFYQQKGFGKGHQS | -46.59 | 51.6 | HHHHHHHHHHHHHHHHHHHHHHHHCCCCCCCC |
| 989 | EEEKKKLLEKINNEEEHKRKQREGFGSGKQN | -46.56 | 38.7 | HHHHHHHHHHHHHHHHHHHHHHHHHHCCCCCC |
| 316 | EEEKRKFLEKYNQESEDKRYQQQGFGKGNSN | -46.55 | 48.4 | HHHHHHHHHHHHHHHHHHHHHHHHCCCCCCCC |
| 257 | EEEKRKFLDKYNQEEEQKRYQREGKGQGHSH | -46.54 | 45.2 | HHHHHHHHHHHHHHHHHHHHHHHHHCCCCCCC |

| | | | | |
|---|---|---|---|---|
| 404 | EEERKKQLDQINNEQEHEEKQRKGRGSGKSR | -46.54 | 41.9 | HHHHHHHHHHHHHHHHHHHHHHHCCCCCCCC |
| 188 | EEERRKQLEKINNEEEDKRYQRYGFGKGHQS | -46.53 | 45.2 | HHHHHHHHHHHHHHHHHHHHHHHCCCCCCCC |
| 675 | EEEARKFQEQYNNEAEDRRYQRYGFGKGHQN | -46.53 | 48.4 | HHHHHHHHHHHHHHHHHHHHHHHCCCCCCCC |
| 323 | EEEKKKFLEQKNNEEEDRRYQQYGMGKGHQP | -46.53 | 48.4 | HHHHHHHHHHHHHHHHHHHHHHHHCCCCCCC |
| 636 | EEEKRKFLEQFNNEFEDRRYQQEGFGKGHQS | -46.53 | 48.4 | HHHHHHHHHHHHHHHHHHHHHHHHCCCCCCC |
| 847 | EEEKRKFLEQYNNEEEDRRYQQQGNGKGHSH | -46.53 | 45.2 | HHHHHHHHHHHCHHHHHHHHHHHCCCCCCCC |
| 345 | EEEKRKLLEKINNEAEDKRYQRQGFGKGDSR | -46.51 | 54.8 | HHHHHHHHHHHHHHHHHHHHHHHCCCCCCCC |
| 310 | EEEKRKLLEKINNEQEDEEYQRKGFGKGDSR | -46.51 | 51.6 | HHHHHHHHHHHHHHHHHHHHHHHCCCCCCCC |
| 576 | EQEARKQLEKYKNEYEQEEYQRKGRGQGHQP | -46.50 | 35.5 | HHHHHHHHHHHHHHHHHHHHHHHHCCCCCCC |
| 194 | EEEKRKMLEQIEQEREDKEYQRQGFGKGHQS | -46.49 | 38.7 | HHHHHHHHHHHHHHHHHHHHHHHHCCCCCCC |
| 577 | EEEKRKMLEKFNNEYEDEEYQRKGRGKGHQS | -46.47 | 48.4 | HHHHHHHHHHHHHHHHHHHHHHHHCCCCCCC |
| 348 | EEEKKKLLEQINNEKEDQEYQRSGMGKGHQP | -46.47 | 48.4 | HHHHHHHHHHHHHHHHHHHHHHHHCCCCCCC |
| 121 | EEEKRKLLEQLNNEKEDKRYQQQGQGKGHQS | -46.47 | 41.9 | HHHHHHHHHHHHHHHHHHHHHHHCCCCCCCC |
| 429 | EEEWKKLMEQYNNEYEDQEYQRNGRGKGHSY | -46.47 | 41.9 | HHHHHHHHHHHHHHHHHHHHHHCCCCCCCCC |
| 589 | EQEYRKWLEKYKNEAEDKRYQRYGMGKGHQP | -46.47 | 41.9 | HHHHHHHHHHHHHHHHHHHHHHHHCCCCCCC |
| 747 | EQERRKFLEQYNNRYEDEEYQKEGMGKGHQP | -46.46 | 38.7 | HHHHHHHHHHHHHHHHHHHHHHHHCCCCCCC |
| 290 | EEEKRKWLEKLNNEKEDEEYQRNGRGKGHSY | -46.46 | 45.2 | HHHHHHHHHHHHHHHHHHHHHHHCCCCCCCC |
| 887 | EEEKRKFLEKFNNEKEDKRYQQQGFGKGDSR | -46.45 | 58.1 | HHHHHHHHHHHHHHHHHHHHHHHCCCCCCCC |
| 437 | EEEKRKFLEKYNNEAEDKRYQQQGFGKGDSR | -46.45 | 58.1 | HHHHHHHHHHHHHHHHHHHHHHHCCCCCCCC |
| 222 | EEEKRKILDQINNRQEQERYQREGQGQGHSS | -46.43 | 35.5 | HHHHHHHHHHHHHHHHHHHHHHHCCCCCCCC |
| 877 | EEEKRKILEKINQEAEDRRYQREGKGKGDSR | -46.42 | 54.8 | HHHHHHHHHHHHHHHHHHHHHHHCCCCCCCC |
| 319 | EEEAKKFEEQFRNEYEDQRYQQEGFGKGHSS | -46.42 | 48.4 | HHHHHHHHHHHHHHHHHHHHHHHHCCCCCCC |
| 459 | EEEARKFKEQYDNEYEDQEYQRTGRGKGHQF | -46.42 | 41.9 | HHHHHHHHHHHHHHHHHHHHHHCCCCCCCCC |
| 456 | EEEERKFQEQYNNESEDQRYQRYGFGKGHQS | -46.42 | 41.9 | CHHHHHHHHHHHCCHHHHHHHHHCCCCCCCC |
| 083 | EEEERKFQEQYNNEYEDEEYQRNGRGKGHSY | -46.42 | 41.9 | CHHHHHHHHHHHHHHHHHHHHHCCCCCCCCC |
| 495 | EEEERKFREQKNNEEEDERYQRQGWGKGHTP | -46.42 | 41.9 | CHHHHHHHHHHHHHHHHHHHHHHCCCCCCCC |
| 399 | EEEERKFREQYNNEAEDQRYQQQGMGKGHQP | -46.42 | 45.2 | CHHHHHHHHHHHHHHHHHHHHHHHCCCCCCC |
| 533 | EEEKKKFLEQFNNEEEDKRYQREGMGKGHQP | -46.42 | 51.6 | HHHHHHHHHHHCHHHHHHHHHHHHCCCCCCC |
| 409 | EEEKKKFLEQFNNEFEDQYYQQKGFGKGHQS | -46.42 | 51.6 | HHHHHHHHHHHHHHHHHHHHHHHCCCCCCCC |
| 712 | EEEKKKFLEQFNNEFEDQYYQQQGFGKGHSN | -46.42 | 51.6 | HHHHHHHHHHHHHHHHHHHHHHHCCCCCCCC |
| 640 | EEEKKKFLEQFNNEYEDKFYQQQGFGKGHSN | -46.42 | 54.8 | HHHHHHHHHHHHHHHHHHHHHHHCCCCCCCC |
| 919 | EEEKKKFLEQFNNEYEDQFYQQKGFGKGHQN | -46.42 | 54.8 | HHHHHHHHHHHHHHHHHHHHHHHCCCCCCCC |
| 115 | EEEKKKFLEQFNNEYEDQYYQQKGQGKGHQP | -46.42 | 51.6 | HHHHHHHHHHHHHHHHHHHHHHHHCCCCCCC |
| 789 | EEEKKKFLEQLNNEKEDEEYQRYGFGKGHQS | -46.42 | 48.4 | HHHHHHHHHHHHHHHHHHHHHHHCCCCCCCC |
| 904 | EEEKKKFLEQYNNEHEDERYQQQGFGKGHQS | -46.42 | 48.4 | HHHHHHHHHHHHHHHHHHHHHHHCCCCCCCC |
| 100 | EEEKKKFLEQYNNEHEDQRYQQQGMGKGHQP | -46.42 | 48.4 | HHHHHHHHHHHHHHHHHHHHHHHHCCCCCCC |
| 811 | EEEKKKFLEQYNNESEDERYQQQGFGKGHQN | -46.42 | 48.4 | HHHHHHHHHHHHHCHHHHHHHHHCCCCCCCC |
| 317 | EEEKKKFLEQYNNESEDKRYQRYGFGKGHQS | -46.42 | 48.4 | HHHHHHHHHHHHHHHHHHHHHHHCCCCCCCC |

| | | | | |
|---|---|---|---|---|
| 623 | EEEKKKFLEQYNNESEDKRYQRYGMGKGHQP | -46.42 | 48.4 | HHHHHHHHHHHHHHHHHHHHHHHHCCCCCCC |
| 757 | EEEKKKFLEQYNNESEDQRYQKEGFGKGHQS | -46.42 | 48.4 | HHHHHHHHHHHHHHHHHHHHHHHHCCCCCCC |
| 851 | EEEKKKFLEQYNNESEDQRYQRYGFGKGHQS | -46.42 | 48.4 | HHHHHHHHHHHHHCHHHHHHHHHCCCCCCCC |
| 540 | EEEKKKFLEQYNNEYEDEEYQRKGMGKGHQP | -46.42 | 48.4 | HHHHHHHHHHHHHHHHHHHHHHHHCCCCCCC |
| 235 | EEEKKKFLEQYNNEYEDEEYQRKGRGKGHQS | -46.42 | 48.4 | HHHHHHHHHHHHHHHHHHHHHHHHCCCCCCC |
| 764 | EEEKKKFLEQYNNEYEDEEYQRNGRGKGHSY | -46.42 | 48.4 | HHHHHHHHHHHHHHHHHHHHHHCCCCCCCCC |
| 551 | EEEKKKFLEQYNNEYEDEQYQRKGFGKGHQS | -46.42 | 48.4 | HHHHHHHHHHHHHHHHHHHHHHHCCCCCCCC |
| 203 | EEEKKKFLEQYNNEYEDQYYQQKGQGKGHQP | -46.42 | 48.4 | HHHHHHHHHHHHHHHHHHHHHHHHCCCCCCC |
| 405 | EEEKRKFLEQFNNEEEDKRYQREGFGKGHQS | -46.42 | 48.4 | HHHHHHHHHHHHHHHHHHHHHHHHCCCCCCC |
| 016 | EEEKRKFLEQFNNEEEDKRYQREGMGKGHQP | -46.42 | 48.4 | HHHHHHHHHHHHHHHHHHHHHHHHCCCCCCC |
| 798 | EEEKRKFLEQFNNEYEDEEYQRKGRGKGHQS | -46.42 | 48.4 | HHHHHHHHHHHHHHHHHHHHHHHHCCCCCCC |
| 788 | EEEKRKFLEQFNNEYEDEEYQRVGRGKGHSF | -46.42 | 48.4 | HHHHHHHHHHHHHHHHHHHHHHHHCCCCCCC |
| 923 | EEEKRKFLEQFNNEYEDKRYQQEGMGKGHQP | -46.42 | 48.4 | HHHHHHHHHHHHHHHHHHHHHHHHCCCCCCC |
| 483 | EEEKRKFLEQFNNEYEDKYYQKEGQGKGHSH | -46.42 | 48.4 | HHHHHHHHHHHHHHHHHHHHHHHCCCCCCCC |
| 036 | EEEKRKFLEQFNNEYEDQFYQQKGFGKGHQS | -46.42 | 51.6 | HHHHHHHHHHHHHHHHHHHHHHHCCCCCCCC |
| 984 | EEEKRKFLEQFNNEYEDQRYQQEGFGKGHQP | -46.42 | 48.4 | HHHHHHHHHHHHHHHHHHHHHHHHCCCCCCC |
| 412 | EEEKRKFLEQFNNEYEDQRYQQEGFGKGHQS | -46.42 | 48.4 | HHHHHHHHHHHHHHHHHHHHHHHHCCCCCCC |
| 059 | EEEKRKFLEQFNNEYEDQRYQQEGMGKGHQP | -46.42 | 48.4 | HHHHHHHHHHHHHHHHHHHHHHHHCCCCCCC |
| 611 | EEEKRKFLEQFNNEYEDQYYQQKGFGKGHQS | -46.42 | 48.4 | HHHHHHHHHHHHHHHHHHHHHHHCCCCCCCC |
| 857 | EEEKRKFLEQFNNEYEDQYYQQKGQGKGHSH | -46.42 | 48.4 | HHHHHHHHHHHHHHHHHHHHHHHHCCCCCCC |
| 500 | EEEKRKFLEQLNNEEEDQRYQKEGFGKGHQS | -46.42 | 45.2 | HHHHHHHHHHHHHHHHHHHHHHHHCCCCCCC |
| 507 | EEEKRKFLEQMKNEKEDEEYQRRGMGKGHQP | -46.42 | 41.9 | HHHHHHHHHHHHHHHHHHHHHHHHHCCCCCC |
| 971 | EEEKRKFLEQQSNEEEDERYQQQGFGKGHQS | -46.42 | 41.9 | HHHHHHHHHHHHHHHHHHHHHHHCCCCCCCC |
| 084 | EEEKRKFLEQYNNEAEDEQYQQRGMGKGHQP | -46.42 | 48.4 | HHHHHHHHHHHHHHHHHHHHHHHHHCCCCCC |
| 631 | EEEKRKFLEQYNNEAEDQRYQQQGFGKGHSS | -46.42 | 48.4 | HHHHHHHHHHHHHHHHHHHHHHHCCCCCCCC |
| 027 | EEEKRKFLEQYNNEAEDQRYQREGMGKGHQP | -46.42 | 48.4 | HHHHHHHHHHHHHHHHHHHHHHHHHCCCCCC |
| 423 | EEEKRKFLEQYNNEAEDQRYQRQGFGKGHQS | -46.42 | 48.4 | HHHHHHHHHHHHHHHHHHHHHHHHCCCCCCC |
| 862 | EEEKRKFLEQYNNEAEDQRYQRYGFGKGHSS | -46.42 | 48.4 | HHHHHHHHHHHHHHHHHHHHHHHCCCCCCCC |
| 424 | EEEKRKFLEQYNNEEEDMKYQREGKGKGHSS | -46.42 | 45.2 | HHHHHHHHHHHHHHHHHHHHHHHCCCCCCCC |
| 322 | EEEKRKFLEQYNNEHEDQRYQQQGMGKGHQP | -46.42 | 45.2 | HHHHHHHHHHHHHHHHHHHHHHHHCCCCCCC |
| 302 | EEEKRKFLEQYNNEHEDQRYQREGFGKGHQS | -46.42 | 45.2 | HHHHHHHHHHHHHHHHHHHHHHHHCCCCCCC |
| 485 | EEEKRKFLEQYNNESEDERYQQQGFGKGHQS | -46.42 | 45.2 | HHHHHHHHHHHHHCHHHHHHHHHCCCCCCCC |
| 924 | EEEKRKFLEQYNNESEDERYQQQGMGKGHQP | -46.42 | 45.2 | HHHHHHHHHHHHHCHHHHHHHHHHCCCCCCC |
| 821 | EEEKRKFLEQYNNESEDQRYQKEGFGKGHSS | -46.42 | 45.2 | HHHHHHHHHHHHHHHHHHHHHHHHCCCCCCC |
| 793 | EEEKRKFLEQYNNESEDQRYQRYGFGKGHQS | -46.42 | 45.2 | HHHHHHHHHHHHHCHHHHHHHHHCCCCCCCC |
| 342 | EEEKRKFLEQYNNESEDQRYQRYGMGKGHQP | -46.42 | 45.2 | HHHHHHHHHHHHHCHHHHHHHHHHCCCCCCC |
| 295 | EEEKRKFLEQYNNESEDSRYQQQGMGKGHQP | -46.42 | 45.2 | HHHHHHHHHHHHHHHHHHHHHHHHCCCCCCC |
| 629 | EEEKRKFLEQYNNEYEDEEYQRKGRGKGHQN | -46.42 | 45.2 | HHHHHHHHHHHHHHHHHHHHHHHHCCCCCCC |

| | | | | |
|---|---|---|---|---|
| 336 | EEEKRKFLEQYNNEYEDEEYQRKGRGKGHQP | -46.42 | 45.2 | HHHHHHHHHHHHHHHHHHHHHHHHCCCCCCC |
| 838 | EEEKRKFLEQYNNEYEDEEYQRKGRGKGHQS | -46.42 | 45.2 | HHHHHHHHHHHHHHHHHHHHHHHHCCCCCCC |
| 963 | EEEKRKFLEQYNNEYEDEEYQRNGRGKGHSY | -46.42 | 45.2 | HHHHHHHHHHHHHHHHHHHHHHCCCCCCCCC |
| 921 | EEEKRKFLEQYNNEYEDQEYQRTGRGKGHSF | -46.42 | 45.2 | HHHHHHHHHHHHHHHHHHHHHHCCCCCCCCC |
| 230 | EEEKRKFLEQYNNEYEDQQYQQQGMGKGHQP | -46.42 | 45.2 | HHHHHHHHHHHHHHHHHHHHHHHHCCCCCCC |
| 669 | EEEKRKFLEQYNNEYEDQYYQQNGQGKGHSY | -46.42 | 45.2 | HHHHHHHHHHHHHHHHHHHHHHHCCCCCCCC |
| 183 | EEEQRKFEEQRNNEEEDSRYQQQGWGKGHTP | -46.42 | 41.9 | HHHHHHHHHHHHHHHHHHHHHHHCCCCCCCC |
| 242 | EEEQRKFLEQYNNEYEDEQYQQQGWGKGHNP | -46.42 | 45.2 | CHHHHHHHHHHHHHHHHHHHHHHCCCCCCCC |
| 063 | EEERRKFLEQQNNEEEDERYQQQGFGKGHSN | -46.42 | 45.2 | HHHHHHHHHHHHHHHHHHHHHHHCCCCCCCC |
| 480 | EQEKRKFLEQFNNEFEDQYYQQKGQGKGHSH | -46.42 | 45.2 | HHHHHHHHHHHHHHHHHHHHHHHHCCCCCCC |
| 907 | EQEKRKFLEQYNNEHEDERYQQQGMGKGHQP | -46.42 | 41.9 | HHHHHHHHHHHHHHHHHHHHHHHHCCCCCCC |
| 740 | EQEKRKFLEQYNNEYEDEEYQQTGRGKGHQF | -46.42 | 41.9 | HHHHHHHHHHHHHHCHHHHHHHHHCCCCCCC |
| 529 | EQEKRKFLEQYNNEYEDEEYQRKGRGKGHQS | -46.42 | 41.9 | HHHHHHHHHHHHHHHHHHHHHHHHCCCCCCC |
| 854 | EQEKRKFLEKMENEREDKEYQKQGFGKGNQN | -46.41 | 41.9 | HHHHHHHHHHHHHHHHHHHHHHHHCCCCCCC |
| 331 | EEEKRKILDQINNEEEQQRYQKEGFGQGHQS | -46.40 | 38.7 | HHHHHHHHHHHCHHHHHHHHHHHCCCCCCCC |
| 545 | EEEKRKLLDQLNQEKEQQRYQRYGFGQGHQS | -46.39 | 38.7 | HHHHHHHHHHHHHHHHHHHHHHHCCCCCCCC |
| 078 | EEEKRKFLQQFNQEYEDQFFQQKGFGKGKQN | -46.39 | 48.4 | HHHHHHHHHHHHHHHHHHHHHHHCCCCCCCC |
| 487 | EEEKRKFLEQMNNRQEDERYQREGKGKGHSH | -46.38 | 41.9 | HHHHHHHHHHHHHHHHHHHHHHHCCCCCCCC |
| 330 | EEQKRKMLEQKNNDEEDQRYQQQGWGKGHNP | -46.36 | 41.9 | HHHHHHHHHHHCCHHHHHHHHHHCCCCCCCC |
| 703 | EEEKRKMLEKIENEREDEEYQRKGRGKGHQS | -46.36 | 41.9 | HHHHHHHHHHHHHHHHHHHHHHHHCCCCCCC |
| 727 | EEEKKKFLEKYNNEYEDKFYQQKGFGKGHQS | -46.36 | 54.8 | HHHHHHHHHHHHHHHHHHHHHHHCCCCCCCC |
| 609 | EEEKKKLLEKINNEAEDRRYQQYGFGKGHQS | -46.36 | 51.6 | HHHHHHHHHHHHHHHHHHHHHHHCCCCCCCC |
| 695 | EEEKKKLLEKINNEAEDRRYQRQGFGKGHQN | -46.36 | 51.6 | HHHHHHHHHHHHHHHHHHHHHHHHCCCCCCC |
| 553 | EEEKRKLLEKINNEEEDRRYQQYGFGKGHSS | -46.36 | 45.2 | HHHHHHHHHHHHHHHHHHHHHHHCCCCCCCC |
| 116 | EEEKRKLLEKLNNEKEDRRYQREGQGKGHQS | -46.36 | 45.2 | HHHHHHHHHHHHHHHHHHHHHHHHCCCCCCC |
| 134 | EEEKRKLLQQINNEKEDEEYQRRGRGKGHQP | -46.35 | 41.9 | HHHHHHHHHHHHHHHHHHHHHHHHCCCCCCC |
| 277 | EEEKKKMLEQYNNESEDQRFQRYGFGKGKQN | -46.34 | 41.9 | HHHHHHHHHHHHHHHHHHHHHHHCCCCCCCC |
| 596 | EEEKRKMLEQINNEAEDKRFQRQGFGKGKQN | -46.34 | 41.9 | HHHHHHHHHHHHHHHHHHHHHHHHCCCCCCC |
| 070 | EEEKRKLLDKINNEEEQKRYQREGMGQGHQP | -46.33 | 41.9 | HHHHHHHHHHHCHHHHHHHHHHHHCCCCCCC |
| 104 | EEEKRKLLDKINNEEEQKRYQRYGFGQGHSN | -46.33 | 41.9 | HHHHHHHHHHHCHHHHHHHHHHHCCCCCCCC |
| 543 | EEEKKKMLEQFNQEYEDEEYQRKGRGKGHQS | -46.32 | 48.4 | HHHHHHHHHHHHHHHHHHHHHHHHCCCCCCC |
| 949 | EEEKKKMLEQINQEKEDEEYQRYGMGKGHQP | -46.32 | 45.2 | HHHHHHHHHHHHHHHHHHHHHHHHHCCCCCC |
| 246 | EEEKKKMLEQYNQESEDQRYQKEGFGKGHQS | -46.32 | 45.2 | HHHHHHHHHHHHHHHHHHHHHHHHCCCCCCC |
| 030 | EEEKRKMLEQYNQESEDQRYQQQGWGKGHNP | -46.32 | 41.9 | HHHHHHHHHHHHHHHHHHHHHHHHCCCCCCC |
| 172 | EQEKRKMLEQYNQEYEDEEYQRKGRGKGHQP | -46.32 | 38.7 | HHHHHHHHHHHHHHHHHHHHHHHHCCCCCCC |
| 982 | EEEKKKWLEQLNNRQEDERYQRYGFGKGHQS | -46.30 | 41.9 | HHHHHHHHHHHHHHHHHHHHHHHCCCCCCCC |
| 777 | EEERRKQLEQINQEEEDQRYQRYGFGKGHQS | -46.30 | 41.9 | HHHHHHHHHHHHHHHHHHHHHHHCCCCCCCC |
| 422 | EEERRKQLEQINQEKEDEEYQRYGFGKGHSS | -46.30 | 41.9 | HHHHHHHHHHHHHHHHHHHHHHHCCCCCCCC |

| | | | | |
|---|---|---|---|---|
| 206 | EEEFRKWMEQLNNEAEDQRYQKEGQGKGHQS | -46.30 | 41.9 | HHHHHHHHHHHHHHHHHHHHHHHHCCCCCCC |
| 264 | EEEKKKWLEQLNNEKEDQRYQREGFGKGHSS | -46.29 | 45.2 | HHHHHHHHHHHHHHHHHHHHHHHHCCCCCCC |
| 625 | EEEKKKWLEQLNNEQEDMRYQRYGFGKGHQS | -46.29 | 45.2 | HHHHHHHHHHHHHHHHHHHHHHHCCCCCCCC |
| 594 | EEEKRKWLEQLSNEKEDERYQQQGFGKGHQS | -46.29 | 38.7 | HHHHHHHHHHHHHHHHHHHHHHHCCCCCCCC |
| 723 | EEEQKKQQEKYSNESEQKRYQQYGMGQGHQP | -46.28 | 35.5 | HHHHHHHHHHHCHHHHHHHHHHHCCCCCCCC |
| 879 | EEEQRKQQEKYNNEYEQKEYQRNGRGQGHSY | -46.28 | 35.5 | HHHHHHHHHHHHHHHHHHHHHHHCCCCCCCC |
| 465 | EEQKRKLLEQINNEQEDEEYQRKGRGKGHQS | -46.28 | 45.2 | HHHHHHHHHHHHHHHHHHHHHHHHCCCCCCC |
| 801 | EEERRKQLEQFNNEFEDKFFQQKGFGKGKSR | -46.28 | 48.4 | HHHHHHHHHHHHHHHHHHHHHHHCCCCCCCC |
| 291 | EEEKRKLLEQINNEAEDKRYQREGFGKGDSR | -46.25 | 51.6 | HHHHHHHHHHHHHHHHHHHHHHHHCCCCCCC |
| 799 | EEEKRKFLDKYNNESEQKRYQRYGFGQGHQS | -46.24 | 45.2 | HHHHHHHHHHHHHHHHHHHHHHHCCCCCCCC |
| 217 | EEEKRKMLDAINNRAEDQRYQRYGFGKGHQS | -46.23 | 45.2 | HHHHHHHHHHHHHHHHHHHHHHHCCCCCCCC |
| 478 | EEEKRKMLEQINNEKEHQEYQRRGQGRGHQS | -46.20 | 35.5 | HHHHHHHHHHHHHHHHHHHHHHHHCCCCCCC |
| 338 | EQEKRKFLEQYNNEYEDEYYQQKGQGKGHSH | -46.19 | 41.9 | HHHHHHHHHHHHHHHHHHHHHHHHCCCCCCC |
| 639 | EEEKRKMLQKFNNEYEDKYYQQKGFGKGHQS | -46.16 | 48.4 | HHHHHHHHHHHHHHHHHHHHHHHCCCCCCCC |
| 035 | EEERRKQLDKINNEAEDRRYQQYGFGKGNSN | -46.15 | 51.6 | HHHHHHHHHHHHHHHHHHHHHHHCCCCCCCC |
| 701 | EEEERKFQEKYNNEYEQEEYQRKGRGRGDSR | -46.14 | 45.2 | CHHHHHHHHHHHHHHHHHHHHHHCCCCCCCC |
| 265 | EEEKKKMLEQFNNEFEDRYYQQQGQGKGHSY | -46.14 | 48.4 | HHHHHHHHHHHHHHHHHHHHHHHHCCCCCCC |
| 554 | EEEKRKMLEQINNEQEDRYYQQYGQGKGHQP | -46.14 | 41.9 | HHHHHHHHHHHHHHHHHHHHHHHHHCCCCCC |
| 024 | EEEKRKMLEQYNNEYEDREYQRTGRGKGHQS | -46.14 | 41.9 | HHHHHHHHHHHHHHHHHHHHHHCCCCCCCCC |
| 168 | EEELKKILEKLNNEKEQKRYQRQGFGQGHQS | -46.11 | 41.9 | HHHHHHHHHHHHHHHHHHHHHHHCCCCCCCC |
| 852 | EEEQRKILEKLNNEKEQKRYQQYGMGQGHQP | -46.11 | 38.7 | HHHHHHHHHHHHHHHHHHHHHHHHCCCCCCC |
| 056 | EEEKRKLLEQINNEEEDQRYQRYGFGKGDSR | -46.09 | 48.4 | HHHHHHHHHHHHHHHHHHHHHHHCCCCCCCC |
| 586 | EEEKRKLLEQINNEKEDQEYQKQGFGKGDSR | -46.09 | 48.4 | HHHHHHHHHHHHHHHHHHHHHHHCCCCCCCC |
| 690 | EEEKKKLLDQINNEEEQQRYQQQGFGQGHQS | -46.09 | 41.9 | HHHHHHHHHHHCHHHHHHHHHHHCCCCCCCC |
| 447 | EEEKKKLLDQINNEYEQKYYQQKGQGQGHSH | -46.09 | 41.9 | HHHHHHHHHHHHHHHHHHHHHHHHCCCCCCC |
| 819 | EEEKRKLLDQINNEEEQQRYQKEGFGQGHQS | -46.09 | 38.7 | HHHHHHHHHHHCHHHHHHHHHHHCCCCCCCC |
| 472 | EEEKRKLLDQINNEKEQEEYQRYGFGQGHQS | -46.09 | 38.7 | HHHHHHHHHHHHHHHHHHHHHHHCCCCCCCC |
| 386 | EEEKRKLLDQINNEQEQKSYQRYGFGQGHQS | -46.09 | 38.7 | HHHHHHHHHHHHHHHHHHHHHHHCCCCCCCC |
| 619 | EEEKRKLLDQINNEYEQEEYQRKGRGQGHQS | -46.09 | 38.7 | HHHHHHHHHHHHHHHHHHHHHHHCCCCCCCC |
| 929 | EEEKRKFLEKYNQEYEDREYQRTGRGKGHQF | -46.09 | 48.4 | HHHHHHHHHHHHHHHHHHHHHHCCCCCCCCC |
| 997 | EEEKRKFLEKYNQEYEDREYQRTGRGKGHSF | -46.09 | 48.4 | HHHHHHHHHHHHHHHHHHHHHHCCCCCCCCC |
| 556 | EEERRKQLEQINQEAEDQRYQREGKGKGHSH | -46.06 | 45.2 | HHHHHHHHHHHHHHHHHHHHHHHCCCCCCCC |
| 525 | EEEKRKMLEKINNEKEDKRYQQQGFGKGDSR | -46.06 | 51.6 | HHHHHHHHHHHHHHHHHHHHHHHCCCCCCCC |
| 996 | EEEKRKLLEQINNEEEDKRYQQYGFGKGNQN | -46.05 | 41.9 | HHHHHHHHHHHHHHHHHHHHHHHHCCCCCCC |
| 305 | EEERRKQLDQYNNEEEHKRYQREGFGQGHSS | -46.05 | 38.7 | HHHHHHHHHHHCCHHHHHHHHHHHCCCCCCC |
| 137 | EEERRKQLEKLNNEKEDKRYQKEGFGKGDSR | -46.03 | 51.6 | HHHHHHHHHHHHHHHHHHHHHHHCCCCCCCC |
| 883 | EEELKKLLEKINNEEEHKEYQRTGRGQGHQF | -46.03 | 41.9 | HHHHHHHHHHHCHHHHHHHHHHHHCCCCCCC |
| 009 | EEERRKQLEKKNNEEEQKRYQRYGFGQGHQS | -46.03 | 38.7 | HHHHHHHHHHHCHHHHHHHHHHHCCCCCCCC |

| | | | | |
|---|---|---|---|---|
| 428 | EEERRKQLEKYNNEAEQKRYQRYGFGQGHQS | -46.03 | 41.9 | HHHHHHHHHHHHHHHHHHHHHHHCCCCCCCC |
| 089 | EEEKKKMLEQFNNEEEDQRYQQEGFGKGHQS | -46.03 | 48.4 | HHHHHHHHHHHCHHHHHHHHHHHHCCCCCCC |
| 421 | EEEKKKMLEQFNNEFEDQYYQQQGQGKGHSY | -46.03 | 48.4 | HHHHHHHHHHHHHHHHHHHHHHHHCCCCCCC |
| 748 | EEEKKKMLEQFNNEKEDQRYQQQGMGKGHQP | -46.03 | 48.4 | HHHHHHHHHHHHHHHHHHHHHHHHCCCCCCC |
| 532 | EEEKKKMLEQFNNEYEDQYYQQKGFGKGHQN | -46.03 | 48.4 | HHHHHHHHHHHHHHHHHHHHHHHCCCCCCCC |
| 496 | EEEKKKMLEQFNNEYEDQYYQQKGFGKGHQS | -46.03 | 48.4 | HHHHHHHHHHHHHHHHHHHHHHHCCCCCCCC |
| 396 | EEEKKKMLEQINNEAEDKRYQREGKGKGHSA | -46.03 | 48.4 | HHHHHHHHHHHHHHHHHHHHHHHCCCCCCCC |
| 034 | EEEKKKMLEQINNEAEDQRYQKEGMGKGHQP | -46.03 | 48.4 | HHHHHHHHHHHHHHHHHHHHHHHHCCCCCCC |
| 805 | EEEKKKMLEQINNEEEDERYQQQGFGKGHQS | -46.03 | 45.2 | HHHHHHHHHHHHHHHHHHHHHHHCCCCCCCC |
| 818 | EEEKKKMLEQINNEEEDERYQQQGMGKGHQP | -46.03 | 45.2 | HHHHHHHHHHHHHHHHHHHHHHHHCCCCCCC |
| 521 | EEEKKKMLEQINNEEEDKKYQREGKGKGHSS | -46.03 | 45.2 | HHHHHHHHHHHHHHHHHHHHHHHCCCCCCCC |
| 092 | EEEKKKMLEQINNEEEDKRYQQQGWGKGHNP | -46.03 | 45.2 | HHHHHHHHHHHHHHHHHHHHHHHCCCCCCCC |
| 125 | EEEKKKMLEQINNEEEDKRYQQYGFGKGHQS | -46.03 | 45.2 | HHHHHHHHHHHHHHHHHHHHHHHCCCCCCCC |
| 367 | EEEKKKMLEQINNEEEDKRYQREGFGKGHQS | -46.03 | 45.2 | HHHHHHHHHHHHHHHHHHHHHHHHCCCCCCC |
| 119 | EEEKKKMLEQINNEEEDKRYQREGMGKGHQP | -46.03 | 45.2 | HHHHHHHHHHHHHHHHHHHHHHHHCCCCCCC |
| 402 | EEEKKKMLEQINNEEEDKRYQRYGFGKGHQS | -46.03 | 45.2 | HHHHHHHHHHHHHHHHHHHHHHHCCCCCCCC |
| 662 | EEEKKKMLEQINNEEEDQRYQRYGFGKGHQS | -46.03 | 45.2 | HHHHHHHHHHHHHHHHHHHHHHHCCCCCCCC |
| 647 | EEEKKKMLEQINNEEEDQRYQRYGMGKGHQP | -46.03 | 45.2 | HHHHHHHHHHHHHHHHHHHHHHHHCCCCCCC |
| 714 | EEEKKKMLEQINNEKEDEEYQQRGMGKGHQP | -46.03 | 45.2 | HHHHHHHHHHHHHHHHHHHHHHHHCCCCCCC |
| 835 | EEEKKKMLEQINNEKEDEEYQRYGFGKGHQS | -46.03 | 45.2 | HHHHHHHHHHHHHHHHHHHHHHHCCCCCCCC |
| 742 | EEEKKKMLEQYNNEAEDQRYQRYGMGKGHQP | -46.03 | 48.4 | HHHHHHHHHHHHHHHHHHHHHHHHCCCCCCC |
| 419 | EEEKKKMLEQYNNEFEDQQYQRTGMGKGHQF | -46.03 | 45.2 | HHHHHHHHHHHHHHHHHHHHHHHCCCCCCCC |
| 899 | EEEKKKMLEQYNNEHEDQRYQQQGFGKGHQS | -46.03 | 45.2 | HHHHHHHHHHHHHHHHHHHHHHHCCCCCCCC |
| 162 | EEEKKKMLEQYNNESEDKRYQQQGFGKGHQS | -46.03 | 45.2 | HHHHHHHHHHHHHHHHHHHHHHHCCCCCCCC |
| 876 | EEEKKKMLEQYNNEYEDEEYQRKGRGKGHQS | -46.03 | 45.2 | HHHHHHHHHHHHHHHHHHHHHHHHCCCCCCC |
| 503 | EEEKKKMLEQYNNEYEDKYYQQKGFGKGHSS | -46.03 | 45.2 | HHHHHHHHHHHHHHHHHHHHHHHCCCCCCCC |
| 987 | EEEKKKMLEQYNNEYEDQFYQQKGMGKGHQP | -46.03 | 48.4 | HHHHHHHHHHHHHHHHHHHHHHHHCCCCCCC |
| 684 | EEEKRKMLEQINNEEEDERYQQQGFGKGHSS | -46.03 | 41.9 | HHHHHHHHHHHHHHHHHHHHHHHCCCCCCCC |
| 564 | EEEKRKMLEQINNEEEDKRYQREGFGKGHQS | -46.03 | 41.9 | HHHHHHHHHHHHHHHHHHHHHHHHCCCCCCC |
| 045 | EEEKRKMLEQINNEEEDKRYQREGMGKGHQP | -46.03 | 41.9 | HHHHHHHHHHHHHHHHHHHHHHHHCCCCCCC |
| 366 | EEEKRKMLEQINNEEEDQRYQQYGFGKGHQS | -46.03 | 41.9 | HHHHHHHHHHHHHHHHHHHHHHHCCCCCCCC |
| 454 | EEEKRKMLEQINNEEEDQRYQQYGMGKGHQP | -46.03 | 41.9 | HHHHHHHHHHHHHHHHHHHHHHHHCCCCCCC |
| 981 | EEEKRKMLEQINNEEEDQRYQRYGFGKGHQS | -46.03 | 41.9 | HHHHHHHHHHHHHHHHHHHHHHHCCCCCCCC |
| 054 | EEEKRKMLEQINNEEEDQRYQRYGFGKGHSS | -46.03 | 41.9 | HHHHHHHHHHHHHHHHHHHHHHHCCCCCCCC |
| 506 | EEEKRKMLEQINNEKEDQEYQRSGQGKGHQS | -46.03 | 45.2 | HHHHHHHHHHHHHHHHHHHHHHHCCCCCCCC |
| 307 | EEEKRKMLEQINNEQEDERYQQQGMGKGHQP | -46.03 | 41.9 | HHHHHHHHHHHHHHHHHHHHHHHHCCCCCCC |
| 720 | EEEKRKMLEQINNEQEDERYQRQGWGKGHNP | -46.03 | 41.9 | HHHHHHHHHHHHHHHHHHHHHHHCCCCCCCC |
| 255 | EEEKRKMLEQINNEQEDEYYQQYGQGKGHQP | -46.03 | 41.9 | HHHHHHHHHHHHHHHHHHHHHHHHHCCCCCC |

| | | | | |
|---|---|---|---|---|
| 538 | EEEKRKMLEQYNNEEEDMKYQREGMGKGHQP | -46.03 | 41.9 | HHHHHHHHHHHHHHHHHHHHHHHHHCCCCCC |
| 767 | EEEKRKMLEQYNNEYEDKQYQKEGFGKGHQS | -46.03 | 41.9 | HHHHHHHHHHHHHHHHHHHHHHHHCCCCCCC |
| 940 | EEEKRKMLEQYNNEYEDQEYQRNGRGKGHSY | -46.03 | 41.9 | HHHHHHHHHHHHHHHHHHHHHHCCCCCCCCC |
| 926 | EEEQRKMLEQINNEAEDKRYQREGMGKGHQP | -46.03 | 45.2 | HHHHHHHHHHHHHHHHHHHHHHHHHCCCCCC |
| 655 | EQEKRKMLEQFNNEYEDKYYQQKGFGKGHQS | -46.03 | 41.9 | HHHHHHHHHHHHHHHHHHHHHHHCCCCCCCC |
| 328 | EQEKRKMLEQINNEEEDKRYQRQGFGKGHQS | -46.03 | 38.7 | HHHHHHHHHHHHHHHHHHHHHHHHCCCCCCC |
| 897 | EEERRKQLEKYNNESEQERYQQQGMGQGHQP | -46.02 | 38.7 | HHHHHHHHHHHHHHHHHHHHHHHHCCCCCCC |
| 816 | EEEKRKFLDKFNNEYEQKYYQQKGFGQGHSN | -46.01 | 48.4 | HHHHHHHHHHHHHHHHHHHHHHHCCCCCCCC |
| 573 | EEEKKKFLEKYEDEYRDEEYQRKGRGKGHQS | -46.01 | 45.2 | HHHHHHHHHHHHHHHHHHHHHHHHCCCCCCC |
| 151 | EEERKKQLEQINNEEEDKRYQREGFGKGHQS | -46.00 | 45.2 | HHHHHHHHHHHHHHHHHHHHHHHHCCCCCCC |
| 262 | EEERKKQLEQINNEEEDKRYQREGVGKGHNP | -46.00 | 45.2 | HHHHHHHHHHHHHHHHHHHHHHHHCCCCCCC |
| 618 | EEERKKQLEQINNEEEDQRYQREGVGKGHQP | -46.00 | 45.2 | HHHHHHHHHHHHHHHHHHHHHHHHCCCCCCC |
| 668 | EEERKKQLEQYNNEHEDERYQQQGFGKGHSS | -46.00 | 45.2 | HHHHHHHHHHHHHHHHHHHHHHHCCCCCCCC |
| 761 | EEERRKQLEQFNNEYEDEEYQRQGRGKGHQP | -46.00 | 45.2 | HHHHHHHHHHHHHHHHHHHHHHHHCCCCCCC |
| 773 | EEERRKQLEQINNEAEDERYQQQGFGKGHQS | -46.00 | 45.2 | HHHHHHHHHHHHHHHHHHHHHHHCCCCCCCC |
| 490 | EEERRKQLEQINNEAEDERYQQQGMGKGHQP | -46.00 | 45.2 | HHHHHHHHHHHHHHHHHHHHHHHHCCCCCCC |
| 197 | EEERRKQLEQINNEAEDKRYQREGMGKGHQP | -46.00 | 45.2 | HHHHHHHHHHHHHHHHHHHHHHHHHCCCCCC |
| 772 | EEERRKQLEQINNEAEDKRYQREGQGKGHQS | -46.00 | 45.2 | HHHHHHHHHHHHHHHHHHHHHHHHCCCCCCC |
| 018 | EEERRKQLEQINNEAEDKRYQRQGFGKGHQS | -46.00 | 45.2 | HHHHHHHHHHHHHHHHHHHHHHHHCCCCCCC |
| 199 | EEERRKQLEQINNEAEDQRYQRQGFGKGHSS | -46.00 | 45.2 | HHHHHHHHHHHHHHHHHHHHHHHCCCCCCCC |
| 585 | EEERRKQLEQINNEAEDQRYQRYGMGKGHQP | -46.00 | 45.2 | HHHHHHHHHHHHHHHHHHHHHHHHCCCCCCC |
| 559 | EEERRKQLEQINNEEEDERYQQQGQGKGHQS | -46.00 | 41.9 | HHHHHHHHHHHHHHHHHHHHHHHHCCCCCCC |
| 646 | EEERRKQLEQINNEEEDKEYQRVGRGKGHQS | -46.00 | 41.9 | HHHHHHHHHHHHHHHHHHHHHHHHCCCCCCC |
| 112 | EEERRKQLEQINNEKEDEEYQRRGMGKGHQP | -46.00 | 41.9 | HHHHHHHHHHHHHHHHHHHHHHHHHCCCCCC |
| 185 | EEERRKQLEQINNEKEDEEYQRYGFGKGHQS | -46.00 | 41.9 | HHHHHHHHHHHHHHHHHHHHHHHCCCCCCCC |
| 663 | EEERRKQLEQINNEQEDEKYQQEGMGKGHQP | -46.00 | 41.9 | HHHHHHHHHHHHHHHHHHHHHHHHCCCCCCC |
| 681 | EEERRKQLEQISNEKEDQEYQKQGRGKGHQP | -46.00 | 38.7 | HHHHHHHHHHHHHHHHHHHHHHHHCCCCCCC |
| 133 | EQERRKQLEQYNNEYEDQYYQQKGQGKGHSH | -46.00 | 38.7 | HHHHHHHHHHHHHHHHHHHHHHHHCCCCCCC |
| 175 | EEEKRKFLEKFNNEYEHKYKQQKGFGSGKQN | -45.99 | 41.9 | HHHHHHHHHHHHHHHHHHHHHHHCCCCCCCC |
| 535 | EEERRKQLEQYNNRSEDERYQQQGMGKGHQP | -45.98 | 38.7 | HHHHHHHHHHHHHCHHHHHHHHHHCCCCCCC |
| 652 | EEEQRKFLEKYNNESEDKRYQQQGFGKGDSR | -45.94 | 54.8 | CHHHHHHHHHHHHHHHHHHHHHHCCCCCCCC |
| 708 | EEEKRKFLDQYRHEYEQEFYQQKGFGQGHSS | -45.94 | 45.2 | HHHHHHHHHHHHHHHHHHHHHHHCCCCCCCC |
| 169 | EEEKRKLLEQINNEAEDQRYQQQGFGKGDSR | -45.93 | 51.6 | HHHHHHHHHHHHHHHHHHHHHHHCCCCCCCC |
| 042 | EEEKRKQLEQYNQEYEQEEYQRVGRGQGHQF | -45.92 | 35.5 | HHHHHHHHHHHHHHHHHHHHHHHHHCCCCCC |
| 970 | EEEKRKLLDQINQEAEQKRYQREGMGQGHQP | -45.92 | 41.9 | HHHHHHHHHHHHHHHHHHHHHHHHHCCCCCC |
| 730 | EQEKRHILEQYNQEYEDEEYQRKGRGKGHQP | -45.92 | 38.7 | HHHHHHHHHHHHHHHHHHHHHHHHCCCCCCC |
| 364 | EEEERKFQEQYNNEYEDEEYQRKGRGKGHQP | -45.90 | 41.9 | CHHHHHHHHHHHHHHHHHHHHHHHCCCCCCC |
| 842 | EEEKRKFLEQFNNEYEDQFYQKEGFGKGHQS | -45.90 | 51.6 | HHHHHHHHHHHHHHHHHHHHHHHHCCCCCCC |

| | | | | |
|---|---|---|---|---|
| 420 | EEEKKKLLDKLNQEKEQKRYQRYGFGQGDSR | -45.90 | 51.6 | HHHHHHHHHHHHHHHHHHHHHHHCCCCCCCC |
| 698 | EEEKRKLLEQIQKDREDREYQREGMGKGHQP | -45.90 | 35.5 | HHHHHHHHHHHHHHHHHHHHHHHHCCCCCCC |
| 650 | EEEKKKMLEKINNEAEDKRYQREGFGKGNQN | -45.86 | 51.6 | HHHHHHHHHHHHHHHHHHHHHHHHCCCCCCC |
| 806 | EEEKKKMLDKINNEAEQKRYQRYGMGQGHQP | -45.85 | 48.4 | HHHHHHHHHHHHHHHHHHHHHHHHCCCCCCC |
| 823 | EEEKKKMLEKFNQEYEDKYYQQKGFGKGDSR | -45.85 | 58.1 | HHHHHHHHHHHHHHHHHHHHHHHCCCCCCCC |
| 654 | EEERRKQLEKINNEQEDKSYQRHGFGKGNSN | -45.83 | 45.2 | HHHHHHHHHHHHHHHHHHHHHHCCCCCCCCC |
| 229 | EEEKKKFLEQYNNESEDQRYQRYGFGKGHQN | -45.82 | 48.4 | HHHHHHHHHHHHHCHHHHHHHHHCCCCCCCC |
| 978 | EEEWKKLLDQIKNEKEQEEYQRRGLGQGHQP | -45.82 | 41.9 | HHHHHHHHHHHHHHHHHHHHHHHHHCCCCCC |
| 278 | EEERRKQLEKINQEEEDKRYQREGFGKGDSR | -45.82 | 51.6 | HHHHHHHHHHHHHHHHHHHHHHHHCCCCCCC |
| 508 | EEEKRKMLEQINNEEEDKRYQREGKGKGHSH | -45.80 | 41.9 | HHHHHHHHHHHHHHHHHHHHHHHCCCCCCCC |
| 012 | EEEKRKMLEQYNNEYEDQQYQKEGFGKGHQS | -45.79 | 41.9 | HHHHHHHHHHHHHHHHHHHHHHHHCCCCCCC |
| 263 | EEEKRKFLDKKNNEEEQKRYQQYGKGQGHSH | -45.77 | 45.2 | HHHHHHHHHHCCHHHHHHHHHHHHCCCCCCC |
| 106 | EEEKRKFLDKYNNEAEQKRYQRQGFGQGHQS | -45.77 | 48.4 | HHHHHHHHHHHHHHHHHHHHHHHCCCCCCCC |
| 985 | EEEQRKQLEQFKNEYEQEFYQQQGFGQGHQS | -45.76 | 38.7 | HHHHHHHHHHHHHHHHHHHHHHHCCCCCCCC |
| 215 | EEEKRKLLEQINNEAEDKRYQRYGFGKGDSR | -45.74 | 51.6 | HHHHHHHHHHHHHHHHHHHHHHHCCCCCCCC |
| 505 | EEEKRKLLEQINNEEEDKRYQREGFGKGDSR | -45.74 | 48.4 | HHHHHHHHHHHHHHHHHHHHHHHHCCCCCCC |
| 470 | EEEKRKLLDQINNEYEQREYQRKGFGQGHQS | -45.72 | 38.7 | HHHHHHHHHHHHHHHHHHHHHHHHCCCCCCC |
| 693 | EEEEKKRQEQYNNEYEDQEYQRTGRGKGHQF | -45.72 | 41.9 | HHHHHHHHHHHHHHHHHHHHHHCCCCCCCCC |
| 174 | EEEKRKLLDKINQEAEQRRYQREGMGQGHQP | -45.71 | 45.2 | HHHHHHHHHHHHHHHHHHHHHHHHCCCCCCC |
| 021 | EEEKRKMLEQINNEAEDQRYQQQGMGKGHQP | -45.70 | 45.2 | HHHHHHHHHHHHHHHHHHHHHHHHCCCCCCC |
| 149 | EEEKRKFLEQFNNEEEDKRYQREGFGKGDSR | -45.69 | 54.8 | HHHHHHHHHHHHHHHHHHHHHHHHCCCCCCC |
| 775 | EEEKRKFLEQYNNEYEDEFYQQKGFGKGDSR | -45.69 | 54.8 | HHHHHHHHHHHHHHHHHHHHHHHCCCCCCCC |
| 885 | EEEKKKMLEQIENEREDEEYQRKGMGKGHQP | -45.67 | 41.9 | HHHHHHHHHHHHHHHHHHHHHHHHCCCCCCC |
| 418 | EEEKRKILEQINQEKEQEEFQRRGMGQGKQP | -45.65 | 32.3 | HHHHHHHHHHHHHHHHHHHHHHHHCCCCCCC |
| 728 | EEEKRKMLDQYNNEYEDEEYQRKGRGKGHQS | -45.64 | 45.2 | HHHHHHHHHHHHHHHHHHHHHHHHCCCCCCC |
| 837 | EEERKKQLEQKNNEEEQERYQQQGFGQGHQS | -45.63 | 38.7 | HHHHHHHHHHHCHHHHHHHHHHHCCCCCCCC |
| 359 | EEERKKQLEQYNNESEQERYQQQGWGQGHNP | -45.63 | 38.7 | HHHHHHHHHHHHHHHHHHHHHHHCCCCCCCC |
| 192 | EEERRKQLEQYNNEAEQQRYQKEGFGQGHQS | -45.63 | 38.7 | HHHHHHHHHHHHHHHHHHHHHHHCCCCCCCC |
| 131 | EEERRKQLEQYNNEAEQQRYQRQGFGQGHQS | -45.63 | 38.7 | HHHHHHHHHHHHHHHHHHHHHHHCCCCCCCC |
| 015 | EEEWKKQMEQYNNEYEQEEYQRKGRGQGHQS | -45.63 | 35.5 | HHHHHHHHHHHHHHHHHHHHHHHHCCCCCCC |
| 026 | EEEKKKLLDQINNEEEQQRYQREGKGQGHSH | -45.62 | 41.9 | HHHHHHHHHHHCHHHHHHHHHHHHCCCCCCC |
| 086 | EEEKRKLLDQINNEKEQEEYQRRGKGQGHSH | -45.62 | 38.7 | HHHHHHHHHHHHHHHHHHHHHHHHHCCCCCC |
| 928 | EEQRRKQLEQINQEQEDKSYQRHGFGKGDSR | -45.60 | 51.6 | HHHHHHHHHHHHHHHHHHHHHHHCCCCCCCC |
| 943 | EEEKKKMLEKINNEEEHKRKQREGFGSGKQN | -45.60 | 38.7 | HHHHHHHHHHHHHHHHHHHHHHHHHCCCCCC |
| 339 | EEQKKKMLEQYNNEYEDQYYQQKGQGKGHSH | -45.59 | 48.4 | HHHHHHHHHHHHHHHHHHHHHHHHCCCCCCC |
| 007 | EEERRKQLEKYNNEAEQRRFQRYGFGQGKSN | -45.56 | 38.7 | HHHHHHHHHHHHHHHHHHHHHHHCCCCCCCC |
| 697 | EEQKKKFLEQFNQEYEDEEYQRKGRGKGDSR | -45.54 | 61.3 | HHHHHHHHHHHHHHHHHHHHHHHCCCCCCCC |
| 351 | EEEKKKFLDQYNNEAEQQRYQQQGFGQGHQS | -45.52 | 48.4 | HHHHHHHHHHHHHHHHHHHHHHHCCCCCCCC |

| | | | | |
|---|---|---|---|---|
| 232 | EEEKKKFLDQYNNEEEQMRYQREGFGQGHQS | -45.52 | 45.2 | HHHHHHHHHHHCCHHHHHHHHHHHHCCCCCC |
| 844 | EEEKKKFLDQYNNESEQQRYQRYGMGQGHQP | -45.52 | 45.2 | HHHHHHHHHHHHCHHHHHHHHHHHCCCCCCC |
| 850 | EEEKRKFLDQYNNEAEQQRYQRYGFGQGHQN | -45.52 | 45.2 | HHHHHHHHHHHHHHHHHHHHHHHCCCCCCCC |
| 118 | EEEKRKFLDQYNNEAEQQRYQRYGFGQGHQS | -45.52 | 45.2 | HHHHHHHHHHHHHHHHHHHHHHHCCCCCCCC |
| 560 | EEEKRKFLDQYNNEHEQQRYQQQGFGQGHQS | -45.52 | 41.9 | HHHHHHHHHHHHHHHHHHHHHHHCCCCCCCC |
| 917 | EEEKRKFLDQYNNEHEQQRYQQQGMGQGHQP | -45.52 | 41.9 | HHHHHHHHHHHHHHHHHHHHHHHHCCCCCCC |
| 517 | EEEKRKFLDQYNNEYEQQYYQQKGFGQGHQS | -45.52 | 41.9 | HHHHHHHHHHHHHHHHHHHHHHHCCCCCCCC |
| 389 | EEEKKKMLEQINNEKEDQEYQRGGFGKGHQP | -45.51 | 45.2 | HHHHHHHHHHHHHHHHHHHHHHCCCCCCCCC |
| 333 | EEEKRKFLDQFNNRYEQEFYQQQGKGQGHSH | -45.50 | 45.2 | HHHHHHHHHHHHHHHHHHHHHHHHCCCCCCC |
| 069 | EEEWKKFIEQYNNEYEDEEYQRKGRGKGNSN | -45.49 | 45.2 | HHHHHHHHHHHHHHHHHHHHHHHCCCCCCCC |
| 898 | EEEKRKFLEQYNNEYEDEEYQRKGRGKGNQN | -45.48 | 45.2 | HHHHHHHHHHHHHHHHHHHHHHHHCCCCCCC |
| 872 | EEEKRKFLEQYNQEAEDKRYQRYGFGKGDSR | -45.47 | 54.8 | HHHHHHHHHHHHHHHHHHHHHHHCCCCCCCC |
| 597 | EEEKRKFLEQKNNREEDERYQQYGFGKGNSN | -45.46 | 41.9 | HHHHHHHHHHHHHHHHHHHHHHHCCCCCCCC |
| 942 | EEEKRKQLEQYNNEYEQEEYQRKGRGQGHQS | -45.43 | 35.5 | HHHHHHHHHHHHHHHHHHHHHHHHCCCCCCC |
| 072 | EEEKRKLLEQLNNEKEDMRYQRYGFGKGDSR | -45.42 | 48.4 | HHHHHHHHHHHHHHHHHHHHHHCCCCCCCCC |
| 497 | EEEKKKMLEKINNEAEDRRYQKEGFGKGHQS | -45.40 | 51.6 | HHHHHHHHHHHHHHHHHHHHHHHHCCCCCCC |
| 273 | EEEKRKMLEKINNEAEDRQYQRNGSGKGHSY | -45.40 | 48.4 | HHHHHHHHHHHHHHHHHHHHHHCCCCCCCCC |
| 184 | EEEKRKMLEKINNEAEDRRYQQQGMGKGHQP | -45.40 | 48.4 | HHHHHHHHHHHHHHHHHHHHHHHHCCCCCCC |
| 223 | EEEWKKILEQLNQEKEQQRYQREGMGQGHQP | -45.40 | 38.7 | HHHHHHHHHHHHHHHHHHHHHHHHHCCCCCC |
| 840 | EEEKRKMLQQINNEEEDSRYQQQGFGKGHQS | -45.39 | 41.9 | HHHHHHHHHHHHHHHHHHHHHHHCCCCCCCC |
| 886 | EEEKRKMLQQINNEKEDEEYQRYGMGKGHQP | -45.39 | 41.9 | HHHHHHHHHHHHHHHHHHHHHHHHHCCCCCC |
| 676 | EEEKRKFLEQYNNEYEHEEYQRKGFGRGDSR | -45.39 | 45.2 | HHHHHHHHHHHHHHHHHHHHHHHCCCCCCCC |
| 324 | EEERRKQLEKYNNEEEQMRYQREGFGQGDSR | -45.39 | 45.2 | HHHHHHHHHHHCCHHHHHHHHHHHCCCCCCC |
| 349 | EEEKRKMLEQINQEAEDERYQQQGFGKGNQN | -45.39 | 45.2 | HHHHHHHHHHHHHHHHHHHHHHHCCCCCCCC |
| 463 | EEEKRKLLEKINQEAEQKQYQRSGKGQGHSS | -45.39 | 45.2 | HHHHHHHHHHHHHHHHHHHHHHHCCCCCCCC |
| 237 | EEEKRKLLEKINQEEEQKRYQRYGFGQGHQS | -45.39 | 38.7 | HHHHHHHHHHHHHHHHHHHHHHHCCCCCCCC |
| 294 | EEERRKQLEKINNEKEDKEYQKQGFGKGDSR | -45.38 | 51.6 | HHHHHHHHHHHHHHHHHHHHHHHCCCCCCCC |
| 202 | EEERRKQLEKLNNEKEDRRYQRYGFGKGHQN | -45.38 | 45.2 | HHHHHHHHHHHHHHHHHHHHHHHCCCCCCCC |
| 888 | EEEKKKMLDKYNNEEEQKRYQRQGFGQGHSS | -45.37 | 45.2 | HHHHHHHHHHHCCHHHHHHHHHHCCCCCCCC |
| 491 | EEEKRKLLDQINNEAEQKRYQRQGFGQGDSR | -45.36 | 48.4 | HHHHHHHHHHHHHHHHHHHHHHHCCCCCCCC |
| 692 | EEEKRKWLEKLNNEKEDRRYQRYGFGKGHQS | -45.34 | 45.2 | HHHHHHHHHHHHHHHHHHHHHHHCCCCCCCC |
| 453 | EEERRKQLEQFNQEYEQKFYQQKGMGQGHQP | -45.31 | 41.9 | HHHHHHHHHHHHHHHHHHHHHHHHCCCCCCC |
| 522 | EEEKRKLLEKINNEKEHKEYQKQGFGQGDSR | -45.30 | 45.2 | HHHHHHHHHHHHHHHHHHHHHHHCCCCCCCC |
| 925 | EEEKKKMLEQINNEAEDKRYQREGFGKGDSR | -45.29 | 54.8 | HHHHHHHHHHHHHHHHHHHHHHHHCCCCCCC |
| 052 | EEEKRKMLEQINNEEEDQRYQQQGFGKGDSR | -45.29 | 48.4 | HHHHHHHHHHHHHHHHHHHHHHHCCCCCCCC |
| 599 | EEEQRKMLEQINNEKEDEEYQRYGFGKGDSR | -45.29 | 48.4 | HHHHHHHHHHHHHHHHHHHHHHHCCCCCCCC |
| 095 | EQEKRKMLEQINNEKEDEEYQRYGFGKGDSR | -45.29 | 45.2 | HHHHHHHHHHHHHHHHHHHHHHHCCCCCCCC |
| 003 | EEEKRKLLEKINNEEEQKRYQQYGFGQGHQS | -45.28 | 38.7 | HHHHHHHHHHHCHHHHHHHHHHHCCCCCCCC |

| | | | | |
|---|---|---|---|---|
| 741 | EEEKRKYLEQYNNEYEDEQYQQQGFGKGDSR | -45.28 | 48.4 | HHHHHHHHHHHHHHHHHHHHHHHCCCCCCCC |
| 004 | EEERRKQLEQINNEEEDERYQQQGFGKGDSR | -45.27 | 48.4 | HHHHHHHHHHHHHHHHHHHHHHHCCCCCCCC |
| 632 | EEEKKKFLEQYEDEYRDQEYQRNGRGKGHSY | -45.24 | 41.9 | HHHHHHHHHHHHHHHHHHHHHHCCCCCCCCC |
| 261 | EEEKRKFLEQYEDEYRDEEYQRKGFGKGHQS | -45.24 | 38.7 | HHHHHHHHHHHHHHHHHHHHHHHCCCCCCCC |
| 848 | EEEKRKFLEQYNNESEHQRKQRYGFGSGKQN | -45.22 | 35.5 | HHHHHHHHHHHHHHHHHHHHHHHCCCCCCCC |
| 077 | EEERRKQLDQINNEEEQRRYQREGFGQGHQS | -45.22 | 38.7 | HHHHHHHHHHHCHHHHHHHHHHHHHCCCCCC |
| 624 | EEEKRKFLEQYNNEEEDMRYQREGFGKGDSR | -45.18 | 51.6 | HHHHHHHHHHHCHHHHHHHHHHHHCCCCCCC |
| 479 | EEEWKKFIEQYNNEYEDEEYQRKGFGKGDSR | -45.18 | 51.6 | HHHHHHHHHHHHHHHHHHHHHHHCCCCCCCC |
| 210 | EQEKRKFLEQYNNRHEDERYQQQGFGKGDSR | -45.15 | 45.2 | HHHHHHHHHHHHHHHHHHHHHHHCCCCCCCC |
| 849 | EEEKKKFLEAYNNRYEDEEYQRNGRGKGHSY | -45.15 | 45.2 | HHHHHHHHHHHHHHHHHHHHHHCCCCCCCCC |
| 208 | EEEKRKFLEQYNNRYEDEEYQRTGRGKGDSR | -45.14 | 48.4 | HHHHHHHHHHHHHHHHHHHHHHCCCCCCCCC |
| 718 | EEEKKKMLDQINNEAEQKRYQQYGFGQGHQS | -45.13 | 45.2 | HHHHHHHHHHHHHHHHHHHHHHHCCCCCCCC |
| 097 | EEEKKKMLDQINNEEEQKRYQRYGFGQGHSS | -45.13 | 41.9 | HHHHHHHHHHHCHHHHHHHHHHHCCCCCCCC |
| 040 | EEEKRKMLDQYNNEEEQMRYQREGFGQGHQS | -45.13 | 38.7 | HHHHHHHHHHHCCHHHHHHHHHHHHCCCCCC |
| 518 | EEEKRKLLEKINNEEEDRRYQQQGFGKGDSR | -45.11 | 51.6 | HHHHHHHHHHHHHHHHHHHHHHHCCCCCCCC |
| 977 | EEERRKQLDQINNEKEQEEYQRYGFGQGHSS | -45.11 | 38.7 | HHHHHHHHHHHHHHHHHHHHHHHCCCCCCCC |
| 082 | EEEKKKILEQINNEAEQKRYQREGMGQGHQP | -45.11 | 41.9 | HHHHHHHHHHHHHHHHHHHHHHHHHCCCCCC |
| 827 | EEEKRKILEQINNEKEQEEYQRRGMGQGHQP | -45.11 | 35.5 | HHHHHHHHHHHHHHHHHHHHHHHHHCCCCCC |
| 245 | EEEKRKILEQINNEKEQQEYQRGGFGQGHQP | -45.11 | 35.5 | HHHHHHHHHHHHHHHHHHHHHHCCCCCCCCC |
| 983 | EEEKRKILEQLNNEKEQQRYQRYGFGQGHQS | -45.11 | 35.5 | HHHHHHHHHHHHHHHHHHHHHHHCCCCCCCC |
| 961 | EQEERKIQEQFNNEYEQEFYQQQGFGQGHQS | -45.11 | 35.5 | HHHHHHHHHHHHHHHHHHHHHHHCCCCCCCC |
| 778 | EEEKRKFLDDYNNRSEDQRYQRYGFGKGHQS | -45.10 | 45.2 | HHHHHHHHHHHHCCCHHHHHHHHCCCCCCCC |
| 950 | EEEKKKMLEQINNEAEDKRYQRYGFGKGNQN | -45.09 | 48.4 | HHHHHHHHHHHHHHHHHHHHHHHCCCCCCCC |
| 043 | EEEKRKMLEQINNEAEDERYQRKGFGKGNQN | -45.09 | 45.2 | HHHHHHHHHHHHHHHHHHHHHHHCCCCCCCC |
| 829 | EEEKRKMLEQYNNEYEDEEYQRKGRGKGNQN | -45.09 | 41.9 | HHHHHHHHHHHHHHHHHHHHHHHHCCCCCCC |
| 239 | EEEERKLQEKLNNEKEQKRYQRYGFGQGHQN | -45.09 | 35.5 | HHHHHHHHHHHHHHHHHHHHHHHCCCCCCCC |
| 715 | EEEERKLQEKLNNEKEQKRYQRYGQGQGHQS | -45.09 | 35.5 | HHHHHHHHHHHHHHHHHHHHHHHHHCCCCCC |
| 988 | EEEKKKLLEKINNEEEQKRYQRYGFGQGHSS | -45.09 | 41.9 | HHHHHHHHHHHCHHHHHHHHHHHCCCCCCCC |
| 411 | EEEKRKLLEKINNEAEQKRYQRQGFGQGHQS | -45.09 | 41.9 | HHHHHHHHHHHHHHHHHHHHHHHCCCCCCCC |
| 947 | EEEKRKLLEKINNEEEQKRYQREGFGQGHQS | -45.09 | 38.7 | HHHHHHHHHHHCHHHHHHHHHHHHCCCCCCC |
| 915 | EEEKRKLLEKLNNEKEQKRYQKEGWGQGHNP | -45.09 | 38.7 | HHHHHHHHHHHHHHHHHHHHHHHHCCCCCCC |
| 583 | EQEKRKLLEKLNNEKEQKRYQREGMGQGHQP | -45.09 | 35.5 | HHHHHHHHHHHHHHHHHHHHHHHHHCCCCCC |
| 064 | EEEKKKFLDQFNNEFEQQYYQQQGFGQGHSN | -45.05 | 48.4 | HHHHHHHHHHHHHHHHHHHHHHHCCCCCCCC |
| 713 | EEEKRKFLDQYNNEYEQEEYQRTGRGQGHQF | -45.05 | 41.9 | HHHHHHHHHHHHHHHHHHHHHHHHCCCCCCC |
| 833 | EEERRKQLEQYNNEAEQERYQQQGMGQGHQP | -45.02 | 38.7 | HHHHHHHHHHHHHHHHHHHHHHHHCCCCCCC |
| 067 | EQEKRHFLEKYNNEAEDKQYQQRGKGKGHSA | -44.95 | 48.4 | HHHHHHHHHHHHHHHHHHHHHHHHCCCCCCC |
| 511 | EEEKRKWLDQLNNEKEQEEYQRYGFGQGHSS | -44.92 | 38.7 | HHHHHHHHHHHHHHHHHHHHHHHCCCCCCCC |
| 385 | EEEKRKMLDQINNEKEQEEYQRRGMGQGHQP | -44.90 | 38.7 | HHHHHHHHHHHHHHHHHHHHHHHHHCCCCCC |

| | | | | |
|---|---|---|---|---|
| 756 | EEERRKQLDKYNQEYEQEEYQRKGRGQGDSR | -44.89 | 48.4 | HHHHHHHHHHHHHHHHHHHHHHHHCCCCCCCC |
| 941 | EEEKRKLLEKINNEEEQKRYQREGFGQGHSS | -44.86 | 38.7 | HHHHHHHHHHHCHHHHHHHHHHHHHCCCCCCC |
| 843 | EEEKRKFLEKYNNEYEDREYQRTGRGKGNQN | -44.86 | 48.4 | HHHHHHHHHHHHHHHHHHHHHHHCCCCCCCCC |
| 598 | EEEKRKFLEKYNNEYEQKEFQKEGFGQGKQN | -44.85 | 38.7 | HHHHHHHHHHHHHHHHHHHHHHHHHCCCCCCC |
| 442 | EEEKKKMLEQINNEKEHEEKQRYGFGSGKQN | -44.83 | 35.5 | HHHHHHHHHHHHHHHHHHHHHHHHCCCCCCCC |
| 111 | EEEKRKMLEQINNEEEHKRKQREGFGSGKQN | -44.83 | 32.3 | HHHHHHHHHHHHHHHHHHHHHHHHHHCCCCCC |
| 687 | EEEKKKFLEKFNQEKEQQRYQQQGFGQGHQS | -44.82 | 48.4 | HHHHHHHHHHHHHHHHHHHHHHHHCCCCCCCC |
| 979 | EEEKRKFLEKYNQEYEQKEYQKEGRGQGHQS | -44.82 | 41.9 | HHHHHHHHHHHHHHHHHHHHHHHHHCCCCCCC |
| 431 | EQEKRKLLEQIENEREQQEYQQKGFGQGTQN | -44.82 | 29.0 | HHHHHHHHHHHHHHHHHHHHHHHHHHCCCCCC |
| 516 | EEEKKKLLEQIEQRREQEEYQQKGQGQGHQS | -44.79 | 32.3 | HHHHHHHHHHHHHHHHHHHHHHHHHHCCCCCC |
| 347 | EEEKKKMLEQINNEEEDQRYQQYGFGKGDSR | -44.78 | 51.6 | HHHHHHHHHHHHHHHHHHHHHHHHCCCCCCCC |
| 256 | EEEKRKMLEQINNEAEDQRYQRYGFGKGDSR | -44.78 | 51.6 | HHHHHHHHHHHHHHHHHHHHHHHHCCCCCCCC |
| 722 | EEEKRKFLDDYNNRSEDERYQRYGMGKGHQP | -44.78 | 45.2 | HHHHHHHHHHHHCCCHHHHHHHHHHCCCCCCC |
| 645 | EEEKKKMLQKINNEAEDRRYQREGKGKGHSH | -44.76 | 51.6 | HHHHHHHHHHHHHHHHHHHHHHHHCCCCCCCC |
| 314 | EEEKKKMLEQINNEAEHKRKQREGQGSGKQS | -44.76 | 38.7 | HHHHHHHHHHHHHHHHHHHHHHHHCCCCCCCC |
| 910 | EEERRKQLEQINNEAEDKRYQRYGFGKGDSR | -44.76 | 51.6 | HHHHHHHHHHHHHHHHHHHHHHHHCCCCCCCC |
| 155 | EEERRKQLQKINNEAEDRRYQRYGFGKGHQN | -44.74 | 48.4 | HHHHHHHHHHHHHHHHHHHHHHHHCCCCCCCC |
| 531 | EEQKRKFLEQFNNEYEDQFYQQKGFGKGDSR | -44.74 | 61.3 | HHHHHHHHHHHHHHHHHHHHHHHHCCCCCCCC |
| 610 | EEERRKQLEQYNNEEEQMRYQREGFGQGHQS | -44.73 | 35.5 | HHHHHHHHHHHCCHHHHHHHHHHHHHCCCCCC |
| 313 | EEQLKKILEQINNEAEQKRYQREGKGQGHSH | -44.67 | 45.2 | HHHHHHHHHHHHHHHHHHHHHHHHHCCCCCCC |
| 417 | EEEKKKLLEQINNEKEQEEFQRYGFGQGKQN | -44.65 | 35.5 | HHHHHHHHHHHHHHHHHHHHHHHHCCCCCCCC |
| 796 | EEELKKILEQLNNEKEQKRYQRYGMGQGHQP | -44.63 | 38.7 | HHHHHHHHHHHHHHHHHHHHHHHHHCCCCCCC |
| 033 | EEEKKKLLEQINQEQEQEEYQRKGRGQGHQP | -44.62 | 38.7 | HHHHHHHHHHHHHHHHHHHHHHHHHCCCCCCC |
| 088 | EEEKRKLLEQINQEEEQQRYQQQGMGQGHQP | -44.62 | 35.5 | HHHHHHHHHHHHHHHHHHHHHHHHHCCCCCCC |
| 797 | EEEKKKLLEKINNEAEQKRYQREGKGQGHSS | -44.62 | 45.2 | HHHHHHHHHHHHHHHHHHHHHHHHCCCCCCCC |
| 383 | EEEKRKLLEKINNEKEQQRYQQQGMGQGHQP | -44.62 | 38.7 | HHHHHHHHHHHHHHHHHHHHHHHHHCCCCCCC |
| 103 | EEEWRKLMEKYNNEEEQKRYQREGMGQGHQP | -44.62 | 35.5 | HHHHHHHHHHHCCHHHHHHHHHHHHCCCCCCC |
| 673 | EEEKRKLLEQINNEAEQKRFQREGKGQGKSH | -44.58 | 35.5 | HHHHHHHHHHHHHHHHHHHHHHHHHCCCCCCC |
| 309 | EEEQRRLYEQLNNEKEDQRYQRYGFGKGHSS | -44.58 | 38.7 | HHHHHHHHHHHHHHHHHHHHHHHHCCCCCCCC |
| 754 | EEEKRKMLDQINNEKEQEEYQRYGFGQGHQS | -44.54 | 38.7 | HHHHHHHHHHHHHHHHHHHHHHHHCCCCCCCC |
| 090 | EEEARKQQEQYNNESEQKRYQQYGFGQGHSN | -44.54 | 35.5 | HHHHHHHHHHHHHHHHHHHHHHHHCCCCCCCC |
| 694 | EEEARKFQEKYSNESEQKRYQQQGFGQGHQS | -44.53 | 38.7 | HHHHHHHHHHHHHHHHHHHHHHHHCCCCCCCC |
| 124 | EEEEKKFQEKYNNESEQKRYQRYGFGQGHQS | -44.53 | 41.9 | CHHHHHHHHHHHHHHHHHHHHHHHCCCCCCCC |
| 820 | EEEERKFQEKYNNEAEQKRYQRYGMGQGHQP | -44.53 | 41.9 | CHHHHHHHHHHHHHHHHHHHHHHHHCCCCCCC |
| 830 | EEEKKKFLEKFNNEYEQEYYQQNGSGQGHQY | -44.53 | 48.4 | HHHHHHHHHHHHHHHHHHHHHHHHCCCCCCCC |
| 604 | EEEKRKFLEKFNNEYEQKYYQQKGFGQGHQS | -44.53 | 45.2 | HHHHHHHHHHHHHHHHHHHHHHHHCCCCCCCC |
| 680 | EEEKRKFLEKFNNEYEQKYYQQKGQGQGHSH | -44.53 | 45.2 | HHHHHHHHHHHHHHHHHHHHHHHHHCCCCCCC |
| 592 | EEEKRKFLEKMKNEKEQEEYQRYGFGQGHSN | -44.53 | 38.7 | HHHHHHHHHHHHHHHHHHHHHHHHCCCCCCCC |

| | | | | |
|---|---|---|---|---|
| 415 | EEEKRKFLEKYNNEAEQKRYQRYGFGQGHQS | -44.53 | 45.2 | HHHHHHHHHHHHHHHHHHHHHHHCCCCCCCC |
| 567 | EEEKKKLLEQIENEREQQEYQRQGFGQGHQN | -44.49 | 35.5 | HHHHHHHHHHHHHHHHHHHHHHHCCCCCCCC |
| 927 | EEEKRKMLQQFNNEFEDQFYQQKGFGKGNSN | -44.46 | 48.4 | HHHHHHHHHHHHHHHHHHHHHHHCCCCCCCC |
| 677 | EEEKRKLLQKINNEEEQKRYQREGQGQGHQS | -44.45 | 38.7 | HHHHHHHHHHHCHHHHHHHHHHHHCCCCCCC |
| 306 | EEEKRKMLQQINQEEEDQRYQRYGFGKGDSR | -44.44 | 48.4 | HHHHHHHHHHHHHHHHHHHHHHHCCCCCCCC |
| 435 | EEEKKKMLEKINQEAEQKRYQREGMGQGHQP | -44.43 | 45.2 | HHHHHHHHHHHHHHHHHHHHHHHHHCCCCCC |
| 606 | EEEKKKMLEKINQEEEQKRYQREGMGQGHQP | -44.43 | 41.9 | HHHHHHHHHHHHHHHHHHHHHHHHCCCCCCC |
| 287 | EEERRKQLEKLNQEKEQKRYQQYGFGQGHQS | -44.40 | 38.7 | HHHHHHHHHHHHHHHHHHHHHHHCCCCCCCC |
| 630 | EEEKRKMLDQYNNESEQQRYQQYGFGQGDSR | -44.40 | 45.2 | HHHHHHHHHHHHHHHHHHHHHHHCCCCCCCC |
| 475 | EEEKKKWLEKLNNEKEQKRYQRYGMGQGHQP | -44.39 | 41.9 | HHHHHHHHHHHHHHHHHHHHHHHHCCCCCCC |
| 275 | EEEKKKWLEKLNNEYEQQEYQRNGRGQGHSY | -44.39 | 41.9 | HHHHHHHHHHHHHHHHHHHHHHHCCCCCCCC |
| 299 | EEEKRKWLEKLNNEKEQERYQQQGMGQGHQP | -44.39 | 38.7 | HHHHHHHHHHHHHHHHHHHHHHHHCCCCCCC |
| 992 | EEEWRKWIEKLNNEKEQEEYQRKGRGQGHQS | -44.39 | 35.5 | HHHHHHHHHHHHHHHHHHHHHHHHCCCCCCC |
| 047 | EEEWRKWIEKLNNEKEQKRYQREGMGQGHQP | -44.39 | 35.5 | HHHHHHHHHHHHHHHHHHHHHHHHHCCCCCC |
| 563 | EQERRKQLEQYNNEAEQQRYQRYGFGQGDSR | -44.39 | 41.9 | HHHHHHHHHHHHHHHHHHHHHHHCCCCCCCC |
| 911 | EEEKRKFLQQYNEEWEQMRYQREGFGQGDSR | -44.35 | 45.2 | HHHHHHHHHHHHHHHHHHHHHHHHCCCCCCC |
| 430 | EEEKKKLLEQINNEAEQKRYQKEGFGQGHQS | -44.32 | 41.9 | HHHHHHHHHHHHHHHHHHHHHHHHCCCCCCC |
| 113 | EEEKKKLLEQINNEAEQKRYQRYGFGQGHQS | -44.32 | 41.9 | HHHHHHHHHHHHHHHHHHHHHHHCCCCCCCC |
| 889 | EEEKKKLLEQINNEEEQKRYQRYGMGQGHQP | -44.32 | 38.7 | HHHHHHHHHHHCHHHHHHHHHHHHCCCCCCC |
| 039 | EEEKKKLLEQINNEKEQEEYQRRGMGQGHQP | -44.32 | 38.7 | HHHHHHHHHHHHHHHHHHHHHHHHHCCCCCC |
| 450 | EEEKKKLLEQINNEQEQERYQQQGWGQGHTP | -44.32 | 38.7 | HHHHHHHHHHHHHHHHHHHHHHHCCCCCCCC |
| 999 | EEEKRKLLEQINNEAEQKRYQREGWGQGHNP | -44.32 | 38.7 | HHHHHHHHHHHHHHHHHHHHHHHHCCCCCCC |
| 374 | EEEKRKLLEQINNEEEQKRYQREGMGQGHQP | -44.32 | 35.5 | HHHHHHHHHHHCHHHHHHHHHHHHCCCCCCC |
| 140 | EEEKRKLLEQINNEEEQKRYQRQGFGQGHQS | -44.32 | 35.5 | HHHHHHHHHHHCHHHHHHHHHHHCCCCCCCC |
| 641 | EEEKRKLLEQINNEEEQQRYQQQGFGQGHQN | -44.32 | 35.5 | HHHHHHHHHHHCHHHHHHHHHHHCCCCCCCC |
| 588 | EEEKRKLLEQINNEKEQEEYQRRGMGQGHQP | -44.32 | 35.5 | HHHHHHHHHHHHHHHHHHHHHHHHHCCCCCC |
| 494 | EEEKRKLLEQINNEKEQEEYQRYGFGQGHQS | -44.32 | 35.5 | HHHHHHHHHHHHHHHHHHHHHHHCCCCCCCC |
| 269 | EEEKRKLLEQINNEKEQQEYQKQGFGQGHQS | -44.32 | 35.5 | HHHHHHHHHHHHHHHHHHHHHHHCCCCCCCC |
| 446 | EEEKRKLLEQINNEQEQEEYQRKGRGQGHQP | -44.32 | 35.5 | HHHHHHHHHHHHHHHHHHHHHHHHCCCCCCC |
| 918 | EEEKRKLLEQINNEQEQKSYQQQGFGQGHQS | -44.32 | 35.5 | HHHHHHHHHHHHHHHHHHHHHHHCCCCCCCC |
| 209 | EEEKRKLLEQINNEQEQKSYQREGMGQGHQP | -44.32 | 35.5 | HHHHHHHHHHHHHHHHHHHHHHHHCCCCCCC |
| 635 | EEEKRKLLEQINNEQEQQYYQQYGFGQGHSS | -44.32 | 35.5 | HHHHHHHHHHHHHHHHHHHHHHHCCCCCCCC |
| 308 | EEEKRKLLEQLNNEKEQQRYQRYGFGQGHQS | -44.32 | 35.5 | HHHHHHHHHHHHHHHHHHHHHHHCCCCCCCC |
| 633 | EEELKKLLEQINNEEEQKRYQRYGFGQGHQS | -44.32 | 38.7 | HHHHHHHHHHHCHHHHHHHHHHHCCCCCCCC |
| 216 | EQEKRKLLEQLNNEKEQQRYQRQGFGQGHQS | -44.32 | 32.3 | HHHHHHHHHHHHHHHHHHHHHHHCCCCCCCC |
| 541 | EEERRKQLEKINNEKEQQRYQQQGFGQGHQS | -44.30 | 38.7 | HHHHHHHHHHHHHHHHHHHHHHHCCCCCCCC |
| 994 | EEEKRKFLEKYNNESEQKRYQQQGFGQGHSS | -44.30 | 41.9 | HHHHHHHHHHHHHHHHHHHHHHHCCCCCCCC |
| 558 | EEEKRKFLDQYNNEAEQQRYQRYGFGQGDSR | -44.29 | 51.6 | HHHHHHHHHHHHHHHHHHHHHHHCCCCCCCC |

| | | | | |
|---|---|---|---|---|
| 998 | EEEWKKFIEQYNNDYEDEEYQRKGRGKGDSR | -44.21 | 48.4 | HHHHHHHHHHHHHHCHHHHHHHHCCCCCCCC |
| 828 | EEEKRKMLEKINNEAEQKRYQRYGFGQGHQS | -44.20 | 41.9 | HHHHHHHHHHHHHHHHHHHHHHHCCCCCCCC |
| 784 | EQEKRHLLEKINNEAEDKRYQREGFGKGDSR | -44.19 | 51.6 | HHHHHHHHHHHHHHHHHHHHHHHHCCCCCCC |
| 176 | EQEKRHFLEQYNNESEDQRYQRYGFGKGHQS | -44.18 | 41.9 | HHHHHHHHHHHHCCHHHHHHHHHCCCCCCCC |
| 171 | EEERRKQLDQINNEAEQQRYQRYGFGQGNSN | -44.17 | 41.9 | HHHHHHHHHHHHHHHHHHHHHHHCCCCCCCC |
| 053 | EEEWRKLMDAVNNREEQERYQQQGWGQGHTP | -44.15 | 32.3 | HHHHHHHHHHHHHHHHHHHHHHHCCCCCCCC |
| 244 | EEEKRKLLEQINQEEEQERYQQQGMGQGHQP | -44.15 | 35.5 | HHHHHHHHHHHHHHHHHHHHHHHHCCCCCCC |
| 153 | EEEKKKMLEKINNEEEQKRYQREGRGQGHQP | -44.13 | 41.9 | HHHHHHHHHHHCHHHHHHHHHHHHCCCCCCC |
| 612 | EEEKKKMLEKINNEEEQKRYQRYGFGQGHSN | -44.13 | 41.9 | HHHHHHHHHHHCHHHHHHHHHHHCCCCCCCC |
| 523 | EEEKRKMLEKFNNEEEQKRYQREGFGQGHQS | -44.13 | 41.9 | HHHHHHHHHHHCCHHHHHHHHHHHCCCCCCC |
| 627 | EEEKRKMLEKFNNEFEQKYYQQKGQGQGHSH | -44.13 | 41.9 | HHHHHHHHHHHHHHHHHHHHHHHHCCCCCCC |
| 178 | EEEKRKMLEKINNEEEQKRYQRYGMGQGHQP | -44.13 | 38.7 | HHHHHHHHHHHCHHHHHHHHHHHHCCCCCCC |
| 335 | EEEKRKMLEKYNNEYEQEEYQRKGRGQGHQP | -44.13 | 38.7 | HHHHHHHHHHHHHHHHHHHHHHHHCCCCCCC |
| 936 | EQEKRKMLEKINNEKEQKEYQRAGFGQGHQS | -44.13 | 35.5 | HHHHHHHHHHHHHHHHHHHHHHHCCCCCCCC |
| 276 | EEERKKQLEKINNEEEQKSYQREGMGQGHQP | -44.11 | 41.9 | HHHHHHHHHHHHHHHHHHHHHHHHCCCCCCC |
| 853 | EEERRKQLEKINNEEEQKRYQRYGFGQGHQS | -44.11 | 38.7 | HHHHHHHHHHHCHHHHHHHHHHHCCCCCCCC |
| 466 | EEERRKQLEKYNNEYEQKEYQRTGRGQGHSF | -44.11 | 38.7 | HHHHHHHHHHHHHHHHHHHHHHHCCCCCCCC |
| 022 | EEEKRKFLEQYNNEYEQEEFQRKGRGQGKQN | -44.08 | 35.5 | HHHHHHHHHHHHHHHHHHHHHHHHHCCCCCC |
| 867 | EEERRKQLDAINNRKEQEEYQRYGFGQGHQS | -44.06 | 35.5 | HHHHHHHHHHHHHHHHHHHHHHHCCCCCCCC |
| 605 | EEEKKKFLEQFNQEYEQQYYQQQGFGQGHSS | -44.06 | 45.2 | HHHHHHHHHHHHHHHHHHHHHHHCCCCCCCC |
| 787 | EEEKRKFLEQYNQEYEQQEYQRNGRGQGHSY | -44.06 | 38.7 | HHHHHHHHHHHHHHHHHHHHHHHCCCCCCCC |
| 191 | EEEQRKFEEQRNQEEEQQRYQQYGFGQGHQS | -44.06 | 35.5 | HHHHHHHHHHHHHHHHHHHHHHHCCCCCCCC |
| 871 | EEEKRKFLEKFNNEFEQKYYQQKGFGQGHSS | -44.05 | 45.2 | HHHHHHHHHHHHHHHHHHHHHHHCCCCCCCC |
| 057 | EEEKRKFLEKYNNESEQKRYQRYGFGQGHQS | -44.05 | 41.9 | HHHHHHHHHHHCHHHHHHHHHHHCCCCCCCC |
| 509 | EEEKRKFLEKYNNEYEQKEYQRTGRGQGHQS | -44.05 | 41.9 | HHHHHHHHHHHHHHHHHHHHHHHCCCCCCCC |
| 432 | EEEKKKLLEDINQRKEDQEYQKQGFGKGHSS | -44.05 | 41.9 | HHHHHHHHHHHHHHHHHHHHHHHCCCCCCCC |
| 211 | EEEKRKFLEQYNQRAEQERYQREGKGQGHSS | -44.03 | 38.7 | HHHHHHHHHHHHHHHHHHHHHHHCCCCCCCC |
| 550 | EEEKRKILEKINNEEEQRRYQQYGFGQGHQN | -44.02 | 38.7 | HHHHHHHHHHHCHHHHHHHHHHHCCCCCCCC |
| 213 | EEEKRKMLEKYNQEEEQKKYQREGKGQGHSH | -43.95 | 38.7 | HHHHHHHHHHHHHHHHHHHHHHHHCCCCCCC |
| 817 | EEEKRKWLEKLNNEKEQKRYQKEGFGQGHQS | -43.92 | 38.7 | HHHHHHHHHHHHHHHHHHHHHHHHCCCCCCC |
| 795 | EEEKRKLLEDIENRREDQEYQQKGRGKGHQP | -43.92 | 35.5 | HHHHHHHHHHHHHHHHHHHHHHHHCCCCCCC |
| 019 | EEEQRKMLEKINNEKEQEEYQRRGMGQGHQP | -43.90 | 38.7 | HHHHHHHHHHHHHHHHHHHHHHHHHCCCCCC |
| 966 | EEEKRKFLQKKNNEEEQKRYQQYGFGQGHQS | -43.89 | 41.9 | HHHHHHHHHHHCHHHHHHHHHHHCCCCCCCC |
| 224 | EQEKRKMLEQFNNEYEDEEYQRQGRGKGHQS | -43.88 | 41.9 | HHHHHHHHHHHHHHHHHHHHHHHHCCCCCCC |
| 117 | EEEKRKMLDQINNEKEQKEYQRAGFGQGDSR | -43.87 | 45.2 | HHHHHHHHHHHHHHHHHHHHHHHCCCCCCCC |
| 570 | EEEKRKYLEQYNNEAEQKRYQREGMGQGHQP | -43.86 | 38.7 | HHHHHHHHHHHHHHHHHHHHHHHHHCCCCCC |
| 855 | EQERRKQLEQINNEAEDKRYQREGMGKGHQP | -43.85 | 41.9 | HHHHHHHHHHHHHHHHHHHHHHHHHCCCCCC |
| 249 | EEEKRKLLEKINNEQEQKRYQREGKGQGDSR | -43.85 | 45.2 | HHHHHHHHHHHHHHHHHHHHHHHCCCCCCCC |

| | | | | |
|---|---|---|---|---|
| 270 | EEEKKKLLEQINNEQEQEEYQRKGRGQGHQP | -43.85 | 38.7 | HHHHHHHHHHHHHHHHHHHHHHHHCCCCCCC |
| 268 | EEEKRKLLEQINNEAEQKRYQREGQGQGHQS | -43.85 | 38.7 | HHHHHHHHHHHHHHHHHHHHHHHHCCCCCCC |
| 110 | EEEKRKLLEQINNEEEQKRYQREGKGQGHSH | -43.85 | 35.5 | HHHHHHHHHHHCHHHHHHHHHHHHCCCCCCC |
| 051 | EEEKRKLLEQLNNEKEQQRYQQYGFGQGHQS | -43.85 | 35.5 | HHHHHHHHHHHHHHHHHHHHHHHCCCCCCCC |
| 128 | EEEKKKLLEQINNEKEQKEYQKQGWGQGHNP | -43.83 | 38.7 | HHHHHHHHHHHHHHHHHHHHHHHCCCCCCCC |
| 220 | EEEKKKFLEKYNNEYEQKEYQRTGRGQGDSR | -43.80 | 51.6 | HHHHHHHHHHHHHHHHHHHHHHHCCCCCCCC |
| 410 | EEEKRKFLEKYNNEYEQKEYQRTGRGQGDSR | -43.80 | 48.4 | HHHHHHHHHHHHHHHHHHHHHHHCCCCCCCC |
| 812 | EQEKRHMLEQINNEKEDEEYQRRGMGKGHQP | -43.79 | 38.7 | HHHHHHHHHHHHHHHHHHHHHHHHHCCCCCC |
| 471 | EEEKKKFLEQKNNEEEQQRYQRYGFGQGHSS | -43.76 | 41.9 | HHHHHHHHHHHCCHHHHHHHHHHCCCCCCCC |
| 449 | EEEKKKFLEQYNNEYEQMFYQQKGFGQGHSS | -43.76 | 45.2 | HHHHHHHHHHHHHHHHHHHHHHHHCCCCCCC |
| 590 | EEEKKKFLEQYNNEYEQQRYQQEGMGQGHQP | -43.76 | 41.9 | HHHHHHHHHHHHHHHHHHHHHHHHHCCCCCC |
| 900 | EEEKKKFLEQYNNEYEQQYYQQNGSGQGHSY | -43.76 | 41.9 | HHHHHHHHHHHHHHHHHHHHHHHCCCCCCCC |
| 568 | EEEKKKFLEQYSNEYEQEEYQRNGRGQGHSY | -43.76 | 38.7 | HHHHHHHHHHHHHHHHHHHHHHHCCCCCCCC |
| 127 | EEEKRKFLEQFNNEYEQMSYQREGKGQGHSH | -43.76 | 41.9 | HHHHHHHHHHHHHHHHHHHHHHHCCCCCCCC |
| 873 | EEEKRKFLEQKNNEEEQQRYQQQGMGQGHQP | -43.76 | 38.7 | HHHHHHHHHHHCCHHHHHHHHHHHCCCCCCC |
| 226 | EEEKRKFLEQMKNEKEQEEYQRRGMGQGHQP | -43.76 | 35.5 | HHHHHHHHHHHHHHHHHHHHHHHHHCCCCCC |
| 228 | EEEKRKFLEQYNNEAEQQQYQRNGSGQGHSY | -43.76 | 41.9 | HHHHHHHHHHHHHHHHHHHHHHHCCCCCCCC |
| 659 | EEEKRKFLEQYNNESEQKRYQRYGFGQGHSS | -43.76 | 38.7 | HHHHHHHHHHHHHHHHHHHHHHHCCCCCCCC |
| 809 | EEEKRKFLEQYNNESEQQRYQQQGQGQGHQS | -43.76 | 38.7 | HHHHHHHHHHHHHHHHHHHHHHHHHCCCCCC |
| 841 | EEEKRKFLEQYNNESEQQRYQRYGFGQGHQS | -43.76 | 38.7 | HHHHHHHHHHHHCHHHHHHHHHHCCCCCCCC |
| 438 | EEEKRKFLEQYNNEWEQEEYQRKGRGQGHQS | -43.76 | 38.7 | HHHHHHHHHHHHHHHHHHHHHHHHCCCCCCC |
| 717 | EEEKRKFLEQYNNEYEQEEYQRNGRGQGHSY | -43.76 | 38.7 | HHHHHHHHHHHHHHHHHHHHHHHCCCCCCCC |
| 661 | EEEKRKFLEQYNNEYEQQEYQRTGRGQGHSF | -43.76 | 38.7 | HHHHHHHHHHHHHHHHHHHHHHHHCCCCCCC |
| 726 | EEEKRKFLEQYSNEAEQQRYQKEGWGQGHNP | -43.76 | 38.7 | HHHHHHHHHHHHHHHHHHHHHHHHCCCCCCC |
| 384 | EEEQKKFEEQRNNEEEQERYQQQGFGQGHQS | -43.76 | 38.7 | CHHHHHHHHHHHHHHHHHHHHHHCCCCCCCC |
| 011 | EQEERKFQEQYNNEEEQQKYQQEGKGQGHSH | -43.76 | 32.3 | CHHHHHHHHHHCHHHHHHHHHHHHCCCCCCC |
| 271 | EQEKRKFLEQYNNEEEQMRYQREGKGQGHSH | -43.76 | 35.5 | HHHHHHHHHHHCHHHHHHHHHHHHCCCCCCC |
| 935 | EQEKRKFLEQYNNEYEQEEYQRTGRGQGHSF | -43.76 | 35.5 | HHHHHHHHHHHHHHHHHHHHHHHHCCCCCCC |
| 959 | EEQKRKMLDQINQEKEQEEYQRYGFGQGDSR | -43.76 | 48.4 | HHHHHHHHHHHHHHHHHHHHHHHCCCCCCCC |
| 613 | EEEKRKLLEDINNRAEDQRYQRYGFGKGHSS | -43.76 | 41.9 | HHHHHHHHHHHHHHHHHHHHHHHCCCCCCCC |
| 227 | EEEKRKFLEQYNNRYEQEYYQQKGFGQGHQS | -43.74 | 35.5 | HHHHHHHHHHHHHHHHHHHHHHHCCCCCCCC |
| 608 | EEEKRKLLEKINNEAEQRRYQREGKGQGHSH | -43.70 | 41.9 | HHHHHHHHHHHHHHHHHHHHHHHHCCCCCCC |
| 678 | EEEKRKMLEQINNEAEQKRFQQQGFGQGKQN | -43.69 | 35.5 | HHHHHHHHHHHHHHHHHHHHHHHCCCCCCCC |
| 575 | EEEKKKMLEKINNEAEDKRYQRYGFGKGHQS | -43.68 | 51.6 | HHHHHHHHHHHHHHHHHHHHHHHCCCCCCCC |
| 710 | EEQRKKQLEKINNEAEQKRYQREGMGQGHQP | -43.67 | 48.4 | HHHHHHHHHHHHHHHHHHHHHHHHHCCCCCC |
| 808 | EEEKRKMLEQINQEAEQQRYQKEGMGQGHQP | -43.66 | 38.7 | HHHHHHHHHHHHHHHHHHHHHHHHCCCCCCC |
| 397 | EEEKRKMLEQINQEEEQKRYQQQGFGQGHQN | -43.66 | 35.5 | HHHHHHHHHHHHHHHHHHHHHHHCCCCCCCC |
| 906 | EQEKRKMLEQYNQEYEQQEYQRNGRGQGHSY | -43.66 | 32.3 | HHHHHHHHHHHHHHHHHHHHHHHCCCCCCCC |

| | | | | |
|---|---|---|---|---|
| 572 | EEEQRKMLEKINNEKEQKEYQKQGWGQGHNP | -43.66 | 38.7 | HHHHHHHHHHHHHHHHHHHHHHHHCCCCCCCC |
| 163 | EQERRKQLEQINQDEEQRRYQQQGMGQGHQP | -43.65 | 29.0 | HHHHHHHHHHHHHHHHHHHHHHHHHCCCCCCC |
| 029 | EEERRKQLEQINQEEEQKRYQREGFGQGHQS | -43.64 | 35.5 | HHHHHHHHHHHHHHHHHHHHHHHHHCCCCCCC |
| 286 | EEERRKQLEQINQEKEQEEYQRYGMGQGHQP | -43.64 | 35.5 | HHHHHHHHHHHHHHHHHHHHHHHHHHCCCCCC |
| 763 | EEEERKFQEKYNNEYEQKYYQQQGFGQGHQS | -43.63 | 38.7 | CHHHHHHHHHHHHHHHHHHHHHHHCCCCCCCC |
| 667 | EEEKKKWLEQLNNEKEQQRYQREGMGQGHQP | -43.63 | 38.7 | HHHHHHHHHHHHHHHHHHHHHHHHHHCCCCCC |
| 065 | EEEKRKWLEQLNNEQEQQRYQRQGFGQGHQS | -43.63 | 35.5 | HHHHHHHHHHHHHHHHHHHHHHHHCCCCCCCC |
| 445 | EEEKRKMLEQINNEAEQEQFQQRGMGQGKQP | -43.62 | 35.5 | HHHHHHHHHHHHHHHHHHHHHHHHHCCCCCCC |
| 457 | EEEERKFQEKYNNEAEQKRYQREGMGQGHQP | -43.61 | 41.9 | CHHHHHHHHHHHHHHHHHHHHHHHHHCCCCCC |
| 736 | EEERRKQLDAFNNRYEQQYYQQKGFGQGHSS | -43.59 | 38.7 | HHHHHHHHHHHHHHHHHHHHHHHHCCCCCCCC |
| 760 | EEELKKFLEKMKQEKEQEEYQRYGFGQGDSR | -43.58 | 48.4 | HHHHHHHHHHHHHHHHHHHHHHHHCCCCCCCC |
| 600 | EEEKRKFLEQYNNEEEQMRYQREGFGQGHQS | -43.53 | 38.7 | HHHHHHHHHHHCHHHHHHHHHHHHHHCCCCCC |
| 143 | EEEKRKLLEQINNEEEQKRYQRYGFGQGHQS | -43.53 | 35.5 | HHHHHHHHHHHCHHHHHHHHHHHHCCCCCCCC |
| 382 | DEERKKMLDDINNRKEDEEYQRYGFGKGDSR | -43.47 | 48.4 | HHHHHHHHHHHHHHHHHHHHHHHHCCCCCCCC |
| 068 | EEELRKLLEQINNEEEQQRYQKEGFGQGHQS | -43.43 | 35.5 | HHHHHHHHHHHCHHHHHHHHHHHHCCCCCCCC |
| 346 | EEEKRKFLEQFNNEYEQRRYQQEGRGQGHQR | -43.39 | 45.2 | HHHHHHHHHHHHHHHHHHHHHHHHHCCCCCCC |
| 074 | EEELKKLLEQINNEKEQKEYQKQGFGQGNQN | -43.39 | 38.7 | HHHHHHHHHHHHHHHHHHHHHHHHCCCCCCCC |
| 061 | EEEKKKMLEQINNEEEQKRYQREGWGQGHNP | -43.37 | 38.7 | HHHHHHHHHHHCHHHHHHHHHHHHHCCCCCCC |
| 930 | EEEKKKMLEQINNEEEQKSYQREGKGQGHSH | -43.37 | 38.7 | HHHHHHHHHHHHHHHHHHHHHHHHCCCCCCCC |
| 281 | EEEKKKMLEQINNEEEQQRYQRYGMGQGHQP | -43.37 | 38.7 | HHHHHHHHHHHCHHHHHHHHHHHHCCCCCCCC |
| 441 | EEEKKKMLEQINNEKEQEEYQRRGKGQGHSS | -43.37 | 38.7 | HHHHHHHHHHHHHHHHHHHHHHHHHCCCCCCC |
| 101 | EEEKKKMLEQINNEKEQEEYQRRGMGQGHQP | -43.37 | 38.7 | HHHHHHHHHHHHHHHHHHHHHHHHHHCCCCCC |
| 114 | EEEKKKMLEQINNEKEQQEYQKMGLGQGHQP | -43.37 | 41.9 | HHHHHHHHHHHHHHHHHHHHHHHHCCCCCCCC |
| 376 | EEEKKKMLEQINNEKEQQEYQRSGMGQGHQP | -43.37 | 41.9 | HHHHHHHHHHHHHHHHHHHHHHHHHCCCCCCC |
| 201 | EEEKRKMLEQINNEEEQERYQRYGFGQGHQS | -43.37 | 35.5 | HHHHHHHHHHHCCHHHHHHHHHHHCCCCCCCC |
| 493 | EEEKRKMLEQINNEEEQQNYQREGKGQGHSH | -43.37 | 35.5 | HHHHHHHHHHHHHHHHHHHHHHHHCCCCCCCC |
| 826 | EEEKRKMLEQINNEEEQQRYQREGMGQGHQP | -43.37 | 35.5 | HHHHHHHHHHHCHHHHHHHHHHHHHHCCCCCC |
| 241 | EQEKRKMLEQINNEAEQKRYQRYGFGQGHSS | -43.37 | 35.5 | HHHHHHHHHHHHHHHHHHHHHHHHCCCCCCCC |
| 860 | EQEKRKMLEQINNEEEQKRYQREGFGQGHQS | -43.37 | 32.3 | HHHHHHHHHHHCHHHHHHHHHHHHHCCCCCCC |
| 182 | EQEKRKMLEQINNEEEQQRYQQQGMGQGHQP | -43.37 | 32.3 | HHHHHHHHHHHCHHHHHHHHHHHHHCCCCCCC |
| 868 | EEERKKQLEQFNNEYEQQFYQKEGFGQGHQS | -43.34 | 45.2 | HHHHHHHHHHHHHHHHHHHHHHHHCCCCCCCC |
| 334 | EEERKKQLEQINNEEEQKRYQREGMGQGHQP | -43.34 | 38.7 | HHHHHHHHHHHCHHHHHHHHHHHHHCCCCCCC |
| 903 | EEERRKQLEQINNEAEQKRYQRYGMGQGHQP | -43.34 | 38.7 | HHHHHHHHHHHHHHHHHHHHHHHHHCCCCCCC |
| 141 | EEERRKQLEQINNEEEQKRYQREGMGQGHQP | -43.34 | 35.5 | HHHHHHHHHHHCHHHHHHHHHHHHHCCCCCCC |
| 055 | EEERRKQLEQINNEEEQQRYQRYGWGQGHNP | -43.34 | 35.5 | HHHHHHHHHHHCHHHHHHHHHHHHCCCCCCCC |
| 548 | EQERRKQLEQINNEAEQKRYQREGMGQGHQP | -43.34 | 35.5 | HHHHHHHHHHHHHHHHHHHHHHHHHHCCCCCC |
| 081 | EQERRKQLEQINNEEEQQRYQRYGFGQGHQS | -43.34 | 32.3 | HHHHHHHHHHHCHHHHHHHHHHHHCCCCCCCC |
| 158 | EEEKRKFLEKYNNEYEQEEYQRKGRGQGDSR | -43.32 | 48.4 | HHHHHHHHHHHHHHHHHHHHHHHHCCCCCCCC |

| | | | | |
|---|---|---|---|---|
| 251 | EEEKKKFLEQFNNEFEQQYYQQKGFGQGHQS | -43.29 | 45.2 | HHHHHHHHHHHHHHHHHHHHHHHCCCCCCCC |
| 752 | EEEKKKFLEQYNNEYEQMEYQRTGRGQGHQF | -43.29 | 41.9 | HHHHHHHHHHHHHHHHHHHHHHHCCCCCCCC |
| 980 | EEEKKKFLEQYNNEYEQQEYQRTGRGQGHQF | -43.29 | 41.9 | HHHHHHHHHHHHHHHHHHHHHHHHCCCCCCC |
| 444 | EEEKRKFLEQFNNEFEQQYYQQKGQGQGHSH | -43.29 | 41.9 | HHHHHHHHHHHHHHHHHHHHHHHHCCCCCCC |
| 709 | EEEKRKFLEQYNNEYEQEEYQRKGRGQGHQS | -43.29 | 38.7 | HHHHHHHHHHHHHHHHHHHHHHHHCCCCCCC |
| 372 | EEEKRKFLEQYNNEYEQQYYQQQGFGQGHSS | -43.29 | 38.7 | HHHHHHHHHHHHHHHHHHHHHHHCCCCCCCC |
| 126 | EEELKEFLKQFNEEFEDKYYQQKGFGKGDSR | -43.25 | 58.1 | HHHHHHHHHHHHHHHHHHHHHHHCCCCCCCC |
| 085 | EEERRKQLEQINNEEEQKRYQREGQGQGHQS | -43.23 | 35.5 | HHHHHHHHHHHCHHHHHHHHHHHHCCCCCCC |
| 381 | EEEKKKMLEKINNEEEQKRYQREGKGQGHSH | -43.22 | 41.9 | HHHHHHHHHHHCHHHHHHHHHHHHCCCCCCC |
| 938 | EEEKKKMLEKFNNEYEQEEYQRKGRGQGNQN | -43.20 | 45.2 | HHHHHHHHHHHHHHHHHHHHHHHHCCCCCCC |
| 743 | EEEKKKMLEKINNEKEQEEYQRYGFGQGNQN | -43.20 | 41.9 | HHHHHHHHHHHHHHHHHHHHHHHCCCCCCCC |
| 846 | EEEKRKFLEDYNNRYEDEFYQQKGFGKGHSS | -43.19 | 45.2 | HHHHHHHHHHHHHHHHHHHHHHHCCCCCCCC |
| 327 | EEEKRKFLEDYNNRYEDMQYQKEGMGKGHQP | -43.19 | 41.9 | HHHHHHHHHHHHHHHHHHHHHHHHHCCCCCC |
| 025 | EEELKKLLEKINNEEEQKRYQRYGFGQGDSR | -43.18 | 48.4 | HHHHHHHHHHHCHHHHHHHHHHHCCCCCCCC |
| 434 | EQERRHQLEKYNNEYEQEEYQRKGRGQGDSR | -43.15 | 41.9 | HHHHHHHHHHHHHHHHHHHHHHHCCCCCCCC |
| 515 | EQERRHQLEKINNEKEDREYQRRGMGKGHQP | -43.14 | 41.9 | HHHHHHHHHHHHHHHHHHHHHHHHCCCCCCC |
| 145 | EEEKRKFLQQYNNESEQQRYQRYGFGQGHQS | -43.12 | 38.7 | HHHHHHHHHHHHHHHHHHHHHHHCCCCCCCC |
| 332 | EEERRKQLEHYNNRSEDQRYQQYGFGKGHQS | -43.12 | 38.7 | HHHHHHHHHHHHCCCHHHHHHHHCCCCCCCC |
| 356 | EEEKRKLLEQLNNEKEQMRYQRYGFGQGDSR | -43.09 | 41.9 | HHHHHHHHHHHHHHHHHHHHHHHCCCCCCCC |
| 388 | EEERKKQLEDINQRAEDQRYQRYGMGKGHQP | -43.07 | 45.2 | HHHHHHHHHHHHHHHHHHHHHHHHCCCCCCC |
| 666 | EEEKKKFLEQYNNDYEQREYQRKGRGQGDSR | -43.04 | 45.2 | HHHHHHHHHHHHHHHHHHHHHHHCCCCCCCC |
| 132 | EEEKKKWLEQLNNEKEQKRYQREGMGQGHQP | -43.03 | 38.7 | HHHHHHHHHHHHHHHHHHHHHHHHHCCCCCC |
| 200 | EQEKRKFLEQFNNEYEQQFYQQKGFGQGDSR | -43.03 | 48.4 | HHHHHHHHHHHHHHHHHHHHHHHCCCCCCCC |
| 716 | EEEKKKFLEKYNNEYEQKYYQQKGFGQGDSR | -43.02 | 51.6 | HHHHHHHHHHHHHHHHHHHHHHHCCCCCCCC |
| 375 | EEEKRKLLDDINNRAEQQRYQREGKGQGHSS | -43.01 | 38.7 | HHHHHHHHHHHHHHHHHHHHHHHCCCCCCCC |
| 557 | EEEKKKMLEQINNEAEQRRYQREGRGQGHQP | -43.00 | 41.9 | HHHHHHHHHHHHHHHHHHHHHHHHCCCCCCC |
| 822 | EEQKRKMLEQINNEEEQKRYQREGVGQGHQP | -42.93 | 38.7 | HHHHHHHHHHHCHHHHHHHHHHHHCCCCCCC |
| 864 | EEEKRKMLEKINNEAEQKRYQRQGFGQGDSR | -42.89 | 48.4 | HHHHHHHHHHHHHHHHHHHHHHHCCCCCCCC |
| 240 | EEEKKKMLEQINNRAEQERYQREGKGQGHSH | -42.87 | 38.7 | HHHHHHHHHHHHHHHHHHHHHHHHCCCCCCC |
| 341 | EEERRKQLEKINNEEEQKRYQRYGFGQGDSR | -42.87 | 45.2 | HHHHHHHHHHHCHHHHHHHHHHHCCCCCCCC |
| 781 | EEERKKQLEQINNEEEQQRYQQYGFGQGHQS | -42.87 | 38.7 | HHHHHHHHHHHCHHHHHHHHHHHCCCCCCCC |
| 504 | EEERRKQLEQINNEAEQKRYQRYGFGQGHSS | -42.87 | 38.7 | HHHHHHHHHHHHHHHHHHHHHHHCCCCCCCC |
| 354 | EEERRKQLEQINNEKEQQRYQQQGFGQGHSS | -42.87 | 35.5 | HHHHHHHHHHHHHHHHHHHHHHHCCCCCCCC |
| 766 | EEEKKKMLQQINNEKEQEEYQRYGFGQGHQS | -42.73 | 38.7 | HHHHHHHHHHHHHHHHHHHHHHHCCCCCCCC |
| 967 | EEEKRRLLEKINNEAEQKRYQQYGFGQGHSS | -42.68 | 41.9 | HHHHHHHHHHHHHHHHHHHHHHHCCCCCCCC |
| 544 | EEEKKKMLEQINNEQEQQRYQQQGFGQGDSR | -42.64 | 45.2 | HHHHHHHHHHHHHHHHHHHHHHHCCCCCCCC |
| 965 | EQEKRKMLEQFNNEYEQEEYQRKGRGQGDSR | -42.64 | 41.9 | HHHHHHHHHHHHHHHHHHHHHHHCCCCCCCC |
| 486 | EEEKRKFLEKFNNEYEQKYYQQKGFGQGDSR | -42.62 | 51.6 | HHHHHHHHHHHHHHHHHHHHHHHCCCCCCCC |

| | | | | |
|---|---|---|---|---|
| 186 | EEEKKKFLEQYNNESEQQRYQRYGFGQGDSR | -42.52 | 48.4 | HHHHHHHHHHHHCHHHHHHHHHHCCCCCCCC |
| 607 | EEEKRKFLEQYNNEYEQEEYQRKGRGQGDSR | -42.52 | 45.2 | HHHHHHHHHHHHHHHHHHHHHHHCCCCCCCC |
| 187 | EEEKRKFLEQYNNEYEQQEYQRTGRGQGDSR | -42.52 | 45.2 | HHHHHHHHHHHHHHHHHHHHHHHCCCCCCCC |
| 165 | EQEKRKFLEQKNNEEEQKRYQQYGFGQGDSR | -42.52 | 41.9 | HHHHHHHHHHHCHHHHHHHHHHHCCCCCCCC |
| 032 | EEEKRKLLEQIEEEERQKRYQREGFGQGHQS | -42.49 | 29.0 | HHHHHHHHHHHHHHHHHHHHHHHHCCCCCCC |
| 050 | EEERKKQLEQINNEEEQKRYQREGFGQGDSR | -42.47 | 45.2 | HHHHHHHHHHHCHHHHHHHHHHHCCCCCCCC |
| 452 | EEEKRKMLEQFNNEYEQEEYQRKGRGQGNQN | -42.43 | 38.7 | HHHHHHHHHHHHHHHHHHHHHHHHCCCCCCC |
| 361 | EEERRKQLEQINNEAEQKRYQREGKGQGNSN | -42.41 | 38.7 | HHHHHHHHHHHHHHHHHHHHHHHCCCCCCCC |
| 896 | EEEYRKWLEQFKNEYEQEFYQQKGFGQGDSR | -42.40 | 45.2 | HHHHHHHHHHHHHHHHHHHHHHHCCCCCCCC |
| 071 | EEEYRKWLEQLKNEKEQEEYQRRGKGQGDSR | -42.40 | 38.7 | HHHHHHHHHHHHHHHHHHHHHHHCCCCCCCC |
| 765 | EEQKRKYLQQYNNEEEQRQYQRSGMGQGHQP | -42.31 | 41.9 | HHHHHHHHHHHCCHHHHHHHHHHHCCCCCCC |
| 707 | EEEKRKFLEDYNNRSEDQRYQKEGFGKGNQN | -42.26 | 41.9 | HHHHHHHHHHHHCCHHHHHHHHHHCCCCCCC |
| 247 | EEEKRKFLEQFNNEYEQKFYQKEGFGQGDSR | -42.20 | 51.6 | HHHHHHHHHHHHHHHHHHHHHHHCCCCCCCC |
| 136 | EEEKRKLLEQINNEQEQQRYQREGFGQGDSR | -42.09 | 41.9 | HHHHHHHHHHHHHHHHHHHHHHHCCCCCCCC |
| 218 | EQEKRHLLEQISNEEEQQRYQRQGFGQGHQS | -42.09 | 29.0 | HHHHHHHHHHHCHHHHHHHHHHHCCCCCCCC |
| 120 | EEEKKKMLEQINNQKEQEEYQRYGFGQGDSR | -42.05 | 41.9 | HHHHHHHHHHHHHHHHHHHHHHHCCCCCCCC |
| 337 | EEEKRKLLEAINNREEQMRYQREGKGQGHSS | -41.99 | 32.3 | HHHHHHHHHHHHHHHHHHHHHHHCCCCCCCC |
| 156 | EQERRKQLEQINQEEEQQRYQQYGFGQGDSR | -41.50 | 38.7 | HHHHHHHHHHHHHHHHHHHHHHHCCCCCCCC |
| 658 | EQEKRHFLEQFNNEKEQQYYQQYGFGQGHSS | -41.05 | 38.7 | HHHHHHHHHHHHHHHHHHHHHHHCCCCCCCC |
| 408 | EEEKKKFLDDYNNRSEQQRYQRYGFGQGDSR | -40.82 | 48.4 | HHHHHHHHHHHHHHHHHHHHHHHCCCCCCCC |

**Table S4**. Summary of 877 peptide sequences designed using EvoEF2 and the evolutionary profile (weight = 0.75).

| #label | #peptide | #binding(EEU) | #SeqID(%) | #Secondary structure |
|---|---|---|---|---|
| WT | EEQAKTFLDKFNHEAEDLFYQSSGLGKGDFR | -46.46 | 100. | HHHHHHHHHHHHHHHHHHHHHHHCCCCCCCC |
| 376 | EEELKKILEKLNQEKEDMRYQRQGFGKGHQS | -49.06 | 48.4 | HHHHHHHHHHHHHHHHHHHHHHHHCCCCCCC |
| 851 | EEELRKILEKINQEKEDMEYQRAGFGKGHSS | -49.06 | 45.2 | HHHHHHHHHHHHHHHHHHHHHHHHCCCCCCC |
| 412 | EEELKKLLDQINQEAEDMRYQRYGFGKGHSS | -49.05 | 51.6 | HHHHHHHHHHHHHHHHHHHHHHHCCCCCCCC |
| 403 | EEELRKLLDQINQEEEDMRYQQQGFGKGHSA | -49.05 | 45.2 | HHHHHHHHHHHHHHHHHHHHHHHHCCCCCCC |
| 207 | EEELRKLLDQINQEEEDMRYQRYGFGKGHSS | -49.05 | 45.2 | HHHHHHHHHHHHHHHHHHHHHHHCCCCCCCC |
| 957 | EEELRKLLDQINQEKEDEEYQRYGFGKGHSA | -49.05 | 45.2 | HHHHHHHHHHHHHHHHHHHHHHHCCCCCCCC |
| 284 | EEELRKLLDKINQEEEDMRYQQYGFGKGDSR | -49.03 | 54.8 | HHHHHHHHHHHHHHHHHHHHHHHCCCCCCCC |
| 434 | EEELKKFLQKYNEEEEDMRYQREGFGKGHQS | -49.02 | 51.6 | HHHHHHHHHHHHHHHHHHHHHHHHHCCCCCC |
| 602 | EQELRKILEKLNQEQEDMRYQRYGFGKGHSA | -48.87 | 41.9 | HHHHHHHHHHHHHHHHHHHHHHHCCCCCCCC |
| 816 | EEELRKLLQQINEEAEDMRYQQQGFGKGHSA | -48.81 | 45.2 | HHHHHHHHHHHHHHHHHHHHHHHHCCCCCCC |
| 843 | EEELKKILEKINNEAEDMRYQQQGFGKGHSS | -48.77 | 51.6 | HHHHHHHHHHHHHHHHHHHHHHHHCCCCCCC |
| 113 | EEELKKILEKINNEAEDMRYQRYGFGKGHSS | -48.77 | 51.6 | HHHHHHHHHHHHHHHHHHHHHHHCCCCCCCC |
| 793 | EEELKKILEKINNEEEDMRYQQYGFGKGHSA | -48.77 | 48.4 | HHHHHHHHHHHHHHHHHHHHHHHCCCCCCCC |
| 426 | EEELKKILEKINNEEEDMRYQQYGFGKGHSS | -48.77 | 48.4 | HHHHHHHHHHHHHHHHHHHHHHHCCCCCCCC |
| 638 | EEELRKILEKINNEEEDMRYQQQGFGKGHSS | -48.77 | 45.2 | HHHHHHHHHHHHHHHHHHHHHHHHCCCCCCC |
| 417 | EEELRKILEKYNNEEEDMRYQREGFGKGHQA | -48.77 | 45.2 | HHHHHHHHHHHCHHHHHHHHHHHHHCCCCCC |
| 611 | EEEWKKILEKLNNEQEDMRYQQNGFGKGHSA | -48.77 | 48.4 | HHHHHHHHHHHHHHHHHHHHHHHHCCCCCCC |
| 815 | EEELKKLLDQINNEEEDMRYQRYGFGKGHQS | -48.77 | 48.4 | HHHHHHHHHHHHHHHHHHHHHHHCCCCCCCC |
| 968 | EESLRKILDQINNEEEDMRFQQYGFGKGKSR | -48.76 | 45.2 | HHHHHHHHHHHHHHHHHHHHHHHCCCCCCCC |
| 574 | EEELKKLLDQLNNEQEDEEYQRKGFGKGHSS | -48.75 | 48.4 | HHHHHHHHHHHHHHHHHHHHHHHCCCCCCCC |
| 139 | EEELRKLLDQINNEAEDMRYQQQGFGKGHSA | -48.75 | 48.4 | HHHHHHHHHHHHHHHHHHHHHHHHCCCCCCC |
| 838 | EEELRKLLDQINNEEEDMRYQQYGFGKGHSS | -48.75 | 45.2 | HHHHHHHHHHHHHHHHHHHHHHHCCCCCCCC |
| 972 | EEELRKLLDQINNEEEDMRYQREGFGKGHSA | -48.75 | 45.2 | HHHHHHHHHHHHHHHHHHHHHHHHHCCCCCC |
| 872 | EEELRKLLDQINNEEEDMRYQRYGFGKGHSA | -48.75 | 45.2 | HHHHHHHHHHHHHHHHHHHHHHHCCCCCCCC |
| 511 | EEELRKLLDQLNNEKEDMRYQRQGFGKGHSA | -48.75 | 45.2 | HHHHHHHHHHHHHHHHHHHHHHHHCCCCCCC |
| 841 | EEELRKILEKINNEEEDMRYQQYGFGKGHSA | -48.50 | 45.2 | HHHHHHHHHHHHHHHHHHHHHHHCCCCCCCC |
| 591 | EEELRKFLDQFNQEYEDMFYQQKGFGKGHSS | -48.49 | 54.8 | HHHHHHHHHHHHHHHHHHHHHHHHCCCCCCC |
| 243 | EEELRKFLDQYNQEEEDMRYQREGFGKGHSA | -48.49 | 48.4 | HHHHHHHHHHHHHHHHHHHHHHHHHCCCCCC |
| 026 | EEELRKFLQQFNEEYEDMFYQQKGFGKGHSS | -48.25 | 51.6 | HHHHHHHHHHHHHHHHHHHHHHHHCCCCCCC |
| 402 | EEELRKFLQQYNEEHEDMRYQQQGFGKGHSA | -48.25 | 45.2 | HHHHHHHHHHHHHHHHHHHHHHHHCCCCCCC |
| 143 | EEELRKILEKINNEEEDMRYQQYGFGKGHQS | -48.25 | 45.2 | HHHHHHHHHHHHHHHHHHHHHHHCCCCCCCC |
| 317 | EEELRKILEKYNNEEEDMRYQREGFGKGHSS | -48.25 | 45.2 | HHHHHHHHHHHCHHHHHHHHHHHHCCCCCCC |
| 125 | EEELKKFLDQLNNEEEDMRYQRYGFGKGHSS | -48.19 | 51.6 | HHHHHHHHHHHHHHHHHHHHHHHCCCCCCCC |
| 080 | EEELKKFLDQYNNEAEDMRYQQQGFGKGHQS | -48.19 | 54.8 | HHHHHHHHHHHHHHHHHHHHHHHHCCCCCCC |

| | | | | |
|---|---|---|---|---|
| 319 | EEELKKFLDQYNNEAEDMRYQQQGFGKGHSA | -48.19 | 54.8 | HHHHHHHHHHHHHHHHHHHHHHHHCCCCCCC |
| 563 | EEELRKFLDQFNNEKEDMRYQQQGFGKGHSS | -48.19 | 51.6 | HHHHHHHHHHHHHHHHHHHHHHHHCCCCCCC |
| 848 | EEELRKFLDQLNNEEEDMRYQQQGFGKGHQS | -48.19 | 48.4 | HHHHHHHHHHHHHHHHHHHHHHHHCCCCCCC |
| 646 | EEELRKFLDQYNNEAEDMRYQRQGFGKGHSA | -48.19 | 51.6 | HHHHHHHHHHHHHHHHHHHHHHHHCCCCCCC |
| 998 | EEELRKFLDQYNNEEEDMRYQREGFGKGHSA | -48.19 | 48.4 | HHHHHHHHHHHHHHHHHHHHHHHHHCCCCCC |
| 095 | EEELRKFLDQYNNEEEDMRYQREGFGKGHSS | -48.19 | 48.4 | HHHHHHHHHHHHHHHHHHHHHHHHCCCCCCC |
| 571 | EEELRKFLDQYNNEYEDMEYQRNGRGKGHSY | -48.19 | 48.4 | HHHHHHHHHHHHHHHHHHHHHHCCCCCCCCC |
| 781 | EEELRKILEKINNEEEDMRYQQYGFGKGHSS | -48.17 | 45.2 | HHHHHHHHHHHHHHHHHHHHHHHCCCCCCCC |
| 610 | EEELRKILEKLNNEKEDMRYQKEGFGKGHSA | -48.17 | 45.2 | HHHHHHHHHHHHHHHHHHHHHHHHCCCCCCC |
| 073 | EEEKRKILEQFNQEYEDMFYQQQGFGKGHSS | -48.06 | 48.4 | HHHHHHHHHHHHHHHHHHHHHHHHCCCCCCC |
| 830 | EEELRKILEQINQEAEDMRYQQQGFGKGHQS | -48.06 | 45.2 | HHHHHHHHHHHHHHHHHHHHHHHHCCCCCCC |
| 804 | EEELKKLLEKINQEAEDMRYQQQGFGKGHSS | -48.05 | 51.6 | HHHHHHHHHHHHHHHHHHHHHHHHCCCCCCC |
| 444 | EEELKKLLEKINQEAEDMRYQREGFGKGHSS | -48.05 | 51.6 | HHHHHHHHHHHHHHHHHHHHHHHHHCCCCCC |
| 428 | EEELKKLLEKINQEEEDMRYQRQGFGKGHSA | -48.05 | 48.4 | HHHHHHHHHHHHHHHHHHHHHHHHHCCCCCC |
| 495 | EEELRKLLEKINQEAEDIRYQQQGFGKGHSS | -48.05 | 48.4 | HHHHHHHHHHHHHHHHHHHHHHHHCCCCCCC |
| 064 | EEELRKLLEKINQEEEDIRYQQQGFGKGHSS | -48.05 | 45.2 | HHHHHHHHHHHHHHHHHHHHHHHHCCCCCCC |
| 301 | EEELRKLLEKINQEEEDMRYQREGFGKGHSS | -48.05 | 45.2 | HHHHHHHHHHHHHHHHHHHHHHHHHCCCCCC |
| 329 | EEELRKLLEKLNQEKEDIRYQQQGFGKGHQS | -48.05 | 45.2 | HHHHHHHHHHHHHHHHHHHHHHHHCCCCCCC |
| 101 | EEELKKLLEKINNEAEDMRFQQQGFGKGKSR | -48.03 | 51.6 | HHHHHHHHHHHHHHHHHHHHHHHHCCCCCCC |
| 390 | EEELRKLLEKINNEAEDMRFQQQGFGKGKSR | -48.03 | 48.4 | HHHHHHHHHHHHHHHHHHHHHHHHCCCCCCC |
| 548 | EEELRKLLEKINNEEEDMRFQREGFGKGKSR | -48.03 | 45.2 | HHHHHHHHHHHHHHHHHHHHHHHHCCCCCCC |
| 628 | EEELKKLLDQINNEEEDMRYQQQGFGKGDSR | -48.01 | 54.8 | HHHHHHHHHHHHHHHHHHHHHHHCCCCCCCC |
| 723 | EEELRKLLDQLNNEKEDMRYQKEGFGKGDSR | -48.01 | 51.6 | HHHHHHHHHHHHHHHHHHHHHHHHCCCCCCC |
| 735 | EEELRKLLDQINQEEEDMRYQQQGFGKGDSR | -47.81 | 51.6 | HHHHHHHHHHHHHHHHHHHHHHHCCCCCCCC |
| 123 | EEEKRKMLDQINNEEEDMRYQREGFGKGHSS | -47.79 | 45.2 | HHHHHHHHHHHHHHHHHHHHHHHHHCCCCCC |
| 869 | EEELRKILEQINNEEEDMRYQQYGFGKGHSS | -47.78 | 41.9 | HHHHHHHHHHHHHHHHHHHHHHHCCCCCCCC |
| 619 | EEELRKILEQYNNEEEDMRYQRQGFGKGHQS | -47.78 | 41.9 | HHHHHHHHHHHCHHHHHHHHHHHHCCCCCCC |
| 286 | EEELKKILEQINNEAEDMRYQRQGFGKGHSA | -47.77 | 48.4 | HHHHHHHHHHHHHHHHHHHHHHHHHCCCCCC |
| 575 | EEELKKILEQINNEEEDMRYQQQGFGKGHQS | -47.77 | 45.2 | HHHHHHHHHHHHHHHHHHHHHHHHCCCCCCC |
| 060 | EEELKKILEQINNEKEDMRYQQQGFGKGHSS | -47.77 | 45.2 | HHHHHHHHHHHHHHHHHHHHHHHHCCCCCCC |
| 202 | EEELRKILEQINNEEEDMRYQQQGFGKGHSS | -47.77 | 41.9 | HHHHHHHHHHHHHHHHHHHHHHHHCCCCCCC |
| 487 | EEELRKILEQINNEEEDMRYQQYGFGKGHSA | -47.77 | 41.9 | HHHHHHHHHHHHHHHHHHHHHHHCCCCCCCC |
| 544 | EEELRKILEQINNEKEDMEYQKQGFGKGHSA | -47.77 | 41.9 | HHHHHHHHHHHHHHHHHHHHHHHHCCCCCCC |
| 140 | EEELRKILEQINNEKEDMEYQKQGFGKGHSS | -47.77 | 41.9 | HHHHHHHHHHHHHHHHHHHHHHHCCCCCCCC |
| 809 | EEELRKILEQLNNEKEDMRYQKEGFGKGHSS | -47.77 | 41.9 | HHHHHHHHHHHHHHHHHHHHHHHHCCCCCCC |
| 001 | EEELRKILEQLNNEKEDMRYQQQGFGKGHSS | -47.77 | 41.9 | HHHHHHHHHHHHHHHHHHHHHHHHCCCCCCC |
| 635 | EEELRKILEQLNNEKEDMRYQRYGFGKGHSA | -47.77 | 41.9 | HHHHHHHHHHHHHHHHHHHHHHHCCCCCCCC |
| 607 | EEELRKILEQLNNEQEDMRYQQQGFGKGHSA | -47.77 | 41.9 | HHHHHHHHHHHHHHHHHHHHHHHHCCCCCCC |

| | | | | |
|---|---|---|---|---|
| 466 | EEEWRKILEQINNEEEDMRYQREGFGKGHSA | -47.77 | 41.9 | HHHHHHHHHHHHHHHHHHHHHHHHHCCCCCC |
| 564 | EEEWRKILEQLNNEKEDMRYQQNGFGKGHSS | -47.77 | 41.9 | HHHHHHHHHHHHHHHHHHHHHHHCCCCCCCC |
| 373 | EEELRKLLEQLNNDKEDMRYQQQGFGKGHSS | -47.76 | 38.7 | HHHHHHHHHHHHHHHHHHHHHHHHCCCCCCC |
| 174 | EEEKRKFLEKYNQESEDIRFQQQGFGKGKSR | -47.76 | 48.4 | HHHHHHHHHHHHHHHHHHHHHHHCCCCCCCC |
| 103 | EEELKKLLEKINNEAEDIRYQQGGFGKGHES | -47.75 | 51.6 | HHHHHHHHHHHHHHHHHHHHHHCCCCCCCCC |
| 570 | EEELKKLLEKINNEAEDIRYQQQGFGKGHSS | -47.75 | 51.6 | HHHHHHHHHHHHHHHHHHHHHHHHCCCCCCC |
| 685 | EEELKKLLEKINNEAEDMRYQQQGFGKGHSA | -47.75 | 51.6 | HHHHHHHHHHHHHHHHHHHHHHHHCCCCCCC |
| 396 | EEELKKLLEKINNEAEDMRYQRQGFGKGHSA | -47.75 | 51.6 | HHHHHHHHHHHHHHHHHHHHHHHHHCCCCCC |
| 171 | EEELKKLLEKINNEAEDMRYQRQGFGKGHSS | -47.75 | 51.6 | HHHHHHHHHHHHHHHHHHHHHHHHCCCCCCC |
| 124 | EEELKKLLEKINNEEEDIRYQQNGMGKGHSY | -47.75 | 48.4 | HHHHHHHHHHHHHHHHHHHHHHHHCCCCCCC |
| 799 | EEELKKLLEKINNEEEDIRYQQQGFGKGHQS | -47.75 | 48.4 | HHHHHHHHHHHHHHHHHHHHHHHHCCCCCCC |
| 480 | EEELKKLLEKINNEEEDIRYQQQGFGKGHSS | -47.75 | 48.4 | HHHHHHHHHHHHHHHHHHHHHHHHCCCCCCC |
| 692 | EEELKKLLEKINNEEEDMRYQQQGFGKGHSA | -47.75 | 48.4 | HHHHHHHHHHHHHHHHHHHHHHHHCCCCCCC |
| 949 | EEELKKLLEKINNEEEDMRYQQYGFGKGHSA | -47.75 | 48.4 | HHHHHHHHHHHHHHHHHHHHHHHCCCCCCCC |
| 092 | EEELKKLLEKINNEEEDMRYQRQGFGKGHSA | -47.75 | 48.4 | HHHHHHHHHHHHHHHHHHHHHHHHCCCCCCC |
| 240 | EEELKKLLEKINNEKEDEEYQRYGFGKGHSA | -47.75 | 48.4 | HHHHHHHHHHHHHHHHHHHHHHHCCCCCCCC |
| 445 | EEELKKLLEKINNEQEDEEYQRVGRGKGHSY | -47.75 | 48.4 | HHHHHHHHHHHHHHHHHHHHHHHHCCCCCCC |
| 332 | EEELKKLLEKLNNEKEDMRYQRYGFGKGHSA | -47.75 | 48.4 | HHHHHHHHHHHHHHHHHHHHHHHCCCCCCCC |
| 496 | EEELKKLLEKLNNEKEDMYYQQKGFGKGHSS | -47.75 | 48.4 | HHHHHHHHHHHHHHHHHHHHHHHHCCCCCCC |
| 795 | EEELKKLLEKLNNEQEDIRYQQGGFGKGHSF | -47.75 | 48.4 | HHHHHHHHHHHHHHHHHHHHHHCCCCCCCCC |
| 274 | EEELKKLLEKLNNEQEDMQYQRAGFGKGHQS | -47.75 | 48.4 | HHHHHHHHHHHHHHHHHHHHHHHHHCCCCCC |
| 231 | EEELRKLLEKINNEAEDIRYQQNGFGKGHSS | -47.75 | 48.4 | HHHHHHHHHHHHHHHHHHHHHHHCCCCCCCC |
| 637 | EEELRKLLEKINNEAEDMRYQRQGFGKGHSS | -47.75 | 48.4 | HHHHHHHHHHHHHHHHHHHHHHHHCCCCCCC |
| 267 | EEELRKLLEKINNEAEDMRYQRYGFGKGHSS | -47.75 | 48.4 | HHHHHHHHHHHHHHHHHHHHHHHCCCCCCCC |
| 612 | EEELRKLLEKINNEEEDMRYQQQGFGKGHSA | -47.75 | 45.2 | HHHHHHHHHHHHHHHHHHHHHHHHCCCCCCC |
| 933 | EEELRKLLEKINNEEEDMRYQQYGFGKGHSS | -47.75 | 45.2 | HHHHHHHHHHHHHHHHHHHHHHHCCCCCCCC |
| 900 | EEELRKLLEKINNEEEDMRYQREGFGKGHSA | -47.75 | 45.2 | HHHHHHHHHHHHHHHHHHHHHHHHHCCCCCC |
| 709 | EEELRKLLEKINNEEEDMRYQRQGFGKGHSA | -47.75 | 45.2 | HHHHHHHHHHHHHHHHHHHHHHHHCCCCCCC |
| 002 | EEELRKLLEKINNEEEDMRYQRYGFGKGHSA | -47.75 | 45.2 | HHHHHHHHHHHHHHHHHHHHHHHCCCCCCCC |
| 779 | EEELRKLLEKINNEQEDMRYQQQGFGKGHQS | -47.75 | 45.2 | HHHHHHHHHHHHHHHHHHHHHHHHCCCCCCC |
| 363 | EEELRKLLEKINNEQEDMSYQQQGFGKGHSS | -47.75 | 45.2 | HHHHHHHHHHHHHHHHHHHHHHHCCCCCCCC |
| 285 | EEELRKLLEKLNNEKEDMRYQQQGFGKGHSA | -47.75 | 45.2 | HHHHHHHHHHHHHHHHHHHHHHHHCCCCCCC |
| 216 | EEELRKLLEKLNNEQEDMRYQRYGFGKGHSA | -47.75 | 45.2 | HHHHHHHHHHHHHHHHHHHHHHHCCCCCCCC |
| 912 | EEEWKKLIEKINNEAEDMRYQQEGFGKGHSA | -47.75 | 48.4 | HHHHHHHHHHHHHHHHHHHHHHHHHCCCCCC |
| 150 | EQELRKLLEKINNEEEDIRYQQQGFGKGHSS | -47.75 | 41.9 | HHHHHHHHHHHHHHHHHHHHHHHHCCCCCCC |
| 328 | EQELRKLLEKINNEEEDMRYQQQGFGKGHQS | -47.75 | 41.9 | HHHHHHHHHHHHHHHHHHHHHHHHCCCCCCC |
| 520 | EEELKKFLDKLNNEEEDMRYQQYGFGKGDSR | -47.66 | 61.3 | HHHHHHHHHHHHHHHHHHHHHHHCCCCCCCC |
| 566 | EEELKKILEKFNNEYEDMFYQQKGFGKGDSR | -47.52 | 61.3 | HHHHHHHHHHHHHHHHHHHHHHHCCCCCCCC |

| | | | | |
|---|---|---|---|---|
| 430 | EEELKKILEKYNNEAEDIRYQQQGFGKGDSR | -47.52 | 58.1 | HHHHHHHHHHHHHHHHHHHHHHHCCCCCCCC |
| 263 | EEELKKLLEKLNNEKEDMRYQQQGFGKGHQS | -47.51 | 48.4 | HHHHHHHHHHHHHHHHHHHHHHHHCCCCCCC |
| 226 | EEELKKFLDKFNNEYEDRFYQQQGFGKGHSS | -47.51 | 61.3 | HHHHHHHHHHHHHHHHHHHHHHHCCCCCCCC |
| 069 | EEELKKLLEKINNEKEDMEYQKQGFGKGHSS | -47.49 | 48.4 | HHHHHHHHHHHHHHHHHHHHHHHCCCCCCCC |
| 419 | EEELKKFLEKFNQEEEDMRYQREGFGKGHQS | -47.48 | 54.8 | HHHHHHHHHHHHHHHHHHHHHHHHHCCCCCC |
| 215 | EEELKKFLEKYNQEAEDMRYQQQGFGKGHSA | -47.48 | 54.8 | HHHHHHHHHHHHHHHHHHHHHHHHCCCCCCC |
| 159 | EEELKKFLEKYNQEEEDMKYQREGKGKGHSS | -47.48 | 51.6 | HHHHHHHHHHHHHHHHHHHHHHHCCCCCCCC |
| 410 | EEELKKFLEKYNQEEEDMRYQRQGFGKGHSS | -47.48 | 51.6 | HHHHHHHHHHHHHHHHHHHHHHHHCCCCCCC |
| 348 | EEELKKFLEKYNQESEDMRYQQQGFGKGHSS | -47.48 | 51.6 | HHHHHHHHHHHHHHHHHHHHHHHHCCCCCCC |
| 895 | EEELRKFLEKFNQEYEDMFYQQKGFGKGHQS | -47.48 | 54.8 | HHHHHHHHHHHHHHHHHHHHHHHHCCCCCCC |
| 812 | EEELRKFLEKFNQEYEDMFYQQKGFGKGHSA | -47.48 | 54.8 | HHHHHHHHHHHHHHHHHHHHHHHHCCCCCCC |
| 126 | EEELRKFLEKYNQEAEDIYYQQQGFGKGHSS | -47.48 | 51.6 | HHHHHHHHHHHHHHHHHHHHHHHHCCCCCCC |
| 706 | EEELRKFLEKYNQESEDIRYQQQGFGKGHSA | -47.48 | 48.4 | HHHHHHHHHHHHHHHHHHHHHHHHCCCCCCC |
| 690 | EEELRKFLEKFNNEEEDIRFQQQGFGKGKSR | -47.46 | 51.6 | HHHHHHHHHHHCCHHHHHHHHHHHCCCCCCC |
| 389 | EEELRKILEQINNEEEDMRYQREGFGKGHSA | -47.44 | 41.9 | HHHHHHHHHHHHHHHHHHHHHHHHHCCCCCC |
| 915 | EEELRKILEKLNNEQEDRRYQQQGFGKGHQS | -47.37 | 45.2 | HHHHHHHHHHHHHHHHHHHHHHHHCCCCCCC |
| 341 | EEEKRKMLEKINQEAEDMRFQKEGFGKGKSR | -47.37 | 48.4 | HHHHHHHHHHHHHHHHHHHHHHHHCCCCCCC |
| 457 | EEELKKILEQLNQEKEDMRYQREGFGKGDSR | -47.33 | 51.6 | HHHHHHHHHHHHHHHHHHHHHHHHCCCCCCC |
| 648 | EEQLRKLLEKINNEEEDIRYQQQGFGKGHQS | -47.31 | 48.4 | HHHHHHHHHHHHHHHHHHHHHHHHCCCCCCC |
| 613 | EEELRKLLEKINQEAEDMRYQREGFGKGDSR | -47.31 | 54.8 | HHHHHHHHHHHHHHHHHHHHHHHHCCCCCCC |
| 164 | EEELKKLLEQINQEEEDERYQQQGFGKGHSS | -47.28 | 45.2 | HHHHHHHHHHHHHHHHHHHHHHHCCCCCCCC |
| 801 | EEELKKLLEQINQEEEDMRYQQQGFGKGHSA | -47.28 | 45.2 | HHHHHHHHHHHHHHHHHHHHHHHHCCCCCCC |
| 919 | EEELKKLLEQINQEEEDMRYQREGFGKGHSS | -47.28 | 45.2 | HHHHHHHHHHHHHHHHHHHHHHHHHCCCCCC |
| 463 | EEELKKLLEQLNQEKEDMRYQQQGFGKGHSA | -47.28 | 45.2 | HHHHHHHHHHHHHHHHHHHHHHHHCCCCCCC |
| 631 | EEELKKLLEQLNQEQEDMRYQRQGFGKGHSA | -47.28 | 45.2 | HHHHHHHHHHHHHHHHHHHHHHHHHCCCCCC |
| 790 | EEELRKLLEQINQEAEDMRYQKEGFGKGHSA | -47.28 | 45.2 | HHHHHHHHHHHHHHHHHHHHHHHHHCCCCCC |
| 553 | EEELRKLLEQINQEAEDMRYQRQGFGKGHSS | -47.28 | 45.2 | HHHHHHHHHHHHHHHHHHHHHHHHCCCCCCC |
| 547 | EEELRKLLEQINQEEEDMRYQKEGFGKGHSS | -47.28 | 41.9 | HHHHHHHHHHHHHHHHHHHHHHHHCCCCCCC |
| 110 | EEELRKLLEQINQEKEDEEYQRYGFGKGHSS | -47.28 | 41.9 | HHHHHHHHHHHHHHHHHHHHHHHCCCCCCCC |
| 976 | EEELRKLLEQINQEKEDMEYQRAGFGKGHSA | -47.28 | 41.9 | HHHHHHHHHHHHHHHHHHHHHHHHHCCCCCC |
| 055 | EEELRKLLEQLNQEKEDERYQQQGFGKGHSA | -47.28 | 41.9 | HHHHHHHHHHHHHHHHHHHHHHHCCCCCCCC |
| 089 | EQELRKLLEQINQEEEDMRYQREGFGKGHSS | -47.28 | 38.7 | HHHHHHHHHHHHHHHHHHHHHHHHHCCCCCC |
| 228 | EEELKKLLEQINNEEEDMRFQQYGFGKGKSR | -47.26 | 45.2 | HHHHHHHHHHHHHHHHHHHHHHHCCCCCCCC |
| 290 | EEELRKLLEQINNEQEDMFFQQKGFGKGKSR | -47.26 | 45.2 | HHHHHHHHHHHHHHHHHHHHHHHHCCCCCCC |
| 157 | EEELRKILEQINNEKEDMEYQRSGMGKGHQP | -47.25 | 45.2 | HHHHHHHHHHHHHHHHHHHHHHHHCCCCCCC |
| 963 | EEELRKLLEKINNEEEDMRYQQYGFGKGHSA | -47.23 | 45.2 | HHHHHHHHHHHHHHHHHHHHHHHCCCCCCCC |
| 526 | EEELRKLLEQLNNEKEDMRFQRQGFGKGKSS | -47.22 | 38.7 | HHHHHHHHHHHHHHHHHHHHHHHHCCCCCCC |
| 759 | EEELRKFLEQYNNDYEDMEYQRNGRGKGHSY | -47.20 | 41.9 | HHHHHHHHHHHHCHHHHHHHHHCCCCCCCCC |

| | | | | |
|---|---|---|---|---|
| 979 | EEEKRKFLEKFNNEYEDMFYQQKGFGKGHSS | -47.19 | 54.8 | HHHHHHHHHHHHHHHHHHHHHHHHCCCCCCC |
| 958 | EEEKRKFLEKKNNEEEDMRYQRYGFGKGHSS | -47.19 | 48.4 | HHHHHHHHHHHCHHHHHHHHHHHCCCCCCCC |
| 038 | EEEKRKFLEKYNNESEDMRYQQQGFGKGHSS | -47.19 | 48.4 | HHHHHHHHHHHHHHHHHHHHHHHHCCCCCCC |
| 021 | EEEKRKFLEKYNNESEDMRYQRYGFGKGHSA | -47.19 | 48.4 | HHHHHHHHHHHHHCHHHHHHHHECCCCCCCC |
| 832 | EEEKRKFLEKYNNEYEDMFYQQKGFGKGHSA | -47.19 | 51.6 | HHHHHHHHHHHHHHHHHHHHHHHHCCCCCCC |
| 238 | EEELKKFLEKFNNEYEDMFYQKEGFGKGHQS | -47.19 | 58.1 | HHHHHHHHHHHHHHHHHHHHHHHHCCCCCCC |
| 822 | EEELKKFLEKFNNEYEDMFYQQKGFGKGHSA | -47.19 | 58.1 | HHHHHHHHHHHHHHHHHHHHHHHHCCCCCCC |
| 922 | EEELKKFLEKFNNEYEDMYYQQKGFGKGHSS | -47.19 | 54.8 | HHHHHHHHHHHHHHHHHHHHHHHHCCCCCCC |
| 825 | EEELKKFLEKFNNEYEDMYYQQQGFGKGHQS | -47.19 | 54.8 | HHHHHHHHHHHHHHHHHHHHHHHHCCCCCCC |
| 046 | EEELKKFLEKLNNEEEDMRYQQQGFGKGHSS | -47.19 | 51.6 | HHHHHHHHHHHHHHHHHHHHHHHHCCCCCCC |
| 314 | EEELKKFLEKYNNEAEDMRYQKEGFGKGHSA | -47.19 | 54.8 | HHHHHHHHHHHHHHHHHHHHHHHHCCCCCCC |
| 302 | EEELKKFLEKYNNEAEDMRYQQQGFGKGHSS | -47.19 | 54.8 | HHHHHHHHHHHHHHHHHHHHHHHHCCCCCCC |
| 188 | EEELKKFLEKYNNEAEDMRYQQYGFGKGHQS | -47.19 | 54.8 | HHHHHHHHHHHHHHHHHHHHHHHCCCCCCCC |
| 955 | EEELKKFLEKYNNEAEDMRYQREGMGKGHQP | -47.19 | 54.8 | HHHHHHHHHHHHHHHHHHHHHHHHHCCCCCC |
| 045 | EEELKKFLEKYNNEAEDMRYQRQGFGKGHSA | -47.19 | 54.8 | HHHHHHHHHHHHHHHHHHHHHHHHCCCCCCC |
| 106 | EEELKKFLEKYNNEAEDMRYQRYGFGKGHQS | -47.19 | 54.8 | HHHHHHHHHHHHHHHHHHHHHHHCCCCCCCC |
| 199 | EEELKKFLEKYNNEAEDMRYQRYGFGKGHSS | -47.19 | 54.8 | HHHHHHHHHHHHHHHHHHHHHHHCCCCCCCC |
| 770 | EEELKKFLEKYNNEEEDMRYQREGFGKGHSA | -47.19 | 51.6 | HHHHHHHHHHHCCHHHHHHHHHHHHCCCCCC |
| 127 | EEELKKFLEKYNNEEEDMYYQQYGFGKGHSA | -47.19 | 51.6 | HHHHHHHHHHHCHHHHHHHHHHHCCCCCCCC |
| 201 | EEELKKFLEKYNNESEDIRYQQQGFGKGHSS | -47.19 | 51.6 | HHHHHHHHHHHHCHHHHHHHHHHHCCCCCCC |
| 599 | EEELKKFLEKYNNESEDMRYQQQGFGKGHSA | -47.19 | 51.6 | HHHHHHHHHHHHHHHHHHHHHHHHCCCCCCC |
| 623 | EEELKKFLEKYNNESEDMRYQRYGFGKGHSS | -47.19 | 51.6 | HHHHHHHHHHHHHHHHHHHHHEECCCCCCCC |
| 650 | EEELKKFLEKYNNEWEDMRYQREGFGKGHSS | -47.19 | 51.6 | HHHHHHHHHHHHHHHHHHHHHHHHHCCCCCC |
| 582 | EEELKKFLEKYNNEYEDMEYQRTGRGKGHSF | -47.19 | 51.6 | HHHHHHHHHHHHHHHHHHHHHHCCCCCCCCC |
| 679 | EEELKKFLEKYNNEYEDMRYQQEGFGKGHSA | -47.19 | 51.6 | HHHHHHHHHHHHHHHHHHHHHHHHHCCCCCC |
| 023 | EEELRKFLEKFNNEEEDMRYQREGFGKGHQS | -47.19 | 51.6 | HHHHHHHHHHHCHHHHHHHHHHHHHCCCCCC |
| 432 | EEELRKFLEKFNNEEEDMRYQREGFGKGHSS | -47.19 | 51.6 | HHHHHHHHHHHCHHHHHHHHHHHHCCCCCCC |
| 535 | EEELRKFLEKFNNEKEDMRYQQQGFGKGHQS | -47.19 | 51.6 | HHHHHHHHHHHHHHHHHHHHHHHHCCCCCCC |
| 643 | EEELRKFLEKFNNEYEDKFYQQKGFGKGHQS | -47.19 | 54.8 | HHHHHHHHHHHHHHHHHHHHHHHCCCCCCCC |
| 050 | EEELRKFLEKFNNEYEDMFYQKEGFGKGHSS | -47.19 | 54.8 | HHHHHHHHHHHHHHHHHHHHHHHHCCCCCCC |
| 367 | EEELRKFLEKFNNEYEDMFYQQKGFGKGHQS | -47.19 | 54.8 | HHHHHHHHHHHHHHHHHHHHHHHHCCCCCCC |
| 697 | EEELRKFLEKLNNEEEDMRYQRYGFGKGHQS | -47.19 | 48.4 | HHHHHHHHHHHHHHHHHHHHHHHCCCCCCCC |
| 006 | EEELRKFLEKYNNEAEDIRYQQQGFGKGHSA | -47.19 | 51.6 | HHHHHHHHHHHHHHHHHHHHHHHHCCCCCCC |
| 590 | EEELRKFLEKYNNEAEDIRYQQQGFGKGHSS | -47.19 | 51.6 | HHHHHHHHHHHHHHHHHHHHHHHHCCCCCCC |
| 083 | EEELRKFLEKYNNEAEDMRYQKEGFGKGHSA | -47.19 | 51.6 | HHHHHHHHHHHHHHHHHHHHHHHHCCCCCCC |
| 391 | EEELRKFLEKYNNEAEDMRYQQQGFGKGHQS | -47.19 | 51.6 | HHHHHHHHHHHHHHHHHHHHHHHHCCCCCCC |
| 528 | EEELRKFLEKYNNEAEDMRYQQQGFGKGHSA | -47.19 | 51.6 | HHHHHHHHHHHHHHHHHHHHHHHHCCCCCCC |
| 028 | EEELRKFLEKYNNEAEDMRYQRQGFGKGHSA | -47.19 | 51.6 | HHHHHHHHHHHHHHHHHHHHHHHHCCCCCCC |

| | | | | |
|---|---|---|---|---|
| 615 | EEELRKFLEKYNNEAEDMRYQRYGFGKGHSS | -47.19 | 51.6 | HHHHHHHHHHHHHHHHHHHHHHHCCCCCCCC |
| 321 | EEELRKFLEKYNNEHEDIRYQQQGFGKGHSS | -47.19 | 48.4 | HHHHHHHHHHHHHHHHHHHHHHHCCCCCCCC |
| 910 | EEELRKFLEKYNNESEDIRYQQQGFGKGHQS | -47.19 | 48.4 | HHHHHHHHHHHHCHHHHHHHHHHHCCCCCCC |
| 220 | EEELRKFLEKYNNESEDIRYQQQGFGKGHSA | -47.19 | 48.4 | HHHHHHHHHHHHCHHHHHHHHHHHCCCCCCC |
| 448 | EEELRKFLEKYNNESEDIRYQQQGFGKGHSS | -47.19 | 48.4 | HHHHHHHHHHHHCHHHHHHHHHHHCCCCCCC |
| 589 | EEELRKFLEKYNNESEDMRYQQQGFGKGHSA | -47.19 | 48.4 | HHHHHHHHHHHHHHHHHHHHHHHHCCCCCCC |
| 746 | EEELRKFLEKYNNEYEDMEYQRNGRGKGHSY | -47.19 | 48.4 | HHHHHHHHHHHHHHHHHHHHHHCCCCCCCCC |
| 997 | EEELRKFLEKYNNEYEDMFYQQKGFGKGHQS | -47.19 | 51.6 | HHHHHHHHHHHHHHHHHHHHHHHHCCCCCCC |
| 120 | EEELRKFLEKYNNEYEDMFYQQKGFGKGHSA | -47.19 | 51.6 | HHHHHHHHHHHHHHHHHHHHHHHHCCCCCCC |
| 194 | EEELRKFLEKYNNEYEDMFYQQKGFGKGHSS | -47.19 | 51.6 | HHHHHHHHHHHHHHHHHHHHHHHHCCCCCCC |
| 058 | EQELRKFLEKFNNEYEDKFYQQKGFGKGHSA | -47.19 | 51.6 | HHHHHHHHHHHHHHHHHHHHHHHCCCCCCCC |
| 808 | EQELRKFLEKFNNEYEDMFYQQKGFGKGHQS | -47.19 | 51.6 | HHHHHHHHHHHHHHHHHHHHHHHHCCCCCCC |
| 508 | EQELRKFLEKFNNEYEDMFYQQKGFGKGHSA | -47.19 | 51.6 | HHHHHHHHHHHHHHHHHHHHHHHHCCCCCCC |
| 657 | EQELRKFLEKFNNEYEDMFYQQKGFGKGHSS | -47.19 | 51.6 | HHHHHHHHHHHHHHHHHHHHHHHHCCCCCCC |
| 162 | EEELKKILEKLNNEKEHMRKQRQGFGSGKQS | -47.17 | 38.7 | HHHHHHHHHHHHHHHHHHHHHHHHHCCCCCC |
| 725 | EEELRKLLEKINNEEEDIRYQQQGFGKGHQS | -47.16 | 45.2 | HHHHHHHHHHHHHHHHHHHHHHHHCCCCCCC |
| 660 | EEELKKILEQLNNREEDERYQRQGFGKGDSR | -47.10 | 48.4 | HHHHHHHHHHHHHHHHHHHHHHHCCCCCCCC |
| 775 | EEELRKLLEQINNEEEDRRYQQYGFGKGHQP | -47.09 | 41.9 | HHHHHHHHHHHHHHHHHHHHHHHCCCCCCCC |
| 947 | EEEKRKMLEKINQEAEDIRYQQQGFGKGHQA | -47.09 | 48.4 | HHHHHHHHHHHHHHHHHHHHHHHHCCCCCCC |
| 765 | EEEKRKMLEKINQEEEDMRYQQQGFGKGHSA | -47.09 | 45.2 | HHHHHHHHHHHHHHHHHHHHHHHHCCCCCCC |
| 901 | EEELKKFLEKFNNEYENMFKQQKGFGSGKSA | -47.07 | 48.4 | HHHHHHHHHHHHHHHHHHHHHHHHCCCCCCC |
| 481 | EEELRKLLEQLNQEKEDMRYQQQGFGKGHQS | -47.06 | 41.9 | HHHHHHHHHHHHHHHHHHHHHHHHCCCCCCC |
| 015 | EEELKKLLEQINQEAEDMRYQQQGFGKGHQS | -47.04 | 48.4 | HHHHHHHHHHHHHHHHHHHHHHHHCCCCCCC |
| 642 | EEELRKILEQINNEEEDMRYQQQGFGKGDSR | -47.03 | 48.4 | HHHHHHHHHHHHHHHHHHHHHHHCCCCCCCC |
| 214 | EEELRKILEQLNNEQEDMRYQQQGFGKGDSR | -47.03 | 48.4 | HHHHHHHHHHHHHHHHHHHHHHHCCCCCCCC |
| 608 | EEELKKLLEKINNEAEDMRYQRQGFGKGDSR | -47.02 | 58.1 | HHHHHHHHHHHHHHHHHHHHHHHCCCCCCCC |
| 368 | EEELRKLLEKLNNEEEDMKYQRYGFGKGDSR | -47.02 | 51.6 | HHHHHHHHHHHHHHHHHHHHHHHCCCCCCCC |
| 578 | EEELKKLLEQINNEEEDMRYQRQGFGKGHSA | -47.00 | 45.2 | HHHHHHHHHHHHHHHHHHHHHHHHCCCCCCC |
| 962 | EEELRKLLEQINNEAEDMRYQREGMGKGHQP | -47.00 | 45.2 | HHHHHHHHHHHHHHHHHHHHHHHHHCCCCCC |
| 454 | EEELRKLLEQINNEEEDMRYQQQGFGKGHSA | -47.00 | 41.9 | HHHHHHHHHHHHHHHHHHHHHHHHCCCCCCC |
| 903 | EEELRKLLEQINNEEEDMRYQQQGFGKGHSS | -47.00 | 41.9 | HHHHHHHHHHHHHHHHHHHHHHHHCCCCCCC |
| 640 | EEELRKLLEQINNEEEDMRYQRYGFGKGHQS | -47.00 | 41.9 | HHHHHHHHHHHHHHHHHHHHHHHCCCCCCCC |
| 183 | EEELRKLLEQINNEQEDMSYQQQGFGKGHQS | -47.00 | 41.9 | HHHHHHHHHHHHHHHHHHHHHHHHCCCCCCC |
| 252 | EEELRKLLEQLNNEQEDMRYQREGFGKGHSA | -47.00 | 41.9 | HHHHHHHHHHHHHHHHHHHHHHHHHCCCCCC |
| 954 | EEEKRKLLEQINNEEEDERYQREGFGKGHSS | -46.99 | 41.9 | HHHHHHHHHHHHHHHHHHHHHHHHCCCCCCC |
| 242 | EEELKKLLEQINNEAEDERYQQQGFGKGHQS | -46.99 | 48.4 | HHHHHHHHHHHHHHHHHHHHHHHCCCCCCCC |
| 733 | EEELKKLLEQINNEAEDMQYQRAGFGKGHSS | -46.99 | 48.4 | HHHHHHHHHHHHHHHHHHHHHHHHHCCCCCC |
| 203 | EEELKKLLEQINNEAEDMRYQKEGFGKGHSS | -46.99 | 48.4 | HHHHHHHHHHHHHHHHHHHHHHHHCCCCCCC |

| | | | | |
|---|---|---|---|---|
| 122 | EEELKKLLEQINNEAEDMRYQQGGFGKGHQS | -46.99 | 48.4 | HHHHHHHHHHHHHHHHHHHHHHCCCCCCCCC |
| 671 | EEELKKLLEQINNEAEDMRYQQQGFGKGHSA | -46.99 | 48.4 | HHHHHHHHHHHHHHHHHHHHHHHHCCCCCCC |
| 806 | EEELKKLLEQINNEAEDMRYQREGFGKGHSS | -46.99 | 48.4 | HHHHHHHHHHHHHHHHHHHHHHHHHCCCCCC |
| 537 | EEELKKLLEQINNEAEDMRYQRYGFGKGHSS | -46.99 | 48.4 | HHHHHHHHHHHHHHHHHHHHHHHCCCCCCCC |
| 737 | EEELKKLLEQINNEEEDIRYQQQGFGKGHSS | -46.99 | 45.2 | HHHHHHHHHHHHHHHHHHHHHHHHCCCCCCC |
| 100 | EEELKKLLEQINNEEEDMRYQQQGFGKGHQA | -46.99 | 45.2 | HHHHHHHHHHHHHHHHHHHHHHHHCCCCCCC |
| 163 | EEELKKLLEQINNEEEDMRYQQQGFGKGHSA | -46.99 | 45.2 | HHHHHHHHHHHHHHHHHHHHHHHHCCCCCCC |
| 031 | EEELKKLLEQINNEEEDMRYQQYGFGKGHSA | -46.99 | 45.2 | HHHHHHHHHHHHHHHHHHHHHHHCCCCCCCC |
| 700 | EEELKKLLEQINNEEEDMRYQREGFGKGHSA | -46.99 | 45.2 | HHHHHHHHHHHHHHHHHHHHHHHHHCCCCCC |
| 493 | EEELKKLLEQINNEEEDMRYQREGFGKGHSS | -46.99 | 45.2 | HHHHHHHHHHHHHHHHHHHHHHHHHCCCCCC |
| 593 | EEELKKLLEQLNNEKEDERYQQQGFGKGHSS | -46.99 | 45.2 | HHHHHHHHHHHHHHHHHHHHHHHCCCCCCCC |
| 625 | EEELKKLLEQLNNEKEDMRYQQQGFGKGHQS | -46.99 | 45.2 | HHHHHHHHHHHHHHHHHHHHHHHHCCCCCCC |
| 826 | EEELKKLLEQLNNEKEDMRYQRQGFGKGHSS | -46.99 | 45.2 | HHHHHHHHHHHHHHHHHHHHHHHHCCCCCCC |
| 736 | EEELKKLLEQLNNEQEDMRYQRYGFGKGHSS | -46.99 | 45.2 | HHHHHHHHHHHHHHHHHHHHHHHCCCCCCCC |
| 916 | EEELRKLLEQINNEAEDIRYQQQGFGKGHSS | -46.99 | 45.2 | HHHHHHHHHHHHHHHHHHHHHHHHCCCCCCC |
| 653 | EEELRKLLEQINNEAEDMRYQQQGFGKGHQS | -46.99 | 45.2 | HHHHHHHHHHHHHHHHHHHHHHHHCCCCCCC |
| 195 | EEELRKLLEQINNEAEDMRYQQQGFGKGHSA | -46.99 | 45.2 | HHHHHHHHHHHHHHHHHHHHHHHHCCCCCCC |
| 117 | EEELRKLLEQINNEAEDMRYQQQGFGKGHSS | -46.99 | 45.2 | HHHHHHHHHHHHHHHHHHHHHHHHCCCCCCC |
| 996 | EEELRKLLEQINNEAEDMRYQQYGFGKGHSS | -46.99 | 45.2 | HHHHHHHHHHHHHHHHHHHHHHHCCCCCCCC |
| 620 | EEELRKLLEQINNEAEDMRYQRQGFGKGHSS | -46.99 | 45.2 | HHHHHHHHHHHHHHHHHHHHHHHHCCCCCCC |
| 345 | EEELRKLLEQINNEAEDMRYQRYGFGKGHSA | -46.99 | 45.2 | HHHHHHHHHHHHHHHHHHHHHHHCCCCCCCC |
| 485 | EEELRKLLEQINNEAEDMRYQRYGFGKGHSS | -46.99 | 45.2 | HHHHHHHHHHHHHHHHHHHHHHHCCCCCCCC |
| 538 | EEELRKLLEQINNEAEDMSYQKEGFGKGHSS | -46.99 | 45.2 | HHHHHHHHHHHHHHHHHHHHHHHHCCCCCCC |
| 427 | EEELRKLLEQINNEEEDERYQQQGFGKGHSA | -46.99 | 41.9 | HHHHHHHHHHHHHHHHHHHHHHHCCCCCCCC |
| 890 | EEELRKLLEQINNEEEDMEYQRNGFGKGHSA | -46.99 | 41.9 | HHHHHHHHHHHHHHHHHHHHHHHHCCCCCCC |
| 491 | EEELRKLLEQINNEEEDMRYQKEGFGKGHSA | -46.99 | 41.9 | HHHHHHHHHHHHHHHHHHHHHHHHCCCCCCC |
| 961 | EEELRKLLEQINNEEEDMRYQKEGFGKGHSS | -46.99 | 41.9 | HHHHHHHHHHHHHHHHHHHHHHHHCCCCCCC |
| 218 | EEELRKLLEQINNEEEDMRYQQYGFGKGHSA | -46.99 | 41.9 | HHHHHHHHHHHHHHHHHHHHHHHCCCCCCCC |
| 834 | EEELRKLLEQINNEEEDMRYQQYGFGKGHSS | -46.99 | 41.9 | HHHHHHHHHHHHHHHHHHHHHHHCCCCCCCC |
| 398 | EEELRKLLEQINNEEEDMRYQREGFGKGHQS | -46.99 | 41.9 | HHHHHHHHHHHHHHHHHHHHHHHHHCCCCCC |
| 007 | EEELRKLLEQINNEEEDMRYQREGFGKGHSA | -46.99 | 41.9 | HHHHHHHHHHHHHHHHHHHHHHHHHCCCCCC |
| 529 | EEELRKLLEQINNEEEDMRYQRQGFGKGHQS | -46.99 | 41.9 | HHHHHHHHHHHHHHHHHHHHHHHHCCCCCCC |
| 439 | EEELRKLLEQINNEEEDMRYQRQGFGKGHSA | -46.99 | 41.9 | HHHHHHHHHHHHHHHHHHHHHHHHCCCCCCC |
| 881 | EEELRKLLEQINNEEEDQRYQQQGFGKGHSS | -46.99 | 41.9 | HHHHHHHHHHHHHHHHHHHHHHHCCCCCCCC |
| 757 | EEELRKLLEQINNEKEDEEYQRRGMGKGHQP | -46.99 | 41.9 | HHHHHHHHHHHHHHHHHHHHHHHHHCCCCCC |
| 761 | EEELRKLLEQINNEKEDMEYQKEGFGKGHQS | -46.99 | 41.9 | HHHHHHHHHHHHHHHHHHHHHHHHCCCCCCC |
| 649 | EEELRKLLEQINNEKEDMRYQQQGFGKGHSA | -46.99 | 41.9 | HHHHHHHHHHHHHHHHHHHHHHHHCCCCCCC |
| 982 | EEELRKLLEQINNEQEDMYYQQKGFGKGHQS | -46.99 | 41.9 | HHHHHHHHHHHHHHHHHHHHHHHHCCCCCCC |

| | | | | |
|---|---|---|---|---|
| 699 | EEELRKLLEQINNEQEDMYYQQKGFGKGHSA | -46.99 | 41.9 | HHHHHHHHHHHHHHHHHHHHHHHHCCCCCCC |
| 034 | EEELRKLLEQLNNEEEDMKYQQQGFGKGHSA | -46.99 | 41.9 | HHHHHHHHHHHHHHHHHHHHHHHHCCCCCCC |
| 057 | EEELRKLLEQLNNEEEDMKYQQQGFGKGHSS | -46.99 | 41.9 | HHHHHHHHHHHHHHHHHHHHHHHHCCCCCCC |
| 743 | EEELRKLLEQLNNEKEDERYQQQGFGKGHSA | -46.99 | 41.9 | HHHHHHHHHHHHHHHHHHHHHHHCCCCCCCC |
| 568 | EEELRKLLEQLNNEKEDERYQQQGFGKGHSS | -46.99 | 41.9 | HHHHHHHHHHHHHHHHHHHHHHHCCCCCCCC |
| 645 | EEELRKLLEQLNNEKEDIRYQQQGFGKGHSS | -46.99 | 41.9 | HHHHHHHHHHHHHHHHHHHHHHHHCCCCCCC |
| 151 | EEELRKLLEQLNNEKEDMRYQQQGFGKGHSS | -46.99 | 41.9 | HHHHHHHHHHHHHHHHHHHHHHHHCCCCCCC |
| 833 | EEELRKLLEQLNNEKEDMRYQQYGFGKGHQS | -46.99 | 41.9 | HHHHHHHHHHHHHHHHHHHHHHHCCCCCCCC |
| 707 | EEELRKLLEQLNNEKEDMRYQRQGFGKGHQS | -46.99 | 41.9 | HHHHHHHHHHHHHHHHHHHHHHHHCCCCCCC |
| 223 | EEELRKLLEQLNNEKEDMRYQRQGFGKGHSA | -46.99 | 41.9 | HHHHHHHHHHHHHHHHHHHHHHHHCCCCCCC |
| 975 | EEELRKLLEQLNNEKEDMRYQRYGFGKGHSA | -46.99 | 41.9 | HHHHHHHHHHHHHHHHHHHHHHHCCCCCCCC |
| 580 | EEELRKLLEQLNNEQEDMRYQKEGFGKGHSA | -46.99 | 41.9 | HHHHHHHHHHHHHHHHHHHHHHHHHCCCCCC |
| 462 | EEELRKLLEQLNNEQEDMRYQQQGFGKGHDP | -46.99 | 41.9 | HHHHHHHHHHHHHHHHHHHHHHHHCCCCCCC |
| 265 | EEEWKKLMEQYNNESEDMRYQQQGFGKGHSS | -46.99 | 41.9 | HHHHHHHHHHHHHHHHHHHHHHHHCCCCCCC |
| 592 | EEEWRKLMEQYNNEEEDMYYQQQGFGKGHSS | -46.99 | 38.7 | HHHHHHHHHHHHHHHHHHHHHHHHCCCCCCC |
| 760 | EQELRKLLEQINNEKEDMEYQKQGFGKGHQS | -46.99 | 38.7 | HHHHHHHHHHHHHHHHHHHHHHHHCCCCCCC |
| 980 | EEEARKMLEKINNEEEDIRYQQQGFGKGHQS | -46.98 | 48.4 | HHHHHHHHHHHHHHHHHHHHHHHHCCCCCCC |
| 239 | EEELRKFLEKLNNEEEDMRYQQQGFGKGHSA | -46.92 | 48.4 | HHHHHHHHHHHHHHHHHHHHHHHHCCCCCCC |
| 415 | EEELRKFLEKYNNESEDMRYQQQGFGKGHSS | -46.92 | 48.4 | HHHHHHHHHHHHHHHHHHHHHHHHCCCCCCC |
| 740 | EEELRKILQKLNNEQEDMRYQQQGFGKGDSR | -46.89 | 51.6 | HHHHHHHHHHHHHHHHHHHHHHHCCCCCCCC |
| 747 | EEELRKILEQLNNEKEDMRYQQQGFGKGDSR | -46.87 | 48.4 | HHHHHHHHHHHHHHHHHHHHHHHCCCCCCCC |
| 020 | EEELKKLLEQINQEAEDRRYQRYGFGKGHQS | -46.87 | 48.4 | HHHHHHHHHHHHHHHHHHHHHHHCCCCCCCC |
| 715 | EEELKKFLEKFNNEYEDMYYQQQGFGKGHSS | -46.86 | 54.8 | HHHHHHHHHHHHHHHHHHHHHHHCCCCCCCC |
| 236 | EEELRKFLEKYNNEAEDMRYQKEGFGKGHQS | -46.86 | 51.6 | HHHHHHHHHHHHHHHHHHHHHHHHCCCCCCC |
| 490 | EEELRKFLEKYNNEEEDMYYQQYGFGKGHQS | -46.86 | 48.4 | HHHHHHHHHHHCHHHHHHHHHHHCCCCCCCC |
| 750 | EEELKKLLDKLNNEKEQMSYQQQGFGQGHSA | -46.80 | 45.2 | HHHHHHHHHHHHHHHHHHHHHHHCCCCCCCC |
| 326 | EEELKKLLEKLNQEQEDMRYQQQGFGKGDSR | -46.80 | 54.8 | HHHHHHHHHHHHHHHHHHHHHHHCCCCCCCC |
| 777 | EEELRKLLEKINQEAEDISYQQQGFGKGDSR | -46.80 | 54.8 | HHHHHHHHHHHHHHHHHHHHHHHCCCCCCCC |
| 627 | EEELRKLLEKINQEEEDMRYQREGFGKGDSR | -46.80 | 51.6 | HHHHHHHHHHHHHHHHHHHHHHHHCCCCCCC |
| 536 | EEELRKLLEKINQEYEDIFYQQKGFGKGDSR | -46.80 | 54.8 | HHHHHHHHHHHHHHHHHHHHHHCCCCCCCCC |
| 068 | EEEKKKMLEKINNEEEDIRYQQQGFGKGHSA | -46.79 | 48.4 | HHHHHHHHHHHHHHHHHHHHHHHHCCCCCCC |
| 865 | EEEKKKMLEKINNEKEDMEYQRAGFGKGHSA | -46.79 | 48.4 | HHHHHHHHHHHHHHHHHHHHHHHHHCCCCCC |
| 727 | EEEKRKMLEKINNEEEDMRYQQYGFGKGHSS | -46.79 | 45.2 | HHHHHHHHHHHHHHHHHHHHHHHCCCCCCCC |
| 965 | EEEKRKMLEKINNEEEDMRYQRQGFGKGHQS | -46.79 | 45.2 | HHHHHHHHHHHHHHHHHHHHHHHHCCCCCCC |
| 527 | EEQLKKFLEKFNNEYEDMFYQQQGFGKGHSA | -46.75 | 61.3 | HHHHHHHHHHHHHHHHHHHHHHHHCCCCCCC |
| 991 | EEELRKFLEQFNNEYEDMFKQQKGFGSGKSR | -46.74 | 48.4 | HHHHHHHHHHHHHHHHHHHHHHHCCCCCCCC |
| 989 | EESLKKILEQLNNEKEDMRYQQQGFGKGHSS | -46.73 | 45.2 | HHHHHHHHHHHHHHHHHHHHHHHHCCCCCCC |
| 762 | EEELKKFLEQYNQEAEDMRYQRYGFGKGHSA | -46.73 | 51.6 | HHHHHHHHHHHHHHHHHHHHHHHCCCCCCCC |

| | | | | |
|---|---|---|---|---|
| 393 | EEELKKFLEQFNQEYEDMFYQKEGFGKGHSA | -46.72 | 54.8 | HHHHHHHHHHHHHHHHHHHHHHHHCCCCCCC |
| 378 | EEELKKFLEQFNQEYEDMFYQQKGFGKGHSS | -46.72 | 54.8 | HHHHHHHHHHHHHHHHHHHHHHHHCCCCCCC |
| 530 | EEELKKFLEQFNQEYEDMYYQQQGFGKGHSS | -46.72 | 51.6 | HHHHHHHHHHHHHHHHHHHHHHHHCCCCCCC |
| 071 | EEELKKFLEQKNQEEEDIRYQQQGFGKGHQS | -46.72 | 48.4 | HHHHHHHHHHHHHHHHHHHHHHHHCCCCCCC |
| 061 | EEELKKFLEQYNQEAEDIYYQQNGFGKGHQS | -46.72 | 51.6 | HHHHHHHHHHHHHHHHHHHHHHHCCCCCCCC |
| 371 | EEELKKFLEQYNQEAEDMRYQRQGFGKGHSA | -46.72 | 51.6 | HHHHHHHHHHHHHHHHHHHHHHHHHCCCCCC |
| 388 | EEELKKFLEQYNQEYEDEEYQRTGRGKGHSF | -46.72 | 48.4 | HHHHHHHHHHHHHHHHHHHHHHHCCCCCCCC |
| 324 | EEELRKFLEQFNQEKEDMRYQQQGFGKGHSS | -46.72 | 48.4 | HHHHHHHHHHHHHHHHHHHHHHHHCCCCCCC |
| 330 | EEELRKFLEQFNQEYEDEEYQRKGFGKGHSA | -46.72 | 48.4 | HHHHHHHHHHHHHHHHHHHHHHHCCCCCCCC |
| 752 | EEELRKFLEQFNQEYEDEEYQRVGRGKGHSY | -46.72 | 48.4 | HHHHHHHHHHHHHHHHHHHHHHHHCCCCCCC |
| 062 | EEELRKFLEQFNQEYEDMYYQQKGFGKGHQS | -46.72 | 48.4 | HHHHHHHHHHHHHHHHHHHHHHHHCCCCCCC |
| 081 | EEELRKFLEQYNQEEEDMEYQRNGRGKGHSY | -46.72 | 45.2 | HHHHHHHHHHHHHHHHHHHHHHHCCCCCCCC |
| 311 | EEELRKFLEQYNQEEEDMRYQREGFGKGHSA | -46.72 | 45.2 | HHHHHHHHHHHHHHHHHHHHHHHHHCCCCCC |
| 136 | EEELRKFLEQYNQESEDMRYQRYGFGKGHQS | -46.72 | 45.2 | HHHHHHHHHHHHHHHHHHHHHHHCCCCCCCC |
| 134 | EEELRKFLEQYNQEYEDMEYQRTGRGKGHSF | -46.72 | 45.2 | HHHHHHHHHHHHHHHHHHHHHHCCCCCCCCC |
| 503 | EEELRKFLEQYNQEYEDMQYQKEGFGKGHQA | -46.72 | 45.2 | HHHHHHHHHHHHHHHHHHHHHHHHCCCCCCC |
| 115 | EEELRKFLEQYNQEYEDMRYQQEGFGKGHSA | -46.72 | 45.2 | HHHHHHHHHHHHHHHHHHHHHHHHHCCCCCC |
| 767 | EEEQRKFEEQRNQEEEDMRYQQQGFGKGHSA | -46.72 | 41.9 | HHHHHHHHHHHHHHHHHHHHHHHHCCCCCCC |
| 555 | EQELRKFLEQFNQEKEDMRYQQQGFGKGHQS | -46.72 | 45.2 | HHHHHHHHHHHHHHHHHHHHHHHHCCCCCCC |
| 051 | EQELRKFLEQMNQEKEDMEYQRNGSGKGHSY | -46.72 | 41.9 | HHHHHHHHHHHHHHHHHHHHHHHCCCCCCCC |
| 224 | EEELRKLLEQIEQEREDMEYQQKGFGKGDSR | -46.71 | 45.2 | HHHHHHHHHHHHHHHHHHHHHHHCCCCCCCC |
| 749 | EEELRKFLEKYNNEAEDMRYQQQGFGKGHSS | -46.67 | 51.6 | HHHHHHHHHHHHHHHHHHHHHHHHCCCCCCC |
| 940 | EEELRKFLEQYNNEEEDMSFQKEGFGKGKSA | -46.66 | 41.9 | HHHHHHHHHHHHHHHHHHHHHHHHCCCCCCC |
| 663 | EEELRKLLEKINQEAEDRRYQRQGFGKGHQS | -46.65 | 48.4 | HHHHHHHHHHHHHHHHHHHHHHHHCCCCCCC |
| 256 | EEELRKLLEQINNEEEQMRYQQYGFGRGHSA | -46.63 | 35.5 | HHHHHHHHHHHCHHHHHHHHHHHCCCCCCCC |
| 251 | EEEKRKFLEKFNNEYEDIFYQQKGFGKGHSA | -46.59 | 54.8 | HHHHHHHHHHHHHHHHHHHHHHHHCCCCCCC |
| 886 | EEEKKKWLEQLNQEQEDMQYQRAGFGKGHSS | -46.58 | 45.2 | HHHHHHHHHHHHHHHHHHHHHHHHHCCCCCC |
| 037 | EEEKKKMLEQINNEAEDMRKQRYGFGSGKSR | -46.56 | 45.2 | HHHHHHHHHHHHHHHHHHHHHHHCCCCCCCC |
| 029 | EEELRKFLQKYNNEAEDIRYQQQGFGKGHSA | -46.55 | 51.6 | HHHHHHHHHHHHHHHHHHHHHHHHCCCCCCC |
| 305 | EEEKKKMLDQINNEAEDMRYQRQGFGKGDSR | -46.55 | 58.1 | HHHHHHHHHHHHHHHHHHHHHHHCCCCCCCC |
| 845 | EEELRKLLEQINQEEEDMRYQQYGFGKGDSR | -46.54 | 48.4 | HHHHHHHHHHHHHHHHHHHHHHHCCCCCCCC |
| 017 | EEELRKFLEQYNNEEEDRRYQRQGFGKGHSS | -46.53 | 45.2 | HHHHHHHHHHHCHHHHHHHHHHHHCCCCCCC |
| 569 | EEELRKILEQLNNEQEDMRYQREGFGKGDSR | -46.52 | 48.4 | HHHHHHHHHHHHHHHHHHHHHHHHCCCCCCC |
| 312 | EEELKKLLEKIENEREDREYQRQGFGKGHSA | -46.52 | 45.2 | HHHHHHHHHHHHHHHHHHHHHHHHCCCCCCC |
| 866 | EEELKKLLEKINNEAEDMRYQQQGFGKGDSR | -46.51 | 58.1 | HHHHHHHHHHHHHHHHHHHHHHHCCCCCCCC |
| 515 | EEELKKLLEKINNEQEDMSYQRYGFGKGDSR | -46.51 | 54.8 | HHHHHHHHHHHHHHHHHHHHHHCCCCCCCCC |
| 325 | EEELRKLLEKINNEAEDERYQQQGFGKGDSR | -46.51 | 54.8 | HHHHHHHHHHHHHHHHHHHHHHHCCCCCCCC |
| 121 | EEELRKLLEKINNEAEDMRYQKEGFGKGDSR | -46.51 | 54.8 | HHHHHHHHHHHHHHHHHHHHHHHHCCCCCCC |

| | | | | |
|---|---|---|---|---|
| 550 | EEELRKLLEKINNEAEDMRYQQQGFGKGDSR | -46.51 | 54.8 | HHHHHHHHHHHHHHHHHHHHHHHCCCCCCCC |
| 855 | EQELRKLLEKLNNEKEDIRYQQQGFGKGDSR | -46.51 | 48.4 | HHHHHHHHHHHHHHHHHHHHHHHCCCCCCCC |
| 664 | EEELKKFLEKYNNEAEDIYYQQNGFGKGDSR | -46.45 | 61.3 | HHHHHHHHHHHHHHHHHHHHHHHCCCCCCCC |
| 512 | EEELRKFLEKFNNEYEDMFYQQKGFGKGDSR | -46.45 | 61.3 | HHHHHHHHHHHHHHHHHHHHHHCCCCCCCCC |
| 413 | EEELRKFLEKYNNESEDMRYQRYGFGKGDSR | -46.45 | 54.8 | HHHHHHHHHHHHHHHHHHHHHECCCCCCCCC |
| 014 | EEELKKFLEQFNNEYEDMFYQQKGFGKGHSS | -46.44 | 54.8 | HHHHHHHHHHHHHHHHHHHHHHHHCCCCCCC |
| 654 | EEELKKFLEQLNNEEEDMRYQQQGFGKGHQS | -46.44 | 48.4 | HHHHHHHHHHHHHHHHHHHHHHHHCCCCCCC |
| 945 | EEELRKFLEQFNNEYEDMFYQQKGFGKGHSA | -46.44 | 51.6 | HHHHHHHHHHHHHHHHHHHHHHHHCCCCCCC |
| 874 | EEELRKFLEQFNNEYEDMYYQQKGFGKGHSS | -46.44 | 48.4 | HHHHHHHHHHHHHHHHHHHHHHHHCCCCCCC |
| 497 | EEELRKFLEQFNNEYEDMYYQQQGFGKGHQS | -46.44 | 48.4 | HHHHHHHHHHHHHHHHHHHHHHHHCCCCCCC |
| 889 | EEELRKFLEQKNNEEEDMRYQQYGFGKGHSS | -46.44 | 45.2 | HHHHHHHHHHHHHHHHHHHHHHHCCCCCCCC |
| 070 | EEELKKILEQLNNEKEDMRYQQYGFGKGDSR | -46.44 | 51.6 | HHHHHHHHHHHHHHHHHHHHHHHCCCCCCCC |
| 248 | EQELRKFLEQLNNDEEDRRYQQYGFGKGHSA | -46.43 | 38.7 | HHHHHHHHHHHHHHHHHHHHHHHCCCCCCCC |
| 695 | EEEKRKFLEQLNNEEEDMRYQQYGFGKGHSS | -46.42 | 45.2 | HHHHHHHHHHHHHHHHHHHHHHHCCCCCCCC |
| 944 | EEEKRKFLEQYNNEEEDMRYQREGFGKGHSS | -46.42 | 45.2 | HHHHHHHHHHHCHHHHHHHHHHHHCCCCCCC |
| 636 | EEELKKFLEQFNNEFEDMYYQQKGFGKGHSS | -46.42 | 51.6 | HHHHHHHHHHHHHHHHHHHHHHHHCCCCCCC |
| 135 | EEELKKFLEQFNNEKEDMRYQQQGFGKGHSA | -46.42 | 51.6 | HHHHHHHHHHHHHHHHHHHHHHHHCCCCCCC |
| 702 | EEELKKFLEQFNNEKEDMRYQQQGFGKGHSS | -46.42 | 51.6 | HHHHHHHHHHHHHHHHHHHHHHHHCCCCCCC |
| 861 | EEELKKFLEQFNNEYEDEFYQQKGFGKGHSA | -46.42 | 54.8 | HHHHHHHHHHHHHHHHHHHHHHHCCCCCCCC |
| 800 | EEELKKFLEQFNNEYEDEFYQQKGFGKGHSS | -46.42 | 54.8 | HHHHHHHHHHHHHHHHHHHHHHHCCCCCCCC |
| 040 | EEELKKFLEQFNNEYEDMFYQKEGFGKGHSA | -46.42 | 54.8 | HHHHHHHHHHHHHHHHHHHHHHHHCCCCCCC |
| 609 | EEELKKFLEQFNNEYEDMFYQKEGFGKGHSS | -46.42 | 54.8 | HHHHHHHHHHHHHHHHHHHHHHHHCCCCCCC |
| 185 | EEELKKFLEQFNNEYEDMFYQQKGFGKGHQS | -46.42 | 54.8 | HHHHHHHHHHHHHHHHHHHHHHHHCCCCCCC |
| 994 | EEELKKFLEQFNNEYEDMFYQQKGFGKGHSA | -46.42 | 54.8 | HHHHHHHHHHHHHHHHHHHHHHHHCCCCCCC |
| 629 | EEELKKFLEQFNNEYEDMKYQEQGFGKGHSA | -46.42 | 51.6 | HHHHHHHHHHHHHHHHHHHHHHHHCCCCCCC |
| 616 | EEELKKFLEQFNNEYEDMYYQQKGFGKGHSS | -46.42 | 51.6 | HHHHHHHHHHHHHHHHHHHHHHHHCCCCCCC |
| 501 | EEELKKFLEQFNNEYEDMYYQQQGFGKGHSS | -46.42 | 51.6 | HHHHHHHHHHHHHHHHHHHHHHHCCCCCCCC |
| 047 | EEELKKFLEQLNNEEEDMRYQQYGFGKGHSA | -46.42 | 48.4 | HHHHHHHHHHHHHHHHHHHHHHHCCCCCCCC |
| 771 | EEELKKFLEQLNNEKEDEEYQRYGFGKGHSA | -46.42 | 48.4 | HHHHHHHHHHHHHHHHHHHHHHHCCCCCCCC |
| 433 | EEELKKFLEQYNNEAEDIRYQQGGFGKGHQS | -46.42 | 51.6 | HHHHHHHHHHHHHHHHHHHHHHCCCCCCCCC |
| 896 | EEELKKFLEQYNNEAEDMRYQQQGFGKGHSA | -46.42 | 51.6 | HHHHHHHHHHHHHHHHHHHHHHHHCCCCCCC |
| 131 | EEELKKFLEQYNNEAEDMRYQRQGFGKGHSA | -46.42 | 51.6 | HHHHHHHHHHHHHHHHHHHHHHHHCCCCCCC |
| 132 | EEELKKFLEQYNNEAEDMRYQRQGFGKGHSS | -46.42 | 51.6 | HHHHHHHHHHHHHHHHHHHHHHHHCCCCCCC |
| 587 | EEELKKFLEQYNNEAEDMRYQRYGFGKGHSA | -46.42 | 51.6 | HHHHHHHHHHHHHHHHHHHHHHHCCCCCCCC |
| 701 | EEELKKFLEQYNNEAEDMRYQRYGFGKGHSS | -46.42 | 51.6 | HHHHHHHHHHHHHHHHHHHHHHHCCCCCCCC |
| 355 | EEELKKFLEQYNNEAEDMYYQRYGFGKGHSS | -46.42 | 51.6 | HHHHHHHHHHHHHHHHHHHHHHHCCCCCCCC |
| 177 | EEELKKFLEQYNNEEEDMEYQRNGFGKGHQS | -46.42 | 48.4 | HHHHHHHHHHHCCHHHHHHHHHHHCCCCCCC |
| 500 | EEELKKFLEQYNNEEEDMRYQREGFGKGHSA | -46.42 | 48.4 | HHHHHHHHHHHCHHHHHHHHHHHHHCCCCCC |

| | | | | |
|---|---|---|---|---|
| 938 | EEELKKFLEQYNNESEDERYQQQGFGKGHQS | -46.42 | 48.4 | HHHHHHHHHHHHCCHHHHHHHHHCCCCCCCC |
| 956 | EEELKKFLEQYNNESEDERYQQQGFGKGHSA | -46.42 | 48.4 | HHHHHHHHHHHHCCHHHHHHHHHCCCCCCCC |
| 827 | EEELKKFLEQYNNESEDMRYQKEGFGKGHSA | -46.42 | 48.4 | HHHHHHHHHHHHHHHHHHHHHHHHCCCCCCC |
| 698 | EEELKKFLEQYNNESEDMRYQQGGFGKGHQS | -46.42 | 48.4 | HHHHHHHHHHHHHHHHHHHHHHCCCCCCCCC |
| 386 | EEELKKFLEQYNNESEDMRYQQQGFGKGHQS | -46.42 | 48.4 | HHHHHHHHHHHHHHHHHHHHHHHHCCCCCCC |
| 172 | EEELKKFLEQYNNESEDMRYQQQGFGKGHSA | -46.42 | 48.4 | HHHHHHHHHHHHHHHHHHHHHHHHCCCCCCC |
| 025 | EEELKKFLEQYNNESEDMRYQQQGFGKGHSS | -46.42 | 48.4 | HHHHHHHHHHHHHHHHHHHHHHHHCCCCCCC |
| 675 | EEELKKFLEQYNNESEDMRYQRYGFGKGHQS | -46.42 | 48.4 | HHHHHHHHHHHHHHHHHHHHHHHCCCCCCCC |
| 016 | EEELKKFLEQYNNESEDMRYQRYGFGKGHSA | -46.42 | 48.4 | HHHHHHHHHHHHHHHHHHHHHHECCCCCCCC |
| 576 | EEELKKFLEQYNNEYEDEEYQRKGFGKGHSA | -46.42 | 48.4 | HHHHHHHHHHHHHHHHHHHHHHHCCCCCCCC |
| 674 | EEELKKFLEQYNNEYEDEEYQRKGFGKGHSS | -46.42 | 48.4 | HHHHHHHHHHHHHHHHHHHHHHHCCCCCCCC |
| 748 | EEELRKFLEQFNNEEEDERYQQQGFGKGHSS | -46.42 | 48.4 | HHHHHHHHHHHCHHHHHHHHHHHCCCCCCCC |
| 783 | EEELRKFLEQFNNEYEDEFYQQKGFGKGHSA | -46.42 | 51.6 | HHHHHHHHHHHHHHHHHHHHHHHCCCCCCCC |
| 192 | EEELRKFLEQFNNEYEDMFYQQKGFGKGHQS | -46.42 | 51.6 | HHHHHHHHHHHHHHHHHHHHHHHHCCCCCCC |
| 782 | EEELRKFLEQFNNEYEDMYYQQKGFGKGHQS | -46.42 | 48.4 | HHHHHHHHHHHHHHHHHHHHHHHHCCCCCCC |
| 460 | EEELRKFLEQFNNEYEDMYYQQKGFGKGHSA | -46.42 | 48.4 | HHHHHHHHHHHHHHHHHHHHHHHHCCCCCCC |
| 277 | EEELRKFLEQFNNEYEDMYYQQNGSGKGHSY | -46.42 | 48.4 | HHHHHHHHHHHHHHHHHHHHHHCCCCCCCCC |
| 624 | EEELRKFLEQFNNEYEDMYYQQQGFGKGHSA | -46.42 | 48.4 | HHHHHHHHHHHHHHHHHHHHHHHHCCCCCCC |
| 254 | EEELRKFLEQLNNEEEDMRYQQQGFGKGHQA | -46.42 | 45.2 | HHHHHHHHHHHHHHHHHHHHHHHHCCCCCCC |
| 438 | EEELRKFLEQLNNEEEDMRYQQQGFGKGHSA | -46.42 | 45.2 | HHHHHHHHHHHHHHHHHHHHHHHHCCCCCCC |
| 049 | EEELRKFLEQLNNEEEDMRYQQYGFGKGHSA | -46.42 | 45.2 | HHHHHHHHHHHHHHHHHHHHHHHCCCCCCCC |
| 447 | EEELRKFLEQLNNEEEDMRYQQYGFGKGHSS | -46.42 | 45.2 | HHHHHHHHHHHHHHHHHHHHHHHCCCCCCCC |
| 091 | EEELRKFLEQYNNEAEDMRYQKEGFGKGHSA | -46.42 | 48.4 | HHHHHHHHHHHHHHHHHHHHHHHHCCCCCCC |
| 175 | EEELRKFLEQYNNEAEDMRYQQQGFGKGHQS | -46.42 | 48.4 | HHHHHHHHHHHHHHHHHHHHHHHHCCCCCCC |
| 138 | EEELRKFLEQYNNEAEDMRYQQQGFGKGHSS | -46.42 | 48.4 | HHHHHHHHHHHHHHHHHHHHHHHHCCCCCCC |
| 966 | EEELRKFLEQYNNEAEDMRYQRQGFGKGHSS | -46.42 | 48.4 | HHHHHHHHHHHHHHHHHHHHHHHHCCCCCCC |
| 494 | EEELRKFLEQYNNEEEDERYQQQGFGKGHSA | -46.42 | 45.2 | HHHHHHHHHHHCCHHHHHHHHHHCCCCCCCC |
| 107 | EEELRKFLEQYNNEEEDMQYQRSGMGKGHQP | -46.42 | 48.4 | HHHHHHHHHHHCHHHHHHHHHHHHCCCCCCC |
| 264 | EEELRKFLEQYNNEEEDMRYQREGFGKGHSA | -46.42 | 45.2 | HHHHHHHHHHHCHHHHHHHHHHHHHCCCCCC |
| 349 | EEELRKFLEQYNNEEEDMRYQREGFGKGHSS | -46.42 | 45.2 | HHHHHHHHHHHCHHHHHHHHHHHHCCCCCCC |
| 272 | EEELRKFLEQYNNEEEDMRYQREGKGKGHSH | -46.42 | 45.2 | HHHHHHHHHHHCHHHHHHHHHHHHCCCCCCC |
| 974 | EEELRKFLEQYNNEEEDMRYQRQGFGKGHSS | -46.42 | 45.2 | HHHHHHHHHHHCHHHHHHHHHHHHCCCCCCC |
| 211 | EEELRKFLEQYNNEHEDMRYQQQGFGKGHSA | -46.42 | 45.2 | HHHHHHHHHHHHHHHHHHHHHHHHCCCCCCC |
| 452 | EEELRKFLEQYNNESEDIRYQQQGFGKGHSS | -46.42 | 45.2 | HHHHHHHHHHHHHHHHHHHHHHHHCCCCCCC |
| 858 | EEELRKFLEQYNNESEDMRYQKEGFGKGHSA | -46.42 | 45.2 | HHHHHHHHHHHHHHHHHHHHHHHHCCCCCCC |
| 810 | EEELRKFLEQYNNESEDMRYQQGGFGKGHSF | -46.42 | 45.2 | HHHHHHHHHHHHHHHHHHHHHHCCCCCCCCC |
| 067 | EEELRKFLEQYNNESEDMRYQQGGFGKGHSS | -46.42 | 45.2 | HHHHHHHHHHHHHHHHHHHHHHCCCCCCCCC |
| 724 | EEELRKFLEQYNNESEDMRYQQQGFGKGHQS | -46.42 | 45.2 | HHHHHHHHHHHHHHHHHHHHHHHHCCCCCCC |

| | | | | |
|---|---|---|---|---|
| 431 | EEELRKFLEQYNNESEDMRYQQQGFGKGHSA | -46.42 | 45.2 | HHHHHHHHHHHHHHHHHHHHHHHHCCCCCCC |
| 588 | EEELRKFLEQYNNESEDMRYQQQGFGKGHSS | -46.42 | 45.2 | HHHHHHHHHHHHHHHHHHHHHHHHCCCCCCC |
| 381 | EEELRKFLEQYNNESEDMRYQRYGFGKGHSA | -46.42 | 45.2 | HHHHHHHHHHHHHHHHHHHHHHECCCCCCCC |
| 883 | EEELRKFLEQYNNEYEDEEYQRKGFGKGHSA | -46.42 | 45.2 | HHHHHHHHHHHHHHHHHHHHHHHCCCCCCCC |
| 842 | EEELRKFLEQYNNEYEDEEYQRKGFGKGHSS | -46.42 | 45.2 | HHHHHHHHHHHHHHHHHHHHHHHCCCCCCCC |
| 418 | EEELRKFLEQYNNEYEDMFYQQKGFGKGHSS | -46.42 | 48.4 | HHHHHHHHHHHHHHHHHHHHHHHHCCCCCCC |
| 729 | EEEWKKFIEQFNNEYEDEEYQRKGFGKGHQS | -46.42 | 48.4 | HHHHHHHHHHHHHHHHHHHHHHHCCCCCCCC |
| 880 | EQELRKFLEQYNNEAEDMRYQQQGFGKGHSS | -46.42 | 45.2 | HHHHHHHHHHHHHHHHHHHHHHHHCCCCCCC |
| 687 | EQELRKFLEQYNNEEEDMRYQREGFGKGHSA | -46.42 | 41.9 | HHHHHHHHHHHHHHHHHHHHHHHHHCCCCCC |
| 399 | EQELRKFLEQYNNEEEDMSYQRYGFGKGHQA | -46.42 | 41.9 | HHHHHHHHHHHHHHHHHHHHHHHCCCCCCCC |
| 546 | EQELRKFLEQYNNESEDERYQQQGFGKGHSS | -46.42 | 41.9 | HHHHHHHHHHHHHCHHHHHHHHHCCCCCCCC |
| 552 | EQELRKFLEQYNNESEDMRYQRYGFGKGHQS | -46.42 | 41.9 | HHHHHHHHHHHHHHHHHHHHHHHCCCCCCCC |
| 300 | EEELKKILEKLNQEKEQMSYQQQGFGQGHSA | -46.40 | 41.9 | HHHHHHHHHHHHHHHHHHHHHHHCCCCCCCC |
| 397 | EEELKKLLEQINNEEEDMRYQQYGFGKGHSS | -46.39 | 45.2 | HHHHHHHHHHHHHHHHHHHHHHHCCCCCCCC |
| 837 | EEELKKLLDKINQEEEQIRYQQQGFGQGDSR | -46.38 | 51.6 | HHHHHHHHHHHHHHHHHHHHHHHCCCCCCCC |
| 295 | EEELKKLLEKINNEAEDRRYQQQGFGKGHSS | -46.36 | 51.6 | HHHHHHHHHHHHHHHHHHHHHHHCCCCCCCC |
| 416 | EEELKKLLEKINNEEEDRRYQREGFGKGHSA | -46.36 | 48.4 | HHHHHHHHHHHHHHHHHHHHHHHHCCCCCCC |
| 008 | EEELRKLLEKINNEQEDRSYQQQGFGKGHSA | -46.36 | 45.2 | HHHHHHHHHHHHHHHHHHHHHHHCCCCCCCC |
| 056 | EEEKKKMLEQINQEAEDMRYQRYGFGKGHSA | -46.32 | 48.4 | HHHHHHHHHHHHHHHHHHHHHHHCCCCCCCC |
| 868 | EEEKKKMLEQINQEEEDMRYQQYGFGKGHSS | -46.32 | 45.2 | HHHHHHHHHHHHHHHHHHHHHHHCCCCCCCC |
| 209 | EEEKKKMLEQINQEQEDMSYQQQGFGKGHSS | -46.32 | 45.2 | HHHHHHHHHHHHHHHHHHHHHHHCCCCCCCC |
| 893 | EEEKKKMLEQYNQESEDMRYQQQGFGKGHSA | -46.32 | 45.2 | HHHHHHHHHHHHHHHHHHHHHHHHCCCCCCC |
| 936 | EEEKRKMLEQINQEAEDMRYQKEGFGKGHSA | -46.32 | 45.2 | HHHHHHHHHHHHHHHHHHHHHHHHHCCCCCC |
| 914 | EEEKRKMLEQINQEEEDMRYQRYGFGKGHSA | -46.32 | 41.9 | HHHHHHHHHHHHHHHHHHHHHHHCCCCCCCC |
| 044 | EEEKRKFLDQYNNDAEQMRYQRQGFGQGHQS | -46.30 | 41.9 | HHHHHHHHHHHHHHHHHHHHHHHHCCCCCCC |
| 857 | EEEWKKWIEQLNNEQEDMRYQRYGFGKGHSS | -46.29 | 41.9 | HHHHHHHHHHHHHHHHHHHHHHHCCCCCCCC |
| 308 | EEELKKLLEQINNEAEDIRYQQQGFGKGDSR | -46.25 | 54.8 | HHHHHHHHHHHHHHHHHHHHHHHCCCCCCCC |
| 197 | EEELKKLLEQINNEEEDERYQQQGFGKGDSR | -46.25 | 51.6 | HHHHHHHHHHHHHHHHHHHHHHHCCCCCCCC |
| 502 | EEELKKLLEQINNEEEDMRYQREGFGKGDSR | -46.25 | 51.6 | HHHHHHHHHHHHHHHHHHHHHHHHCCCCCCC |
| 259 | EEELKKLLEQLNNEKEDMRYQQQGFGKGDSR | -46.25 | 51.6 | HHHHHHHHHHHHHHHHHHHHHHHCCCCCCCC |
| 165 | EEELRKLLEQINNEAEDMRYQQQGFGKGDSR | -46.25 | 51.6 | HHHHHHHHHHHHHHHHHHHHHHHCCCCCCCC |
| 605 | EEELKKLLEKLNNEKEDMYYQQKGFGKGDSR | -46.24 | 54.8 | HHHHHHHHHHHHHHHHHHHHHHHCCCCCCCC |
| 269 | EEELRKFLEKLNNEEEQIRYQQQGFGRGHSA | -46.23 | 41.9 | HHHHHHHHHHHCHHHHHHHHHHHHCCCCCCC |
| 212 | EEELKKFLEQFNQEYEDMRYQQEGFGKGHSA | -46.20 | 51.6 | HHHHHHHHHHHHHHHHHHHHHHHHHCCCCCC |
| 105 | EEELRKFLEQMNNEKEDMEYQRAGFGKGHSA | -46.18 | 45.2 | HHHHHHHHHHHHHHHHHHHHHHHHHCCCCCC |
| 877 | EEELRKFLEQYNNEEEDMRYQREGFGKGHQS | -46.18 | 45.2 | HHHHHHHHHHHCHHHHHHHHHHHHHCCCCCC |
| 784 | EEELKKILEKFNNEYEQMFYQQKGFGQGHSA | -46.11 | 48.4 | HHHHHHHHHHHHHHHHHHHHHHHHCCCCCCC |
| 086 | EEELKKILEKFNNEYEQMFYQQQGFGQGHSS | -46.11 | 48.4 | HHHHHHHHHHHHHHHHHHHHHHHHCCCCCCC |

| | | | | |
|---|---|---|---|---|
| 852 | EEELRKILEKINNEEEQIRYQQQGFGQGHSA | -46.11 | 38.7 | HHHHHHHHHHHCHHHHHHHHHHHHCCCCCCC |
| 584 | EEELRKILEKINNEKEQMEYQRNGSGQGHSY | -46.11 | 38.7 | HHHHHHHHHHHHHHHHHHHHHHHCCCCCCCC |
| 716 | EEELRKILEKLNNEKEQMSYQQQGFGQGHSA | -46.11 | 38.7 | HHHHHHHHHHHHHHHHHHHHHHHCCCCCCCC |
| 534 | EQELRKILEKINNEQEQMSYQQQGFGQGHSS | -46.11 | 35.5 | HHHHHHHHHHHHHHHHHHHHHHHCCCCCCCC |
| 791 | EEEWRKFLEQYNNESEDMRYQQYGFGKGHSA | -46.10 | 45.2 | HHHHHHHHHHHHHHHHHHHHHHHCCCCCCCC |
| 887 | EEELKKFLEQYNNEAEDMRYQQQGFGKGHQA | -46.10 | 51.6 | HHHHHHHHHHHHHHHHHHHHHHHHCCCCCCC |
| 315 | EEEKRKFLDKYNQESEQIRYQQQGFGQGHSS | -46.07 | 45.2 | HHHHHHHHHHHHHHHHHHHHHHHHCCCCCCC |
| 666 | EEELRKFLEQFNNEKEQMRYQQQGFGRGHSS | -46.06 | 41.9 | HHHHHHHHHHHHHHHHHHHHHHHHCCCCCCC |
| 985 | EEELKKLLEQINNEAEDERYQQQGFGKGNSN | -46.05 | 48.4 | HHHHHHHHHHHHHHHHHHHHHHHCCCCCCCC |
| 365 | EEELRKLLEQINNEAEDERYQQQGFGKGNSN | -46.05 | 45.2 | HHHHHHHHHHHHHHHHHHHHHHHCCCCCCCC |
| 995 | EEELRKFLEQFNQEYEDMFYQQKGFGKGHSA | -46.05 | 51.6 | HHHHHHHHHHHHHHHHHHHHHHHHCCCCCCC |
| 661 | EEELKKLLEQINQEEEDMRYQQQGFGKGDSR | -46.04 | 51.6 | HHHHHHHHHHHHHHHHHHHHHHHCCCCCCCC |
| 232 | EEELKKLLEQLNQEKEDMRYQREGFGKGDSR | -46.04 | 51.6 | HHHHHHHHHHHHHHHHHHHHHHHHCCCCCCC |
| 353 | EEELRKLLEQINQEKEDEEYQRYGFGKGDSR | -46.04 | 48.4 | HHHHHHHHHHHHHHHHHHHHHHHCCCCCCCC |
| 506 | EEEARKMLEQINNEEEDMRYQREGFGKGHSA | -46.03 | 45.2 | HHHHHHHHHHHHHHHHHHHHHHHHHCCCCCC |
| 347 | EEEARKMLEQINNEQEDMSYQRHGFGKGHSS | -46.03 | 45.2 | HHHHHHHHHHHHHHHHHHHHHHHCCCCCCCC |
| 854 | EEEKKKMLEQINNEAEDMRYQREGFGKGHQS | -46.03 | 48.4 | HHHHHHHHHHHHHHHHHHHHHHHHHCCCCCC |
| 054 | EEEKKKMLEQINNEAEDMRYQRQGFGKGHSA | -46.03 | 48.4 | HHHHHHHHHHHHHHHHHHHHHHHHHCCCCCC |
| 310 | EEEKKKMLEQINNEEEDKRYQREGVGKGHSS | -46.03 | 45.2 | HHHHHHHHHHHHHHHHHHHHHHHHCCCCCCC |
| 831 | EEEKKKMLEQINNEEEDMRYQQQGFGKGHQS | -46.03 | 45.2 | HHHHHHHHHHHHHHHHHHHHHHHHCCCCCCC |
| 713 | EEEKKKMLEQINNEEEDMRYQQQGFGKGHSS | -46.03 | 45.2 | HHHHHHHHHHHHHHHHHHHHHHHHCCCCCCC |
| 042 | EEEKKKMLEQINNEEEDMRYQQYGFGKGHSA | -46.03 | 45.2 | HHHHHHHHHHHHHHHHHHHHHHHCCCCCCCC |
| 549 | EEEKKKMLEQYNNEEEDMRYQREGFGKGHSA | -46.03 | 45.2 | HHHHHHHHHHHCHHHHHHHHHHHHHCCCCCC |
| 902 | EEEKRKMLEQINNEEEDKRYQREGFGKGHSS | -46.03 | 41.9 | HHHHHHHHHHHHHHHHHHHHHHHHCCCCCCC |
| 828 | EEEKKKMLEQINNREEDERYQQYGFGKGHQP | -46.01 | 41.9 | HHHHHHHHHHHHHHHHHHHHHHHCCCCCCCC |
| 085 | EEQLRKFLEQYNNEEEDMRYQREGFGKGHQS | -45.98 | 48.4 | HHHHHHHHHHHHHHHHHHHHHHHHHCCCCCC |
| 449 | EEQLRKFLEQYNNEEEDMRYQRQGFGKGHSA | -45.98 | 48.4 | HHHHHHHHHHHHHHHHHHHHHHHHCCCCCCC |
| 387 | EEELRKFLEQYNQESEDMRYQRYGFGKGDSR | -45.98 | 51.6 | HHHHHHHHHHHHHHHHHHHHHHCCCCCCCCC |
| 179 | EEELKKFLEKFNNEYEDMFYQKEGFGKGDSR | -45.94 | 64.5 | HHHHHHHHHHHHHHHHHHHHHHHCCCCCCCC |
| 287 | EEELKKFLEKYNNEAEDIYYQQQGFGKGDSR | -45.94 | 61.3 | HHHHHHHHHHHHHHHHHHHHHHHCCCCCCCC |
| 802 | EEELKKFLEKYNNEAEDMRYQQQGFGKGDSR | -45.94 | 61.3 | HHHHHHHHHHHHHHHHHHHHHHHCCCCCCCC |
| 173 | EEELKKFLEKYNNESEDMRYQRYGFGKGDSR | -45.94 | 58.1 | HHHHHHHHHHHHHHHHHHHHHECCCCCCCCC |
| 617 | EEELKKFLEKYNNEYEDMEYQRKGFGKGDSR | -45.94 | 58.1 | HHHHHHHHHHHHHHHHHHHHHHHCCCCCCCC |
| 181 | EEELRKFLEKFNNEKEDMRYQQQGFGKGDSR | -45.94 | 58.1 | HHHHHHHHHHHHHHHHHHHHHHHCCCCCCCC |
| 406 | EEELRKFLEKYNNEAEDIRYQQGGFGKGDSR | -45.94 | 58.1 | HHHHHHHHHHHHHHHHHHHHHHCCCCCCCCC |
| 283 | EEELRKFLEKYNNEAEDMRYQRQGFGKGDSR | -45.94 | 58.1 | HHHHHHHHHHHHHHHHHHHHHHHCCCCCCCC |
| 763 | EEELRKFLEKYNNESEDMRYQKEGFGKGDSR | -45.94 | 54.8 | HHHHHHHHHHHHHHHHHHHHHHHCCCCCCCC |
| 717 | EEELRKFLEKYNNEYEDMEYQRKGFGKGDSR | -45.94 | 54.8 | HHHHHHHHHHHHHHHHHHHHHHHCCCCCCCC |

| | | | | |
|---|---|---|---|---|
| 469 | EEELKKFLEKYNNESEDMRYQQQGFGKGDSR | -45.93 | 58.1 | HHHHHHHHHHHHHHHHHHHHHHHCCCCCCCC |
| 988 | EEELKKFLEKYNNEYEDMFYQQKGFGKGDSR | -45.93 | 61.3 | HHHHHHHHHHHHHHHHHHHHHHCCCCCCCCC |
| 614 | EEEWKKFIEKYNNESEDMRYQQQGFGKGDSR | -45.93 | 54.8 | HHHHHHHHHHHHHHHHHHHHHHHCCCCCCCC |
| 382 | EEELRKILDQINQEEEQMRYQQYGFGQGHSS | -45.93 | 38.7 | HHHHHHHHHHHHHHHHHHHHHHHCCCCCCCC |
| 155 | EEELRKFLEKFNNEEEHKRKQREGFGSGKSA | -45.92 | 41.9 | HHHHHHHHHHHCHHHHHHHHHHHHCCCCCCC |
| 156 | EEELRKILEKINNEQEQMSYQQQGFGQGHSA | -45.88 | 38.7 | HHHHHHHHHHHHHHHHHHHHHHHCCCCCCCC |
| 864 | EEELRKILEKLNNEQEQMRYQRQGFGQGHSS | -45.88 | 38.7 | HHHHHHHHHHHHHHHHHHHHHHHHCCCCCCC |
| 543 | EEEKKKMLEQIENEREDEEYQQNGMGKGHQP | -45.87 | 41.9 | HHHHHHHHHHHHHHHHHHHHHHHCCCCCCCC |
| 187 | EEEARKMLDKINNEEEQMRYQREGFGQGHSS | -45.85 | 45.2 | HHHHHHHHHHHHHHHHHHHHHHHHCCCCCCC |
| 850 | EEEKKKMLDKINNEAEQMRYQREGMGQGHQP | -45.85 | 48.4 | HHHHHHHHHHHHHHHHHHHHHHHHHCCCCCC |
| 343 | EEEKRKFLEQFNNEYEDMFYQQKGFGKGHSS | -45.82 | 51.6 | HHHHHHHHHHHHHHHHHHHHHHHHCCCCCCC |
| 898 | EEELKKFLEKYNNEAEDRRYQRYGFGKGHQS | -45.79 | 54.8 | HHHHHHHHHHHHHHHHHHHHHHHCCCCCCCC |
| 705 | EEELKKFLEKYNNESEDRRYQQQGFGKGHSS | -45.79 | 51.6 | HHHHHHHHHHHHHHHHHHHHHHHHCCCCCCC |
| 429 | EEELKKFLEKYNNESEDRRYQRYGFGKGHSA | -45.79 | 51.6 | HHHHHHHHHHHHHHHHHHHHHHHCCCCCCCC |
| 696 | EEELRKFLEKFNNEYEDRFYQQKGFGKGHSS | -45.79 | 54.8 | HHHHHHHHHHHHHHHHHHHHHHHCCCCCCCC |
| 119 | EEELRKFLEKLNNEKEDREYQRAGFGKGHSS | -45.79 | 48.4 | HHHHHHHHHHHHHHHHHHHHHHHHCCCCCCC |
| 153 | EEELRKFLEKYNNEAEDRRYQRQGFGKGHQS | -45.79 | 51.6 | HHHHHHHHHHHHHHHHHHHHHHHHCCCCCCC |
| 875 | EQELRKFLEKYNNEAEDRRYQRYGFGKGHSS | -45.79 | 48.4 | HHHHHHHHHHHHHHHHHHHHHHHCCCCCCCC |
| 776 | EEEKRKMLEQFNNEKEDMRYQQQGFGKGHQA | -45.79 | 45.2 | HHHHHHHHHHHHHHHHHHHHHHHHCCCCCCC |
| 551 | EEELRKFLQQFNNEYEDMFYQQKGFGKGHSS | -45.78 | 51.6 | HHHHHHHHHHHHHHHHHHHHHHHHCCCCCCC |
| 074 | EEELKKLLEQINNEAEHMRKQQQGFGSGKSR | -45.76 | 41.9 | HHHHHHHHHHHHHHHHHHHHHHHCCCCCCCC |
| 036 | EEELKKLLEQINNEAEDMRYQQQGFGKGDSR | -45.74 | 54.8 | HHHHHHHHHHHHHHHHHHHHHHHCCCCCCCC |
| 052 | EEELKKLLEQINNEAEDMRYQRYGFGKGDSR | -45.74 | 54.8 | HHHHHHHHHHHHHHHHHHHHHHHCCCCCCCC |
| 200 | EEELKKLLEQINNEKEDEEYQRYGFGKGDSR | -45.74 | 51.6 | HHHHHHHHHHHHHHHHHHHHHHHCCCCCCCC |
| 147 | EEELKKLLEQLNNEKEDMRYQREGFGKGDSR | -45.74 | 51.6 | HHHHHHHHHHHHHHHHHHHHHHHHCCCCCCC |
| 990 | EEELKKLLEQLNNEKEDMRYQRQGFGKGDSR | -45.74 | 51.6 | HHHHHHHHHHHHHHHHHHHHHHHCCCCCCCC |
| 769 | EEELRKLLEQINNEEEDMRYQQQGFGKGDSR | -45.74 | 48.4 | HHHHHHHHHHHHHHHHHHHHHHHCCCCCCCC |
| 193 | EEELRKLLEQINNEEEDMRYQREGFGKGDSR | -45.74 | 48.4 | HHHHHHHHHHHHHHHHHHHHHHHHCCCCCCC |
| 983 | EEELRKLLEQINNEKEDMEYQRAGFGKGDSR | -45.74 | 48.4 | HHHHHHHHHHHHHHHHHHHHHHHHCCCCCCC |
| 788 | EEELRKLLEQLNNEKEDMRYQQQGFGKGDSR | -45.74 | 48.4 | HHHHHHHHHHHHHHHHHHHHHHHCCCCCCCC |
| 774 | EQELRKLLEQINNEAEDERYQQQGFGKGDSR | -45.74 | 48.4 | HHHHHHHHHHHHHHHHHHHHHHHCCCCCCCC |
| 208 | EEEKRKFLEQYNNEAEDMRYQQYGFGKGDSR | -45.69 | 54.8 | HHHHHHHHHHHHHHHHHHHHHHHCCCCCCCC |
| 595 | EEELKKFLEQFNNEYEDMFYQQKGFGKGDSR | -45.69 | 61.3 | HHHHHHHHHHHHHHHHHHHHHHHCCCCCCCC |
| 688 | EEELKKFLEQFNNEYEDMYYQQQGFGKGDSR | -45.69 | 58.1 | HHHHHHHHHHHHHHHHHHHHHHHCCCCCCCC |
| 009 | EEELKKFLEQYNNEAEDIYYQQNGFGKGDSR | -45.69 | 58.1 | HHHHHHHHHHHHHHHHHHHHHHHCCCCCCCC |
| 093 | EEELKKFLEQYNNESEDERYQQNGFGKGDSR | -45.69 | 54.8 | HHHHHHHHHHHHCCHHHHHHHHHCCCCCCCC |
| 540 | EEELRKFLEQFNNEYEDMFYQQKGFGKGDSR | -45.69 | 58.1 | HHHHHHHHHHHHHHHHHHHHHHHCCCCCCCC |
| 065 | EEELRKFLEQLNNEKEDMEYQRAGFGKGDSR | -45.69 | 51.6 | HHHHHHHHHHHHHHHHHHHHHHHHCCCCCCC |

| | | | | |
|---|---|---|---|---|
| 178 | EEELRKFLEQYNNEEEDMRYQREGFGKGDSR | -45.69 | 51.6 | HHHHHHHHHHCHHHHHHHHHHHHHCCCCCCC |
| 755 | EEELRKFLEQYNNEYEDMFYQQKGFGKGDSR | -45.69 | 54.8 | HHHHHHHHHHHHHHHHHHHHHHCCCCCCCCC |
| 805 | EQELRKFLEQFNNEYEDEFYQQKGFGKGDSR | -45.69 | 54.8 | HHHHHHHHHHHHHHHHHHHHHHHCCCCCCCC |
| 474 | EQELRKILEKLNNEKEQMSYQQQGFGQGHQS | -45.65 | 35.5 | HHHHHHHHHHHHHHHHHHHHHHHCCCCCCCC |
| 186 | EEELKKILEQLNNEQEDMRYQRQGFGKGHSS | -45.63 | 45.2 | HHHHHHHHHHHHHHHHHHHHHHHHCCCCCCC |
| 924 | EEELKKILEKLNNEKEQMRYQQQGFGQGHSA | -45.63 | 41.9 | HHHHHHHHHHHHHHHHHHHHHHHHCCCCCCC |
| 278 | EEEKRKMLEQINQEEEDQRYQQQGFGKGDSR | -45.59 | 48.4 | HHHHHHHHHHHHHHHHHHHHHHHCCCCCCCC |
| 728 | EEELRKFLEKFNNEYEDRFYQQKGFGKGHSA | -45.56 | 54.8 | HHHHHHHHHHHHHHHHHHHHHHHHCCCCCCC |
| 768 | EEELRKFLDQYNNEEEQMRYQREGFGQGHSS | -45.54 | 41.9 | HHHHHHHHHHCHHHHHHHHHHHHHCCCCCCC |
| 823 | EEELKKFLDQYNQEEEQMRFQREGFGQGKSS | -45.53 | 41.9 | HHHHHHHHHHHHHHHHHHHHHHHHCCCCCCC |
| 670 | EEEARKFLDQYNNEYEQMYYQQQGFGQGHSS | -45.52 | 45.2 | HHHHHHHHHHHHHHHHHHHHHHHCCCCCCCC |
| 622 | EEELRKFLDQYNNEAEQIYYQQNGFGQGHSA | -45.52 | 45.2 | HHHHHHHHHHHHHHHHHHHHHHHCCCCCCCC |
| 421 | EEEARKMLQQINNEAEDRRYQREGFGKGHSA | -45.50 | 48.4 | HHHHHHHHHHHHHHHHHHHHHHHHCCCCCCC |
| 836 | EEELRKFLEQYNNEEEDMRYQREGFGKGNSN | -45.48 | 45.2 | HHHHHHHHHHCHHHHHHHHHHHHHCCCCCCC |
| 921 | EEELKKFLEQYNQEEEDMRYQRQGFGKGDSR | -45.47 | 54.8 | HHHHHHHHHHHHHHHHHHHHHHHCCCCCCCC |
| 532 | EEELRKFLEQFNQEYEDMFYQQKGFGKGDSR | -45.47 | 58.1 | HHHHHHHHHHHHHHHHHHHHHHHCCCCCCCC |
| 558 | EEELRKFLEQYNQESEDMRYQQQGFGKGDSR | -45.47 | 51.6 | HHHHHHHHHHHHHHHHHHHHHHHCCCCCCCC |
| 813 | EEELRKFLEQYNQEYEDEEYQRKGFGKGDSR | -45.47 | 51.6 | HHHHHHHHHHHHHHHHHHHHHHHCCCCCCCC |
| 853 | EEELKKFLDSLNNREEDMRYQRYGFGKGDSR | -45.40 | 54.8 | HHHHHHHHHHHHHHHHHHHHHHHCCCCCCCC |
| 423 | EEELRKILEQINQEEEQMRYQQQGFGQGHQS | -45.40 | 35.5 | HHHHHHHHHHHHHHHHHHHHHHHCCCCCCCC |
| 539 | EEELKKLLEKINQEAEQMRYQKEGFGQGHSA | -45.39 | 45.2 | HHHHHHHHHHHHHHHHHHHHHHHHCCCCCCC |
| 796 | EEELKKLLEKLNQEKEQMRYQRYGFGQGHSA | -45.39 | 41.9 | HHHHHHHHHHHHHHHHHHHHHHHCCCCCCCC |
| 379 | EEELRKLLEKLNQEKEQKRYQRQGFGQGHSA | -45.39 | 38.7 | HHHHHHHHHHHHHHHHHHHHHHHCCCCCCCC |
| 230 | EEEWKKLMEKYNQESEQMRYQRYGFGQGHSA | -45.39 | 38.7 | HHHHHHHHHHHHHHHHHHHHHHHCCCCCCCC |
| 255 | EQELRKLLEKINQEQEQKSYQQQGFGQGHQS | -45.39 | 35.5 | HHHHHHHHHHHHHHHHHHHHHHHCCCCCCCC |
| 392 | EEELRKLLEKLNNEKEQMSFQQQGFGQGKSR | -45.38 | 38.7 | HHHHHHHHHHHHHHHHHHHHHHHHCCCCCCC |
| 184 | EEELKKILEKINNEEEQMRYQQYGFGQGDSR | -45.38 | 48.4 | HHHHHHHHHHCHHHHHHHHHHHHCCCCCCCC |
| 694 | EEELRKILEKLNNEKEQMRYQRYGFGQGDSR | -45.35 | 45.2 | HHHHHHHHHHHHHHHHHHHHHHHCCCCCCCC |
| 710 | EEELRKFLEKYNQESEDRRYQRYGFGKGDSR | -45.35 | 54.8 | HHHHHHHHHHHHHHHHHHHHHHCCCCCCCCC |
| 732 | EEEKRKMLEQINNEKEDMEYQRAGFGKGDSR | -45.29 | 48.4 | HHHHHHHHHHHHHHHHHHHHHHHHCCCCCCC |
| 356 | EEQKRKMLEQINNEAEQMRYQRQGFGRGHSS | -45.23 | 41.9 | HHHHHHHHHHHHHHHHHHHHHHHHCCCCCCC |
| 521 | EEELRKILEKINNEEEQMRYQREGFGQGHSA | -45.21 | 38.7 | HHHHHHHHHHCHHHHHHHHHHHHHHCCCCCC |
| 370 | EEEKRKFLEQYNNEAEHMRKQQQGFGSGKSR | -45.19 | 41.9 | HHHHHHHHHHHHHHHHHHHHHHHCCCCCCCC |
| 712 | EEEKRKFLEQYNNESEDMRYQQQGFGKGDSR | -45.18 | 51.6 | HHHHHHHHHHHHHHHHHHHHHHHCCCCCCCC |
| 453 | EEELKKFLEQLNNEEEDMRYQQYGFGKGDSR | -45.18 | 54.8 | HHHHHHHHHHHHHHHHHHHHHHHCCCCCCCC |
| 484 | EEELKKFLEQYNNEAEDMRYQQQGFGKGDSR | -45.18 | 58.1 | HHHHHHHHHHHHHHHHHHHHHHHCCCCCCCC |
| 486 | EEELKKFLEQYNNEAEDMRYQQYGFGKGDSR | -45.18 | 58.1 | HHHHHHHHHHHHHHHHHHHHHHHCCCCCCCC |
| 087 | EEELKKFLEQYNNESEDMRYQQQGFGKGDSR | -45.18 | 54.8 | HHHHHHHHHHHHHHHHHHHHHHHCCCCCCCC |

| | | | | |
|---|---|---|---|---|
| 943 | EEELKKFLEQYNNEYEDMFYQQKGFGKGDSR | -45.18 | 58.1 | HHHHHHHHHHHHHHHHHHHHHHHCCCCCCCCC |
| 191 | EEELRKFLEQFNNEYEDEEYQRKGFGKGDSR | -45.18 | 54.8 | HHHHHHHHHHHHHHHHHHHHHHHHCCCCCCCC |
| 778 | EEELRKFLEQYNNEAEDMRYQQQGFGKGDSR | -45.18 | 54.8 | HHHHHHHHHHHHHHHHHHHHHHHHCCCCCCCC |
| 294 | EEELRKFLEQYNNESEDERYQQQGFGKGDSR | -45.18 | 51.6 | HHHHHHHHHHHHHCHHHHHHHHHHCCCCCCCC |
| 400 | EEELRKFLEQYNNESEDMRYQQQGFGKGDSR | -45.18 | 51.6 | HHHHHHHHHHHHHHHHHHHHHHHHCCCCCCCC |
| 084 | EEELRKFLEQYNNEYEDEEYQRKGFGKGDSR | -45.18 | 51.6 | HHHHHHHHHHHHHHHHHHHHHHHHCCCCCCCC |
| 280 | EEELRKFLEQYNNEYEDEEYQRQGFGKGDSR | -45.18 | 51.6 | HHHHHHHHHHHHHHHHHHHHHHHHCCCCCCCC |
| 253 | EEELRKFLEQYNNEYEDMYYQQQGFGKGDSR | -45.18 | 51.6 | HHHHHHHHHHHHHHHHHHHHHHHHCCCCCCCC |
| 794 | EEELKKLLDQINQEKEQMEYQRAGFGQGDSR | -45.15 | 48.4 | HHHHHHHHHHHHHHHHHHHHHHHHHCCCCCCC |
| 930 | EEELKKLLDDINNRAEDMRYQKEGFGKGHSS | -45.15 | 48.4 | HHHHHHHHHHHHHHHHHHHHHHHHHCCCCCCC |
| 668 | EEELKKILEQFNNEYEQMFYQQKGFGQGHSA | -45.12 | 45.2 | HHHHHHHHHHHHHHHHHHHHHHHHHCCCCCCC |
| 651 | EEELKKILEQLNNEKEQMRYQQQGFGQGHQS | -45.12 | 38.7 | HHHHHHHHHHHHHHHHHHHHHHHHHCCCCCCC |
| 137 | EEELRKILEQYNNEYEQMEYQRNGRGQGHSY | -45.12 | 35.5 | HHHHHHHHHHHHHHHHHHHHHHHHCCCCCCCC |
| 357 | EEELKKLLEKINNEAEDRRYQRQGFGKGDSR | -45.11 | 58.1 | HHHHHHHHHHHHHHHHHHHHHHHHCCCCCCCC |
| 160 | EEELRKLLEKINNEAEDRRYQRYGFGKGDSR | -45.11 | 54.8 | HHHHHHHHHHHHHHHHHHHHHHHCCCCCCCCC |
| 097 | EEELRKLLEKINNEEEDRRYQREGFGKGDSR | -45.11 | 51.6 | HHHHHHHHHHHHHHHHHHHHHHHHHCCCCCCC |
| 939 | EEELKKILEQINNEEEQMRYQQYGFGQGHQS | -45.11 | 38.7 | HHHHHHHHHHHCHHHHHHHHHHHHCCCCCCCC |
| 116 | EEELKKILEQINNEEEQMRYQQYGFGQGHSA | -45.11 | 38.7 | HHHHHHHHHHHCHHHHHHHHHHHHCCCCCCCC |
| 678 | EEELRKILEQINNEKEQMEYQKQGFGQGHSA | -45.11 | 35.5 | HHHHHHHHHHHHHHHHHHHHHHHHHCCCCCCC |
| 950 | EEELRKILEQLNNEKEQMRYQQYGFGQGHSA | -45.11 | 35.5 | HHHHHHHHHHHHHHHHHHHHHHHHCCCCCCCC |
| 144 | EEEWRKILEQINNEKEQMRYQQQGFGQGHSS | -45.11 | 35.5 | HHHHHHHHHHHHHHHHHHHHHHHHCCCCCCCC |
| 533 | EEELRKLLEQLNNDKEQMRYQREGFGQGHQS | -45.10 | 32.3 | HHHHHHHHHHHHHHHHHHHHHHHHHHCCCCCC |
| 573 | EEEWKKLMEQFNNDYEQMYYQQKGFGQGHQS | -45.10 | 35.5 | HHHHHHHHHHHHHHHHHHHHHHHHHCCCCCCC |
| 364 | EEELKKLLEKINNEAEQIRYQQQGFGQGHSS | -45.09 | 45.2 | HHHHHHHHHHHHHHHHHHHHHHHHHCCCCCCC |
| 504 | EEELKKLLEKINNEAEQKRYQRQGFGQGHSA | -45.09 | 45.2 | HHHHHHHHHHHHHHHHHHHHHHHHCCCCCCCC |
| 722 | EEELKKLLEKINNEAEQMRYQREGMGQGHQP | -45.09 | 45.2 | HHHHHHHHHHHHHHHHHHHHHHHHHHCCCCCC |
| 579 | EEELKKLLEKINNEAEQMRYQRYGFGQGHSS | -45.09 | 45.2 | HHHHHHHHHHHHHHHHHHHHHHHHCCCCCCCC |
| 888 | EEELKKLLEKINNEAEQMSYQQQGFGQGHSA | -45.09 | 45.2 | HHHHHHHHHHHHHHHHHHHHHHHHCCCCCCCC |
| 372 | EEELKKLLEKINNEEEQIRYQQQGFGQGHSS | -45.09 | 41.9 | HHHHHHHHHHHCHHHHHHHHHHHHHCCCCCCC |
| 519 | EEELKKLLEKINNEEEQKRYQREGFGQGHSA | -45.09 | 41.9 | HHHHHHHHHHHCHHHHHHHHHHHHHCCCCCCC |
| 807 | EEELKKLLEKINNEEEQMRYQREGFGQGHSS | -45.09 | 41.9 | HHHHHHHHHHHCHHHHHHHHHHHHHCCCCCCC |
| 876 | EEELKKLLEKINNEEEQMRYQRYGFGQGHSA | -45.09 | 41.9 | HHHHHHHHHHHCHHHHHHHHHHHHCCCCCCCC |
| 967 | EEELKKLLEKINNEKEQMEYQRAGFGQGHQA | -45.09 | 41.9 | HHHHHHHHHHHHHHHHHHHHHHHHHHCCCCCC |
| 785 | EEELKKLLEKINNEKEQMEYQRRGMGQGHQP | -45.09 | 41.9 | HHHHHHHHHHHHHHHHHHHHHHHHHHCCCCCC |
| 385 | EEELKKLLEKINNEKEQMEYQRYGFGQGHSS | -45.09 | 41.9 | HHHHHHHHHHHHHHHHHHHHHHHHCCCCCCCC |
| 260 | EEELKKLLEKLNNEKEQMRYQQQGFGQGHSS | -45.09 | 41.9 | HHHHHHHHHHHHHHHHHHHHHHHHCCCCCCCC |
| 129 | EEELRKLLEKINNEAEQIRYQQQGFGQGHSS | -45.09 | 41.9 | HHHHHHHHHHHHHHHHHHHHHHHHHCCCCCCC |
| 470 | EEELRKLLEKINNEAEQMRYQQYGFGQGHSS | -45.09 | 41.9 | HHHHHHHHHHHHHHHHHHHHHHHHCCCCCCCC |

| | | | | |
|---|---|---|---|---|
| 322 | EEELRKLLEKINNEEEQKRYQREGFGQGHSA | -45.09 | 38.7 | HHHHHHHHHHHCHHHHHHHHHHHHCCCCCCC |
| 464 | EEELRKLLEKINNEEEQMRYQQYGFGQGHQS | -45.09 | 38.7 | HHHHHHHHHHHCHHHHHHHHHHHCCCCCCCC |
| 476 | EEELRKLLEKINNEKEQMEYQRAGFGQGHSA | -45.09 | 38.7 | HHHHHHHHHHHHHHHHHHHHHHHHHCCCCCC |
| 693 | EEELRKLLEKINNEQEQKSYQQQGFGQGHSA | -45.09 | 38.7 | HHHHHHHHHHHHHHHHHHHHHHHCCCCCCCC |
| 905 | EEELRKLLEKINNEQEQKSYQQQGFGQGHSS | -45.09 | 38.7 | HHHHHHHHHHHHHHHHHHHHHHHCCCCCCCC |
| 934 | EEELRKLLEKLNNEKEQIRYQQQGFGQGHSS | -45.09 | 38.7 | HHHHHHHHHHHHHHHHHHHHHHHHCCCCCCC |
| 780 | EEELRKLLEKLNNEKEQMSYQQQGFGQGHQP | -45.09 | 38.7 | HHHHHHHHHHHHHHHHHHHHHHHHCCCCCCC |
| 222 | EEELRKFLDQYNNEYEQEEYQRKGFGQGHSA | -45.05 | 41.9 | HHHHHHHHHHHHHHHHHHHHHHHCCCCCCCC |
| 987 | EEELKKILEQLNQEKEQMRYQRQGFGQGHSA | -44.93 | 38.7 | HHHHHHHHHHHHHHHHHHHHHHHHCCCCCCC |
| 971 | EEELRKLLEKLNNEKEQMSYQQQGFGQGHSA | -44.86 | 38.7 | HHHHHHHHHHHHHHHHHHHHHHHCCCCCCCC |
| 522 | EEELKKFLEKYNQEAEQIYYQQQGFGQGHSS | -44.82 | 48.4 | HHHHHHHHHHHHHHHHHHHHHHHHCCCCCCC |
| 003 | EEELKKFLEKYNQEEEQMRYQREGFGQGHSS | -44.82 | 45.2 | HHHHHHHHHHHCHHHHHHHHHHHHCCCCCCC |
| 309 | EEELRKFLEKFNQEYEQMFYQQKGFGQGHSA | -44.82 | 48.4 | HHHHHHHHHHHHHHHHHHHHHHHHCCCCCCC |
| 170 | EEELRKFLEKYNQEEEQMSYQRYGFGQGHSS | -44.82 | 41.9 | HHHHHHHHHHHHHHHHHHHHHHHCCCCCCCC |
| 369 | EEELRKFLEKFNNEYEQIFFQQKGFGQGKSR | -44.82 | 48.4 | HHHHHHHHHHHHHHHHHHHHHHHCCCCCCCC |
| 719 | EEELRKFLEKYNNEEEQMSFQKEGFGQGKSR | -44.82 | 41.9 | HHHHHHHHHHHCHHHHHHHHHHHHCCCCCCC |
| 471 | EEELRKLLEKLNNEQEQMRYQQQGFGQGHQA | -44.80 | 38.7 | HHHHHHHHHHHHHHHHHHHHHHHHCCCCCCC |
| 059 | EEEKKKMLEQINNEAEDMRYQQQGFGKGDSR | -44.78 | 54.8 | HHHHHHHHHHHHHHHHHHHHHHHCCCCCCCC |
| 772 | EEEKRKMLEQINNEAEDMRYQRYGFGKGDSR | -44.78 | 51.6 | HHHHHHHHHHHHHHHHHHHHHHHCCCCCCCC |
| 441 | EEEKRKMLEQINNEQEDEEYQRKGFGKGDSR | -44.78 | 48.4 | HHHHHHHHHHHHHHHHHHHHHHHCCCCCCCC |
| 141 | EEELKKILEQLNNEKEQKRYQKEGFGQGHSA | -44.78 | 38.7 | HHHHHHHHHHHHHHHHHHHHHHHHCCCCCCC |
| 738 | EEELKKILEQLNNEQEQERYQQQGFGQGHSS | -44.78 | 38.7 | HHHHHHHHHHHHHHHHHHHHHHHCCCCCCCC |
| 455 | EEELKKLLEKINNEQEQMSYQQQGFGQGHQA | -44.77 | 41.9 | HHHHHHHHHHHHHHHHHHHHHHHHCCCCCCC |
| 847 | EQELRKLLEKINQEEEQIRYQQQGFGQGDSR | -44.66 | 41.9 | HHHHHHHHHHHHHHHHHHHHHHHCCCCCCCC |
| 839 | EEEARKMLEKINNEEEDMRYQREGFGKGHSA | -44.64 | 48.4 | HHHHHHHHHHHHHHHHHHHHHHHHHCCCCCC |
| 359 | EEELKKLLEQINQEQEQMYYQQKGFGQGHQS | -44.62 | 38.7 | HHHHHHHHHHHHHHHHHHHHHHHHCCCCCCC |
| 219 | EEELRKLLEQINQEAEQEFYQQKGFGQGHQS | -44.62 | 41.9 | HHHHHHHHHHHHHHHHHHHHHHHCCCCCCCC |
| 090 | EEELRKLLEQINQEAEQIRYQQQGFGQGHSS | -44.62 | 38.7 | HHHHHHHHHHHHHHHHHHHHHHHCCCCCCCC |
| 676 | EEELRKLLEQINQEEEQMRYQQQGFGQGHSS | -44.62 | 35.5 | HHHHHHHHHHHHHHHHHHHHHHHCCCCCCCC |
| 517 | EEELRKLLEQINQEEEQMRYQQYGFGQGHSS | -44.62 | 35.5 | HHHHHHHHHHHHHHHHHHHHHHHCCCCCCCC |
| 358 | EEELRKLLEQLNQEEEQMKYQRYGFGQGHSS | -44.62 | 35.5 | HHHHHHHHHHHHHHHHHHHHHHHCCCCCCCC |
| 745 | EEELKKLLEKINNEAEQIRYQQQGFGQGHSA | -44.62 | 45.2 | HHHHHHHHHHHHHHHHHHHHHHHHCCCCCCC |
| 405 | EEELKKLLEKLNNEQEQMRYQQNGFGQGHQS | -44.62 | 41.9 | HHHHHHHHHHHHHHHHHHHHHHHHCCCCCCC |
| 146 | EEELRKLLEKINNEEEQMFYQQKGFGQGHSS | -44.62 | 41.9 | HHHHHHHHHHHHHHHHHHHHHHHHCCCCCCC |
| 111 | EEELRKLLEKINNEKEQMEYQKQGFGQGHSS | -44.62 | 38.7 | HHHHHHHHHHHHHHHHHHHHHHHCCCCCCCC |
| 870 | EEELRKLLEKLNNEKEQMRYQQYGFGQGHSA | -44.62 | 38.7 | HHHHHHHHHHHHHHHHHHHHHHHCCCCCCCC |
| 394 | EEELKKLLEQINNEEEQMRFQQQGFGQGKSR | -44.62 | 38.7 | HHHHHHHHHHHCHHHHHHHHHHHHCCCCCCC |
| 018 | EEEKRKFLDQFNQEYEQMFYQQKGFGQGDSR | -44.59 | 54.8 | HHHHHHHHHHHHHHHHHHHHHHHCCCCCCCC |

| | | | | |
|---|---|---|---|---|
| 509 | EEELRKILEQINNEAEQMFYQQKGFGQGHSS | -44.59 | 41.9 | HHHHHHHHHHHHHHHHHHHHHHHHCCCCCCC |
| 499 | EEELRRLLEQINNEAEDMRYQQQGFGKGHQS | -44.58 | 45.2 | HHHHHHHHHHHHHHHHHHHHHHHHCCCCCCC |
| 773 | EQELRKFLEKFNNEYEDRFYQQKGFGKGDSR | -44.55 | 58.1 | HHHHHHHHHHHHHHHHHHHHHHHCCCCCCCC |
| 565 | EEELKKFLEKFNNEEEQMSYQRQGFGQGHSA | -44.53 | 48.4 | HHHHHHHHHHHCHHHHHHHHHHHCCCCCCCC |
| 221 | EEELKKFLEKFNNEYEQMYYQQKGFGQGHSS | -44.53 | 48.4 | HHHHHHHHHHHHHHHHHHHHHHHHCCCCCCC |
| 189 | EEELKKFLEKYNNEAEQIRYQQQGFGQGHSA | -44.53 | 48.4 | HHHHHHHHHHHHHHHHHHHHHHHHCCCCCCC |
| 744 | EEELKKFLEKYNNEAEQKRYQQQGFGQGHSS | -44.53 | 48.4 | HHHHHHHHHHHHHHHHHHHHHHHCCCCCCCC |
| 918 | EEELKKFLEKYNNEAEQMRYQQQGFGQGHSS | -44.53 | 48.4 | HHHHHHHHHHHHHHHHHHHHHHHCCCCCCCC |
| 408 | EEELKKFLEKYNNEEEQMSYQKEGFGQGHSA | -44.53 | 45.2 | HHHHHHHHHHHCHHHHHHHHHHHHCCCCCCC |
| 296 | EEELKKFLEKYNNEYEQKFYQQKGFGQGHQS | -44.53 | 48.4 | HHHHHHHHHHHHHHHHHHHHHHHCCCCCCCC |
| 012 | EEELKKFLEKYNNEYEQMEYQRTGRGQGHSF | -44.53 | 45.2 | HHHHHHHHHHHHHHHHHHHHHHHCCCCCCCC |
| 984 | EEELKKFLEKYNNEYEQMQYQKEGFGQGHSS | -44.53 | 45.2 | HHHHHHHHHHHHHHHHHHHHHHHHCCCCCCC |
| 245 | EEELRKFLEKFNNEYEQMFYQQKGFGQGHSA | -44.53 | 48.4 | HHHHHHHHHHHHHHHHHHHHHHHHCCCCCCC |
| 510 | EEELRKFLEKFNNEYEQMFYQQKGFGQGHSS | -44.53 | 48.4 | HHHHHHHHHHHHHHHHHHHHHHHHCCCCCCC |
| 004 | EEELRKFLEKFNNEYEQMYYQQQGFGQGHSA | -44.53 | 45.2 | HHHHHHHHHHHHHHHHHHHHHHHCCCCCCCC |
| 033 | EEELRKFLEKYNNEAEQIRYQQQGMGQGHQP | -44.53 | 45.2 | HHHHHHHHHHHHHHHHHHHHHHHHCCCCCCC |
| 247 | EEELRKFLEKYNNEAEQMRYQKEGFGQGHQS | -44.53 | 45.2 | HHHHHHHHHHHHHHHHHHHHHHHHCCCCCCC |
| 456 | EEELRKFLEKYNNEAEQMRYQQQGFGQGHQS | -44.53 | 45.2 | HHHHHHHHHHHHHHHHHHHHHHHCCCCCCCC |
| 149 | EEELRKFLEKYNNEAEQMRYQRQGFGQGHSA | -44.53 | 45.2 | HHHHHHHHHHHHHHHHHHHHHHHHCCCCCCC |
| 473 | EEELRKFLEKYNNEEEQMRYQREGFGQGHQS | -44.53 | 41.9 | HHHHHHHHHHHCCHHHHHHHHHHHHCCCCCC |
| 814 | EEELRKFLEKYNNEEEQMRYQREGKGQGHSH | -44.53 | 41.9 | HHHHHHHHHHHCCHHHHHHHHHHHHCCCCCC |
| 266 | EEELRKFLEKYNNEEEQMRYQRQGFGQGHSS | -44.53 | 41.9 | HHHHHHHHHHHCCHHHHHHHHHHHHCCCCCC |
| 626 | EEELRKFLEKYNNESEQIRYQQQGFGQGHSA | -44.53 | 41.9 | HHHHHHHHHHHHHHHHHHHHHHHHCCCCCCC |
| 227 | EEELRKFLEKYNNESEQMRYQQQGFGQGHSS | -44.53 | 41.9 | HHHHHHHHHHHHHHHHHHHHHHHHCCCCCCC |
| 867 | EEELRKFLEKYNNEYEQMEYQRKGFGQGHQS | -44.53 | 41.9 | HHHHHHHHHHHHHHHHHHHHHHHHCCCCCCC |
| 380 | EEELRKLLEKINQEEEQMRYQQQGFGQGHSS | -44.49 | 38.7 | HHHHHHHHHHHHHHHHHHHHHHHCCCCCCCC |
| 011 | EEEKKKMLEKINNEEEQMRFQQYGFGQGKSR | -44.43 | 41.9 | HHHHHHHHHHHCHHHHHHHHHHHCCCCCCCC |
| 703 | EEELRKILEQINNREEQERYQQYGFGQGDSR | -44.41 | 38.7 | HHHHHHHHHHHHHHHHHHHHHHHCCCCCCCC |
| 479 | EEQLRKFLEKYNQEEEQMRYQREGFGQGHQS | -44.38 | 45.2 | HHHHHHHHHHHHHHHHHHHHHHHHHCCCCCC |
| 333 | EEELKKLLEKINNEAEQMSYQRHGFGQGHSS | -44.36 | 45.2 | HHHHHHHHHHHHHHHHHHHHHHHCCCCCCCC |
| 518 | EEELKKLLEKINNEKEQMEYQKQGFGQGDSR | -44.36 | 48.4 | HHHHHHHHHHHHHHHHHHHHHHHCCCCCCCC |
| 477 | EEELRKLLEKINNEEEQMRYQQQGFGQGDSR | -44.36 | 45.2 | HHHHHHHHHHHCHHHHHHHHHHHCCCCCCCC |
| 217 | EEELRKLLEKINNEEEQMRYQRYGFGQGDSR | -44.36 | 45.2 | HHHHHHHHHHHCHHHHHHHHHHHCCCCCCCC |
| 586 | EEELRKFLEKFNQEEEQMSYQRHGFGQGHSS | -44.35 | 45.2 | HHHHHHHHHHHHHHHHHHHHHHHCCCCCCCC |
| 291 | EEELKKLLEQINNEAEQERYQQQGFGQGHQS | -44.32 | 41.9 | HHHHHHHHHHHHHHHHHHHHHHHCCCCCCCC |
| 894 | EEELKKLLEQINNEAEQERYQQQGFGQGHSS | -44.32 | 41.9 | HHHHHHHHHHHHHHHHHHHHHHHCCCCCCCC |
| 246 | EEELKKLLEQINNEAEQMRYQQQGFGQGHSA | -44.32 | 41.9 | HHHHHHHHHHHHHHHHHHHHHHHCCCCCCCC |
| 925 | EEELKKLLEQINNEAEQMRYQRYGFGQGHSA | -44.32 | 41.9 | HHHHHHHHHHHHHHHHHHHHHHHCCCCCCCC |

| | | | | |
|---|---|---|---|---|
| 929 | EEELKKLLEQINNEAEQMRYQRYGFGQGHSS | -44.32 | 41.9 | HHHHHHHHHHHHHHHHHHHHHHHHCCCCCCCC |
| 354 | EEELKKLLEQINNEEEQMRYQQYGFGQGHSS | -44.32 | 38.7 | HHHHHHHHHHHCHHHHHHHHHHHHCCCCCCCC |
| 680 | EEELKKLLEQINNEEEQMRYQREGFGQGHSA | -44.32 | 38.7 | HHHHHHHHHHHCHHHHHHHHHHHHHHCCCCCC |
| 213 | EEELKKLLEQINNEQEQESYQQQGFGQGHQS | -44.32 | 38.7 | HHHHHHHHHHHHHHHHHHHHHHHHCCCCCCCC |
| 754 | EEELKKLLEQINNEQEQMSYQQQGFGQGHSS | -44.32 | 38.7 | HHHHHHHHHHHHHHHHHHHHHHHHCCCCCCCC |
| 489 | EEELKKLLEQLNNEKEQMRYQQQGFGQGHQA | -44.32 | 38.7 | HHHHHHHHHHHHHHHHHHHHHHHHHCCCCCCC |
| 981 | EEELKKLLEQLNNEKEQMRYQRQGFGQGHSA | -44.32 | 38.7 | HHHHHHHHHHHHHHHHHHHHHHHHHCCCCCCC |
| 331 | EEELKKLLEQLNNEKEQMRYQRYGFGQGHQS | -44.32 | 38.7 | HHHHHHHHHHHHHHHHHHHHHHHHCCCCCCCC |
| 030 | EEELRKLLEQINNEAEQMRYQQQGFGQGHSA | -44.32 | 38.7 | HHHHHHHHHHHHHHHHHHHHHHHHCCCCCCCC |
| 306 | EEELRKLLEQINNEAEQMRYQRYGFGQGHQS | -44.32 | 38.7 | HHHHHHHHHHHHHHHHHHHHHHHHCCCCCCCC |
| 161 | EEELRKLLEQINNEEEQERYQQQGFGQGHSA | -44.32 | 35.5 | HHHHHHHHHHHCHHHHHHHHHHHHCCCCCCCC |
| 993 | EEELRKLLEQINNEEEQERYQQQGFGQGHSS | -44.32 | 35.5 | HHHHHHHHHHHCHHHHHHHHHHHHCCCCCCCC |
| 297 | EEELRKLLEQINNEEEQMRYQQQGFGQGHQS | -44.32 | 35.5 | HHHHHHHHHHHCHHHHHHHHHHHHCCCCCCCC |
| 741 | EEELRKLLEQINNEEEQMRYQQQGFGQGHSA | -44.32 | 35.5 | HHHHHHHHHHHCHHHHHHHHHHHHCCCCCCCC |
| 891 | EEELRKLLEQINNEEEQMRYQQQGFGQGHSP | -44.32 | 35.5 | HHHHHHHHHHHCHHHHHHHHHHHHHCCCCCCC |
| 053 | EEELRKLLEQINNEEEQMRYQQQGFGQGHSS | -44.32 | 35.5 | HHHHHHHHHHHCHHHHHHHHHHHHCCCCCCCC |
| 307 | EEELRKLLEQINNEEEQMRYQQYGFGQGHQS | -44.32 | 35.5 | HHHHHHHHHHHCHHHHHHHHHHHHCCCCCCCC |
| 304 | EEELRKLLEQINNEEEQMRYQQYGFGQGHSA | -44.32 | 35.5 | HHHHHHHHHHHCHHHHHHHHHHHHCCCCCCCC |
| 821 | EEELRKLLEQINNEEEQMRYQQYGFGQGHSS | -44.32 | 35.5 | HHHHHHHHHHHCHHHHHHHHHHHHCCCCCCCC |
| 352 | EEELRKLLEQINNEEEQMRYQREGFGQGHQS | -44.32 | 35.5 | HHHHHHHHHHHCHHHHHHHHHHHHHHCCCCCC |
| 766 | EEELRKLLEQINNEKEQEEYQRKGFGQGHSS | -44.32 | 35.5 | HHHHHHHHHHHHHHHHHHHHHHHHCCCCCCCC |
| 492 | EEELRKLLEQINNEQEQMSYQQQGFGQGHQS | -44.32 | 35.5 | HHHHHHHHHHHHHHHHHHHHHHHHCCCCCCCC |
| 633 | EEELRKLLEQLNNEKEQMRYQRQGFGQGHSA | -44.32 | 35.5 | HHHHHHHHHHHHHHHHHHHHHHHHHCCCCCCC |
| 288 | EEELRKLLEQLNNEKEQMRYQRYGFGQGHSS | -44.32 | 35.5 | HHHHHHHHHHHHHHHHHHHHHHHHCCCCCCCC |
| 340 | EEELKKFLDQYNNESEQMRYQQQGFGQGDSR | -44.30 | 51.6 | HHHHHHHHHHHHHHHHHHHHHHHHCCCCCCCC |
| 268 | EEELRKFLEKYNNEEEQMRYQREGFGQGHSA | -44.30 | 41.9 | HHHHHHHHHHHCCHHHHHHHHHHHHHCCCCCC |
| 641 | EEELKKLLEQINQEEEQMRYQQYGFGQGHSA | -44.30 | 38.7 | HHHHHHHHHHHHHHHHHHHHHHHHCCCCCCCC |
| 899 | EEELKKFLDQLNNEKEQMEYQRAGFGQGDSR | -44.29 | 51.6 | HHHHHHHHHHHHHHHHHHHHHHHHHCCCCCCC |
| 856 | EEELRKFLEQYNQEYEQMEYQRQGFGQGSSY | -44.27 | 38.7 | HHHHHHHHHHHHHHHHHHHHHHHHHCCCCCCC |
| 063 | EQELRKFLEKFNNEEEQMRYQREGFGQGHSS | -44.24 | 41.9 | HHHHHHHHHHHCHHHHHHHHHHHHHCCCCCCC |
| 276 | EEELRKLLEKINNEAEQMRYQREGFGQGHSA | -44.21 | 41.9 | HHHHHHHHHHHHHHHHHHHHHHHHHHCCCCCC |
| 873 | EEELRKILEKYNNEEEQMRYQQQGFGQGDSR | -44.20 | 45.2 | HHHHHHHHHHHCCHHHHHHHHHHHCCCCCCCC |
| 862 | EEELKKFLEKYNNEAEEMRKQQQGFGSGKSR | -44.14 | 48.4 | HHHHHHHHHHHHHHHHHHHHHHHHCCCCCCCC |
| 142 | EEEARKMLEKINNEAEQMRYQQQGFGQGHQS | -44.13 | 45.2 | HHHHHHHHHHHHHHHHHHHHHHHHCCCCCCCC |
| 669 | EEEKKKMLEKINNEAEQIRYQQQGFGQGHSS | -44.13 | 45.2 | HHHHHHHHHHHHHHHHHHHHHHHHHCCCCCCC |
| 206 | EEEKKKMLEKINNEAEQMRYQQQGFGQGHSA | -44.13 | 45.2 | HHHHHHHHHHHHHHHHHHHHHHHHCCCCCCCC |
| 632 | EEEKKKMLEKINNEEEQERYQQQGFGQGHSA | -44.13 | 41.9 | HHHHHHHHHHHCHHHHHHHHHHHHCCCCCCCC |
| 414 | EEEKKKMLEKINNEQEQMSYQQQGFGQGHQS | -44.13 | 41.9 | HHHHHHHHHHHHHHHHHHHHHHHHCCCCCCCC |

| | | | | |
|---|---|---|---|---|
| 190 | EEEKRKMLEKINNEAEQIRYQQNGFGQGHSS | -44.13 | 41.9 | HHHHHHHHHHHHHHHHHHHHHHHCCCCCCCC |
| 554 | EEEKRKMLEKINNEEEQMRYQQYGFGQGHSS | -44.13 | 38.7 | HHHHHHHHHHHCHHHHHHHHHHHCCCCCCCC |
| 560 | EEELKKFLEKYNQEAEQMRYQRQGFGQGDSR | -44.09 | 54.8 | HHHHHHHHHHHHHHHHHHHHHHHCCCCCCCC |
| 112 | EEELRKFLEKYNQEAEQIRYQQQGFGQGDSR | -44.09 | 51.6 | HHHHHHHHHHHHHHHHHHHHHHHCCCCCCCC |
| 819 | EEELRKFLEKYNQEAEQMRYQKEGFGQGDSR | -44.09 | 51.6 | HHHHHHHHHHHHHHHHHHHHHHHHCCCCCCC |
| 878 | EEQLKKFLEKMNNEKEQMEYQRAGFGQGHSS | -44.09 | 48.4 | HHHHHHHHHHHHHHHHHHHHHHHHHCCCCCC |
| 360 | EEELRKLLEQINNEEEQMRYQRYGFGQGHSA | -44.06 | 35.5 | HHHHHHHHHHHCHHHHHHHHHHHCCCCCCCC |
| 718 | EEELKKFLEQFNQEYEQMFYQQQGFGQGHSS | -44.06 | 48.4 | HHHHHHHHHHHHHHHHHHHHHHHHCCCCCCC |
| 952 | EEELKKFLEQYNQEAEQMRYQQQGFGQGHSA | -44.06 | 45.2 | HHHHHHHHHHHHHHHHHHHHHHHCCCCCCCC |
| 964 | EEELRKFLEQMNQEKEQMEYQRSGFGQGHSY | -44.06 | 41.9 | HHHHHHHHHHHHHHHHHHHHHHHHHCCCCCC |
| 835 | EEEKRKFLEKYNNEEEQMRYQREGFGQGHSS | -44.05 | 41.9 | HHHHHHHHHHHCCHHHHHHHHHHHCCCCCCC |
| 742 | EEELRKFLEKYNNESEQMRYQQQGFGQGHSA | -44.05 | 41.9 | HHHHHHHHHHHHHHHHHHHHHHHHCCCCCCC |
| 647 | EEELKKLLDQLNNEKEQMRYQQQGFGQGHSS | -43.96 | 41.9 | HHHHHHHHHHHHHHHHHHHHHHHCCCCCCCC |
| 926 | EEELKKFLQKFNNEYEDRFYQKEGFGKGDSR | -43.91 | 64.5 | HHHHHHHHHHHHHHHHHHHHHHHCCCCCCCC |
| 292 | EEELRKFLEAFNNRYEDMYYQQQGFGKGDSR | -43.91 | 51.6 | HHHHHHHHHHHHHHHHHHHHHHHCCCCCCCC |
| 336 | EEEKKKMLDQINNEAEQMRYQRYGFGQGDSR | -43.89 | 51.6 | HHHHHHHHHHHHHHHHHHHHHHHCCCCCCCC |
| 275 | EEELKKFLQKYNNEEEQMSYQQNGSGQGHSY | -43.89 | 45.2 | HHHHHHHHHHHHHHHHHHHHHHHCCCCCCCC |
| 523 | EEELKKILEQINNEEEQMRYQREGFGQGDSR | -43.87 | 45.2 | HHHHHHHHHHHCHHHHHHHHHHHHCCCCCCC |
| 338 | EEELRKILEQLNNEKEQMRYQRYGFGQGDSR | -43.87 | 41.9 | HHHHHHHHHHHHHHHHHHHHHHHCCCCCCCC |
| 005 | EEELKKLLEKINNEAEQIRYQQQGFGQGDSR | -43.85 | 51.6 | HHHHHHHHHHHHHHHHHHHHHHHCCCCCCCC |
| 424 | EEELKKLLEKINNEEEQMRYQQYGFGQGDSR | -43.85 | 48.4 | HHHHHHHHHHHCHHHHHHHHHHHCCCCCCCC |
| 130 | EEELRKLLEKINNEKEQMEYQRYGFGQGDSR | -43.85 | 45.2 | HHHHHHHHHHHHHHHHHHHHHHHCCCCCCCC |
| 279 | EEELRKLLEKLNNEKEQMRYQRQGFGQGDSR | -43.85 | 45.2 | HHHHHHHHHHHHHHHHHHHHHHHCCCCCCCC |
| 764 | EEELKKLLEQINNEEEQMRYQQYGFGQGHSA | -43.85 | 38.7 | HHHHHHHHHHHCHHHHHHHHHHHCCCCCCCC |
| 909 | EEELKKFLEKYNNEEEQMRYQREGFGQGDSR | -43.80 | 51.6 | HHHHHHHHHHHCCHHHHHHHHHHHCCCCCCC |
| 099 | EEEWKKILEQLNNEQEQMRYQRYGFGQGDSR | -43.78 | 45.2 | HHHHHHHHHHHHHHHHHHHHHHHCCCCCCCC |
| 789 | EEELKKFLEQYNNEAEQMRYQRYGFGQGHSA | -43.78 | 45.2 | HHHHHHHHHHHHHHHHHHHHHHHCCCCCCCC |
| 992 | EEELKKFLEQYNNEHEQMRYQQQGFGQGHQS | -43.78 | 41.9 | HHHHHHHHHHHHHHHHHHHHHHHHCCCCCCC |
| 884 | EEEKRKFLEQYNNEAEQMRYQREGKGQGHSH | -43.76 | 41.9 | HHHHHHHHHHHHHHHHHHHHHHHHCCCCCCC |
| 844 | EEEKRKFLEQYNNEEEQMRYQREGFGQGHQS | -43.76 | 38.7 | HHHHHHHHHHHCHHHHHHHHHHHHHCCCCCC |
| 659 | EEEKRKFLEQYNNEEEQMRYQREGFGQGHSS | -43.76 | 38.7 | HHHHHHHHHHHCHHHHHHHHHHHHCCCCCCC |
| 786 | EEELKKFLEQFNNEAEQMQYQRGGFGQGHQP | -43.76 | 48.4 | HHHHHHHHHHHHHHHHHHHHHHHCCCCCCCC |
| 820 | EEELKKFLEQFNNEYEQMFYQQKGFGQGHQS | -43.76 | 48.4 | HHHHHHHHHHHHHHHHHHHHHHHHCCCCCCC |
| 273 | EEELKKFLEQLNNEKEQEEYQRYGFGQGHQS | -43.76 | 41.9 | HHHHHHHHHHHHHHHHHHHHHHHCCCCCCCC |
| 249 | EEELKKFLEQYNNEAEQMRYQKEGFGQGHSS | -43.76 | 45.2 | HHHHHHHHHHHHHHHHHHHHHHHHCCCCCCC |
| 606 | EEELKKFLEQYNNEAEQMRYQQQGFGQGHSS | -43.76 | 45.2 | HHHHHHHHHHHHHHHHHHHHHHHCCCCCCCC |
| 948 | EEELKKFLEQYNNEEEQERYQQQGFGQGHSA | -43.76 | 41.9 | HHHHHHHHHHHCCHHHHHHHHHHCCCCCCCC |
| 298 | EEELKKFLEQYNNEEEQMRYQREGFGQGHQS | -43.76 | 41.9 | HHHHHHHHHHHCHHHHHHHHHHHHHCCCCCC |

| | | | | |
|---|---|---|---|---|
| 960 | EEELKKFLEQYNNEEEQMRYQREGFGQGHSA | -43.76 | 41.9 | HHHHHHHHHHHCHHHHHHHHHHHHHCCCCCC |
| 840 | EEELKKFLEQYNNESEQIRYQQQGFGQGHSS | -43.76 | 41.9 | HHHHHHHHHHHHHHHHHHHHHHHHCCCCCCC |
| 384 | EEELKKFLEQYNNESEQMRYQKEGFGQGHSS | -43.76 | 41.9 | HHHHHHHHHHHHHHHHHHHHHHHHCCCCCCC |
| 282 | EEELKKFLEQYNNESEQMRYQQNGSGQGHSY | -43.76 | 41.9 | HHHHHHHHHHHHHHHHHHHHHHCCCCCCCCC |
| 927 | EEELKKFLEQYNNESEQMRYQQQGFGQGHSA | -43.76 | 41.9 | HHHHHHHHHHHHHHHHHHHHHHHHCCCCCCC |
| 024 | EEELKKFLEQYNNESEQMRYQRYGFGQGHSA | -43.76 | 41.9 | HHHHHHHHHHHHHHHHHHHHHHHCCCCCCCC |
| 404 | EEELKKFLEQYNNEYEQMQYQKEGFGQGHSA | -43.76 | 41.9 | HHHHHHHHHHHHHHHHHHHHHHHHCCCCCCC |
| 731 | EEELRKFLEQFNNEQEQMSYQQQGFGQGHSS | -43.76 | 41.9 | HHHHHHHHHHHHHHHHHHHHHHHCCCCCCCC |
| 468 | EEELRKFLEQLNNEEEQMRYQQYGFGQGHQS | -43.76 | 38.7 | HHHHHHHHHHHHHHHHHHHHHHHCCCCCCCC |
| 758 | EEELRKFLEQYNNEAEQMRYQQQGFGQGHQS | -43.76 | 41.9 | HHHHHHHHHHHHHHHHHHHHHHHCCCCCCCC |
| 409 | EEELRKFLEQYNNEAEQMRYQQQGFGQGHSA | -43.76 | 41.9 | HHHHHHHHHHHHHHHHHHHHHHHHCCCCCCC |
| 420 | EEELRKFLEQYNNEAEQMRYQRQGFGQGHQS | -43.76 | 41.9 | HHHHHHHHHHHHHHHHHHHHHHHHCCCCCCC |
| 210 | EEELRKFLEQYNNEAEQMRYQRQGFGQGHSA | -43.76 | 41.9 | HHHHHHHHHHHHHHHHHHHHHHHHCCCCCCC |
| 730 | EEELRKFLEQYNNEEEQMRYQREGFGQGHSS | -43.76 | 38.7 | HHHHHHHHHHHCHHHHHHHHHHHHCCCCCCC |
| 906 | EEELRKFLEQYNNEHEQMRYQQQGFGQGHSS | -43.76 | 38.7 | HHHHHHHHHHHHHHHHHHHHHHHHCCCCCCC |
| 229 | EEELRKFLEQYNNESEQERYQRKGFGQGHSS | -43.76 | 38.7 | HHHHHHHHHHHCCHHHHHHHHHHCCCCCCCC |
| 829 | EEELRKFLEQYNNESEQMRYQQQGFGQGHSA | -43.76 | 38.7 | HHHHHHHHHHHHHHHHHHHHHHHHCCCCCCC |
| 913 | EEELRKFLEQYNNESEQMRYQQQGFGQGHSS | -43.76 | 38.7 | HHHHHHHHHHHHHHHHHHHHHHHHCCCCCCC |
| 885 | EEELRKFLEQYNNEYEQEEYQRKGFGQGHSS | -43.76 | 38.7 | HHHHHHHHHHHHHHHHHHHHHHHCCCCCCCC |
| 959 | EEELRKFLEQYNNEYEQMFYQQKGFGQGHSS | -43.76 | 41.9 | HHHHHHHHHHHHHHHHHHHHHHHHCCCCCCC |
| 475 | EEELRKFLEQYNNEYEQMYYQQKGFGQGHSS | -43.76 | 38.7 | HHHHHHHHHHHHHHHHHHHHHHHHCCCCCCC |
| 618 | EQELRKFLEQYNNESEQMRYQRYGFGQGHSS | -43.76 | 35.5 | HHHHHHHHHHHHHHHHHHHHHHHCCCCCCCC |
| 148 | EEELRKFLEQYNNRAEQERYQQQGFGQGHSA | -43.74 | 38.7 | HHHHHHHHHHHHHHHHHHHHHHHCCCCCCCC |
| 953 | EEEKKKMLEQINQEEEQMRYQQQGFGQGHSA | -43.66 | 38.7 | HHHHHHHHHHHHHHHHHHHHHHHCCCCCCCC |
| 818 | EEEKKKMLEQYNQEEEQMRYQREGFGQGHSA | -43.66 | 38.7 | HHHHHHHHHHHHHHHHHHHHHHHHHCCCCCC |
| 531 | EEEKKKMLEKINNEEEQIRYQQQGFGQGHSS | -43.66 | 41.9 | HHHHHHHHHHHCHHHHHHHHHHHHCCCCCCC |
| 908 | EEEKKKMLEKINNEEEQMRYQRYGFGQGHSA | -43.66 | 41.9 | HHHHHHHHHHHCHHHHHHHHHHHCCCCCCCC |
| 482 | EEELRKFLEKYNNESEQMRYQRYGFGQGHSA | -43.63 | 41.9 | HHHHHHHHHHHHHHHHHHHHHHHCCCCCCCC |
| 986 | EEEKRKFLDQMNNEKEQMEYQRAGFGQGDSR | -43.62 | 48.4 | HHHHHHHHHHHHHHHHHHHHHHHHCCCCCCC |
| 443 | EEELRKFLEQFNQEYEQMFYQQKGFGQGHSA | -43.60 | 45.2 | HHHHHHHHHHHHHHHHHHHHHHHHCCCCCCC |
| 545 | EEELRKILDDLNNRKEQMRYQRYGFGQGHSA | -43.58 | 35.5 | HHHHHHHHHHHHHHHHHHHHHHHCCCCCCCC |
| 721 | EEEKRKFLEQFNQEYEQMFYQQKGFGQGHSA | -43.58 | 45.2 | HHHHHHHHHHHHHHHHHHHHHHHHCCCCCCC |
| 440 | EEELKKFLEQYNQESEQMRYQQQGFGQGHQS | -43.58 | 41.9 | HHHHHHHHHHHHHHHHHHHHHHHHCCCCCCC |
| 498 | EEELRKFLEQYNQEEEQMRYQREGFGQGHQS | -43.58 | 38.7 | HHHHHHHHHHHHHHHHHHHHHHHHHCCCCCC |
| 225 | EEELRKFLEQYNQEYEQMEYQRNGRGQGHSY | -43.58 | 38.7 | HHHHHHHHHHHHHHHHHHHHHHHCCCCCCCC |
| 897 | EEELRKFLEQYNNEYEQMFYQQKGFGQGHQS | -43.53 | 41.9 | HHHHHHHHHHHHHHHHHHHHHHHHCCCCCCC |
| 567 | EEELKKILEQINQEEEQMRYQREGFGQGDSR | -43.49 | 45.2 | HHHHHHHHHHHHHHHHHHHHHHHHCCCCCCC |
| 542 | EEELKKLLEQINNEEEQMRYQQQGFGQGHSA | -43.43 | 38.7 | HHHHHHHHHHHCHHHHHHHHHHHCCCCCCCC |

| | | | | |
|---|---|---|---|---|
| 262 | EEEKRKMLEKINNEQEQMSYQRHGFGQGDSR | -43.40 | 45.2 | HHHHHHHHHHHHHHHHHHHHHHHHCCCCCCCC |
| 313 | EEELKKLLEQINQEEEQMRYQREGFGQGDSR | -43.38 | 45.2 | HHHHHHHHHHHHHHHHHHHHHHHHHCCCCCCC |
| 293 | EEELRKLLEQLNQEKEQMSYQQQGFGQGDSR | -43.38 | 41.9 | HHHHHHHHHHHHHHHHHHHHHHHHCCCCCCCC |
| 907 | EEELRKLLEQLNQEQEQMRYQQQGFGQGDSR | -43.38 | 41.9 | HHHHHHHHHHHHHHHHHHHHHHHHCCCCCCCC |
| 824 | EEEARKMLEQINNEEEQMRYQRYGFGQGHSA | -43.37 | 38.7 | HHHHHHHHHHHHHHHHHHHHHHHHCCCCCCCC |
| 597 | EEEKKKMLEQINNEEEQMRYQQQGFGQGHSA | -43.37 | 38.7 | HHHHHHHHHHHCHHHHHHHHHHHHCCCCCCCC |
| 751 | EEEKKKMLEQINNEEEQMRYQREGFGQGHSS | -43.37 | 38.7 | HHHHHHHHHHHCHHHHHHHHHHHHHCCCCCCC |
| 361 | EEEKKKMLEQINNEEEQMRYQRYGFGQGHSS | -43.37 | 38.7 | HHHHHHHHHHHCHHHHHHHHHHHHCCCCCCCC |
| 467 | EEEKRKMLEQINNEAEQMRYQKEGFGQGHSS | -43.37 | 38.7 | HHHHHHHHHHHHHHHHHHHHHHHHHCCCCCCC |
| 516 | EEEKRKMLEQINNEAEQMRYQRQGFGQGHSS | -43.37 | 38.7 | HHHHHHHHHHHHHHHHHHHHHHHHCCCCCCCC |
| 684 | EEEKRKMLEQINNEAEQMRYQRYGFGQGHSA | -43.37 | 38.7 | HHHHHHHHHHHHHHHHHHHHHHHHCCCCCCCC |
| 082 | EEEKRKMLEQINNEEEQMRYQQYGFGQGHSS | -43.37 | 35.5 | HHHHHHHHHHHHHHHHHHHHHHHHCCCCCCCC |
| 923 | EEEKRKMLEQINNEQEQMSYQQQGFGQGHSS | -43.37 | 35.5 | HHHHHHHHHHHHHHHHHHHHHHHHCCCCCCCC |
| 505 | EEEKRKFLEKYNNESEQKRYQQQGFGQGDSR | -43.29 | 48.4 | HHHHHHHHHHHHHHHHHHHHHHHHCCCCCCCC |
| 594 | EEEKRKFLEKYNNEYEQMYYQQKGFGQGDSR | -43.29 | 48.4 | HHHHHHHHHHHHHHHHHHHHHHHHCCCCCCCC |
| 686 | EEELKKFLEKLNNEEEQMRYQQYGFGQGDSR | -43.29 | 51.6 | HHHHHHHHHHHCHHHHHHHHHHHHCCCCCCCC |
| 931 | EEELKKFLEKYNNESEQIRYQQQGFGQGDSR | -43.29 | 51.6 | HHHHHHHHHHHHHHHHHHHHHHHHCCCCCCCC |
| 342 | EEELRKFLEKYNNEAEQIYYQQNGFGQGDSR | -43.29 | 51.6 | HHHHHHHHHHHHHHHHHHHHHHHHCCCCCCCC |
| 656 | EEELRKFLEKYNNEYEQMFYQQKGFGQGDSR | -43.29 | 51.6 | HHHHHHHHHHHHHHHHHHHHHHHHCCCCCCCC |
| 303 | EEELKKFLEQLNNEEEQMRYQQYGFGQGHSS | -43.29 | 41.9 | HHHHHHHHHHHHHHHHHHHHHHHHCCCCCCCC |
| 422 | EEELKKFLEQYNNESEQMRYQQQGFGQGHQS | -43.29 | 41.9 | HHHHHHHHHHHHHHHHHHHHHHHHHCCCCCCC |
| 734 | EEELKKFLEQYNNESEQMRYQQQGFGQGHSS | -43.29 | 41.9 | HHHHHHHHHHHHHHHHHHHHHHHHHCCCCCCC |
| 334 | EEELRKFLEQFNNEYEQMFYQQKGFGQGHSS | -43.29 | 45.2 | HHHHHHHHHHHHHHHHHHHHHHHHHCCCCCCC |
| 320 | EEELRKFLEQYNNESEQMRYQQQGFGQGHQA | -43.29 | 38.7 | HHHHHHHHHHHHHHHHHHHHHHHHHCCCCCCC |
| 241 | EEELRKFLEQYNNESEQMRYQQQGFGQGHQS | -43.29 | 38.7 | HHHHHHHHHHHHHHHHHHHHHHHHHCCCCCCC |
| 072 | EEELRKFLEQYNNESEQMRYQRYGFGQGHQS | -43.29 | 38.7 | HHHHHHHHHHHHHHHHHHHHHHHHCCCCCCCC |
| 234 | EEELRKFLEQYNNEYEQEFYQQKGFGQGHSA | -43.29 | 41.9 | HHHHHHHHHHHHHHHHHHHHHHHHCCCCCCCC |
| 951 | EEELRRFLEQLNNEEEDMRYQQQGFGKGDSR | -43.28 | 51.6 | HHHHHHHHHHHHHHHHHHHHHHHHCCCCCCCC |
| 817 | EEELRKLLEKINNEAEQRRYQRYGFGQGHSS | -43.22 | 41.9 | HHHHHHHHHHHHHHHHHHHHHHHHCCCCCCCC |
| 027 | EEELRKFLEDYNNRAEDMRYQRQGFGKGHQS | -43.19 | 45.2 | HHHHHHHHHHHHHHHHHHHHHHHHHCCCCCCC |
| 077 | EEEKRKMLEKINQEAEQMRYQQYGFGQGDSR | -43.19 | 48.4 | HHHHHHHHHHHHHHHHHHHHHHHHCCCCCCCC |
| 904 | EEELKKLLEKINNEAEQMRYQKEGFGQGDSR | -43.18 | 51.6 | HHHHHHHHHHHHHHHHHHHHHHHHHCCCCCCC |
| 483 | EEELKKLLEKLNNEQEQIRYQQQGFGQGDSR | -43.18 | 48.4 | HHHHHHHHHHHHHHHHHHHHHHHHCCCCCCCC |
| 970 | EEEWKKLIEKINNEEEQKRYQRYGFGQGDSR | -43.18 | 45.2 | HHHHHHHHHHHCHHHHHHHHHHHHCCCCCCCC |
| 667 | EEELKKFLQKYNNESEDMRYQKEGFGKGDSR | -43.17 | 58.1 | HHHHHHHHHHHHHCHHHHHHHHHHCCCCCCCC |
| 088 | EEEKKKMLEQINNEAEQMRYQRYGFGQGHSA | -43.14 | 41.9 | HHHHHHHHHHHHHHHHHHHHHHHHCCCCCCCC |
| 726 | EEELRKFLEKFNNEYEQRFYQQKGFGQGHSA | -43.13 | 48.4 | HHHHHHHHHHHHHHHHHHHHHHHHCCCCCCCC |
| 811 | EEELKKLLEQINNEAEQMRYQQGGFGQGDSR | -43.09 | 48.4 | HHHHHHHHHHHHHHHHHHHHHHHCCCCCCCCC |

| | | | | |
|---|---|---|---|---|
| 634 | EQELRKLLEQINNEEEQMRYQQYGFGQGDSR | -43.09 | 38.7 | HHHHHHHHHHHHHHHHHHHHHHHCCCCCCCC |
| 289 | EEELKKFLEQYNNEEEQMRYQREGFGQGDSR | -43.03 | 48.4 | HHHHHHHHHHHHCHHHHHHHHHHHCCCCCCC |
| 596 | EEELKKFLEQYNNESEQMRYQRYGFGQGDSR | -43.03 | 48.4 | HHHHHHHHHHHHHHHHHHHHHHHCCCCCCCC |
| 078 | EEELRKFLEQFNNEYEQMFYQQKGFGQGDSR | -43.03 | 51.6 | HHHHHHHHHHHHHHHHHHHHHHHCCCCCCCC |
| 407 | EEELRKFLEQLNNEEEQMRYQQYGFGQGDSR | -43.03 | 45.2 | HHHHHHHHHHHHHHHHHHHHHHHCCCCCCCC |
| 681 | EEELRKFLEQYNNEEEQMRYQREGFGQGDSR | -43.03 | 45.2 | HHHHHHHHHHHHCHHHHHHHHHHHCCCCCCC |
| 451 | EEEWKKFIEQYNNEAEQMRYQRQGFGQGDSR | -43.03 | 48.4 | HHHHHHHHHHHHHHHHHHHHHHHCCCCCCCC |
| 109 | EQELRKFLEQFNNEYEQMFYQQKGFGQGDSR | -43.03 | 48.4 | HHHHHHHHHHHHHHHHHHHHHHHCCCCCCCC |
| 281 | EEELRKILEQFNNEEEQMRYQREGFGQGDSR | -42.97 | 45.2 | HHHHHHHHHHHHCHHHHHHHHHHHCCCCCCC |
| 401 | EEELKKLLEKLNNEKEQRRYQQQGFGQGDSR | -42.97 | 48.4 | HHHHHHHHHHHHHHHHHHHHHHHCCCCCCCC |
| 299 | EEELKKFLEQFNNEQEQMSYQRHGFGQGHQS | -42.96 | 45.2 | HHHHHHHHHHHHHHHHHHHHHHHCCCCCCCC |
| 581 | EEQLRKLLEQLNQEKEQMSYQRYGFGQGDSR | -42.94 | 45.2 | HHHHHHHHHHHHHHHHHHHHHHHCCCCCCCC |
| 205 | EEEKRKFLEKYNQEAEQMRYQRYGFGQGDSR | -42.91 | 51.6 | HHHHHHHHHHHHHHHHHHHHHHHCCCCCCCC |
| 098 | EEELKKFLEKFNQEYEQMFYQKEGFGQGDSR | -42.91 | 58.1 | HHHHHHHHHHHHHHHHHHHHHHHCCCCCCCC |
| 798 | EEELRKFLEKLNQEEEQMRYQQYGFGQGDSR | -42.91 | 48.4 | HHHHHHHHHHHHHHHHHHHHHHHCCCCCCCC |
| 166 | EEQLKKFLEQYNNESEQMRYQQQGFGQGHSA | -42.85 | 45.2 | HHHHHHHHHHHHHHHHHHHHHHHHCCCCCCC |
| 662 | EEELRKFLEQYNQRSEQERYQQQGFGQGDSR | -42.79 | 41.9 | HHHHHHHHHHHHHHHHHHHHHHHCCCCCCCC |
| 585 | EEELKKFLEDYNNRAEDMRYQQNGFGKGHSA | -42.67 | 48.4 | HHHHHHHHHHHHHHHHHHHHHHHHCCCCCCC |
| 911 | EEELRKFLEKYNNEAEQIRYQQQGFGQGDSR | -42.62 | 51.6 | HHHHHHHHHHHHHHHHHHHHHHHCCCCCCCC |
| 458 | EEELRKFLEQYNQEYEQMFYQQKGFGQGDSR | -42.60 | 48.4 | HHHHHHHHHHHHHHHHHHHHHHHCCCCCCCC |
| 437 | EEEKKKMLEKINQEEEQMRYQREGFGQGDSR | -42.59 | 48.4 | HHHHHHHHHHHHHHHHHHHHHHHHCCCCCCC |
| 167 | EEELRKFLEQFNNEYEQMFYQQKGFGQGTSR | -42.58 | 48.4 | HHHHHHHHHHHHHHHHHHHHHHHHCCCCCCC |
| 344 | EEEKKKMLEQFNNEYEQMYYQQNGQGQGHSY | -42.57 | 41.9 | HHHHHHHHHHHHHHHHHHHHHHHCCCCCCCC |
| 478 | EEELKKFLEQFNNEYEQMFYQQKGFGQGDSR | -42.52 | 54.8 | HHHHHHHHHHHHHHHHHHHHHHHCCCCCCCC |
| 860 | EEELKKFLEQYNNEAEQMRYQQQGFGQGDSR | -42.52 | 51.6 | HHHHHHHHHHHHHHHHHHHHHHHCCCCCCCC |
| 327 | EEELKKFLEQYNNEAEQMRYQRQGFGQGDSR | -42.52 | 51.6 | HHHHHHHHHHHHHHHHHHHHHHHCCCCCCCC |
| 010 | EEELKKFLEQYNNESEQMRYQQQGFGQGDSR | -42.52 | 48.4 | HHHHHHHHHHHHHHHHHHHHHHHCCCCCCCC |
| 435 | EEELRKFLEQYNNESEQMRYQQQGFGQGDSR | -42.52 | 45.2 | HHHHHHHHHHHHHHHHHHHHHHHCCCCCCCC |
| 022 | EEELRKFLEQYNNEYEQEEYQRKGFGQGDSR | -42.52 | 45.2 | HHHHHHHHHHHHHHHHHHHHHHHCCCCCCCC |
| 318 | EEEWKKFIEQYNNEEEQMRYQREGFGQGDSR | -42.52 | 45.2 | HHHHHHHHHHHHCHHHHHHHHHHHCCCCCCC |
| 792 | EEEWRKFLEQFNNEYEQMFYQQQGFGQGDSR | -42.44 | 51.6 | HHHHHHHHHHHHHHHHHHHHHHHCCCCCCCC |
| 169 | EEEWRKFIEQLNNEEEQMRYQQQGFGQGDSR | -42.43 | 41.9 | HHHHHHHHHHHHHHHHHHHHHHHCCCCCCCC |
| 244 | EEEKRKMLEQINQEEEQMRYQQQGFGQGDSR | -42.42 | 41.9 | HHHHHHHHHHHHHHHHHHHHHHHCCCCCCCC |
| 377 | EEELKKLLEQINNEQEQEEYQRKGFGQGDSR | -42.42 | 45.2 | HHHHHHHHHHHHHHHHHHHHHHHCCCCCCCC |
| 846 | EEELKKLLEQLNNEKEQMRYQRQGFGQGDSR | -42.42 | 45.2 | HHHHHHHHHHHHHHHHHHHHHHHCCCCCCCC |
| 572 | EEEKRKFLEKYNNESEQRRYQRYGFGQGDSR | -42.40 | 48.4 | HHHHHHHHHHHHCHHHHHHHHHHCCCCCCCC |
| 383 | EEEARKMLEQINNEEEQMRYQQQGFGQGDSR | -42.13 | 45.2 | HHHHHHHHHHHHHHHHHHHHHHHCCCCCCCC |
| 258 | EEELRKFLEKYNNESEQRRYQQEGFGQGDSR | -41.89 | 48.4 | HHHHHHHHHHHHHHHHHHHHHHHCCCCCCCC |

| | | | | |
|---|---|---|---|---|
| 600 | EEELKKFLEQYNNEEEQMRYQRQGFGQGDSR | -41.85 | 48.4 | HHHHHHHHHHHCHHHHHHHHHHHCCCCCCCC |
| 019 | EEELKKFLEQYNNEYEQMYYQQKGFGQGDSR | -41.85 | 48.4 | HHHHHHHHHHHHHHHHHHHHHHHCCCCCCCC |
| 756 | EEEKKKMLEQYEEEERQMRYQREGFGQGHSS | -41.54 | 32.3 | HHHHHHHHHHHHHHHHHHHHHHHHCCCCCCC |
| 350 | EEEAKKMLEQINNEAEQMRYQRYGFGQGDSR | -41.46 | 51.6 | HHHHHHHHHHHHHHHHHHHHHHHCCCCCCCC |
| 639 | EEEKRKMLEQINNEEEQMRYQREGFGQGDSR | -41.46 | 41.9 | HHHHHHHHHHHCHHHHHHHHHHHHCCCCCCC |
| 917 | EEEKRKMLEQINNEEEQERYQQQGFGQGDSR | -41.19 | 41.9 | HHHHHHHHHHHCHHHHHHHHHHHCCCCCCCC |
| 928 | EEELKKFLEDYNNRYEQEEYQRKGFGQGHSS | -40.53 | 38.7 | HHHHHHHHHHHHHHHHHHHHHHHCCCCCCCC |

**Table S5**. Summary of 695 peptide sequences designed using EvoEF2 and the evolutionary profile (weight = 1.00).

| #label | #peptide | #binding(EEU) | #SeqID(%) | #Secondary structure |
|---|---|---|---|---|
| WT | EEQAKTFLDKFNHEAEDLFYQSSGLGKGDFR | -46.46 | 100. | HHHHHHHHHHHHHHHHHHHHHHHCCCCCCCC |
| 322 | EEELKKILEKINQEEEDMRYQQQGFGKGHDA | -49.06 | 48.4 | HHHHHHHHHHHHHHHHHHHHHHHHCCCCCCC |
| 591 | EEELKKILEKLNQEQEDMRYQREGFGKGHSA | -49.06 | 48.4 | HHHHHHHHHHHHHHHHHHHHHHHHHCCCCCC |
| 276 | EEEWRKILEKLNQEEEDIKYQQNGFGKGHDA | -49.06 | 45.2 | HHHHHHHHHHHHHHHHHHHHHHHHCCCCCCC |
| 783 | EEELKKLLDQLNQEQEDMRYQQNGFGKGHDA | -49.05 | 48.4 | HHHHHHHHHHHHHHHHHHHHHHHHCCCCCCC |
| 773 | EEELKKILEKINNEEEDIRYQQQGFGKGHDA | -48.77 | 48.4 | HHHHHHHHHHHHHHHHHHHHHHHHCCCCCCC |
| 255 | EEELRKILEKLNNEQEDMRYQQNGFGKGHDA | -48.77 | 45.2 | HHHHHHHHHHHHHHHHHHHHHHHHCCCCCCC |
| 199 | EEEWRKILEKLNNEEEDMKYQKEGFGKGHDA | -48.77 | 45.2 | HHHHHHHHHHHHHHHHHHHHHHHHCCCCCCC |
| 604 | EEELRKILEKLNNEQEDMRYQRYGFGKGHDA | -48.53 | 45.2 | HHHHHHHHHHHHHHHHHHHHHHHCCCCCCCC |
| 895 | EEELKKLLDQINNEEEDMRYQQQGFGKGHDA | -48.23 | 48.4 | HHHHHHHHHHHHHHHHHHHHHHHHCCCCCCC |
| 237 | EEEARKFLDQFNNEYEDMFYQQKGFGKGHDA | -48.19 | 58.1 | HHHHHHHHHHHHHHHHHHHHHHHHCCCCCCC |
| 615 | EEELKKLLEKLNQEEEDIKYQQNGFGKGHDA | -48.05 | 48.4 | HHHHHHHHHHHHHHHHHHHHHHHHCCCCCCC |
| 703 | EEELKKLLEKLNQEKEDMTYQRHGFGKGHDA | -48.05 | 48.4 | HHHHHHHHHHHHHHHHHHHHHHHCCCCCCCC |
| 900 | EEELRKLLEKINQEAEDIRYQQNGFGKGHDA | -48.05 | 48.4 | HHHHHHHHHHHHHHHHHHHHHHHHCCCCCCC |
| 758 | EEELRKLLEKLNQEQEDMRYQKEGFGKGHDA | -48.05 | 45.2 | HHHHHHHHHHHHHHHHHHHHHHHHHCCCCCC |
| 496 | EEELKKILEQLNNEKEDMSFQQQGFGKGKDA | -48.00 | 41.9 | HHHHHHHHHHHHHHHHHHHHHHHHCCCCCCC |
| 651 | EEELKKLLEKINNEEEDIRFQQQGFGKGKDA | -47.99 | 45.2 | HHHHHHHHHHHHHHHHHHHHHHHHCCCCCCC |
| 355 | EEELRKLLEKLNNEQEDIRYQQQGFGKGHDA | -47.94 | 45.2 | HHHHHHHHHHHHHHHHHHHHHHHHCCCCCCC |
| 638 | EEELKKFLEQYNNEAEDMRYQRQGFGKGHDA | -47.92 | 51.6 | HHHHHHHHHHHHHHHHHHHHHHHHCCCCCCC |
| 806 | EEELRKLLEKIENEREDITYQQNGFGKGHDA | -47.91 | 41.9 | HHHHHHHHHHHHHHHHHHHHHHHCCCCCCCC |
| 936 | EEELRKLLEKLNQEQEDIRYQQNGFGKGHDA | -47.81 | 45.2 | HHHHHHHHHHHHHHHHHHHHHHHHCCCCCCC |
| 091 | EEELKKLLEKINNEAEDMRYQQQGFGKGHDA | -47.77 | 51.6 | HHHHHHHHHHHHHHHHHHHHHHHHCCCCCCC |
| 263 | EEELKKILEQINNEAEDMRYQQGGFGKGHDF | -47.77 | 48.4 | HHHHHHHHHHHHHHHHHHHHHHCCCCCCCCC |
| 075 | EEELKKILEQINNEEEDMRYQQQGFGKGHDA | -47.77 | 45.2 | HHHHHHHHHHHHHHHHHHHHHHHHCCCCCCC |
| 934 | EEELKKILEQINNEEEDMRYQQYGFGKGHDA | -47.77 | 45.2 | HHHHHHHHHHHHHHHHHHHHHHHCCCCCCCC |
| 415 | EEELKKILEQLNNEEEDMKYQQNGFGKGHDA | -47.77 | 45.2 | HHHHHHHHHHHHHHHHHHHHHHHHCCCCCCC |
| 512 | EEELKKILEQLNNEQEDMRYQQQGFGKGHDA | -47.77 | 45.2 | HHHHHHHHHHHHHHHHHHHHHHHHCCCCCCC |
| 623 | EEELKKILEQLNNEQEDMRYQRYGFGKGHDA | -47.77 | 45.2 | HHHHHHHHHHHHHHHHHHHHHHHCCCCCCCC |
| 316 | EEELRKILEQINNEEEDMRYQQYGFGKGHDA | -47.77 | 41.9 | HHHHHHHHHHHHHHHHHHHHHHHCCCCCCCC |
| 886 | EEELRKILEQLNNEQEDMRYQQQGFGKGHDA | -47.77 | 41.9 | HHHHHHHHHHHHHHHHHHHHHHHHCCCCCCC |
| 239 | EEELKKLLEKINNEAEDIRYQQNGFGKGHDA | -47.75 | 51.6 | HHHHHHHHHHHHHHHHHHHHHHHHCCCCCCC |
| 910 | EEELKKLLEKINNEEEDIRYQQQGFGKGHDA | -47.75 | 48.4 | HHHHHHHHHHHHHHHHHHHHHHHHCCCCCCC |
| 413 | EEELKKLLEKINNEEEDIRYQQQGFGKGHSA | -47.75 | 48.4 | HHHHHHHHHHHHHHHHHHHHHHHHCCCCCCC |
| 720 | EEELKKLLEKINNEEEDMRYQQYGFGKGHDA | -47.75 | 48.4 | HHHHHHHHHHHHHHHHHHHHHHHCCCCCCCC |
| 999 | EEELKKLLEKINNEEEDMRYQREGFGKGHDA | -47.75 | 48.4 | HHHHHHHHHHHHHHHHHHHHHHHHHCCCCCC |

| | | | | |
|---|---|---|---|---|
| 526 | EEELKKLLEKLNNEKEDMRYQQQGFGKGHDA | -47.75 | 48.4 | HHHHHHHHHHHHHHHHHHHHHHHHCCCCCCC |
| 733 | EEELKKLLEKLNNEQEDIRYQQNGFGKGHDA | -47.75 | 48.4 | HHHHHHHHHHHHHHHHHHHHHHHHCCCCCCC |
| 979 | EEELKKLLEKLNNEQEDMRYQQQGFGKGHDA | -47.75 | 48.4 | HHHHHHHHHHHHHHHHHHHHHHHHCCCCCCC |
| 363 | EEELRKLLEKINNEAEDIRYQQNGFGKGHDA | -47.75 | 48.4 | HHHHHHHHHHHHHHHHHHHHHHHHCCCCCCC |
| 045 | EEELRKLLEKINNEAEDIRYQQNGFGKGHSA | -47.75 | 48.4 | HHHHHHHHHHHHHHHHHHHHHHHHCCCCCCC |
| 849 | EEELRKLLEKINNEAEDMRYQQQGFGKGHDA | -47.75 | 48.4 | HHHHHHHHHHHHHHHHHHHHHHHHCCCCCCC |
| 819 | EEELRKLLEKINNEAEDMRYQRQGFGKGHDA | -47.75 | 48.4 | HHHHHHHHHHHHHHHHHHHHHHHHCCCCCCC |
| 361 | EEELRKLLEKINNEEEDIRYQQQGFGKGHDA | -47.75 | 45.2 | HHHHHHHHHHHHHHHHHHHHHHHHCCCCCCC |
| 454 | EEELRKLLEKINNEEEDIRYQQQGFGKGHSA | -47.75 | 45.2 | HHHHHHHHHHHHHHHHHHHHHHHHCCCCCCC |
| 275 | EEELRKLLEKINNEEEDMRYQQYGFGKGHDA | -47.75 | 45.2 | HHHHHHHHHHHHHHHHHHHHHHHCCCCCCCC |
| 326 | EEELRKLLEKLNNEKEDMRYQQQGFGKGHDA | -47.75 | 45.2 | HHHHHHHHHHHHHHHHHHHHHHHHCCCCCCC |
| 285 | EEELRKLLEKLNNEKEDMSYQQQGFGKGHDA | -47.75 | 45.2 | HHHHHHHHHHHHHHHHHHHHHHHCCCCCCCC |
| 107 | EEELRKLLEKLNNEQEDIRYQQNGFGKGHDA | -47.75 | 45.2 | HHHHHHHHHHHHHHHHHHHHHHHHCCCCCCC |
| 510 | EEELRKLLEKLNNEQEDMRYQQNGFGKGHDA | -47.75 | 45.2 | HHHHHHHHHHHHHHHHHHHHHHHHCCCCCCC |
| 567 | EEELRKLLEKLNNEQEDMRYQRQGFGKGHDA | -47.75 | 45.2 | HHHHHHHHHHHHHHHHHHHHHHHHCCCCCCC |
| 469 | EEEWKKLMEKYNNEEEDMRYQREGFGKGHDA | -47.75 | 45.2 | HHHHHHHHHHHCCHHHHHHHHHHHHCCCCCC |
| 582 | EQELRKLLEKLNNEQEDIRYQQNGFGKGHDA | -47.75 | 41.9 | HHHHHHHHHHHHHHHHHHHHHHHHCCCCCCC |
| 959 | EEEARKFLEKYNQEEEDMRYQQQGFGKGHSA | -47.67 | 51.6 | HHHHHHHHHHHHHHHHHHHHHHHHCCCCCCC |
| 833 | EEELRKILEQLNNEQEDMRYQQGGFGKGHDA | -47.53 | 41.9 | HHHHHHHHHHHHHHHHHHHHHHCCCCCCCCC |
| 788 | EEEARKFLEKYNQEEEDMRYQREGFGKGHSA | -47.49 | 51.6 | HHHHHHHHHHHHHHHHHHHHHHHHHCCCCCC |
| 228 | EEEARKFLEKFNQEEEDMRYQRQGFGKGHDA | -47.48 | 54.8 | HHHHHHHHHHHHHHHHHHHHHHHHCCCCCCC |
| 042 | EEEARKFLEKYNQEAEDIRYQQNGFGKGHDA | -47.48 | 54.8 | HHHHHHHHHHHHHHHHHHHHHHHHCCCCCCC |
| 694 | EEEARKFLEKYNQEAEDIYYQQNGFGKGHDA | -47.48 | 54.8 | HHHHHHHHHHHHHHHHHHHHHHHCCCCCCCC |
| 301 | EEEARKFLEKYNQEAEDMRYQQQGFGKGHDA | -47.48 | 54.8 | HHHHHHHHHHHHHHHHHHHHHHHHCCCCCCC |
| 486 | EEEARKFLEKYNQEEEDMRYQQQGFGKGHDA | -47.48 | 51.6 | HHHHHHHHHHHHHHHHHHHHHHHHCCCCCCC |
| 006 | EEEARKFLEKYNQEEEDMRYQREGFGKGHDA | -47.48 | 51.6 | HHHHHHHHHHHHHHHHHHHHHHHHHCCCCCC |
| 547 | EEEARKFLEKYNQESEDIRYQQNGFGKGHDA | -47.48 | 51.6 | HHHHHHHHHHHHHHHHHHHHHHHCCCCCCCC |
| 101 | EEEARKFLEKYNQESEDIRYQQQGFGKGHDA | -47.48 | 51.6 | HHHHHHHHHHHHHHHHHHHHHHHHCCCCCCC |
| 504 | EEEARKFLEKYNQESEDMRYQQQGFGKGHDA | -47.48 | 51.6 | HHHHHHHHHHHHHHHHHHHHHHHHCCCCCCC |
| 712 | EEEARKFLEKYNQESEDMRYQRYGFGKGHDA | -47.48 | 51.6 | HHHHHHHHHHHHHHHHHHHHHHHCCCCCCCC |
| 995 | EEELKKFLEKFNQEYEDMFYQQKGFGKGHDA | -47.48 | 58.1 | HHHHHHHHHHHHHHHHHHHHHHHHCCCCCCC |
| 704 | EEELKKFLEKLNQEEEDIRYQQQGFGKGHDA | -47.48 | 51.6 | HHHHHHHHHHHHHHHHHHHHHHHHCCCCCCC |
| 551 | EEELKKFLEKYNQEAEDIYYQQNGFGKGHDA | -47.48 | 54.8 | HHHHHHHHHHHHHHHHHHHHHHHCCCCCCCC |
| 865 | EEELKKFLEKYNQEEEDMRYQREGFGKGHDA | -47.48 | 51.6 | HHHHHHHHHHHHHHHHHHHHHHHHHCCCCCC |
| 648 | EEELKKFLEKYNQESEDIRYQQNGFGKGHDA | -47.48 | 51.6 | HHHHHHHHHHHHHHHHHHHHHHHCCCCCCCC |
| 259 | EEELRKLLEKINQEEEDMRYQQYGFGKGHDA | -47.45 | 45.2 | HHHHHHHHHHHHHHHHHHHHHHHCCCCCCCC |
| 874 | EEEAKKFLEKYNQEEEDMYYQQKGFGKGHDA | -47.43 | 54.8 | HHHHHHHHHHHHHHHHHHHHHHHHCCCCCCC |
| 687 | EEEARKFLEKYNNEAEDIRFQQNGFGKGKSA | -47.42 | 51.6 | HHHHHHHHHHHHHHHHHHHHHHHCCCCCCCC |

| | | | | |
|---|---|---|---|---|
| 291 | EEELKKFLEKYNNEAEDIYFQQNGFGKGKDA | -47.42 | 51.6 | HHHHHHHHHHHHHHHHHHHHHHHCCCCCCCC |
| 965 | EEELKKFLEKYNNESEDMRFQQQGFGKGKDA | -47.42 | 48.4 | HHHHHHHHHHHHHHHHHHHHHHHHCCCCCCC |
| 553 | EEELKKFLEKLNNEEEDMRYQQQGFGKGHDA | -47.37 | 51.6 | HHHHHHHHHHHHHHHHHHHHHHHHCCCCCCC |
| 216 | EEELKKFLEKYNNESEDIRYQQNGFGKGHDA | -47.37 | 51.6 | HHHHHHHHHHHHCHHHHHHHHHHCCCCCCCC |
| 057 | EEELRKLLEQINNEAEDRTFQQNGFGKGKDA | -47.33 | 41.9 | HHHHHHHHHHHHHHHHHHHHHHHCCCCCCCC |
| 323 | EEELKKLLEQINQEEEDMRYQQQGFGKGHDA | -47.30 | 45.2 | HHHHHHHHHHHHHHHHHHHHHHHHCCCCCCC |
| 164 | EEELKKLLEQINQEAEDMRYQQQGFGKGHDA | -47.28 | 48.4 | HHHHHHHHHHHHHHHHHHHHHHHHCCCCCCC |
| 892 | EEELKKLLEQINQEEEDMRYQREGFGKGHDA | -47.28 | 45.2 | HHHHHHHHHHHHHHHHHHHHHHHHHCCCCCC |
| 552 | EEELKKLLEQLNQEEEDRRYQQNGFGKGHDA | -47.28 | 45.2 | HHHHHHHHHHHHHHHHHHHHHHHCCCCCCCC |
| 675 | EEELKKLLEQLNQEKEDMRYQQQGFGKGHDA | -47.28 | 45.2 | HHHHHHHHHHHHHHHHHHHHHHHHCCCCCCC |
| 668 | EEELKKLLEQLNQEQEDMRYQQNGFGKGHDA | -47.28 | 45.2 | HHHHHHHHHHHHHHHHHHHHHHHHCCCCCCC |
| 158 | EEELKKLLEQLNQEQEDMRYQRQGFGKGHDA | -47.28 | 45.2 | HHHHHHHHHHHHHHHHHHHHHHHHCCCCCCC |
| 197 | EEELRKLLEQINQEAEDIRYQQNGFGKGHDA | -47.28 | 45.2 | HHHHHHHHHHHHHHHHHHHHHHHHCCCCCCC |
| 559 | EEELRKLLEQINQEEEDMFYQQKGFGKGHSA | -47.28 | 45.2 | HHHHHHHHHHHHHHHHHHHHHHHHCCCCCCC |
| 973 | EEELRKLLEQINQEEEDMRYQQQGFGKGHDA | -47.28 | 41.9 | HHHHHHHHHHHHHHHHHHHHHHHHCCCCCCC |
| 889 | EEELRKLLEQINQEEEDMRYQREGFGKGHDA | -47.28 | 41.9 | HHHHHHHHHHHHHHHHHHHHHHHHHCCCCCC |
| 795 | EEELRKLLEQINQEEEDMRYQRYGFGKGHDA | -47.28 | 41.9 | HHHHHHHHHHHHHHHHHHHHHHHCCCCCCCC |
| 459 | EEELRKLLEQLNQEKEDMSYQQQGFGKGHDA | -47.28 | 41.9 | HHHHHHHHHHHHHHHHHHHHHHHCCCCCCCC |
| 610 | EEELRKLLEQLNQEQEDMRYQQNGFGKGHDA | -47.28 | 41.9 | HHHHHHHHHHHHHHHHHHHHHHHHCCCCCCC |
| 398 | EQELRKLLEQINQEEEDMRYQQQGFGKGHDA | -47.28 | 38.7 | HHHHHHHHHHHHHHHHHHHHHHHHCCCCCCC |
| 458 | EEELRKILEQFNNEEEDMYYQQYGFGKGHDA | -47.25 | 45.2 | HHHHHHHHHHHHHHHHHHHHHHHCCCCCCCC |
| 426 | EEELKKLLEQINNEAEDMRFQKEGFGKGKDA | -47.22 | 45.2 | HHHHHHHHHHHHHHHHHHHHHHHHCCCCCCC |
| 605 | EEELRKLLEQINNEEEDMRFQQQGFGKGKDA | -47.22 | 38.7 | HHHHHHHHHHHHHHHHHHHHHHHHCCCCCCC |
| 757 | EEELRKLLEQLNNEQEDMRFQQQGFGKGKDA | -47.22 | 38.7 | HHHHHHHHHHHHHHHHHHHHHHHHCCCCCCC |
| 393 | EEELKKLLEKINNEEEDIRYQQQGFGKGDSR | -47.20 | 54.8 | HHHHHHHHHHHHHHHHHHHHHHHCCCCCCCC |
| 794 | EEELKKLLEKLNNEQEDIRYQQNGFGKGDSR | -47.20 | 54.8 | HHHHHHHHHHHHHHHHHHHHHHHCCCCCCCC |
| 626 | EEELRKILEQYNNEEEDMRYQREGFGKGHDA | -47.20 | 41.9 | HHHHHHHHHHHCHHHHHHHHHHHHHCCCCCC |
| 825 | EEEWKKFLEKLNNEAEDMQYQRAGFGKGHDA | -47.19 | 54.8 | HHHHHHHHHHHHHHHHHHHHHHHHHCCCCCC |
| 520 | EEEARKFLEKFNNEAEDMQYQRAGFGKGHDA | -47.19 | 58.1 | HHHHHHHHHHHHHHHHHHHHHHHHHCCCCCC |
| 894 | EEEARKFLEKFNNEEEDMSYQQQGFGKGHDA | -47.19 | 54.8 | HHHHHHHHHHHHHHHHHHHHHHHCCCCCCCC |
| 441 | EEEARKFLEKFNNEYEDMFYQQKGFGKGHDA | -47.19 | 58.1 | HHHHHHHHHHHHHHHHHHHHHHHHCCCCCCC |
| 971 | EEEARKFLEKFNNEYEDMFYQQQGFGKGHDA | -47.19 | 58.1 | HHHHHHHHHHHHHHHHHHHHHHHHCCCCCCC |
| 240 | EEEARKFLEKLNNEEEDIRYQQQGFGKGHDA | -47.19 | 51.6 | HHHHHHHHHHHHHHHHHHHHHHHHCCCCCCC |
| 770 | EEEARKFLEKLNNEEEDMRYQQQGFGKGHDA | -47.19 | 51.6 | HHHHHHHHHHHHHHHHHHHHHHHHCCCCCCC |
| 400 | EEEARKFLEKYNNEAEDIRYQQGGFGKGHDA | -47.19 | 54.8 | HHHHHHHHHHHHHHHHHHHHHHCCCCCCCCC |
| 753 | EEEARKFLEKYNNEAEDIRYQQQGFGKGHDA | -47.19 | 54.8 | HHHHHHHHHHHHHHHHHHHHHHHHCCCCCCC |
| 406 | EEEARKFLEKYNNEAEDIYYQQNGFGKGHDA | -47.19 | 54.8 | HHHHHHHHHHHHHHHHHHHHHHHCCCCCCCC |
| 994 | EEEARKFLEKYNNEAEDMRYQQQGFGKGHDA | -47.19 | 54.8 | HHHHHHHHHHHHHHHHHHHHHHHHCCCCCCC |

| | | | | |
|---|---|---|---|---|
| 569 | EEEARKFLEKYNNESEDIRYQQGGFGKGHDA | -47.19 | 51.6 | HHHHHHHHHHHCHHHHHHHHHCCCCCCCCCC |
| 683 | EEEARKFLEKYNNESEDIRYQQQGFGKGHDA | -47.19 | 51.6 | HHHHHHHHHHHCHHHHHHHHHHHHCCCCCCC |
| 131 | EEEARKFLEKYNNESEDIRYQQQGFGKGHSA | -47.19 | 51.6 | HHHHHHHHHHHCHHHHHHHHHHHHCCCCCCC |
| 980 | EEELKKFLEKFNNEYEDIFYQQKGFGKGHDA | -47.19 | 58.1 | HHHHHHHHHHHHHHHHHHHHHHHHCCCCCCC |
| 546 | EEELKKFLEKFNNEYEDIFYQQQGFGKGHDA | -47.19 | 58.1 | HHHHHHHHHHHHHHHHHHHHHHHCCCCCCCC |
| 089 | EEELKKFLEKFNNEYEDMFYQQKGFGKGHDA | -47.19 | 58.1 | HHHHHHHHHHHHHHHHHHHHHHHHCCCCCCC |
| 620 | EEELKKFLEKFNNEYEDMFYQQKGFGKGHSA | -47.19 | 58.1 | HHHHHHHHHHHHHHHHHHHHHHHHCCCCCCC |
| 126 | EEELKKFLEKLNNEEEDMRYQREGFGKGHDA | -47.19 | 51.6 | HHHHHHHHHHHHHHHHHHHHHHHHHCCCCCC |
| 207 | EEELKKFLEKYNNEAEDIRYQQNGFGKGHDA | -47.19 | 54.8 | HHHHHHHHHHHHHHHHHHHHHHHHCCCCCCC |
| 144 | EEELKKFLEKYNNEAEDIRYQQQGFGKGHDA | -47.19 | 54.8 | HHHHHHHHHHHHHHHHHHHHHHHHCCCCCCC |
| 961 | EEELKKFLEKYNNEAEDIYYQQNGFGKGHDA | -47.19 | 54.8 | HHHHHHHHHHHHHHHHHHHHHHHCCCCCCCC |
| 976 | EEELKKFLEKYNNEAEDIYYQQNGFGKGHSA | -47.19 | 54.8 | HHHHHHHHHHHHHHHHHHHHHHHCCCCCCCC |
| 409 | EEELKKFLEKYNNEAEDMRYQQQGFGKGHDA | -47.19 | 54.8 | HHHHHHHHHHHHHHHHHHHHHHHHCCCCCCC |
| 008 | EEELKKFLEKYNNEEEDMRYQREGFGKGHDA | -47.19 | 51.6 | HHHHHHHHHHHCCHHHHHHHHHHHHCCCCCC |
| 984 | EEELKKFLEKYNNEEEDMSYQRYGFGKGHDA | -47.19 | 51.6 | HHHHHHHHHHHHHHHHHHHHHHCCCCCCCCC |
| 090 | EEELKKFLEKYNNESEDIRYQQGGFGKGHDF | -47.19 | 51.6 | HHHHHHHHHHHHCHHHHHHHHCCCCCCCCCC |
| 442 | EEELKKFLEKYNNESEDIRYQQQGFGKGHDA | -47.19 | 51.6 | HHHHHHHHHHHHCHHHHHHHHHHHCCCCCCC |
| 803 | EEELKKFLEKYNNEYEDIFYQQKGFGKGHDA | -47.19 | 54.8 | HHHHHHHHHHHHHHHHHHHHHHHHCCCCCCC |
| 310 | EEELKKFLEKYNNEYEDMEYQRNGRGKGHDY | -47.19 | 51.6 | HHHHHHHHHHHHHHHHHHHHHHHCCCCCCCC |
| 784 | EEELKKFLEKYNNEYEDMFYQQKGFGKGHDA | -47.19 | 54.8 | HHHHHHHHHHHHHHHHHHHHHHHHCCCCCCC |
| 129 | EEEWKKFIEKYNNEAEDIYYQQNGFGKGHDA | -47.19 | 51.6 | HHHHHHHHHHHHHHHHHHHHHHHCCCCCCCC |
| 117 | EQEARKFLEKYNNESEDIRYQQGGFGKGHDF | -47.19 | 48.4 | HHHHHHHHHHHHHHHHHHHHHCCCCCCCCCC |
| 989 | EEEARKFLEKFNQEYEDMFYQQKGFGKGHDA | -47.16 | 58.1 | HHHHHHHHHHHHHHHHHHHHHHHHCCCCCCC |
| 755 | EEELKKLLEKLNQEQEDMYYQQKGFGKGDSR | -47.15 | 54.8 | HHHHHHHHHHHHHHHHHHHHHHHCCCCCCCC |
| 283 | EEELKKLLEQINQEAEDMRYQQNGFGKGHDA | -47.04 | 48.4 | HHHHHHHHHHHHHHHHHHHHHHHHCCCCCCC |
| 587 | EEEARKFLEKYNQEAEQIRYQQQGFGRGHDA | -47.03 | 48.4 | HHHHHHHHHHHHHHHHHHHHHHHCCCCCCCC |
| 289 | EEELKKLLEQINNEEEDMRYQQQGFGKGHDA | -47.00 | 45.2 | HHHHHHHHHHHHHHHHHHHHHHHHCCCCCCC |
| 431 | EEELKKLLEQINNEEEDMRYQREGFGKGHDA | -47.00 | 45.2 | HHHHHHHHHHHHHHHHHHHHHHHHHCCCCCC |
| 530 | EEELRKLLEQINNEKEDMEYQRAGFGKGHDA | -47.00 | 41.9 | HHHHHHHHHHHHHHHHHHHHHHHHCCCCCCC |
| 066 | EEELKKFLEKYNQEAENIYKQQNGFGSGKDA | -46.99 | 45.2 | HHHHHHHHHHHHHHHHHHHHHHHCCCCCCCC |
| 108 | EEELKKLLEQINNEAEDIRYQQNGFGKGHDA | -46.99 | 48.4 | HHHHHHHHHHHHHHHHHHHHHHHHCCCCCCC |
| 109 | EEELKKLLEQINNEAEDIRYQQNGFGKGHSA | -46.99 | 48.4 | HHHHHHHHHHHHHHHHHHHHHHHHCCCCCCC |
| 529 | EEELKKLLEQINNEAEDMRYQQNGFGKGHDA | -46.99 | 48.4 | HHHHHHHHHHHHHHHHHHHHHHHHCCCCCCC |
| 831 | EEELKKLLEQINNEAEDMRYQQQGFGKGHDA | -46.99 | 48.4 | HHHHHHHHHHHHHHHHHHHHHHHHCCCCCCC |
| 843 | EEELKKLLEQINNEEEDMRYQQQGFGKGHSA | -46.99 | 45.2 | HHHHHHHHHHHHHHHHHHHHHHHHCCCCCCC |
| 058 | EEELKKLLEQINNEEEDMRYQQYGFGKGHDA | -46.99 | 45.2 | HHHHHHHHHHHHHHHHHHHHHHHCCCCCCCC |
| 014 | EEELKKLLEQINNEKEDMEYQRAGFGKGHDA | -46.99 | 45.2 | HHHHHHHHHHHHHHHHHHHHHHHHCCCCCCC |
| 966 | EEELKKLLEQLNNEEEDKQYQRAGFGKGHDA | -46.99 | 45.2 | HHHHHHHHHHHHHHHHHHHHHHHHCCCCCCC |

| | | | | |
|---|---|---|---|---|
| 223 | EEELKKLLEQLNNEEEDMKYQQNGFGKGHDA | -46.99 | 45.2 | HHHHHHHHHHHHHHHHHHHHHHHHCCCCCCC |
| 281 | EEELKKLLEQLNNEEEDRRYQQGGFGKGHDA | -46.99 | 45.2 | HHHHHHHHHHHHHHHHHHHHHHCCCCCCCCC |
| 438 | EEELKKLLEQLNNEKEDMRYQQQGFGKGHDA | -46.99 | 45.2 | HHHHHHHHHHHHHHHHHHHHHHHHCCCCCCC |
| 991 | EEELKKLLEQLNNEKEDMSYQQQGFGKGHDA | -46.99 | 45.2 | HHHHHHHHHHHHHHHHHHHHHHHCCCCCCCC |
| 226 | EEELKKLLEQLNNEQEDIRYQQGGFGKGHDF | -46.99 | 45.2 | HHHHHHHHHHHHHHHHHHHHHHCCCCCCCCC |
| 955 | EEELKKLLEQLNNEQEDMRYQQNGFGKGHDA | -46.99 | 45.2 | HHHHHHHHHHHHHHHHHHHHHHHHCCCCCCC |
| 192 | EEELKKLLEQLNNEQEDMRYQQQGFGKGHDA | -46.99 | 45.2 | HHHHHHHHHHHHHHHHHHHHHHHHCCCCCCC |
| 368 | EEELKKLLEQLNNEQEDMRYQRQGFGKGHDA | -46.99 | 45.2 | HHHHHHHHHHHHHHHHHHHHHHHHCCCCCCC |
| 815 | EEELRKLLEQINNEAEDMQYQRAGFGKGHDA | -46.99 | 45.2 | HHHHHHHHHHHHHHHHHHHHHHHHHCCCCCC |
| 203 | EEELRKLLEQINNEAEDMRYQQNGFGKGHDA | -46.99 | 45.2 | HHHHHHHHHHHHHHHHHHHHHHHHCCCCCCC |
| 210 | EEELRKLLEQINNEAEDMSYQKEGFGKGHDA | -46.99 | 45.2 | HHHHHHHHHHHHHHHHHHHHHHHHCCCCCCC |
| 187 | EEELRKLLEQINNEEEDMRYQQQGFGKGHDA | -46.99 | 41.9 | HHHHHHHHHHHHHHHHHHHHHHHHCCCCCCC |
| 399 | EEELRKLLEQINNEEEDMRYQQYGFGKGHDA | -46.99 | 41.9 | HHHHHHHHHHHHHHHHHHHHHHHCCCCCCCC |
| 201 | EEELRKLLEQINNEEEDMRYQREGFGKGHDA | -46.99 | 41.9 | HHHHHHHHHHHHHHHHHHHHHHHHHCCCCCC |
| 484 | EEELRKLLEQINNEKEDMEYQRAGFGKGHSA | -46.99 | 41.9 | HHHHHHHHHHHHHHHHHHHHHHHHHCCCCCC |
| 295 | EEELRKLLEQLNNEEEDMKYQKEGFGKGHDA | -46.99 | 41.9 | HHHHHHHHHHHHHHHHHHHHHHHHHCCCCCC |
| 649 | EEELRKLLEQLNNEEEDMKYQQNGFGKGHDA | -46.99 | 41.9 | HHHHHHHHHHHHHHHHHHHHHHHHCCCCCCC |
| 483 | EEELRKLLEQLNNEEEDMRYQQNGFGKGHDA | -46.99 | 41.9 | HHHHHHHHHHHHHHHHHHHHHHHHCCCCCCC |
| 159 | EEELRKLLEQLNNEKEDMSYQQQGFGKGHDA | -46.99 | 41.9 | HHHHHHHHHHHHHHHHHHHHHHHCCCCCCCC |
| 983 | EEELRKLLEQLNNEQEDMRYQQNGFGKGHDA | -46.99 | 41.9 | HHHHHHHHHHHHHHHHHHHHHHHHCCCCCCC |
| 196 | EEELRKLLEQLNNEQEDMRYQQQGFGKGHDA | -46.99 | 41.9 | HHHHHHHHHHHHHHHHHHHHHHHHCCCCCCC |
| 284 | EEELRKLLEQLNNEQEDMRYQRYGFGKGHDA | -46.99 | 41.9 | HHHHHHHHHHHHHHHHHHHHHHHCCCCCCCC |
| 402 | EEELKKFLEKFNNEYEDMYYQQQGFGKGHDA | -46.95 | 54.8 | HHHHHHHHHHHHHHHHHHHHHHHHCCCCCCC |
| 044 | EEELKKFLEQYNQEAEDMRFQQGGFGKGKDA | -46.95 | 48.4 | HHHHHHHHHHHHHHHHHHHHHHCCCCCCCCC |
| 902 | EEELKKFLEKLNQEEEDMRYQQYGFGKGHDA | -46.89 | 51.6 | HHHHHHHHHHHHHHHHHHHHHHHCCCCCCCC |
| 848 | EEEARKFLEKYNNEAEQIYYQQNGFGRGHDA | -46.83 | 48.4 | HHHHHHHHHHHHHHHHHHHHHHHCCCCCCCC |
| 723 | EEELKKLLDKINNEEEQIRYQQQGFGQGHDA | -46.80 | 45.2 | HHHHHHHHHHHHHHHHHHHHHHHCCCCCCCC |
| 065 | EEELKKLLEKINQEAEDIRYQQQGFGKGDSR | -46.80 | 58.1 | HHHHHHHHHHHHHHHHHHHHHHHCCCCCCCC |
| 561 | EEELRKLLEKINQEEEDIRYQQQGFGKGDSR | -46.80 | 51.6 | HHHHHHHHHHHHHHHHHHHHHHHCCCCCCCC |
| 329 | EEEARKFLEKFNQEYEDMFYQQKGFGKGDSR | -46.75 | 64.5 | HHHHHHHHHHHHHHHHHHHHHHHCCCCCCCC |
| 796 | EEEARKFLEKYNQEAEDMRYQRQGFGKGDSR | -46.75 | 61.3 | HHHHHHHHHHHHHHHHHHHHHHHCCCCCCCC |
| 935 | EEELKKFLEKYNQEAEDIRYQQNGFGKGDSR | -46.75 | 61.3 | HHHHHHHHHHHHHHHHHHHHHHHCCCCCCCC |
| 296 | EEEAKKFLEQYNQEYEDMFYQQKGFGKGHDA | -46.72 | 54.8 | HHHHHHHHHHHHHHHHHHHHHHHHCCCCCCC |
| 088 | EEEARKFLEQFNQEYEDMFYQQKGFGKGHDA | -46.72 | 54.8 | HHHHHHHHHHHHHHHHHHHHHHHHCCCCCCC |
| 855 | EEEARKFLEQFNQEYEDMFYQQKGFGKGHSA | -46.72 | 54.8 | HHHHHHHHHHHHHHHHHHHHHHHHCCCCCCC |
| 920 | EEEARKFLEQFNQEYEDMYYQQQGFGKGHDA | -46.72 | 51.6 | HHHHHHHHHHHHHHHHHHHHHHHHCCCCCCC |
| 969 | EEEARKFLEQLNQEEEDMRYQQQGFGKGHSA | -46.72 | 48.4 | HHHHHHHHHHHHHHHHHHHHHHHHCCCCCCC |
| 506 | EEEARKFLEQLNQEEEDMRYQQYGFGKGHDA | -46.72 | 48.4 | HHHHHHHHHHHHHHHHHHHHHHHCCCCCCCC |

| | | | | |
|---|---|---|---|---|
| 345 | EEEARKFLEQYNQEAEDIYYQQNGFGKGHDA | -46.72 | 51.6 | HHHHHHHHHHHHHHHHHHHHHHHCCCCCCCC |
| 437 | EEEARKFLEQYNQEAEDMRYQQNGFGKGHDA | -46.72 | 51.6 | HHHHHHHHHHHHHHHHHHHHHHHHCCCCCCC |
| 171 | EEEARKFLEQYNQEAEDMRYQREGFGKGHDA | -46.72 | 51.6 | HHHHHHHHHHHHHHHHHHHHHHHHHCCCCCC |
| 248 | EEEARKFLEQYNQEEEDMRYQREGFGKGHDA | -46.72 | 48.4 | HHHHHHHHHHHHHHHHHHHHHHHHHCCCCCC |
| 858 | EEEARKFLEQYNQESEDIRYQQNGFGKGHDA | -46.72 | 48.4 | HHHHHHHHHHHHHHHHHHHHHHHCCCCCCCC |
| 766 | EEEARKFLEQYNQESEDIRYQQQGFGKGHDA | -46.72 | 48.4 | HHHHHHHHHHHHHHHHHHHHHHHHCCCCCCC |
| 279 | EEEARKFLEQYNQESEDMRYQQQGFGKGHDA | -46.72 | 48.4 | HHHHHHHHHHHHHHHHHHHHHHHHCCCCCCC |
| 360 | EEELKKFLEQLNQEAEDMRYQQNGFGKGHDA | -46.72 | 51.6 | HHHHHHHHHHHHHHHHHHHHHHHHCCCCCCC |
| 321 | EEELKKFLEQYNQEAEDIRYQQQGFGKGHDA | -46.72 | 51.6 | HHHHHHHHHHHHHHHHHHHHHHHHCCCCCCC |
| 083 | EEELKKFLEQYNQEEEDMRYQREGFGKGHSA | -46.72 | 48.4 | HHHHHHHHHHHHHHHHHHHHHHHHHCCCCCC |
| 948 | EEELKKFLEQYNQESEDMRYQQQGFGKGHDA | -46.72 | 48.4 | HHHHHHHHHHHHHHHHHHHHHHHHCCCCCCC |
| 143 | EEELKKFLEQYNQEYEDMFYQQQGFGKGHSA | -46.72 | 51.6 | HHHHHHHHHHHHHHHHHHHHHHHHCCCCCCC |
| 715 | EEELRKFLEQFNQEYEDMFYQQKGFGKGHDA | -46.72 | 51.6 | HHHHHHHHHHHHHHHHHHHHHHHHCCCCCCC |
| 808 | EEELRKLLEKINNEEEQMRYQREGFGRGDSR | -46.71 | 45.2 | HHHHHHHHHHHCHHHHHHHHHHHCCCCCCCC |
| 769 | EEELKKLLEQLNQEEEDMKYQQNGFGKGHDA | -46.68 | 45.2 | HHHHHHHHHHHHHHHHHHHHHHHHCCCCCCC |
| 069 | EEEARKFLEKYNNEAEDIRYQQNGFGKGHDA | -46.67 | 54.8 | HHHHHHHHHHHHHHHHHHHHHHHHCCCCCCC |
| 156 | EEEAKKFLEQYNNEEEDMFFQQKGFGKGKDA | -46.66 | 51.6 | HHHHHHHHHHHHHHHHHHHHHHHHCCCCCCC |
| 667 | EEEARKFLEQFNNEEEDMQFQRSGFGKGKDA | -46.66 | 51.6 | HHHHHHHHHHHHHHHHHHHHHHHHCCCCCCC |
| 601 | EEEARKFLEQFNNEYEDMFFQQKGFGKGKDA | -46.66 | 51.6 | HHHHHHHHHHHHHHHHHHHHHHHHCCCCCCC |
| 468 | EEEARKFLEQYNNEAEDMRFQQQGFGKGKDA | -46.66 | 48.4 | HHHHHHHHHHHHHHHHHHHHHHHHCCCCCCC |
| 374 | EEELKKFLEKFNNEYEDMFYQQKGFGKGDSR | -46.64 | 64.5 | HHHHHHHHHHHHHHHHHHHHHHCCCCCCCCC |
| 634 | EEELKKFLEQYNNEAEDMRYQQQGFGKGSDY | -46.63 | 51.6 | HHHHHHHHHHHHHHHHHHHHHHHHCCCCCCC |
| 016 | EEQWKKLIEQINNEKEDMEYQRAGFGKGHDA | -46.55 | 45.2 | HHHHHHHHHHHHHHHHHHHHHHHHCCCCCCC |
| 749 | EEELKKLLEQINQEAEDMQYQRAGFGKGDSR | -46.54 | 54.8 | HHHHHHHHHHHHHHHHHHHHHHHHCCCCCCC |
| 046 | EEELKKLLEQLNQEQEDMRYQQNGFGKGDSR | -46.54 | 51.6 | HHHHHHHHHHHHHHHHHHHHHHHCCCCCCCC |
| 876 | EEELRKLLEQLNQEQEDMRYQQNGFGKGDSR | -46.54 | 48.4 | HHHHHHHHHHHHHHHHHHHHHHHCCCCCCCC |
| 568 | EESLKKFLDQYNNESEQMRYQQNGFGRGHDA | -46.54 | 45.2 | HHHHHHHHHHHHHHHHHHHHHHHHCCCCCCC |
| 740 | EEELKKILEQINNEEEDMRYQQQGFGKGDSR | -46.52 | 51.6 | HHHHHHHHHHHHHHHHHHHHHHHCCCCCCCC |
| 273 | EEELKKILEQLNNEQEDMRYQQNGFGKGDSR | -46.52 | 51.6 | HHHHHHHHHHHHHHHHHHHHHHHCCCCCCCC |
| 339 | EEELKKLLEKINNEAEDIRYQQQGFGKGDSR | -46.51 | 58.1 | HHHHHHHHHHHHHHHHHHHHHHHCCCCCCCC |
| 152 | EEELKKLLEKINNEAEDMRYQQQGFGKGDSR | -46.51 | 58.1 | HHHHHHHHHHHHHHHHHHHHHHHCCCCCCCC |
| 761 | EEELKKLLEKLNNEQEDMRYQQQGFGKGDSR | -46.51 | 54.8 | HHHHHHHHHHHHHHHHHHHHHHHCCCCCCCC |
| 049 | EEELRKLLEKINNEAEDIRYQQNGFGKGDSR | -46.51 | 54.8 | HHHHHHHHHHHHHHHHHHHHHHHCCCCCCCC |
| 186 | EEELRKLLEKINNEAEDMRYQQQGFGKGDSR | -46.51 | 54.8 | HHHHHHHHHHHHHHHHHHHHHHHCCCCCCCC |
| 899 | EEELRKLLEKINNEAEDMRYQREGFGKGDSR | -46.51 | 54.8 | HHHHHHHHHHHHHHHHHHHHHHHHCCCCCCC |
| 004 | EEELRKLLEKLNNEEEDMKYQQGGFGKGDSR | -46.51 | 51.6 | HHHHHHHHHHHHHHHHHHHHHHCCCCCCCCC |
| 691 | EEELRKLLEKLNNEKEDMSYQQQGFGKGDSR | -46.51 | 51.6 | HHHHHHHHHHHHHHHHHHHHHHHCCCCCCCC |
| 050 | EEELKKFLEQFNQEYEDMFYQQKGFGKGHDA | -46.50 | 54.8 | HHHHHHHHHHHHHHHHHHHHHHHHCCCCCCC |

| | | | | |
|---|---|---|---|---|
| 738 | EEELKKFLEQLNQEEEDMRYQQQGFGKGHSA | -46.48 | 48.4 | HHHHHHHHHHHHHHHHHHHHHHHHCCCCCCC |
| 252 | EEEAKKMLEKINNEEEDIRYQQQGFGKGHDA | -46.47 | 51.6 | HHHHHHHHHHHHHHHHHHHHHHHHCCCCCCC |
| 521 | EEEARKFLEKFNNEYEDMFYQQKGFGKGDSR | -46.45 | 64.5 | HHHHHHHHHHHHHHHHHHHHHHCCCCCCCCC |
| 507 | EEEARKFLEKYNNEEEDMRYQRQGFGKGDDR | -46.45 | 58.1 | HHHHHHHHHHHCHHHHHHHHHHHCCCCCCCC |
| 377 | EEEARKFLEKYNNESEDIRYQQQGFGKGDSR | -46.45 | 58.1 | HHHHHHHHHHHCHHHHHHHHHHHCCCCCCCC |
| 175 | EEELKKFLEKYNNEAEDIYYQQNGFGKGDSR | -46.45 | 61.3 | HHHHHHHHHHHHHHHHHHHHHHHCCCCCCCC |
| 891 | EEELKKFLEKYNNESEDIRYQQNGFGKGDSR | -46.45 | 58.1 | HHHHHHHHHHHHCHHHHHHHHHHCCCCCCCC |
| 531 | EEELKKFLEKYNNEYEDMFYQQKGFGKGDSR | -46.45 | 61.3 | HHHHHHHHHHHHHHHHHHHHHHCCCCCCCCC |
| 417 | EEEARKFLEQFNNEYEDMFYQQKGFGKGHDA | -46.44 | 54.8 | HHHHHHHHHHHHHHHHHHHHHHHHCCCCCCC |
| 452 | EEEARKFLEQYNNEEEDMRYQRQGFGKGHDA | -46.44 | 48.4 | HHHHHHHHHHHCHHHHHHHHHHHHCCCCCCC |
| 318 | EEELKKFLEQYNNEAEDMRYQQQGFGKGHDA | -46.44 | 51.6 | HHHHHHHHHHHHHHHHHHHHHHHHCCCCCCC |
| 225 | EEEWRKFLEQLNNEEEDMRYQQQGFGKGHSA | -46.43 | 45.2 | HHHHHHHHHHHHHHHHHHHHHHHHCCCCCCC |
| 023 | EEEWRKFLEQLNNEEEDMRYQREGFGKGHDA | -46.43 | 45.2 | HHHHHHHHHHHHHHHHHHHHHHHHHCCCCCC |
| 370 | EEEAKKFLEQFNNEYEDMFYQQKGFGKGHSA | -46.42 | 58.1 | HHHHHHHHHHHHHHHHHHHHHHHHCCCCCCC |
| 586 | EEEAKKFLEQFNNEYEDMFYQQQGFGKGHDA | -46.42 | 58.1 | HHHHHHHHHHHHHHHHHHHHHHHHCCCCCCC |
| 963 | EEEAKKFLEQYNNEAEDIRYQQGGFGKGHDY | -46.42 | 54.8 | HHHHHHHHHHHHHHHHHHHHHHCCCCCCCCC |
| 905 | EEEAKKFLEQYNNEAEDIYYQQNGFGKGHDA | -46.42 | 54.8 | HHHHHHHHHHHHHHHHHHHHHHHCCCCCCCC |
| 832 | EEEAKKFLEQYNNESEDIRYQQNGFGKGHDA | -46.42 | 51.6 | HHHHHHHHHHHHHHHHHHHHHHHCCCCCCCC |
| 222 | EEEAKKFLEQYNNESEDIRYQQQGFGKGHDA | -46.42 | 51.6 | HHHHHHHHHHHHHHHHHHHHHHHHCCCCCCC |
| 937 | EEEAKKFLEQYNNEYEDMFYQQQGFGKGHDA | -46.42 | 54.8 | HHHHHHHHHHHHHHHHHHHHHHHHCCCCCCC |
| 919 | EEEARKFLEQFNNEKEDMRYQQQGFGKGHDA | -46.42 | 51.6 | HHHHHHHHHHHHHHHHHHHHHHHHCCCCCCC |
| 932 | EEEARKFLEQLNNEEEDMQYQRSGFGKGHDA | -46.42 | 51.6 | HHHHHHHHHHHHHHHHHHHHHHHHCCCCCCC |
| 637 | EEEARKFLEQLNNEEEDMRYQQYGFGKGHDA | -46.42 | 48.4 | HHHHHHHHHHHHHHHHHHHHHHHCCCCCCCC |
| 149 | EEEARKFLEQYNNEAEDIYYQQNGFGKGHDA | -46.42 | 51.6 | HHHHHHHHHHHHHHHHHHHHHHHCCCCCCCC |
| 555 | EEEARKFLEQYNNEAEDIYYQQNGFGKGHSA | -46.42 | 51.6 | HHHHHHHHHHHHHHHHHHHHHHHCCCCCCCC |
| 000 | EEEARKFLEQYNNEAEDMRYQQNGFGKGHDA | -46.42 | 51.6 | HHHHHHHHHHHHHHHHHHHHHHHHCCCCCCC |
| 112 | EEEARKFLEQYNNEAEDMRYQQNGFGKGHSA | -46.42 | 51.6 | HHHHHHHHHHHHHHHHHHHHHHHHCCCCCCC |
| 317 | EEEARKFLEQYNNEAEDMRYQQQGFGKGHDA | -46.42 | 51.6 | HHHHHHHHHHHHHHHHHHHHHHHHCCCCCCC |
| 621 | EEEARKFLEQYNNEAEDMRYQQQGFGKGHSA | -46.42 | 51.6 | HHHHHHHHHHHHHHHHHHHHHHHHCCCCCCC |
| 781 | EEEARKFLEQYNNEAEDMRYQRQGFGKGHDA | -46.42 | 51.6 | HHHHHHHHHHHHHHHHHHHHHHHHCCCCCCC |
| 081 | EEEARKFLEQYNNEEEDMFYQQKGFGKGHDA | -46.42 | 51.6 | HHHHHHHHHHHHHHHHHHHHHHHHCCCCCCC |
| 121 | EEEARKFLEQYNNEEEDMKYQQEGFGKGHDA | -46.42 | 48.4 | HHHHHHHHHHHHHHHHHHHHHHHHHCCCCCC |
| 047 | EEEARKFLEQYNNEEEDMRYQQQGFGKGHSA | -46.42 | 48.4 | HHHHHHHHHHHHHHHHHHHHHHHHCCCCCCC |
| 879 | EEEARKFLEQYNNEEEDMRYQREGFGKGHDA | -46.42 | 48.4 | HHHHHHHHHHHCHHHHHHHHHHHHHCCCCCC |
| 881 | EEEARKFLEQYNNESEDIRYQQGGFGKGHSA | -46.42 | 48.4 | HHHHHHHHHHHHHHHHHHHHHCCCCCCCCCC |
| 294 | EEEARKFLEQYNNESEDIRYQQNGFGKGHDA | -46.42 | 48.4 | HHHHHHHHHHHHHHHHHHHHHHHCCCCCCCC |
| 463 | EEEARKFLEQYNNESEDMRYQQNGFGKGHDA | -46.42 | 48.4 | HHHHHHHHHHHHHHHHHHHHHHHHCCCCCCC |
| 839 | EEEARKFLEQYNNESEDMRYQQNGFGKGHSA | -46.42 | 48.4 | HHHHHHHHHHHHHHHHHHHHHHHHCCCCCCC |

| | | | | |
|---|---|---|---|---|
| 772 | EEEARKFLEQYNNESEDMRYQQQGFGKGHDA | -46.42 | 48.4 | HHHHHHHHHHHHHHHHHHHHHHHHCCCCCCC |
| 525 | EEEARKFLEQYNNEYEDMEYQRSGFGKGHDA | -46.42 | 51.6 | HHHHHHHHHHHHHHHHHHHHHHHHCCCCCCC |
| 732 | EEEARKFLEQYNNEYEDMFYQQKGFGKGHDA | -46.42 | 51.6 | HHHHHHHHHHHHHHHHHHHHHHHHCCCCCCC |
| 235 | EEELKKFLEQFNNEAEDMRYQQQGFGKGHDA | -46.42 | 54.8 | HHHHHHHHHHHHHHHHHHHHHHHHCCCCCCC |
| 341 | EEELKKFLEQFNNEEEDMRYQREGFGKGHDA | -46.42 | 51.6 | HHHHHHHHHHHHHHHHHHHHHHHHHCCCCCC |
| 153 | EEELKKFLEQFNNEYEDMFYQKEGFGKGHDA | -46.42 | 54.8 | HHHHHHHHHHHHHHHHHHHHHHHHCCCCCCC |
| 166 | EEELKKFLEQFNNEYEDMFYQQKGFGKGHDA | -46.42 | 54.8 | HHHHHHHHHHHHHHHHHHHHHHHHCCCCCCC |
| 111 | EEELKKFLEQFNNEYEDMFYQQKGFGKGHSA | -46.42 | 54.8 | HHHHHHHHHHHHHHHHHHHHHHHHCCCCCCC |
| 650 | EEELKKFLEQFNNEYEDMYYQQKGFGKGHDA | -46.42 | 51.6 | HHHHHHHHHHHHHHHHHHHHHHHHCCCCCCC |
| 274 | EEELKKFLEQLNNEEEDMRYQQQGFGKGHDA | -46.42 | 48.4 | HHHHHHHHHHHHHHHHHHHHHHHHCCCCCCC |
| 600 | EEELKKFLEQLNNEEEDMRYQQQGFGKGHSA | -46.42 | 48.4 | HHHHHHHHHHHHHHHHHHHHHHHHCCCCCCC |
| 688 | EEELKKFLEQLNNEEEDMRYQQYGFGKGHDA | -46.42 | 48.4 | HHHHHHHHHHHHHHHHHHHHHHHCCCCCCCC |
| 103 | EEELKKFLEQLNNEEEDMRYQREGFGKGHDA | -46.42 | 48.4 | HHHHHHHHHHHHHHHHHHHHHHHHHCCCCCC |
| 516 | EEELKKFLEQYNNEAEDIRYQQNGFGKGHDA | -46.42 | 51.6 | HHHHHHHHHHHHHHHHHHHHHHHHCCCCCCC |
| 258 | EEELKKFLEQYNNEAEDIYYQQNGFGKGHDA | -46.42 | 51.6 | HHHHHHHHHHHHHHHHHHHHHHHCCCCCCCC |
| 033 | EEELKKFLEQYNNEAEDIYYQQQGFGKGHDA | -46.42 | 51.6 | HHHHHHHHHHHHHHHHHHHHHHHHCCCCCCC |
| 653 | EEELKKFLEQYNNEAEDMRYQQEGFGKGHDA | -46.42 | 51.6 | HHHHHHHHHHHHHHHHHHHHHHHHHCCCCCC |
| 763 | EEELKKFLEQYNNEAEDMRYQQGGFGKGHDA | -46.42 | 51.6 | HHHHHHHHHHHHHHHHHHHHHHCCCCCCCCC |
| 475 | EEELKKFLEQYNNEAEDMRYQQNGFGKGHDA | -46.42 | 51.6 | HHHHHHHHHHHHHHHHHHHHHHHHCCCCCCC |
| 288 | EEELKKFLEQYNNEAEDMRYQRYGFGKGHDA | -46.42 | 51.6 | HHHHHHHHHHHHHHHHHHHHHHHCCCCCCCC |
| 022 | EEELKKFLEQYNNEEEDMKYQQEGFGKGHSA | -46.42 | 48.4 | HHHHHHHHHHHHHHHHHHHHHHHHHCCCCCC |
| 043 | EEELKKFLEQYNNEEEDMRYQREGFGKGHDA | -46.42 | 48.4 | HHHHHHHHHHHCHHHHHHHHHHHHHCCCCCC |
| 594 | EEELKKFLEQYNNESEDMRYQQNGFGKGHDA | -46.42 | 48.4 | HHHHHHHHHHHHHHHHHHHHHHHHCCCCCCC |
| 508 | EEELKKFLEQYNNESEDMRYQQQGFGKGHDA | -46.42 | 48.4 | HHHHHHHHHHHHHHHHHHHHHHHHCCCCCCC |
| 800 | EEELKKFLEQYNNEYEDMEYQRSGFGKGHDA | -46.42 | 51.6 | HHHHHHHHHHHHHHHHHHHHHHHHCCCCCCC |
| 933 | EEELKKFLEQYNNEYEDMFYQQKGFGKGHDA | -46.42 | 51.6 | HHHHHHHHHHHHHHHHHHHHHHHHCCCCCCC |
| 071 | EEEWKKFIEQYNNEAEDIYYQQNGFGKGHDA | -46.42 | 48.4 | HHHHHHHHHHHHHHHHHHHHHHHCCCCCCCC |
| 070 | EQEARKFLEQFNNEQEDMTYQKEGFGKGHDA | -46.42 | 48.4 | HHHHHHHHHHHHHHHHHHHHHHHHCCCCCCC |
| 102 | EQEARKFLEQYNNEAEDMRYQQQGFGKGHDA | -46.42 | 48.4 | HHHHHHHHHHHHHHHHHHHHHHHHCCCCCCC |
| 533 | EQEARKFLEQYNNEYEDMFYQKEGFGKGHDA | -46.42 | 48.4 | HHHHHHHHHHHHHHHHHHHHHHHHCCCCCCC |
| 334 | EEELKKFLEQFNNEYEDMYFQQKGFGKGKDA | -46.39 | 48.4 | HHHHHHHHHHHHHHHHHHHHHHHHCCCCCCC |
| 663 | EEELRKLLDQINQEEEQMRYQREGFGQGHDA | -46.39 | 38.7 | HHHHHHHHHHHHHHHHHHHHHHHHHCCCCCC |
| 038 | EEELKKLLEKINNEEEDRRYQQYGFGKGHDA | -46.36 | 48.4 | HHHHHHHHHHHHHHHHHHHHHHHCCCCCCCC |
| 303 | EEELKKFLQKLNEEEEQMRYQQYGFGQGHDA | -46.35 | 45.2 | HHHHHHHHHHHHHHHHHHHHHHHCCCCCCCC |
| 491 | EEELRKLLEQINNEAEDMRYQQQGFGKGDSR | -46.27 | 51.6 | HHHHHHHHHHHHHHHHHHHHHHHCCCCCCCC |
| 366 | EEELRKLLEQINNEEEDMRYQQQGFGKGDSR | -46.27 | 48.4 | HHHHHHHHHHHHHHHHHHHHHHHCCCCCCCC |
| 188 | EEELKKLLEQINNEAEDMRYQQNGFGKGDSR | -46.25 | 54.8 | HHHHHHHHHHHHHHHHHHHHHHHCCCCCCCC |
| 141 | EEELKKLLEQLNNEEEDMKYQQQGFGKGDSR | -46.25 | 51.6 | HHHHHHHHHHHHHHHHHHHHHHHCCCCCCCC |

| | | | | |
|---|---|---|---|---|
| 975 | EEELKKLLEQLNNEQEDMRYQQNGFGKGDSR | -46.25 | 51.6 | HHHHHHHHHHHHHHHHHHHHHHHCCCCCCCC |
| 764 | EEELKKLLEQLNNEQEDMRYQQQGFGKGDSR | -46.25 | 51.6 | HHHHHHHHHHHHHHHHHHHHHHHCCCCCCCC |
| 896 | EEELKKLLEQLNNEQEDMRYQREGFGKGDSR | -46.25 | 51.6 | HHHHHHHHHHHHHHHHHHHHHHHHCCCCCCC |
| 233 | EEELRKLLEQLNNEQEDMRYQQNGFGKGDSR | -46.25 | 48.4 | HHHHHHHHHHHHHHHHHHHHHHHCCCCCCCC |
| 570 | EEELKKFLEKFNQEQEDMTYQRHGFGKGDSR | -46.24 | 61.3 | HHHHHHHHHHHHHHHHHHHHHHHCCCCCCCC |
| 774 | EEELKKFLEKYNQEAEDIRYQQQGFGKGDDR | -46.24 | 61.3 | HHHHHHHHHHHHHHHHHHHHHHHCCCCCCCC |
| 606 | EEELKKFLEKYNQEAEDMRYQKEGFGKGDSR | -46.24 | 61.3 | HHHHHHHHHHHHHHHHHHHHHHHHCCCCCCC |
| 351 | EEELRKFLEQYNNESEDMRYQQQGFGKGHSA | -46.21 | 45.2 | HHHHHHHHHHHHHHHHHHHHHHHHCCCCCCC |
| 389 | EEEWRKILEQLNNEQEDMRYQQNGFGKGDSR | -46.19 | 48.4 | HHHHHHHHHHHHHHHHHHHHHHHCCCCCCCC |
| 782 | EEELKKFLEQYNNDAEDMRYQRQGFGKGHDA | -46.13 | 48.4 | HHHHHHHHHHHHHHHHHHHHHHHHCCCCCCC |
| 950 | EEELKKFLEQYNQESEDIRYQQNGFGKGHDA | -46.12 | 48.4 | HHHHHHHHHHHHHHHHHHHHHHHCCCCCCCC |
| 655 | EEELKKILEKINNEAEQIRYQQNGFGQGHDA | -46.11 | 45.2 | HHHHHHHHHHHHHHHHHHHHHHHHCCCCCCC |
| 424 | EEELKKILEKLNNEQEQMQYQRAGFGQGHDA | -46.11 | 41.9 | HHHHHHHHHHHHHHHHHHHHHHHHHCCCCCC |
| 394 | EEELKKLLDQINNEAEQIRYQQNGFGQGHDA | -46.09 | 45.2 | HHHHHHHHHHHHHHHHHHHHHHHHCCCCCCC |
| 513 | EESLKKLLEQINQEAEDMFYQQKGFGKGHDA | -46.06 | 51.6 | HHHHHHHHHHHHHHHHHHHHHHHHCCCCCCC |
| 092 | EEELKKLLEQINQEEEDMRYQQQGFGKGDSR | -46.04 | 51.6 | HHHHHHHHHHHHHHHHHHHHHHHCCCCCCCC |
| 115 | EEELKKLLEQLNQEEEDMKYQQNGFGKGDSR | -46.04 | 51.6 | HHHHHHHHHHHHHHHHHHHHHHHCCCCCCCC |
| 260 | EEELRKLLEQINQEEEDMRYQQYGFGKGDSR | -46.04 | 48.4 | HHHHHHHHHHHHHHHHHHHHHHHCCCCCCCC |
| 056 | EEELRKLLEQLNQEEEDMKYQQNGFGKGDSR | -46.04 | 48.4 | HHHHHHHHHHHHHHHHHHHHHHHCCCCCCCC |
| 061 | EEEARKMLEQINNEEEDMRYQREGFGKGHDA | -46.03 | 45.2 | HHHHHHHHHHHHHHHHHHHHHHHHHCCCCCC |
| 079 | EEEARKFLEQYNQEEEDMRYQREGFGKGDSR | -45.98 | 54.8 | HHHHHHHHHHHHHHHHHHHHHHHHCCCCCCC |
| 372 | EEELKKFLEKLNNEEEDRRYQQYGFGKGHDA | -45.98 | 51.6 | HHHHHHHHHHHHHHHHHHHHHHHCCCCCCCC |
| 505 | EEEWRKFLEKYNNEAEDIRYQQQGFGKGDSR | -45.95 | 58.1 | HHHHHHHHHHHHHHHHHHHHHHHCCCCCCCC |
| 138 | EEEAKKFLEKFNNEEEDMRYQREGFGKGDSR | -45.94 | 64.5 | HHHHHHHHHHHCHHHHHHHHHHHHCCCCCCC |
| 747 | EEEARKFLEKYNNEAEDIRYQQNGFGKGDSR | -45.94 | 61.3 | HHHHHHHHHHHHHHHHHHHHHHHCCCCCCCC |
| 481 | EEELKKFLEKLNNEKEDMEYQKQGFGKGDSR | -45.94 | 58.1 | HHHHHHHHHHHHHHHHHHHHHHHCCCCCCCC |
| 574 | EEELKKFLEKYNNEAEDIRYQQNGFGKGDSR | -45.94 | 61.3 | HHHHHHHHHHHHHHHHHHHHHHHCCCCCCCC |
| 719 | EEELKKFLEKYNNEAEDIRYQQQGFGKGDSR | -45.94 | 61.3 | HHHHHHHHHHHHHHHHHHHHHHHCCCCCCCC |
| 631 | EEELKKFLEKYNNEAEDIYYQQQGFGKGDSR | -45.94 | 61.3 | HHHHHHHHHHHHHHHHHHHHHHHCCCCCCCC |
| 717 | EEELKKFLEKYNNEAEDMRYQQQGFGKGDSR | -45.94 | 61.3 | HHHHHHHHHHHHHHHHHHHHHHHCCCCCCCC |
| 212 | EEELKKILEKLNQEQEQIRYQQNGFGQGHDA | -45.88 | 41.9 | HHHHHHHHHHHHHHHHHHHHHHHHCCCCCCC |
| 018 | EEELKKFLEKYNNEYERMFFQQKGFGKGKDA | -45.88 | 48.4 | HHHHHHHHHHHHHHHHHHHHHHHCCCCCCCC |
| 474 | EEELRKLLDKLNQEQEQMSYQQQGFGQGDSR | -45.87 | 48.4 | HHHHHHHHHHHHHHHHHHHHHHHCCCCCCCC |
| 416 | EEEARKFLEQFNNEYEDMYYQQQGFGKGHSA | -45.82 | 51.6 | HHHHHHHHHHHHHHHHHHHHHHHHCCCCCCC |
| 028 | EEEARKFLEKFNNEEEDRSYQQQGFGKGHDA | -45.79 | 54.8 | HHHHHHHHHHHHHHHHHHHHHHHCCCCCCCC |
| 037 | EEEARKFLEKFNNEYEDRFYQQKGFGKGHDA | -45.79 | 58.1 | HHHHHHHHHHHHHHHHHHHHHHHCCCCCCCC |
| 257 | EEELKKFLEKYNNEAEDRRYQKEGFGKGHDA | -45.79 | 54.8 | HHHHHHHHHHHHHHHHHHHHHHHHCCCCCCC |
| 309 | EESLKKLLEQINNEEEDMRYQQYGFGKGHDA | -45.78 | 45.2 | HHHHHHHHHHHHHHHHHHHHHHHCCCCCCCC |

| | | | | |
|---|---|---|---|---|
| 554 | EEELKKLLEQINNEAEDMRYQQQGFGKGDSR | -45.74 | 54.8 | HHHHHHHHHHHHHHHHHHHHHHHCCCCCCCC |
| 269 | EEELKKLLEQINNEEEDIRYQQQGFGKGDSR | -45.74 | 51.6 | HHHHHHHHHHHHHHHHHHHHHHHCCCCCCCC |
| 589 | EEELKKLLEQINNEEEDMRYQQQGFGKGDSR | -45.74 | 51.6 | HHHHHHHHHHHHHHHHHHHHHHHCCCCCCCC |
| 861 | EEELKKLLEQINNEEEDMRYQREGFGKGDSR | -45.74 | 51.6 | HHHHHHHHHHHHHHHHHHHHHHHHCCCCCCC |
| 206 | EEELRKLLEQINNEKEDMEYQRAGFGKGDSR | -45.74 | 48.4 | HHHHHHHHHHHHHHHHHHHHHHHHCCCCCCC |
| 177 | EEELRKLLEQLNNEQEDMRYQQQGFGKGDSR | -45.74 | 48.4 | HHHHHHHHHHHHHHHHHHHHHHHCCCCCCCC |
| 644 | EEELKKLLEAINNREEDMRYQQYGFGKGHDA | -45.72 | 41.9 | HHHHHHHHHHHHHHHHHHHHHHHCCCCCCCC |
| 443 | EEELKKFLEQFNNEYEDMFYQQKGFGKGDSR | -45.70 | 61.3 | HHHHHHHHHHHHHHHHHHHHHHHCCCCCCCC |
| 221 | EEEARKFLEQFNNEYEDMFYQQKGFGKGDSR | -45.69 | 61.3 | HHHHHHHHHHHHHHHHHHHHHHHCCCCCCCC |
| 182 | EEEARKFLEQLNNEEEDMRYQQQGFGKGDSR | -45.69 | 54.8 | HHHHHHHHHHHHHHHHHHHHHHHCCCCCCCC |
| 997 | EEEARKFLEQYNNEAEDIRYQQGGFGKGDSR | -45.69 | 58.1 | HHHHHHHHHHHHHHHHHHHHHHCCCCCCCCC |
| 348 | EEEARKFLEQYNNEAEDMRYQQQGFGKGDSR | -45.69 | 58.1 | HHHHHHHHHHHHHHHHHHHHHHHCCCCCCCC |
| 660 | EEEARKFLEQYNNEEEDMRYQRQGFGKGDSR | -45.69 | 54.8 | HHHHHHHHHHHCHHHHHHHHHHHCCCCCCCC |
| 305 | EEELKKFLEQLNNEEEDIRYQQQGFGKGDSR | -45.69 | 54.8 | HHHHHHHHHHHHHHHHHHHHHHHCCCCCCCC |
| 450 | EEELKKFLEQYNNEAEDMRYQRYGFGKGDSR | -45.69 | 58.1 | HHHHHHHHHHHHHHHHHHHHHHHCCCCCCCC |
| 939 | EEEWKKFIEQYNNEEEDMYYQQQGFGKGDSR | -45.69 | 51.6 | HHHHHHHHHHHHHHHHHHHHHHHCCCCCCCC |
| 867 | EEELKKLLQQLNEEEEQMQYQRAGFGQGHDA | -45.68 | 38.7 | HHHHHHHHHHHHHHHHHHHHHHHHCCCCCCC |
| 246 | EEELKKFLEQFNNVYEDMFYQQKGFGKGDSR | -45.63 | 58.1 | HHHHHHHHHHHHHHHHHHHHHHCCCCCCCCC |
| 497 | EEELKKLLDQINNEEEQMRYQQQGFGQGHDA | -45.62 | 41.9 | HHHHHHHHHHHHHHHHHHHHHHHCCCCCCCC |
| 378 | EEELKKFLEKLNNEEEDRRYQQQGFGKGHDA | -45.59 | 51.6 | HHHHHHHHHHHHHHHHHHHHHHHHCCCCCCC |
| 238 | EEELRKLLEKINQEAEQIRYQQNGFGQGHSA | -45.57 | 41.9 | HHHHHHHHHHHHHHHHHHHHHHHHCCCCCCC |
| 696 | EEEARKMLEKINNEEEDIRYQQQGFGKGDSR | -45.55 | 54.8 | HHHHHHHHHHHHHHHHHHHHHHHCCCCCCCC |
| 189 | EEEARKFLQQYNNEYEDMFYQQKGFGKGHDA | -45.55 | 51.6 | HHHHHHHHHHHHHHHHHHHHHHHHCCCCCCC |
| 790 | EEELRKFLEQFNQEYEDMFYQQQGFGKGDSR | -45.49 | 58.1 | HHHHHHHHHHHHHHHHHHHHHHHCCCCCCCC |
| 888 | EEEARKFLEQYNQEAEDMRYQQQGFGKGDSR | -45.47 | 58.1 | HHHHHHHHHHHHHHHHHHHHHHHCCCCCCCC |
| 412 | EEEARKFLEQYNQEEEDMFYQQKGFGKGDSR | -45.47 | 58.1 | HHHHHHHHHHHHHHHHHHHHHHHCCCCCCCC |
| 266 | EEEARKFLEQYNQEEEDMRYQQQGFGKGDSR | -45.47 | 54.8 | HHHHHHHHHHHHHHHHHHHHHHHCCCCCCCC |
| 654 | EEELKKFLEQLNQEEEDMRYQQQGFGKGDSR | -45.47 | 54.8 | HHHHHHHHHHHHHHHHHHHHHHHCCCCCCCC |
| 232 | EEELKKFLEQYNQEAEDMRYQQNGFGKGDSR | -45.47 | 58.1 | HHHHHHHHHHHHHHHHHHHHHHHCCCCCCCC |
| 870 | EEELKKFLEQYNQEEEDMRYQREGFGKGDSR | -45.47 | 54.8 | HHHHHHHHHHHHHHHHHHHHHHHHCCCCCCC |
| 528 | EQEARKFLEQYNQEEEDMRYQREGFGKGDSR | -45.47 | 51.6 | HHHHHHHHHHHHHHHHHHHHHHHHCCCCCCC |
| 661 | EEELRKLLEQINQEEEDMRYQQQGFGKGDSR | -45.44 | 48.4 | HHHHHHHHHHHHHHHHHHHHHHHCCCCCCCC |
| 195 | EEELKKLLEKINQEAEQIRYQQNGFGQGHDA | -45.39 | 45.2 | HHHHHHHHHHHHHHHHHHHHHHHCCCCCCCC |
| 032 | EEELKKLLEKINQEEEQIRYQQQGFGQGHDA | -45.39 | 41.9 | HHHHHHHHHHHHHHHHHHHHHHHCCCCCCCC |
| 713 | EEELKKLLEKINQEEEQMRYQKEGFGQGHDA | -45.39 | 41.9 | HHHHHHHHHHHHHHHHHHHHHHHHCCCCCCC |
| 168 | EEELKKLLEKLNQEKEQIRYQQNGFGQGHDA | -45.39 | 41.9 | HHHHHHHHHHHHHHHHHHHHHHHCCCCCCCC |
| 541 | EEELKKLLEKLNQEKEQMSYQQQGFGQGHSA | -45.39 | 41.9 | HHHHHHHHHHHHHHHHHHHHHHHCCCCCCCC |
| 119 | EEELRKLLEKINQEAEQMRYQRYGFGQGHDA | -45.39 | 41.9 | HHHHHHHHHHHHHHHHHHHHHHHCCCCCCCC |

| | | | | |
|---|---|---|---|---|
| 160 | EEELRKLLEKINQEAEQMSYQQQGFGQGHSA | -45.39 | 41.9 | HHHHHHHHHHHHHHHHHHHHHHHCCCCCCCC |
| 864 | EEELRKLLEKLNQEKEQMRYQQQGFGQGHDA | -45.39 | 38.7 | HHHHHHHHHHHHHHHHHHHHHHHHCCCCCCC |
| 346 | EEELRKLLEKLNQEQEQMSYQQQGFGQGHDA | -45.39 | 38.7 | HHHHHHHHHHHHHHHHHHHHHHHCCCCCCCC |
| 060 | EEEARKFLEKYNNEAEDMRYQRQGFGKGDSR | -45.35 | 61.3 | HHHHHHHHHHHHHHHHHHHHHHHCCCCCCCC |
| 679 | EEELKKLLEKINNEAEQIRFQQNGFGQGKDA | -45.34 | 41.9 | HHHHHHHHHHHHHHHHHHHHHHHHCCCCCCC |
| 515 | EEELRKLLEKIEQEREQITYQQNGFGQGHDA | -45.32 | 35.5 | HHHHHHHHHHHHHHHHHHHHHHHCCCCCCCC |
| 746 | EEELRKLLEKLNNEQEQIRYQQNGFGQGHSA | -45.28 | 38.7 | HHHHHHHHHHHHHHHHHHHHHHHHCCCCCCC |
| 540 | EEEARKFLEQLNNEEEQMRYQQYGFGQGHDA | -45.26 | 41.9 | HHHHHHHHHHHHHHHHHHHHHHHCCCCCCCC |
| 518 | EEEARKFLEQYNQEAEDMRYQQQGFGKGDDR | -45.25 | 58.1 | HHHHHHHHHHHHHHHHHHHHHHHCCCCCCCC |
| 234 | EEQARKFLEQYNNESEDMRYQQQGFGKGDSR | -45.25 | 58.1 | HHHHHHHHHHHHHHHHHHHHHHHCCCCCCCC |
| 673 | EESLKKFLEQYNNESEDIRYQQNGFGKGHDA | -45.20 | 48.4 | HHHHHHHHHHHHHHHHHHHHHHHCCCCCCCC |
| 184 | EEEAKKFLEQLNNEEEDMRYQQQGFGKGDSR | -45.18 | 58.1 | HHHHHHHHHHHHHHHHHHHHHHHCCCCCCCC |
| 853 | EEEAKKFLEQLNNEEEDMRYQQYGFGKGDSR | -45.18 | 58.1 | HHHHHHHHHHHHHHHHHHHHHHHCCCCCCCC |
| 502 | EEEARKFLEQLNNEEEDMRYQQNGFGKGDSR | -45.18 | 54.8 | HHHHHHHHHHHHHHHHHHHHHHHCCCCCCCC |
| 272 | EEEARKFLEQLNNEEEDMRYQQYGFGKGDSR | -45.18 | 54.8 | HHHHHHHHHHHHHHHHHHHHHHHCCCCCCCC |
| 593 | EEEARKFLEQYNNEAEDIRYQQNGFGKGDSR | -45.18 | 58.1 | HHHHHHHHHHHHHHHHHHHHHHHCCCCCCCC |
| 154 | EEEARKFLEQYNNEAEDMRYQQNGFGKGDDR | -45.18 | 58.1 | HHHHHHHHHHHHHHHHHHHHHHHCCCCCCCC |
| 324 | EEEARKFLEQYNNEEEDMFYQQKGFGKGDSR | -45.18 | 58.1 | HHHHHHHHHHHHHHHHHHHHHHHCCCCCCCC |
| 801 | EEEARKFLEQYNNEEEDMRYQQEGFGKGDSR | -45.18 | 54.8 | HHHHHHHHHHHHHHHHHHHHHHHHCCCCCCC |
| 822 | EEELKKFLEQFNNEYEDMFYQQKGFGKGDDR | -45.18 | 61.3 | HHHHHHHHHHHHHHHHHHHHHHHCCCCCCCC |
| 074 | EEELKKFLEQLNNEEEDMRYQQYGFGKGDSR | -45.18 | 54.8 | HHHHHHHHHHHHHHHHHHHHHHHCCCCCCCC |
| 151 | EEELKKFLEQYNNEAEDIYYQQNGFGKGDSR | -45.18 | 58.1 | HHHHHHHHHHHHHHHHHHHHHHHCCCCCCCC |
| 472 | EEELKKFLEQYNNEAEDMRYQQGGFGKGDSR | -45.18 | 58.1 | HHHHHHHHHHHHHHHHHHHHHHCCCCCCCCC |
| 382 | EEELKKFLEQYNNEAEDMRYQQNGFGKGDSR | -45.18 | 58.1 | HHHHHHHHHHHHHHHHHHHHHHHCCCCCCCC |
| 786 | EEELKKFLEQYNNEAEDMRYQQQGFGKGDSR | -45.18 | 58.1 | HHHHHHHHHHHHHHHHHHHHHHHCCCCCCCC |
| 780 | EEELKKFLEQYNNEAEDMRYQRQGFGKGDSR | -45.18 | 58.1 | HHHHHHHHHHHHHHHHHHHHHHHCCCCCCCC |
| 519 | EEELKKFLEQYNNESEDIRYQQNGFGKGDSR | -45.18 | 54.8 | HHHHHHHHHHHHHHHHHHHHHHHCCCCCCCC |
| 838 | EEELKKFLEQYNNEYEDMFYQQKGFGKGDSR | -45.18 | 58.1 | HHHHHHHHHHHHHHHHHHHHHHCCCCCCCCC |
| 304 | EEELKKFLEQYNNESEDMRYQQQGFGKGDSR | -45.17 | 54.8 | HHHHHHHHHHHHHHHHHHHHHHHCCCCCCCC |
| 328 | EEEARKFLEQYNQEAEDMRYQQNGFGKGDSR | -45.15 | 58.1 | HHHHHHHHHHHHHHHHHHHHHHHCCCCCCCC |
| 422 | EEELRKLLEKINNEEEDRRYQQQGFGKGDSR | -45.11 | 51.6 | HHHHHHHHHHHHHHHHHHHHHHHCCCCCCCC |
| 179 | EEELKKLLEKINNEAEQIRYQQNGFGQGHDA | -45.09 | 45.2 | HHHHHHHHHHHHHHHHHHHHHHHHCCCCCCC |
| 215 | EEELKKLLEKINNEAEQITYQQNGFGQGHDA | -45.09 | 45.2 | HHHHHHHHHHHHHHHHHHHHHHHCCCCCCCC |
| 522 | EEELKKLLEKINNEAEQITYQQNGFGQGHSA | -45.09 | 45.2 | HHHHHHHHHHHHHHHHHHHHHHHCCCCCCCC |
| 635 | EEELKKLLEKINNEEEQMRYQQQGFGQGHDA | -45.09 | 41.9 | HHHHHHHHHHHCHHHHHHHHHHHCCCCCCCC |
| 172 | EEELKKLLEKINNEQEQMSYQQQGFGQGHDA | -45.09 | 41.9 | HHHHHHHHHHHHHHHHHHHHHHHCCCCCCCC |
| 353 | EEELKKLLEKLNNEEEQMKYQQNGFGQGHDA | -45.09 | 41.9 | HHHHHHHHHHHCHHHHHHHHHHHHCCCCCCC |
| 388 | EEELKKLLEKLNNEEEQMKYQQQGFGQGHDA | -45.09 | 41.9 | HHHHHHHHHHHCHHHHHHHHHHHCCCCCCCC |

| | | | | |
|---|---|---|---|---|
| 928 | EEELKKLLEKLNNEEEQMKYQRYGFGQGHDA | -45.09 | 41.9 | HHHHHHHHHHHCHHHHHHHHHHHCCCCCCCC |
| 901 | EEELKKLLEKLNNEKEQITYQQNGFGQGHDA | -45.09 | 41.9 | HHHHHHHHHHHHHHHHHHHHHHHCCCCCCCC |
| 549 | EEELKKLLEKLNNEQEQIRYQQGGFGQGHDF | -45.09 | 41.9 | HHHHHHHHHHHHHHHHHHHHHHCCCCCCCCC |
| 162 | EEELKKLLEKLNNEQEQIRYQQNGFGQGHDA | -45.09 | 41.9 | HHHHHHHHHHHHHHHHHHHHHHHHCCCCCCC |
| 941 | EEELKKLLEKLNNEQEQMRYQQNGFGQGHDA | -45.09 | 41.9 | HHHHHHHHHHHHHHHHHHHHHHHHCCCCCCC |
| 509 | EEELKKLLEKLNNEQEQMRYQRQGFGQGHDA | -45.09 | 41.9 | HHHHHHHHHHHHHHHHHHHHHHHHCCCCCCC |
| 331 | EEELRKLLEKINNEAEQIRYQQGGFGQGHDF | -45.09 | 41.9 | HHHHHHHHHHHHHHHHHHHHHHCCCCCCCCC |
| 478 | EEELRKLLEKINNEAEQIRYQQNGFGQGHDA | -45.09 | 41.9 | HHHHHHHHHHHHHHHHHHHHHHHHCCCCCCC |
| 306 | EEELRKLLEKINNEEEQIRYQQQGFGQGHDA | -45.09 | 38.7 | HHHHHHHHHHHCHHHHHHHHHHHCCCCCCCC |
| 700 | EEELRKLLEKINNEKEQMEYQRAGFGQGHDA | -45.09 | 38.7 | HHHHHHHHHHHHHHHHHHHHHHHHCCCCCCC |
| 191 | EEELRKLLEKLNNEEEQIKYQQNGFGQGHDA | -45.09 | 38.7 | HHHHHHHHHHHCHHHHHHHHHHHHCCCCCCC |
| 242 | EEELRKLLEKLNNEEEQMKYQQNGFGQGHDA | -45.09 | 38.7 | HHHHHHHHHHHCHHHHHHHHHHHHCCCCCCC |
| 227 | EEELRKLLEKLNNEEEQMKYQQQGFGQGHDA | -45.09 | 38.7 | HHHHHHHHHHHCHHHHHHHHHHHCCCCCCCC |
| 856 | EEELRKLLEKLNNEKEQMSYQQQGFGQGHDA | -45.09 | 38.7 | HHHHHHHHHHHHHHHHHHHHHHHCCCCCCCC |
| 173 | EEELRKLLEKLNNEQEQMRYQQGGFGQGHDF | -45.09 | 38.7 | HHHHHHHHHHHHHHHHHHHHHHCCCCCCCCC |
| 862 | EEELRKLLEKLNNEQEQMRYQQQGFGQGHDA | -45.09 | 38.7 | HHHHHHHHHHHHHHHHHHHHHHHCCCCCCCC |
| 447 | EEELRKLLEKLNNEQEQMRYQRQGFGQGHDA | -45.09 | 38.7 | HHHHHHHHHHHHHHHHHHHHHHHHCCCCCCC |
| 007 | EEELKKFLEKYNQEAEQMRFQQQGFGQGKDA | -45.07 | 45.2 | HHHHHHHHHHHHHHHHHHHHHHHCCCCCCCC |
| 931 | EEEARKFLEKYNNESEDIRYQQNGFGKGHDA | -45.04 | 51.6 | HHHHHHHHHHHCCHHHHHHHHHHCCCCCCCC |
| 051 | EESLKKLLEQINNEEEDMRYQQQGFGKGDSR | -45.03 | 51.6 | HHHHHHHHHHHHHHHHHHHHHHHCCCCCCCC |
| 407 | EESARKFLDKFNNEYEQMFYQKEGFGQGHDA | -45.02 | 54.8 | HHHHHHHHHHHHHHHHHHHHHHHHCCCCCCC |
| 595 | EEELRKLLEKINQEAEEMRKQQQGFGSGKDA | -44.96 | 38.7 | HHHHHHHHHHHHHHHHHHHHHHHHCCCCCCC |
| 883 | EEELKKLLEKLNQEQEQMRYQRQGFGQGHDA | -44.91 | 41.9 | HHHHHHHHHHHHHHHHHHHHHHHHCCCCCCC |
| 629 | EEEARKFLEQYNNEEEDMYYQQKGFGKGDSR | -44.86 | 54.8 | HHHHHHHHHHHHHHHHHHHHHHHCCCCCCCC |
| 990 | EEEARKFLEKFNQEAEQITYQQNGFGQGHDA | -44.82 | 51.6 | HHHHHHHHHHHHHHHHHHHHHHHCCCCCCCC |
| 658 | EEEARKFLEKYNQESEQIRYQQNGFGQGHDA | -44.82 | 45.2 | HHHHHHHHHHHHHHHHHHHHHHHCCCCCCCC |
| 599 | EEEARKFLEKYNQESEQIRYQQQGFGQGHDA | -44.82 | 45.2 | HHHHHHHHHHHHHHHHHHHHHHHHCCCCCCC |
| 369 | EEELKKFLEKFNQEYEQMFYQQKGFGQGHDA | -44.82 | 51.6 | HHHHHHHHHHHHHHHHHHHHHHHHCCCCCCC |
| 802 | EEELKKFLEKLNQEEEQIRYQQQGFGQGHSA | -44.82 | 45.2 | HHHHHHHHHHHHHHHHHHHHHHHHCCCCCCC |
| 503 | EEELKKFLEKLNQEEEQMRYQQQGFGQGHDA | -44.82 | 45.2 | HHHHHHHHHHHHHHHHHHHHHHHCCCCCCCC |
| 851 | EEELKKFLEKYNQEAEQIYYQQNGFGQGHDA | -44.82 | 48.4 | HHHHHHHHHHHHHHHHHHHHHHHCCCCCCCC |
| 205 | EEELKKFLEKYNQEAEQLRYQQKGFGQGHDA | -44.82 | 51.6 | HHHHHHHHHHHHHHHHHHHHHHHHCCCCCCC |
| 456 | EEELKKFLEKYNQESEQIRYQQNGFGQGHDA | -44.82 | 45.2 | HHHHHHHHHHHHHHHHHHHHHHHCCCCCCCC |
| 217 | EEEARKMLEQINNEEEDMRYQQYGFGKGDSR | -44.80 | 51.6 | HHHHHHHHHHHHHHHHHHHHHHHCCCCCCCC |
| 110 | EEELKKLLEKINQEEEQMRYQQYGFGQGHDA | -44.79 | 41.9 | HHHHHHHHHHHHHHHHHHHHHHHCCCCCCCC |
| 985 | EEEARKMLEQINNEAEDMRYQQNGFGKGDSR | -44.78 | 54.8 | HHHHHHHHHHHHHHHHHHHHHHHCCCCCCCC |
| 873 | EEEARKFLEKFNNEYEQMFYQQKGFGQGHSA | -44.71 | 51.6 | HHHHHHHHHHHHHHHHHHHHHHHHCCCCCCC |
| 500 | EEELRKILEQINQEEEQMRYQQYGFGQGDSR | -44.67 | 41.9 | HHHHHHHHHHHHHHHHHHHHHHHCCCCCCCC |

| | | | | |
|---|---|---|---|---|
| 672 | EQEAREFLKQFNEEYEDMFYQQKGFGKGHDA | -44.65 | 51.6 | HHHHHHHHHHHHHHHHHHHHHHHHCCCCCCC |
| 596 | EEELRKILEQINNEEEQMRYQQQGFGQGHSA | -44.63 | 35.5 | HHHHHHHHHHHCHHHHHHHHHHHCCCCCCCC |
| 477 | EEELKKLLEQINQEAEQMRYQKEGFGQGHDA | -44.62 | 41.9 | HHHHHHHHHHHHHHHHHHHHHHHHCCCCCCC |
| 542 | EEELKKLLEQINQEAEQMRYQQNGFGQGHDA | -44.62 | 41.9 | HHHHHHHHHHHHHHHHHHHHHHHCCCCCCCC |
| 445 | EEELKKLLEQINQEAEQMRYQQQGFGQGHDA | -44.62 | 41.9 | HHHHHHHHHHHHHHHHHHHHHHHCCCCCCCC |
| 760 | EEELKKLLEQLNQEKEQMSYQRYGFGQGHDA | -44.62 | 38.7 | HHHHHHHHHHHHHHHHHHHHHHHCCCCCCCC |
| 249 | EEELRKLLEQINQEAEQMRYQQNGFGQGHDA | -44.62 | 38.7 | HHHHHHHHHHHHHHHHHHHHHHHCCCCCCCC |
| 434 | EEELRKLLEQINQEEEQMRYQREGFGQGHDA | -44.62 | 35.5 | HHHHHHHHHHHHHHHHHHHHHHHHHCCCCCC |
| 677 | EEELRKLLEQLNQEEEQMKYQQQGFGQGHDA | -44.62 | 35.5 | HHHHHHHHHHHHHHHHHHHHHHHCCCCCCCC |
| 293 | EEELKKLLEQINQEEEQMRYQREGFGQGHDA | -44.60 | 38.7 | HHHHHHHHHHHHHHHHHHHHHHHHHCCCCCC |
| 639 | EEEARKFLEQFNNEEEDMRYQQQGFGKGDSR | -44.58 | 58.1 | HHHHHHHHHHHHHHHHHHHHHHHCCCCCCCC |
| 752 | EEELRKFLEKYNNEAEDRRYQQQGFGKGDSR | -44.55 | 58.1 | HHHHHHHHHHHHHHHHHHHHHHHCCCCCCCC |
| 583 | EEELKKFLEQLNNDKEQMEYQRAGFGQGHDA | -44.54 | 38.7 | HHHHHHHHHHHHHHHHHHHHHHHHCCCCCCC |
| 868 | EEEAKKFLEKYNNEYEQMEYQRTGRGQGHDF | -44.53 | 48.4 | HHHHHHHHHHHHHHHHHHHHHHHCCCCCCCC |
| 840 | EEEARKFLEKFNNEAEQMQYQRAGFGQGHDA | -44.53 | 51.6 | HHHHHHHHHHHHHHHHHHHHHHHHCCCCCCC |
| 470 | EEEARKFLEKFNNEEEQMRYQQEGFGQGHSA | -44.53 | 48.4 | HHHHHHHHHHHCHHHHHHHHHHHHCCCCCCC |
| 118 | EEEARKFLEKFNNEYEQMFYQQKGFGQGHDA | -44.53 | 51.6 | HHHHHHHHHHHHHHHHHHHHHHHHCCCCCCC |
| 607 | EEEARKFLEKLNNEEEQMRYQQQGFGQGHDA | -44.53 | 45.2 | HHHHHHHHHHHHHHHHHHHHHHHCCCCCCCC |
| 165 | EEEARKFLEKLNNEEEQMRYQQQGFGQGHSA | -44.53 | 45.2 | HHHHHHHHHHHHHHHHHHHHHHHCCCCCCCC |
| 039 | EEEARKFLEKLNNEEEQMRYQQYGFGQGHDA | -44.53 | 45.2 | HHHHHHHHHHHHHHHHHHHHHHHCCCCCCCC |
| 414 | EEEARKFLEKYNNEAEQIRYQQNGFGQGHDA | -44.53 | 48.4 | HHHHHHHHHHHHHHHHHHHHHHHHCCCCCCC |
| 087 | EEEARKFLEKYNNEAEQIYYQQNGFGQGHDA | -44.53 | 48.4 | HHHHHHHHHHHHHHHHHHHHHHHCCCCCCCC |
| 548 | EEEARKFLEKYNNEEEQMFYQQKGFGQGHDA | -44.53 | 48.4 | HHHHHHHHHHHCHHHHHHHHHHHHCCCCCCC |
| 903 | EEEARKFLEKYNNESEQIRYQQNGFGQGHDA | -44.53 | 45.2 | HHHHHHHHHHHCHHHHHHHHHHHCCCCCCCC |
| 633 | EEEARKFLEKYNNEYEQIFYQQKGFGQGHDA | -44.53 | 48.4 | HHHHHHHHHHHHHHHHHHHHHHHHCCCCCCC |
| 678 | EEELKKFLEKFNNEEEQMSYQQQGFGQGHDA | -44.53 | 48.4 | HHHHHHHHHHHCHHHHHHHHHHHCCCCCCCC |
| 829 | EEELKKFLEKFNNEYEQIFYQQKGFGQGHDA | -44.53 | 51.6 | HHHHHHHHHHHHHHHHHHHHHHHHCCCCCCC |
| 956 | EEELKKFLEKFNNEYEQMFYQQKGFGQGHDA | -44.53 | 51.6 | HHHHHHHHHHHHHHHHHHHHHHHHCCCCCCC |
| 493 | EEELKKFLEKFNNEYEQMFYQQKGFGQGHSA | -44.53 | 51.6 | HHHHHHHHHHHHHHHHHHHHHHHHCCCCCCC |
| 768 | EEELKKFLEKLNNEEEQMRYQQYGFGQGHDA | -44.53 | 45.2 | HHHHHHHHHHHCHHHHHHHHHHHCCCCCCCC |
| 737 | EEELKKFLEKYNNEAEQIRYQQNGFGQGHDA | -44.53 | 48.4 | HHHHHHHHHHHHHHHHHHHHHHHHCCCCCCC |
| 671 | EEELKKFLEKYNNEAEQIRYQQQGFGQGHDA | -44.53 | 48.4 | HHHHHHHHHHHHHHHHHHHHHHHHCCCCCCC |
| 395 | EEELKKFLEKYNNEAEQIYYQQNGFGQGHDA | -44.53 | 48.4 | HHHHHHHHHHHHHHHHHHHHHHHCCCCCCCC |
| 392 | EEELKKFLEKYNNEAEQIYYQQNGFGQGHSA | -44.53 | 48.4 | HHHHHHHHHHHHHHHHHHHHHHHCCCCCCCC |
| 077 | EEELKKFLEKYNNEAEQMRYQQQGFGQGHDA | -44.53 | 48.4 | HHHHHHHHHHHHHHHHHHHHHHHHCCCCCCC |
| 198 | EEELKKFLEKYNNEEEQMRYQREGFGQGHDA | -44.53 | 45.2 | HHHHHHHHHHHCCHHHHHHHHHHHHCCCCCC |
| 992 | EEELKKFLEKYNNEEEQMSYQQQGFGQGHDA | -44.53 | 45.2 | HHHHHHHHHHHCHHHHHHHHHHHCCCCCCCC |
| 202 | EEELKKFLEKYNNESEQIRYQQNGFGQGHDA | -44.53 | 45.2 | HHHHHHHHHHHHHHHHHHHHHHHCCCCCCCC |

| | | | | |
|---|---|---|---|---|
| 946 | EEELKKFLEKYNNESEQIRYQQQGFGQGHDA | -44.53 | 45.2 | HHHHHHHHHHHHHHHHHHHHHHHHCCCCCCC |
| 571 | EEELKKFLEKYNNESEQMRYQQQGFGQGHDA | -44.53 | 45.2 | HHHHHHHHHHHHHHHHHHHHHHHHCCCCCCC |
| 243 | EEELKKFLEKYNNEYEQMFYQQKGFGQGHDA | -44.53 | 48.4 | HHHHHHHHHHHHHHHHHHHHHHHHCCCCCCC |
| 435 | EEELRKFLEKYNNEYEQIFYQQKGFGQGHDA | -44.53 | 45.2 | HHHHHHHHHHHHHHHHHHHHHHHHCCCCCCC |
| 480 | EQEARKFLEKYNNEAEQIYYQQNGFGQGHDA | -44.53 | 45.2 | HHHHHHHHHHHHHHHHHHHHHHHCCCCCCCC |
| 183 | EEELKKFLEQYNQESEEMRKQQNGFGSGKDA | -44.52 | 38.7 | HHHHHHHHHHHHHHHHHHHHHHHHCCCCCCC |
| 034 | EEELKKILEQLNQEEEQMKYQRYGFGQGHDA | -44.48 | 38.7 | HHHHHHHHHHHHHHHHHHHHHHHCCCCCCCC |
| 730 | EEELKKLLEKLNQEQEQMSKQQQGFGSGKDA | -44.46 | 38.7 | HHHHHHHHHHHHHHHHHHHHHHHHCCCCCCC |
| 914 | EEEARKFLEKYNQEAEEIYKQQNGFGSGKDA | -44.40 | 45.2 | HHHHHHHHHHHHHHHHHHHHHHHCCCCCCCC |
| 573 | EEELKKILEQLNNEKEQMSYQQQGFGQGDSR | -44.38 | 45.2 | HHHHHHHHHHHHHHHHHHHHHHHCCCCCCCC |
| 375 | EEELKKLLEKINNEAEQIRYQQNGFGQGDSR | -44.36 | 51.6 | HHHHHHHHHHHHHHHHHHHHHHHCCCCCCCC |
| 964 | EEELKKLLEKLNNEEEQMKYQQQGFGQGDSR | -44.36 | 48.4 | HHHHHHHHHHHCHHHHHHHHHHHCCCCCCCC |
| 136 | EEELRKLLEKINNEQEQISYQQNGFGQGDSR | -44.36 | 45.2 | HHHHHHHHHHHHHHHHHHHHHHHCCCCCCCC |
| 418 | EEELRKLLEKLNNEQEQMRYQQNGFGQGDSR | -44.36 | 45.2 | HHHHHHHHHHHHHHHHHHHHHHHCCCCCCCC |
| 645 | EEEARKFLEKFNQEYEQIFYQQKGFGQGHDA | -44.35 | 51.6 | HHHHHHHHHHHHHHHHHHHHHHHHCCCCCCC |
| 734 | EEEARKFLEKYNQEEEQMRYQRQGFGQGHDA | -44.35 | 45.2 | HHHHHHHHHHHHHHHHHHHHHHHHCCCCCCC |
| 127 | EEELKKLLEQINNEEEQMRYQQQGFGQGHDA | -44.34 | 38.7 | HHHHHHHHHHHCHHHHHHHHHHHCCCCCCCC |
| 878 | EEELRKLLEQINNEEEQMRYQQYGFGQGHDA | -44.34 | 35.5 | HHHHHHHHHHHCHHHHHHHHHHHCCCCCCCC |
| 564 | EEELKKLLEQINNEAEQIRYQQNGFGQGHDA | -44.32 | 41.9 | HHHHHHHHHHHHHHHHHHHHHHHHCCCCCCC |
| 665 | EEELKKLLEQINNEAEQIRYQQNGFGQGHSA | -44.32 | 41.9 | HHHHHHHHHHHHHHHHHHHHHHHHCCCCCCC |
| 190 | EEELKKLLEQINNEAEQMRYQQNGFGQGHDA | -44.32 | 41.9 | HHHHHHHHHHHHHHHHHHHHHHHHCCCCCCC |
| 550 | EEELKKLLEQINNEEEQIRYQQQGFGQGHDA | -44.32 | 38.7 | HHHHHHHHHHHCHHHHHHHHHHHCCCCCCCC |
| 335 | EEELKKLLEQINNEEEQMRYQQYGFGQGHDA | -44.32 | 38.7 | HHHHHHHHHHHCHHHHHHHHHHHCCCCCCCC |
| 214 | EEELKKLLEQINNEKEQMEYQRAGFGQGHDA | -44.32 | 38.7 | HHHHHHHHHHHHHHHHHHHHHHHHCCCCCCC |
| 837 | EEELKKLLEQLNNEEEQMKYQQGGFGQGHDF | -44.32 | 38.7 | HHHHHHHHHHHCHHHHHHHHHHCCCCCCCCC |
| 024 | EEELKKLLEQLNNEEEQMKYQQNGFGQGHDA | -44.32 | 38.7 | HHHHHHHHHHHCHHHHHHHHHHHHCCCCCCC |
| 875 | EEELKKLLEQLNNEKEQMSYQQQGFGQGHDA | -44.32 | 38.7 | HHHHHHHHHHHHHHHHHHHHHHHCCCCCCCC |
| 557 | EEELKKLLEQLNNEKEQMSYQQQGFGQGHSA | -44.32 | 38.7 | HHHHHHHHHHHHHHHHHHHHHHHCCCCCCCC |
| 457 | EEELKKLLEQLNNEQEQMRYQQNGFGQGHDA | -44.32 | 38.7 | HHHHHHHHHHHHHHHHHHHHHHHHCCCCCCC |
| 597 | EEELKKLLEQLNNEQEQMRYQQQGFGQGHDA | -44.32 | 38.7 | HHHHHHHHHHHHHHHHHHHHHHHCCCCCCCC |
| 880 | EEELKKLLEQLNNEQEQMRYQRQGFGQGHDA | -44.32 | 38.7 | HHHHHHHHHHHHHHHHHHHHHHHHCCCCCCC |
| 949 | EEELKKLLEQLNNEQEQMRYQRYGFGQGHDA | -44.32 | 38.7 | HHHHHHHHHHHHHHHHHHHHHHHCCCCCCCC |
| 735 | EEELRKLLEQINNEAEQMRYQQNGFGQGHDA | -44.32 | 38.7 | HHHHHHHHHHHHHHHHHHHHHHHHCCCCCCC |
| 690 | EEELRKLLEQINNEAEQMSYQQQGFGQGHDA | -44.32 | 38.7 | HHHHHHHHHHHHHHHHHHHHHHHCCCCCCCC |
| 710 | EEELRKLLEQINNEEEQMRYQQQGFGQGHDA | -44.32 | 35.5 | HHHHHHHHHHHCHHHHHHHHHHHCCCCCCCC |
| 791 | EEELRKLLEQINNEEEQMRYQREGFGQGHDA | -44.32 | 35.5 | HHHHHHHHHHHCHHHHHHHHHHHHHCCCCCC |
| 410 | EEELRKLLEQLNNEEEQMKYQQNGFGQGHDA | -44.32 | 35.5 | HHHHHHHHHHHCHHHHHHHHHHHHCCCCCCC |
| 517 | EEELRKLLEQLNNEKEQMRYQRQGFGQGHDA | -44.32 | 35.5 | HHHHHHHHHHHHHHHHHHHHHHHHCCCCCCC |

| | | | | |
|---|---|---|---|---|
| 947 | EEELRKLLEQLNNEKEQMSYQQQGFGQGHDA | -44.32 | 35.5 | HHHHHHHHHHHHHHHHHHHHHHHCCCCCCCC |
| 927 | EEELRKLLEQLNNEQEQMRYQREGFGQGHDA | -44.32 | 35.5 | HHHHHHHHHHHHHHHHHHHHHHHHHCCCCCC |
| 996 | EQELRKLLEQINNEEEQMRYQQQGFGQGHDA | -44.32 | 32.3 | HHHHHHHHHHHHHHHHHHHHHHHCCCCCCCC |
| 330 | EEEWKKLIEQINNDEEQMRYQQQGFGQGHDA | -44.32 | 32.3 | HHHHHHHHHHHCCHHHHHHHHHHCCCCCCCC |
| 167 | EEEWKKFLEKYNNEAEQIYYQQNGFGQGHSA | -44.21 | 48.4 | HHHHHHHHHHHHHHHHHHHHHHHCCCCCCCC |
| 471 | EEEARKFLEKFNNEEEQMSYQQQGFGQGHDA | -44.20 | 48.4 | HHHHHHHHHHHHHHHHHHHHHHHCCCCCCCC |
| 277 | EEELKKILEQLNQEQEQMSYQKEGFGQGDSR | -44.16 | 45.2 | HHHHHHHHHHHHHHHHHHHHHHHCCCCCCCC |
| 793 | EEELKKLLEKLNQEEEQMKYQQNGFGQGDSR | -44.15 | 48.4 | HHHHHHHHHHHHHHHHHHHHHHHCCCCCCCC |
| 124 | EEELRKLLEQINQEEEQMRYQQQGFGQGHDA | -44.15 | 35.5 | HHHHHHHHHHHHHHHHHHHHHHHCCCCCCCC |
| 492 | EEELKKFLEKFNNEYEEMFKQQKGFGSGKDA | -44.10 | 48.4 | HHHHHHHHHHHHHHHHHHHHHHHCCCCCCCC |
| 383 | EEELKKFLEKYNNEAEEIRKQQGGFGSGKDA | -44.10 | 45.2 | HHHHHHHHHHHHHHHHHHHHHHCCCCCCCCC |
| 813 | EEELKKFLEKYNNEAEEIRKQQNGFGSGKDA | -44.10 | 45.2 | HHHHHHHHHHHHHHHHHHHHHHHCCCCCCCC |
| 181 | EEELKKFLEKFNQEYEQMFYQQKGFGQGDSR | -44.09 | 58.1 | HHHHHHHHHHHHHHHHHHHHHHHCCCCCCCC |
| 643 | EEEARKFLEQYNQEAEQMRYQQQGFGQGHSA | -44.07 | 45.2 | HHHHHHHHHHHHHHHHHHHHHHHCCCCCCCC |
| 930 | EEEARKFLEQFNQEYEQMFYQQKGFGQGHDA | -44.06 | 48.4 | HHHHHHHHHHHHHHHHHHHHHHHHCCCCCCC |
| 106 | EEEARKFLEQFNQEYEQMYYQQQGFGQGHDA | -44.06 | 45.2 | HHHHHHHHHHHHHHHHHHHHHHHCCCCCCCC |
| 408 | EEEARKFLEQLNQEEEQMRYQQQGFGQGHDA | -44.06 | 41.9 | HHHHHHHHHHHHHHHHHHHHHHHCCCCCCCC |
| 290 | EEEARKFLEQYNQESEQMRYQQQGFGQGHDA | -44.06 | 41.9 | HHHHHHHHHHHHHHHHHHHHHHHHCCCCCCC |
| 439 | EEELKKFLEQFNQEYEQMFYQQKGFGQGHDA | -44.06 | 48.4 | HHHHHHHHHHHHHHHHHHHHHHHHCCCCCCC |
| 913 | EEELKKFLEQYNQEAEQIYYQQNGFGQGHDA | -44.06 | 45.2 | HHHHHHHHHHHHHHHHHHHHHHHCCCCCCCC |
| 967 | EEELKKFLEQYNQEAEQMRYQQNGFGQGHSA | -44.06 | 45.2 | HHHHHHHHHHHHHHHHHHHHHHHHCCCCCCC |
| 137 | EEELKKFLEQYNQEEEQMRYQREGFGQGHDA | -44.06 | 41.9 | HHHHHHHHHHHHHHHHHHHHHHHHHCCCCCC |
| 847 | EEELKKFLEQYNQESEQMRYQQQGFGQGHDA | -44.06 | 41.9 | HHHHHHHHHHHHHHHHHHHHHHHHCCCCCCC |
| 527 | EEELKKFLEKFNNEEEQMRYQRQGFGQGHDA | -44.05 | 48.4 | HHHHHHHHHHHCHHHHHHHHHHHHCCCCCCC |
| 142 | EEELKKFLEKYNNEAEQMRYQQQGFGQGHSA | -44.05 | 48.4 | HHHHHHHHHHHHHHHHHHHHHHHHCCCCCCC |
| 728 | EEEWKKFIEKLNNEEEQMRYQQQGFGQGHDA | -44.05 | 41.9 | HHHHHHHHHHHCHHHHHHHHHHHCCCCCCCC |
| 804 | EEELKKLLEKLNNEQEQIRYQQGGFGQGDSR | -44.04 | 48.4 | HHHHHHHHHHHHHHHHHHHHHHCCCCCCCCC |
| 693 | EEELRKLLEKINNEEEQIRYQQQGFGQGDSR | -44.04 | 45.2 | HHHHHHHHHHHCHHHHHHHHHHHCCCCCCCC |
| 729 | EEEARKFLEKYNNESEQMRYQQQGFGQGDSR | -43.98 | 51.6 | HHHHHHHHHHHHHHHHHHHHHHHCCCCCCCC |
| 657 | EEEARKFLEKYNNEAEQMRYQQQGFGQGHDA | -43.93 | 48.4 | HHHHHHHHHHHHHHHHHHHHHHHHCCCCCCC |
| 627 | EEEAKKFLEKYNQEEEQMSYQQQGFGQGHDA | -43.93 | 48.4 | HHHHHHHHHHHHHHHHHHHHHHHCCCCCCCC |
| 180 | EEELKKLLEQINNEKEEMEKQKQGFGSGKDA | -43.90 | 35.5 | HHHHHHHHHHHHHHHHHHHHHHHHCCCCCCC |
| 872 | EEELKKLLEQLNNEEEEMKKQQNGFGSGKDA | -43.90 | 35.5 | HHHHHHHHHHHHHHHHHHHHHHHHCCCCCCC |
| 725 | EEQLKKLLEQINNEEEQIRYQQQGFGQGHDA | -43.89 | 41.9 | HHHHHHHHHHHCHHHHHHHHHHHCCCCCCCC |
| 224 | EESLKKLLEKINNEAEQIRYQQNGFGQGHDA | -43.87 | 45.2 | HHHHHHHHHHHHHHHHHHHHHHHHCCCCCCC |
| 585 | EEELKKLLEKLNNEKEQMSYQQQGFGQGDSR | -43.85 | 48.4 | HHHHHHHHHHHHHHHHHHHHHHHCCCCCCCC |
| 325 | EEELRKLLEKINNEEEQMRYQQQGFGQGDSR | -43.85 | 45.2 | HHHHHHHHHHHCHHHHHHHHHHHCCCCCCCC |
| 436 | EEELRKLLEKINNEQEQMSYQQQGFGQGDSR | -43.85 | 45.2 | HHHHHHHHHHHHHHHHHHHHHHHCCCCCCCC |

| | | | | |
|---|---|---|---|---|
| 908 | EEELRKLLEKLNNEEEQMKYQKEGFGQGDSR | -43.85 | 45.2 | HHHHHHHHHHHCHHHHHHHHHHHCCCCCCCC |
| 386 | EEELRKLLEKLNNEQEQMRYQRYGFGQGDSR | -43.85 | 45.2 | HHHHHHHHHHHHHHHHHHHHHHHCCCCCCCC |
| 662 | EEELRKLLEQLNNEQEQMRYQQNGFGQGHDA | -43.85 | 35.5 | HHHHHHHHHHHHHHHHHHHHHHHHCCCCCCC |
| 026 | EEELRKLLEQLNNEQEQMRYQRQGFGQGHDA | -43.85 | 35.5 | HHHHHHHHHHHHHHHHHHHHHHHHCCCCCCC |
| 314 | EEELKKFLEKLNNEQEQMSYQQQGFGQGDSR | -43.80 | 51.6 | HHHHHHHHHHHHHHHHHHHHHHHCCCCCCCC |
| 256 | EEELKKFLEKYNNEEEQMRYQRQGFGQGDDR | -43.80 | 51.6 | HHHHHHHHHHHCCHHHHHHHHHHCCCCCCCC |
| 499 | EEELKKFLEKYNNEEEQMRYQREGFGQGHSA | -43.79 | 45.2 | HHHHHHHHHHHCCHHHHHHHHHHHHCCCCCC |
| 646 | EEEARKFLENYNNRYEDMEYQRTGRGKGHDF | -43.78 | 45.2 | HHHHHHHHHHHHHHHHHHHHHHCCCCCCCCC |
| 957 | EEEAKKFLEQFNNEEEQMSYQQQGFGQGHDA | -43.76 | 48.4 | HHHHHHHHHHHHHHHHHHHHHHHCCCCCCCC |
| 863 | EEEAKKFLEQYNNEYEQMFYQQKGFGQGHDA | -43.76 | 48.4 | HHHHHHHHHHHHHHHHHHHHHHHHCCCCCCC |
| 376 | EEEARKFLEQFNNEEEQMSYQQNGFGQGHDA | -43.76 | 45.2 | HHHHHHHHHHHHHHHHHHHHHHHHCCCCCCC |
| 280 | EEEARKFLEQFNNEEEQMSYQQQGFGQGHDA | -43.76 | 45.2 | HHHHHHHHHHHHHHHHHHHHHHHCCCCCCCC |
| 204 | EEEARKFLEQFNNEYEQMFYQQKGFGQGHDA | -43.76 | 48.4 | HHHHHHHHHHHHHHHHHHHHHHHHCCCCCCC |
| 307 | EEEARKFLEQLNNEEEQMRYQQQGFGQGHDA | -43.76 | 41.9 | HHHHHHHHHHHHHHHHHHHHHHHCCCCCCCC |
| 411 | EEEARKFLEQLNNEKEQMEYQRAGFGQGHDA | -43.76 | 41.9 | HHHHHHHHHHHHHHHHHHHHHHHHCCCCCCC |
| 857 | EEEARKFLEQYNNEAEQIYYQQNGFGQGHDA | -43.76 | 45.2 | HHHHHHHHHHHHHHHHHHHHHHHCCCCCCCC |
| 584 | EEEARKFLEQYNNEAEQMRYQQQGFGQGHDA | -43.76 | 45.2 | HHHHHHHHHHHHHHHHHHHHHHHHCCCCCCC |
| 271 | EEEARKFLEQYNNEAEQMYYQQQGFGQGHDA | -43.76 | 45.2 | HHHHHHHHHHHHHHHHHHHHHHHCCCCCCCC |
| 745 | EEEARKFLEQYNNEEEQMFYQQKGFGQGHDA | -43.76 | 45.2 | HHHHHHHHHHHCHHHHHHHHHHHHCCCCCCC |
| 860 | EEEARKFLEQYNNEEEQMRYQQQGFGQGHDA | -43.76 | 41.9 | HHHHHHHHHHHCHHHHHHHHHHHCCCCCCCC |
| 013 | EEEARKFLEQYNNEEEQMRYQREGFGQGHDA | -43.76 | 41.9 | HHHHHHHHHHHCHHHHHHHHHHHHHCCCCCC |
| 739 | EEEARKFLEQYNNEEEQMRYQREGFGQGHSA | -43.76 | 41.9 | HHHHHHHHHHHCHHHHHHHHHHHHHCCCCCC |
| 576 | EEEARKFLEQYNNEEEQMSYQQQGFGQGHDA | -43.76 | 41.9 | HHHHHHHHHHHHHHHHHHHHHHHCCCCCCCC |
| 617 | EEEARKFLEQYNNESEQIRYQQNGFGQGHDA | -43.76 | 41.9 | HHHHHHHHHHHHHHHHHHHHHHHCCCCCCCC |
| 797 | EEEARKFLEQYNNESEQIRYQQQGFGQGHDA | -43.76 | 41.9 | HHHHHHHHHHHHHHHHHHHHHHHHCCCCCCC |
| 721 | EEEARKFLEQYNNESEQMRYQQQGFGQGHDA | -43.76 | 41.9 | HHHHHHHHHHHHHHHHHHHHHHHHCCCCCCC |
| 904 | EEEARKFLEQYNNEYEQMFYQQKGFGQGHDA | -43.76 | 45.2 | HHHHHHHHHHHHHHHHHHHHHHHHCCCCCCC |
| 628 | EEELKKFLEQFNNEQEQMTYQRYGFGQGHDA | -43.76 | 45.2 | HHHHHHHHHHHHHHHHHHHHHHHCCCCCCCC |
| 017 | EEELKKFLEQFNNEYEQMFYQQKGFGQGHDA | -43.76 | 48.4 | HHHHHHHHHHHHHHHHHHHHHHHHCCCCCCC |
| 479 | EEELKKFLEQFNNEYEQMFYQQQGFGQGHDA | -43.76 | 48.4 | HHHHHHHHHHHHHHHHHHHHHHHHCCCCCCC |
| 286 | EEELKKFLEQLNNEEEQMRYQQQGFGQGHDA | -43.76 | 41.9 | HHHHHHHHHHHHHHHHHHHHHHHCCCCCCCC |
| 332 | EEELKKFLEQLNNEEEQMRYQQYGFGQGHDA | -43.76 | 41.9 | HHHHHHHHHHHHHHHHHHHHHHHCCCCCCCC |
| 776 | EEELKKFLEQYNNEAEQIYYQQNGFGQGHDA | -43.76 | 45.2 | HHHHHHHHHHHHHHHHHHHHHHHCCCCCCCC |
| 489 | EEELKKFLEQYNNEAEQMRYQKEGFGQGHDA | -43.76 | 45.2 | HHHHHHHHHHHHHHHHHHHHHHHHCCCCCCC |
| 253 | EEELKKFLEQYNNEAEQMRYQQNGFGQGHDA | -43.76 | 45.2 | HHHHHHHHHHHHHHHHHHHHHHHHCCCCCCC |
| 379 | EEELKKFLEQYNNEAEQMRYQQQGFGQGHDA | -43.76 | 45.2 | HHHHHHHHHHHHHHHHHHHHHHHHCCCCCCC |
| 419 | EEELKKFLEQYNNEAEQMRYQRQGFGQGHDA | -43.76 | 45.2 | HHHHHHHHHHHHHHHHHHHHHHHHCCCCCCC |
| 692 | EEELKKFLEQYNNEEEQMFYQQKGFGQGHDA | -43.76 | 45.2 | HHHHHHHHHHHCHHHHHHHHHHHHCCCCCCC |

| | | | | |
|---|---|---|---|---|
| 598 | EEELKKFLEQYNNEEEQMRYQQQGFGQGHDA | -43.76 | 41.9 | HHHHHHHHHHHCHHHHHHHHHHHCCCCCCCC |
| 921 | EEELKKFLEQYNNEEEQMRYQREGFGQGHEA | -43.76 | 41.9 | HHHHHHHHHHHCHHHHHHHHHHHHHCCCCCC |
| 556 | EEELKKFLEQYNNEEEQMSYQQQGFGQGHDA | -43.76 | 41.9 | HHHHHHHHHHHCHHHHHHHHHHHCCCCCCCC |
| 890 | EEELKKFLEQYNNESEQIRYQQNGFGQGHDA | -43.76 | 41.9 | HHHHHHHHHHHHHHHHHHHHHHHCCCCCCCC |
| 104 | EEELKKFLEQYNNESEQMRYQQGGFGQGHDF | -43.76 | 41.9 | HHHHHHHHHHHHHHHHHHHHHHCCCCCCCCC |
| 292 | EEELKKFLEQYNNESEQMRYQQQGFGQGHDA | -43.76 | 41.9 | HHHHHHHHHHHHHHHHHHHHHHHHCCCCCCC |
| 019 | EQEARKFLEQYNNEEEQMRYQREGFGQGHDA | -43.76 | 38.7 | HHHHHHHHHHHCHHHHHHHHHHHHHCCCCCC |
| 938 | EEELKKLLEQINQEAEQMRYQRYGFGQGHDA | -43.70 | 41.9 | HHHHHHHHHHHHHHHHHHHHHHHCCCCCCCC |
| 707 | EEEARKFLEQFNQEYEEMFKQQKGFGSGKDA | -43.63 | 45.2 | HHHHHHHHHHHHHHHHHHHHHHHCCCCCCCC |
| 231 | EEELKKLLEQINNEAEQIRYQQNGFGQGDSR | -43.59 | 48.4 | HHHHHHHHHHHHHHHHHHHHHHHCCCCCCCC |
| 401 | EEELKKLLEQINNEEEQMRYQQQGFGQGDSR | -43.59 | 45.2 | HHHHHHHHHHHCHHHHHHHHHHHCCCCCCCC |
| 974 | EEELKKLLEQINNEQEQMSYQQQGFGQGDSR | -43.59 | 45.2 | HHHHHHHHHHHHHHHHHHHHHHHCCCCCCCC |
| 680 | EEELRKLLEQLNNEQEQMRYQQQGFGQGDSR | -43.59 | 41.9 | HHHHHHHHHHHHHHHHHHHHHHHCCCCCCCC |
| 724 | EEEARKFLEKYNQEEEQMRYQREGFGQGDSR | -43.58 | 51.6 | HHHHHHHHHHHHHHHHHHHHHHHHCCCCCCC |
| 311 | EEELKKFLEKLNQEEEQMRYQQQGFGQGDSR | -43.58 | 51.6 | HHHHHHHHHHHHHHHHHHHHHHHCCCCCCCC |
| 912 | EEEARKFLEQLNQEEEQMRYQQYGFGQGHDA | -43.58 | 41.9 | HHHHHHHHHHHHHHHHHHHHHHHCCCCCCCC |
| 298 | EEEARKFLEQYNQEYEQMEYQRSGFGQGHDA | -43.58 | 45.2 | HHHHHHHHHHHHHHHHHHHHHHHHCCCCCCC |
| 343 | EEELKKFLEQYNQEEEQMRYQREGFGQGHSA | -43.58 | 41.9 | HHHHHHHHHHHHHHHHHHHHHHHHHCCCCCC |
| 473 | EEELKKLLEKLNQEQEQIRYQQNGFGQGDSR | -43.48 | 48.4 | HHHHHHHHHHHHHHHHHHHHHHHCCCCCCCC |
| 811 | EEELRKLLEKLNQEQEQIRYQQNGFGQGDSR | -43.48 | 45.2 | HHHHHHHHHHHHHHHHHHHHHHHCCCCCCCC |
| 697 | EEEARKFLEKYNQEAEQRRYQQNGFGQGHDA | -43.43 | 48.4 | HHHHHHHHHHHHHHHHHHHHHHHCCCCCCCC |
| 208 | EEELKKLLEQINQEAEQITYQQNGFGQGDSR | -43.38 | 48.4 | HHHHHHHHHHHHHHHHHHHHHHHCCCCCCCC |
| 923 | EEELRKLLEQLNQEKEQMSYQQQGFGQGDSR | -43.38 | 41.9 | HHHHHHHHHHHHHHHHHHHHHHHCCCCCCCC |
| 063 | EEEARKMLEQINNEEEQIRYQQQGFGQGHDA | -43.37 | 38.7 | HHHHHHHHHHHHHHHHHHHHHHHCCCCCCCC |
| 320 | EEEARKMLEQINNEEEQMRYQQQGFGQGHDA | -43.37 | 38.7 | HHHHHHHHHHHHHHHHHHHHHHHCCCCCCCC |
| 592 | EEEARKFLEQYNNEAEEIYKQQNGFGSGKDA | -43.34 | 41.9 | HHHHHHHHHHHHHHHHHHHHHHHCCCCCCCC |
| 350 | EEEARKFLEKFNNEYEQMYYQQKGFGQGDSR | -43.29 | 54.8 | HHHHHHHHHHHHHHHHHHHHHHHCCCCCCCC |
| 871 | EEEARKFLEKLNNEEEQMRYQREGFGQGDSR | -43.29 | 51.6 | HHHHHHHHHHHHHHHHHHHHHHHHCCCCCCC |
| 025 | EEEARKFLEKYNNEAEQIRYQQNGFGQGDSR | -43.29 | 54.8 | HHHHHHHHHHHHHHHHHHHHHHHCCCCCCCC |
| 005 | EEEARKFLEKYNNEAEQIYYQQNGFGQGDDR | -43.29 | 54.8 | HHHHHHHHHHHHHHHHHHHHHHHCCCCCCCC |
| 178 | EEELKKFLEKLNNEEEQMQYQRSGFGQGDSR | -43.29 | 54.8 | HHHHHHHHHHHCHHHHHHHHHHHHCCCCCCC |
| 924 | EEELKKFLEKLNNEEEQMRYQQQGFGQGDSR | -43.29 | 51.6 | HHHHHHHHHHHCHHHHHHHHHHHCCCCCCCC |
| 779 | EEELKKFLEKLNNEKEQIEYQRKGFGQGDSR | -43.29 | 51.6 | HHHHHHHHHHHHHHHHHHHHHHHCCCCCCCC |
| 603 | EEELKKFLEKYNNEAEQIYYQQNGFGQGDSR | -43.29 | 54.8 | HHHHHHHHHHHHHHHHHHHHHHHCCCCCCCC |
| 100 | EEELKKFLEKYNNESEQIRYQQQGFGQGDDR | -43.29 | 51.6 | HHHHHHHHHHHHHHHHHHHHHHHCCCCCCCC |
| 267 | EEELKKFLEQYNNEAEQIYYQQNGCGQGHDY | -43.29 | 45.2 | HHHHHHHHHHHHHHHHHHHHHHHCCCCCCCC |
| 951 | EEELKKFLEQYNNEAEQMRYQQGGFGQGHDF | -43.29 | 45.2 | HHHHHHHHHHHHHHHHHHHHHHCCCCCCCCC |
| 229 | EEELKKFLEQLNQEEEQMRYQQQGFGQGDSR | -43.25 | 48.4 | HHHHHHHHHHHHHHHHHHHHHHHCCCCCCCC |

| | | | | |
|---|---|---|---|---|
| 236 | EEELKEFLKQYNEEAEDIYYQQNGFGKGDSR | -43.25 | 58.1 | HHHHHHHHHHHHHHHHHHHHHHHHCCCCCCCC |
| 669 | EEEARKFLEKYNNESEQIRYQQQGFGQGDSR | -43.20 | 51.6 | HHHHHHHHHHHCHHHHHHHHHHHHCCCCCCCC |
| 887 | EEELKKLLEKLNNEEEQIKYQQNGFGQGDSR | -43.18 | 48.4 | HHHHHHHHHHHCHHHHHHHHHHHHCCCCCCCC |
| 614 | EEELKKLLEKLNNEQEQMRYQQNGFGQGDDR | -43.18 | 48.4 | HHHHHHHHHHHHHHHHHHHHHHHHCCCCCCCC |
| 333 | EEEARKFLQQYNNEAEQMRYQQQGFGQGHDA | -43.12 | 45.2 | HHHHHHHHHHHHHHHHHHHHHHHHHCCCCCCC |
| 942 | EEELKKFLQQYNNEEEQMSYQKEGFGQGHDA | -43.12 | 41.9 | HHHHHHHHHHHHHHHHHHHHHHHHHCCCCCCC |
| 613 | EQELRKLLEQINNEEEQMRYQQQGFGQGDSR | -43.10 | 38.7 | HHHHHHHHHHHHHHHHHHHHHHHHCCCCCCCC |
| 030 | EESLRKLLEQINNEAEQMSYQQQGFGQGHDA | -43.10 | 38.7 | HHHHHHHHHHHHHHHHHHHHHHHHCCCCCCCC |
| 268 | EEELKKLLEQINNEAEQMRYQQNGFGQGDSR | -43.09 | 48.4 | HHHHHHHHHHHHHHHHHHHHHHHHCCCCCCCC |
| 609 | EEELKKLLEQINNEEEQMRYQQYGFGQGDSR | -43.09 | 45.2 | HHHHHHHHHHHCHHHHHHHHHHHHCCCCCCCC |
| 099 | EEELKKLLEQLNNEKEQMSYQQQGFGQGDSR | -43.09 | 45.2 | HHHHHHHHHHHHHHHHHHHHHHHHCCCCCCCC |
| 359 | EEELRKLLEQINNEAEQMRYQRQGFGQGDSR | -43.09 | 45.2 | HHHHHHHHHHHHHHHHHHHHHHHHCCCCCCCC |
| 105 | EEELRKLLEQINNEEEQMRYQQQGFGQGDSR | -43.09 | 41.9 | HHHHHHHHHHHCHHHHHHHHHHHHCCCCCCCC |
| 759 | EEELRKLLEQINNEEEQMRYQQYGFGQGDSR | -43.09 | 41.9 | HHHHHHHHHHHCHHHHHHHHHHHHCCCCCCCC |
| 396 | EEELRKLLEQLNNEKEQMSYQQQGFGQGDSR | -43.09 | 41.9 | HHHHHHHHHHHHHHHHHHHHHHHHCCCCCCCC |
| 929 | EEELKKFLEKLNNEAEQMQYQRAGFGQGDSR | -43.08 | 54.8 | HHHHHHHHHHHHHHHHHHHHHHHHHCCCCCCC |
| 176 | EEEARKFLEQFNNEEEQMSYQQQGFGQGDSR | -43.03 | 51.6 | HHHHHHHHHHHHHHHHHHHHHHHHCCCCCCCC |
| 140 | EEEARKFLEQYNNEAEQMRYQQQGFGQGDSR | -43.03 | 51.6 | HHHHHHHHHHHHHHHHHHHHHHHHCCCCCCCC |
| 812 | EEELKKFLEQYNNEAEQMRYQQNGFGQGDSR | -43.03 | 51.6 | HHHHHHHHHHHHHHHHHHHHHHHHCCCCCCCC |
| 344 | EEELKKFLEQYNNEEEQMSYQQNGFGQGDSR | -43.03 | 48.4 | HHHHHHHHHHHCHHHHHHHHHHHHCCCCCCCC |
| 347 | EEELKKFLEQYNNEYEQMEYQRSGFGQGDSR | -43.03 | 51.6 | HHHHHHHHHHHHHHHHHHHHHHHHCCCCCCCC |
| 695 | EEELKKFLEKYNQEEEQRRYQQQGFGQGHDA | -42.95 | 45.2 | HHHHHHHHHHHCHHHHHHHHHHHHCCCCCCCC |
| 666 | EEEARKFLEQYNNEYEQMEYQRTGRGQGHDF | -42.84 | 41.9 | HHHHHHHHHHHHHHHHHHHHHHHHCCCCCCCC |
| 841 | EEEARKFLEQFNQEEEQMSYQQNGFGQGDSR | -42.82 | 51.6 | HHHHHHHHHHHHHHHHHHHHHHHHCCCCCCCC |
| 420 | EEEARKFLEQYNQEEEQMSYQQQGFGQGDSR | -42.82 | 48.4 | HHHHHHHHHHHHHHHHHHHHHHHHCCCCCCCC |
| 220 | EEELKKFLEQFNQEYEQMFYQQKGFGQGDSR | -42.82 | 54.8 | HHHHHHHHHHHHHHHHHHHHHHHHCCCCCCCC |
| 041 | EEELKKFLEQYNQEEEQMSYQKEGFGQGDSR | -42.82 | 48.4 | HHHHHHHHHHHHHHHHHHHHHHHHCCCCCCCC |
| 792 | EEELKKFLEQYNQEEEQMSYQQQGFGQGDSR | -42.82 | 48.4 | HHHHHHHHHHHHHHHHHHHHHHHHCCCCCCCC |
| 270 | EEELKKLLEQLNQEQEQMRYQQQGFGQGDSR | -42.71 | 45.2 | HHHHHHHHHHHHHHHHHHHHHHHHCCCCCCCC |
| 476 | EEELKKLLEQINQEEEQMRYQQQGFGQGDSR | -42.69 | 45.2 | HHHHHHHHHHHHHHHHHHHHHHHHCCCCCCCC |
| 132 | EEEARKFLQQYNNEAEQMRYQQNGFGQGHDA | -42.65 | 45.2 | HHHHHHHHHHHHHHHHHHHHHHHHHCCCCCCC |
| 560 | EEEARKFLEKFNNEEEQMRYQREGFGQGDSR | -42.62 | 54.8 | HHHHHHHHHHHCHHHHHHHHHHHHHCCCCCCC |
| 319 | EEELKKFLEKFNNEAEQMSYQQQGFGQGDSR | -42.62 | 58.1 | HHHHHHHHHHHHHHHHHHHHHHHHCCCCCCCC |
| 095 | EEEARKFLEQLNNEEEQMRYQQYGFGQGDSR | -42.55 | 48.4 | HHHHHHHHHHHHHHHHHHHHHHHHCCCCCCCC |
| 072 | EEELKKFLEQYNNEYEQMFYQQKGFGQGDSR | -42.55 | 51.6 | HHHHHHHHHHHHHHHHHHHHHHHHCCCCCCCC |
| 581 | EEEARKFLEQFNNEYEQMFYQQKGFGQGDSR | -42.52 | 54.8 | HHHHHHHHHHHHHHHHHHHHHHHHCCCCCCCC |
| 059 | EEEARKFLEQYNNEEEQMRYQRQGFGQGDSR | -42.52 | 48.4 | HHHHHHHHHHHCHHHHHHHHHHHHCCCCCCCC |
| 155 | EEELKKFLEQFNNEYEQMFYQQKGFGQGDSR | -42.52 | 54.8 | HHHHHHHHHHHHHHHHHHHHHHHHCCCCCCCC |

| | | | | |
|---|---|---|---|---|
| 495 | EEELKKFLEQYNNEAEQIYYQQNGFGQGDSR | -42.52 | 51.6 | HHHHHHHHHHHHHHHHHHHHHHHCCCCCCCC |
| 076 | EEELKKFLEQYNNEAEQMRYQQQGFGQGDSR | -42.52 | 51.6 | HHHHHHHHHHHHHHHHHHHHHHHCCCCCCCC |
| 170 | EEELKKFLEQYNNEYEQMEYQRKGFGQGDSR | -42.52 | 48.4 | HHHHHHHHHHHHHHHHHHHHHHHCCCCCCCC |
| 536 | EQEARKFLEQFNNEEEQMSYQQQGFGQGDSR | -42.52 | 48.4 | HHHHHHHHHHHHHHHHHHHHHHHCCCCCCCC |
| 064 | EEEARKFLEKYNNEEEQMRYQREGFGQGDSR | -42.30 | 51.6 | HHHHHHHHHHHCCHHHHHHHHHHHCCCCCCC |
| 367 | EEEARKFLEQFNQEYEQMFYQQKGFGQGDSR | -42.15 | 54.8 | HHHHHHHHHHHHHHHHHHHHHHHCCCCCCCC |
| 150 | EEELKKFLEQYNQEEEQMRYQREGFGQGDSR | -42.15 | 48.4 | HHHHHHHHHHHHHHHHHHHHHHHHCCCCCCC |
| 998 | EEELKKFLEQYNQESEQIRYQQQGFGQGDSR | -42.15 | 48.4 | HHHHHHHHHHHHHHHHHHHHHHHCCCCCCCC |
| 125 | EEELKKFLEQYNQESEQMRYQQGGFGQGDSR | -42.15 | 48.4 | HHHHHHHHHHHHHHHHHHHHHHCCCCCCCCC |
| 139 | EEELRRLLEKINNEEEQMRYQQYGFGQGDSR | -41.95 | 45.2 | HHHHHHHHHHHCHHHHHHHHHHHCCCCCCCC |
| 357 | EEEARKFLEQLNNEEEQMRYQQQGFGQGDSR | -41.85 | 48.4 | HHHHHHHHHHHHHHHHHHHHHHHCCCCCCCC |
| 145 | EEEARKFLEQYNNESEQMRYQKEGFGQGDSR | -41.85 | 48.4 | HHHHHHHHHHHHHHHHHHHHHHHHCCCCCCC |
| 944 | EEELKKFLEQLNNEEEQMRYQQQGFGQGDSR | -41.85 | 48.4 | HHHHHHHHHHHHHHHHHHHHHHHCCCCCCCC |
| 926 | EEEARKFLEQYNNEAEQMRYQQNGFGQGDSR | -41.62 | 51.6 | HHHHHHHHHHHHHHHHHHHHHHHCCCCCCCC |
| 364 | EESLKKFLEQYNNEEEQMSYQQQGFGQGDSR | -41.30 | 48.4 | HHHHHHHHHHHHHHHHHHHHHHHCCCCCCCC |

### Supporting Figures

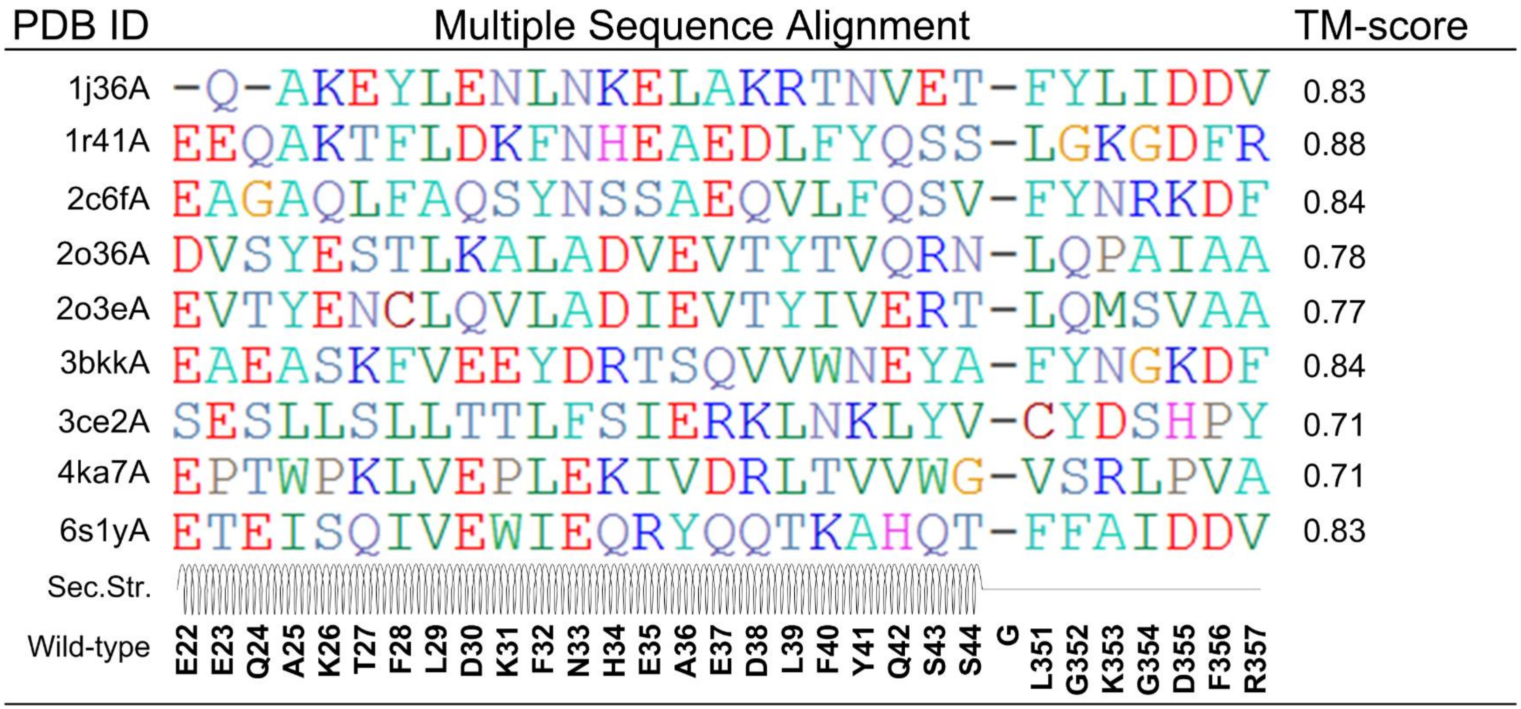


**Figure S1**. Peptide multiple sequence alignment for evolutionary profile construction.
